# Selective sweeps under dominance and inbreeding

**DOI:** 10.1101/318410

**Authors:** Matthew Hartfield, Thomas Bataillon

## Abstract

A major research goal in evolutionary genetics is to uncover loci experiencing positive selection. One approach involves finding ‘selective sweeps’ patterns, which can either be ‘hard sweeps’ formed by *de novo* mutation, or ‘soft sweeps’ arising from recurrent mutation or existing standing variation. Existing theory generally assumes outcrossing populations, and it is unclear how dominance affects soft sweeps. We consider how arbitrary dominance and inbreeding via self-fertilisation affect hard and soft sweep signatures. With increased self-fertilisation, they are maintained over longer map distances due to reduced effective recombination and faster beneficial allele fixation times. Dominance can affect sweep patterns in outcrossers if the derived variant originates from either a single novel allele, or from recurrent mutation. These models highlight the challenges in distinguishing hard and soft sweeps, and propose methods to differentiate between scenarios.

## Introduction

Inferring adaptive mutations from nucleotide polymorphism data is a major research goal in evolutionary genetics, and has been subject to extensive modelling work to determine the footprints they leave in genome data (Stephan 2019). The earliest models focussed on a scenario where a beneficial mutation arose as a single copy before rapidly fixing. Linked neutral mutations then ‘hitchhike’ to fixation with the adaptive variant, reducing diversity around the selected locus (Maynard Smith and Haigh 1974; Kaplan *et al*. 1989). Hitchhiking also increases linkage disequilibrium at regions flanking the selected site, by raising the haplotype carrying the selected allele to high frequency. It is minimal when measured at sites either side of the selected mutation (Thomson 1977; Innan and Nordborg 2003; McVean 2007). These theoretical expectations have spurred the creation of summary statistics for detecting sweeps, usually based on finding genetic regions exhibiting extended haplotype homozygosity (Sabeti *et al*. 2002; Kim and Nielsen 2004; Voight *et al*. 2006; Ferrer-Admetlla *et al*. 2014; Vatsiou *et al*. 2016), or an increase in high frequency derived variants (Fay and Wu 2000; Kim and Stephan 2002; Nielsen 2005; Boitard *et al*. 2009; Yang *et al*. 2018; Fujito *et al*. 2018).

Classic hitchhiking models consider ‘hard’ sweeps, where the common ancestor of an adaptive allele occurs after the onset of selection (Hermisson and Pennings 2017). Recent years have seen a focus on ‘soft’ sweeps, where the most recent common ancestor of a beneficial allele appeared before it became selected for (reviewed by Barrett and Schluter (2008); Messer and Petrov (2013); Hermisson and Pennings (2017)). Soft sweeps can originate from beneficial mutations being introduced by recurrent mutation at the target locus (Pennings and Hermisson 2006a,b), or originating from existing standing variation that was either neutral or deleterious (Orr and Betancourt 2001; Innan and Kim 2004; Przeworski *et al*. 2005; Hermisson and Pennings 2005; Wilson *et al*. 2014; Berg and Coop 2015; Wilson *et al*. 2017). A key property of soft sweeps is that the beneficial variant is present on multiple genetic backgrounds as it sweeps to fixation, so different haplotypes may carry the derived allele. This property is often used to detect soft sweeps in genetic data (Peter *et al*. 2012; Vitti *et al*. 2013; Garud *et al*. 2015; Garud and Petrov 2016;Schrider and Kern 2016; Sheehan and Song 2016; Harris *et al*. 2018a; Kern and Schrider 2018; Harris and DeGiorgio 2018, 2019). Soft sweeps have been reported in *Drosophila* (Karasov *et al*. 2010; Garud *et al*. 2015; Garud and Petrov 2016; Vy *et al*. 2017), humans (Peter *et al*. 2012; Schrider and Kern 2017), maize (Fustier *et al*. 2017), *Anopheles* mosquitoes (Xue *et al*. 2019), and pathogens including *Plasmodium falciparum* (Anderson *et al*. 2016) and HIV (Pennings *et al*. 2014;Williams and Pennings 2019). Yet determining how extensive soft sweeps are in nature remains a contentious issue (Jensen 2014; Harris *et al*. 2018b).

Up to now, there have only been a few investigations into how dominance affects sweep signatures. In a simulation study, Teshima and Przeworski (2006) explored how recessive mutations spend long periods of time at low frequencies, increasing the amount of recombination that acts on derived haplotypes, weakening signatures of hard sweeps. Fully recessive mutations may need a long time to reach a significantly high frequency to be detectable by genome scans (Teshima *et al*. 2006). Ewing *et al*. (2011) have carried out a general mathematical analysis of how dominance affects hard sweeps. Yet the impact of dominance on soft sweeps has yet to be explored in depth.

In addition, existing models have so far focussed on randomly mating populations, with haplotypes freely mixing between individuals over generations. Different reproductive modes alter how alleles are inherited, affecting the hitchhiking effect. Self-fertilisation, where male and female gametes produced from the same individual can fertilise one another, can alter adaptation rates and selection signatures (Hartfield *et al*. 2017). This mating system is prevalent amongst angiosperms (Igic and Kohn 2006), some animals (Jarne and Auld 2006) and fungi (Billiard *et al*. 2011). As the effects of dominance and self-fertilisation become strongly intertwined, it is important to consider both together. Dominant mutations are more likely to fix than recessive ones in outcrossers, as they have a higher initial selection advantage (Haldane 1927). Yet recessive alleles can fix more easily in selfers than in outcrossers as homozygote mutations are created more rapidly (Charlesworth 1992; Glémin 2012). Furthermore, a decrease in effective recombination rates in selfers (Nordborg *et al*. 1996; Nordborg 2000; Charlesworth and Charlesworth 2010) can interfere with selection acting at linked sites, making it likelier that deleterious mutations hitchhike to fixation with adaptive alleles (Hartfield and Glémin 2014), or competition between adaptive mutations at closely-linked loci increases the probability that rare mutations are lost by drift (Hartfield and Glémin 2016).

In a constant-sized population, beneficial mutations can be less likely to fix from standing variation (either neutral or deleterious) in selfers as they maintain lower diversity levels (Glémin and Ronfort 2013). Yet adaptation from standing variation becomes likelier in selfers compared to outcrossers under ‘evolutionary rescue’ scenarios, where swift adaptation is needed to prevent population extinction following environmental change. Here, rescue mutations are only present in standing variation as the population size otherwise becomes too small (Glémin and Ronfort 2013). Self-fertilisation further aids this process by creating beneficial homozygotes more rapidly than in outcrossing populations (Uecker 2017).

Little data currently exists on the extent of soft sweeps in self-fertilisers. Many selfing organisms exhibit sweep-like patterns, including *Arabidopsis thaliana* (Long *et al*. 2013; Huber *et al*. 2014; Fulgione *et al*. 2018; Price *et al*. 2018); Caenorhabditis elegans (Andersen *et al*. 2012); *Medicago truncatula* (Bonhomme *et al*. 2015); and *Microbotryum* fungi (Badouin *et al*. 2017). Soft sweeps have also been reported in soya bean (Zhong *et al*. 2017). Detailed analyses of these cases has been hampered by a lack of theory on how hard and soft sweep signatures should manifest themselves under different self-fertilisation and dominance levels. Previous studies have only focussed on special cases; Hedrick (1980) analysed linkage disequilibrium caused by a hard sweep under self-fertilisation, while Schoen *et al*. (1996) modelled sweep patterns caused by modifiers that altered the mating system in different ways.

To this end, we develop a selective sweep model that accounts for dominance and inbreeding via self–fertilisation. We determine the genetic diversity present following a sweep from either a *de novo* mutation, or from standing variation. We also determine the number of segregating sites and the site frequency spectrum, while comparing results to an alternative soft-sweep model where adaptive alleles arise via recurrent mutation. Note that we focus here on single sweep events, rather than characterising how sweeps affect genome-wide diversity (Elyashiv *et al*. 2016;Campos *et al*. 2017; Booker and Keightley 2018; Rettelbach *et al*. 2019).

## Results

### Model Outline

We consider a diploid population of size *N* (carrying 2*N* haplotypes in total). Individuals reproduce by self-fertilisation with probability *σ*, and outcross with probability 1 − *σ*. A derived allele arises at a locus, and we are interested in determining the population history of neutral regions that are linked to it, with a recombination rate *r* between them. We principally look at the case where the beneficial allele arises from previously–neutral standing variation, and subsequently look at a sweep arising from recurrent mutation. The derived allele initially segregates neutrally for a period of time, then becomes advantageous with selective advantage 1 + *hs* when heterozygous and 1 + *s* when homozygous, with 0 < *h* < 1 and *s* > 0. We further assume that the population size is large and selection is large enough so that the beneficial allele’s change in frequency can be modelled deterministically (i.e., *N_e_hs* ≫ 1 and 1/*N_e_* ≪ *s* ≪ 1). Table 1 lists the notation used in the analysis.

**Table 1.** Glossary of Notation.

| Symbol | Usage |
| --- | --- |
| $N$ | Population size (with $2N$ haplotypes) |
| $\sigma$ | Proportion of matings that are self-fertilising |
| $F$ | Wright's inbreeding coefficient, probability of identity-by-descent at a single gene, equal to $\sigma/(2 - \sigma)$ at steady-state |
| $\Phi$ | Joint probability of identity-by-descent at two loci (Equation 1) |
| $N_e$ | Effective population size, equal to $N/(1 + F)$ with selfing |
| $r$ | Recombination rate between loci $A$ and $B$ |
| $r_{eff}$ | 'Effective' recombination rate, approximately equal to $r(1 - 2F + \Phi)$ with selfing |
| $R$ | $2Nr$, the population-level recombination rate |
| $p_0$ | Frequency at which the derived allele at $B$ becomes advantageous |
| $p_{0,A}$ | Accelerated (effective) starting frequency of $B$ appearing as a single copy, conditional on fixation |
| $s$ | Selective advantage of derived allele at $B$ |
| $h$ | Dominance coefficient of derived allele at $B$ |
| $t$ | Number of generations in the past from the present day |
| $\tau_{p_0}$ | Time in the past when derived locus became beneficial |
| $p(t)$ | Frequency of beneficial allele at time $t$ |
| $P_c$ | Probability of coalescence at time $t$ |
| $P_r$ | Probability of recombination at time $t$ |
| $P_m$ | Probability of mutation at time $t$ |
| $P_{NE}$ | Probability that neutral marker does not coalesce or recombine during sweep phase |
| $P_{R,Sw}$ | Probability that neutral marker recombines during sweep phase |
| $P_{R,Sd}$ | Probability that neutral marker recombines during standing phase |
| $P_{M,Sw}$ | Probability that a lineage mutates during sweep phase |
| $P_{M,Sd}$ | Probability that a lineage mutates during standing phase |
| $H_l, H_h$ | 'Effective' dominance coefficient for allele at low, high frequency |
| $\pi$ | Pairwise diversity at site ($\pi_0$ is expected value without a sweep) |
| $\pi_{SV}$ | Pairwise diversity following sweep from standing variation |
| $\pi_M$ | Pairwise diversity following sweep from recurrent mutation |
| $\mu$ | Probability of neutral mutation occurring per site per generation |
| $\mu_b$ | Probability of beneficial mutation occurring at target locus per generation |
| $\theta = 4N_e\mu$ | Population level neutral mutation rate |
| $\Theta_b = 2N_e\mu_b$ | Population level beneficial mutation rate |

Our goal is to determine how the spread of the derived, adaptive allele affects genealogies at linked neutral regions. For a sweep originating from standing variation, we follow the approach of Berg and Coop (2015) and, looking backwards in time, break down the selected allele history into two phases. The first phase (the ‘sweep phase’) considers the derived allele being selectively favoured from an initial frequency *p*_0_ and spreading through the population. The second phase (the ‘standing phase’) assumes that the derived allele is present at a fixed frequency *p*_0_. During both phases, a pair of haplotypes can either coalesce, or one of them recombines onto the ancestral background. A schematic is shown in Figure 1.

**Figure 1.**
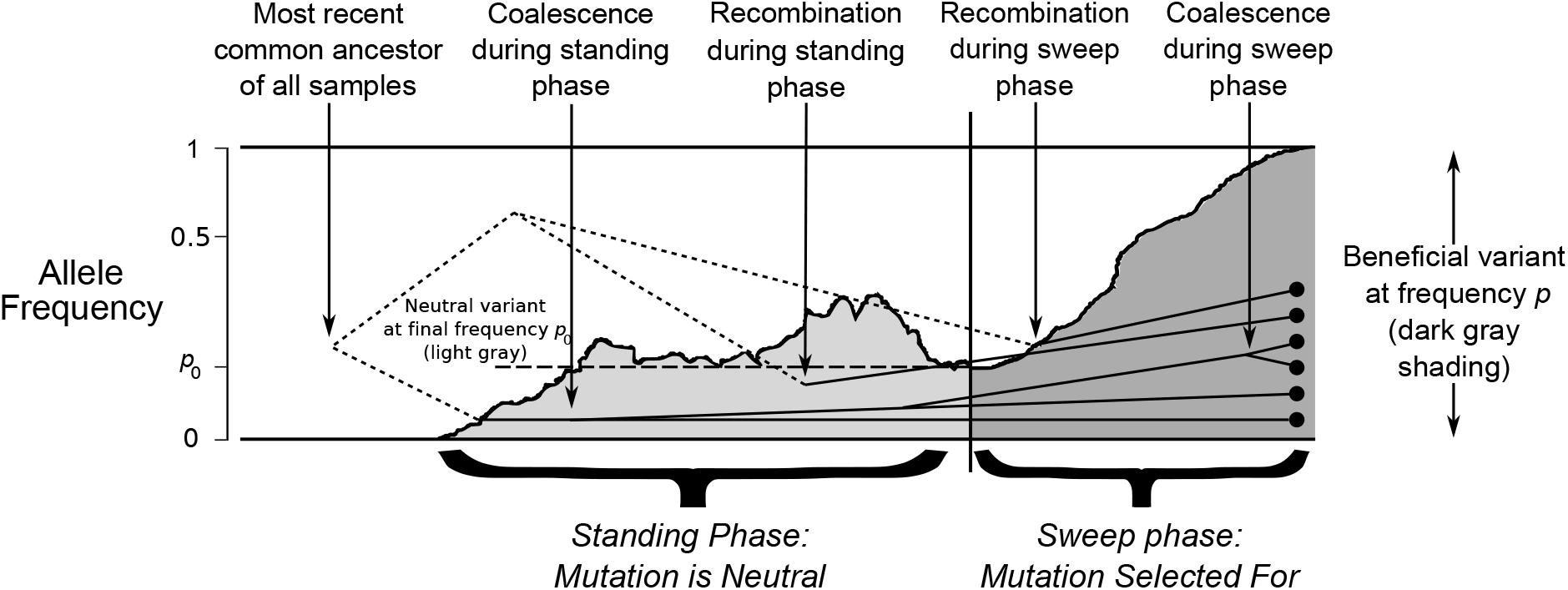
A schematic of the model. The history of the derived variant is separated into two phases; the ‘standing phase’ (shown in light gray), and the ‘sweep phase’ (shown in dark gray). Axis on the left-hand side show allele frequency on a log-scale. Dots on the right-hand side represent a sample of haplotypes taken at the present day, with lines representing their genetic histories. Solid lines represent coalescent histories for the derived genetic background; dotted lines represent coalescent histories for the ancestral, neutral background.

During the sweep phase, the derived allele will also cause the spread of linked haplotypes that it appeared on. Over the course of the sweep, haplotypes are broken down by recombination; the total number of recombination events is proportional to *rτ*_*p*_0__, where *τ*_*p*_0__ is the fixation time of the beneficial allele, given an initial frequency *p*_0_ (Maynard Smith and Haigh 1974). Dominance and self–fertilisation have different effects on *τ*_*p*_0__, and therefore the number of fixing haplotypes. If *p*_0_ is low (~1/2*N*) then highly recessive or dominant mutations take longer to go to fixation (Glémin 2012), which can increase the number of recombination events. Dominance also affects the nature of the sweep trajectory. For example, recessive mutations spend more time at a low frequency compared to dominant mutations. These different sweep trajectories can also affect the final sweep profile (Teshima and Przeworski 2006). Self–fertilisation leads to decreased fixation time of adaptive mutations through converting heterozygotes to homozygotes (Glémin 2012). Recombination is likelier to act between homozygotes under self-fertilisation, so its effective rate is reduced by a factor 1 − 2*F* + Φ, for *F* = *σ*/(2 − *σ*) the inbreeding coefficient (Nordborg *et al*. 1996; Nordborg 2000) and Φ the joint probability of identity-by-descent at the two loci (Roze 2009, 2016; Hartfield and Glémin 2016), defined as:

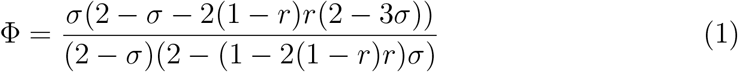

Note that 1 − 2*F* + Φ approximates to 1 − *F* (as Φ ≈ *F*), unless *σ* is close to one and *r* is high (approximately greater than 0.1).

During the standing phase, the amount of initial recombinant haplotypes that are swept to fixation depend on the relative rates of recombination and coalescence. The latter occurs with probability proportional to 1/2*N_e_* for *N_e_* the effective population size. Under self–fertilisation *N_e_* = *N*/(1 + *F*) (Wright 1951; Pollak 1987;Charlesworth 1992; Caballero and Hill 1992; Nordborg and Donnelly 1997), so self–fertilisation increases the coalescence probability. This scaling factor remains a good approximation if there is non-Poisson variation in offspring, unless female fitness strongly affects reproduction number (Laporte and Charlesworth 2002). Although we focus on inbreeding via self-fertilisation, the scalings *N_e_* = *N*/(1 + *F*) and *r_e_* ≈ *r*(1 − *F*) should also hold under other systems of regular inbreeding (Caballero and Hill 1992; Charlesworth and Charlesworth 2010, Box 8.4).

We will outline how both coalescence and recombination act during both of these phases, and use these calculations to determine selective sweep properties. Previous models tended to only determine how lineages recombine away from the derived background during the sweep phase, without considering how two lineages coalesce during the sweep phase. If lineages coalesce during the sweep, then the total number of unique recombination events, and hence the number of linked haplotypes, are reduced. Barton (1998) showed that these coalescent events are negligible only for very strong selection (log(*Ns*) ≫ 1; and B. Charlesworth, un-published results). Hence, accounting for these coalescent events is important for producing accurate matches with simulation results.

Throughout, analytical solutions are compared to results from Wright-Fisher forward-in-time stochastic simulations that were ran using SLiM version 3.3 (Haller and Messer 2019). Results for outcrossing populations were also tested using coalescent simulations ran with *msms* (Ewing and Hermisson 2010). The simulation methods are outlined in Supplementary File S2.

#### Data Availability

File S1 is a *Mathematica* notebook of analytical derivations and simulation results. File S2 contains additional methods, results and figures. File S3 contains copies of the simulation scripts, which are also available from https://github.com/MattHartfield/SweepDomSelf. Supplemental material has also been uploaded to Figshare.

### Probability of events during sweep phase

We first look at the probability of events (coalescence or recombination) acting during the sweep phase for the simplest case of two alleles. Looking back in time following the fixation of the derived mutation, sites linked to the beneficial allele can either coalesce or recombine onto the ancestral genetic background. Let *p*(*t*) be the adaptive mutation frequency at time *t*, defined as the number of generations prior to the present day. Further define *p*(0) = 1 (i.e., the allele is fixed at the present day), and *τ*_*p*_0__ the time in the past when the derived variant became beneficial (i.e., *p*(*τ*_*p*_0__) = *p*_0_).

For a pair of haplotype samples carrying the derived allele, if it is at frequency *p*(*t*) at time *t*, this lineage pair can either coalesce or one of the haplotypes recombine onto the ancestral background. Each event occurs with probability:

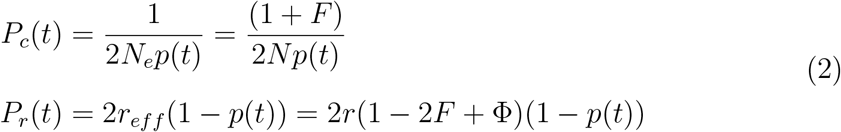

Equation 2 is based on those obtained by Kaplan *et al*. (1989), assuming that *N_e_* = *N*/(1 + *F*) due to self-fertilisation (Pollak 1987; Charlesworth 1992; Caballero and Hill 1992; Nordborg and Donnelly 1997), and *r_eff_* = *r*(1 − 2*F* + Φ) is the ‘effective’ recombination rate after correcting for increased homozygosity due to self-fertilisation (Nordborg *et al*. 1996; Nordborg 2000; Charlesworth and Charlesworth 2010; Roze 2009, 2016; Hartfield and Glémin 2016). Equation 2 demonstrates how each event is differently influenced by *p*. In particular, the per– generation coalescence probability *P_c_* can be small unless *p* is close to 1/2*N*. The total probability that coalescence occurs during the sweep phase increases if the beneficial allele spends a sizeable time at low frequency, e.g., when it is recessive. The terms in Equation 2 can also be defined as functions of *p*.

We are interested in calculating (i) the probability *P_NE_* that no coalescence or recombination occurs in the sweep phase; (ii) the probability *P_R,Sw_* that recombination acts on a lineage to transfer it to the neutral background that is linked to the ancestral allele, assuming that no more than one recombination event occurs per generation (see Campos and Charlesworth (2019) for derivations assuming multiple recombination events). We will go through these probabilities in turn to determine expected pairwise diversity. For *P_NE_*, the total probability that the two lineages do not coalesce or recombine over *τ*_*p*_0__ generations equals:

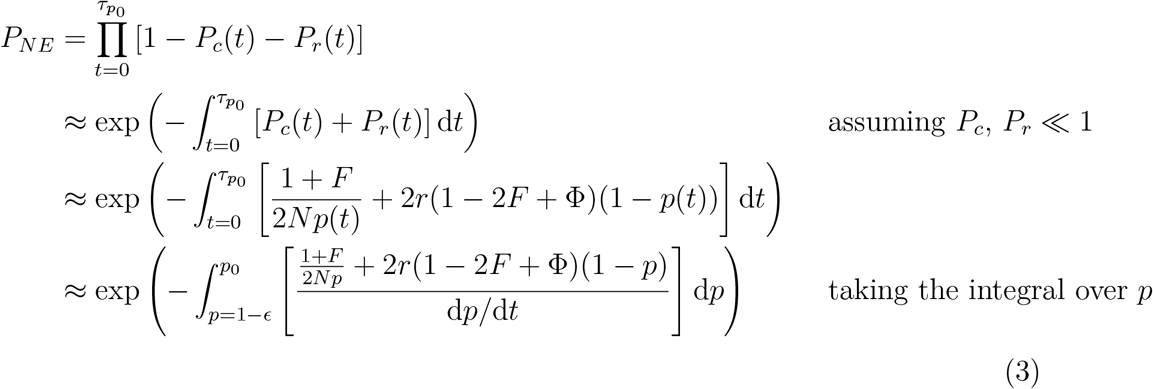

Here *ϵ* is a small term and 1 − *ϵ* is the upper limit of the deterministic spread of the beneficial allele. We will discuss in the section ‘Effective starting frequency from a *de novo* mutation’ what a reasonable value for *ϵ* should be. Also note that we switch from a discrete–time calculation to a continuous–time calculation, which can give simplifying results. To calculate *P_NE_* we insert the deterministic change in allele frequency *p* (Glémin 2012):

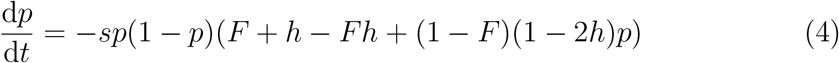

Note the negative factor in Equation 4 since we are looking back in time. By substituting Equation 4 into Equation 3, we obtain an analytical solution for *P_NE_*, although the resulting expression is complicated (Section A of Supplementary File S1).

To calculate *P_R,Sw_*, the probability that recombination acts during the sweep, we first calculate the probability that recombination occurs when the beneficial allele is at frequency *p*′. Here, no events occur in the time leading up to *p*′, then a recombination event occurs with probability *P_r_*(*p*′) = 2*r*(1 − 2*F* + Φ)(1 − *p*′). *P_R,Sw_* is obtained by integrating this probability over the entire sweep from time 0 to *τ*_*p*_0__:

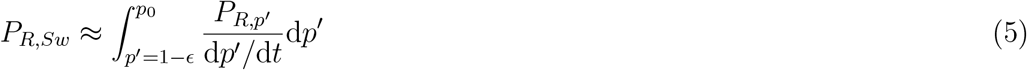

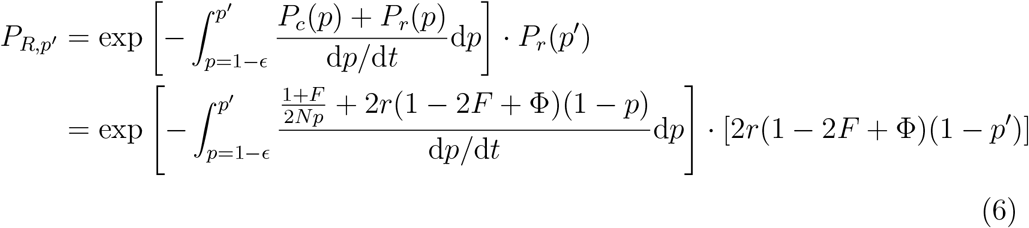

Note that the exponential term of *P_R,p′_* is different from *P_NE_* (Equation 3) since the upper integral limit is to *p′* rather than *p*_0_. That is, it only covers part of the sweep phase. Equation 5 is evaluated numerically. In Supplementary File S2, we provide a ‘star–like’ analytical approximation to *P_NE_* that assumes no coalescence during the sweep phase.

### Probability of coalescence from standing variation

The variant becomes advantageous at frequency *p*_0_. We assume that *p*_0_, and hence event probabilities, remain fixed over time. Berg and Coop (2015) have shown this assumption provides a good approximation to coalescent rates during the standing phase. The outcome during the standing phase is thus determined by competing Poisson processes. The two haplotypes could coalesce, with an exponentially-distributed waiting time with rate *P_c_*(*p*_0_) = (1 + *F*)/(2*N*_*p*_0__). Alternatively, one of the two haplotypes could recombine onto the ancestral background with mean waiting time *P_r_*(*p*_0_) = 2*r_eff_*(1 − *p*_0_). For two competing exponential distributions with rates λ_1_ and λ_2_, the probability of the first event occurring given an event happens equals λ_1_/(λ_1_ + λ_2_) (Wakeley 2009). Hence the probability that recombination occurs instead of coalescence equals:

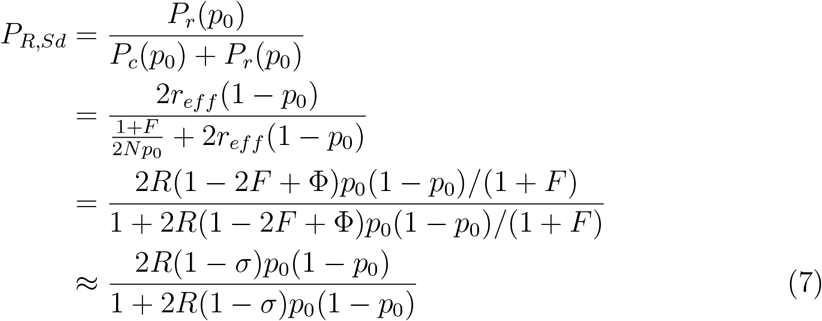

The probability of coalescence rather than recombination is *P_C,S_d__* = 1 − *P_R, Sd_*. Here *R* = 2*Nr* is the population-scaled recombination rate. The final approximation arises as (1 − 2*F* + Φ)/(1 + *F*) ≈ (1 − *F*)/(1 + *F*) = (1 − *σ*) if Φ ≈ *F*. This term reflects how increased homozygosity reduces both effective recombination and *N_e_*, with the latter making coalescence more likely. In addition, it also highlights how the signature of a sweep from standing variation, as characterised by the spread of different initial recombinant haplotypes, is spread over an increased distance of 1/(1 − *σ*) under self–fertilisation.

### Effective starting frequency for a *de novo* mutation

When a new beneficial mutation appears at a single copy, it is highly likely to go extinct by chance (Fisher 1922; Haldane 1927). Beneficial mutations that increase in frequency faster than expected when rare are more able to overcome this stochastic loss and reach fixation. These beneficial mutations will hence display an apparent ‘acceleration’ in their logistic growth, equivalent to having a starting frequency that is greater than 1/(2*N*) (Maynard Smith 1976; Barton 1998; Desai and Fisher 2007; Martin and Lambert 2015). Correcting for this acceleration is important to accurately model hard sweep signatures, and inform on the minimum level of standing variation needed to differentiate a hard sweep from one originating from standing variation.

In Section B of Supplementary File S1, we determine that hard sweeps that go to fixation have the following effective starting frequency:

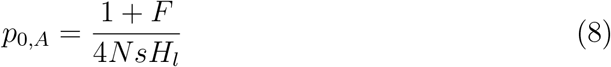

where *H_l_* = *F* + *h* − *Fh* is the effective dominance coefficient for mutations at a low frequency. This result is consistent with those of Martin and Lambert (2015), who obtained a distribution of effective starting frequencies using stochastic differential equations. This acceleration effect can create substantial increases in the effective *p*_0_, especially for recessive mutations (Figure 2).

**Figure 2.**
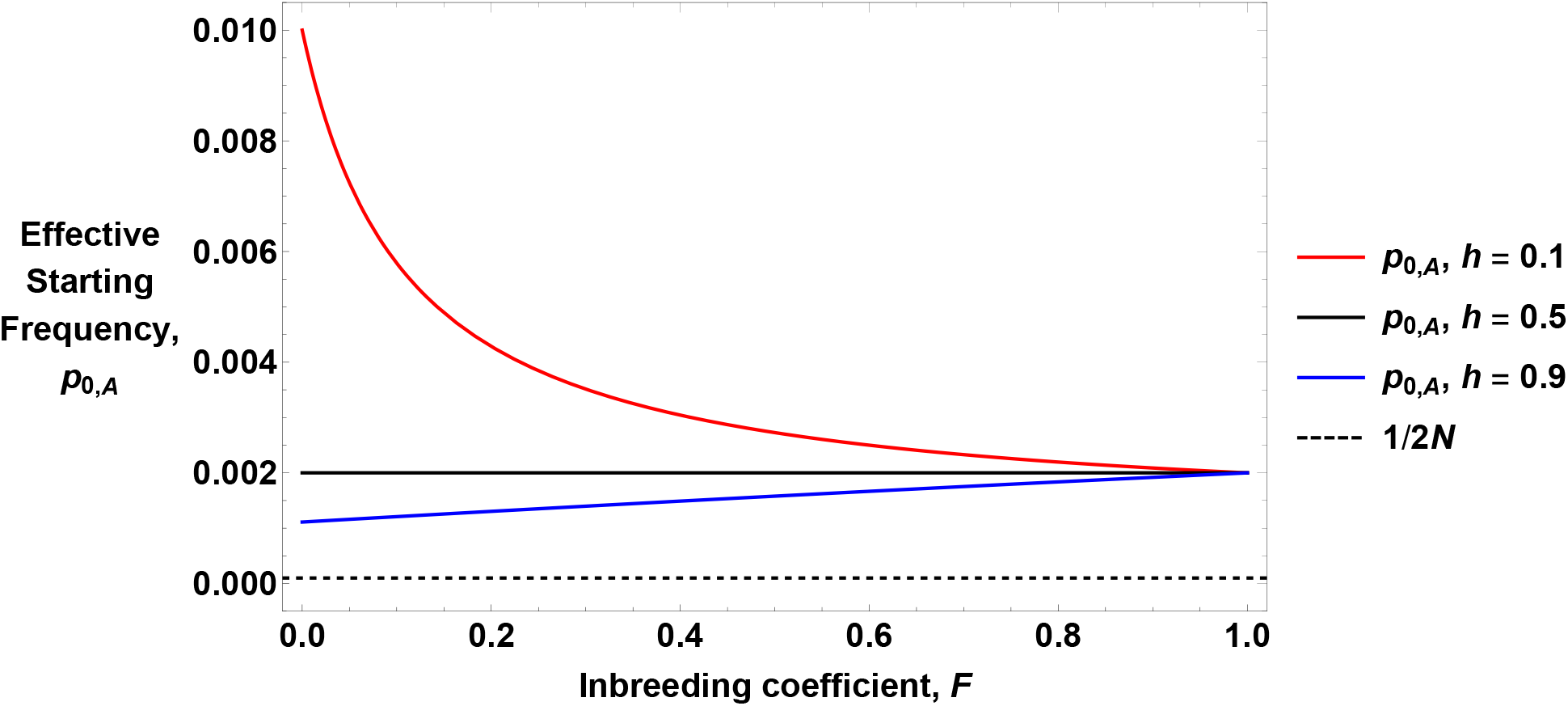
Examples of the effective starting frequency. Equation 8 is plotted as a function of *F* for different dominance values, as shown in the legend. Other parameters are *N* = 5, 000, *s* = 0.05. The dashed line shows the actual starting frequency, 1/2*N*.

#### Effective final frequency

The effective final frequency of the derived allele 1 − *ϵ*, at which its spread is no longer deterministic, can be obtained by setting *ϵ* = *p*_0,*A*_(1 − *h*); that is, by substituting *H_l_* to *H_h_* = 1 − *h* + *Fh* in Equation 8. Van Herwaarden and Van der Wal (2002) determined that the sojourn time for an allele with dominance coefficient *h* that is increasing in frequency, is the same for an allele decreasing in frequency with dominance 1 − *h*. Glémin (2012) showed that this result also holds under any inbreeding value *F* (and B. Charlesworth, unpublished results).

### Expected Pairwise Diversity

We use *P_NE, P_R,sw__* and *P_R,sd_* to calculate the expected pairwise diversity (denoted *π*) present around a sweep. During the sweep phase, the two neutral sites could either coalesce, or one of them recombines onto the ancestral background. If coalescence occurs, since it does so in the recent past then it is assumed that no diversity exist between samples, i.e., *π* ≈ 0 for *π* the average number of differences between two alleles (Tajima 1983). In reality there may be some residual diversity caused by appearance of mutations during the sweep phase; we do not account for these mutations while calculating *π* but will do so when calculating the site-frequency spectrum. Alternatively, if one of the two samples recombines onto the neutral background, they will have the same pairwise diversity between them as the background population (*π*_0_). If the two samples trace back to the standing phase (with probability *P_NE_*) then the same logic applies. Hence the expected diversity following a sweep *π_SV_*, relative to the background value *π*_0_, equals:

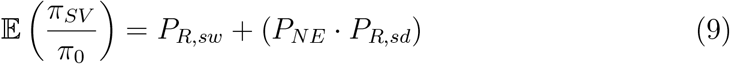

The full solution to Equation 9 can be obtained by plugging in the relevant parts from Equations 3, 5 and 7, which we evaluate numerically. Equation 9 is undefined for *h* = 0 or 1 with *σ* = 0; these cases can be derived separately.

Figure 3 plots Equation 9 with different dominance, self-fertilisation, and standing frequency values. The analytical solution fits well compared to forward-in-time simulations, yet slightly overestimates them for high self-fertilisation frequencies. It is unclear why this mismatch arises. One explanation could be that drift effects are magnified under self–fertilisation, which causes a quicker sweep fixation time than expected from deterministic spread, if conditioning on a sweep going to fixation. Although *p*_0,*A*_ (Equation 8) captures these drift effects for rare alleles, there may be additional effects that are not accounted for. Under complete outcrossing, baseline diversity is restored (i.e., 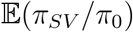 goes to 1) closer to the sweep origin for recessive mutations (*h* = 0.1), compared to semidominant (*h* = 0.5) or dominant (*h* = 0.9) mutations. Sweeps caused by dominant and semidominant mutations result in a similar genetic diversity, so these cases may be hard to differentiate from diversity data alone.

**Figure 3.**
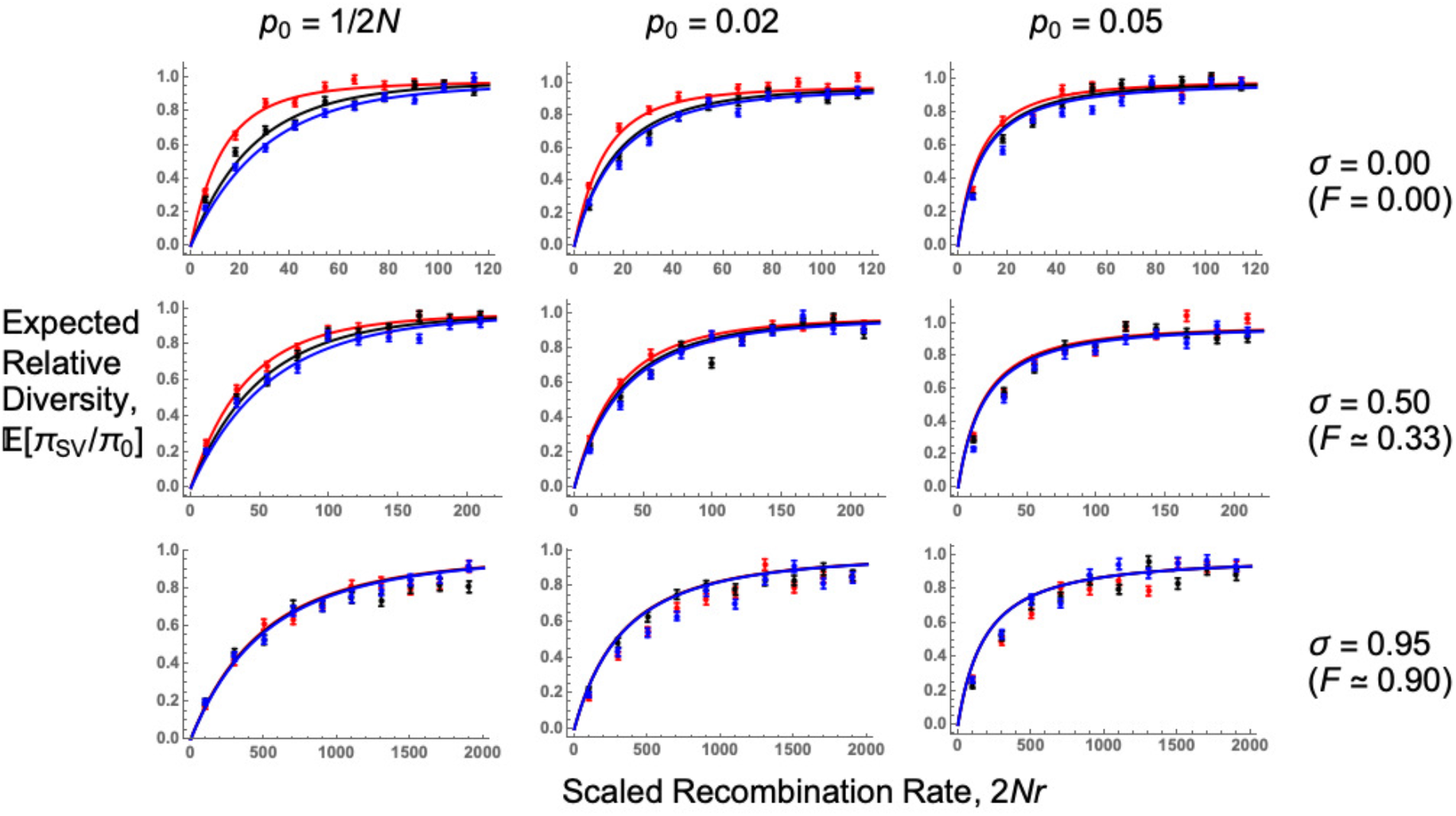
Expected relative pairwise diversity following a selective sweep. Plots of 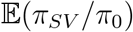 as a function of the recombination rate scaled to population size 2*Nr*. Lines are analytical solutions (Equation 9), points are forward-in-time simulation results. *N* = 5,000, *s* = 0.05, 4*Nμ* = 40 (note *μ* is scaled by *N*, not *N_e_*), and dominance coefficient *h* = 0.1 (red lines, points), 0.5 (black lines, points), or 0.9 (blue lines, points). Values of *p*_0_ and self-fertilisation rates *σ* used are shown for the relevant row and column; note the *x*—axis range changes with the self-fertilisation rate. For *p*_0_ = 1/2*N* we use *p*_0,*A*_ in our model, as given by Equation 8. Further results are plotted in Section C of Supplementary File S1.

These results can be better understood by examining the underlying allele trajectories, using logic described by Teshima and Przeworski (2006) (Figure 4). For outcrossing populations, recessive mutations spend most of the sojourn time at low frequencies, maximising recombination events and restoring neutral variation. These trajectories mimic sweeps from standing variation, which spend extended periods of time at low frequencies in the standing phase. Conversely, dominant mutations spend most of their time at high frequencies, reducing the chance for neutral markers to recombine onto the ancestral background.

**Figure 4.**
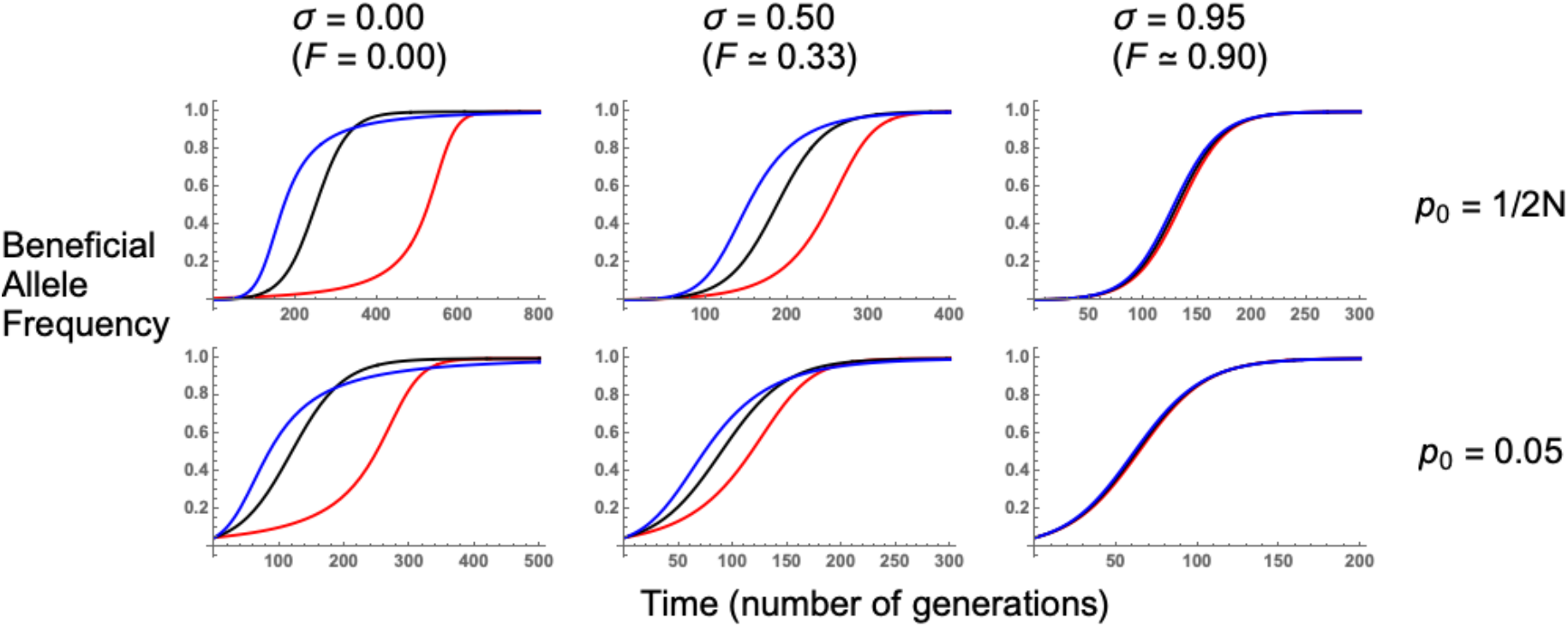
Beneficial allele trajectories. These were obtained by numerically evaluating the negative of Equation 4 forward in time. *N* = 5,000, *s* = 0.05, and *h* equals either 0.1 (red lines), 0.5 (black lines), or 0.9 (blue lines). Values of *p*_0_ and self-fertilisation rates *σ* used are shown for the relevant row and column. Note the different *x*—axis scales used in each panel. Further results are plotted in Section C of Supplementary File S1.

As self-fertilisation increases, sweep signatures become similar to the co-dominant case as the derived allele is more likely to spread as a homozygote, weakening the influence that dominance exerts over beneficial allele trajectories. Increasing *p*_0_ also causes sweeps with different dominance coefficients to produce comparable signatures, as beneficial mutation trajectories become similar after conditioning on starting at an elevated frequency.

#### Star–like approximation

An analytical approximation can be obtained by using the ‘star-like’ result for *P_NE_* (described in Supplementary Files S1, S2). In this case the expected pairwise diversity approximates to:

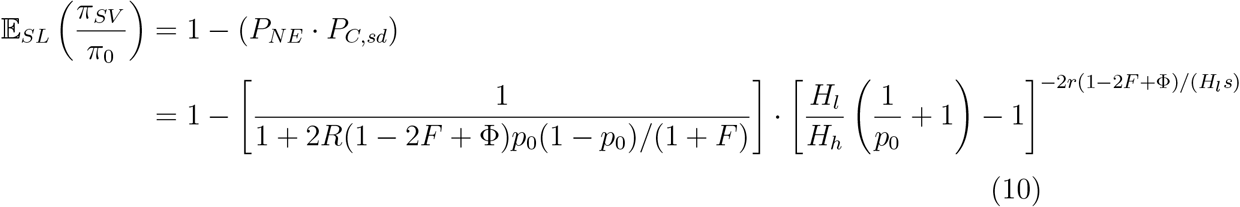

Note that Equation 10 instead uses the probability of coalescence during the standing phase, *P_C,sd_* = 1 − *P_R,sd_*. This approximation reflects similar formulas for diversity following soft sweeps in haploid outcrossing populations (Pennings and Hermisson 2006b; Berg and Coop 2015). There is a factor of two in the power term to account for two lineages. In Supplementary File S2 we demonstrate that this equation overestimates the relative diversity following a selective sweep. This mismatch arises since the star-like assumption of no coalescence during the sweep phase is only accurate for very strongly selected mutations (Barton 1998; B. Charlesworth, unpublished results). Hence it is important to consider coalescence during the sweep phase to accurately model selective sweeps that do not have an extremely high selection coefficient.

### Site Frequency Spectrum

The star-like approximation can be used to obtain analytical solutions for the number of segregating sites and the site frequency spectrum (i.e., the probability that *l* = 1, 2 … *n* − 1 of *n* alleles carry derived variants). The full derivation for these statistics are outlined in Supplementary File S2, which uses the star-like approximation. Figure 5 plots the SFS (Equation A12 in Supplementary File S2) alongside simulation results. Analytical results fit the simulation data well after including an adjusted singleton class, which accounts for recent mutations that arise on the derived background during both the standing and sweep phases (Berg and Coop 2015). Including this new singleton class improves the model fit, but there remains a tendency for analytical results to underestimate the proportion of low- and high-frequency classes (*l* = 1 and 9 in Figure 5), and overestimate the proportion of intermediate-frequency classes. Additional inaccuracies could have arisen due to the use of the star-like approximation, which assumes that there is no coalescence during the sweep phase.

**Figure 5.**
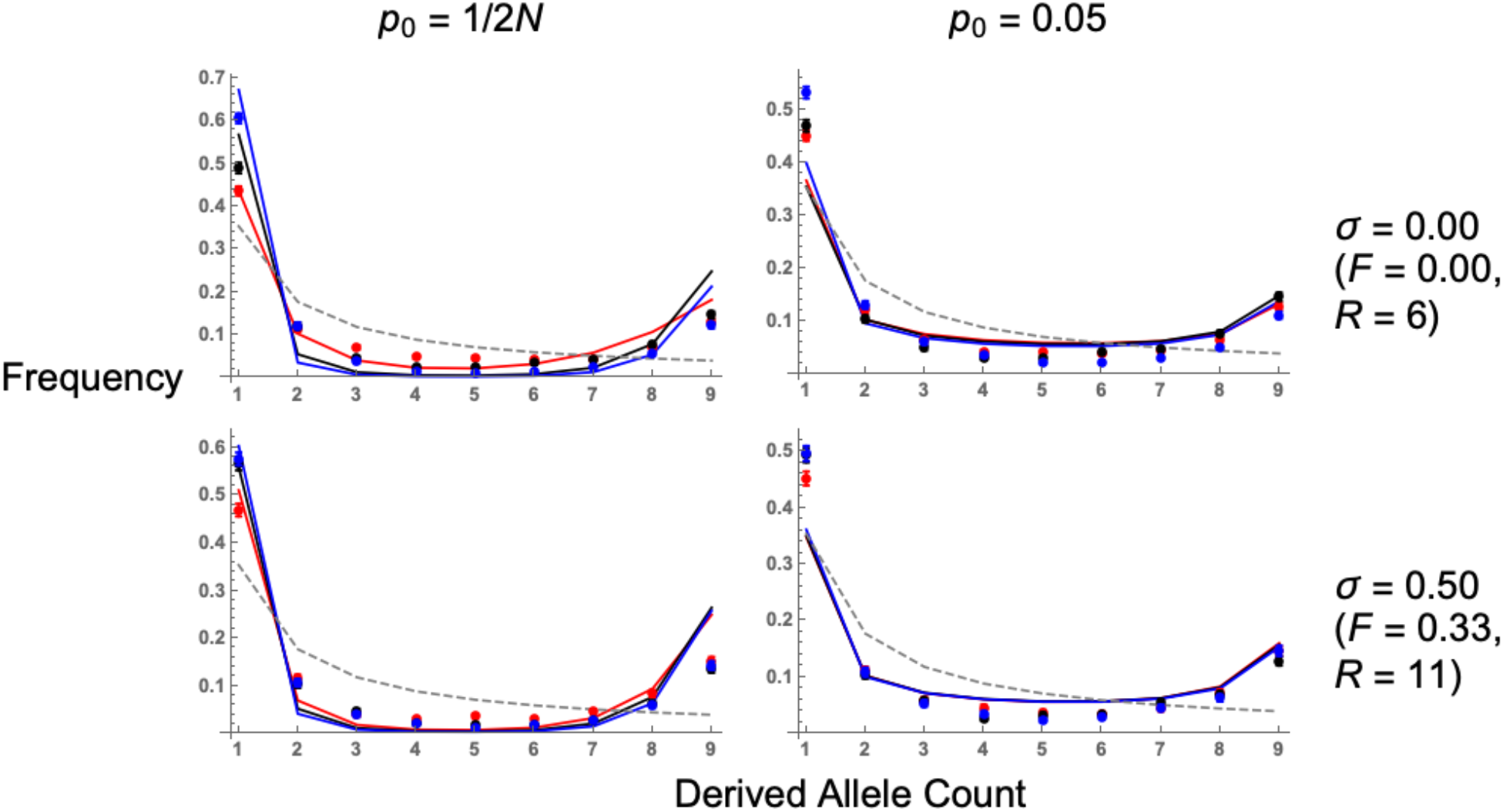
Expected site frequency spectrum, in flanking regions to the adaptive mutation, following a selective sweep. Lines are analytical solutions (Equation A12 in Supplementary File S2), points are simulation results. *N* = 5,000, *s* = 0.05, 4*Nμ* = 40, and dominance coefficient *h* = 0.1 (red lines, points), 0.5 (black lines, points), or 0.9 (blue lines, points). The neutral SFS is also included for comparisons (grey dashed line). Values of *p*_0_, self-fertilisation rates *σ* and recombination distances *R* are shown for the relevant row and column. Results for other recombination distances are in Section E of Supplementary File S1.

Hard sweeps in either outcrossers or partial selfers are characterised by a large number of singletons and highly-derived variants (Figure 5), which is a typical selective sweep signature (Braverman *et al*. 1995; Barton 1998; Kim and Stephan 2002). As the initial frequency *p*_0_ increases, so does the number of intermediate frequency variants (Figure 5). This signature is often seen as a characteristic of soft sweeps (Pennings and Hermisson 2006b; Berg and Coop 2015). Recessive hard sweeps (*h* = 0.1 and *p*_0_ = 1/2*N*) can produce SFS profiles that are similar to sweeps from standing variation, as there are an increased number of recombination events occurring since the allele is at a low frequency for long time periods (Figure 4). With increased self-fertilisation, both hard and soft sweep signatures (e.g., increased number of intermediate-frequency alleles) are recovered when measuring the SFS at a longer recombination distance than in outcrossers (Figure 5, bottom row). This is an example of how signatures of sweeps from standing variation are extended over an increased recombination distance of around 1/(1 − *σ*), as demonstrated by Equation 7.

### Soft sweeps from recurrent mutation

So far, we have only focussed on a soft sweep that arises from standing variation. An alternative type of soft sweep is one where recurrent mutation at the selected locus introduces the beneficial allele onto different genetic backgrounds. We can examine this case by modifying existing results. Below we derive the expected relative diversity between two alleles following this type of soft sweep, and outline the SFS for more than two samples in Supplementary File S2.

In this model, derived alleles arise from recurrent mutation and are instantaneously beneficial (i.e., there is no ‘standing phase’). During the sweep phase, lineages can escape the derived background by recombination, or if they are derived from a mutation event. If the beneficial allele is at frequency *p* then the probability of being descended from an ancestral allele by mutation is *P_m_*(*p*) = 2*μb*(1 − *p*)*/p*, for *μ_b_* the mutation probability (Pennings and Hermisson 2006b). Denote the probability of a lineage experiencing recombination or mutation during this sweep phase by *P_R,sw_*, *P_M,sw_* respectively. In both these cases the expected diversity present at linked sites is *π*_0_. If none of these events arise with probability *P_NE_*, then remaining lineages can either coalesce, or they arise from independent mutation events. If they coalesce then they have approximately zero pairwise diversity between them; alternatively, they have different origins and thus exhibit the same pairwise diversity *π*_0_ as the neutral background. Let *P_M,sd_* denote the probability that mutation occurs at the sweep origin, as opposed to coalescence.

Following this logic, the expected relative diversity for a sweep arising from recurrent mutation equals (with additional details in Supplementary File S1):

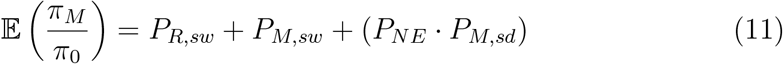

*π_M_* denotes the diversity around a soft sweep from recurrent mutation. *P_R,sw_*, *P_NE_* are similar to the equations used when modelling a sweep from standing variation. They are both modified to account for additional beneficial mutation arising during the sweep phase:

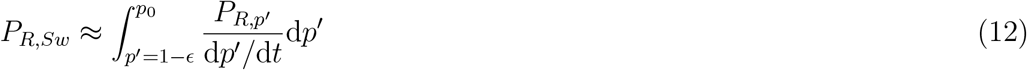

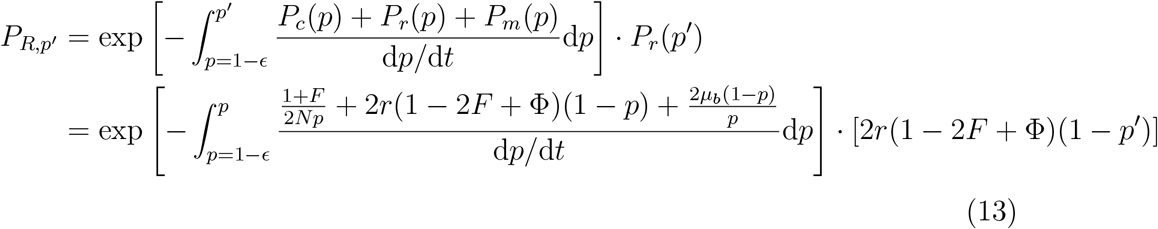

and:

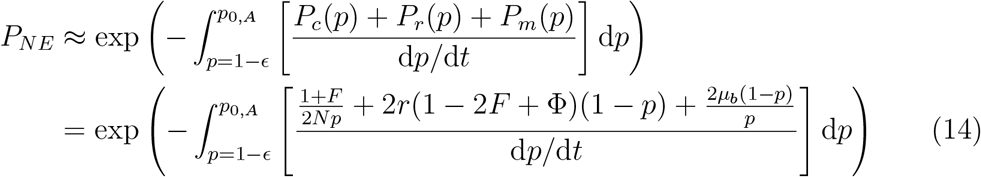

Note that Equation 14 has an upper integral limit of *p*_0,*A*_, as opposed to a general *p*_0_ used in the sweep from standing variation model, reflecting that there is no standing phase.

*P_M,sw_* is the mutation probability during the sweep phase, and is similar to Equation 13 except that 2*r*(1 − 2*F* + Φ)(1 − *p*′) is replaced by 2*μ_b_*(1 − *p*′)/*p*′, for *p*′ is the derived allele frequency when the event occurs. *PM,sd* is the probability that, at the sweep origin, the derived allele appears by mutation instead of coalescing, and is defined in a similar manner to *P_R,sd_* (Equation 7):

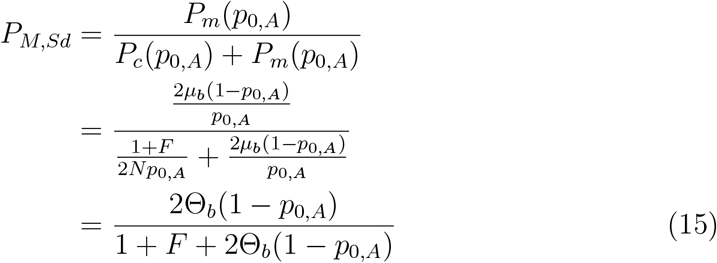

where Θ_*b*_ = 2*Nμ_b_*. The coalescence probability is 1 − *P_M,Sd_*. Equation 15 implies that self–fertilisation makes it more likely for beneficial mutations to coalesce at the start of a sweep, rather than arising from independent mutation events. Hence the signatures of soft sweeps via recurrent mutation will be weakened under inbreeding.

Figure 6 compares 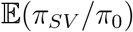 in the standing variation case, and 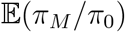 for the recurrent mutation case, under different levels of self-fertilisation. While dominance only weakly affects sweep signatures arising from standing variation under outcrossing, it more strongly affects sweeps from recurrent mutation in outcrossing populations, as each variant arises from an initial frequency close to 1/(2*N*) (Figure 4). Second, the two models exhibit different behaviour close to the selected locus (*R* close to zero). The recurrent mutation model has non–zero diversity levels, while the standing variation model exhibits zero diversity. As *R* increases, diversity eventually becomes higher for the standing variation case compared to the recurrent mutation case. We can heuristically determine when this transition occurs as follows. Assume a large population size but weak recombination and mutation rates. Hence, it is unlikely that any events occur during the sweep phase, so *P_R,sw_, P_M,sw_* ≈ 0 and *P_NE_* ≈ 1. Then the expected relative diversity (Equation 11) equals *P_R,sd_* for a sweep from standing variation, and *P_M,sd_* for one from recurrent mutation. To find the recombination rate *R_lim_* at which a sweep from recurrent mutation yields higher diversity than one from standing variation, we find the *R* value needed to equate the two probabilities, giving:

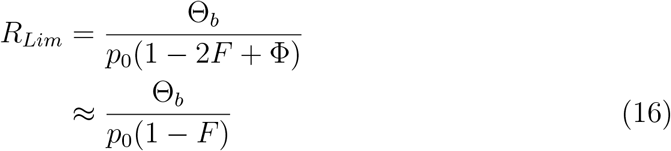

**Figure 6.**
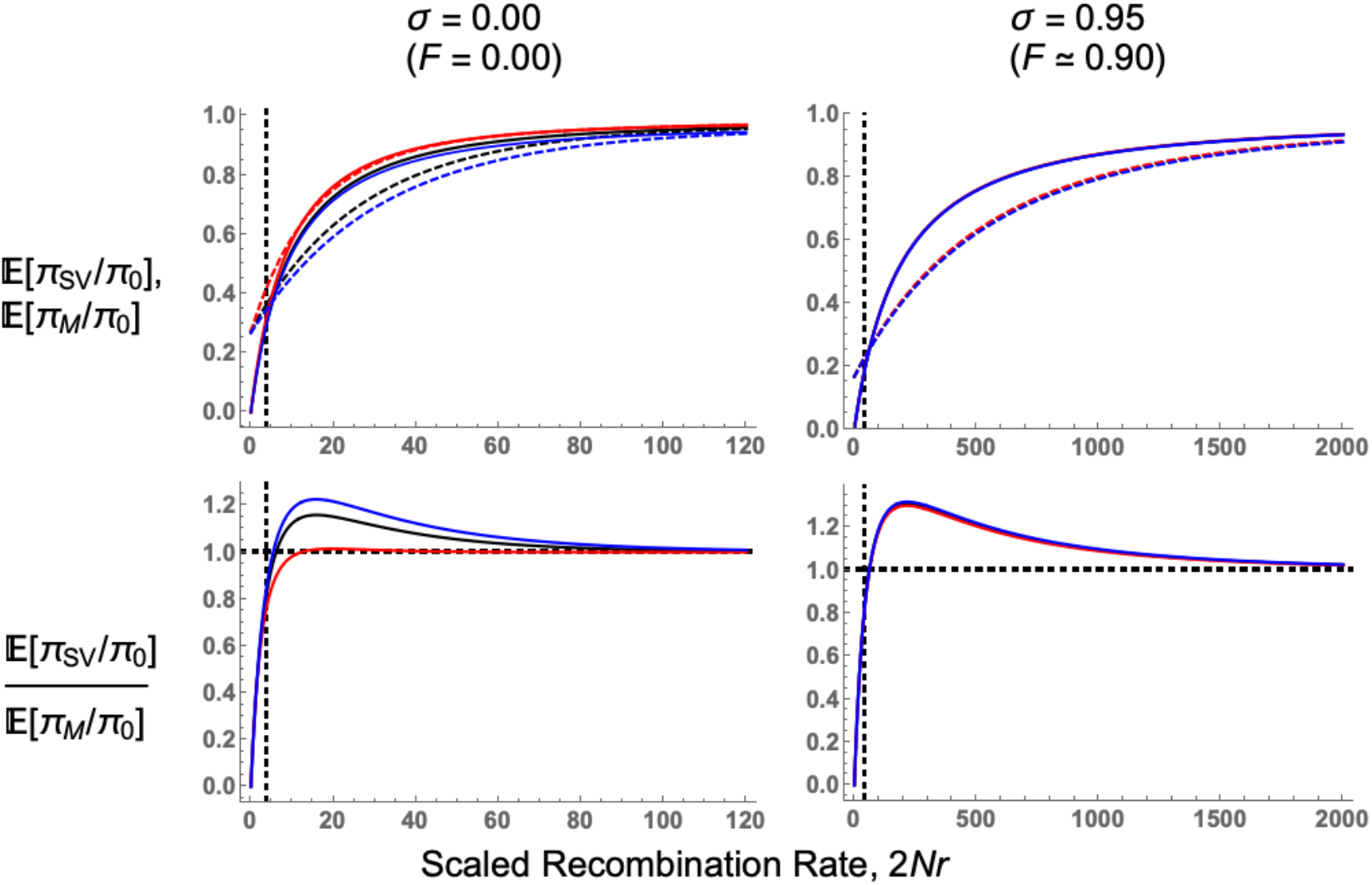
Comparing sweeps from recurrent mutation to those from standing variation. Top row: comparing relative diversity following a soft sweep, from either standing variation (Equation 9 with *p*_0_ = 0.05, solid lines) or recurrent mutation (using Equation 11 with Θ_*b*_ = 0.2, dashed lines). *N* = 5,000, *s* = 0.05, and dominance coefficient *h* = 0.1 (red lines), 0.5 (black lines), or 0.9 (blue lines). Bottom row: the ratio of the diversity following a sweep from standing variation to one from recurrent mutation. Parameters for each panel are as in the respective plot for the top row. Vertical dashed black line indicates *R_Lim_* (the approximate form of Equation 16); horizontal dashed line in the bottom-row plots show when the ratio equals 1. Note the different *x*—axis between left- and right-hand panels. Results are also plotted in Section F of Supplementary File S1.

The last approximation arises as Φ ≈ *F*. Hence for a fixed Θ_*b*_, the window where recurrent mutations create higher diversity near the selected locus increases for lower *p*_0_ or higher *F*, since both these factors reduces the potential for recombination to create new haplotypes during the standing phase. Equation 16 is generally accurate when sweeps from standing variation have higher diversity than sweeps with recurrent mutations (Figure 6, bottom row), but becomes inaccurate for *h* = 0.1 in outcrossing populations, as some events are likely to occur during the sweep phase. In Supplementary File S2 we show how similar results apply to the SFS.

## Discussion

### Summary of Theoretical Findings

While there has been many investigations into how different sweep processes can be detected from next-generation sequence data (Pritchard and Di Rienzo 2010; Messer and Petrov 2013; Stephan 2016; Hermisson and Pennings 2017), these models generally assumed idealised randomly mating populations and beneficial mutations that are semidominant (*h* = 0.5). Here we have created a more general selective sweep model, with arbitrary self-fertilisation and dominance levels. Our principal focus is on comparing a hard sweep arising from a single allele copy to a soft sweep arising from standing variation, but we also consider the case of recurrent mutation (Figure 6).

We find that the qualitative patterns of different selective sweeps under selfing remain similar to expectations from outcrossing models. In particular, a sweep from standing variation still creates an elevated number of intermediate-frequency variants compared to a sweep from *de novo* mutation (Figures 5, 6). This pattern is standard for soft sweeps (Pennings and Hermisson 2006b; Messer and Petrov 2013;Berg and Coop 2015; Hermisson and Pennings 2017) so existing statistical methods for detecting them (e.g., observing an higher than expected number of haplotypes; Vitti *et al*. (2013); Garud *et al*. (2015)) can, in principle, also be applied to selfing organisms. Under self-fertilisation, these signatures are stretched over longer physical regions than in outcrossers. These extensions arise as self-fertilisation affects gene genealogies during both the sweep and standing phases in different ways. During the sweep phase, beneficial alleles fix more rapidly under higher self-fertilisation as homozygous mutations are created more rapidly (Charlesworth 1992; Glémin 2012). In addition, the effective recombination rate is reduced by approximately 1 − *F* (Nordborg *et al*. 1996; Nordborg 2000; Charlesworth and Charlesworth 2010), and slightly more for highly inbred populations (Roze 2009, 2016). These two effects mean that neutral variants linked to an adaptive allele are less likely to recombine onto the neutral background during the sweep phase, as reflected in Equation 3 for *P_NE_*. During the standing phase, two haplotypes are more likely to coalesce under high levels of self-fertilisation since *N_e_* is decreased by a factor 1/(1 + *F*) (Pollak 1987; Charlesworth 1992; Caballero and Hill 1992; Nordborg and Donnelly 1997). This effect, combined with a reduced effective recombination rate, means that the overall recombination probability during the standing phase is reduced by a factor (1−*σ*) (Equation 7). Hence intermediate-frequency variants, which could provide evidence of adaptation from standing variation, will be spread out over longer genomic regions (this result can be seen in the site–frequency spectrum results, Figure 5). The elongation of sweep signatures means sweeps from standing variation can be easier to detect in selfing organisms than in outcrossers. Conversely, sweeps from recurrent mutation will have weakened signatures under self–fertilisation. This result is due to a reduced effective population size, making it likelier that lineages trace back to a common ancestor rather than independent mutation events.

We have also investigated how dominance affects soft sweep signatures, since previous analyses have only focussed on how dominance affects hard sweeps (Teshima and Przeworski 2006; Teshima *et al*. 2006; Ewing *et al*. 2011). In outcrossing organisms, recessive mutations leave weaker sweep signatures than additive or dominant mutations as they spend more time at low frequencies, increasing the amount of recombination that restores neutral variation (Figures 3, 4). With increased self-fertilisation, dominance has a weaker impact on sweep signatures as most mutations are homozygous (Figure 4). We also show that the SFS for recessive alleles can resemble a soft sweep, with a higher number of intermediate-frequency variants than for other hard sweeps (Figure 5). Dominance only weakly affects sweeps from standing variation, as trajectories of beneficial alleles become similar once the variant’s initial frequency exceeds 1/(2*N*) (Figures 3, 4). Yet different dominance levels can affect sweep signatures if the beneficial allele is reintroduced by recurrent mutation (Figure 6). Hence if one wishes to understand how dominance affects sweep signatures, it is also important to consider which processes underlie observed patterns of genetic diversity.

These results also demonstrate that the effects of dominance on sweeps are not necessarily intuitive. For example, both highly dominant and recessive mutations have elongated fixation times compared to co–dominant mutations (Glémin 2012). Based on this intuition, one could expect both dominant and recessive mutations to both produce weaker sweep signatures than co-dominant ones. In practice, dominant mutations have similar sweep signatures to co–dominant mutations (Figures 3, 5), and recessive sweeps could produce similar signatures to sweeps to standing variation (Figure 5). Dominance also has a weaker impact on sweeps on standing variation (Figures 3, 5).

### Soft sweeps from recurrent mutation or standing variation?

These theoretical results shed light onto how to distinguish between soft sweeps that arise either from standing variation, or from recurrent mutation. Both models are characterised by an elevated number of intermediate-frequency variants, in comparison to a hard sweep. Yet sweeps arising from recurrent mutation have non–zero diversity at the selected locus, whereas a sweep from standing variation exhibits approximately zero diversity. Hence a sweep from recurrent mutation shows intermediate-frequency variants closer to the beneficial locus, compared to sweeps from standing variation (Figures 6 and C in Supplementary File S2). Further from the selected locus, a sweep from standing variation exhibits greater variation than one from recurrent mutation, due to recombinant haplotypes being created during the standing phase. Equation 16 provides a simple condition for *R_Lim_*, the recombination distance needed for a sweep from standing variation to exhibit higher diversity than one from recurrent mutation; from this equation, we see that the size of this region increases under higher self-fertilisation. Hence it may be easier to differentiate between these two sweep scenarios in self–fertilising organisms.

Differences in haplotype structure between sweeps from either standing variation or recurrent mutation should be more pronounced in self-fertilising organisms, due to the reduction in effective recombination rates. However, when investigating sweep patterns over broad genetic regions, it becomes likelier that genetic diversity will be affected by multiple beneficial mutations spreading throughout the genome. Competing selective sweeps can lead to elevated diversity near a target locus for two reasons. First, selection interference increases the fixation time of individual mutations, allowing more recombination that can restore neutral diversity (Kim and Stephan 2003). In addition, competing selective sweeps can drag different sets of neutral variation to fixation, creating asymmetric diversity levels around a substitution (Chevin *et al*. 2008). Further investigations of selective sweep patterns across long genetic distances will prove to be a rich area of future research.

Finally, we have assumed a fixed population size, and that sweeps from standing variation arose from neutral variation. The resulting signatures could differ if the population size has changed over time, or if the beneficial allele was previously deleterious. Both issues could also affect our ability to discriminate between soft and hard sweeps.

### Potential applications to self-fertilising organisms

Existing methods for finding sweep signatures in nucleotide polymorphism data are commonly based on finding regions with a site-frequency spectrum matching what is expected under a selective sweep (Nielsen *et al*. 2005; Boitard *et al*. 2009; Pavlidis *et al*. 2013; DeGiorgio *et al*. 2016; Huber *et al*. 2016). The more general models developed here can be used to create more specific sweep-detection methods that include self-fertilisation. However, a recent analysis found that soft-sweep signatures can be incorrectly inferred if analysing genetic regions that flank hard sweeps, which was named the ‘soft shoulder’ effect (Schrider *et al*. 2015). Due to the reduction in recombination in selfers, these model results indicate that ‘soft-shoulder’ footprints can arise over long genetic distances and should be taken into account. One remedy to this problem is to not just classify genetic regions as being subject to either a hard or soft sweep, but also as being linked to a region subject to one of these sweeps (Schrider and Kern 2016). These more general calculations can also be extended to quantify to what extent background selection and sweeps jointly shape genome-wide diversity in self-fertilising organisms (Elyashiv *et al*. 2016; Campos *et al*. 2017; Booker and Keightley 2018; Rettelbach *et al*. 2019), or detect patterns of introgression (Setter *et al*. 2019).

## Acknowledgments

We would like to thank Sally Otto for providing information on the elevated effective starting frequency of beneficial mutations; Brian Charlesworth on providing advice on modelling selective sweeps, sharing unpublished results, and providing comments on the manuscript; Ben Haller for answering questions about SLiM; Nick Barton and other anonymous referees for providing feedback on the manuscript. MH was supported by a Marie Curie International Outgoing Fellowship (MC-IOF-622936) and a NERC Independent Research Fellowship (NE/R015686/1). MH and TB also acknowledge financial support from the European Research Council under the European Union’s Seventh Framework Program (FP7/20072013, ERC Grant 311341).

