## Supplemental File S1 for "Selective sweeps under dominance and inbreeding": File_S1.pdf

##### Clearing workspace and setting up libraries

Note this notebook uses the “ErrorBarPlots” package, which has to be installed before use.

```
ClearAll["`*"];
Needs["ErrorBarPlots`"]
```

#### Section A: Derivation of $\mathbb{E}[\pi_{SV}/\pi_0]$ while considering coalescence during the sweep phase

In this scenario, there are four possible outcomes:
(1) Coalescence during the sweep phase;
(2) Recombination during the sweep phase;
(3) Coalescence during the standing phase;
(4) Recombination during the standing phase.

If events (1), (3) occur then $\pi \simeq 0$.
If events (2), (4) occur then $\pi \simeq \pi_0$, the background levels of diversity.
Hence $\mathbb{E}(\pi/\pi_0)$ = P(Event 2) + P(Event 4).

Going through these in turn;
To calculate P(Event 2), we need to determine the probabilities that (i) no event (coalescence or recombination) occurs in the sweep phase when the derived allele is between a frequency of $p$ to 1; (ii) recombination occurs at frequency $p$; (iii) integrating this solution over all frequencies $p0$ to 1.

To calculate P(Event 4), we need to determine the probabilities that (i) no event (coalescence or recombination) occurs in the sweep phase when the derived allele is between a frequency of *p0* to 1; (ii) recombination occurs during the standing phase where the derived allele is at a fixed frequency of *p0*.

Let's look at the relative probability of each event over each timestep.
If the frequency of the beneficial allele is *p* at a certain time, then the probability of coalescence is $\frac{1}{2\,\mathrm{Ne}\,p} = \frac{1+F}{2Np}$. The probability of one of the two samples recombining out is 2r(1-2F+Φ)(1-p).

The probability of no action occurring at any timepoint is $1 - \frac{1}{2\,\mathrm{Ne}\,p}$ – 2r(1-2F+Φ)(1-p). The total probability of neither action over the entire sweep phase is
$\simeq \mathrm{Exp}\left[-\int_{t=0}^{\tau f}\left(\frac{1+F}{2Np} + 2r(1-2F+\Phi)(1-p)\right)dt\right]$ for fixation time τf, or
$\int_{p=1-\epsilon 1}^{p0} \frac{\left(\frac{1+F}{2Np} + 2r(1-2F+\Phi)\,(1-p)\right)}{dp/dt}\,dp$.

The sum of the coalescence and recombination probabilities are:

```
2 r (1 - 2 F + Φ) (1 - p) + (1 + F)/(2 Na p) // Together
```

$$\frac{1}{2\,\mathrm{Na}\,p}\left(1 + F + 4\,\mathrm{Na}\,p\,r - 8\,F\,\mathrm{Na}\,p\,r - 4\,\mathrm{Na}\,p^2\,r + 8\,F\,\mathrm{Na}\,p^2\,r + 4\,\mathrm{Na}\,p\,r\,\Phi - 4\,\mathrm{Na}\,p^2\,r\,\Phi\right)$$

Tidying up:

```
(1 + F + 4 Na p (1 - p) (1 - 2 F + Φ) r)/(2 Na p) -
  {1/(2 Na p) (1 + F + 4 Na p r - 8 F Na p r - 4 Na p^2 r + 8 F Na p^2 r + 4 Na p r Φ - 4 Na p^2 r Φ)} //
  FullSimplify
```

{0}

Dividing by $-s\,(1-p)\,p\,(F + h - F\,h + (1-F)\,(1-2\,h)\,p)$ and taking the indefinite integral;

```
Integrate[-((1 + F + 4 Na p (1 - p) (1 - 2 F + Φ) r) /
    ((2 Na p) (s (1 - p) p (F + h - F h + (1 - F) (1 - 2 h) p)))), p] // Simplify
```

$$-\frac{1}{2\,Na\,s}\left(-\frac{1+F}{(F+h-F\,h)\,p}-\frac{(1+F)\,\mathrm{Log}[1-p]}{1+(-1+F)\,h}+\left(\left(-1+F+F^2\,(2-8\,Na\,r+h\,(-3+8\,Na\,r))+h\,(3+4\,Na\,r\,(1+\Phi))-4\,F\,Na\,r\,(-1-\Phi+h\,(3+\Phi))\right)\mathrm{Log}[p]\right)\Big/(F+h-F\,h)^2-\frac{1}{(1+(-1+F)\,h)\,(F+h-F\,h)^2}\left(-1+F^3\left(-1+h\,(4-8\,Na\,r)+h^2\,(-4+8\,Na\,r)\right)+4\,h\,(1+Na\,r\,(1+\Phi))-4\,h^2\,(1+Na\,r\,(1+\Phi))+F\left(1+4\,Na\,r\,(1+\Phi)-4\,h\,(1+2\,Na\,r\,(2+\Phi))+4\,h^2\,(1+2\,Na\,r\,(2+\Phi))\right)+F^2\left(1-8\,Na\,r+4\,h\,(-1+Na\,r\,(5+\Phi))-4\,h^2\,(-1+Na\,r\,(5+\Phi))\right)\right)\mathrm{Log}[F+h+p-F\,p-2\,h\,p+F\,h\,(-1+2\,p)]\right)$$

We can use this integral to calculate (A) the probability of recombination during the sweep phase; (B) the probability of recombination during the standing phase, given no actions during the sweep phase.

The total probability of no event occurring for frequency $p$ is the difference of this integral between p and 1 - $\epsilon$ (where $\epsilon$ is the effective fixed frequency of the derived allele). This term is then multiplied by (2 r (1-2F+Φ)(1-p)) to determine the probability that recombination acts at frequency *p.*

(A) can be calculated by integrating the total recombination probability between p0 and 1 - $\epsilon$, again after diving by $dp/dt$ to convert from a time integral into a frequency integral. It does not appear that this integral has an analytical solution, so this integral is instead integrated numerically.

(B) can be calculated by working out (i) the probability of no event acting between frequency *p*0 and 1 − $\epsilon$, and (ii) multiplying it by the probability that a recombination event takes precedence over coalescence in the standing phase.

(i) is obtained by taking the integral solution between p0 and 1. For (ii), the probability is equal to:

$$\frac{2\,r\,(1-2\,F+\Phi)\,(1-p0)}{\frac{1+F}{2\,Na\,p0}+2\,r\,(1-2\,F+\Phi)\,(1-p0)}\ \text{// FullSimplify}$$

$$(4\,Na\,(-1+p0)\,p0\,r\,(-1+2\,F-\Phi))\,/\,(1+F+8\,F\,Na\,(-1+p0)\,p0\,r-4\,Na\,(-1+p0)\,p0\,r\,(1+\Phi))$$

Which can be rewritten as:

$$\frac{4\,Na\,r\,(1-2\,F+\Phi)\,(1-p0)\,p0}{1+F+4\,Na\,r\,(1-2\,F+\Phi)\,(1-p0)\,p0}-\{(4\,Na\,(-1+p0)\,p0\,r\,(-1+2\,F-\Phi))\,/\,(1+F+8\,F\,Na\,(-1+p0)\,p0\,r-4\,Na\,(-1+p0)\,p0\,r\,(1+\Phi))\}\ \text{// Simplify}$$

{0}

#### Deriving the star–like approximation for $P_{NR}$

Next we derive $P_{NR} = \mathrm{Exp}[-r_{eff}\int_{1-\epsilon}^{p0}\frac{1-p}{-dp/dt}dp]$ for a small factor $\epsilon$ denoting the upper limit of the deterministic equation, and the starting allele frequency p0. Below is $\frac{1-p}{-dp/dt}$:

$$\frac{1-p}{-s\,(1-p)\,p\,(F+h-F\,h+(1-F)\,(1-2\,h)\,p)}\;\text{// Simplify}$$

$$-\frac{1}{p\,(F+h-F\,h+(-1+F)\,(-1+2\,h)\,p)\,s}$$

Separating out the fraction after removing the $\frac{1}{s}$ term:

$$\text{Apart}\left[-\frac{1}{p\,(F+h-F\,h+(-1+F)\,(-1+2\,h)\,p)}\right]$$

$$\frac{1}{(-F-h+F\,h)\,p}+(-1+F+2\,h-2\,F\,h)\,/\,((-F-h+F\,h)\,(F+h-F\,h+(1-F-2\,h+2\,F\,h)\,p))$$

The first fraction can be written as:

$$-\frac{1}{(F+h-F\,h)\,p}-\left\{\frac{1}{(-F-h+F\,h)\,p}\right\}\;\text{// Simplify}$$

{0}

The second fraction can be written as:

$$\frac{1-F-2\,h\,(1-F)}{(F+h-F\,h)\,(F+h-F\,h+(1-F)\,(1-2\,h)\,p)}-\{(-1+F+2\,h-2\,F\,h)\,/\,((-F-h+F\,h)\,(F+h-F\,h+(1-F-2\,h+2\,F\,h)\,p))\}\;\text{// FullSimplify}$$

{0}

Integrating the first fraction over p:

$$\text{Integrate}\left[-\frac{1}{(F+h-F\,h)\,p},\,p\right]$$

$$-\frac{\text{Log}[p]}{F+h-F\,h}$$

Integrating the second fraction over p, after tidying it up:

```
(1 - 2 h - F (1 - 2 h)) / ((F + h - F h) (F + h - F h + (1 - 2 h - F (1 - 2 h)) p)) -
   {(-1 + F + 2 h - 2 F h) / ((-F - h + F h) (F + h - F h + (1 - F - 2 h + 2 F h) p))} //
  FullSimplify
```

{0}

```
Integrate[(-1 + F + 2 h - 2 F h) / ((-F - h + F h) (F + h - F h + (1 - F - 2 h + 2 F h) p)), p] //
  FullSimplify
```

$$\frac{\text{Log}[F+h-F\,h+(-1+F)\,(-1+2\,h)\,p]}{F+h-F\,h}$$

Putting the two parts together and tidying up:

$$-\frac{\text{Log}[p]}{F+h-F h}+\frac{\text{Log}[F+h-F h+(-1+F)(-1+2 h) p]}{F+h-F h} \text{ // FullSimplify}$$

$$-\frac{\text{Log}[p]-\text{Log}[F+h-F h+(-1+F)(-1+2 h) p]}{F+h-F h}$$

$$\frac{\text{Log}[F+h-F h+(1-F)(1-2 h) p]-\text{Log}[p]}{F+h-F h}-\left\{-\frac{1}{F+h-F h}(\text{Log}[p]-\text{Log}[F+h-F h+(-1+F)(-1+2 h) p])\right\} \text{ // FullSimplify}$$

$$\{0\}$$

Integrating between p0 and 1–$\epsilon$:

$$\left(\frac{1}{F+h-F h}(\text{Log}[F+h-F h+(1-F)(1-2 h) p]-\text{Log}[p]) \text{ /. } p\to p0\right)-\left(\frac{1}{F+h-F h}(\text{Log}[F+h-F h+(1-F)(1-2 h) p]-\text{Log}[p])\right) \text{ /. } p\to 1-\epsilon \text{ // FullSimplify}$$

$$-\frac{1}{F+h-F h}(\text{Log}[p0]-\text{Log}[F+h-F h+(-1+F)(-1+2 h) p0]-\text{Log}[1-\epsilon]+\text{Log}[1+(-1+F)\epsilon+h(-1+F+2\epsilon-2 F\epsilon)])$$

This solution can be rewritten as a single Log term:

$$\frac{1}{F+h-F h} * \text{Log}\left[\frac{(F+h-F h-(1-F)(1-2 h) p0)(1-\epsilon)}{p0(1-(1-F)(\epsilon+h(1-2\epsilon)))}\right]$$

$$\frac{\text{Log}\left[\frac{(F+h-F h-(1-F)(1-2 h) p0)(1-\epsilon)}{p0(1-(1-F)(h(1-2\epsilon)+\epsilon))}\right]}{F+h-F h}$$

$P_{NR}$ follows by multiplying by $-\frac{r_{eff}}{s}$ and taking the exponential:

$$\text{Exp}\left[-\frac{r_{eff}}{s(F+h-F h)} \text{Log}\left[\frac{(F+h-F h-(1-F)(1-2 h) p0)(1-\epsilon)}{p0(1-(1-F)(\epsilon+h(1-2\epsilon)))}\right]\right] \text{ // Simplify}$$

$$(((F+h-F h-(-1+F)(-1+2 h) p0)(1-\epsilon))/(p0-(1-F) p0(h+\epsilon-2 h\epsilon)))^{-\frac{r_{eff}}{(F+h-F h) s}}$$

We can write this equation in a neater form by using HL = F + h - Fh, HH = 1 - h + Fh:

```
Solve[HL == F + h - F h && HH == 1 - h + F h, {F, h}] // FullSimplify
```

$$\left\{\left\{F\to -1+HH+HL,\ h\to \frac{-1+HH}{-2+HH+HL}\right\}\right\}$$

$$(((F+h-F h-(-1+F)(-1+2 h) p0)(1-\epsilon))/(p0-(1-F) p0(h+\epsilon-2 h\epsilon)))^{-\frac{r_{eff}}{(F+h-F h) s}} \text{ /. } \left\{F\to -1+HH+HL,\ h\to \frac{-1+HH}{-2+HH+HL}\right\} \text{ // FullSimplify}$$

$$\left(\frac{(HL-HH\ p0+HL\ p0)(-1+\epsilon)}{p0(HH(-1+\epsilon)-HL\ \epsilon)}\right)^{-\frac{r_{eff}}{HL\ s}}$$

Tidying up:

$$\left(\frac{(HL + p0\,(HL - HH))\,(1-\epsilon)}{p0\,(HH + \epsilon\,(HL - HH))}\right)^{-\frac{r_{eff}}{HL\,s}} - \left\{\left(\frac{(HL - HH\,p0 + HL\,p0)\,(-1+\epsilon)}{p0\,(HH\,(-1+\epsilon) - HL\,\epsilon)}\right)^{-\frac{r_{eff}}{HL\,s}}\right\} \text{ // Simplify}$$

{0}

If using $\epsilon = 0$ this solution simplifies to:

$$\left(\frac{(HL + p0\,(HL - HH))\,(1-\epsilon)}{p0\,(HH + \epsilon\,(HL - HH))}\right)^{-\frac{r_{eff}}{HL\,s}} \text{ /. } \epsilon \to 0$$

$$\left(\frac{HL + (-HH + HL)\,p0}{HH\,p0}\right)^{-\frac{r_{eff}}{HL\,s}}$$

Which can be written in the desired form after some rewriting:

$$\frac{HL + (-HH + HL)\,p0}{HH\,p0} \text{ // Apart}$$

$$-\frac{HH - HL}{HH} + \frac{HL}{HH\,p0}$$

$$-\frac{HH - HL}{HH} \text{ // Apart}$$

$$-1 + \frac{HL}{HH}$$

$$\left(\frac{HL}{HH}\left(\frac{1}{p0} + 1\right) - 1\right)^{-\frac{r_{eff}}{HL\,s}} - \left\{\left(\frac{HL + (-HH + HL)\,p0}{HH\,p0}\right)^{-\frac{r_{eff}}{HL\,s}}\right\} \text{ // Simplify}$$

{0}

### Section B: Derivation of effective starting frequency from a *de novo* mutation

This code was based on a *Mathematica* notebook provided by Sarah Otto (University of British Columbia).

#### Original haploid derivation

Following Ewens' (2004) derivation of the fixation probability in his book for the haploid model we have:

```
ϕ[x_] = Exp[-Simplify[Integrate[2 ((1 - x) x ψ) / ((1 - x) x), x]]]
```

$$e^{-2\,x\,\psi}$$

where $\psi$=Ns and

```
Sx[x_] = Simplify[Integrate[ϕ[x], x]]
```

$$-\frac{e^{-2\,x\,\psi}}{2\,\psi}$$

which are used in the fixation probability (Eq. 4.17):

```
fix[p0_, ψ_] = Simplify[(Sx[x0] - Sx[0]) / (Sx[1] - Sx[0]) /. x0 → p0]
```

$$\frac{e^{2\psi}\left(1-e^{-2\,p0\,\psi}\right)}{-1+e^{2\psi}}$$

The time spent at frequency x < p (conditional on fixation) is then given by (4.22):

```
Simplify[2 (1 - fix[p0, ψ]) / (x (1 - x) ϕ[x]) Integrate[ϕ[y], {y, 0, x}]]
```

$$\frac{e^{-2\,p0\,\psi}\left(-e^{2\psi}+e^{2\,p0\,\psi}\right)\left(-1+e^{2\,x\,\psi}\right)}{\left(-1+e^{2\psi}\right)(-1+x)\,x\,\psi}$$

while the time spent at frequency x > p (conditional on fixation) is given by (4.23)

```
Simplify[2 fix[p0, ψ] / (x (1 - x) ϕ[x]) Integrate[ϕ[y], {y, x, 1}]]
```

$$\frac{e^{-2\,p0\,\psi}\left(-1+e^{2\,p0\,\psi}\right)\left(-e^{2\psi}+e^{2\,x\,\psi}\right)}{\left(-1+e^{2\psi}\right)(-1+x)\,x\,\psi}$$

Assuming p0=1/N, only the latter is relevant (the system cannot spend time below this frequency if it will fix).

By comparison, the time spent at frequency x in the deterministic process is given by: $\frac{1}{\psi\,(1-x)\,x}$

(this is obtained by rearranging dx/dt = sx(1-x) and then measuring time in units of N generations).

A simple estimate of the acceleration is given by taking the time spent at x=p0 in the diffusion process and equating this to the time in the deterministic process at an accelerated position x=$\alpha$ p0.

```
e^(-2 p0 ψ) (-1 + e^(2 p0 ψ)) (-e^(2 ψ) + e^(2 x ψ)) / ((-1 + e^(2 ψ)) (-1 + x) x ψ) /. x → p0
```

$$\frac{e^{-2\,p0\,\psi}\left(-1+e^{2\,p0\,\psi}\right)\left(-e^{2\psi}+e^{2\,p0\,\psi}\right)}{\left(-1+e^{2\psi}\right)(-1+p0)\,p0\,\psi}$$

```
(1 / (ψ (1 - x) x) /. x → α p0)
```

$$\frac{1}{p0\,\alpha\,(1-p0\,\alpha)\,\psi}$$

Given that p0 is very small, the latter can be approximated as $\frac{1}{p0\,\alpha\,\psi}$, allowing us to solve for the acceleration:

```
Solve[e^(-2 p0 ψ) (-1 + e^(2 p0 ψ)) (-e^(2 ψ) + e^(2 p0 ψ)) / ((-1 + e^(2 ψ)) (-1 + p0) p0 ψ) == 1 / (p0 α ψ), α] // Flatten
```

$$\left\{\alpha\to\frac{e^{2\,p0\,\psi}\left(-1+e^{2\psi}\right)(-1+p0)}{\left(-1+e^{2\,p0\,\psi}\right)\left(-e^{2\psi}+e^{2\,p0\,\psi}\right)}\right\}$$

To leading order in p0, this is:

```
Normal[Series[(α /. %), {p0, 0, -1}]]
```

$$\frac{1}{2\ p0\ \psi}$$

Setting p0=1/N and $\psi$=Ns, we get that the acceleration is approximately equivalent to starting at an allele frequency that is $\frac{1}{2\ s}$ times higher than the initial allele frequency, as used in the simulations of Otto and Barton (1997).

$$\frac{1}{2\ p0\ \psi}\ /.\ \left\{p0 \to \frac{1}{Na},\ \psi \to Na\ s\right\}$$

$$\frac{1}{2\ s}$$

#### Diploid derivation (including dominance and selfing)

Following Ewens' (2004) derivation of the fixation probability in his book for the haploid model we have:

```
ϕ[x_] = Exp[
  -Simplify[Integrate[2 ((1 - x) x (F + h - F h + (1 - F) (1 - 2 h) x) ψ) / ((1 - x) x), x]]]
```

$$e^{-x\ (2\ F+2\ h-2\ F\ h+(-1+F)\ (-1+2\ h)\ x)\ \psi}$$

where $\psi = 2\ Ne\ s$ and

```
Sx[x_] = Simplify[Integrate[ϕ[x], x]]
```

$$\left(e^{\frac{(F+h-F\ h)^2\ \psi}{(-1+F)\ (-1+2\ h)}}\ \sqrt{\pi}\ \mathrm{Erf}\left[\left((F+h+x-F\ x-2\ h\ x+F\ h\ (-1+2\ x))\ \sqrt{\psi}\right)\Big/\left(\sqrt{-1+F}\ \sqrt{-1+2\ h}\right)\right]\right)\Big/\left(2\ \sqrt{-1+F}\ \sqrt{-1+2\ h}\ \sqrt{\psi}\right)$$

which are used in the fixation probability (Eq 4.17):

```
fix[p0_, ψ_] =
 Simplify[Integrate[ϕ[x], {x, 0, x0}] / Integrate[ϕ[x], {x, 0, 1}] /. x0 → p0]
```

$$\left(-\mathrm{Erf}\left[\frac{(F+h-F\ h)\ \sqrt{\psi}}{\sqrt{-1+F}\ \sqrt{-1+2\ h}}\right]+\mathrm{Erf}\left[\frac{(F+h-F\ h+(-1+F)\ (-1+2\ h)\ p0)\ \sqrt{\psi}}{\sqrt{-1+F}\ \sqrt{-1+2\ h}}\right]\right)\Big/\left(\mathrm{Erf}\left[\frac{(1+(-1+F)\ h)\ \sqrt{\psi}}{\sqrt{-1+F}\ \sqrt{-1+2\ h}}\right]-\mathrm{Erf}\left[\frac{(F+h-F\ h)\ \sqrt{\psi}}{\sqrt{-1+F}\ \sqrt{-1+2\ h}}\right]\right)$$

However, Pfix here is undefined for $F = 1$ and/or $h = \frac{1}{2}$. As with the outcrossing case an approximation can be obtained by instead using $e^{-2\ (F+h-F\ h)\ x\ \psi}$ to obtain Pfix = $\frac{-1+e^{2\ (F\ (-1+h)-h)\ p0\ \psi}}{-1+e^{2\ (F\ (-1+h)-h)\ \psi}}$. Note this equation becomes inaccurate for x0 >> $\frac{1}{2N}$ (see Glémin 2012 Theor. Popul. Biol. for a more complete derivation).

The time spent at frequency $x < p$ (conditional on fixation) is then given by (4.22):

```
Simplify[2 (1 - fix[p0, ψ])/(x (1 - x) ϕ[x]) Integrate[ϕ[y], {y, 0, x}]]
```

$$\left(e^{\frac{(F+h+x-F\,x-2\,h\,x+F\,h\,(-1+2\,x))^2\,\psi}{(-1+F)\,(-1+2\,h)}}\sqrt{\pi}\left(\mathrm{Erf}\left[\frac{(1+(-1+F)\,h)\,\sqrt{\psi}}{\sqrt{-1+F}\,\sqrt{-1+2\,h}}\right]-\mathrm{Erf}\left[\left((F+h-F\,h+(-1+F)\,(-1+2\,h)\,p0)\,\sqrt{\psi}\right)\Big/\left(\sqrt{-1+F}\,\sqrt{-1+2\,h}\right)\right]\right)\left(\mathrm{Erf}\left[\frac{(F+h-F\,h)\,\sqrt{\psi}}{\sqrt{-1+F}\,\sqrt{-1+2\,h}}\right]-\mathrm{Erf}\left[\left((F+h-F\,h+(-1+F)\,(-1+2\,h)\,x)\,\sqrt{\psi}\right)\Big/\left(\sqrt{-1+F}\,\sqrt{-1+2\,h}\right)\right]\right)\right)\Big/\left(\sqrt{-1+F}\,\sqrt{-1+2\,h}\,(-1+x)\,x\,\sqrt{\psi}\left(\mathrm{Erf}\left[\frac{(1+(-1+F)\,h)\,\sqrt{\psi}}{\sqrt{-1+F}\,\sqrt{-1+2\,h}}\right]-\mathrm{Erf}\left[\frac{(F+h-F\,h)\,\sqrt{\psi}}{\sqrt{-1+F}\,\sqrt{-1+2\,h}}\right]\right)\right)$$

while the time spent at frequency x > p (conditional on fixation) is given by (4.23)

```
Simplify[2 fix[p0, ψ]/(x (1 - x) ϕ[x]) Integrate[ϕ[y], {y, x, 1}]]
```

$$\left(e^{\frac{(F+h+x-F\,x-2\,h\,x+F\,h\,(-1+2\,x))^2\,\psi}{(-1+F)\,(-1+2\,h)}}\sqrt{\pi}\left(\mathrm{Erf}\left[\frac{(F+h-F\,h)\,\sqrt{\psi}}{\sqrt{-1+F}\,\sqrt{-1+2\,h}}\right]-\mathrm{Erf}\left[\left((F+h-F\,h+(-1+F)\,(-1+2\,h)\,p0)\,\sqrt{\psi}\right)\Big/\left(\sqrt{-1+F}\,\sqrt{-1+2\,h}\right)\right]\right)\left(\mathrm{Erf}\left[\frac{(1+(-1+F)\,h)\,\sqrt{\psi}}{\sqrt{-1+F}\,\sqrt{-1+2\,h}}\right]-\mathrm{Erf}\left[\left((F+h-F\,h+(-1+F)\,(-1+2\,h)\,x)\,\sqrt{\psi}\right)\Big/\left(\sqrt{-1+F}\,\sqrt{-1+2\,h}\right)\right]\right)\right)\Big/\left(\sqrt{-1+F}\,\sqrt{-1+2\,h}\,(-1+x)\,x\,\sqrt{\psi}\left(\mathrm{Erf}\left[\frac{(1+(-1+F)\,h)\,\sqrt{\psi}}{\sqrt{-1+F}\,\sqrt{-1+2\,h}}\right]-\mathrm{Erf}\left[\frac{(F+h-F\,h)\,\sqrt{\psi}}{\sqrt{-1+F}\,\sqrt{-1+2\,h}}\right]\right)\right)$$

Assuming p0=1/2N, only the latter is relevant (the system cannot spend time below this frequency if it will fix).

By comparison, the time spent at frequency x in the deterministic process is given by:

$$\frac{1}{((1-x)\,x\,(F+h-F\,h+(1-F)\,(1-2\,h)\,x)\,\psi)}$$

A simple estimate of the acceleration is given by taking the time spent at x=p0 in the diffusion process and equating this to the time in the deterministic process at an accelerated position x=$\alpha$ p0.

$$\left(e^{\frac{(F+h+x-F\,x-2\,h\,x+F\,h\,(-1+2\,x))^2\,\psi}{(-1+F)\,(-1+2\,h)}}\sqrt{\pi}\left(\mathrm{Erf}\left[\frac{(F+h-F\,h)\sqrt{\psi}}{\sqrt{-1+F}\,\sqrt{-1+2\,h}}\right]-\mathrm{Erf}\left[\left((F+h-F\,h+(-1+F)\,(-1+2\,h)\,p0)\sqrt{\psi}\right)\Big/\left(\sqrt{-1+F}\,\sqrt{-1+2\,h}\right)\right]\right)\left(\mathrm{Erf}\left[\frac{(1+(-1+F)\,h)\sqrt{\psi}}{\sqrt{-1+F}\,\sqrt{-1+2\,h}}\right]-\mathrm{Erf}\left[\left((F+h-F\,h+(-1+F)\,(-1+2\,h)\,x)\sqrt{\psi}\right)\Big/\left(\sqrt{-1+F}\,\sqrt{-1+2\,h}\right)\right]\right)\right)\Big/\left(\sqrt{-1+F}\,\sqrt{-1+2\,h}\,(-1+x)\,x\,\sqrt{\psi}\left(\mathrm{Erf}\left[\frac{(1+(-1+F)\,h)\sqrt{\psi}}{\sqrt{-1+F}\,\sqrt{-1+2\,h}}\right]-\mathrm{Erf}\left[\frac{(F+h-F\,h)\sqrt{\psi}}{\sqrt{-1+F}\,\sqrt{-1+2\,h}}\right]\right)\right)\;/.\;x\to p0$$

$$\left(e^{\frac{(F+h+p0-F\,p0-2\,h\,p0+F\,h\,(-1+2\,p0))^2\,\psi}{(-1+F)\,(-1+2\,h)}}\sqrt{\pi}\left(\mathrm{Erf}\left[\frac{(1+(-1+F)\,h)\sqrt{\psi}}{\sqrt{-1+F}\,\sqrt{-1+2\,h}}\right]-\mathrm{Erf}\left[\frac{(F+h-F\,h+(-1+F)\,(-1+2\,h)\,p0)\sqrt{\psi}}{\sqrt{-1+F}\,\sqrt{-1+2\,h}}\right]\right)\left(\mathrm{Erf}\left[\frac{(F+h-F\,h)\sqrt{\psi}}{\sqrt{-1+F}\,\sqrt{-1+2\,h}}\right]-\mathrm{Erf}\left[\frac{(F+h-F\,h+(-1+F)\,(-1+2\,h)\,p0)\sqrt{\psi}}{\sqrt{-1+F}\,\sqrt{-1+2\,h}}\right]\right)\right)\Big/\left(\sqrt{-1+F}\,\sqrt{-1+2\,h}\,(-1+p0)\,p0\,\sqrt{\psi}\left(\mathrm{Erf}\left[\frac{(1+(-1+F)\,h)\sqrt{\psi}}{\sqrt{-1+F}\,\sqrt{-1+2\,h}}\right]-\mathrm{Erf}\left[\frac{(F+h-F\,h)\sqrt{\psi}}{\sqrt{-1+F}\,\sqrt{-1+2\,h}}\right]\right)\right)$$

$$\left(\frac{1}{(1-x)\,x\,(F+h-F\,h+(1-F)\,(1-2\,h)\,x)\,\psi}\;/.\;x\to\alpha\,p0\right)$$

$$1/(p0\,\alpha\,(1-p0\,\alpha)\,(F+h-F\,h+(1-F)\,(1-2\,h)\,p0\,\alpha)\,\psi)$$

```
Series[1 / (p0 α (1 - p0 α) (F + h - F h + (1 - F) (1 - 2 h) p0 α) ψ), {p0, 0, -1}]
```

$$\frac{1}{(F+h-F\,h)\,\alpha\,\psi\,p0}+O[p0]^0$$

Given that p0 is very small, the latter can be approximated as $\frac{1}{(F+h-F\,h)\,\alpha\,\psi\,p0}$, allowing us to solve for the acceleration:

```
Solve[(e^(((F+h+p0-F p0-2 h p0+F h (-1+2 p0))^2 ψ)/((-1+F) (-1+2 h))) Sqrt[π] (Erf[((1 + (-1 + F) h) Sqrt[ψ])/(Sqrt[-1 + F] Sqrt[-1 + 2 h])] -
      Erf[((F + h - F h + (-1 + F) (-1 + 2 h) p0) Sqrt[ψ])/(Sqrt[-1 + F] Sqrt[-1 + 2 h])])
    (Erf[((F + h - F h) Sqrt[ψ])/(Sqrt[-1 + F] Sqrt[-1 + 2 h])] - Erf[((F + h - F h + (-1 + F) (-1 + 2 h) p0) Sqrt[ψ])/
      (Sqrt[-1 + F] Sqrt[-1 + 2 h])]))/(Sqrt[-1 + F] Sqrt[-1 + 2 h] (-1 + p0)
    p0 Sqrt[ψ] (Erf[((1 + (-1 + F) h) Sqrt[ψ])/(Sqrt[-1 + F] Sqrt[-1 + 2 h])] - Erf[((F + h - F h) Sqrt[ψ])/(Sqrt[-1 + F] Sqrt[-1 + 2 h])])) ==
  1/((F + h - F h) α ψ p0), α] // Flatten
```

```
{α → (Sqrt[-1 + F] Sqrt[-1 + 2 h] Erf[((1 + (-1 + F) h) Sqrt[ψ])/(Sqrt[-1 + F] Sqrt[-1 + 2 h])] -
    Sqrt[-1 + F] Sqrt[-1 + 2 h] p0 Erf[((1 + (-1 + F) h) Sqrt[ψ])/(Sqrt[-1 + F] Sqrt[-1 + 2 h])] - Sqrt[-1 + F] Sqrt[-1 + 2 h]
    Erf[((F + h - F h) Sqrt[ψ])/(Sqrt[-1 + F] Sqrt[-1 + 2 h])] + Sqrt[-1 + F] Sqrt[-1 + 2 h] p0 Erf[((F + h - F h) Sqrt[ψ])/(Sqrt[-1 + F] Sqrt[-1 + 2 h])])/
   (-e^(((F+h+p0-F p0-2 h p0+F h (-1+2 p0))^2 ψ)/((-1+F) (-1+2 h))) F Sqrt[π] Sqrt[ψ] Erf[((1 + (-1 + F) h) Sqrt[ψ])/(Sqrt[-1 + F] Sqrt[-1 + 2 h])] Erf[((F + h - F h) Sqrt[ψ])/(Sqrt[-1 + F] Sqrt[-1 + 2 h])] -
    e^(((F+h+p0-F p0-2 h p0+F h (-1+2 p0))^2 ψ)/((-1+F) (-1+2 h))) h Sqrt[π] Sqrt[ψ] Erf[((1 + (-1 + F) h) Sqrt[ψ])/(Sqrt[-1 + F] Sqrt[-1 + 2 h])] Erf[((F + h - F h) Sqrt[ψ])/(Sqrt[-1 + F] Sqrt[-1 + 2 h])] +
    e^(((F+h+p0-F p0-2 h p0+F h (-1+2 p0))^2 ψ)/((-1+F) (-1+2 h))) F h Sqrt[π] Sqrt[ψ] Erf[((1 + (-1 + F) h) Sqrt[ψ])/(Sqrt[-1 + F] Sqrt[-1 + 2 h])] Erf[
      ((F + h - F h) Sqrt[ψ])/(Sqrt[-1 + F] Sqrt[-1 + 2 h])] + e^(((F+h+p0-F p0-2 h p0+F h (-1+2 p0))^2 ψ)/((-1+F) (-1+2 h))) F Sqrt[π] Sqrt[ψ] Erf[((1 + (-1 + F) h) Sqrt[ψ])/(Sqrt[-1 + F] Sqrt[-1 + 2 h])]
    Erf[((F + h - F h + (-1 + F) (-1 + 2 h) p0) Sqrt[ψ])/(Sqrt[-1 + F] Sqrt[-1 + 2 h])] +
    e^(((F+h+p0-F p0-2 h p0+F h (-1+2 p0))^2 ψ)/((-1+F) (-1+2 h))) h Sqrt[π] Sqrt[ψ] Erf[((1 + (-1 + F) h) Sqrt[ψ])/(Sqrt[-1 + F] Sqrt[-1 + 2 h])]
    Erf[((F + h - F h + (-1 + F) (-1 + 2 h) p0) Sqrt[ψ])/(Sqrt[-1 + F] Sqrt[-1 + 2 h])] -
    e^(((F+h+p0-F p0-2 h p0+F h (-1+2 p0))^2 ψ)/((-1+F) (-1+2 h))) F h Sqrt[π] Sqrt[ψ] Erf[((1 + (-1 + F) h) Sqrt[ψ])/(Sqrt[-1 + F] Sqrt[-1 + 2 h])]
    Erf[((F + h - F h + (-1 + F) (-1 + 2 h) p0) Sqrt[ψ])/(Sqrt[-1 + F] Sqrt[-1 + 2 h])] +
    e^(((F+h+p0-F p0-2 h p0+F h (-1+2 p0))^2 ψ)/((-1+F) (-1+2 h))) F Sqrt[π] Sqrt[ψ] Erf[((F + h - F h) Sqrt[ψ])/(Sqrt[-1 + F] Sqrt[-1 + 2 h])]
```

$$\mathrm{Erf}\left[\left((F+h-F\,h+(-1+F)\,(-1+2\,h)\,p0)\,\sqrt{\psi}\right)\Big/\left(\sqrt{-1+F}\,\sqrt{-1+2\,h}\right)\right]+$$

$$e^{\frac{(F+h+p0-F\,p0-2\,h\,p0+F\,h\,(-1+2\,p0))^2\,\psi}{(-1+F)\,(-1+2\,h)}}\,h\,\sqrt{\pi}\,\sqrt{\psi}\,\mathrm{Erf}\left[\frac{(F+h-F\,h)\,\sqrt{\psi}}{\sqrt{-1+F}\,\sqrt{-1+2\,h}}\right]$$

$$\mathrm{Erf}\left[\left((F+h-F\,h+(-1+F)\,(-1+2\,h)\,p0)\,\sqrt{\psi}\right)\Big/\left(\sqrt{-1+F}\,\sqrt{-1+2\,h}\right)\right]-$$

$$e^{\frac{(F+h+p0-F\,p0-2\,h\,p0+F\,h\,(-1+2\,p0))^2\,\psi}{(-1+F)\,(-1+2\,h)}}\,F\,h\,\sqrt{\pi}\,\sqrt{\psi}\,\mathrm{Erf}\left[\frac{(F+h-F\,h)\,\sqrt{\psi}}{\sqrt{-1+F}\,\sqrt{-1+2\,h}}\right]$$

$$\mathrm{Erf}\left[\left((F+h-F\,h+(-1+F)\,(-1+2\,h)\,p0)\,\sqrt{\psi}\right)\Big/\left(\sqrt{-1+F}\,\sqrt{-1+2\,h}\right)\right]-$$

$$e^{\frac{(F+h+p0-F\,p0-2\,h\,p0+F\,h\,(-1+2\,p0))^2\,\psi}{(-1+F)\,(-1+2\,h)}}\,F\,\sqrt{\pi}\,\sqrt{\psi}$$

$$\mathrm{Erf}\left[\left((F+h-F\,h+(-1+F)\,(-1+2\,h)\,p0)\,\sqrt{\psi}\right)\Big/\left(\sqrt{-1+F}\,\sqrt{-1+2\,h}\right)\right]^2-$$

$$e^{\frac{(F+h+p0-F\,p0-2\,h\,p0+F\,h\,(-1+2\,p0))^2\,\psi}{(-1+F)\,(-1+2\,h)}}\,h\,\sqrt{\pi}\,\sqrt{\psi}$$

$$\mathrm{Erf}\left[\left((F+h-F\,h+(-1+F)\,(-1+2\,h)\,p0)\,\sqrt{\psi}\right)\Big/\left(\sqrt{-1+F}\,\sqrt{-1+2\,h}\right)\right]^2+$$

$$e^{\frac{(F+h+p0-F\,p0-2\,h\,p0+F\,h\,(-1+2\,p0))^2\,\psi}{(-1+F)\,(-1+2\,h)}}\,F\,h\,\sqrt{\pi}\,\sqrt{\psi}$$

$$\left.\mathrm{Erf}\left[\left((F+h-F\,h+(-1+F)\,(-1+2\,h)\,p0)\,\sqrt{\psi}\right)\Big/\left(\sqrt{-1+F}\,\sqrt{-1+2\,h}\right)\right]^2\right)\Bigg\}$$

To leading order in p0, this is:

```
Normal[Series[(α /. %), {p0, 0, -1}]]
```

$$-\frac{1}{2\,(-F-h+F\,h)\,p0\,\psi}$$

Rewriting as:

$$\frac{1}{2\,(F+h-F\,h)\,p0\,\psi}-\left\{-\frac{1}{2\,(-F-h+F\,h)\,p0\,\psi}\right\}\ \texttt{// FullSimplify}$$

{0}

Setting p0 = 1/2N and $\psi = 2\,Ne\,s = \frac{2Ns}{1+F}$, we get that the acceleration is approximately equivalent to starting at an allele frequency that exhibits the following boost:

$$\frac{1}{2\,(F+h-F\,h)\,p0\,\psi}\ \texttt{/.}\ \left\{p0\to\frac{1}{2\,Na},\ \psi\to\frac{2\,Na\,s}{1+F}\right\}$$

$$\frac{1+F}{2\,(F+h-F\,h)\,s}$$

In the limit of no selfing (F = 0) this boost reduces to the diploid outcrossing result:

$$\frac{1+F}{2\,(F+h-F\,h)\,s}\ \texttt{/.}\ \{F\to 0\}$$

$$\frac{1}{2\,h\,s}$$

With complete selfing the result reduces to 1/s:

```
(1 + F)/(2 (F + h - F h) s) /. {F → 1}
```

$$\frac{1}{s}$$

Furthermore for additive dominance, the accelerative effect equals 1/s for all F values, which is the 'establishment frequency' obtained by Desai and Fisher (2007). Beneficial alleles spread deterministically once above this frequency in their haploid model.

```
(1 + F)/(2 (F + h - F h) s) /. {h → 1/2} // FullSimplify
```

$$\frac{1}{s}$$

#### Consistency with Martin and Lambert (2015)

Martin and Lambert (2015 Theor. Pop. Biol.) used a Feller diffusion to demonstrate that a selective sweep the originated in *n* copies has an effective starting frequency drawn from a Gamma distribution with shape parameter *n* and scale parameter $1/[2N(1-e^{-2N_e s(h+F-hF)/N})]$, or $1/[2N(1-e^{-2s(h+F-hF)/(1+F)})]$ if assuming $N_e = \frac{N}{1+F}$.

The mean of such a distribution is the product of the shape and scale parameters, i.e. $1/[2N(1-e^{-2s(h+F-hF)/(1+F)})]$. This value is approximately (1/2Nx) for $x = \frac{2(F+h-Fh)s}{1+F}$ if s << 1 (see code below). Hence their results are consistent with the above results, where the deterministic starting frequency is elevated by a factor $\frac{1+F}{2(F+h-Fh)s}$.

```
Series[1/(2 N (1 - Exp[-(2 (F+h-F h) s)/(1+F)])), {s, 0, -1}]
```

$$\frac{1+F}{4(F+h-Fh)Ns} + O[s]^0$$

#### Example plot of boost

```
Boostp0[Na_, s_, h_, F_] := (1 + F)/(4 Na s (F + h - F h))
```

Red, black, blue lines are boosting values for $h = 0.1, 0.5, 0.9$ respectively, plotted below as a function of F. The effect is strongest for recessive mutants in outcrossers.

```
BoostPlot = Legended[Plot[{Boostp0[5000, 0.05, 0.1, F],
    Boostp0[5000, 0.05, 0.5, F], Boostp0[5000, 0.05, 0.9, F]}, {F, 0, 1},
   PlotStyle → {{Red, Thick}, {Black, Thick}, {Blue, Thick}}, FrameLabel →
    {"Inbreeding coefficient, F", "Effective\nStarting\nFrequency,\np0,A"},
   Frame → True, Axes → True, RotateLabel → {False, True} ,
   LabelStyle → {FontFamily → "Arial", FontSize → 20, FontColor → Black},
   PlotRange → {Automatic, {-0.0005, 0.0105}}, GridLines → {None, {1/10 000}},
   GridLinesStyle → {Thick, Dashed}, ImageSize → 750],
  LineLegend[{Directive[{Red, Thick}], Directive[{Black, Thick}],
    Directive[{Blue, Thick}], Directive[{Black, Dashed, Thick}]},
   {"p0,A, h = 0.1", "p0,A, h = 0.5", "p0,A, h = 0.9", "1/2N"},
   LabelStyle → {FontSize → 20}]]
```

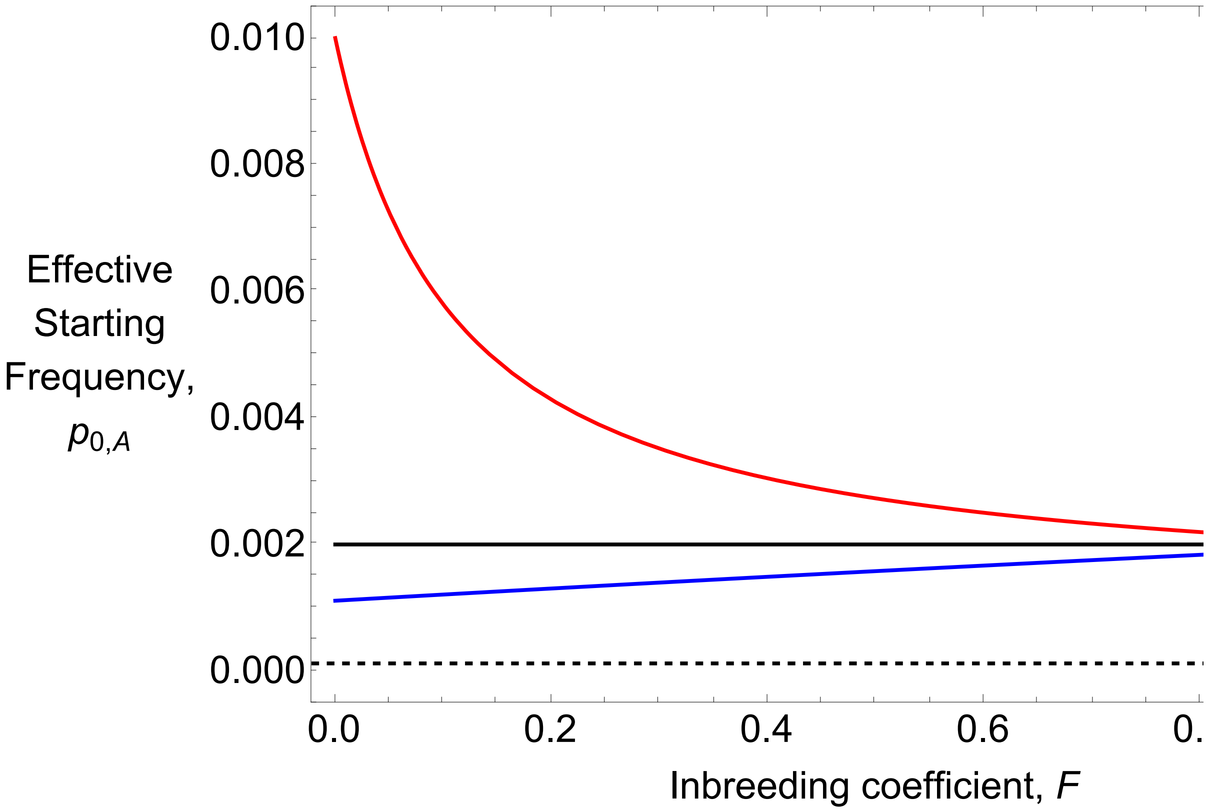


### Section C: Simulation comparisons, $\mathbb{E}[\pi_{SV}/\pi_0]$ and allele trajectory plots

#### Equations

Here are the equations for *F*, Φ at steady state, as a function of the self–fertilisation fraction and recombination probability:

$$F[\sigma\_] := \frac{\sigma}{2 - \sigma}$$

$$\Phi[r\_, \sigma\_] := \frac{\sigma\ (2 - \sigma - 2\ (1 - r)\ r\ (2 - 3\ \sigma))}{(2 - \sigma)\ (2 - (1 - 2\ (1 - r)\ r)\ \sigma)}$$

Below are the ‘star-like’ approximation results.

PWR2B[Na_, s_, h_, σ_, R_, p0_] :=

$$\left(1 - \left(\frac{1}{1 + \frac{2 R}{1 + F[\sigma]}\ (1 - F[\sigma])\ p0\ (1 - p0)}\right)\left(\left(\frac{(F[\sigma] + h - F[\sigma]\ h)}{(1 - h + F[\sigma]\ h)}\left(\frac{1}{p0} + 1\right) - 1\right)^{-\frac{R\ (1 - F[\sigma])}{Na\ (F[\sigma] + h - F[\sigma]\ h)\ s}}\right)\right)$$

The following is the ‘effective’ starting frequency of a beneficial allele at initial frequency $\frac{1}{2\,\mathrm{Na}}$, given that it goes to fixation:

$$\mathrm{Boostp0}[Na\_, s\_, h\_, \sigma\_] := \frac{1 + F[\sigma]}{4\ Na\ s\ (F[\sigma] + h - F[\sigma]\ h)}$$

Below are the equations for the solution that accounts for coalescence in the sweep phase:

SweepIntFH[Na_, s_, h_, F_, Φ_, r_, p_] :=

$$-\frac{1}{2\ Na\ s}\left(-\frac{1 + F}{(F + h - F\ h)\ p} - \frac{(1 + F)\ \mathrm{Log}[1 - p]}{1 + (-1 + F)\ h} + \left(\left(-1 + F + F^2\ (2 - 8\ Na\ r + h\ (-3 + 8\ Na\ r)) + h\ (3 + 4\ Na\ r\ (1 + \Phi)) - 4\ F\ Na\ r\ (-1 - \Phi + h\ (3 + \Phi))\right)\ \mathrm{Log}[p]\right) / (F + h - F\ h)^2 - \frac{1}{(1 + (-1 + F)\ h)\ (F + h - F\ h)^2}\left(-1 + F^3\left(-1 + h\ (4 - 8\ Na\ r) + h^2\ (-4 + 8\ Na\ r)\right) + 4\ h\ (1 + Na\ r\ (1 + \Phi)) - 4\ h^2\ (1 + Na\ r\ (1 + \Phi)) + F\left(1 + 4\ Na\ r\ (1 + \Phi) - 4\ h\ (1 + 2\ Na\ r\ (2 + \Phi)) + 4\ h^2\ (1 + 2\ Na\ r\ (2 + \Phi))\right) + F^2\left(1 - 8\ Na\ r + 4\ h\ (-1 + Na\ r\ (5 + \Phi)) - 4\ h^2\ (-1 + Na\ r\ (5 + \Phi))\right)\right)\ \mathrm{Log}[F + h + p - F\ p - 2\ h\ p + F\ h\ (-1 + 2\ p)]\right)$$

```
PRecPFH[Na_, s_, h_, F_, Φ_, r_, p_] := (2 r (1 - 2 F + Φ) (1 - p))
  Exp[-(SweepIntFH[Na, s, h, F, Φ, r, p] -
      SweepIntFH[Na, s, h, F, Φ, r, 1 - Boostp0[Na, s, 1 - h, F]])]
PRecFHp0[Na_, s_, h_, F_, Φ_, r_, p0_] := NIntegrate[PRecPFH[Na, s, h, F, Φ, r, p] /
   (s (1 - p) p (F + h - F h + (1 - F) (1 - 2 h) p)), {p, p0, 1 - Boostp0[Na, s, 1 - h, F]}]
PNoActFH[Na_, s_, h_, F_, Φ_, r_, p0_] := Exp[-(SweepIntFH[Na, s, h, F, Φ, r, p0] -
     SweepIntFH[Na, s, h, F, Φ, r, 1 - Boostp0[Na, s, 1 - h, F]])]
PRecp0FH[Na_, s_, h_, F_, Φ_, r_, p0_] := PNoActFH[Na, s, h, F, Φ, r, p0] *
```

$$\left(\frac{4\ Na\ r\ (1 - 2\ F + \Phi)\ (1 - p0)\ p0}{1 + F + 4\ Na\ r\ (1 - 2\ F + \Phi)\ (1 - p0)\ p0}\right)$$

Here’s the equation for $\mathbb{E}[\pi/\pi_0]$, note it is a function of $\sigma$ rather than F or Φ

```
ExPiSV[Na_, s_, h_, σ_, r_, p0_] :=
 PRecp0FH[Na, s, h, F[σ], Φ[r, σ], r, p0] + PRecFHp0[Na, s, h, F[σ], Φ[r, σ], r, p0]
```

### Outcrossing case ($\sigma = F = 0$)

#### Simulation comparisons, from initial frequency $p_0 = 1/2N$

Loading Data

```
SetDirectory[NotebookDirectory[]];
Rin = Table[6 + 12 * i, {i, 0, 9}];
```

h = 0.5

SLiM simulation data

```
PiRelh05 = Import["SLIM_F0/StatsProc_R_120_h_0.5_self_0_f_1e-04_10b_SLIM.dat",
    "Table"][[10]];
PiRelh05CIB = Import[
    "SLIM_F0/StatsProc_R_120_h_0.5_self_0_f_1e-04_10b_SLIM.dat",
    "Table"][[11]];
PiRelh05CIT = Import[
    "SLIM_F0/StatsProc_R_120_h_0.5_self_0_f_1e-04_10b_SLIM.dat",
    "Table"][[12]];
PiRelh05S = Partition[Riffle[Partition[Riffle[Rin, PiRelh05], 2],
    Map[ErrorBar, Partition[Riffle[-PiRelh05CIB, PiRelh05CIT], 2]]], 2];
```

MSMS simulation data (Note that throughout, the ‘R’ file index value is twice that for SLiM. This is because MSMS uses a scaling of 4Nr, while custom SLiM code uses a scaling of 2Nr).

```
PiRelh05MSMS =
  Import["MSMS_SV_5K/StatsProc_R_240_h_0.5_self_0_f_1e-04_10b_MSMS.dat",
    "Table"][[10]];
PiRelh05MSMSCIB = Import[
    "MSMS_SV_5K/StatsProc_R_240_h_0.5_self_0_f_1e-04_10b_MSMS.dat",
    "Table"][[11]];
PiRelh05MSMSCIT = Import[
    "MSMS_SV_5K/StatsProc_R_240_h_0.5_self_0_f_1e-04_10b_MSMS.dat",
    "Table"][[12]];
PiRelh05MSMSS = Partition[Riffle[Partition[Riffle[Rin, PiRelh05MSMS], 2],
    Map[ErrorBar, Partition[Riffle[-PiRelh05MSMSCIB, PiRelh05MSMSCIT], 2]]], 2];
```

Analytical results;
From here on we use both the star-like results and the coalescence-in-sweep-phase results. Note that for the latter the input is unscaled recombination probability, so it has to be divided by 2N = 10,000 for these parameters. Unless stated otherwise, the x-axis is the scaled recombination rate 2Nr; the y-axis is $\mathbb{E}[\pi/\pi_0]$. Axes labels will be added at the end when plots are grouped together.

```
Ph05 = Plot[{PWR2B[5000, 0.05, 0.5, 0, R, Boostp0[5000, 0.05, 0.5, 0]],
   ExPiSV[5000, 0.05, 0.5, 0, R/10 000, Boostp0[5000, 0.05, 0.5, 0]]},
  {R, 0, 120}, PlotStyle → {{Black, Thick, Dashed}, {Black, Thick}},
  PlotRange → All, Frame → {{True, False}, {True, False}},
  BaseStyle → {FontWeight → "Bold", FontColor → Black, FontSize → 12}]
```

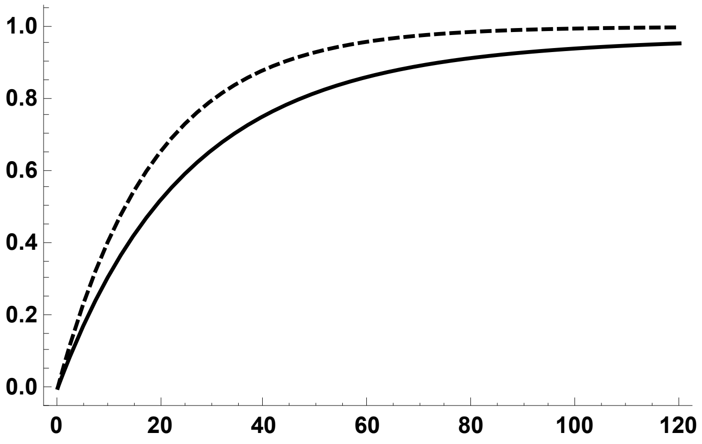


```
Ph05A = Plot[{ExPiSV[5000, 0.05, 0.5, 0, R/10 000, Boostp0[5000, 0.05, 0.5, 0]]},
   {R, 0, 120}, PlotStyle → {{Black, Thick}},
   PlotRange → {All, {0, 1.1}}, Frame → {{True, False}, {True, False}},
   BaseStyle → {FontWeight → "Bold", FontColor → Black, FontSize → 12}];
```

Plot of SLiM simulation data

```
Ph05S = ErrorListPlot[PiRelh05S,
  PlotStyle → {Black, PointSize[0.02]}, PlotRange → {{0, 120}, {0, 1.1}}]
```

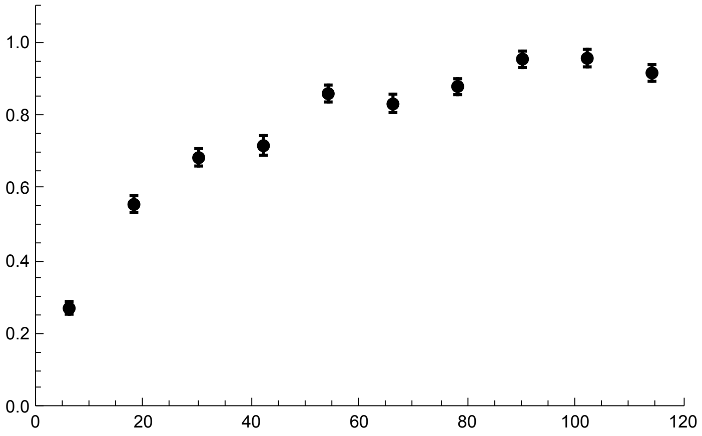


Plot of MSMS simulation data

```
Ph05MSMSS = ErrorListPlot[PiRelh05MSMSS,
  PlotStyle → {Gray, PointSize[0.02]}, PlotRange → {{0, 120}, {0, 1.1}}]
```

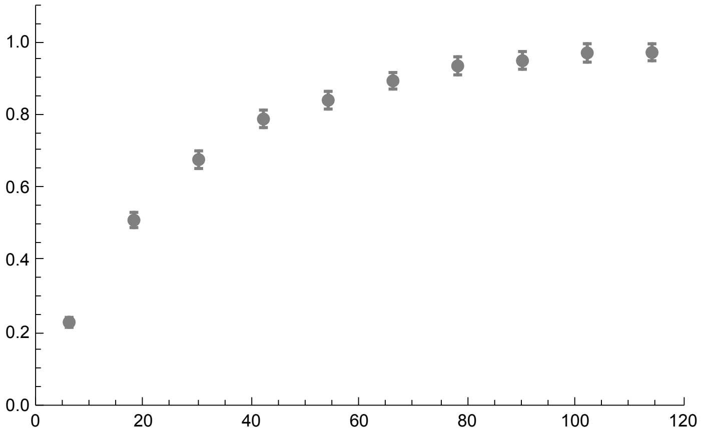

Analytical solutions against both simulation data

```
h05MSMS = Show[Ph05, Ph05S, Ph05MSMSS, ImageSize → 250]
```

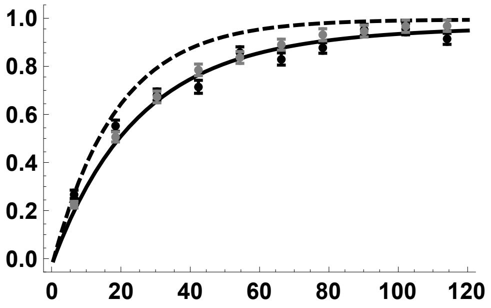

h = 0.1

SLiM simulation data

```
PiRelh01 = Import["SLIM_F0/StatsProc_R_120_h_0.1_self_0_f_1e-04_10b_SLIM.dat",
    "Table"][[10]];
PiRelh01CIB = Import[
    "SLIM_F0/StatsProc_R_120_h_0.1_self_0_f_1e-04_10b_SLIM.dat",
    "Table"][[11]];
PiRelh01CIT = Import[
    "SLIM_F0/StatsProc_R_120_h_0.1_self_0_f_1e-04_10b_SLIM.dat",
    "Table"][[12]];
PiRelh01S = Partition[Riffle[Partition[Riffle[Rin, PiRelh01], 2],
    Map[ErrorBar, Partition[Riffle[-PiRelh01CIB, PiRelh01CIT], 2]]], 2];
```

MSMS simulation data

```
PiRelh01MSMS =
  Import["MSMS_SV_5K/StatsProc_R_240_h_0.1_self_0_f_1e-04_10b_MSMS.dat",
    "Table"][[10]];
PiRelh01MSMSCIB = Import[
    "MSMS_SV_5K/StatsProc_R_240_h_0.1_self_0_f_1e-04_10b_MSMS.dat",
    "Table"][[11]];
PiRelh01MSMSCIT = Import[
    "MSMS_SV_5K/StatsProc_R_240_h_0.1_self_0_f_1e-04_10b_MSMS.dat",
    "Table"][[12]];
PiRelh01MSMSS = Partition[Riffle[Partition[Riffle[Rin, PiRelh01MSMS], 2],
    Map[ErrorBar, Partition[Riffle[-PiRelh01MSMSCIB, PiRelh01MSMSCIT], 2]]], 2];
```

Analytical results

```
Ph01 = Plot[{PWR2B[5000, 0.05, 0.1, 0, R, Boostp0[5000, 0.05, 0.1, 0]],
   ExPiSV[5000, 0.05, 0.1, 0, R/10000, Boostp0[5000, 0.05, 0.1, 0]]},
  {R, 0, 120}, PlotStyle → {{Red, Thick, Dashed}, {Red, Thick}},
  PlotRange → All, Frame → {{True, False}, {True, False}},
  BaseStyle → {FontWeight → "Bold", FontColor → Black, FontSize → 12}]
```

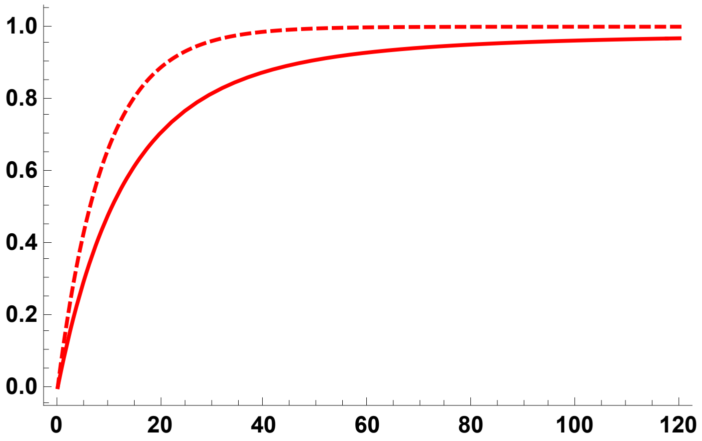


```
Ph01A = Plot[{ExPiSV[5000, 0.05, 0.1, 0, R/10000, Boostp0[5000, 0.05, 0.1, 0]]},
   {R, 0, 120}, PlotStyle → {{Red, Thick}},
   PlotRange → All, Frame → {{True, False}, {True, False}},
   BaseStyle → {FontWeight → "Bold", FontColor → Black, FontSize → 12}];
```

SLiM simulation data

```
Ph01S = ErrorListPlot[PiRelh01S,
  PlotStyle → {Red, PointSize[0.02]}, PlotRange → {{0, 120}, {0, 1.1}}]
```

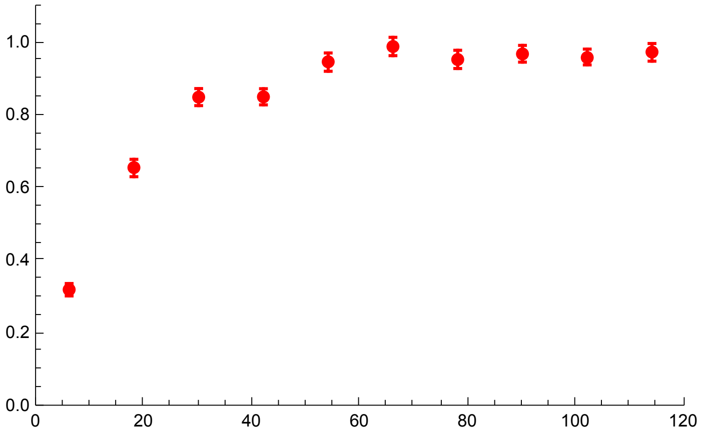

MSMS simulation data

```
Ph01MSMSS = ErrorListPlot[PiRelh01MSMSS,
  PlotStyle → {Pink, PointSize[0.02]}, PlotRange → {{0, 120}, {0, 1.1}}]
```

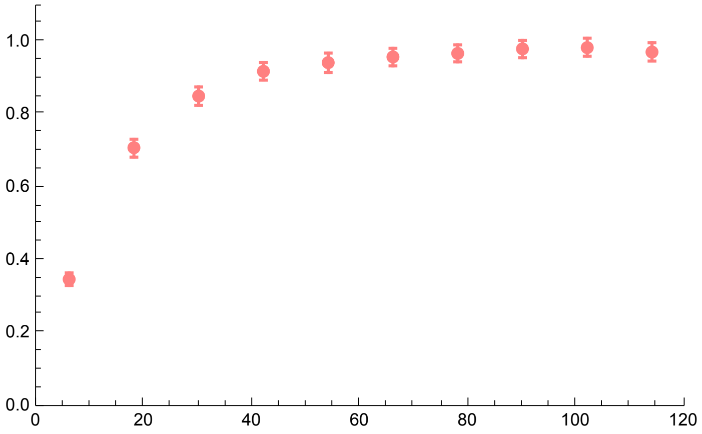

Analytical solutions against both types of simulation data

```
h01MSMS = Show[Ph01, Ph01S, Ph01MSMSS, ImageSize → 250]
```

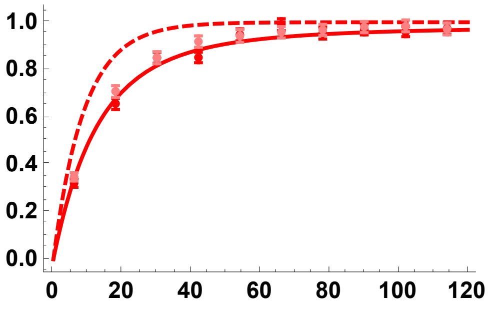

h = 0.9

SLiM simulation data

```
PiRelh09 = Import["SLIM_F0/StatsProc_R_120_h_0.9_self_0_f_1e-04_10b_SLIM.dat",
    "Table"][[10]];
PiRelh09CIB = Import[
    "SLIM_F0/StatsProc_R_120_h_0.9_self_0_f_1e-04_10b_SLIM.dat",
    "Table"][[11]];
PiRelh09CIT = Import[
    "SLIM_F0/StatsProc_R_120_h_0.9_self_0_f_1e-04_10b_SLIM.dat",
    "Table"][[12]];
PiRelh09S = Partition[Riffle[Partition[Riffle[Rin, PiRelh09], 2],
    Map[ErrorBar, Partition[Riffle[-PiRelh09CIB, PiRelh09CIT], 2]]], 2];
```

MSMS simulation data

```
PiRelh09MSMS =
  Import["MSMS_SV_5K/StatsProc_R_240_h_0.9_self_0_f_1e-04_10b_MSMS.dat",
    "Table"][[10]];
PiRelh09MSMSCIB = Import[
    "MSMS_SV_5K/StatsProc_R_240_h_0.9_self_0_f_1e-04_10b_MSMS.dat",
    "Table"][[11]];
PiRelh09MSMSCIT = Import[
    "MSMS_SV_5K/StatsProc_R_240_h_0.9_self_0_f_1e-04_10b_MSMS.dat",
    "Table"][[12]];
PiRelh09MSMSS = Partition[Riffle[Partition[Riffle[Rin, PiRelh09MSMS], 2],
    Map[ErrorBar, Partition[Riffle[-PiRelh09MSMSCIB, PiRelh09MSMSCIT], 2]]], 2];
```

Analytical results

```
Ph09 = Plot[{PWR2B[5000, 0.05, 0.9, 0, R, Boostp0[5000, 0.05, 0.9, 0]],
   ExPiSV[5000, 0.05, 0.9, 0, R/10 000, Boostp0[5000, 0.05, 0.9, 0]]},
  {R, 0, 120}, PlotStyle → { {Blue, Thick, Dashed}, {Blue, Thick}},
  PlotRange → All, Frame → {{True, False}, {True, False}},
  BaseStyle → {FontWeight → "Bold", FontColor → Black, FontSize → 12}]
```

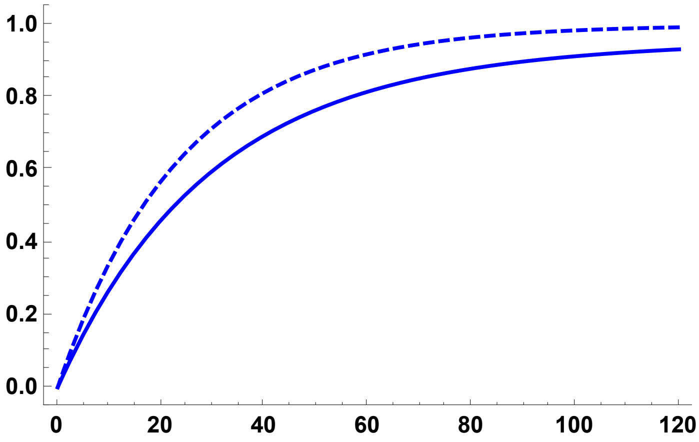

```
Ph09A = Plot[{ExPiSV[5000, 0.05, 0.9, 0, R/10000], Boostp0[5000, 0.05, 0.9, 0]},
   {R, 0, 120}, PlotStyle → {{Blue, Thick}}, PlotRange → All,
   Frame → {{True, False}, {True, False}}, FrameLabel →
    {{"π/π0", None}, {"Recombination Rate, 2Nr", None}}, RotateLabel → False];
```

Plot of SLiM simulation data

```
Ph09S = ErrorListPlot[PiRelh09S,
  PlotStyle → {Blue, PointSize[0.02]}, PlotRange → {{0, 120}, {0, 1.1}}]
```

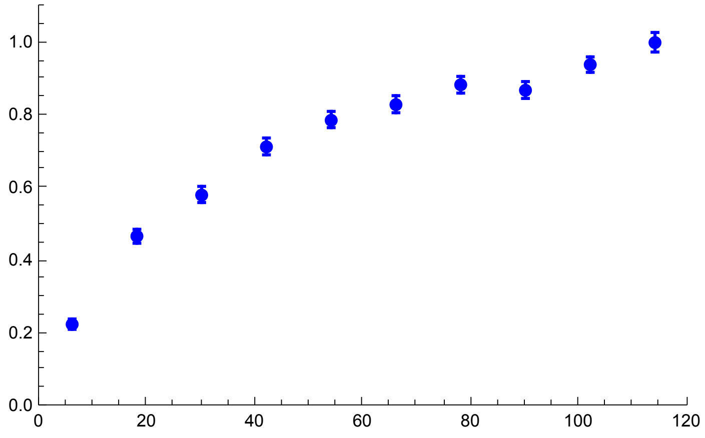


Plot of MSMS simulation data

```
Ph09MSMSS = ErrorListPlot[PiRelh09MSMSS,
  PlotStyle → {Cyan, PointSize[0.02]}, PlotRange → {{0, 120}, {0, 1.1}}]
```

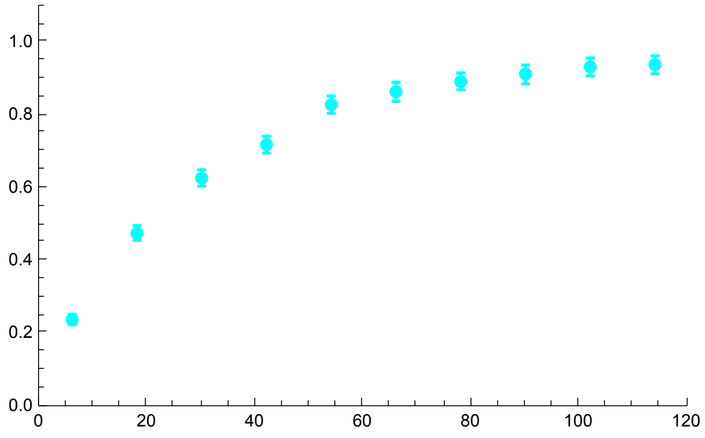


Analytical solutions against both types of simulation data

```
h09MSMS = Show[Ph09, Ph09S, Ph09MSMSS, ImageSize → 250]
```

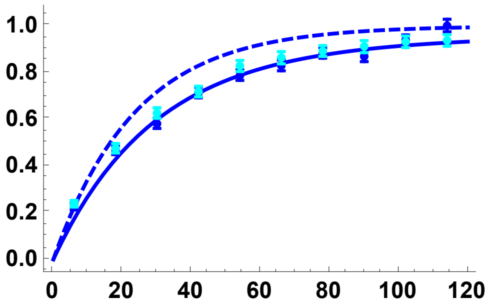

All h values, using the coalescence-in-sweep-phase results and SLiM simulations:

```
Sims0DN = Show[Ph01A, Ph01S, Ph05A, Ph05S, Ph09A, Ph09S, ImageSize → 250]
```

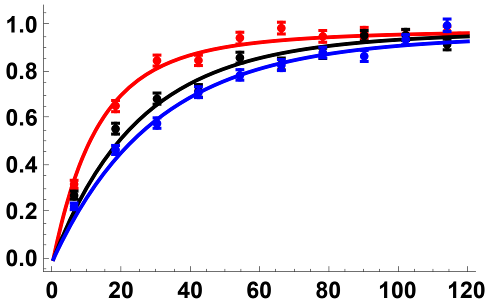


All h values, comparing star-like approximation with coalescence-in-sweep-phase results:

```
Sims0DN2 = Show[Ph01, Ph05, Ph09, ImageSize → 250]
```

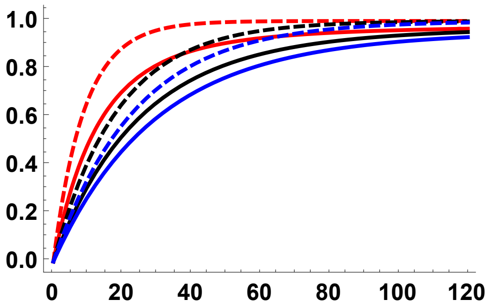


#### Simulation comparisons, $p_0$ = 0.02

Loading Data

```
SetDirectory[NotebookDirectory[]];
Rin = Table[6 + 12 * i, {i, 0, 9}];
```

h = 0.5

SLiM simulation results

```
PiRelh05p002 =
  Import["SLIM_F0/StatsProc_R_120_h_0.5_self_0_f_0.02_10b_SLIM.dat", "Table"][[
   10]];
PiRelh05p002CIB = Import[
    "SLIM_F0/StatsProc_R_120_h_0.5_self_0_f_0.02_10b_SLIM.dat",
    "Table"][[11]];
PiRelh05p002CIT = Import[
    "SLIM_F0/StatsProc_R_120_h_0.5_self_0_f_0.02_10b_SLIM.dat",
    "Table"][[12]];
PiRelh05p002S = Partition[Riffle[Partition[Riffle[Rin, PiRelh05p002], 2],
    Map[ErrorBar, Partition[Riffle[-PiRelh05p002CIB, PiRelh05p002CIT], 2]]], 2];
```

MSMS simulation results

```
PiRelh05p002MSMS =
  Import["MSMS_SV_5K/StatsProc_R_240_h_0.5_self_0_f_0.02_10b_MSMS.dat",
    "Table"][[10]];
PiRelh05p002MSMSCIB = Import[
    "MSMS_SV_5K/StatsProc_R_240_h_0.5_self_0_f_0.02_10b_MSMS.dat",
    "Table"][[11]];
PiRelh05p002MSMSCIT = Import[
    "MSMS_SV_5K/StatsProc_R_240_h_0.5_self_0_f_0.02_10b_MSMS.dat",
    "Table"][[12]];
PiRelh05p002MSMSS = Partition[Riffle[
    Partition[Riffle[Rin, PiRelh05p002MSMS], 2], Map[ErrorBar,
     Partition[Riffle[-PiRelh05p002MSMSCIB, PiRelh05p002MSMSCIT], 2]]], 2];
```

Analytical results

```
Ph05p002 = Plot[
  {PWR2B[5000, 0.05, 0.5, 0, R, 0.02], ExPiSV[5000, 0.05, 0.5, 0, R/10000, 0.02]},
  {R, 0, 120}, PlotStyle → {{Black, Thick, Dashed}, {Black, Thick}},
  PlotRange → All, Frame → {{True, False}, {True, False}},
  BaseStyle → {FontWeight → "Bold", FontColor → Black, FontSize → 12}]
```

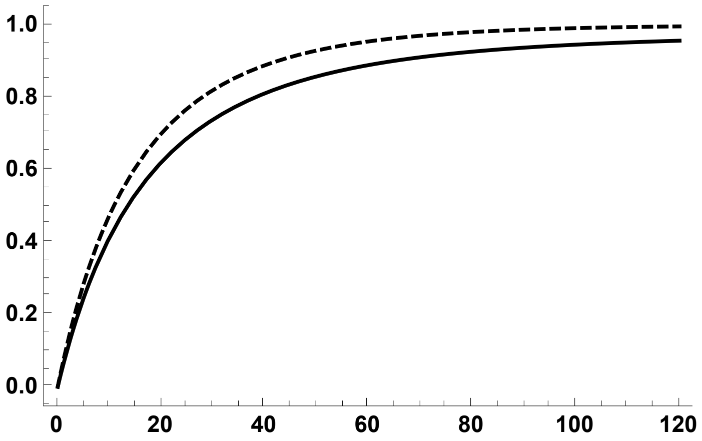


```
Ph05p002A = Plot[{ExPiSV[5000, 0.05, 0.5, 0, R/10000, 0.02]},
   {R, 0, 120}, PlotStyle → {{Black, Thick}},
   PlotRange → All, Frame → {{True, False}, {True, False}},
   BaseStyle → {FontWeight → "Bold", FontColor → Black, FontSize → 12}];
```

SLiM simulation res plot

```
Ph05p002S = ErrorListPlot[PiRelh05p002S,
  PlotStyle → {Black, PointSize[0.02]}, PlotRange → {{0, 120}, {0, 1.1}}]
```

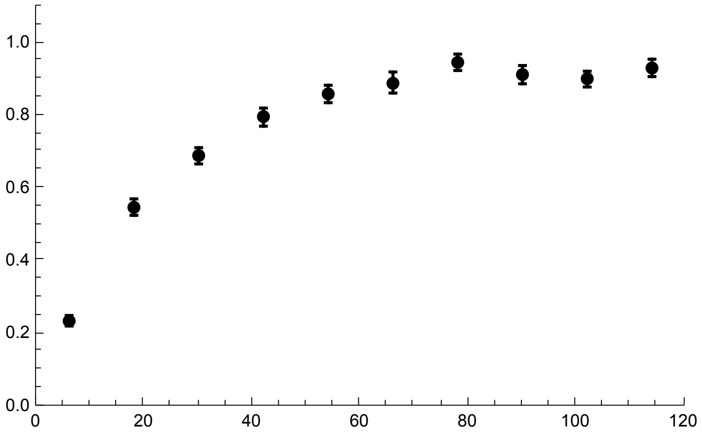

MSMS simulation res plot

```
Ph05p002MSMSS = ErrorListPlot[PiRelh05p002MSMSS,
  PlotStyle → {Gray, PointSize[0.02]}, PlotRange → {{0, 120}, {0, 1.1}}]
```

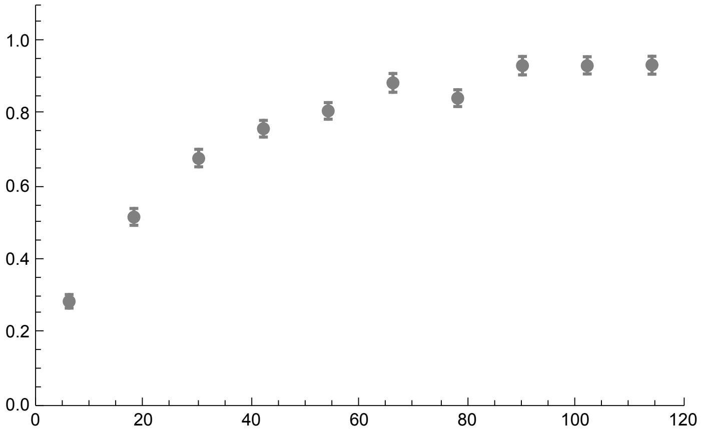

Analytical solutions against both simulation data

```
h05MSMS2pc = Show[Ph05p002, Ph05p002S, Ph05p002MSMSS, ImageSize → 250]
```

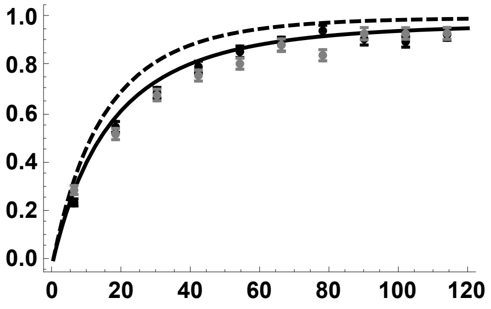

h = 0.1

SLiM simulation results

```
PiRelh01p002 =
  Import["SLIM_F0/StatsProc_R_120_h_0.1_self_0_f_0.02_10b_SLIM.dat", "Table"][[
   10]];
PiRelh01p002CIB = Import[
    "SLIM_F0/StatsProc_R_120_h_0.1_self_0_f_0.02_10b_SLIM.dat",
    "Table"][[11]];
PiRelh01p002CIT = Import[
    "SLIM_F0/StatsProc_R_120_h_0.1_self_0_f_0.02_10b_SLIM.dat",
    "Table"][[12]];
PiRelh01p002S = Partition[Riffle[Partition[Riffle[Rin, PiRelh01p002], 2],
    Map[ErrorBar, Partition[Riffle[-PiRelh01p002CIB, PiRelh01p002CIT], 2]]], 2];
```

MSMS simulation results

```
PiRelh01p002MSMS =
  Import["MSMS_SV_5K/StatsProc_R_240_h_0.1_self_0_f_0.02_10b_MSMS.dat",
    "Table"][[10]];
PiRelh01p002MSMSCIB = Import[
    "MSMS_SV_5K/StatsProc_R_240_h_0.1_self_0_f_0.02_10b_MSMS.dat",
    "Table"][[11]];
PiRelh01p002MSMSCIT = Import[
    "MSMS_SV_5K/StatsProc_R_240_h_0.1_self_0_f_0.02_10b_MSMS.dat",
    "Table"][[12]];
PiRelh01p002MSMSS = Partition[Riffle[
    Partition[Riffle[Rin, PiRelh01p002MSMS], 2], Map[ErrorBar,
     Partition[Riffle[-PiRelh01p002MSMSCIB, PiRelh01p002MSMSCIT], 2]]], 2];
```

Analytical results

```
Ph01p002 = Plot[
  {PWR2B[5000, 0.05, 0.1, 0, R, 0.02], ExPiSV[5000, 0.05, 0.1, 0, R/10 000, 0.02]},
  {R, 0, 120}, PlotStyle → {{Red, Thick, Dashed}, {Red, Thick}},
  PlotRange → All, Frame → {{True, False}, {True, False}},
  BaseStyle → {FontWeight → "Bold", FontColor → Black, FontSize → 12}]
```

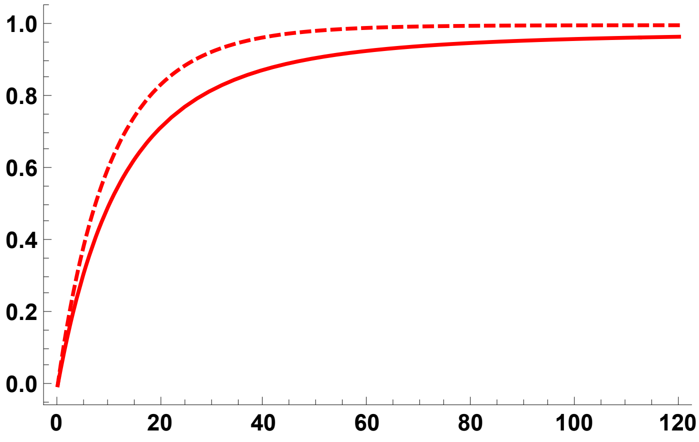

```
Ph01p002A = Plot[{ExPiSV[5000, 0.05, 0.1, 0, R/10000, 0.02]},
    {R, 0, 120}, PlotStyle → {{Red, Thick}},
    PlotRange → All, Frame → {{True, False}, {True, False}},
    BaseStyle → {FontWeight → "Bold", FontColor → Black, FontSize → 12}];
```

SLiM simulation res plot

```
Ph01p002S = ErrorListPlot[PiRelh01p002S,
  PlotStyle → {Red, PointSize[0.02]}, PlotRange → {{0, 120}, {0, 1.1}}]
```

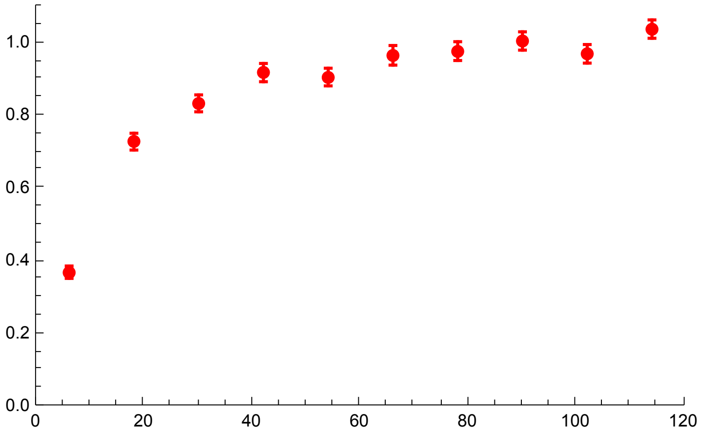


MSMS simulation res plot

```
Ph01p002MSMSS = ErrorListPlot[PiRelh01p002MSMSS,
  PlotStyle → {Pink, PointSize[0.02]}, PlotRange → {{0, 120}, {0, 1.1}}]
```

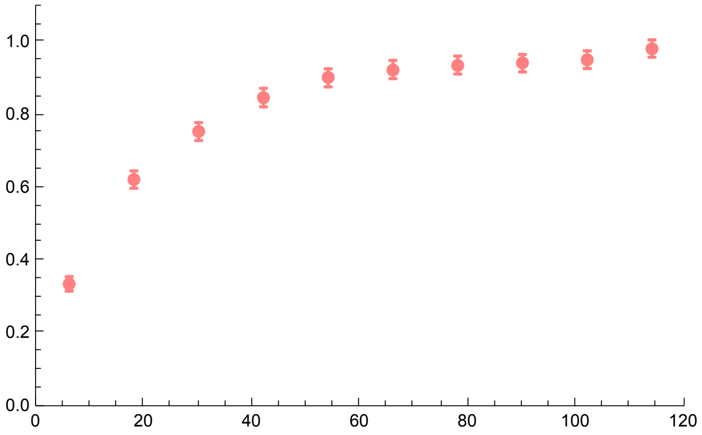


Analytical solutions against both simulation data

```
h01MSMS2pc = Show[Ph01p002, Ph01p002S, Ph01p002MSMSS, ImageSize → 250]
```

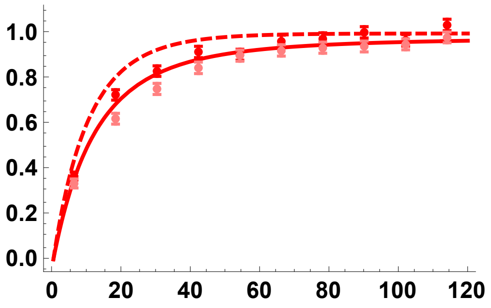

h = 0.9

SLiM simulation results

```
PiRelh09p002 =
  Import["SLIM_F0/StatsProc_R_120_h_0.9_self_0_f_0.02_10b_SLIM.dat", "Table"][[
   10]];
PiRelh09p002CIB = Import[
    "SLIM_F0/StatsProc_R_120_h_0.9_self_0_f_0.02_10b_SLIM.dat",
    "Table"][[11]];
PiRelh09p002CIT = Import[
    "SLIM_F0/StatsProc_R_120_h_0.9_self_0_f_0.02_10b_SLIM.dat",
    "Table"][[12]];
PiRelh09p002S = Partition[Riffle[Partition[Riffle[Rin, PiRelh09p002], 2],
    Map[ErrorBar, Partition[Riffle[-PiRelh09p002CIB, PiRelh09p002CIT], 2]]], 2];
```

MSMS simulation results

```
PiRelh09p002MSMS =
  Import["MSMS_SV_5K/StatsProc_R_240_h_0.9_self_0_f_0.02_10b_MSMS.dat",
    "Table"][[10]];
PiRelh09p002MSMSCIB = Import[
    "MSMS_SV_5K/StatsProc_R_240_h_0.9_self_0_f_0.02_10b_MSMS.dat",
    "Table"][[11]];
PiRelh09p002MSMSCIT = Import[
    "MSMS_SV_5K/StatsProc_R_240_h_0.9_self_0_f_0.02_10b_MSMS.dat",
    "Table"][[12]];
PiRelh09p002MSMSS = Partition[Riffle[
    Partition[Riffle[Rin, PiRelh09p002MSMS], 2], Map[ErrorBar,
     Partition[Riffle[-PiRelh09p002MSMSCIB, PiRelh09p002MSMSCIT], 2]]], 2];
```

Analytical results

```
Ph09p002 = Plot[
  {PWR2B[5000, 0.05, 0.9, 0, R, 0.02], ExPiSV[5000, 0.05, 0.9, 0, R/10 000, 0.02]},
  {R, 0, 120}, PlotStyle → { {Blue, Thick, Dashed}, {Blue, Thick}},
  PlotRange → All, Frame → {{True, False}, {True, False}},
  BaseStyle → {FontWeight → "Bold", FontColor → Black, FontSize → 12}]
```

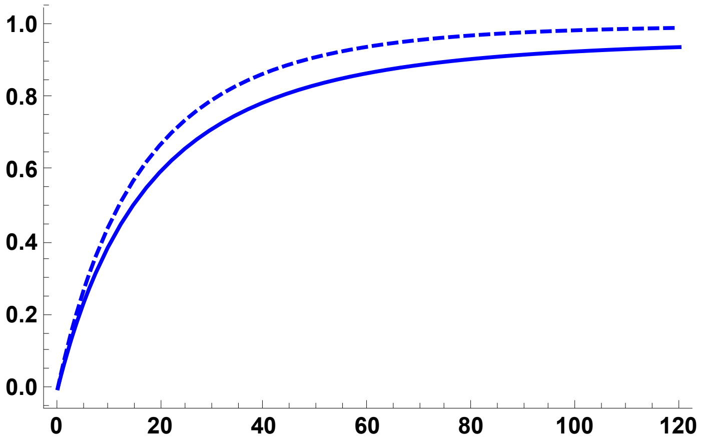


```
Ph09p002A = Plot[{ExPiSV[5000, 0.05, 0.9, 0, R/10 000, 0.02]},
   {R, 0, 120}, PlotStyle → {{Blue, Thick}},
   PlotRange → All, Frame → {{True, False}, {True, False}},
   BaseStyle → {FontWeight → "Bold", FontColor → Black, FontSize → 12}];
```

SLiM simulation plot

```
Ph09p002S = ErrorListPlot[PiRelh09p002S,
  PlotStyle → {Blue, PointSize[0.02]}, PlotRange → {{0, 120}, {0, 1.1}}]
```

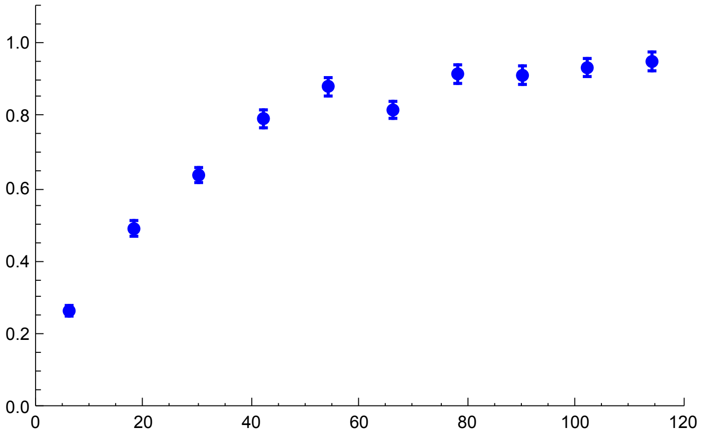


MSMS simulation plot

```
Ph09p002MSMSS = ErrorListPlot[PiRelh09p002MSMSS,
  PlotStyle → {Cyan, PointSize[0.02]}, PlotRange → {{0, 120}, {0, 1.1}}]
```

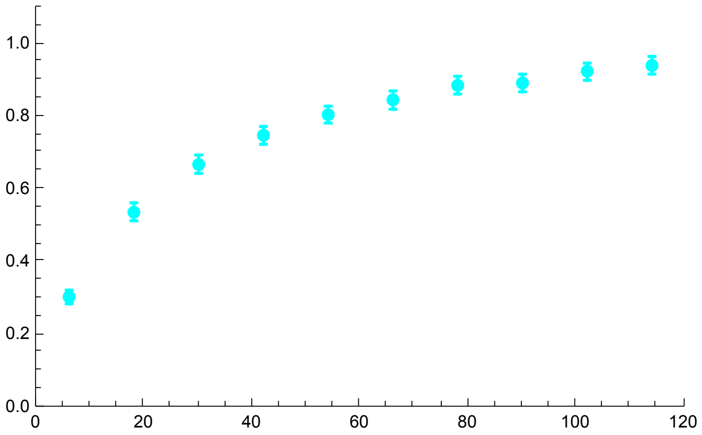

Analytical solutions against both simulation data

```
h09MSMS2pc = Show[Ph09p002, Ph09p002S, Ph09p002MSMSS, ImageSize → 250]
```

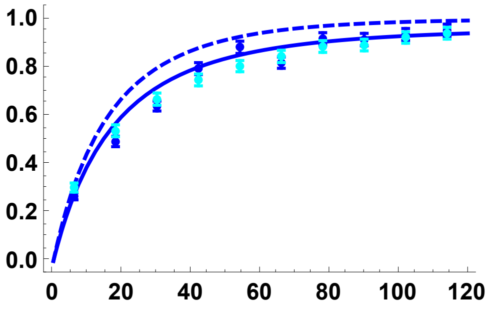

All h values, using the coalescence-in-sweep-phase results and SLiM simulations:

```
Sims02pc = Show[Ph01p002A, Ph01p002S,
  Ph05p002A, Ph05p002S, Ph09p002A, Ph09p002S, ImageSize → 250]
```

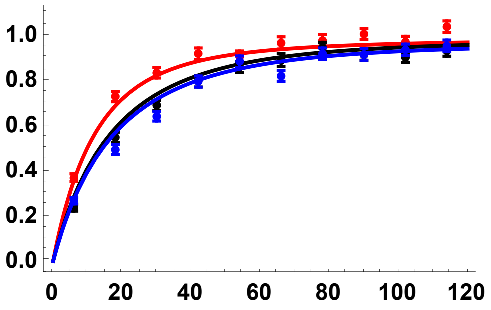

All h values, comparing star-like approximation with coalescence-in-sweep-phase results:

```
Sims02pc2 = Show[Ph01p002, Ph05p002, Ph09p002, ImageSize → 250]
```

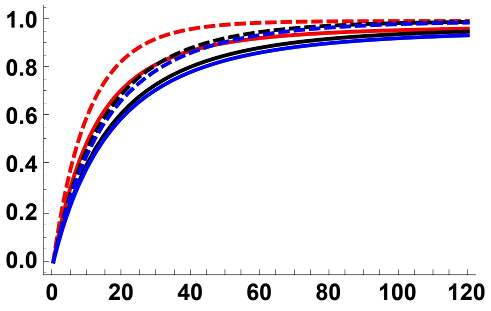


#### Simulation comparisons, $p_0$ = 0.05

h = 0.5

SLiM sims

```
PiRelh05p005 =
  Import["SLIM_F0/StatsProc_R_120_h_0.5_self_0_f_0.05_10b_SLIM.dat", "Table"][[
   10]];
PiRelh05p005CIB = Import[
    "SLIM_F0/StatsProc_R_120_h_0.5_self_0_f_0.05_10b_SLIM.dat",
    "Table"][[11]];
PiRelh05p005CIT = Import[
    "SLIM_F0/StatsProc_R_120_h_0.5_self_0_f_0.05_10b_SLIM.dat",
    "Table"][[12]];
PiRelh05p005S = Partition[Riffle[Partition[Riffle[Rin, PiRelh05p005], 2],
    Map[ErrorBar, Partition[Riffle[-PiRelh05p005CIB, PiRelh05p005CIT], 2]]], 2];
```

MSMS Sims

```
PiRelh05p005MSMS =
  Import["MSMS_SV_5K/StatsProc_R_240_h_0.5_self_0_f_0.05_10b_MSMS.dat",
    "Table"][[10]];
PiRelh05p005MSMSCIB = Import[
    "MSMS_SV_5K/StatsProc_R_240_h_0.5_self_0_f_0.05_10b_MSMS.dat",
    "Table"][[11]];
PiRelh05p005MSMSCIT = Import[
    "MSMS_SV_5K/StatsProc_R_240_h_0.5_self_0_f_0.05_10b_MSMS.dat",
    "Table"][[12]];
PiRelh05p005MSMSS = Partition[Riffle[
    Partition[Riffle[Rin, PiRelh05p005MSMS], 2], Map[ErrorBar,
     Partition[Riffle[-PiRelh05p005MSMSCIB, PiRelh05p005MSMSCIT], 2]]], 2];
```

Analytical solutions

```
Ph05p005 = Plot[
  {PWR2B[5000, 0.05, 0.5, 0, R, 0.05], ExPiSV[5000, 0.05, 0.5, 0, R/10000, 0.05]},
  {R, 0, 120}, PlotStyle → {{Black, Thick, Dashed}, {Black, Thick}},
  PlotRange → All, Frame → {{True, False}, {True, False}},
  BaseStyle → {FontWeight → "Bold", FontColor → Black, FontSize → 12}]
```

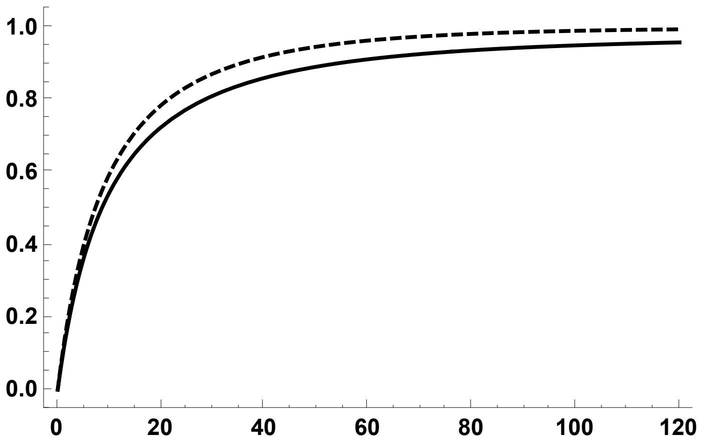


```
Ph05p005A = Plot[{ExPiSV[5000, 0.05, 0.5, 0, R/10000, 0.05]},
   {R, 0, 120}, PlotStyle → {{Black, Thick}},
   PlotRange → All, Frame → {{True, False}, {True, False}},
   BaseStyle → {FontWeight → "Bold", FontColor → Black, FontSize → 12}];
```

SLiM sims

```
Ph05p005S = ErrorListPlot[PiRelh05p005S,
  PlotStyle → {Black, PointSize[0.02]}, PlotRange → {{0, 120}, {0, 1.1}}]
```

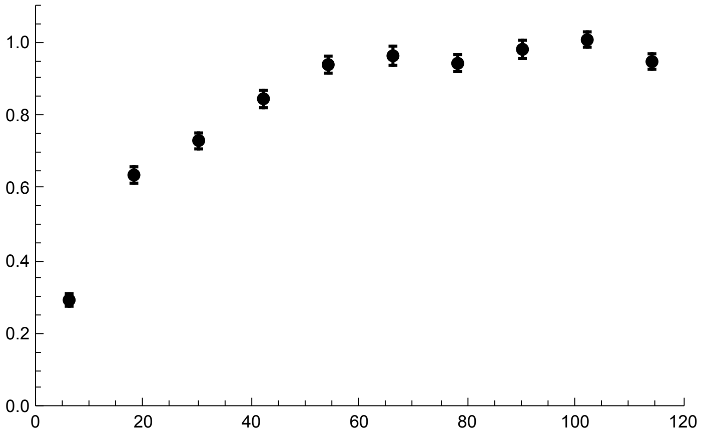


MSMS Sims

```
Ph05p005MSMSS = ErrorListPlot[PiRelh05p005MSMSS,
  PlotStyle → {Gray, PointSize[0.02]}, PlotRange → {{0, 120}, {0, 1.1}}]
```

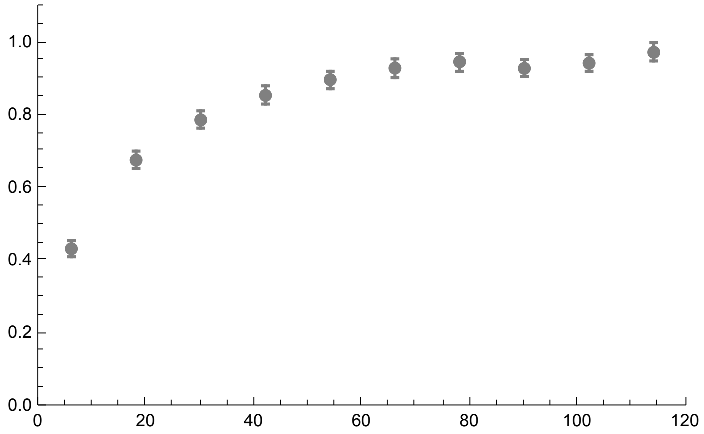


Analytical solutions against both types of simulation data

```
h05MSMS5pc = Show[Ph05p005, Ph05p005S, Ph05p005MSMSS, ImageSize → 250]
```

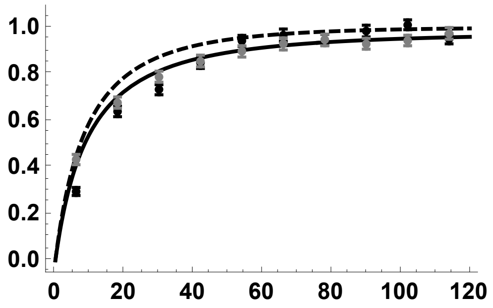


h = 0.1

SLiM simulation data

```
PiRelh01p005 =
  Import["SLIM_F0/StatsProc_R_120_h_0.1_self_0_f_0.05_10b_SLIM.dat", "Table"][[
   10]];
PiRelh01p005CIB = Import[
    "SLIM_F0/StatsProc_R_120_h_0.1_self_0_f_0.05_10b_SLIM.dat",
    "Table"][[11]];
PiRelh01p005CIT = Import[
    "SLIM_F0/StatsProc_R_120_h_0.1_self_0_f_0.05_10b_SLIM.dat",
    "Table"][[12]];
PiRelh01p005S = Partition[Riffle[Partition[Riffle[Rin, PiRelh01p005], 2],
    Map[ErrorBar, Partition[Riffle[-PiRelh01p005CIB, PiRelh01p005CIT], 2]]], 2];
```

MSMS simulation data

```
PiRelh01p005MSMS =
  Import["MSMS_SV_5K/StatsProc_R_240_h_0.1_self_0_f_0.05_10b_MSMS.dat",
    "Table"][[10]];
PiRelh01p005MSMSCIB = Import[
    "MSMS_SV_5K/StatsProc_R_240_h_0.1_self_0_f_0.05_10b_MSMS.dat",
    "Table"][[11]];
PiRelh01p005MSMSCIT = Import[
    "MSMS_SV_5K/StatsProc_R_240_h_0.1_self_0_f_0.05_10b_MSMS.dat",
    "Table"][[12]];
PiRelh01p005MSMSS = Partition[Riffle[
    Partition[Riffle[Rin, PiRelh01p005MSMS], 2], Map[ErrorBar,
     Partition[Riffle[-PiRelh01p005MSMSCIB, PiRelh01p005MSMSCIT], 2]]], 2];
```

Analytical results

```
Ph01p005 = Plot[
  {PWR2B[5000, 0.05, 0.1, 0, R, 0.05], ExPiSV[5000, 0.05, 0.1, 0, R/10 000, 0.05]},
  {R, 0, 120}, PlotStyle → {{Red, Thick, Dashed}, {Red, Thick}},
  PlotRange → All, Frame → {{True, False}, {True, False}},
  BaseStyle → {FontWeight → "Bold", FontColor → Black, FontSize → 12}]
```

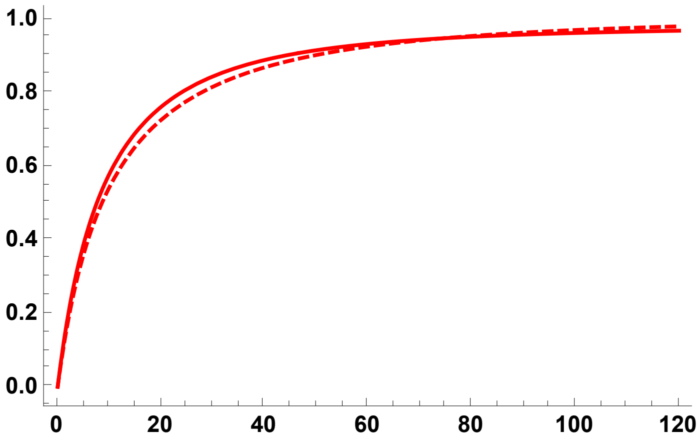


```
Ph01p005A = Plot[{ExPiSV[5000, 0.05, 0.1, 0, R/10 000, 0.05]},
   {R, 0, 120}, PlotStyle → {{Red, Thick}},
   PlotRange → All, Frame → {{True, False}, {True, False}},
   BaseStyle → {FontWeight → "Bold", FontColor → Black, FontSize → 12}];
```

SLiM simulation data plot

```
Ph01p005S = ErrorListPlot[PiRelh01p005S,
  PlotStyle → {Red, PointSize[0.02]}, PlotRange → {{0, 120}, {0, 1.1}}]
```

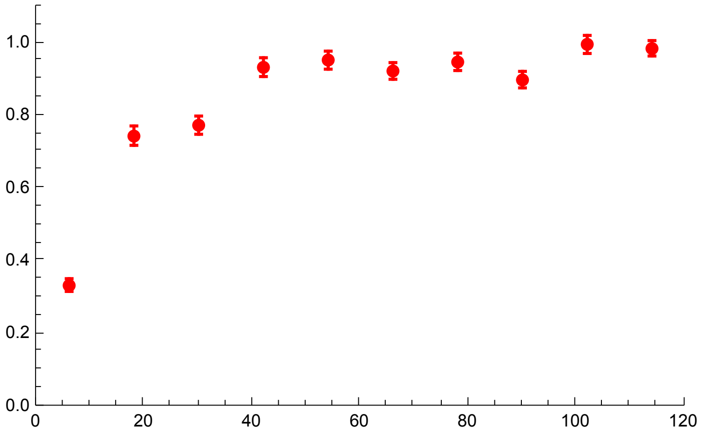


MSMS simulation data plot

```
Ph01p005MSMSS = ErrorListPlot[PiRelh01p005MSMSS,
  PlotStyle → {Pink, PointSize[0.02]}, PlotRange → {{0, 120}, {0, 1.1}}]
```

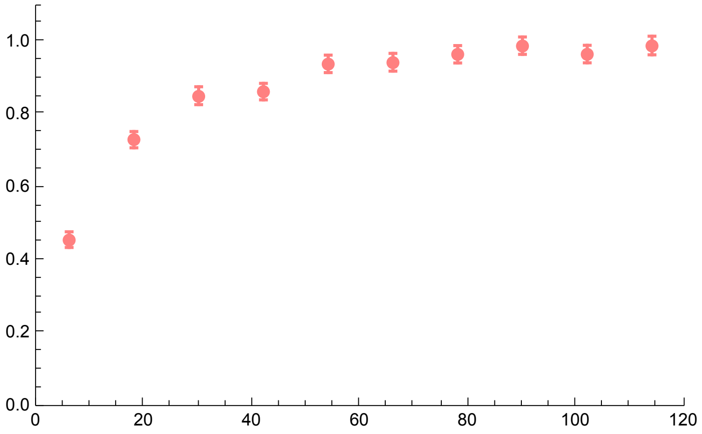


Analytical solutions against both types of simulation data

```
h01MSMS5pc = Show[Ph01p005, Ph01p005S, Ph01p005MSMSS, ImageSize → 250]
```

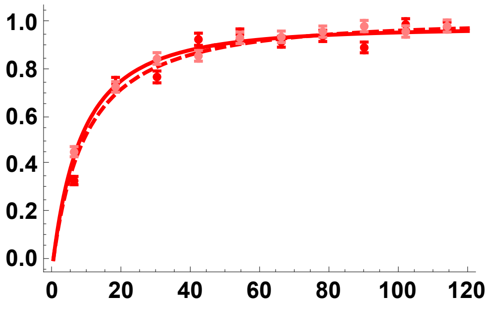


h = 0.9

SLiM simulation data

```
PiRelh09p005 =
  Import["SLIM_F0/StatsProc_R_120_h_0.9_self_0_f_0.05_10b_SLIM.dat", "Table"][[
   10]];
PiRelh09p005CIB = Import[
    "SLIM_F0/StatsProc_R_120_h_0.9_self_0_f_0.05_10b_SLIM.dat",
    "Table"][[11]];
PiRelh09p005CIT = Import[
    "SLIM_F0/StatsProc_R_120_h_0.9_self_0_f_0.05_10b_SLIM.dat",
    "Table"][[12]];
PiRelh09p005S = Partition[Riffle[Partition[Riffle[Rin, PiRelh09p005], 2],
    Map[ErrorBar, Partition[Riffle[-PiRelh09p005CIB, PiRelh09p005CIT], 2]]], 2];
```

MSMS simulation data

```
PiRelh09p005MSMS =
  Import["MSMS_SV_5K/StatsProc_R_240_h_0.9_self_0_f_0.05_10b_MSMS.dat",
    "Table"][[10]];
PiRelh09p005MSMSCIB = Import[
    "MSMS_SV_5K/StatsProc_R_240_h_0.9_self_0_f_0.05_10b_MSMS.dat",
    "Table"][[11]];
PiRelh09p005MSMSCIT = Import[
    "MSMS_SV_5K/StatsProc_R_240_h_0.9_self_0_f_0.05_10b_MSMS.dat",
    "Table"][[12]];
PiRelh09p005MSMSS = Partition[Riffle[
    Partition[Riffle[Rin, PiRelh09p005MSMS], 2], Map[ErrorBar,
     Partition[Riffle[-PiRelh09p005MSMSCIB, PiRelh09p005MSMSCIT], 2]]], 2];
```

Analytical solutions

```
Ph09p005 = Plot[
  {PWR2B[5000, 0.05, 0.9, 0, R, 0.05], ExPiSV[5000, 0.05, 0.9, 0, R/10 000, 0.05]},
  {R, 0, 120}, PlotStyle → { {Blue, Thick, Dashed}, {Blue, Thick}},
  PlotRange → All, Frame → {{True, False}, {True, False}},
  BaseStyle → {FontWeight → "Bold", FontColor → Black, FontSize → 12}]
```

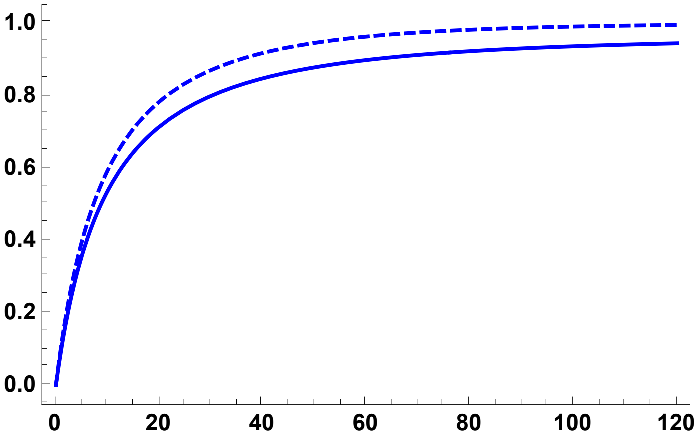

```
Ph09p005A = Plot[{ExPiSV[5000, 0.05, 0.9, 0, R/10 000, 0.05]},
   {R, 0, 120}, PlotStyle → {{Blue, Thick}},
   PlotRange → All, Frame → {{True, False}, {True, False}},
   BaseStyle → {FontWeight → "Bold", FontColor → Black, FontSize → 12}];
```

SLiM simulation data plot

```
Ph09p005S = ErrorListPlot[PiRelh09p005S,
  PlotStyle → {Blue, PointSize[0.02]}, PlotRange → {{0, 120}, {0, 1.1}}]
```

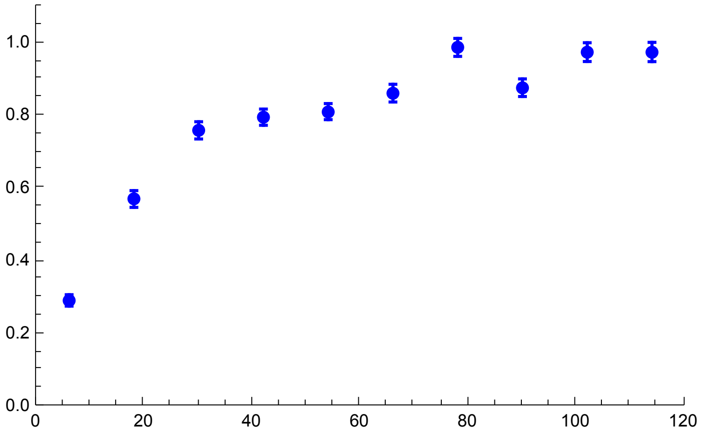

MSMS simulation data plot

```
Ph09p005MSMSS = ErrorListPlot[PiRelh09p005MSMSS,
  PlotStyle → {Cyan, PointSize[0.02]}, PlotRange → {{0, 120}, {0, 1.1}}]
```

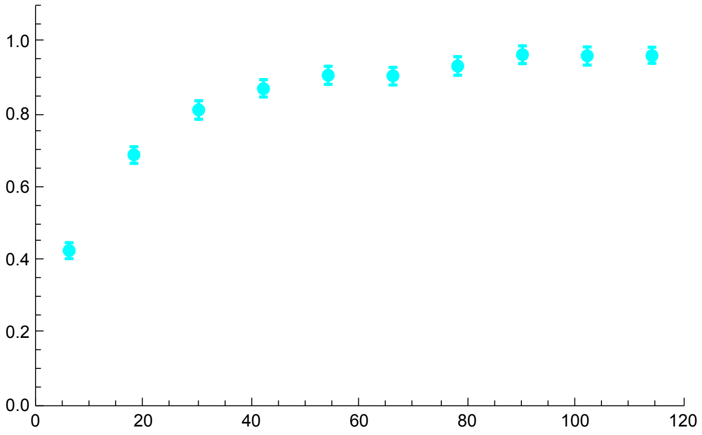

Analytical results against both types of simulation data

```
h09MSMS5pc = Show[Ph09p005, Ph09p005S, Ph09p005MSMSS, ImageSize → 250]
```

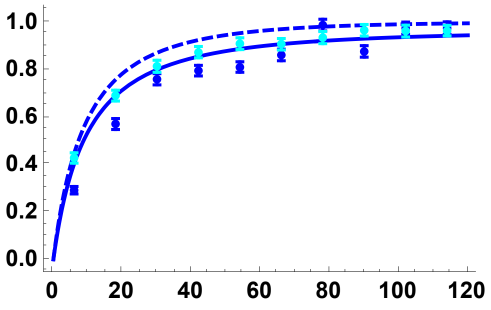

All h values, using the coalescence-in-sweep-phase results and SLiM simulations:

```
Sims05pc = Show[Ph01p005A, Ph01p005S,
  Ph05p005A, Ph05p005S, Ph09p005A, Ph09p005S, ImageSize → 250]
```

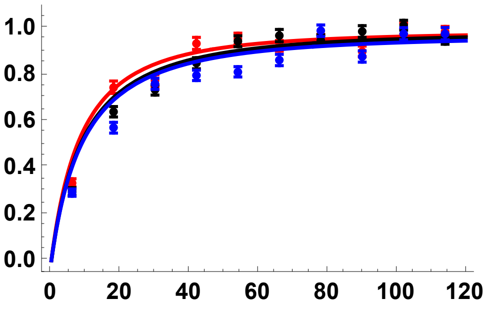

All h values, comparing star-like approximation with coalescence-in-sweep-phase results:

```
Sims05pc2 = Show[Ph01p005, Ph05p005, Ph09p005, ImageSize → 250]
```

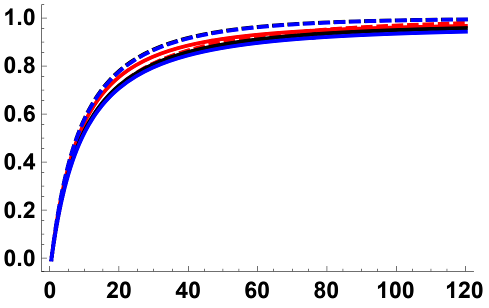

#### Results Grid: comparing SLiM and MSMS Simulations

```
SLiMandMSMSRes = Labeled[
  Grid[{{Text@TraditionalForm@Style["h = 0.1", 24, TextAlignment → Center],
     Text@TraditionalForm@Style["h = 0.5", 24], Text@
      TraditionalForm@Style["h = 0.9", 24],}, {Text@TraditionalForm@Style[
        "Red points: forward-in-time\nPink points: coalescent simulations",
        14, TextAlignment → Center], Text@TraditionalForm@Style[
        "Black points: forward-in-time\nGrey points: coalescent simulations",
        14, TextAlignment → Center], Text@TraditionalForm@Style[
        "Blue points: forward-in-time\nCyan points: coalescent simulations",
        14, TextAlignment → Center],},
    {h01MSMS, h05MSMS, h09MSMS, Text@TraditionalForm@Style["p0 = 1/2N", 24]},
    {h01MSMS2pc, h05MSMS2pc, h09MSMS2pc,
     Text@TraditionalForm@Style["p0 = 0.02", 24]},
    {h01MSMS5pc, h05MSMS5pc, h09MSMS5pc,
     Text@TraditionalForm@Style["p0 = 0.05", 24]}}, Spacings → {2, 1}],
  {Text@TraditionalForm@Style["Expected\nRelative\nDiversity,\n𝔼[πSV/π0]", 24],
   Text@TraditionalForm@Style["Scaled Recombination Rate, 2Nr", 24]}, {Left,
   Bottom}]
```

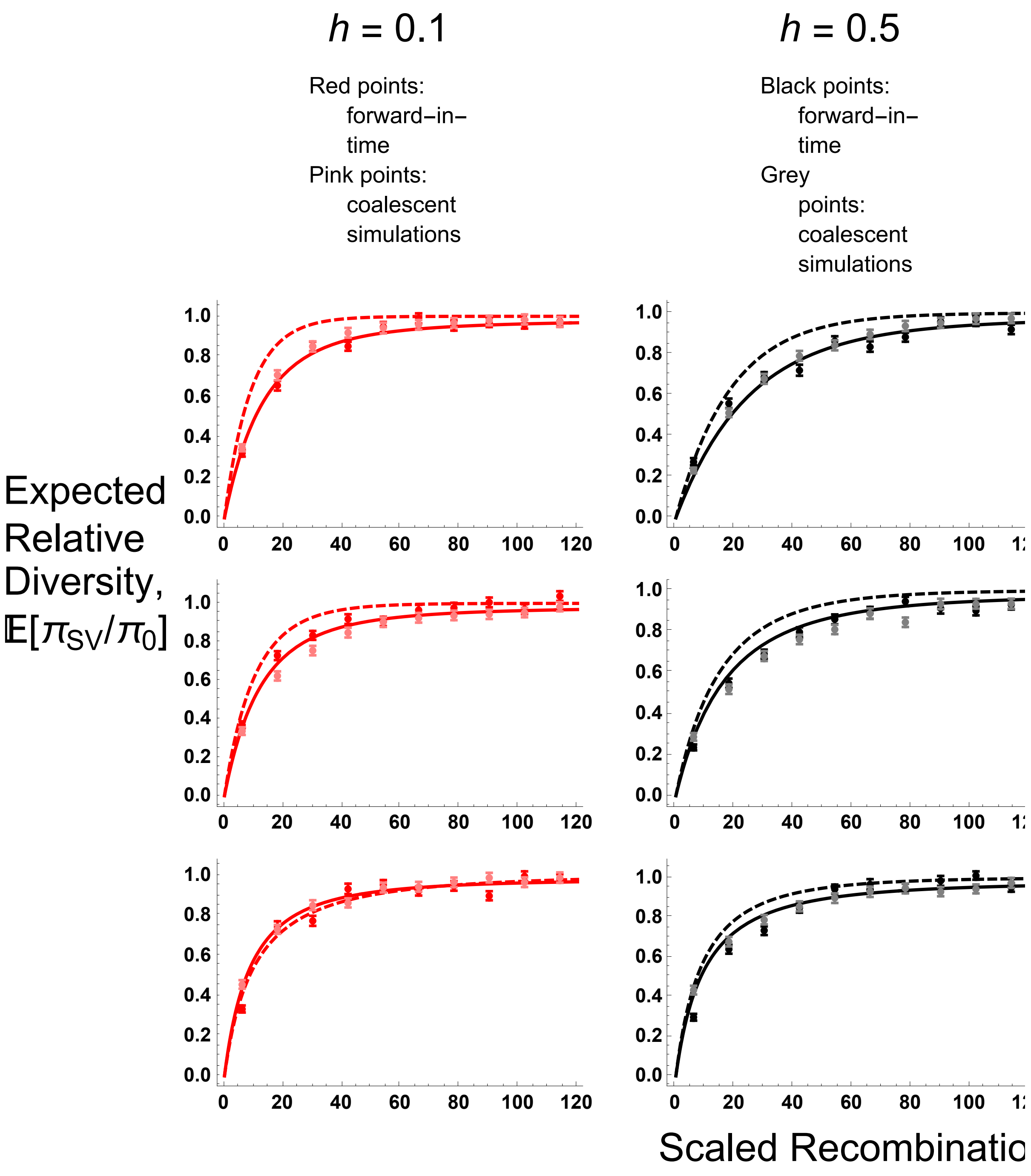


#### $\sigma = 1/2$ ($F = 1/3$)

All simulation data from here on is from SLiM.

Preliminaries

```
SetDirectory[NotebookDirectory[]];
Rin = Table[11 + 22 * i, {i, 0, 9}];
```

##### Simulation comparisons, from initial frequency $p_0 = 1/2N$

h = 0.5

```
PiRelh05s05 =
  Import["SLIM_F033/StatsProc_R_220_h_0.5_self_0.5_f_1e-04_10b_SLIM.dat",
    "Table"][[10]];
PiRelh05s05CIB = Import[
    "SLIM_F033/StatsProc_R_220_h_0.5_self_0.5_f_1e-04_10b_SLIM.dat",
    "Table"][[11]];
PiRelh05s05CIT = Import[
    "SLIM_F033/StatsProc_R_220_h_0.5_self_0.5_f_1e-04_10b_SLIM.dat",
    "Table"][[12]];
PiRelh05s05S = Partition[Riffle[Partition[Riffle[Rin, PiRelh05s05], 2],
    Map[ErrorBar, Partition[Riffle[-PiRelh05s05CIB, PiRelh05s05CIT], 2]]], 2];
```

```
Ph05s05 = Plot[{PWR2B[5000, 0.05, 0.5, 0.5, R, Boostp0[5000, 0.05, 0.5, 0.5]],
   ExPiSV[5000, 0.05, 0.5, 0.5, R/10 000, Boostp0[5000, 0.05, 0.5, 0.5]]},
  {R, 0, 220}, PlotStyle → {{Black, Thick, Dashed}, {Black, Thick}},
  PlotRange → All, Frame → {{True, False}, {True, False}},
  BaseStyle → {FontWeight → "Bold", FontColor → Black, FontSize → 12}]
```

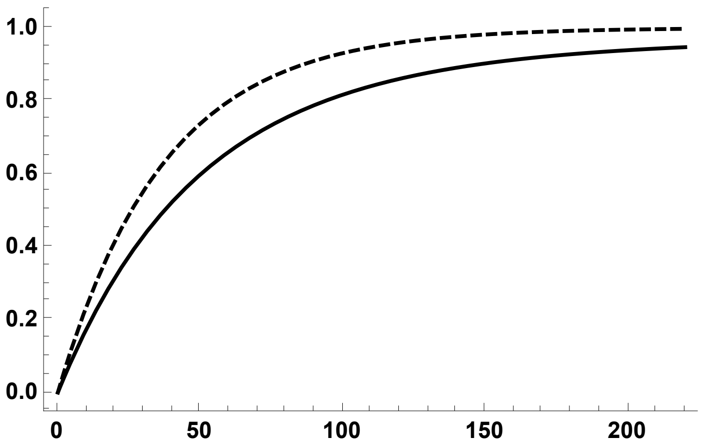


```
Ph05s05A =
  Plot[{ExPiSV[5000, 0.05, 0.5, 0.5, R/10 000, Boostp0[5000, 0.05, 0.5, 0.5]]},
   {R, 0, 220}, PlotStyle → {{Black, Thick}},
   PlotRange → All, Frame → {{True, False}, {True, False}},
   BaseStyle → {FontWeight → "Bold", FontColor → Black, FontSize → 12}];
```

```
Ph05s05S = ErrorListPlot[PiRelh05s05S,
  PlotStyle → {Black, PointSize[0.02]}, PlotRange → {{0, 220}, {0, 1.1}}]
```

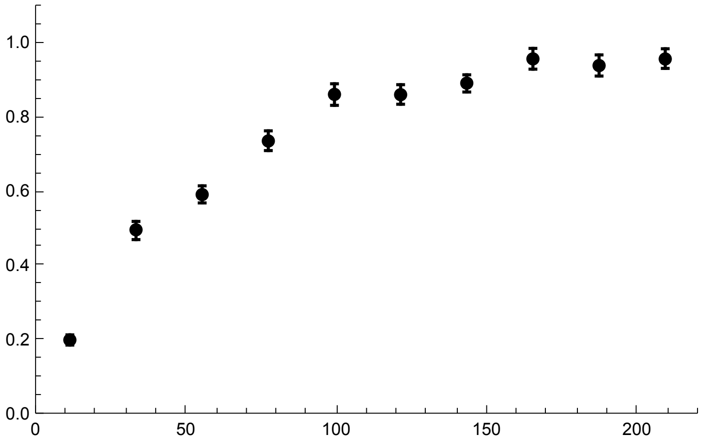

```
Show[Ph05s05, Ph05s05S]
```

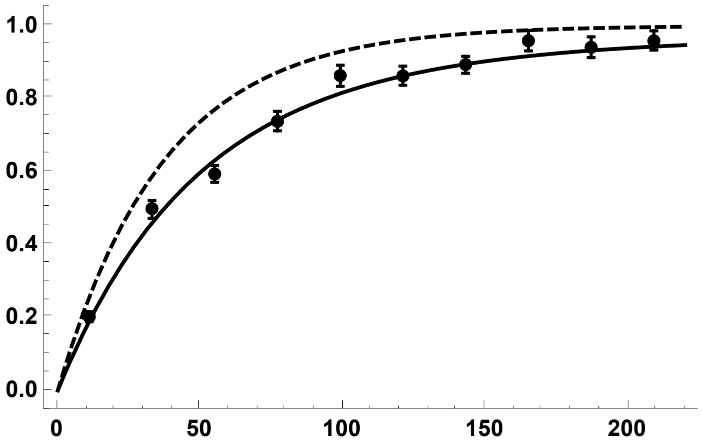

h = 0.1

```
PiRelh01s05 =
  Import["SLIM_F033/StatsProc_R_220_h_0.1_self_0.5_f_1e-04_10b_SLIM.dat",
    "Table"][[10]];
PiRelh01s05CIB = Import[
    "SLIM_F033/StatsProc_R_220_h_0.1_self_0.5_f_1e-04_10b_SLIM.dat",
    "Table"][[11]];
PiRelh01s05CIT = Import[
    "SLIM_F033/StatsProc_R_220_h_0.1_self_0.5_f_1e-04_10b_SLIM.dat",
    "Table"][[12]];
PiRelh01s05S = Partition[Riffle[Partition[Riffle[Rin, PiRelh01s05], 2],
    Map[ErrorBar, Partition[Riffle[-PiRelh01s05CIB, PiRelh01s05CIT], 2]]], 2];
```

```
Ph01s05 = Plot[{PWR2B[5000, 0.05, 0.1, 0.5, R, Boostp0[5000, 0.05, 0.1, 0.5]],
   ExPiSV[5000, 0.05, 0.1, 0.5, R/10 000, Boostp0[5000, 0.05, 0.1, 0.5]]},
  {R, 0, 220}, PlotStyle → {{Red, Thick, Dashed}, {Red, Thick}},
  PlotRange → All, Frame → {{True, False}, {True, False}},
  BaseStyle → {FontWeight → "Bold", FontColor → Black, FontSize → 12}]
```

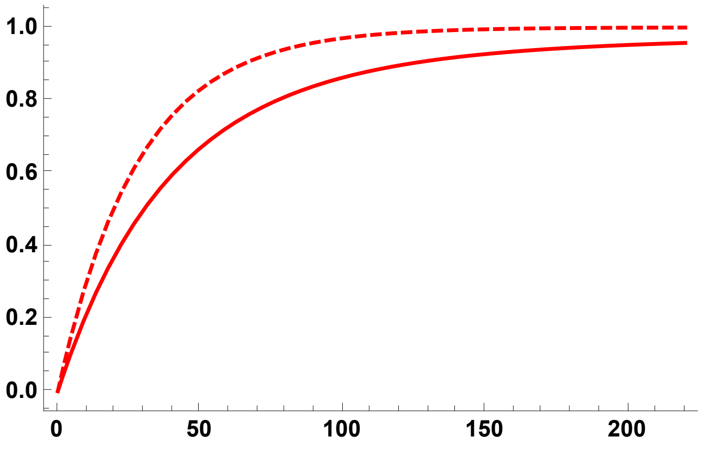


```
Ph01s05A =
  Plot[{ExPiSV[5000, 0.05, 0.1, 0.5, R/10 000, Boostp0[5000, 0.05, 0.1, 0.5]]},
   {R, 0, 220}, PlotStyle → {{Red, Thick}},
   PlotRange → All, Frame → {{True, False}, {True, False}},
   BaseStyle → {FontWeight → "Bold", FontColor → Black, FontSize → 12}];

Ph01s05S = ErrorListPlot[PiRelh01s05S,
  PlotStyle → {Red, PointSize[0.02]}, PlotRange → {{0, 220}, {0, 1.1}}]
```

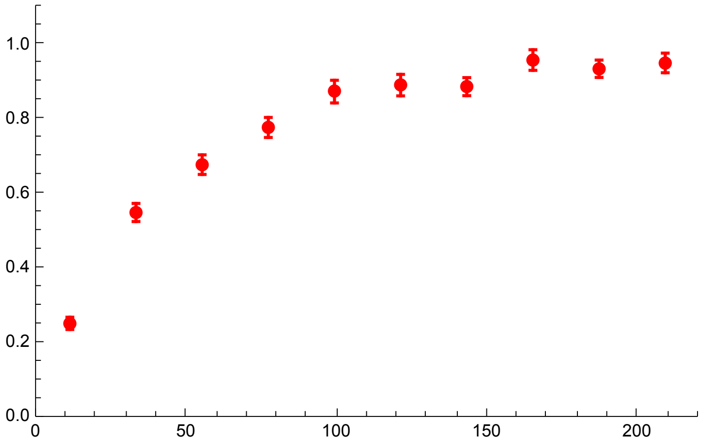

```
Show[Ph01s05, Ph01s05S]
```

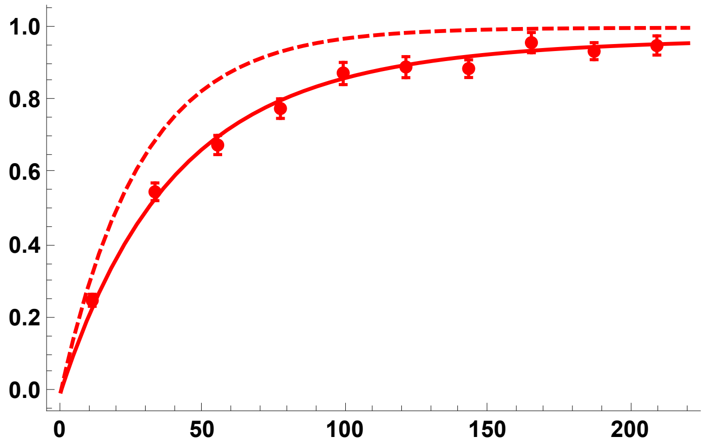


h = 0.9

```
PiRelh09s05 =
  Import["SLIM_F033/StatsProc_R_220_h_0.9_self_0.5_f_1e-04_10b_SLIM.dat",
    "Table"][[10]];
PiRelh09s05CIB = Import[
    "SLIM_F033/StatsProc_R_220_h_0.9_self_0.5_f_1e-04_10b_SLIM.dat",
    "Table"][[11]];
PiRelh09s05CIT = Import[
    "SLIM_F033/StatsProc_R_220_h_0.9_self_0.5_f_1e-04_10b_SLIM.dat",
    "Table"][[12]];
PiRelh09s05S = Partition[Riffle[Partition[Riffle[Rin, PiRelh09s05], 2],
    Map[ErrorBar, Partition[Riffle[-PiRelh09s05CIB, PiRelh09s05CIT], 2]]], 2];
```

```
Ph09s05 = Plot[{PWR2B[5000, 0.05, 0.9, 0.5, R, Boostp0[5000, 0.05, 0.9, 0.5]],
   ExPiSV[5000, 0.05, 0.9, 0.5, R/10 000, Boostp0[5000, 0.05, 0.9, 0.5]]},
  {R, 0, 220}, PlotStyle → { {Blue, Thick, Dashed}, {Blue, Thick}},
  PlotRange → All, Frame → {{True, False}, {True, False}},
  BaseStyle → {FontWeight → "Bold", FontColor → Black, FontSize → 12}]
```

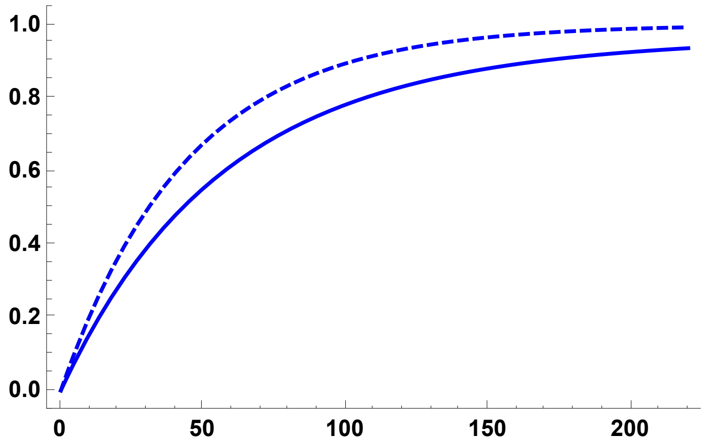

```
Ph09s05A =
  Plot[{ExPiSV[5000, 0.05, 0.9, 0.5, R/10 000, Boostp0[5000, 0.05, 0.9, 0.5]]},
   {R, 0, 220}, PlotStyle → {{Blue, Thick}},
   PlotRange → All, Frame → {{True, False}, {True, False}},
   BaseStyle → {FontWeight → "Bold", FontColor → Black, FontSize → 12}];

Ph09s05S = ErrorListPlot[PiRelh09s05S,
  PlotStyle → {Blue, PointSize[0.02]}, PlotRange → {{0, 220}, {0, 1}}]
```

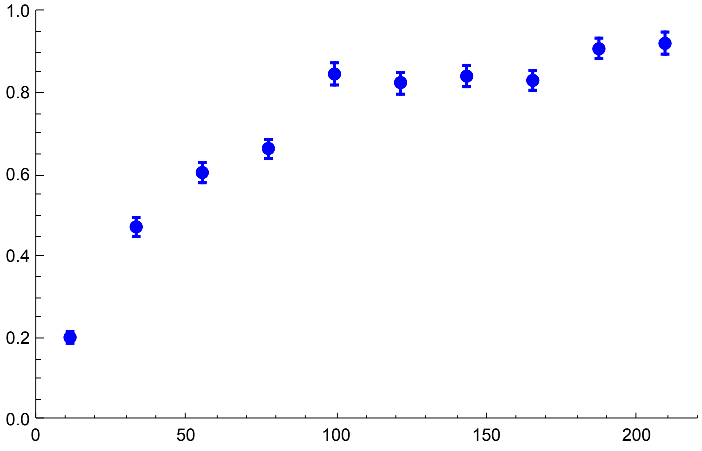


```
Show[Ph09s05, Ph09s05S]
```

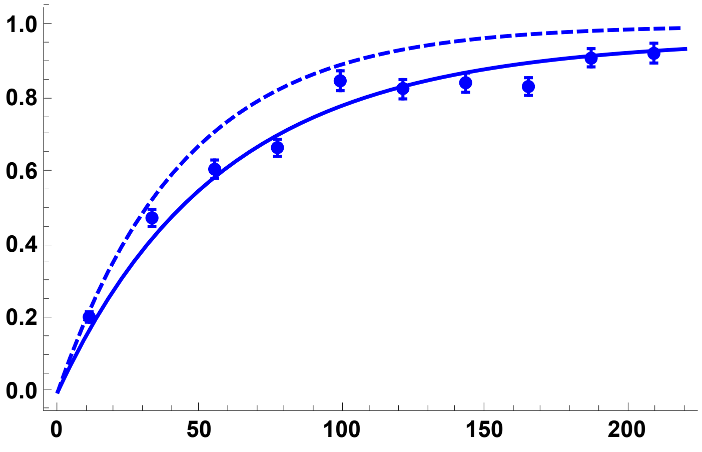


All h values, using the coalescence-in-sweep-phase results and simulations:

```
Sims05DN =
 Show[Ph01s05A, Ph01s05S, Ph05s05A, Ph05s05S, Ph09s05A, Ph09s05S, ImageSize → 250]
```

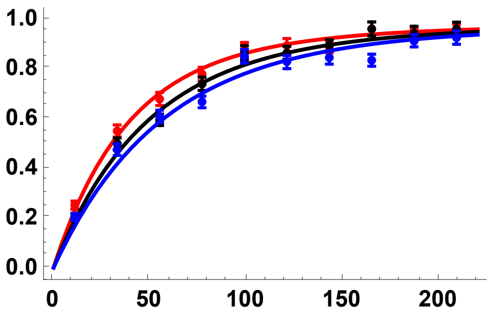

All h values, comparing star-like approximation with coalescence-in-sweep-phase results:

```
Sims05DN2 = Show[Ph01s05, Ph05s05, Ph09s05, ImageSize → 250]
```

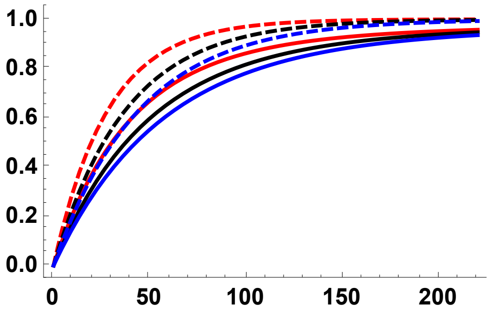


#### Simulation comparisons, $p_0$ = 0.02

h = 0.5

```
PiRelh05p002s05 =
  Import["SLIM_F033/StatsProc_R_220_h_0.5_self_0.5_f_0.02_10b_SLIM.dat",
    "Table"][[10]];
PiRelh05p002s05CIB = Import[
    "SLIM_F033/StatsProc_R_220_h_0.5_self_0.5_f_0.02_10b_SLIM.dat",
    "Table"][[11]];
PiRelh05p002s05CIT = Import[
    "SLIM_F033/StatsProc_R_220_h_0.5_self_0.5_f_0.02_10b_SLIM.dat",
    "Table"][[12]];
PiRelh05p002s05S = Partition[Riffle[
    Partition[Riffle[Rin, PiRelh05p002s05], 2], Map[ErrorBar,
     Partition[Riffle[-PiRelh05p002s05CIB, PiRelh05p002s05CIT], 2]]], 2];
```

```
Ph05p002s05 = Plot[{PWR2B[5000, 0.05, 0.5, 0.5, R, 0.02],
   ExPiSV[5000, 0.05, 0.5, 0.5, R/10 000, 0.02]}, {R, 0, 220},
  PlotStyle → {{Black, Thick, Dashed}, {Black, Thick}},
  PlotRange → All, Frame → {{True, False}, {True, False}},
  BaseStyle → {FontWeight → "Bold", FontColor → Black, FontSize → 12}]
```

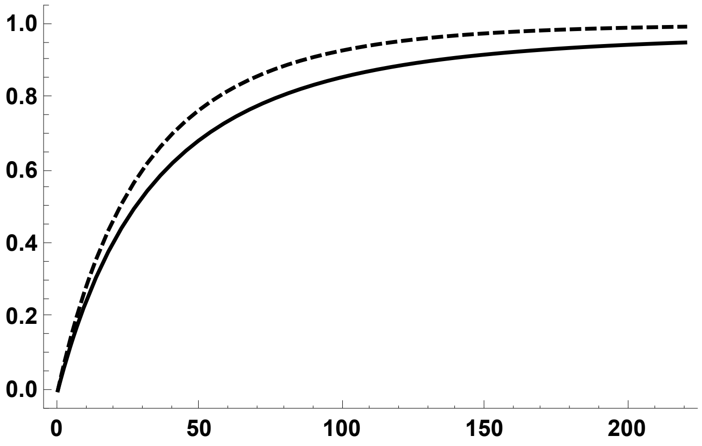

```
Ph05p002s05A = Plot[{ExPiSV[5000, 0.05, 0.5, 0.5, R/10000, 0.02]},
   {R, 0, 220}, PlotStyle → {{Black, Thick}},
   PlotRange → All, Frame → {{True, False}, {True, False}},
   BaseStyle → {FontWeight → "Bold", FontColor → Black, FontSize → 12}];

Ph05p002s05S = ErrorListPlot[PiRelh05p002s05S,
  PlotStyle → {Black, PointSize[0.02]}, PlotRange → {{0, 220}, {0, 1.1}}]
```

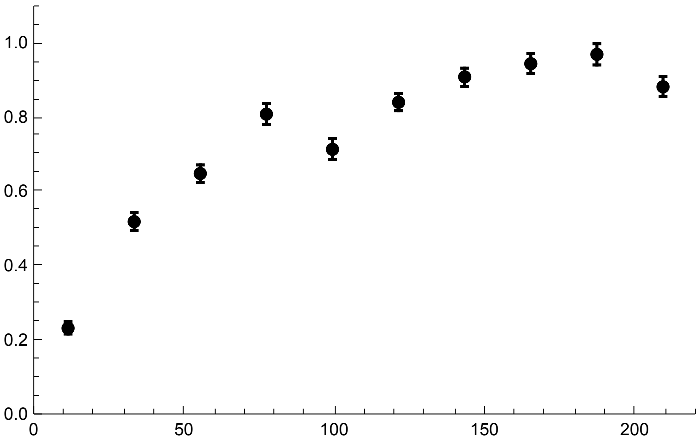


```
Show[Ph05p002s05, Ph05p002s05S]
```

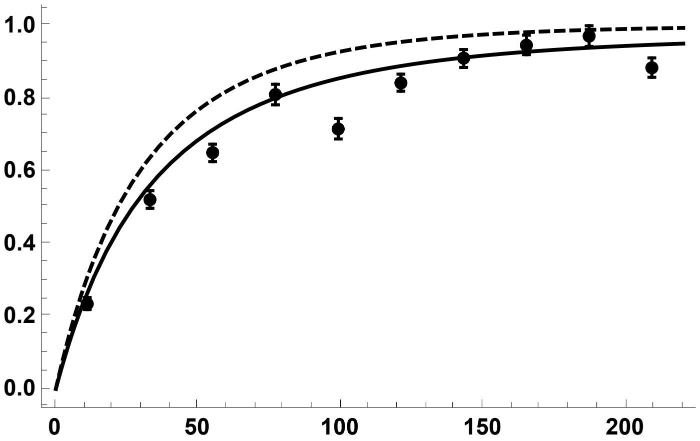


h = 0.1

```
PiRelh01p002s05 =
  Import["SLIM_F033/StatsProc_R_220_h_0.1_self_0.5_f_0.02_10b_SLIM.dat",
    "Table"][[10]];
PiRelh01p002s05CIB = Import[
    "SLIM_F033/StatsProc_R_220_h_0.1_self_0.5_f_0.02_10b_SLIM.dat",
    "Table"][[11]];
PiRelh01p002s05CIT = Import[
    "SLIM_F033/StatsProc_R_220_h_0.1_self_0.5_f_0.02_10b_SLIM.dat",
    "Table"][[12]];
PiRelh01p002s05S = Partition[Riffle[
    Partition[Riffle[Rin, PiRelh01p002s05], 2], Map[ErrorBar,
     Partition[Riffle[-PiRelh01p002s05CIB, PiRelh01p002s05CIT], 2]]], 2];
```

```
Ph01p002s05 = Plot[{PWR2B[5000, 0.05, 0.1, 0.5, R, 0.02],
    ExPiSV[5000, 0.05, 0.1, 0.5, R/10000, 0.02]},
   {R, 0, 220}, PlotStyle → {{Red, Thick, Dashed}, {Red, Thick}},
   PlotRange → All, Frame → {{True, False}, {True, False}},
   BaseStyle → {FontWeight → "Bold", FontColor → Black, FontSize → 12}]
```

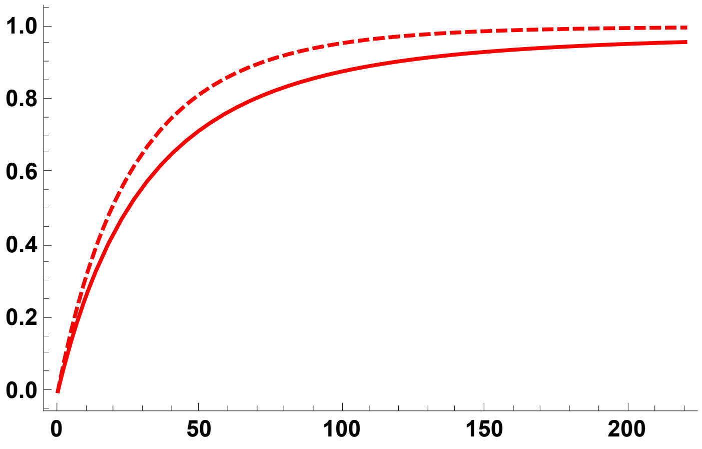


```
Ph01p002s05A = Plot[{ExPiSV[5000, 0.05, 0.1, 0.5, R/10000, 0.02]},
    {R, 0, 220}, PlotStyle → {{Red, Thick}},
    PlotRange → All, Frame → {{True, False}, {True, False}},
    BaseStyle → {FontWeight → "Bold", FontColor → Black, FontSize → 12}];

Ph01p002s05S = ErrorListPlot[PiRelh01p002s05S,
   PlotStyle → {Red, PointSize[0.02]}, PlotRange → {{0, 220}, {0, 1.1}}]
```

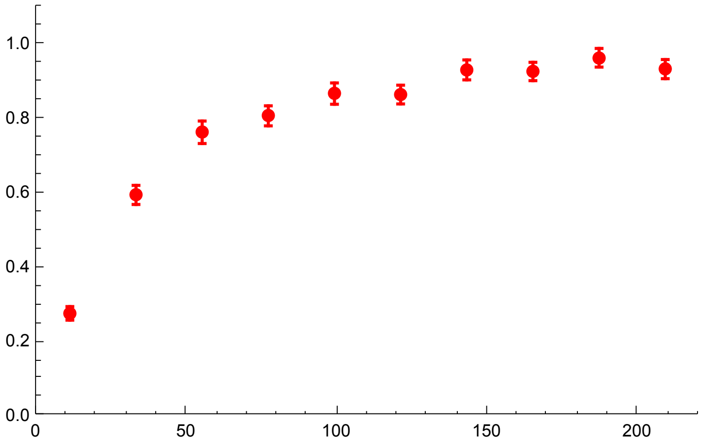

```
Show[Ph01p002s05, Ph01p002s05S]
```

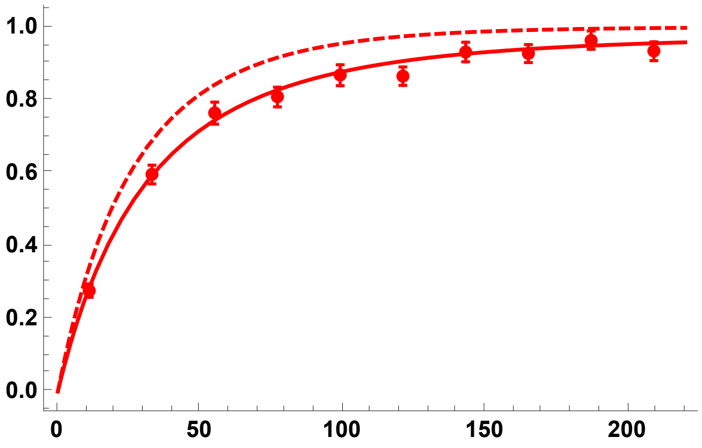


h = 0.9

```
PiRelh09p002s05 =
  Import["SLIM_F033/StatsProc_R_220_h_0.9_self_0.5_f_0.02_10b_SLIM.dat",
    "Table"][[10]];
PiRelh09p002s05CIB = Import[
    "SLIM_F033/StatsProc_R_220_h_0.9_self_0.5_f_0.02_10b_SLIM.dat",
    "Table"][[11]];
PiRelh09p002s05CIT = Import[
    "SLIM_F033/StatsProc_R_220_h_0.9_self_0.5_f_0.02_10b_SLIM.dat",
    "Table"][[12]];
PiRelh09p002s05S = Partition[Riffle[
    Partition[Riffle[Rin, PiRelh09p002s05], 2], Map[ErrorBar,
     Partition[Riffle[-PiRelh09p002s05CIB, PiRelh09p002s05CIT], 2]]], 2];
```

```
Ph09p002s05 = Plot[{PWR2B[5000, 0.05, 0.9, 0.5, R, 0.02],
   ExPiSV[5000, 0.05, 0.9, 0.5, R/10000, 0.02]},
  {R, 0, 220}, PlotStyle → { {Blue, Thick, Dashed}, {Blue, Thick}},
  PlotRange → All, Frame → {{True, False}, {True, False}},
  BaseStyle → {FontWeight → "Bold", FontColor → Black, FontSize → 12}]
```

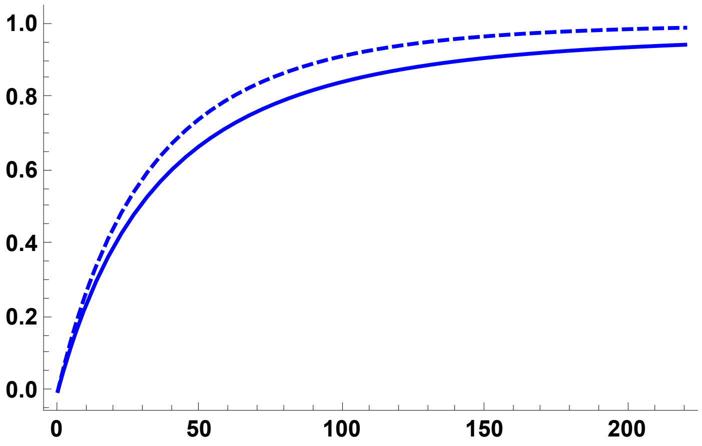

```
Ph09p002s05A = Plot[{ExPiSV[5000, 0.05, 0.9, 0.5, R/10 000, 0.02]},
    {R, 0, 220}, PlotStyle → { {Blue, Thick}},
    PlotRange → All, Frame → {{True, False}, {True, False}},
    BaseStyle → {FontWeight → "Bold", FontColor → Black, FontSize → 12}];
```

```
Ph09p002s05S = ErrorListPlot[PiRelh09p002s05S,
   PlotStyle → {Blue, PointSize[0.02]}, PlotRange → {{0, 220}, {0, 1.1}}]
```

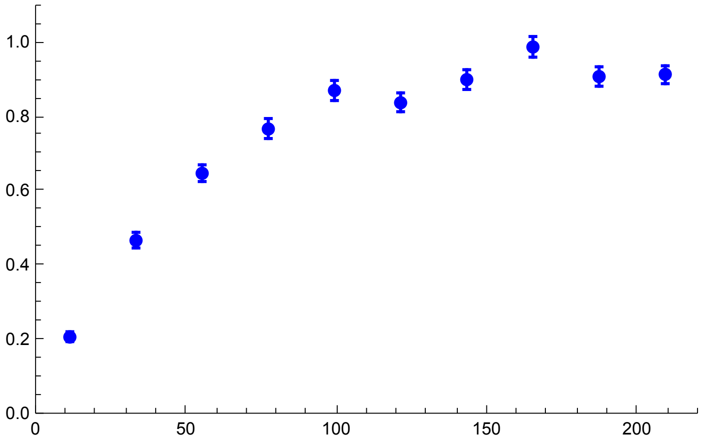


```
Show[Ph09p002s05, Ph09p002s05S]
```

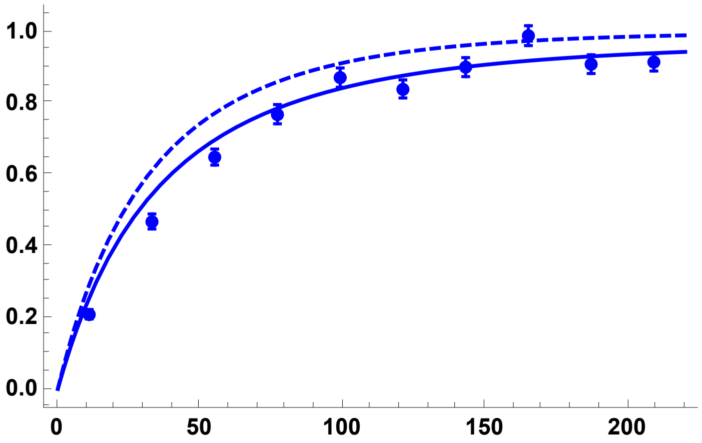


All h values, using the coalescence-in-sweep-phase results and simulations:

```
Sims052pc = Show[Ph01p002s05A, Ph01p002s05S, Ph05p002s05A,
   Ph05p002s05S, Ph09p002s05A, Ph09p002s05S, ImageSize → 250]
```

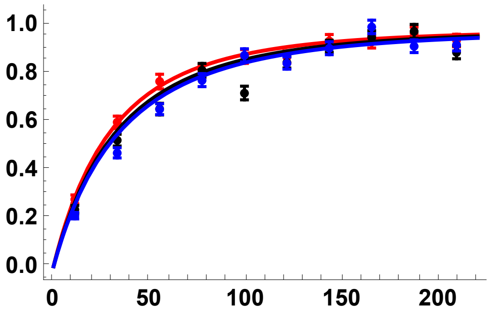


All h values, comparing star-like approximation with coalescence-in-sweep-phase results:

```
Sims052pc2 = Show[Ph01p002s05, Ph05p002s05, Ph09p002s05, ImageSize → 250]
```

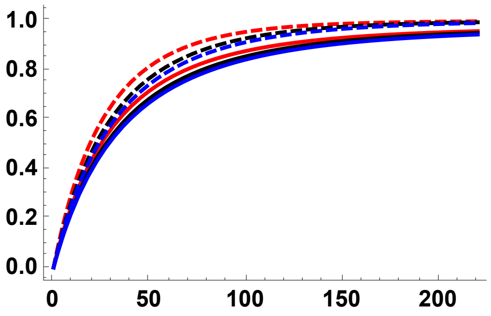


#### Simulation comparisons, $p_0$ = 0.05

h = 0.5

```
PiRelh05p005s05 =
  Import["SLIM_F033/StatsProc_R_220_h_0.5_self_0.5_f_0.05_10b_SLIM.dat",
    "Table"][[10]];
PiRelh05p005s05CIB = Import[
    "SLIM_F033/StatsProc_R_220_h_0.5_self_0.5_f_0.05_10b_SLIM.dat",
    "Table"][[11]];
PiRelh05p005s05CIT = Import[
    "SLIM_F033/StatsProc_R_220_h_0.5_self_0.5_f_0.05_10b_SLIM.dat",
    "Table"][[12]];
PiRelh05p005s05S = Partition[Riffle[
    Partition[Riffle[Rin, PiRelh05p005s05], 2], Map[ErrorBar,
     Partition[Riffle[-PiRelh05p005s05CIB, PiRelh05p005s05CIT], 2]]], 2];
```

```
Ph05p005s05 = Plot[{PWR2B[5000, 0.05, 0.5, 0.5, R, 0.05],
   ExPiSV[5000, 0.05, 0.5, 0.5, R/10 000, 0.05]}, {R, 0, 220},
  PlotStyle → {{Black, Thick, Dashed}, {Black, Thick}},
  PlotRange → All, Frame → {{True, False}, {True, False}},
  BaseStyle → {FontWeight → "Bold", FontColor → Black, FontSize → 12}]
```

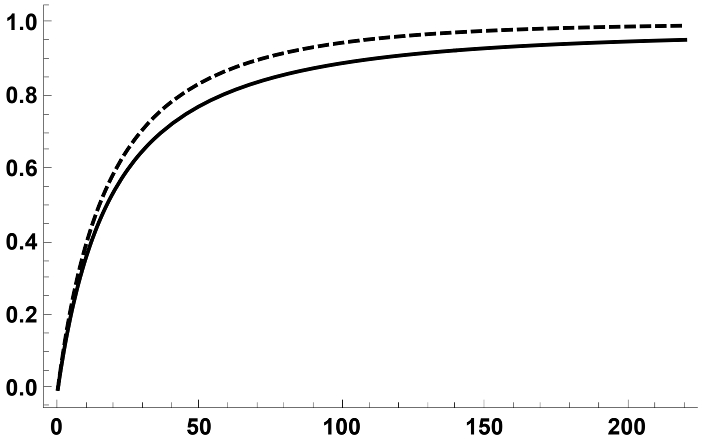

```
Ph05p005s05A = Plot[{ExPiSV[5000, 0.05, 0.5, 0.5, R/10000, 0.05]},
  {R, 0, 220}, PlotStyle → {{Black, Thick}},
  PlotRange → All, Frame → {{True, False}, {True, False}},
  BaseStyle → {FontWeight → "Bold", FontColor → Black, FontSize → 12}];

Ph05p005s05S = ErrorListPlot[PiRelh05p005s05S,
  PlotStyle → {Black, PointSize[0.02]}, PlotRange → {{0, 220}, {0, 1.1}}]
```

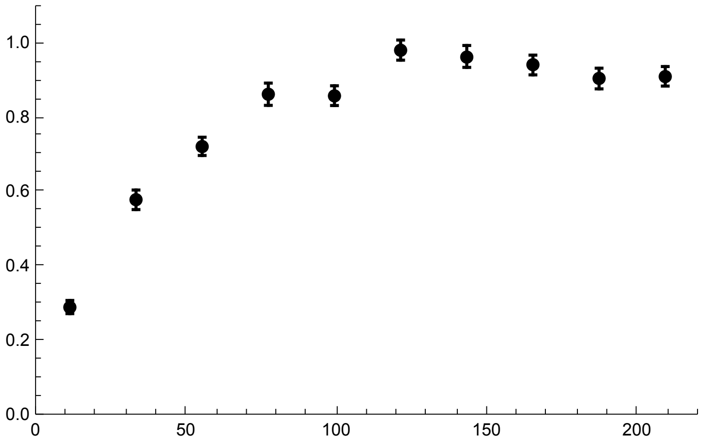


```
Show[Ph05p005s05, Ph05p005s05S]
```

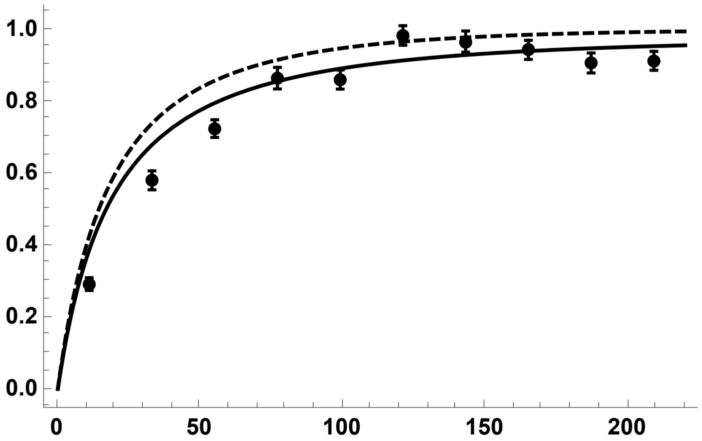


h = 0.1

```
PiRelh01p005s05 =
  Import["SLIM_F033/StatsProc_R_220_h_0.1_self_0.5_f_0.05_10b_SLIM.dat",
    "Table"][[10]];
PiRelh01p005s05CIB = Import[
    "SLIM_F033/StatsProc_R_220_h_0.1_self_0.5_f_0.05_10b_SLIM.dat",
    "Table"][[11]];
PiRelh01p005s05CIT = Import[
    "SLIM_F033/StatsProc_R_220_h_0.1_self_0.5_f_0.05_10b_SLIM.dat",
    "Table"][[12]];
PiRelh01p005s05S = Partition[Riffle[
    Partition[Riffle[Rin, PiRelh01p005s05], 2], Map[ErrorBar,
     Partition[Riffle[-PiRelh01p005s05CIB, PiRelh01p005s05CIT], 2]]], 2];
```

```
Ph01p005s05 = Plot[{PWR2B[5000, 0.05, 0.1, 0.5, R, 0.05],
   ExPiSV[5000, 0.05, 0.1, 0.5, R/10000, 0.05]},
  {R, 0, 220}, PlotStyle → {{Red, Thick, Dashed}, {Red, Thick}},
  PlotRange → All, Frame → {{True, False}, {True, False}},
  BaseStyle → {FontWeight → "Bold", FontColor → Black, FontSize → 12}]
```

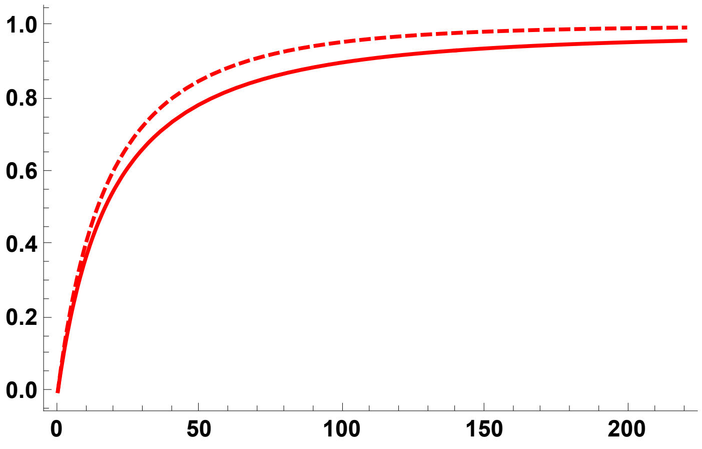


```
Ph01p005s05A = Plot[{ExPiSV[5000, 0.05, 0.1, 0.5, R/10000, 0.05]},
   {R, 0, 220}, PlotStyle → {{Red, Thick}},
   PlotRange → All, Frame → {{True, False}, {True, False}},
   BaseStyle → {FontWeight → "Bold", FontColor → Black, FontSize → 12}];

Ph01p005s05S = ErrorListPlot[PiRelh01p005s05S,
  PlotStyle → {Red, PointSize[0.02]}, PlotRange → {{0, 220}, {0, 1.1}}]
```

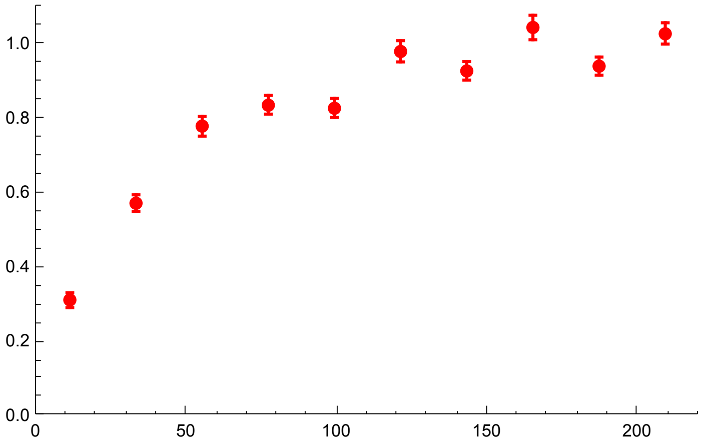

```
Show[Ph01p005s05, Ph01p005s05S]
```

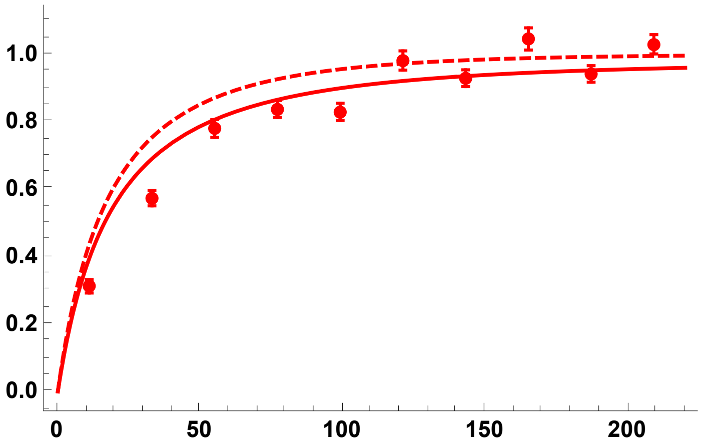


h = 0.9

```
PiRelh09p005s05 =
  Import["SLIM_F033/StatsProc_R_220_h_0.9_self_0.5_f_0.05_10b_SLIM.dat",
    "Table"][[10]];
PiRelh09p005s05CIB = Import[
    "SLIM_F033/StatsProc_R_220_h_0.9_self_0.5_f_0.05_10b_SLIM.dat",
    "Table"][[11]];
PiRelh09p005s05CIT = Import[
    "SLIM_F033/StatsProc_R_220_h_0.9_self_0.5_f_0.05_10b_SLIM.dat",
    "Table"][[12]];
PiRelh09p005s05S = Partition[Riffle[
    Partition[Riffle[Rin, PiRelh09p005s05], 2], Map[ErrorBar,
     Partition[Riffle[-PiRelh09p005s05CIB, PiRelh09p005s05CIT], 2]]], 2];
```

```
Ph09p005s05 = Plot[{PWR2B[5000, 0.05, 0.9, 0.5, R, 0.05],
   ExPiSV[5000, 0.05, 0.9, 0.5, R/10000, 0.05]},
  {R, 0, 220}, PlotStyle → { {Blue, Thick, Dashed}, {Blue, Thick}},
  PlotRange → All, Frame → {{True, False}, {True, False}},
  BaseStyle → {FontWeight → "Bold", FontColor → Black, FontSize → 12}]
```

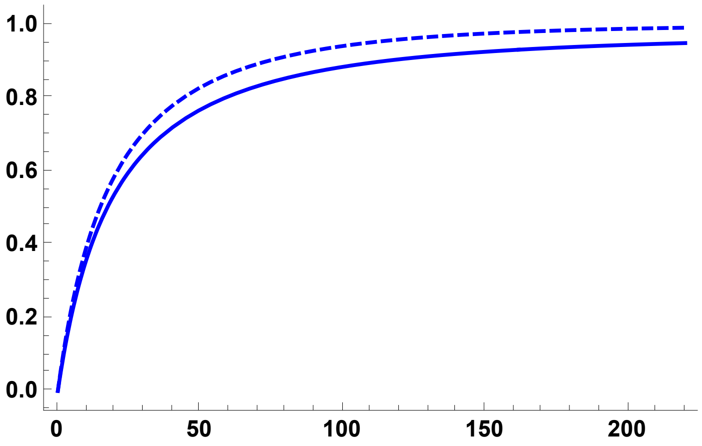

```
Ph09p005s05A = Plot[{ExPiSV[5000, 0.05, 0.9, 0.5, R/10000, 0.05]},
   {R, 0, 220}, PlotStyle → { {Blue, Thick}},
   PlotRange → All, Frame → {{True, False}, {True, False}},
   BaseStyle → {FontWeight → "Bold", FontColor → Black, FontSize → 12}];
```

```
Ph09p005s05S = ErrorListPlot[PiRelh09p005s05S,
  PlotStyle → {Blue, PointSize[0.02]}, PlotRange → {{0, 220}, {0, 1.1}}]
```

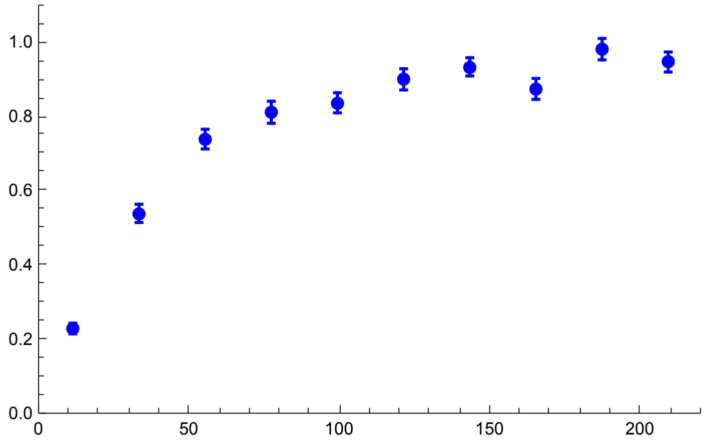


```
Show[Ph09p005s05, Ph09p005s05S]
```

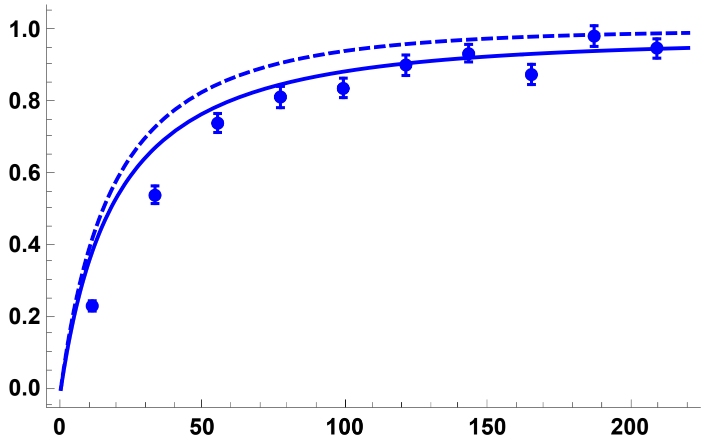


All h values, using the coalescence-in-sweep-phase results and simulations:

```
Sims055pc = Show[Ph01p005s05A, Ph01p005s05S, Ph05p005s05A,
  Ph05p005s05S, Ph09p005s05A, Ph09p005s05S, ImageSize → 250]
```

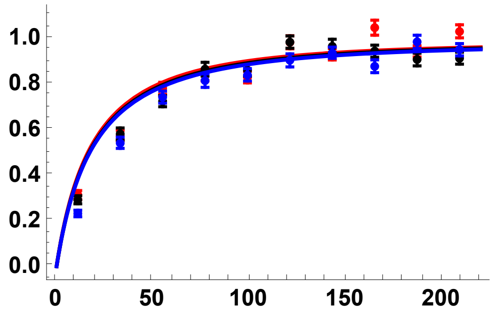


All h values, comparing star-like approximation with coalescence-in-sweep-phase results:

```
Sims055pc2 = Show[Ph01p005s05, Ph05p005s05, Ph09p005s05, ImageSize → 250]
```

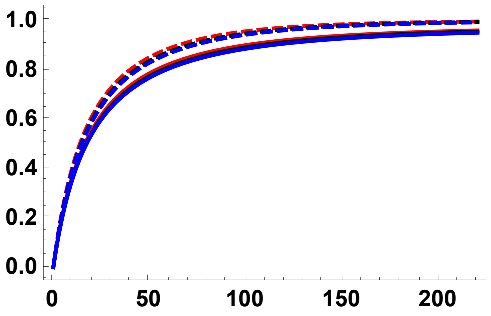


#### $\sigma = 0.95$ ($F \simeq 0.904$)

Preliminaries

```
SetDirectory[NotebookDirectory[]];
Rin = Table[100 + 200 * i, {i, 0, 9}];
```

##### Simulation comparisons, from initial frequency $p_0 = 1/2N$

h = 0.5

```
PiRelh05s095 =
  Import["SLIM_F09/StatsProc_R_2000_h_0.5_self_0.95_f_1e-04_10b_SLIM.dat",
    "Table"][[10]];
PiRelh05s095CIB = Import[
    "SLIM_F09/StatsProc_R_2000_h_0.5_self_0.95_f_1e-04_10b_SLIM.dat",
    "Table"][[11]];
PiRelh05s095CIT = Import[
    "SLIM_F09/StatsProc_R_2000_h_0.5_self_0.95_f_1e-04_10b_SLIM.dat",
    "Table"][[12]];
PiRelh05s095S = Partition[Riffle[Partition[Riffle[Rin, PiRelh05s095], 2],
    Map[ErrorBar, Partition[Riffle[-PiRelh05s095CIB, PiRelh05s095CIT], 2]]], 2];
```

```
Ph05s095 = Plot[{PWR2B[5000, 0.05, 0.5, 0.95, R, Boostp0[5000, 0.05, 0.5, 0.95]],
   ExPiSV[5000, 0.05, 0.5, 0.95, R/10 000, Boostp0[5000, 0.05, 0.5, 0.95]]},
  {R, 0, 2000}, PlotStyle → {{Black, Thick, Dashed}, {Black, Thick}},
  PlotRange → All, Frame → {{True, False}, {True, False}},
  BaseStyle → {FontWeight → "Bold", FontColor → Black, FontSize → 12}]
```

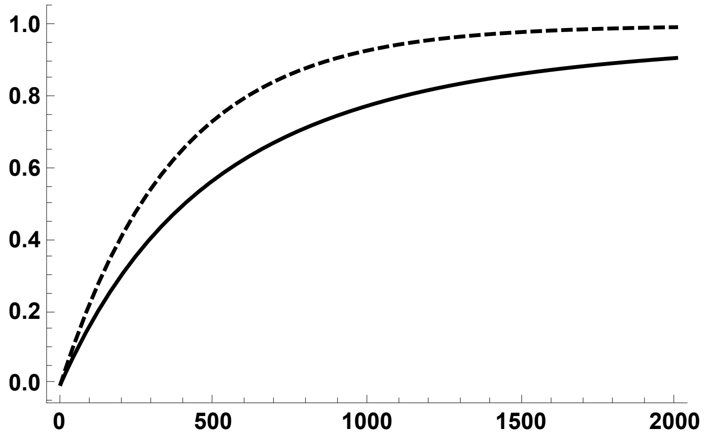


```
Ph05s095A =
  Plot[{ExPiSV[5000, 0.05, 0.5, 0.95, R/10 000, Boostp0[5000, 0.05, 0.5, 0.95]]},
   {R, 0, 2000}, PlotStyle → {{Black, Thick}},
   PlotRange → All, Frame → {{True, False}, {True, False}},
   BaseStyle → {FontWeight → "Bold", FontColor → Black, FontSize → 12}];

Ph05s095S = ErrorListPlot[PiRelh05s095S,
  PlotStyle → {Black, PointSize[0.02]}, PlotRange → {{0, 2000}, {0, 1.1}}]
```

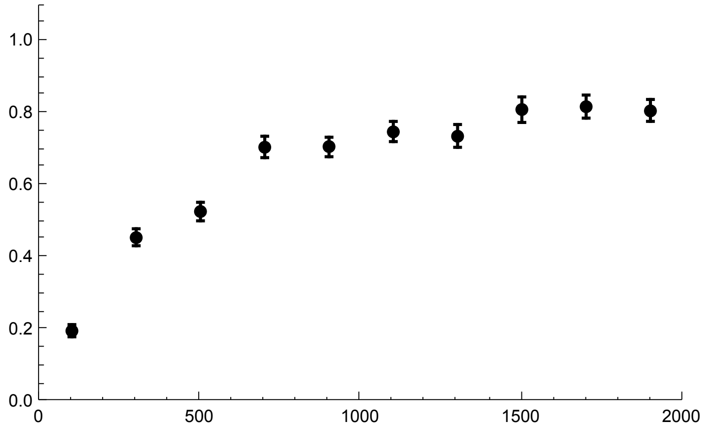

```
Show[Ph05s095, Ph05s095S]
```

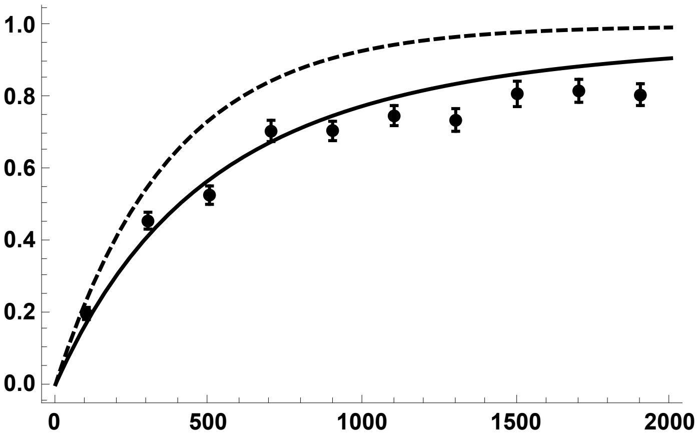


h = 0.1

```
PiRelh01s095 =
  Import["SLIM_F09/StatsProc_R_2000_h_0.1_self_0.95_f_1e-04_10b_SLIM.dat",
    "Table"][[10]];
PiRelh01s095CIB = Import[
    "SLIM_F09/StatsProc_R_2000_h_0.1_self_0.95_f_1e-04_10b_SLIM.dat",
    "Table"][[11]];
PiRelh01s095CIT = Import[
    "SLIM_F09/StatsProc_R_2000_h_0.1_self_0.95_f_1e-04_10b_SLIM.dat",
    "Table"][[12]];
PiRelh01s095S = Partition[Riffle[Partition[Riffle[Rin, PiRelh01s095], 2],
    Map[ErrorBar, Partition[Riffle[-PiRelh01s095CIB, PiRelh01s095CIT], 2]]], 2];
```

```
Ph01s095 = Plot[{PWR2B[5000, 0.05, 0.1, 0.95, R, Boostp0[5000, 0.05, 0.1, 0.95]],
   ExPiSV[5000, 0.05, 0.1, 0.95, R/10000, Boostp0[5000, 0.05, 0.1, 0.95]]},
  {R, 0, 2000}, PlotStyle → {{Red, Thick, Dashed}, {Red, Thick}},
  PlotRange → {All, {0, 1}}, Frame → {{True, False}, {True, False}},
  BaseStyle → {FontWeight → "Bold", FontColor → Black, FontSize → 12}]
```

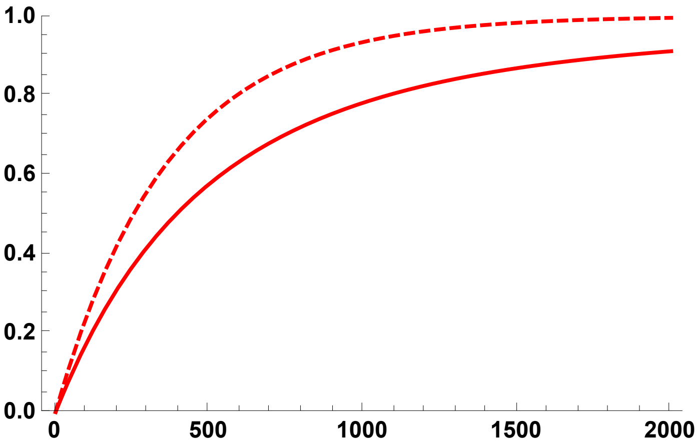

```
Ph01s095A =
  Plot[{ExPiSV[5000, 0.05, 0.1, 0.95, R/10000], Boostp0[5000, 0.05, 0.1, 0.95]},
   {R, 0, 2000}, PlotStyle → {{Red, Thick}},
   PlotRange → {All, {0, 1}}, Frame → {{True, False}, {True, False}},
   BaseStyle → {FontWeight → "Bold", FontColor → Black, FontSize → 12}];

Ph01s095S = ErrorListPlot[PiRelh01s095S,
  PlotStyle → {Red, PointSize[0.02]}, PlotRange → {{0, 2000}, {0, 1.1}}]
```

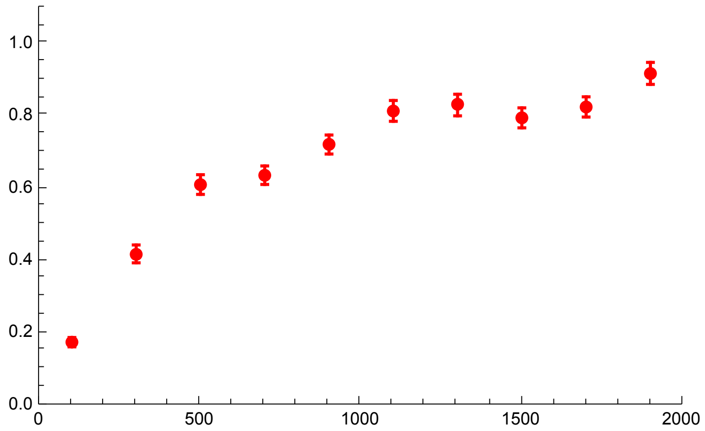


```
Show[Ph01s095, Ph01s095S]
```

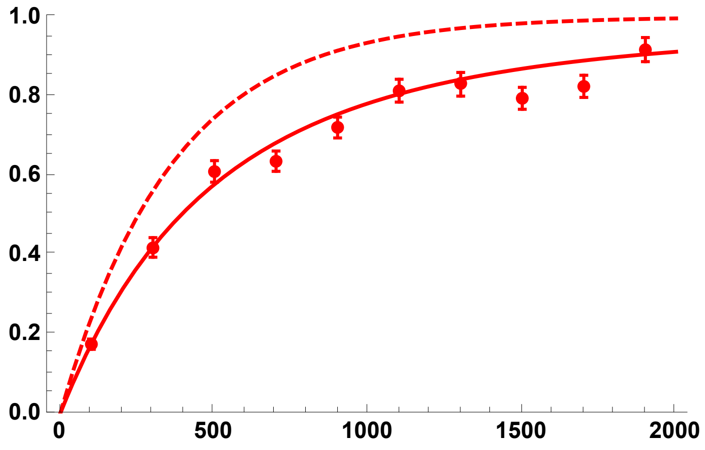


h = 0.9

```
PiRelh09s095 =
  Import["SLIM_F09/StatsProc_R_2000_h_0.9_self_0.95_f_1e-04_10b_SLIM.dat",
    "Table"][[10]];
PiRelh09s095CIB = Import[
    "SLIM_F09/StatsProc_R_2000_h_0.9_self_0.95_f_1e-04_10b_SLIM.dat",
    "Table"][[11]];
PiRelh09s095CIT = Import[
    "SLIM_F09/StatsProc_R_2000_h_0.9_self_0.95_f_1e-04_10b_SLIM.dat",
    "Table"][[12]];
PiRelh09s095S = Partition[Riffle[Partition[Riffle[Rin, PiRelh09s095], 2],
    Map[ErrorBar, Partition[Riffle[-PiRelh09s095CIB, PiRelh09s095CIT], 2]]], 2];
```

```
Ph09s095 = Plot[{PWR2B[5000, 0.05, 0.9, 0.95, R, Boostp0[5000, 0.05, 0.9, 0.95]],
   ExPiSV[5000, 0.05, 0.9, 0.95, R/10 000, Boostp0[5000, 0.05, 0.9, 0.95]]},
  {R, 0, 2000}, PlotStyle → { {Blue, Thick, Dashed}, {Blue, Thick}},
  PlotRange → {All, {0, 1}}, Frame → {{True, False}, {True, False}},
  BaseStyle → {FontWeight → "Bold", FontColor → Black, FontSize → 12}]
```

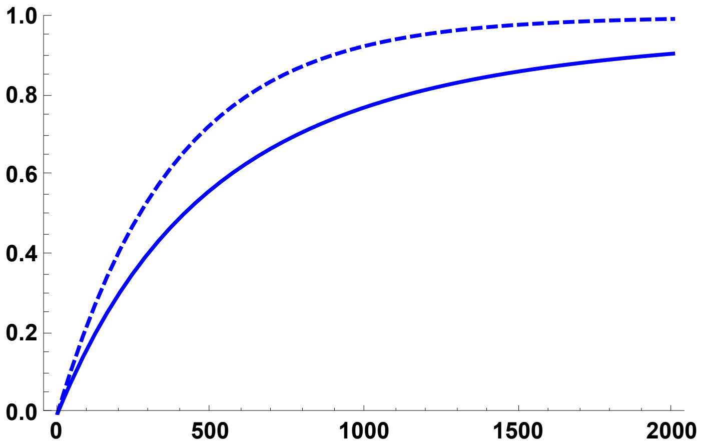


```
Ph09s095A =
  Plot[{ExPiSV[5000, 0.05, 0.9, 0.95, R/10 000, Boostp0[5000, 0.05, 0.9, 0.95]]},
   {R, 0, 2000}, PlotStyle → { {Blue, Thick}},
   PlotRange → {All, {0, 1}}, Frame → {{True, False}, {True, False}},
   BaseStyle → {FontWeight → "Bold", FontColor → Black, FontSize → 12}];

Ph09s095S = ErrorListPlot[PiRelh09s095S,
  PlotStyle → {Blue, PointSize[0.02]}, PlotRange → {{0, 2000}, {0, 1}}]
```

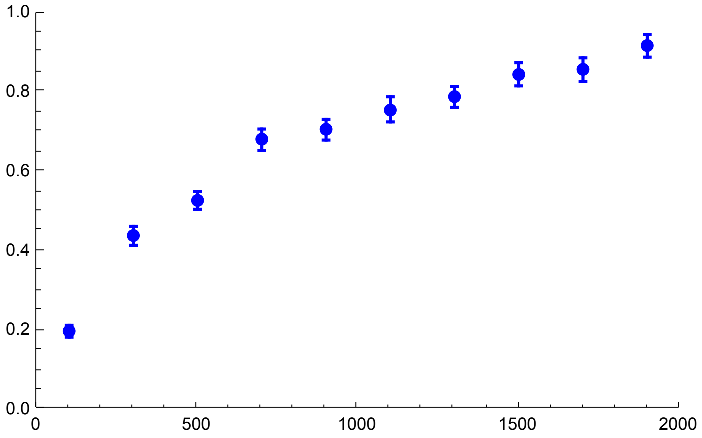

```
Show[Ph09s095, Ph09s095S]
```

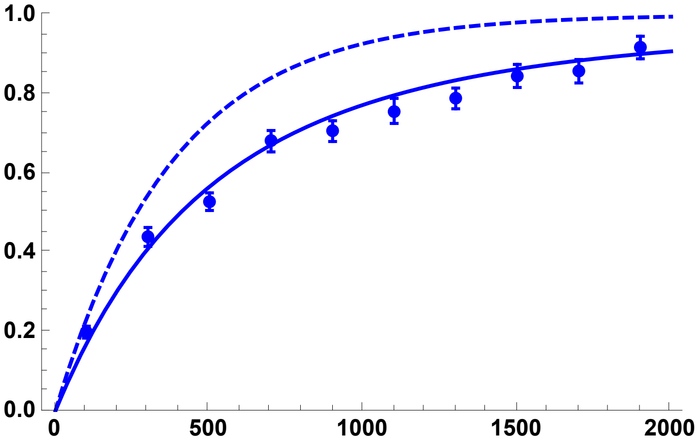


All h values, using the coalescence-in-sweep-phase results and simulations:

```
Sims095DN = Show[Ph01s095A, Ph01s095S,
  Ph05s095A, Ph05s095S, Ph09s095A, Ph09s095S, ImageSize → 250]
```

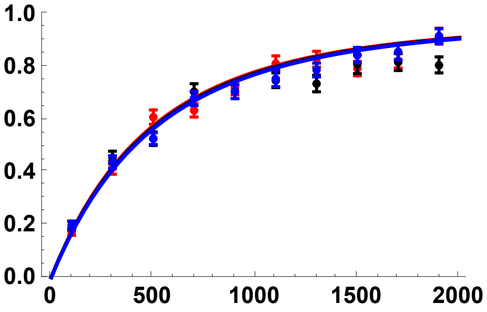


All h values, using the coalescence-in-sweep-phase results and simulations:

```
Sims095DN2 = Show[Ph01s095, Ph05s095, Ph09s095, ImageSize → 250]
```

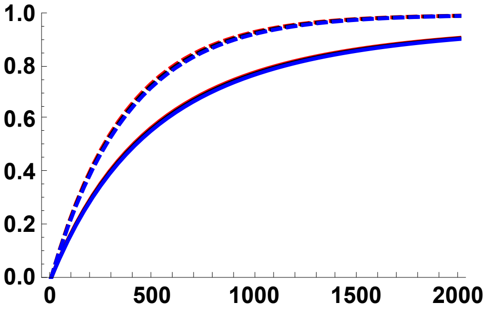


#### Simulation comparisons, $p_0$ = 0.02

h = 0.5

```
PiRelh05p002s095 =
  Import["SLIM_F09/StatsProc_R_2000_h_0.5_self_0.95_f_0.02_10b_SLIM.dat",
    "Table"][[10]];
PiRelh05p002s095CIB = Import[
    "SLIM_F09/StatsProc_R_2000_h_0.5_self_0.95_f_0.02_10b_SLIM.dat",
    "Table"][[11]];
PiRelh05p002s095CIT = Import[
    "SLIM_F09/StatsProc_R_2000_h_0.5_self_0.95_f_0.02_10b_SLIM.dat",
    "Table"][[12]];
PiRelh05p002s095S = Partition[Riffle[
    Partition[Riffle[Rin, PiRelh05p002s095], 2], Map[ErrorBar,
     Partition[Riffle[-PiRelh05p002s095CIB, PiRelh05p002s095CIT], 2]]], 2];
```

```
Ph05p002s095 = Plot[{PWR2B[5000, 0.05, 0.5, 0.95, R, 0.02],
   ExPiSV[5000, 0.05, 0.5, 0.95, R/10000, 0.02]}, {R, 0, 2000},
  PlotStyle → {{Black, Thick, Dashed}, {Black, Thick}},
  PlotRange → All, Frame → {{True, False}, {True, False}},
  BaseStyle → {FontWeight → "Bold", FontColor → Black, FontSize → 12}]
```

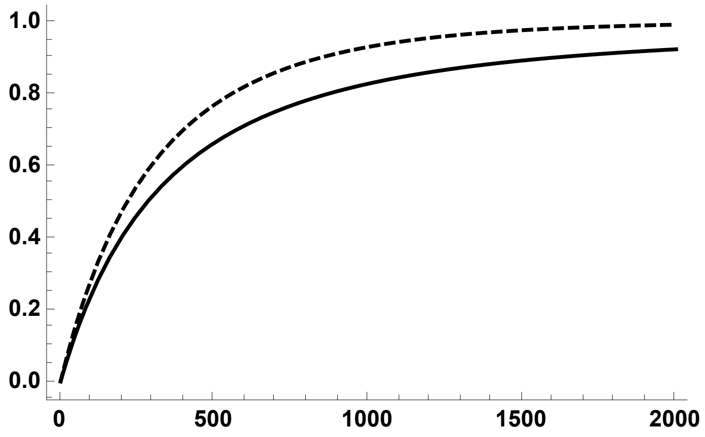


```
Ph05p002s095A = Plot[{ExPiSV[5000, 0.05, 0.5, 0.95, R/10000, 0.02]},
   {R, 0, 2000}, PlotStyle → {{Black, Thick}},
   PlotRange → All, Frame → {{True, False}, {True, False}},
   BaseStyle → {FontWeight → "Bold", FontColor → Black, FontSize → 12}];
```

```
Ph05p002s095S = ErrorListPlot[PiRelh05p002s095S,
  PlotStyle → {Black, PointSize[0.02]}, PlotRange → {{0, 2000}, {0, 1.1}}]
```

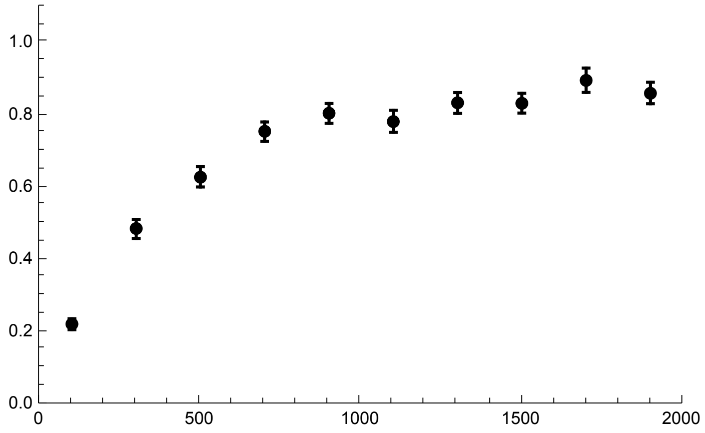


```
Show[Ph05p002s095, Ph05p002s095S]
```

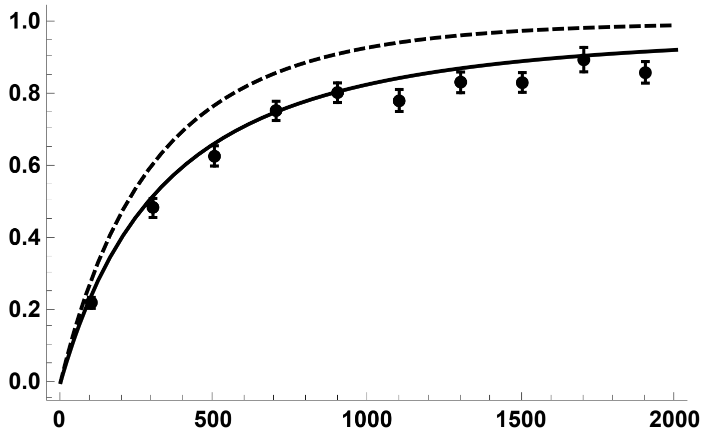


h = 0.1

```
PiRelh01p002s095 =
  Import["SLIM_F09/StatsProc_R_2000_h_0.1_self_0.95_f_0.02_10b_SLIM.dat",
    "Table"][[10]];
PiRelh01p002s095CIB = Import[
    "SLIM_F09/StatsProc_R_2000_h_0.1_self_0.95_f_0.02_10b_SLIM.dat",
    "Table"][[11]];
PiRelh01p002s095CIT = Import[
    "SLIM_F09/StatsProc_R_2000_h_0.1_self_0.95_f_0.02_10b_SLIM.dat",
    "Table"][[12]];
PiRelh01p002s095S = Partition[Riffle[
    Partition[Riffle[Rin, PiRelh01p002s095], 2], Map[ErrorBar,
     Partition[Riffle[-PiRelh01p002s095CIB, PiRelh01p002s095CIT], 2]]], 2];
```

```
Ph01p002s095 = Plot[{PWR2B[5000, 0.05, 0.1, 0.95, R, 0.02],
    ExPiSV[5000, 0.05, 0.1, 0.95, R/10000, 0.02]},
  {R, 0, 2000}, PlotStyle → {{Red, Thick, Dashed}, {Red, Thick}},
  PlotRange → All, Frame → {{True, False}, {True, False}},
  BaseStyle → {FontWeight → "Bold", FontColor → Black, FontSize → 12}]
```

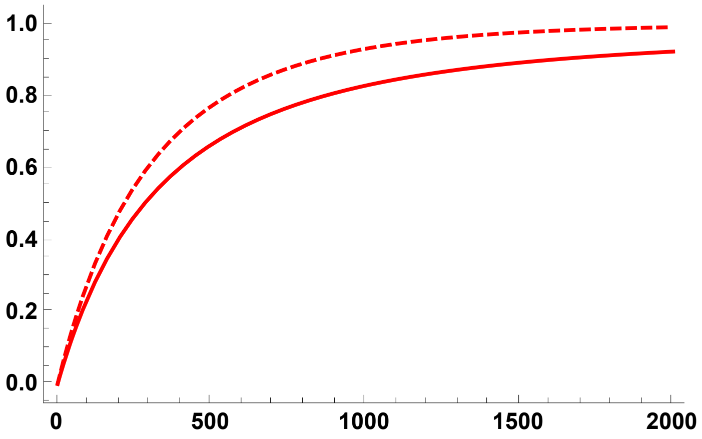

```
Ph01p002s095A = Plot[{ExPiSV[5000, 0.05, 0.1, 0.95, R/10000, 0.02]},
   {R, 0, 2000}, PlotStyle → {{Red, Thick}},
   PlotRange → All, Frame → {{True, False}, {True, False}},
   BaseStyle → {FontWeight → "Bold", FontColor → Black, FontSize → 12}];

Ph01p002s095S = ErrorListPlot[PiRelh01p002s095S,
  PlotStyle → {Red, PointSize[0.02]}, PlotRange → {{0, 2000}, {0, 1.1}}]
```

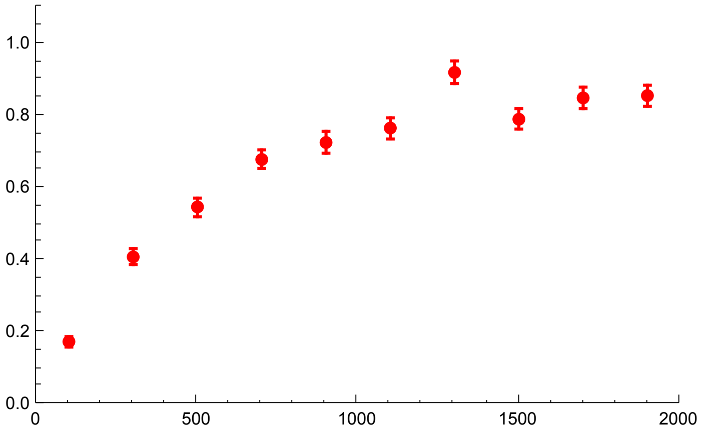

```
Show[Ph01p002s095, Ph01p002s095S]
```

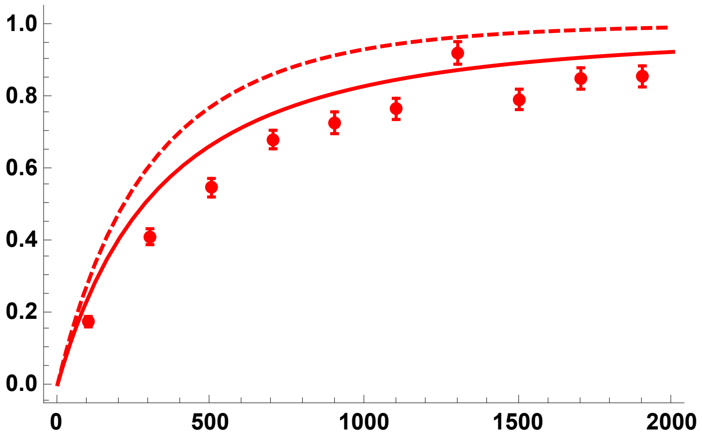


h = 0.9

```
PiRelh09p002s095 =
  Import["SLIM_F09/StatsProc_R_2000_h_0.9_self_0.95_f_0.02_10b_SLIM.dat",
    "Table"][[10]];
PiRelh09p002s095CIB = Import[
    "SLIM_F09/StatsProc_R_2000_h_0.9_self_0.95_f_0.02_10b_SLIM.dat",
    "Table"][[11]];
PiRelh09p002s095CIT = Import[
    "SLIM_F09/StatsProc_R_2000_h_0.9_self_0.95_f_0.02_10b_SLIM.dat",
    "Table"][[12]];
PiRelh09p002s095S = Partition[Riffle[
    Partition[Riffle[Rin, PiRelh09p002s095], 2], Map[ErrorBar,
     Partition[Riffle[-PiRelh09p002s095CIB, PiRelh09p002s095CIT], 2]]], 2];
```

```
Ph09p002s095 = Plot[{PWR2B[5000, 0.05, 0.9, 0.95, R, 0.02],
   ExPiSV[5000, 0.05, 0.9, 0.95, R/10000, 0.02]},
  {R, 0, 2000}, PlotStyle → { {Blue, Thick, Dashed}, {Blue, Thick}},
  PlotRange → All, Frame → {{True, False}, {True, False}},
  BaseStyle → {FontWeight → "Bold", FontColor → Black, FontSize → 12}]
```

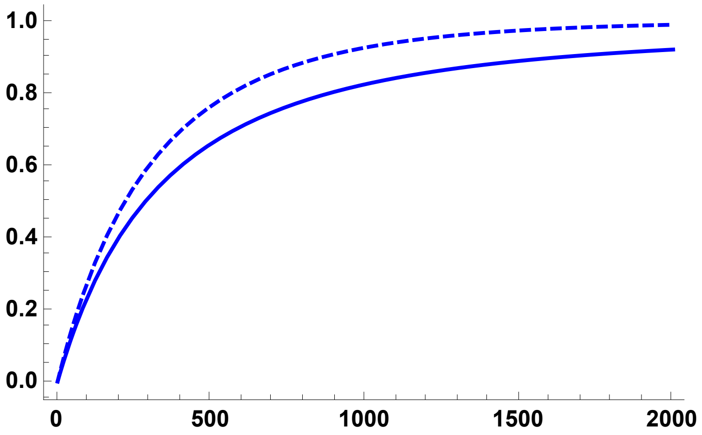

```
Ph09p002s095A = Plot[{ExPiSV[5000, 0.05, 0.9, 0.95, R/10000, 0.02]},
    {R, 0, 2000}, PlotStyle → { {Blue, Thick}},
    PlotRange → All, Frame → {{True, False}, {True, False}},
    BaseStyle → {FontWeight → "Bold", FontColor → Black, FontSize → 12}];
```

```
Ph09p002s095S = ErrorListPlot[PiRelh09p002s095S,
  PlotStyle → {Blue, PointSize[0.02]}, PlotRange → {{0, 2000}, {0, 1.1}}]
```

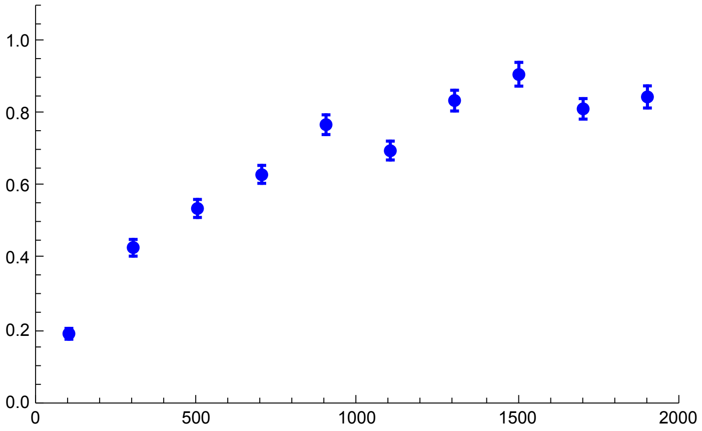


```
Show[Ph09p002s095, Ph09p002s095S]
```

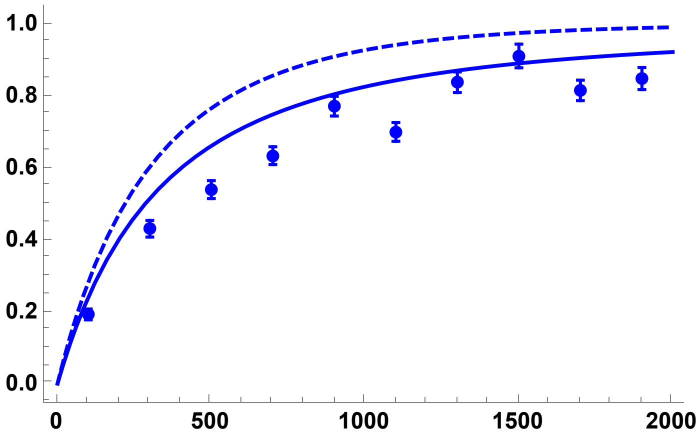


All h values, using the coalescence-in-sweep-phase results and simulations:

```
Sims0952pc = Show[Ph01p002s095A, Ph01p002s095S, Ph05p002s095A,
  Ph05p002s095S, Ph09p002s095A, Ph09p002s095S, ImageSize → 250]
```

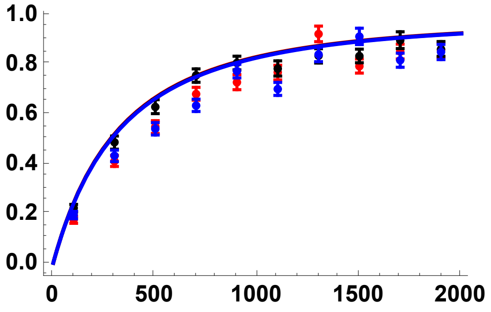


All h values, using the coalescence-in-sweep-phase results and simulations:

```
Sims0952pc2 = Show[Ph01p002s095, Ph05p002s095, Ph09p002s095, ImageSize → 250]
```

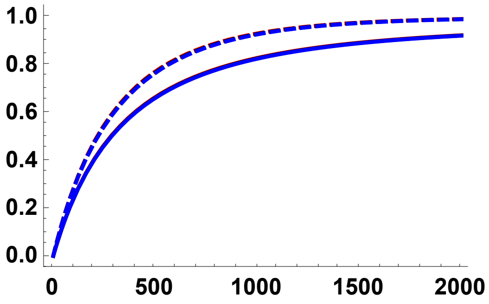


#### Simulation comparisons, $p_0$ = 0.05

h = 0.5

```
PiRelh05p005s095 =
  Import["SLIM_F09/StatsProc_R_2000_h_0.5_self_0.95_f_0.05_10b_SLIM.dat",
    "Table"][[10]];
PiRelh05p005s095CIB = Import[
    "SLIM_F09/StatsProc_R_2000_h_0.5_self_0.95_f_0.05_10b_SLIM.dat",
    "Table"][[11]];
PiRelh05p005s095CIT = Import[
    "SLIM_F09/StatsProc_R_2000_h_0.5_self_0.95_f_0.05_10b_SLIM.dat",
    "Table"][[12]];
PiRelh05p005s095S = Partition[Riffle[
    Partition[Riffle[Rin, PiRelh05p005s095], 2], Map[ErrorBar,
     Partition[Riffle[-PiRelh05p005s095CIB, PiRelh05p005s095CIT], 2]]], 2];
```

```
Ph05p005s095 = Plot[{PWR2B[5000, 0.05, 0.5, 0.95, R, 0.05],
   ExPiSV[5000, 0.05, 0.5, 0.95, R/10 000, 0.05]}, {R, 0, 2000},
  PlotStyle → {{Black, Thick, Dashed}, {Black, Thick}},
  PlotRange → {All, {0, 1.1}},
  BaseStyle → {FontWeight → "Bold", FontColor → Black, FontSize → 12}]
```

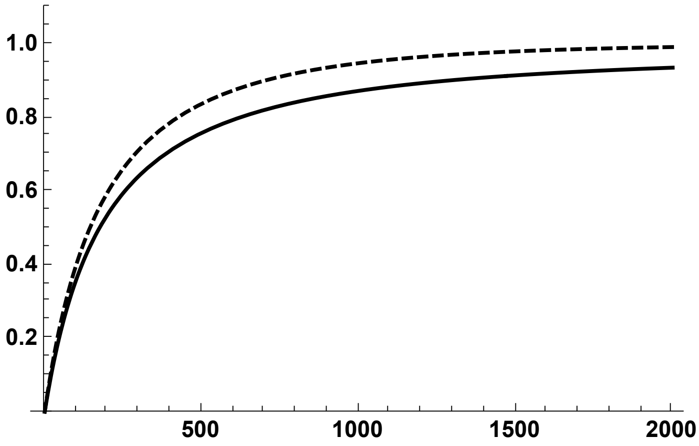

```
Ph05p005s095A = Plot[{ExPiSV[5000, 0.05, 0.5, 0.95, R/10000, 0.05]},
   {R, 0, 2000}, PlotStyle → {{Black, Thick}},
   PlotRange → {All, {0, 1.1}}, Frame → {{True, False}, {True, False}},
   BaseStyle → {FontWeight → "Bold", FontColor → Black, FontSize → 12}];

Ph05p005s095S = ErrorListPlot[PiRelh05p005s095S,
  PlotStyle → {Black, PointSize[0.02]}, PlotRange → {{0, 2000}, {0, 1.1}}]
```

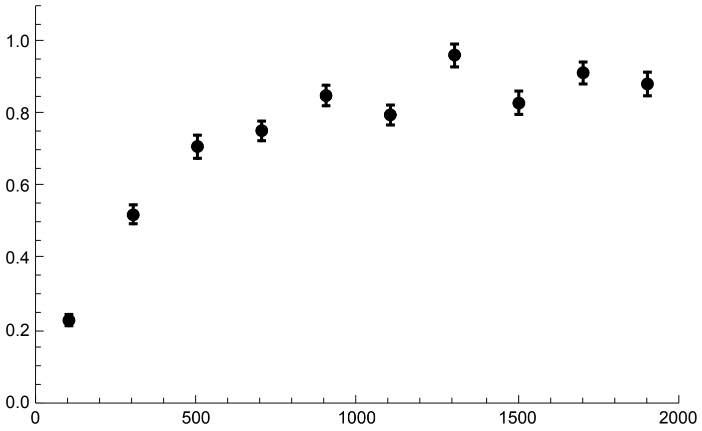


```
Show[Ph05p005s095, Ph05p005s095S]
```

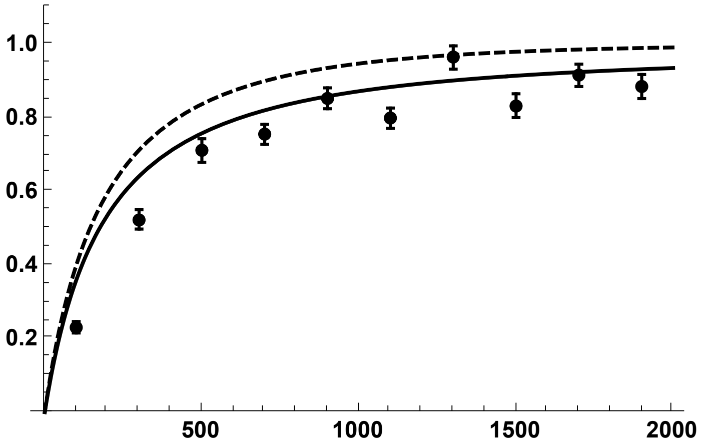


h = 0.1

```
PiRelh01p005s095 =
  Import["SLIM_F09/StatsProc_R_2000_h_0.1_self_0.95_f_0.05_10b_SLIM.dat",
    "Table"][[10]];
PiRelh01p005s095CIB = Import[
    "SLIM_F09/StatsProc_R_2000_h_0.1_self_0.95_f_0.05_10b_SLIM.dat",
    "Table"][[11]];
PiRelh01p005s095CIT = Import[
    "SLIM_F09/StatsProc_R_2000_h_0.1_self_0.95_f_0.05_10b_SLIM.dat",
    "Table"][[12]];
PiRelh01p005s095S = Partition[Riffle[
    Partition[Riffle[Rin, PiRelh01p005s095], 2], Map[ErrorBar,
     Partition[Riffle[-PiRelh01p005s095CIB, PiRelh01p005s095CIT], 2]]], 2];
```

```
Ph01p005s095 = Plot[{PWR2B[5000, 0.05, 0.1, 0.95, R, 0.05],
    ExPiSV[5000, 0.05, 0.1, 0.95, R/10 000, 0.05]},
   {R, 0, 2000}, PlotStyle → {{Red, Thick, Dashed}, {Red, Thick}},
   PlotRange → All, Frame → {{True, False}, {True, False}},
   BaseStyle → {FontWeight → "Bold", FontColor → Black, FontSize → 12}]
```

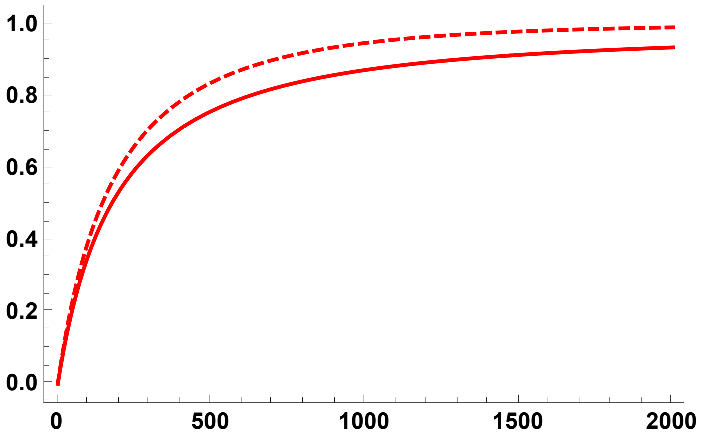


```
Ph01p005s095A = Plot[{ExPiSV[5000, 0.05, 0.1, 0.95, R/10 000, 0.05]},
    {R, 0, 2000}, PlotStyle → {{Red, Thick}},
    PlotRange → All, Frame → {{True, False}, {True, False}},
    BaseStyle → {FontWeight → "Bold", FontColor → Black, FontSize → 12}];

Ph01p005s095S = ErrorListPlot[PiRelh01p005s095S,
   PlotStyle → {Red, PointSize[0.02]}, PlotRange → {{0, 2000}, {0, 1.1}}]
```

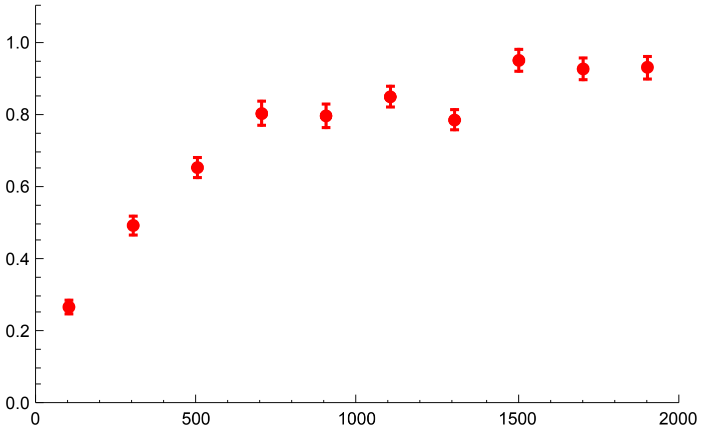

```
Show[Ph01p005s095, Ph01p005s095S]
```

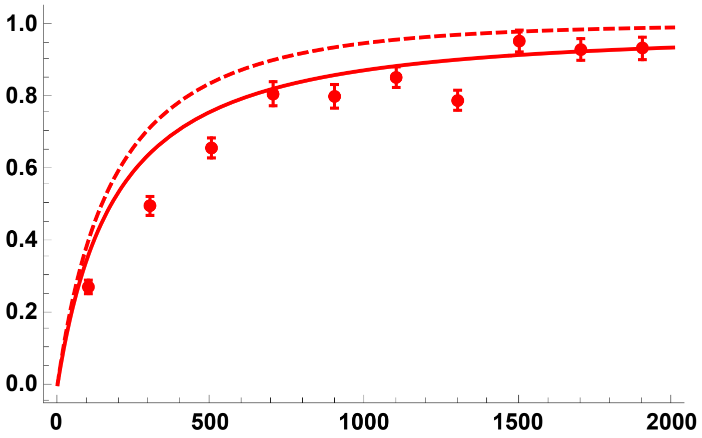


h = 0.9

```
PiRelh09p005s095 =
  Import["SLIM_F09/StatsProc_R_2000_h_0.9_self_0.95_f_0.05_10b_SLIM.dat",
    "Table"][[10]];
PiRelh09p005s095CIB = Import[
    "SLIM_F09/StatsProc_R_2000_h_0.9_self_0.95_f_0.05_10b_SLIM.dat",
    "Table"][[11]];
PiRelh09p005s095CIT = Import[
    "SLIM_F09/StatsProc_R_2000_h_0.9_self_0.95_f_0.05_10b_SLIM.dat",
    "Table"][[12]];
PiRelh09p005s095S = Partition[Riffle[
    Partition[Riffle[Rin, PiRelh09p005s095], 2], Map[ErrorBar,
     Partition[Riffle[-PiRelh09p005s095CIB, PiRelh09p005s095CIT], 2]]], 2];
```

```
Ph09p005s095 = Plot[{PWR2B[5000, 0.05, 0.9, 0.95, R, 0.05],
   ExPiSV[5000, 0.05, 0.9, 0.95, R/10 000, 0.05]},
  {R, 0, 2000}, PlotStyle → { {Blue, Thick, Dashed}, {Blue, Thick}},
  PlotRange → All, Frame → {{True, False}, {True, False}},
  BaseStyle → {FontWeight → "Bold", FontColor → Black, FontSize → 12}]
```

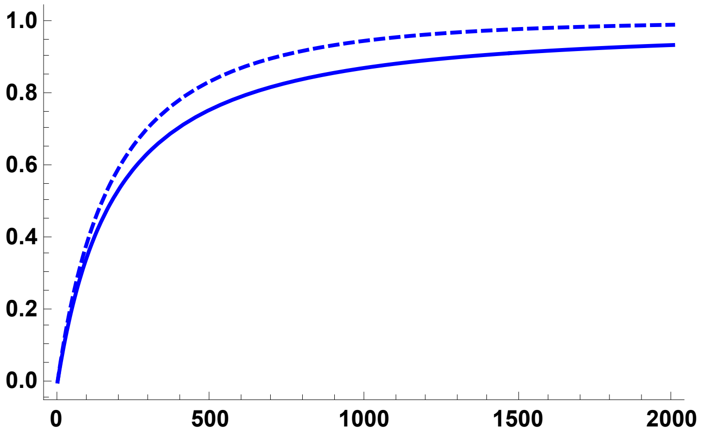

```
Ph09p005s095A = Plot[{ExPiSV[5000, 0.05, 0.9, 0.95, R/10 000, 0.05]},
    {R, 0, 2000}, PlotStyle → {{Blue, Thick}},
    PlotRange → All, Frame → {{True, False}, {True, False}},
    BaseStyle → {FontWeight → "Bold", FontColor → Black, FontSize → 12}];
```

```
Ph09p005s095S = ErrorListPlot[PiRelh09p005s095S,
   PlotStyle → {Blue, PointSize[0.02]}, PlotRange → {{0, 2000}, {0, 1.1}}]
```

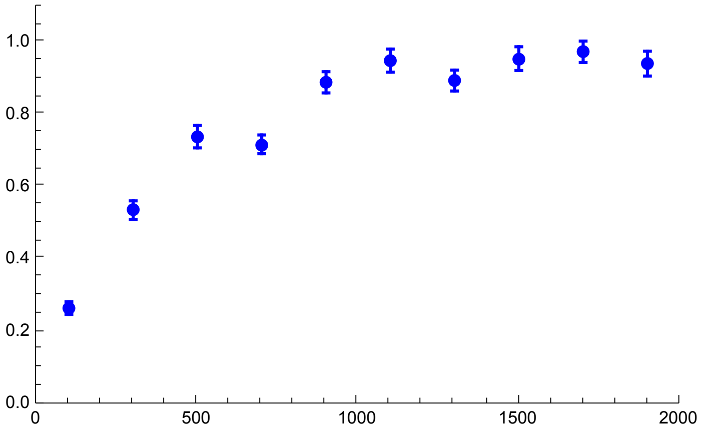


```
Show[Ph09p005s095, Ph09p005s095S]
```

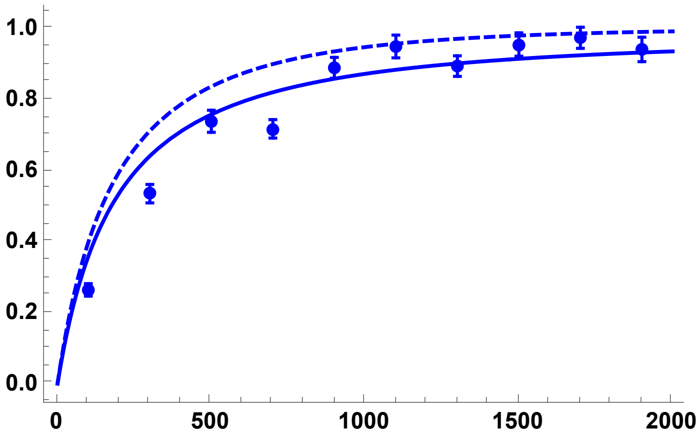


All h values, using the coalescence-in-sweep-phase results and simulations:

```
Sims0955pc = Show[Ph01p005s095A, Ph01p005s095S, Ph05p005s095A,
   Ph05p005s095S, Ph09p005s095A, Ph09p005s095S, ImageSize → 250]
```

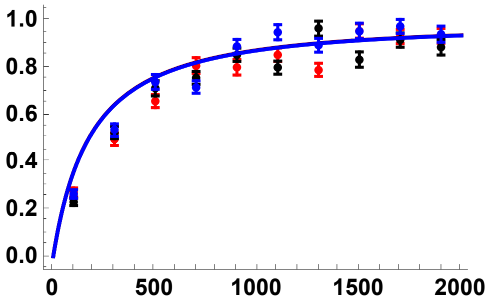


All h values, using the coalescence-in-sweep-phase results and simulations:

```
Sims0955pc2 = Show[Ph01p005s095, Ph05p005s095, Ph09p005s095, ImageSize → 250]
```

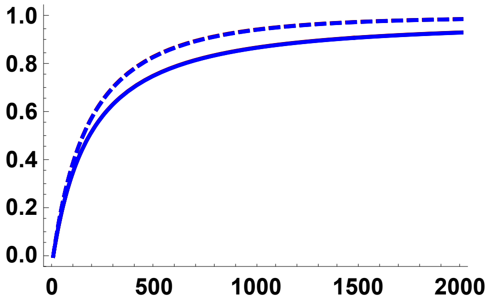


#### Plots of sweep trajectories

Function to numerically evaluate trajectories

```
PSolve[Na_, σ_, s_, h_, tmax_, pin_] :=
 NDSolve[{p'[t] == (1 - p[t]) p[t] s ((F[σ] + h - F[σ] h + (1 - F[σ]) (1 - 2 h) p[t])),
   p[0] == pin}, p[t], {t, 0, tmax}]
```

```
Boostp0[Na_, s_, h_, σ_] := (1 + F[σ]) / (4 Na s (F[σ] + h - F[σ] h))
```

In the following plots, x-axis is the time (number of generations) and y-axis is the selected allele frequency.

##### $\sigma = 0, f_0 = 1/2N$

```
p1h01 = Plot[
  Evaluate[p[t] /. PSolve[5000, 0, 0.05, 0.1, 800, Boostp0[5000, 0.05, 0.1, 0]]],
  {t, 0, 800}, PlotStyle → {Red, Thick}, PlotRange → Full, AxesOrigin → {0, 0},
  BaseStyle → {FontWeight → "Bold", FontColor → Black, FontSize → 12}]
```

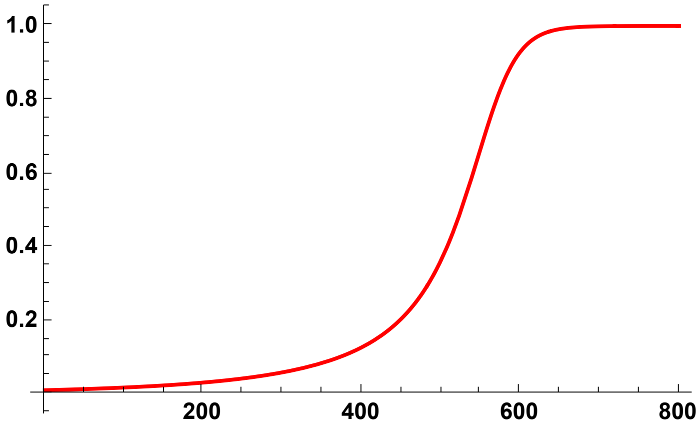

```
p1h05 = Plot[
  Evaluate[p[t] /. PSolve[5000, 0, 0.05, 0.5, 800, Boostp0[5000, 0.05, 0.5, 0]]],
  {t, 0, 800}, PlotStyle → {Black, Thick}, AxesOrigin → {0, 0},
  BaseStyle → {FontWeight → "Bold", FontColor → Black, FontSize → 12}]
```

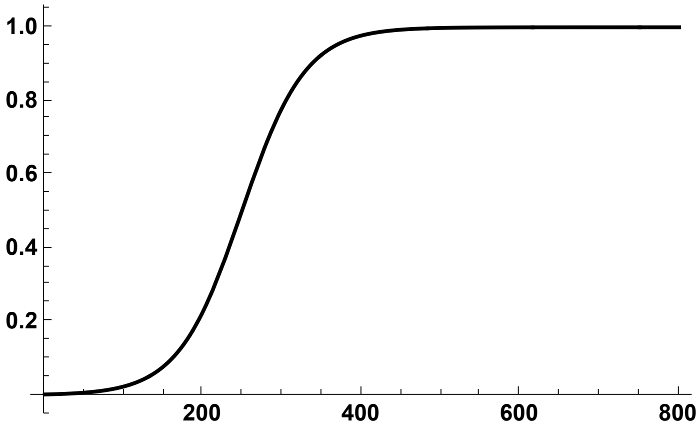


```
p1h09 = Plot[
  Evaluate[p[t] /. PSolve[5000, 0, 0.05, 0.9, 800, Boostp0[5000, 0.05, 0.9, 0]]],
  {t, 0, 800}, PlotStyle → {Blue, Thick}, AxesOrigin → {0, 0},
  BaseStyle → {FontWeight → "Bold", FontColor → Black, FontSize → 12}]
```

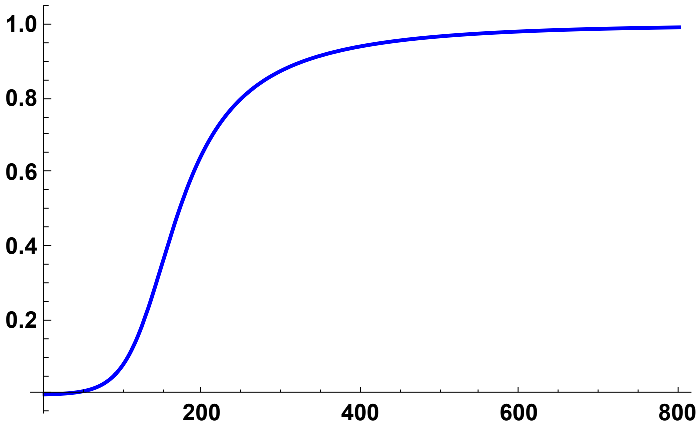


Composite plot

```
Trajs0DN = Show[p1h01, p1h05, p1h09, ImageSize → 250]
```

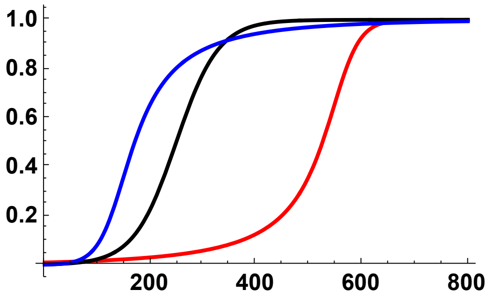

#### $\sigma = 0, f_0 = 0.02$

```
p1h01p02 = Plot[Evaluate[p[t] /. PSolve[5000, 0, 0.05, 0.1, 600, 0.02]],
  {t, 0, 600}, PlotStyle → {Red, Thick}, PlotRange → Full, AxesOrigin → {0, 0},
  BaseStyle → {FontWeight → "Bold", FontColor → Black, FontSize → 12}]
```

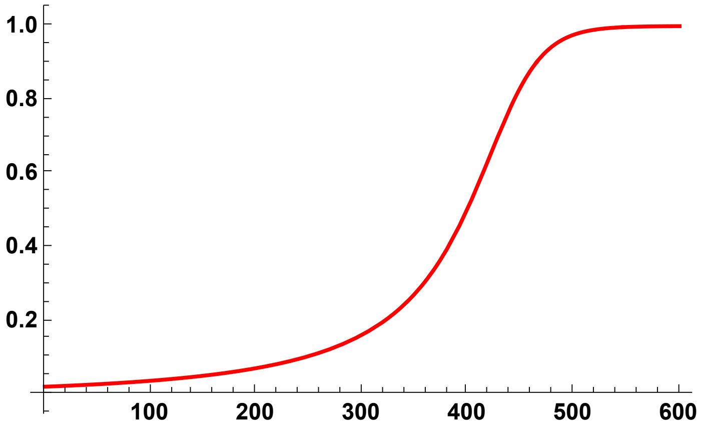


```
p1h05p02 = Plot[Evaluate[p[t] /. PSolve[5000, 0, 0.05, 0.5, 600, 0.02]],
  {t, 0, 600}, PlotStyle → {Black, Thick}, AxesOrigin → {0, 0},
  BaseStyle → {FontWeight → "Bold", FontColor → Black, FontSize → 12}]
```

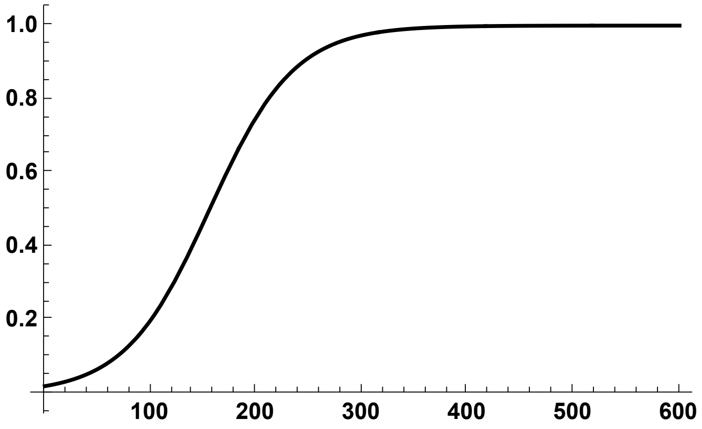


```
p1h09p02 = Plot[Evaluate[p[t] /. PSolve[5000, 0, 0.05, 0.9, 600, 0.02]],
  {t, 0, 600}, PlotStyle → {Blue, Thick}, AxesOrigin → {0, 0},
  BaseStyle → {FontWeight → "Bold", FontColor → Black, FontSize → 12}]
```

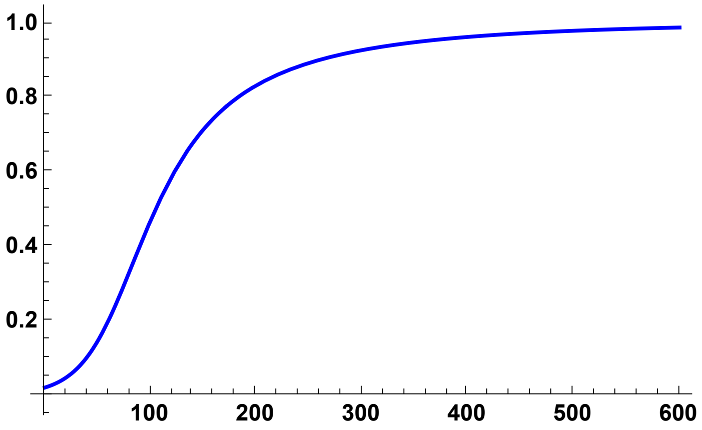

Composite plot

```
Trajs02pc = Show[p1h01p02, p1h05p02, p1h09p02, ImageSize → 250]
```

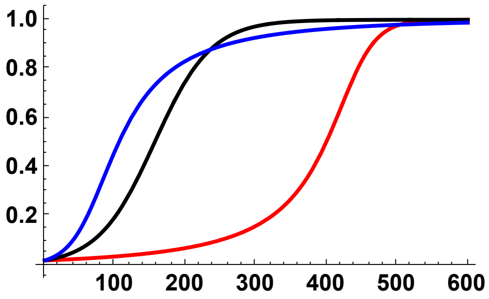


#### $\sigma = 0, f_0 = 0.05$

```
p1h01p05 = Plot[Evaluate[p[t] /. PSolve[5000, 0, 0.05, 0.1, 500, 0.05]],
  {t, 0, 500}, PlotStyle → {Red, Thick}, PlotRange → Full, AxesOrigin → {0, 0},
  BaseStyle → {FontWeight → "Bold", FontColor → Black, FontSize → 12}]
```

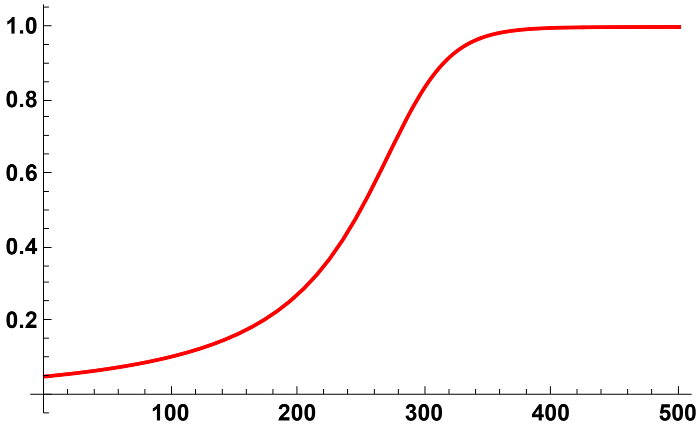


```
p1h05p05 = Plot[Evaluate[p[t] /. PSolve[5000, 0, 0.05, 0.5, 500, 0.05]],
  {t, 0, 500}, PlotStyle → {Black, Thick}, AxesOrigin → {0, 0},
  BaseStyle → {FontWeight → "Bold", FontColor → Black, FontSize → 12}]
```

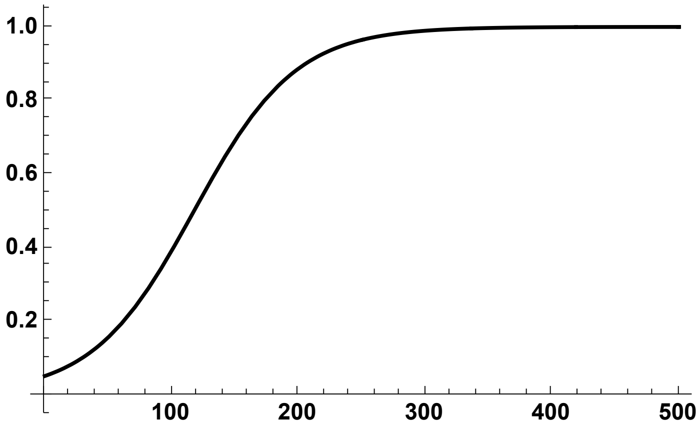

```
p1h09p05 = Plot[Evaluate[p[t] /. PSolve[5000, 0, 0.05, 0.9, 500, 0.05]],
  {t, 0, 500}, PlotStyle → {Blue, Thick}, AxesOrigin → {0, 0},
  BaseStyle → {FontWeight → "Bold", FontColor → Black, FontSize → 12}]
```

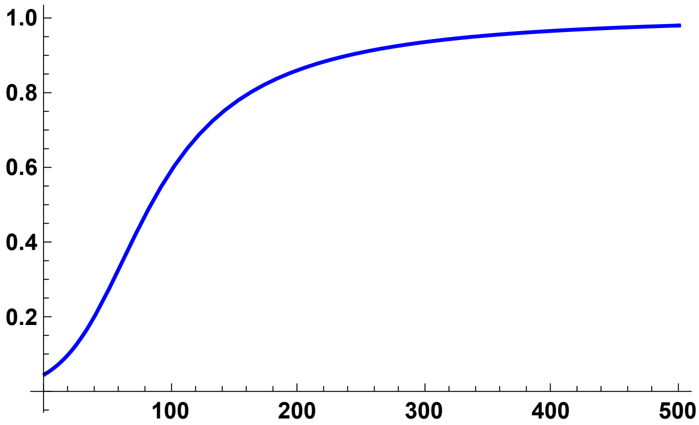


Composite plot

```
Trajs05pc = Show[p1h01p05, p1h05p05, p1h09p05, ImageSize → 250]
```

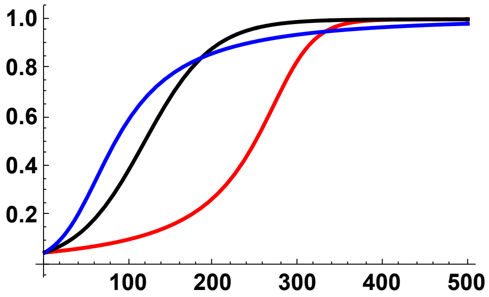


#### $\sigma = 0.50, f_0 = 1/2\,N$

```
p2h01 = Plot[Evaluate[
    p[t] /. PSolve[5000, 0.5, 0.05, 0.1, 400, Boostp0[5000, 0.05, 0.1, 0.5]]],
  {t, 0, 400}, PlotStyle → {Red, Thick}, PlotRange → Full, AxesOrigin → {0, 0},
  BaseStyle → {FontWeight → "Bold", FontColor → Black, FontSize → 12}]
```

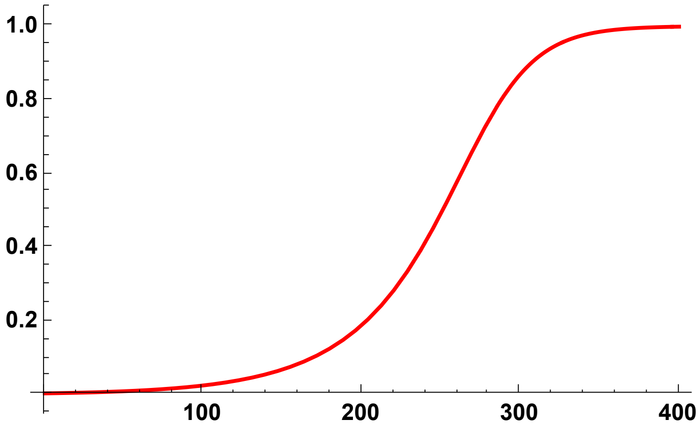

```
p2h05 = Plot[Evaluate[
   p[t] /. PSolve[5000, 0.5, 0.05, 0.5, 400, Boostp0[5000, 0.05, 0.5, 0.5]]],
  {t, 0, 400}, PlotStyle → {Black, Thick}, AxesOrigin → {0, 0},
  BaseStyle → {FontWeight → "Bold", FontColor → Black, FontSize → 12}]
```

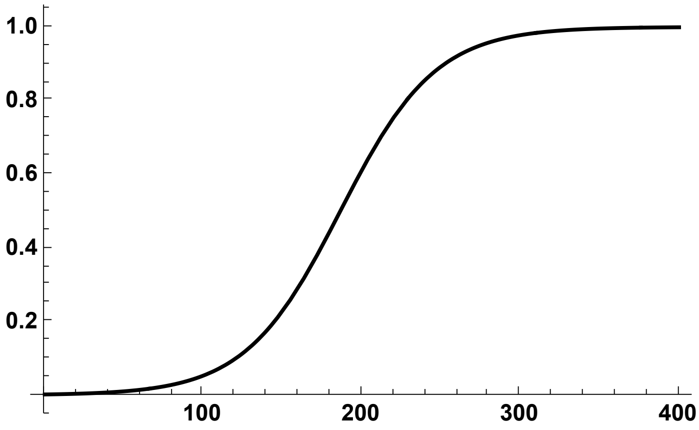


```
p2h09 = Plot[Evaluate[
   p[t] /. PSolve[5000, 0.5, 0.05, 0.9, 400, Boostp0[5000, 0.05, 0.9, 0.5]]],
  {t, 0, 400}, PlotStyle → {Blue, Thick}, AxesOrigin → {0, 0},
  BaseStyle → {FontWeight → "Bold", FontColor → Black, FontSize → 12}]
```

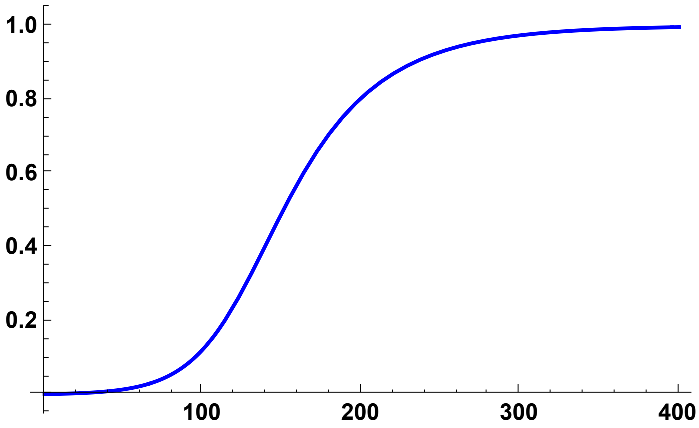


Composite plot

```
Trajs05DN = Show[p2h01, p2h05, p2h09, ImageSize → 250]
```

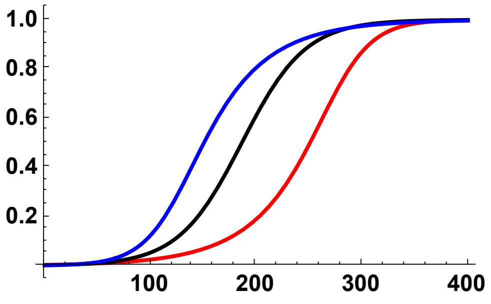

#### $\sigma = 0.50, f_0 = 0.02$

```
p2h01p02 = Plot[Evaluate[p[t] /. PSolve[5000, 0.5, 0.05, 0.1, 400, 0.02]],
  {t, 0, 400}, PlotStyle → {Red, Thick}, PlotRange → Full, AxesOrigin → {0, 0},
  BaseStyle → {FontWeight → "Bold", FontColor → Black, FontSize → 12}]
```

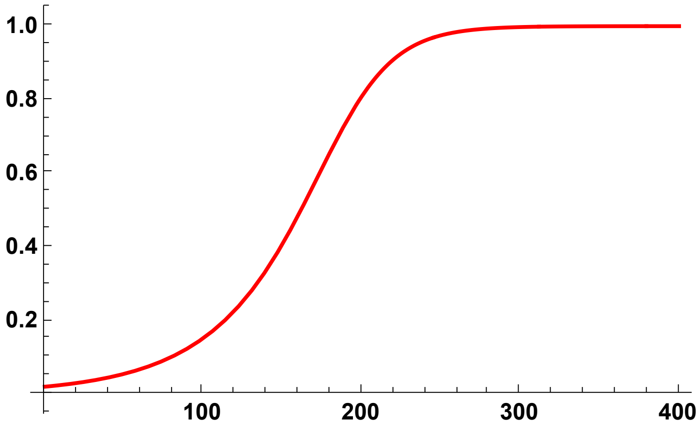


```
p2h05p02 = Plot[Evaluate[p[t] /. PSolve[5000, 0.5, 0.05, 0.5, 400, 0.02]],
  {t, 0, 400}, PlotStyle → {Black, Thick}, AxesOrigin → {0, 0},
  BaseStyle → {FontWeight → "Bold", FontColor → Black, FontSize → 12}]
```

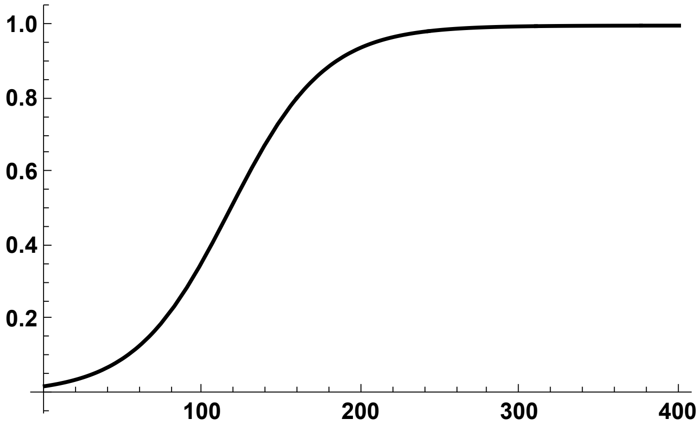


```
p2h09p02 = Plot[Evaluate[p[t] /. PSolve[5000, 0.5, 0.05, 0.9, 400, 0.02]],
  {t, 0, 400}, PlotStyle → {Blue, Thick}, AxesOrigin → {0, 0},
  BaseStyle → {FontWeight → "Bold", FontColor → Black, FontSize → 12}]
```

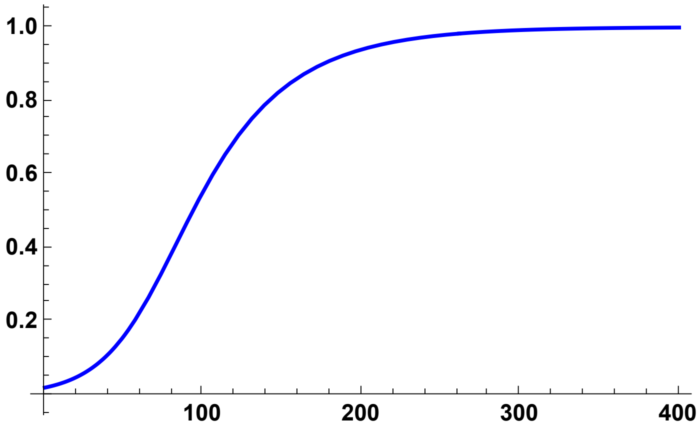

Composite plot

```
Trajs052pc = Show[p2h01p02, p2h05p02, p2h09p02, ImageSize → 250]
```

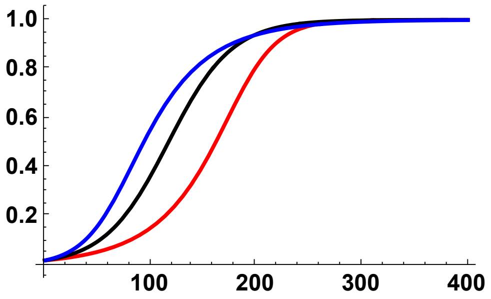


#### $\sigma = 0.50, f_0 = 0.05$

```
p2h01p05 = Plot[Evaluate[p[t] /. PSolve[5000, 0.5, 0.05, 0.1, 400, 0.05]],
  {t, 0, 300}, PlotStyle → {Red, Thick}, PlotRange → Full, AxesOrigin → {0, 0},
  BaseStyle → {FontWeight → "Bold", FontColor → Black, FontSize → 12}]
```

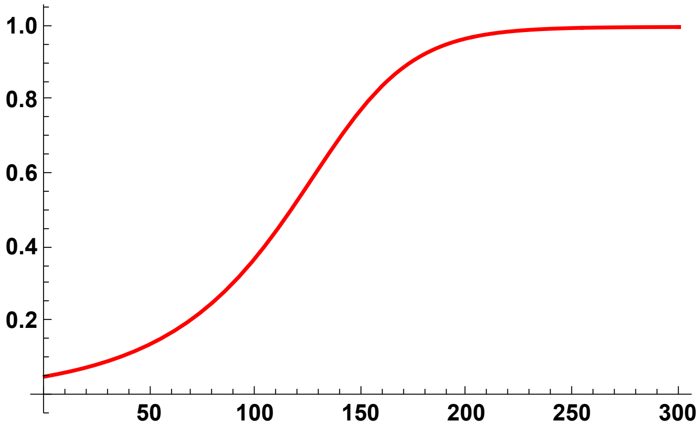


```
p2h05p05 = Plot[Evaluate[p[t] /. PSolve[5000, 0.5, 0.05, 0.5, 400, 0.05]],
  {t, 0, 300}, PlotStyle → {Black, Thick}, AxesOrigin → {0, 0},
  BaseStyle → {FontWeight → "Bold", FontColor → Black, FontSize → 12}]
```

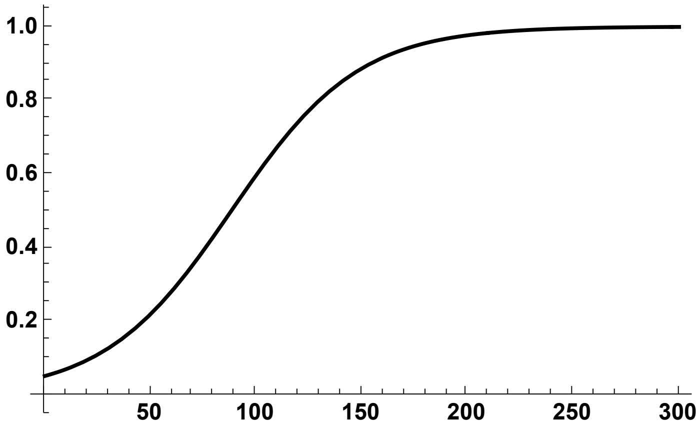

```
p2h09p05 = Plot[Evaluate[p[t] /. PSolve[5000, 0.5, 0.05, 0.9, 400, 0.05]],
  {t, 0, 300}, PlotStyle → {Blue, Thick}, AxesOrigin → {0, 0},
  BaseStyle → {FontWeight → "Bold", FontColor → Black, FontSize → 12}]
```

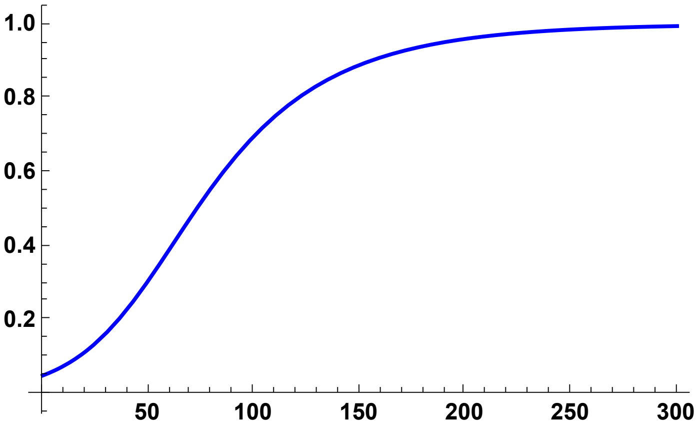


Composite plot

```
Trajs055pc = Show[p2h01p05, p2h05p05, p2h09p05, ImageSize → 250]
```

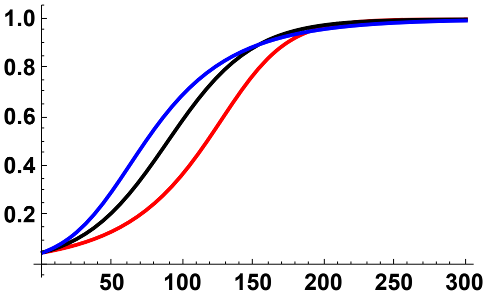


#### $\sigma = 0.95, f_0 = 1/2\,N$

```
p3h01 = Plot[Evaluate[
    p[t] /. PSolve[5000, 0.95, 0.05, 0.1, 300, Boostp0[5000, 0.05, 0.1, 0.95]]],
  {t, 0, 300}, PlotStyle → {Red, Thick}, PlotRange → Full, AxesOrigin → {0, 0},
  BaseStyle → {FontWeight → "Bold", FontColor → Black, FontSize → 12}]
```

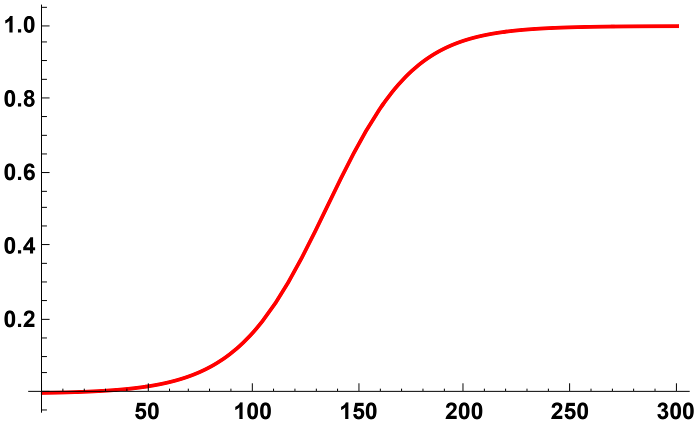

```
p3h05 = Plot[Evaluate[
   p[t] /. PSolve[5000, 0.95, 0.05, 0.5, 300, Boostp0[5000, 0.05, 0.5, 0.95]]],
  {t, 0, 300}, PlotStyle → {Black, Thick}, AxesOrigin → {0, 0},
  BaseStyle → {FontWeight → "Bold", FontColor → Black, FontSize → 12}]
```

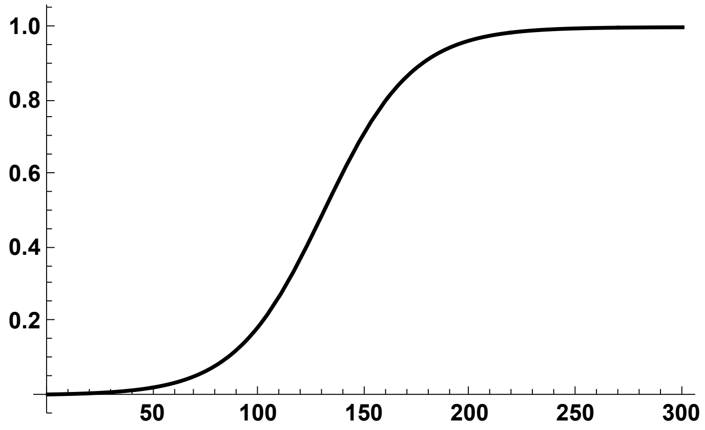


```
p3h09 = Plot[Evaluate[
   p[t] /. PSolve[5000, 0.95, 0.05, 0.9, 300, Boostp0[5000, 0.05, 0.9, 0.95]]],
  {t, 0, 300}, PlotStyle → {Blue, Thick}, AxesOrigin → {0, 0},
  BaseStyle → {FontWeight → "Bold", FontColor → Black, FontSize → 12}]
```

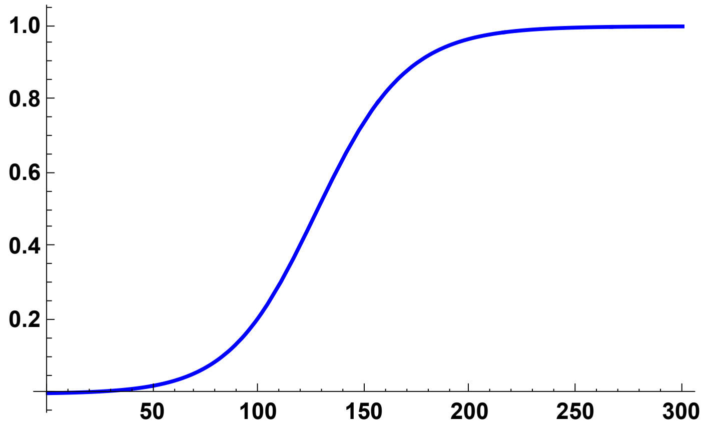


Composite plot

```
Trajs095DN = Show[p3h01, p3h05, p3h09, ImageSize → 250]
```

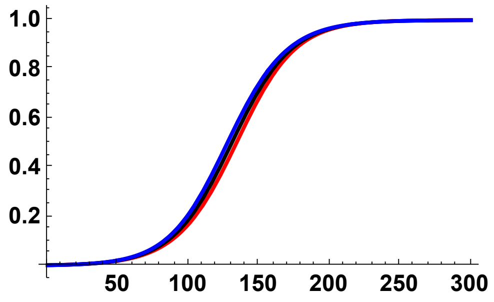

#### $\sigma = 0.95, f_0 = 0.02$

```
p3h01p02 = Plot[Evaluate[p[t] /. PSolve[5000, 0.95, 0.05, 0.1, 400, 0.02]],
  {t, 0, 200}, PlotStyle → {Red, Thick}, PlotRange → Full, AxesOrigin → {0, 0},
  BaseStyle → {FontWeight → "Bold", FontColor → Black, FontSize → 12}]
```

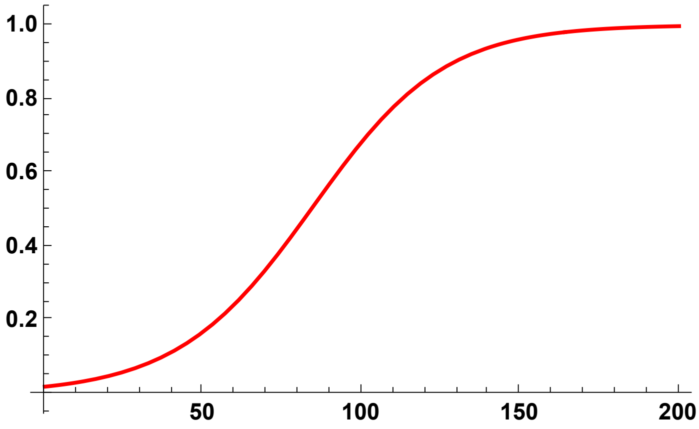


```
p3h05p02 = Plot[Evaluate[p[t] /. PSolve[5000, 0.95, 0.05, 0.5, 400, 0.02]],
  {t, 0, 200}, PlotStyle → {Black, Thick}, AxesOrigin → {0, 0},
  BaseStyle → {FontWeight → "Bold", FontColor → Black, FontSize → 12}]
```

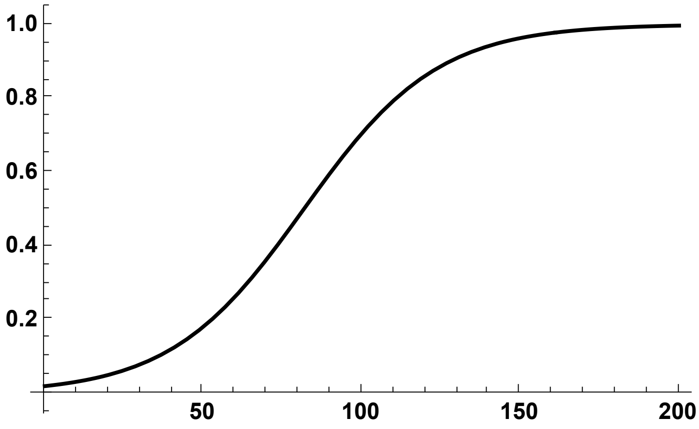


```
p3h09p02 = Plot[Evaluate[p[t] /. PSolve[5000, 0.95, 0.05, 0.9, 400, 0.02]],
  {t, 0, 200}, PlotStyle → {Blue, Thick}, AxesOrigin → {0, 0},
  BaseStyle → {FontWeight → "Bold", FontColor → Black, FontSize → 12}]
```

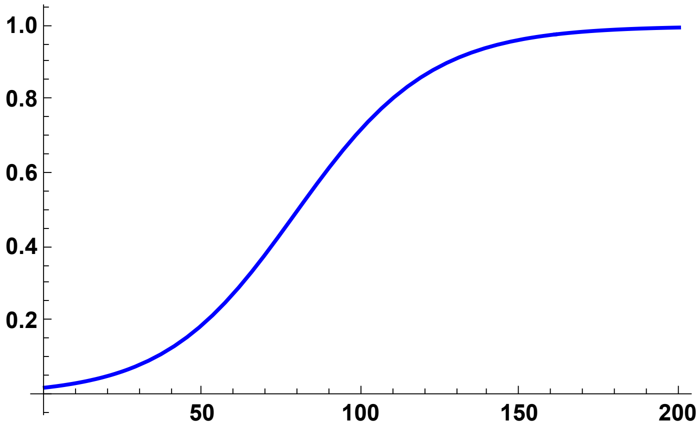

Composite plot

```
Trajs0952pc = Show[p3h01p02, p3h05p02, p3h09p02, ImageSize → 250]
```

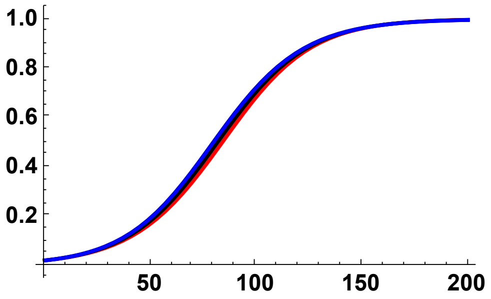


#### $\sigma = 0.95, f_0 = 0.05$

```
p3h01p05 = Plot[Evaluate[p[t] /. PSolve[5000, 0.95, 0.05, 0.1, 400, 0.05]],
  {t, 0, 200}, PlotStyle → {Red, Thick}, PlotRange → Full, AxesOrigin → {0, 0},
  BaseStyle → {FontWeight → "Bold", FontColor → Black, FontSize → 12}]
```

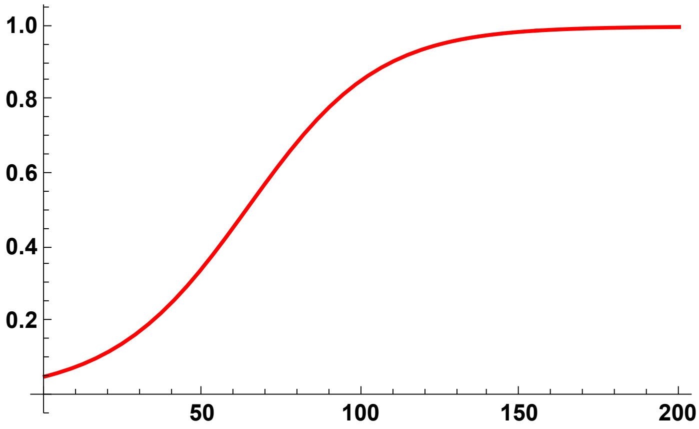


```
p3h05p05 = Plot[Evaluate[p[t] /. PSolve[5000, 0.95, 0.05, 0.5, 400, 0.05]],
  {t, 0, 200}, PlotStyle → {Black, Thick}, AxesOrigin → {0, 0},
  BaseStyle → {FontWeight → "Bold", FontColor → Black, FontSize → 12}]
```

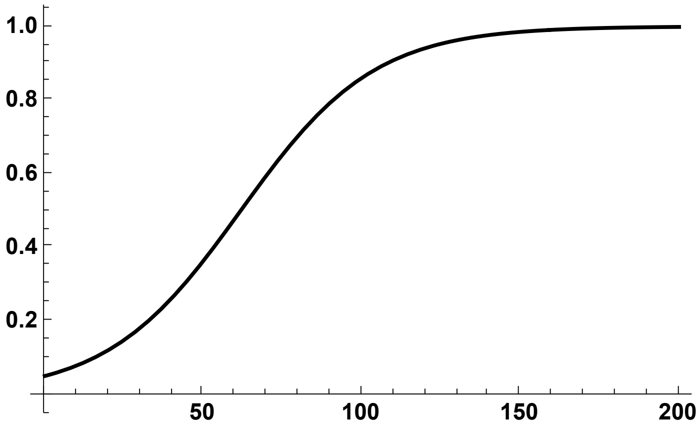

```
p3h09p05 = Plot[Evaluate[p[t] /. PSolve[5000, 0.95, 0.05, 0.9, 400, 0.05]],
  {t, 0, 200}, PlotStyle → {Blue, Thick}, AxesOrigin → {0, 0},
  BaseStyle → {FontWeight → "Bold", FontColor → Black, FontSize → 12}]
```

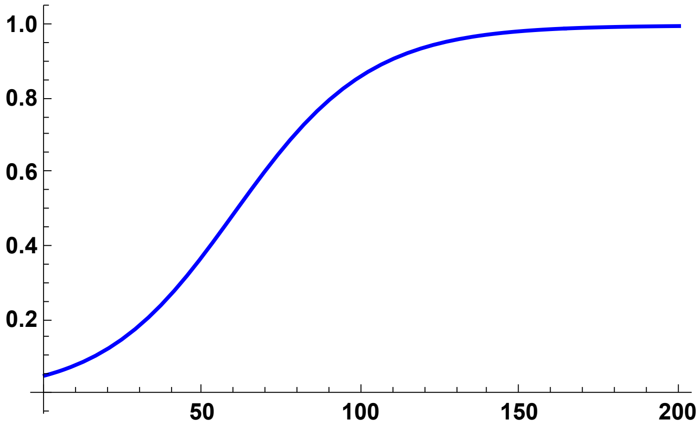


Composite plot

```
Trajs0955pc = Show[p3h01p05, p3h05p05, p3h09p05, ImageSize → 250]
```

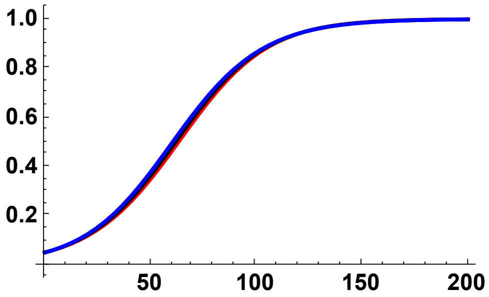

### Grid of key results

#### Comparing coalescence-in-sweep-phase results and SLiM simulations

```
SimComp = Labeled[Grid[{{Text@TraditionalForm@Style["p0 = 1/2N", 24],
     Text@TraditionalForm@Style["p0 = 0.02", 24],
     Text@TraditionalForm@Style["p0 = 0.05", 24],}, {Sims0DN, Sims02pc,
     Sims05pc, Text@TraditionalForm@Style["σ = 0.00\n(F = 0.00)", 24]},
    {Sims05DN, Sims052pc, Sims055pc,
     Text@TraditionalForm@Style["σ = 0.50\n(F ≃ 0.33)", 24]},
    {Sims095DN, Sims0952pc, Sims0955pc, Text@
      TraditionalForm@Style["σ = 0.95\n(F ≃ 0.90)", 24]}}, Spacings → {2, 1}],
  {Text@TraditionalForm@Style["Expected\nRelative\nDiversity,\n𝔼[πSV/π0]", 24],
   Text@TraditionalForm@Style["Scaled Recombination Rate, 2Nr", 24]}, {Left,
   Bottom}]
```

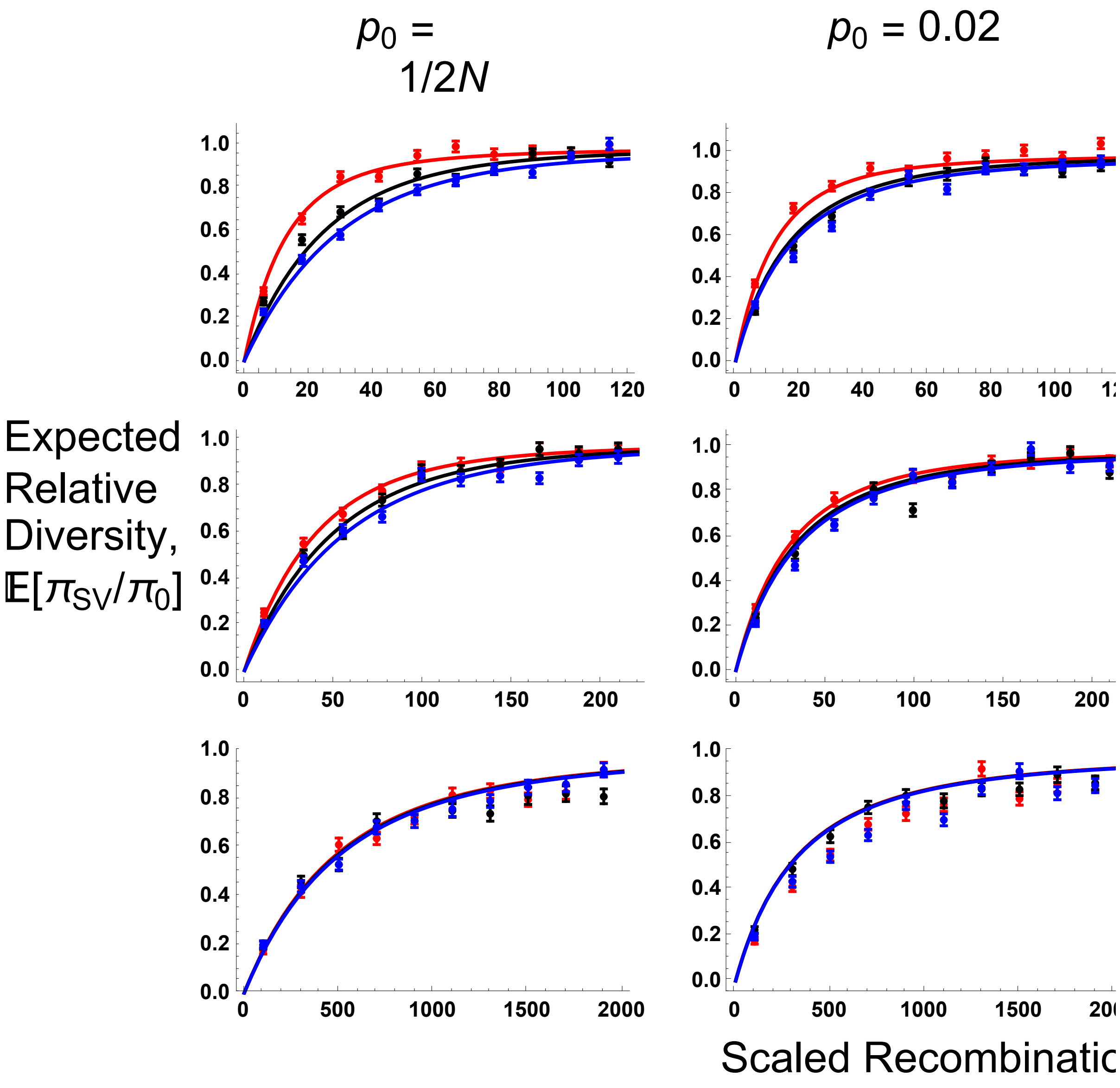

#### Comparing coalescence-in-sweep-phase results and star-like approximation

```
AnalyticComp = Labeled[Grid[{{Text@TraditionalForm@Style["p₀ = 1/2N", 24],
     Text@TraditionalForm@Style["p₀ = 0.02", 24],
     Text@TraditionalForm@Style["p₀ = 0.05", 24],}, {Sims0DN2, Sims02pc2,
     Sims05pc2, Text@TraditionalForm@Style["σ = 0.00\n(F = 0.00)", 24]},
    {Sims05DN2, Sims052pc2, Sims055pc2,
     Text@TraditionalForm@Style["σ = 0.50\n(F ≃ 0.33)", 24]},
    {Sims095DN2, Sims0952pc2, Sims0955pc2, Text@
      TraditionalForm@Style["σ = 0.95\n(F ≃ 0.90)", 24]}}, Spacings → {2, 1}],
  {Text@TraditionalForm@Style["Expected\nRelative\nDiversity,\n𝔼[πSV/π0]", 24],
   Text@TraditionalForm@Style["Scaled Recombination Rate, 2Nr", 24]}, {Left,
   Bottom}]
```

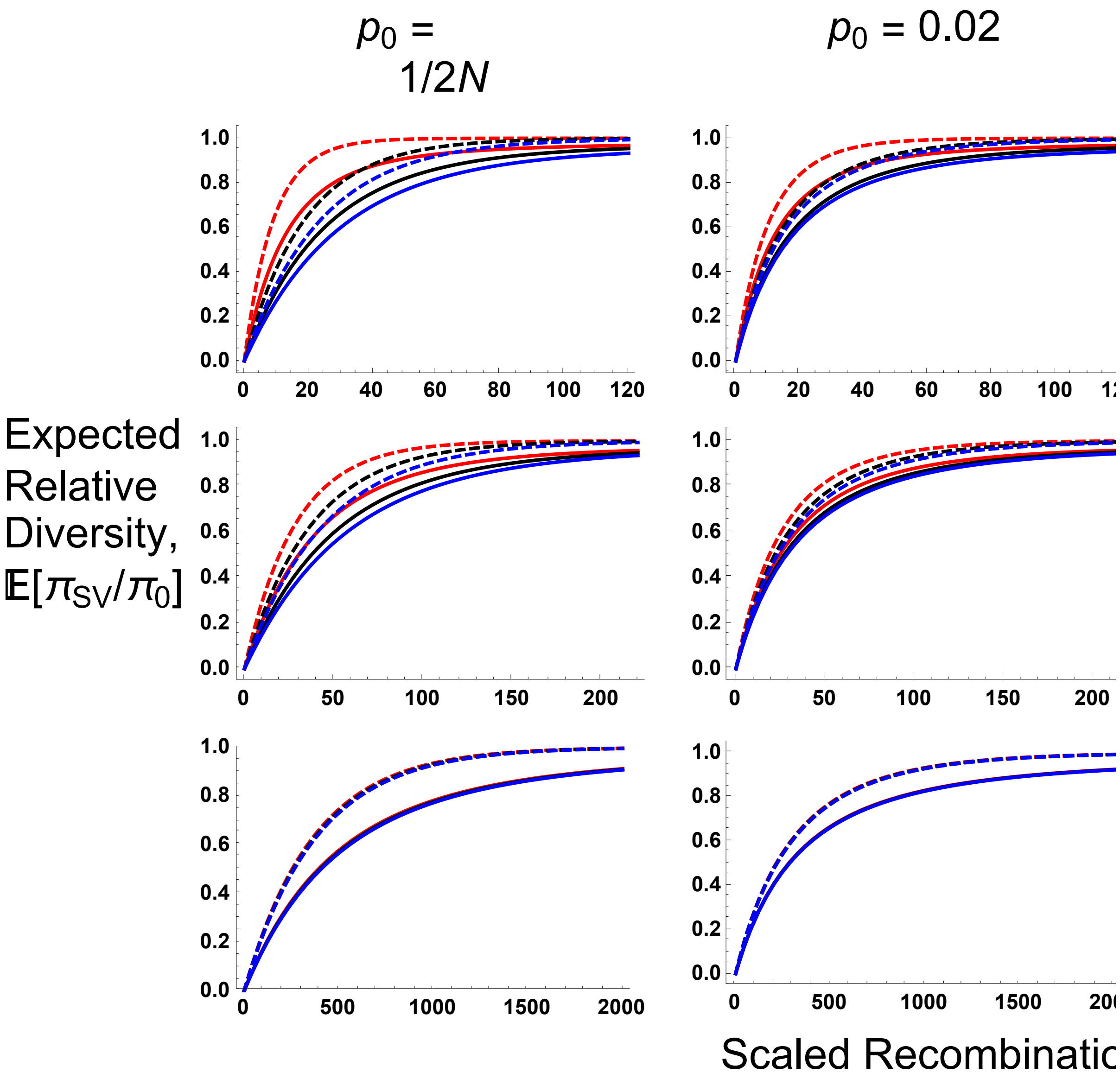

##### Plot of sweep trajectories

```
SweepTrajs =
 Labeled[Grid[{{Text@TraditionalForm@Style["σ = 0.00\n(F = 0.00)", 24],
     Text@TraditionalForm@Style["σ = 0.50\n(F ≃ 0.33)", 24],
     Text@TraditionalForm@Style["σ = 0.95\n(F ≃ 0.90)", 24],}, {Trajs0DN,
     Trajs05DN, Trajs095DN, Text@TraditionalForm@Style["p0 = 1/2N", 24]},
    {Trajs05pc, Trajs055pc, Trajs0955pc,
     Text@TraditionalForm@Style["p0 = 0.05", 24]}}, Spacings → {2, 1}],
  {Text@TraditionalForm@Style["Beneficial\nAllele\nFrequency", 24], Text@
    TraditionalForm@Style["Time (number of generations)", 24]}, {Left, Bottom}]
```

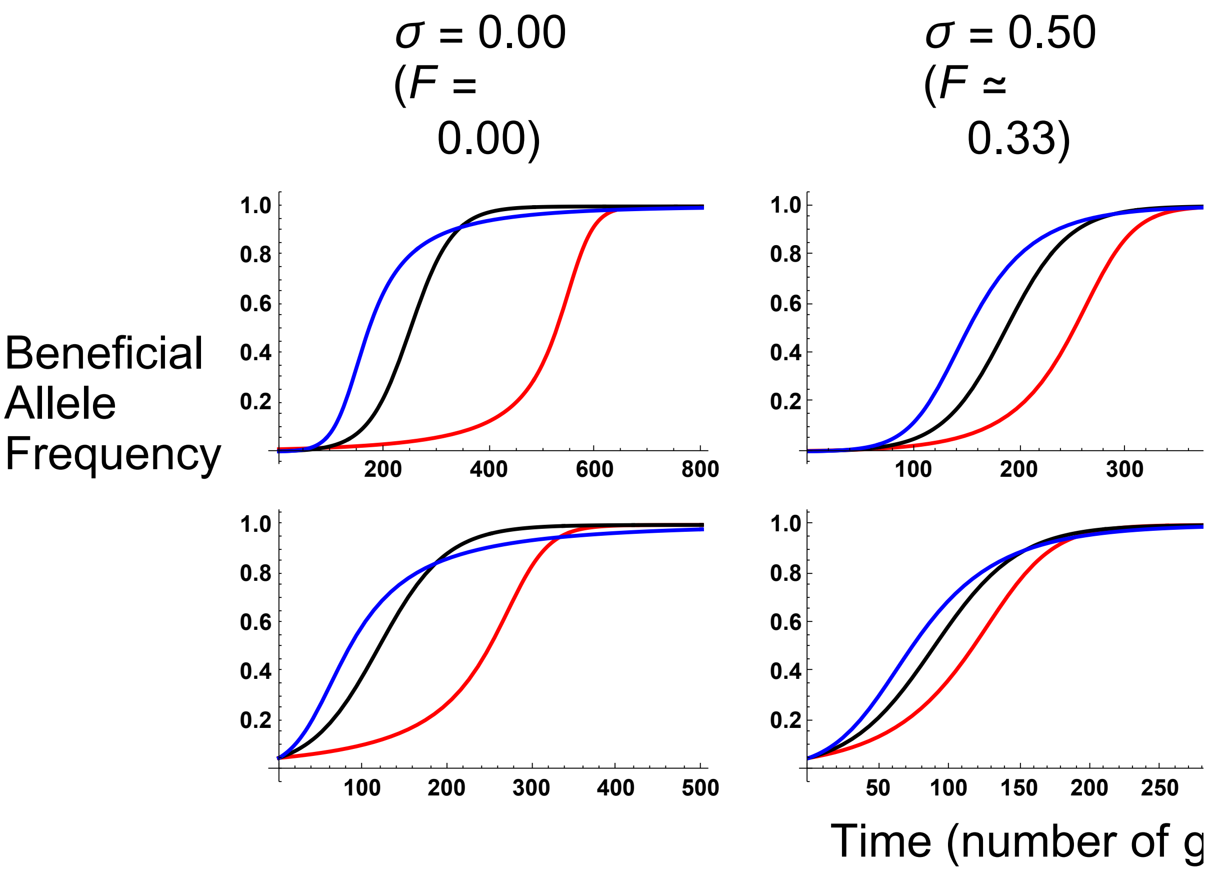


### Section D: Simulation comparisons, $\mathbb{E}[S]$ (Number of segregating sites)

##### Equations

$$F[\sigma\_] := \frac{\sigma}{2 - \sigma}$$

$$\Phi[r\_, \sigma\_] := \frac{\sigma\,(2 - \sigma - 2\,(1 - r)\,r\,(2 - 3\,\sigma))}{(2 - \sigma)\,(2 - (1 - 2\,(1 - r)\,r)\,\sigma)}$$

```
PNR[Na_, F_, Φ_, s_, h_, R_, p0_] := (((F + h - F h)/(1 - h + F h)) (1/p0 + 1) - 1)^(-R (1-F)/(2 Na (F+h-F h) s))

PNRI[Na_, F_, Φ_, s_, h_, R_, p0_, i_, n_] :=
 PDF[BinomialDistribution[n, PNR[Na, F, Φ, s, h, R, p0]], i]

PESF[k_, F_, Φ_, R_, i_, p0_] :=
 (((2 R)/(1 + F) (1 - 2 F + Φ) p0 (1 - p0))^k Abs[StirlingS1[i, k]]) /
  Product[(((2 R)/(1 + F) (1 - 2 F + Φ) p0 (1 - p0)) + l), {l, 0, i - 1}]

JS[k_] := Sum[1/j, {j, 1, k - 1}]
```

𝔼[s] is given as:

```
ESS2[Na_, σ_, s_, h_, R_, θ_, p0_, n_] :=
 (Sum[(PNRI[Na, F[σ], Φ[R/(2 Na), σ], s, h, R, p0, i, n] * Sum[
      PESF[k, F[σ], Φ[R/(2 Na), σ], R, i, p0] * JS[k + n - i], {k, 0, i}]), {i, 0, n}]) θ
```

The following is the ‘effective’ starting frequency of a beneficial allele at initial frequency $\frac{1}{2\,Na}$, given that it goes to fixation:

```
Boostp0[Na_, s_, h_, σ_] := (1 + F[σ])/(4 Na s (F[σ] + h - F[σ] h))
```

#### Outcrossing case ($\sigma = F = 0$)

Preliminaries

```
SetDirectory[NotebookDirectory[]];
Rin = Table[6 + 12 * i, {i, 0, 9}];
```

##### Simulation comparisons, from initial frequency $p_0 = 1/2\,N$

h = 0.5

SLiM simulation data

```
SSh05 = Import["SLIM_F0/StatsProc_R_120_h_0.5_self_0_f_1e-04_10b_SLIM.dat",
    "Table"][[1]];
SSh05CIB = Import["SLIM_F0/StatsProc_R_120_h_0.5_self_0_f_1e-04_10b_SLIM.dat",
    "Table"][[2]];
SSh05CIT = Import["SLIM_F0/StatsProc_R_120_h_0.5_self_0_f_1e-04_10b_SLIM.dat",
    "Table"][[3]];
SSh05S = Partition[Riffle[Partition[Riffle[Rin, SSh05], 2],
    Map[ErrorBar, Partition[Riffle[-SSh05CIB, SSh05CIT], 2]]], 2];
```

MSMS simulation data

```
SSh05MSMS =
  Import["MSMS_SV_5K/StatsProc_R_240_h_0.5_self_0_f_1e-04_10b_MSMS.dat",
    "Table"][[1]];
SSh05MSMSCIB = Import[
    "MSMS_SV_5K/StatsProc_R_240_h_0.5_self_0_f_1e-04_10b_MSMS.dat",
    "Table"][[2]];
SSh05MSMSCIT = Import[
    "MSMS_SV_5K/StatsProc_R_240_h_0.5_self_0_f_1e-04_10b_MSMS.dat",
    "Table"][[3]];
SSh05MSMSS = Partition[Riffle[Partition[Riffle[Rin, SSh05MSMS], 2],
    Map[ErrorBar, Partition[Riffle[-SSh05MSMSCIB, SSh05MSMSCIT], 2]]], 2];
```

Analytical result

```
Sh05 = Plot[ESS2[5000, 0, 0.05, 0.5, R, 4, Boostp0[5000, 0.05, 0.5, 0], 10],
  {R, 0, 120}, PlotStyle → {{Black, Thick}},
  PlotRange → {All, {-0.5, 12.5}}, Frame → {{True, False}, {True, False}},
  BaseStyle → {FontWeight → "Bold", FontColor → Black, FontSize → 12}]
```

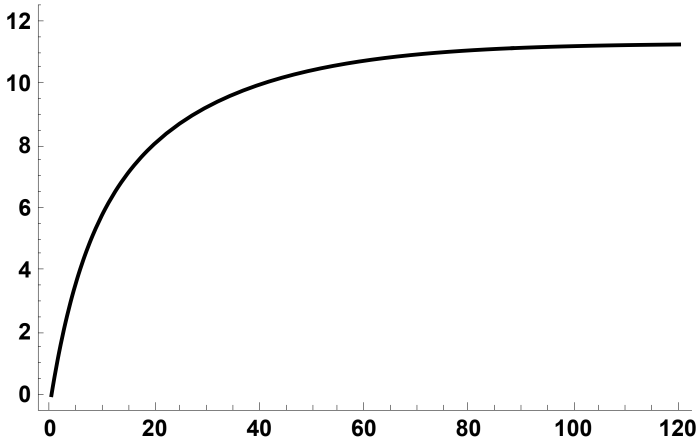


SLiM simulation plot

```
Sh05S = ErrorListPlot[SSh05S,
  PlotStyle → {Black, PointSize[0.02]}, PlotRange → {{0, 120}, {0, 12}}]
```

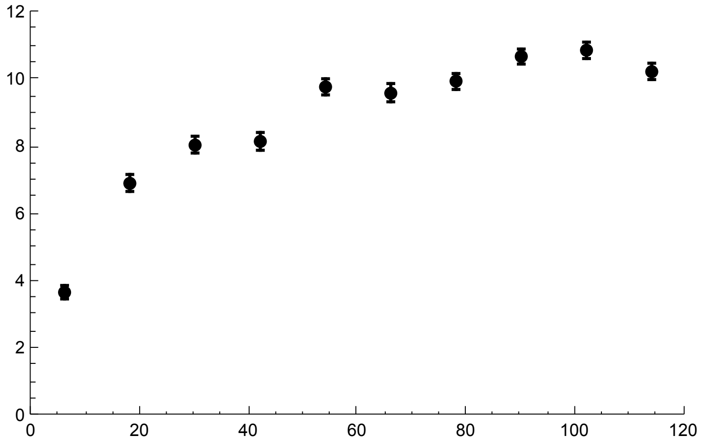


MSMS simulation plot

```
Sh05MSMSS = ErrorListPlot[SSh05MSMSS,
  PlotStyle → {Gray, PointSize[0.02]}, PlotRange → {{0, 120}, {0, 12}}]
```

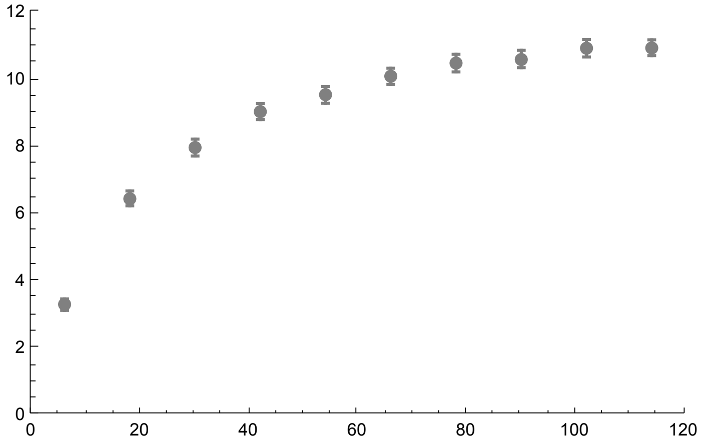


All together

```
h05MSMSSegsites = Show[Sh05, Sh05S, Sh05MSMSS, ImageSize → 200]
```

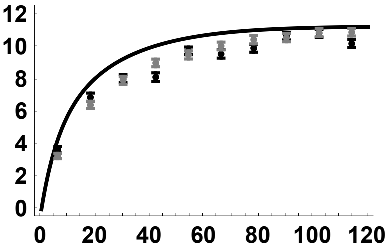


h = 0.1

SLiM simulation data

```
SSh01 = Import["SLIM_F0/StatsProc_R_120_h_0.1_self_0_f_1e-04_10b_SLIM.dat",
    "Table"][[1]];
SSh01CIB = Import["SLIM_F0/StatsProc_R_120_h_0.5_self_0_f_1e-04_10b_SLIM.dat",
    "Table"][[2]];
SSh01CIT = Import["SLIM_F0/StatsProc_R_120_h_0.5_self_0_f_1e-04_10b_SLIM.dat",
    "Table"][[3]];
SSh01S = Partition[Riffle[Partition[Riffle[Rin, SSh01], 2],
    Map[ErrorBar, Partition[Riffle[-SSh01CIB, SSh01CIT], 2]]], 2];
```

MSMS simulation data

```
SSh01MSMS =
  Import["MSMS_SV_5K/StatsProc_R_240_h_0.1_self_0_f_1e-04_10b_MSMS.dat",
    "Table"][[1]];
SSh01MSMSCIB = Import[
    "MSMS_SV_5K/StatsProc_R_240_h_0.1_self_0_f_1e-04_10b_MSMS.dat",
    "Table"][[2]];
SSh01MSMSCIT = Import[
    "MSMS_SV_5K/StatsProc_R_240_h_0.1_self_0_f_1e-04_10b_MSMS.dat",
    "Table"][[3]];
SSh01MSMSS = Partition[Riffle[Partition[Riffle[Rin, SSh01MSMS], 2],
    Map[ErrorBar, Partition[Riffle[-SSh01MSMSCIB, SSh01MSMSCIT], 2]]], 2];
```

Analytical result

```
Sh01 = Plot[ESS2[5000, 0, 0.05, 0.1, R, 4, Boostp0[5000, 0.05, 0.1, 0], 10],
  {R, 0, 120}, PlotStyle → {{Red, Thick}},
  PlotRange → {All, {-0.5, 12.5}}, Frame → {{True, False}, {True, False}},
  BaseStyle → {FontWeight → "Bold", FontColor → Black, FontSize → 12}]
```

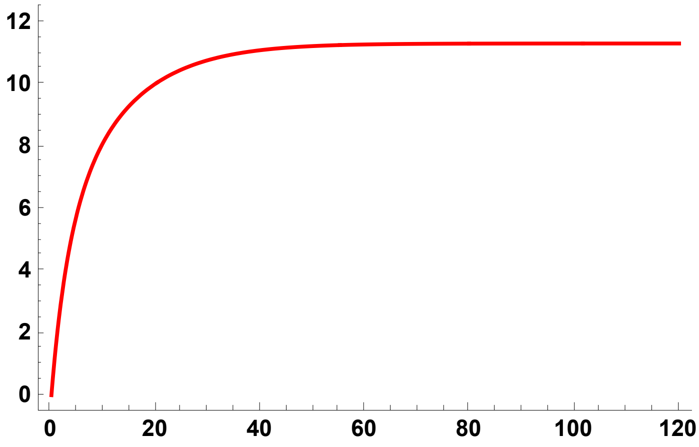


SLiM simulation plot

```
Sh01S = ErrorListPlot[SSh01S,
  PlotStyle → {Red, PointSize[0.02]}, PlotRange → {{0, 120}, {0, 12}}]
```

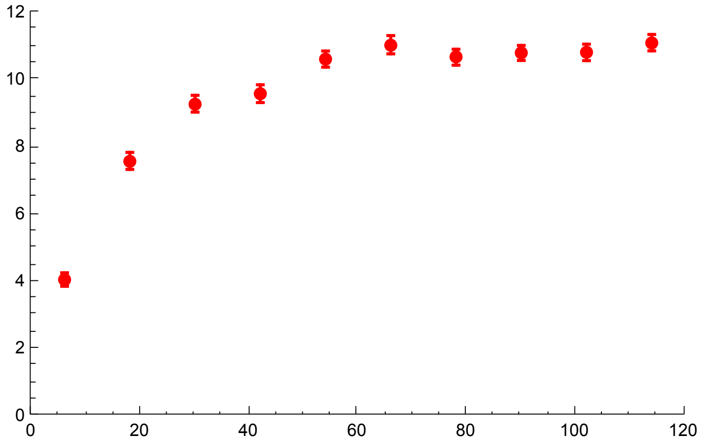


MSMS simulation plot

```
Sh01MSMSS = ErrorListPlot[SSh01MSMSS,
  PlotStyle → {Pink, PointSize[0.02]}, PlotRange → {{0, 120}, {0, 12}}]
```

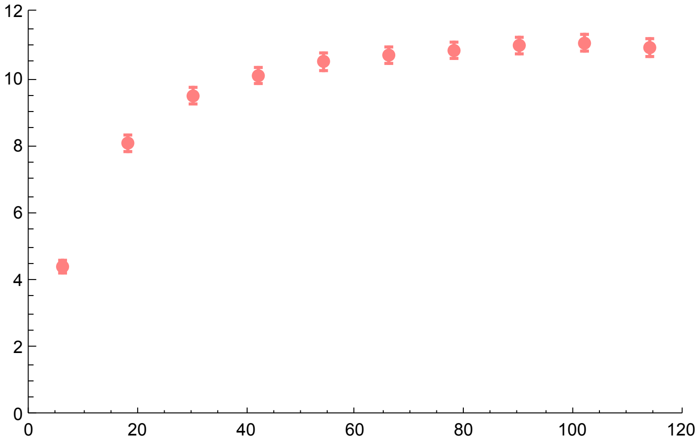


All together

```
h01MSMSSegsites = Show[Sh01, Sh01S, Sh01MSMSS, ImageSize → 200]
```

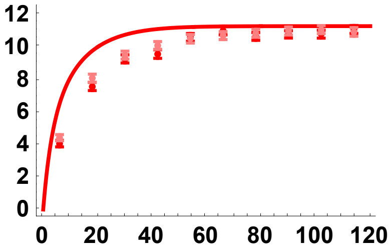


h = 0.9

SLiM simulation data

```
SSh09 = Import["SLIM_F0/StatsProc_R_120_h_0.9_self_0_f_1e-04_10b_SLIM.dat",
    "Table"][[1]];
SSh09CIB = Import["SLIM_F0/StatsProc_R_120_h_0.9_self_0_f_1e-04_10b_SLIM.dat",
    "Table"][[2]];
SSh09CIT = Import["SLIM_F0/StatsProc_R_120_h_0.9_self_0_f_1e-04_10b_SLIM.dat",
    "Table"][[3]];
SSh09S = Partition[Riffle[Partition[Riffle[Rin, SSh09], 2],
    Map[ErrorBar, Partition[Riffle[-SSh09CIB, SSh09CIT], 2]]], 2];
```

MSMS simulation data

```
SSh09MSMS =
  Import["MSMS_SV_5K/StatsProc_R_240_h_0.9_self_0_f_1e-04_10b_MSMS.dat",
    "Table"][[1]];
SSh09MSMSCIB = Import[
    "MSMS_SV_5K/StatsProc_R_240_h_0.9_self_0_f_1e-04_10b_MSMS.dat",
    "Table"][[2]];
SSh09MSMSCIT = Import[
    "MSMS_SV_5K/StatsProc_R_240_h_0.9_self_0_f_1e-04_10b_MSMS.dat",
    "Table"][[3]];
SSh09MSMSS = Partition[Riffle[Partition[Riffle[Rin, SSh09MSMS], 2],
    Map[ErrorBar, Partition[Riffle[-SSh09MSMSCIB, SSh09MSMSCIT], 2]]], 2];
```

Analytical result

```
Sh09 = Plot[ESS2[5000, 0, 0.05, 0.9, R, 4, Boostp0[5000, 0.05, 0.9, 0], 10],
  {R, 0, 120}, PlotStyle → {{Blue, Thick}},
  PlotRange → {All, {-0.5, 12.5}}, Frame → {{True, False}, {True, False}},
  BaseStyle → {FontWeight → "Bold", FontColor → Black, FontSize → 12}]
```

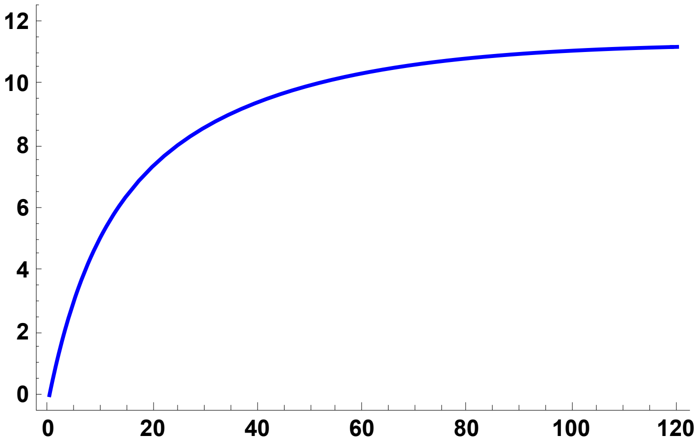


SLiM simulation plot

```
Sh09S = ErrorListPlot[SSh09S,
  PlotStyle → {Blue, PointSize[0.02]}, PlotRange → {{0, 120}, {0, 12}}]
```

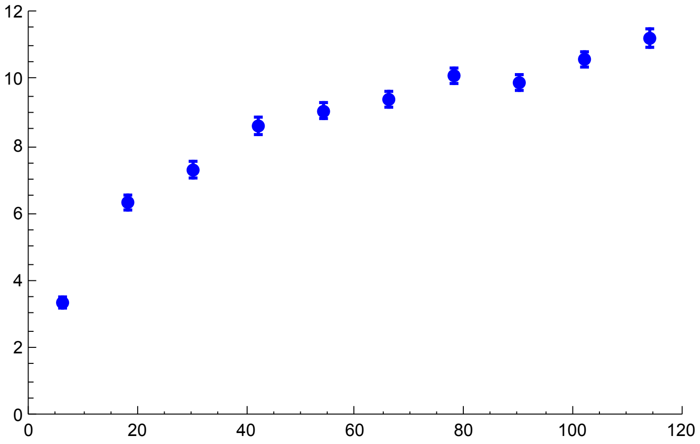


MSMS simulation plot

```
Sh09MSMSS = ErrorListPlot[SSh09MSMSS,
  PlotStyle → {Cyan, PointSize[0.02]}, PlotRange → {{0, 120}, {0, 12}}]
```

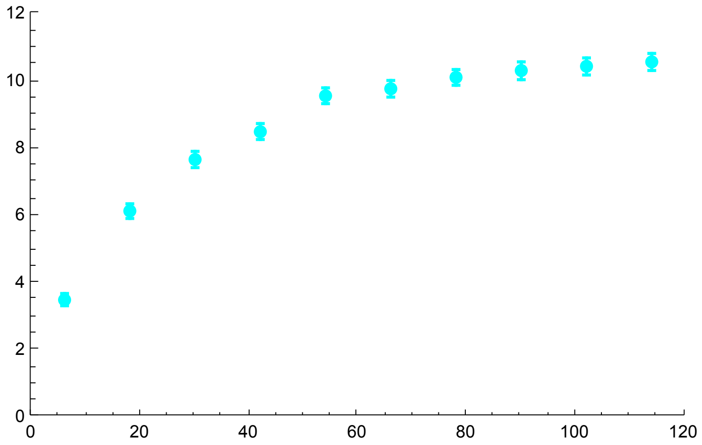


All together

```
h09MSMSSegsites = Show[Sh09, Sh09S, Sh09MSMSS, ImageSize → 200]
```

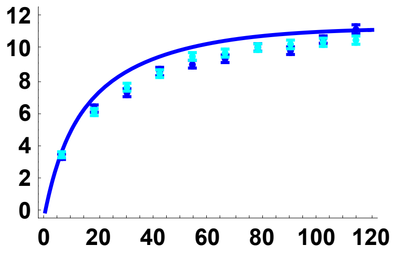


All *h* results, analytical solution and SLiM results

```
SimSegSitess0DN = Show[Sh01, Sh01S, Sh05, Sh05S, Sh09, Sh09S, ImageSize → 200]
```

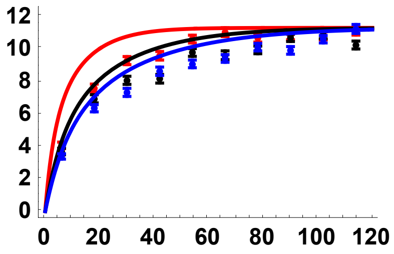


#### Simulation comparisons, $p_0$ = 0.02

h = 0.5

SLiM simulation data

```
SSh05p002 = Import[
    "SLIM_F0/StatsProc_R_120_h_0.5_self_0_f_0.02_10b_SLIM.dat", "Table"][[1]];
SSh05p002CIB = Import[
    "SLIM_F0/StatsProc_R_120_h_0.5_self_0_f_0.02_10b_SLIM.dat", "Table"][[2]];
SSh05p002CIT = Import[
    "SLIM_F0/StatsProc_R_120_h_0.5_self_0_f_0.02_10b_SLIM.dat", "Table"][[3]];
SSh05p002S = Partition[Riffle[Partition[Riffle[Rin, SSh05p002], 2],
    Map[ErrorBar, Partition[Riffle[-SSh05p002CIB, SSh05p002CIT], 2]]], 2];
```

MSMS simulation data

```
SSh05p002MSMS =
  Import["MSMS_SV_5K/StatsProc_R_240_h_0.5_self_0_f_0.02_10b_MSMS.dat",
    "Table"][[1]];
SSh05p002MSMSCIB = Import[
    "MSMS_SV_5K/StatsProc_R_240_h_0.5_self_0_f_0.02_10b_MSMS.dat",
    "Table"][[2]];
SSh05p002MSMSCIT = Import[
    "MSMS_SV_5K/StatsProc_R_240_h_0.5_self_0_f_0.02_10b_MSMS.dat",
    "Table"][[3]];
SSh05p002MSMSS = Partition[Riffle[Partition[Riffle[Rin, SSh05p002MSMS], 2], Map[
     ErrorBar, Partition[Riffle[-SSh05p002MSMSCIB, SSh05p002MSMSCIT], 2]]], 2];
```

Analytical result

```
SSPh05p002 = Plot[ESS2[5000, 0, 0.05, 0.5, R, 4, 0.02, 10],
  {R, 0, 120}, PlotStyle → {{Black, Thick}},
  PlotRange → {All, {-0.5, 12.5}}, Frame → {{True, False}, {True, False}},
  BaseStyle → {FontWeight → "Bold", FontColor → Black, FontSize → 12}]
```

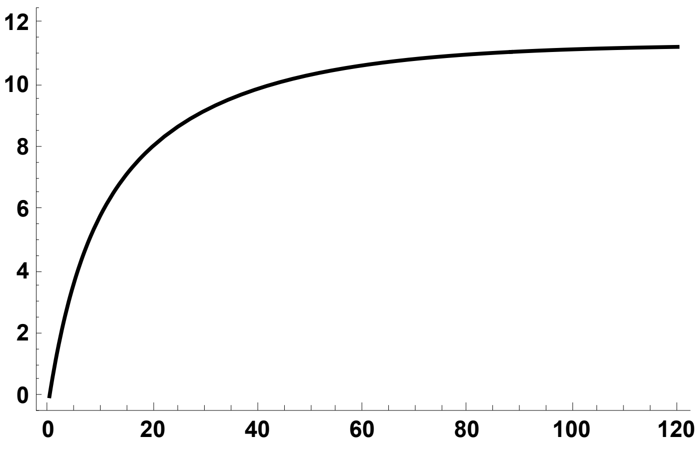


SLiM simulation plot

```
SPh05p002S = ErrorListPlot[SSh05p002S,
  PlotStyle → {Black, PointSize[0.02]}, PlotRange → {{0, 120}, {0, 12}}]
```

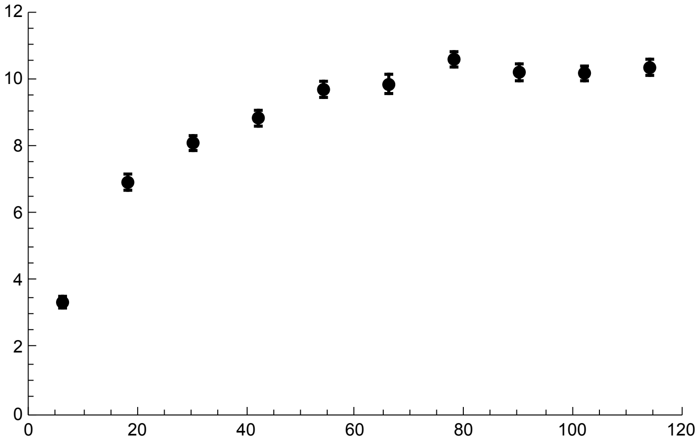


MSMS simulation plot

```
SPh05p002MSMSS = ErrorListPlot[SSh05p002MSMSS,
  PlotStyle → {Gray, PointSize[0.02]}, PlotRange → {{0, 120}, {0, 12}}]
```

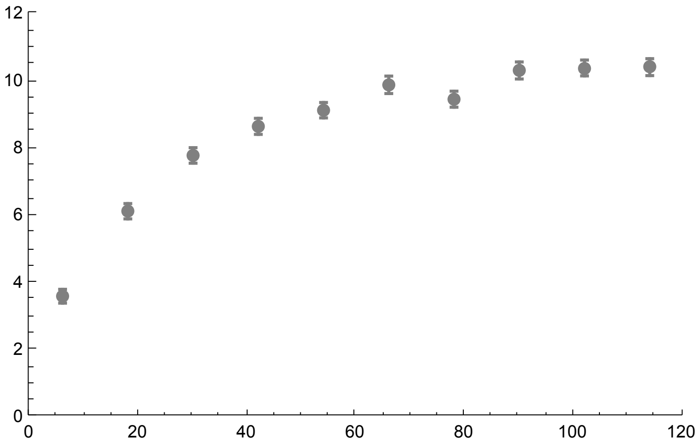


All together

```
h05MSMS2pcSegsites = Show[SSPh05p002, SPh05p002S, SPh05p002MSMSS, ImageSize → 200]
```

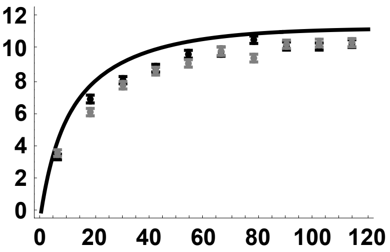


h = 0.1

SLiM simulation data

```
SSh01p002 = Import[
    "SLIM_F0/StatsProc_R_120_h_0.1_self_0_f_0.02_10b_SLIM.dat", "Table"][[1]];
SSh01p002CIB = Import[
    "SLIM_F0/StatsProc_R_120_h_0.1_self_0_f_0.02_10b_SLIM.dat", "Table"][[2]];
SSh01p002CIT = Import[
    "SLIM_F0/StatsProc_R_120_h_0.1_self_0_f_0.02_10b_SLIM.dat", "Table"][[3]];
SSh01p002S = Partition[Riffle[Partition[Riffle[Rin, SSh01p002], 2],
    Map[ErrorBar, Partition[Riffle[-SSh01p002CIB, SSh01p002CIT], 2]]], 2];
```

MSMS simulation data

```
SSh01p002MSMS =
  Import["MSMS_SV_5K/StatsProc_R_240_h_0.1_self_0_f_0.02_10b_MSMS.dat",
    "Table"][[1]];
SSh01p002MSMSCIB = Import[
    "MSMS_SV_5K/StatsProc_R_240_h_0.1_self_0_f_0.02_10b_MSMS.dat",
    "Table"][[2]];
SSh01p002MSMSCIT = Import[
    "MSMS_SV_5K/StatsProc_R_240_h_0.1_self_0_f_0.02_10b_MSMS.dat",
    "Table"][[3]];
SSh01p002MSMSS = Partition[Riffle[Partition[Riffle[Rin, SSh01p002MSMS], 2], Map[
     ErrorBar, Partition[Riffle[-SSh01p002MSMSCIB, SSh01p002MSMSCIT], 2]]], 2];
```

Analytical result

```
SSPh01p002 = Plot[ESS2[5000, 0, 0.05, 0.1, R, 4, 0.02, 10],
  {R, 0, 120}, PlotStyle → {{Red, Thick}},
  PlotRange → {All, {-0.5, 12.5}}, Frame → {{True, False}, {True, False}},
  BaseStyle → {FontWeight → "Bold", FontColor → Black, FontSize → 12}]
```

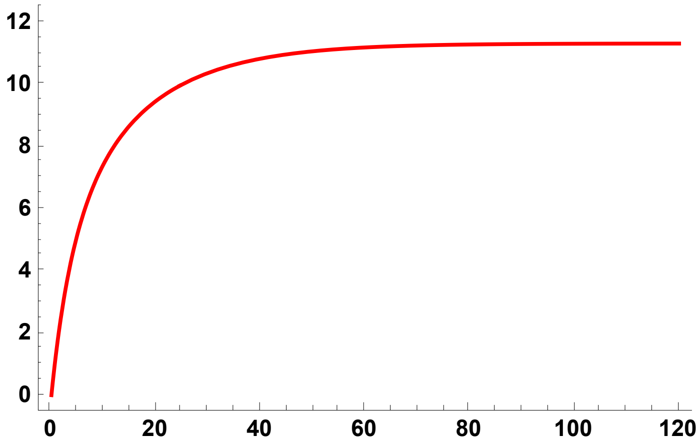


SLiM simulation plot

```
SPh01p002S = ErrorListPlot[SSh01p002S,
  PlotStyle → {Red, PointSize[0.02]}, PlotRange → {{0, 120}, {0, 12}}]
```

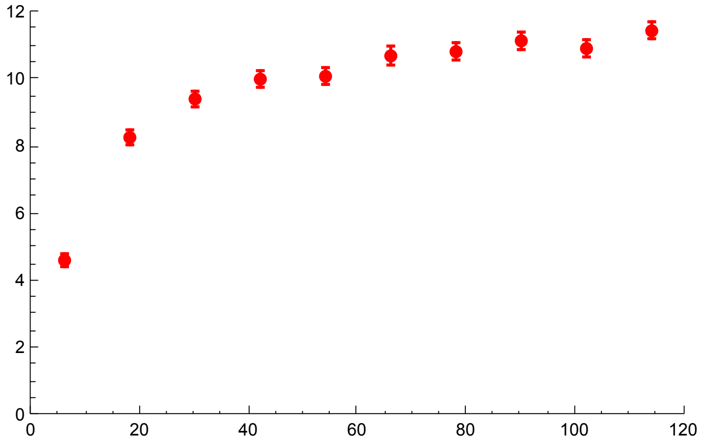


MSMS simulation plot

```
SPh01p002MSMSS = ErrorListPlot[SSh01p002MSMSS,
  PlotStyle → {Pink, PointSize[0.02]}, PlotRange → {{0, 120}, {0, 12}}]
```

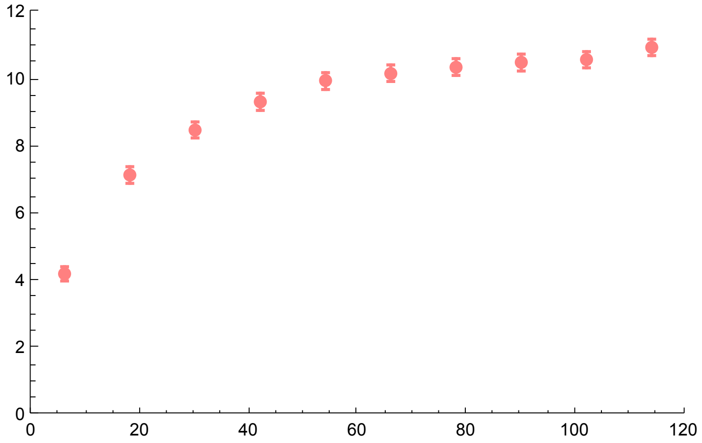


All together

```
h01MSMS2pcSegsites = Show[SSPh01p002, SPh01p002S, SPh01p002MSMSS, ImageSize → 200]
```

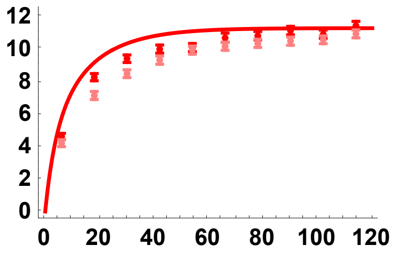


h = 0.9

SLIM simulation data

```
SSh09p002 = Import[
    "SLIM_F0/StatsProc_R_120_h_0.9_self_0_f_0.02_10b_SLIM.dat", "Table"][[1]];
SSh09p002CIB = Import[
    "SLIM_F0/StatsProc_R_120_h_0.9_self_0_f_0.02_10b_SLIM.dat", "Table"][[2]];
SSh09p002CIT = Import[
    "SLIM_F0/StatsProc_R_120_h_0.9_self_0_f_0.02_10b_SLIM.dat", "Table"][[3]];
SSh09p002S = Partition[Riffle[Partition[Riffle[Rin, SSh09p002], 2],
    Map[ErrorBar, Partition[Riffle[-SSh09p002CIB, SSh09p002CIT], 2]]], 2];
```

MSMS simulation data

```
SSh09p002MSMS =
  Import["MSMS_SV_5K/StatsProc_R_240_h_0.9_self_0_f_0.02_10b_MSMS.dat",
    "Table"][[1]];
SSh09p002MSMSCIB = Import[
    "MSMS_SV_5K/StatsProc_R_240_h_0.9_self_0_f_0.02_10b_MSMS.dat",
    "Table"][[2]];
SSh09p002MSMSCIT = Import[
    "MSMS_SV_5K/StatsProc_R_240_h_0.9_self_0_f_0.02_10b_MSMS.dat",
    "Table"][[3]];
SSh09p002MSMSS = Partition[Riffle[Partition[Riffle[Rin, SSh09p002MSMS], 2], Map[
     ErrorBar, Partition[Riffle[-SSh09p002MSMSCIB, SSh09p002MSMSCIT], 2]]], 2];
```

Analytical result

```
SSPh09p002 = Plot[ESS2[5000, 0, 0.05, 0.9, R, 4, 0.02, 10],
  {R, 0, 120}, PlotStyle → {{Blue, Thick}},
  PlotRange → {All, {-0.5, 12.5}}, Frame → {{True, False}, {True, False}},
  BaseStyle → {FontWeight → "Bold", FontColor → Black, FontSize → 12}]
```

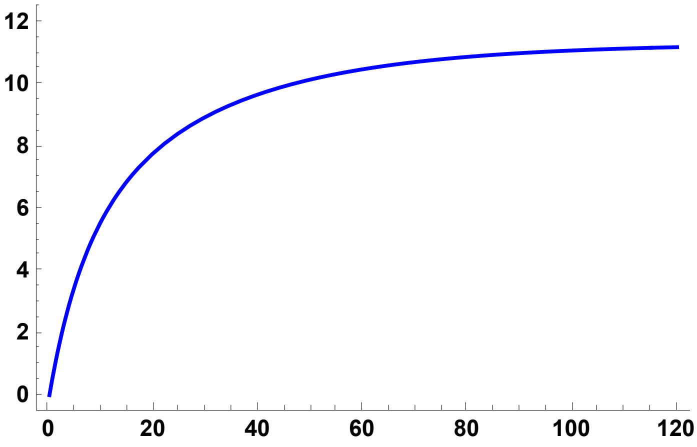


SLiM simulation plot

```
SPh09p002S = ErrorListPlot[SSh09p002S,
  PlotStyle → {Blue, PointSize[0.02]}, PlotRange → {{0, 120}, {0, 12}}]
```

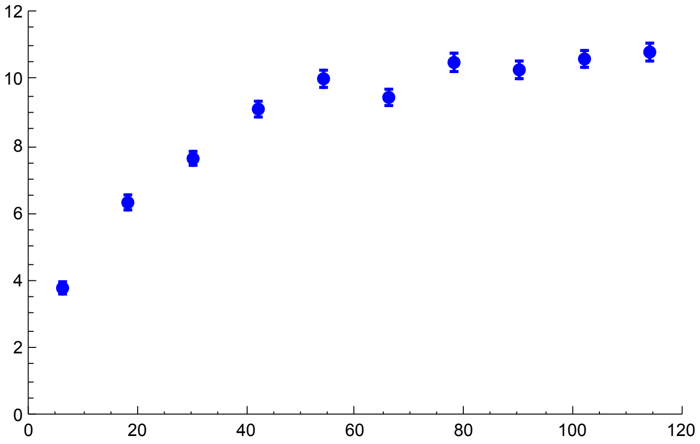


MSMS simulation plot

```
SPh09p002MSMSS = ErrorListPlot[SSh09p002MSMSS,
  PlotStyle → {Cyan, PointSize[0.02]}, PlotRange → {{0, 120}, {0, 12}}]
```

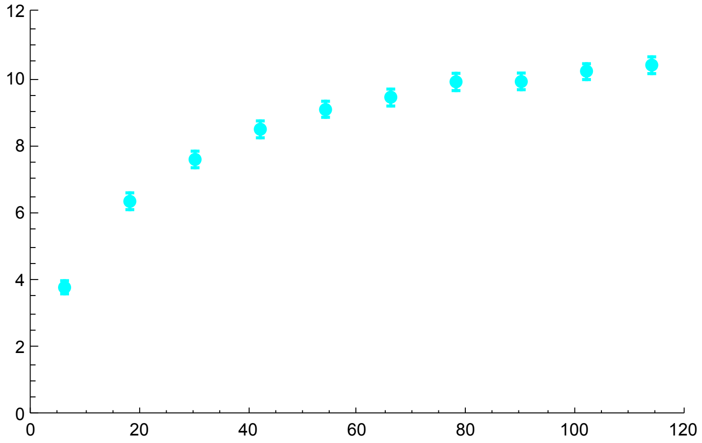

All together

```
h09MSMS2pcSegsites = Show[SSPh09p002, SPh09p002S, SPh09p002MSMSS, ImageSize → 200]
```

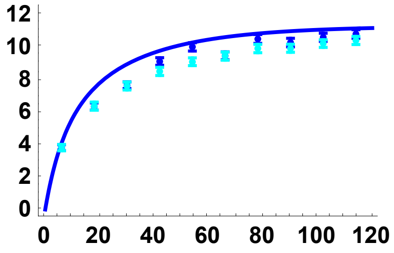

All *h* results, analytical solution and SLiM results

```
SimSegSitess02pc = Show[SSPh01p002, SPh01p002S,
  SSPh05p002, SPh05p002S, SSPh09p002, SPh09p002S, ImageSize → 200]
```

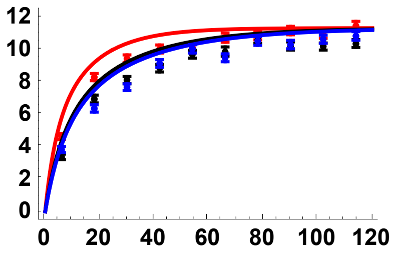

#### Simulation comparisons, $p_0$ = 0.05

h = 0.5

SLiM simulation data

```
SSh05p005 = Import[
    "SLIM_F0/StatsProc_R_120_h_0.5_self_0_f_0.05_10b_SLIM.dat", "Table"][[1]];
SSh05p005CIB = Import[
    "SLIM_F0/StatsProc_R_120_h_0.5_self_0_f_0.05_10b_SLIM.dat", "Table"][[2]];
SSh05p005CIT = Import[
    "SLIM_F0/StatsProc_R_120_h_0.5_self_0_f_0.05_10b_SLIM.dat", "Table"][[3]];
SSh05p005S = Partition[Riffle[Partition[Riffle[Rin, SSh05p005], 2],
    Map[ErrorBar, Partition[Riffle[-SSh05p005CIB, SSh05p005CIT], 2]]], 2];
```

MSMS simulation data

```
SSh05p005MSMS =
  Import["MSMS_SV_5K/StatsProc_R_240_h_0.5_self_0_f_0.05_10b_MSMS.dat",
    "Table"][[1]];
SSh05p005MSMSCIB = Import[
    "MSMS_SV_5K/StatsProc_R_240_h_0.5_self_0_f_0.05_10b_MSMS.dat",
    "Table"][[2]];
SSh05p005MSMSCIT = Import[
    "MSMS_SV_5K/StatsProc_R_240_h_0.5_self_0_f_0.05_10b_MSMS.dat",
    "Table"][[3]];
SSh05p005MSMSS = Partition[Riffle[Partition[Riffle[Rin, SSh05p005MSMS], 2], Map[
      ErrorBar, Partition[Riffle[-SSh05p005MSMSCIB, SSh05p005MSMSCIT], 2]]], 2];
```

Analytical result

```
SSPh05p005 = Plot[ESS2[5000, 0, 0.05, 0.5, R, 4, 0.05, 10],
  {R, 0, 120}, PlotStyle → {{Black, Thick}},
  PlotRange → {All, {-0.5, 12.5}}, Frame → {{True, False}, {True, False}},
  BaseStyle → {FontWeight → "Bold", FontColor → Black, FontSize → 12}]
```

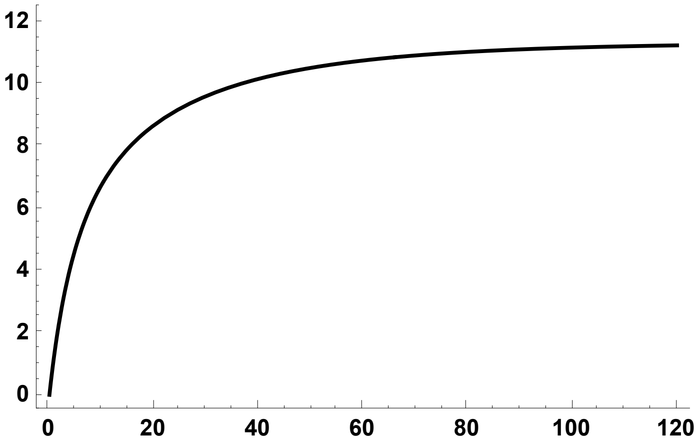


SLiM simulation plot

```
SPh05p005S = ErrorListPlot[SSh05p005S,
  PlotStyle → {Black, PointSize[0.02]}, PlotRange → {{0, 120}, {0, 12.5}}]
```

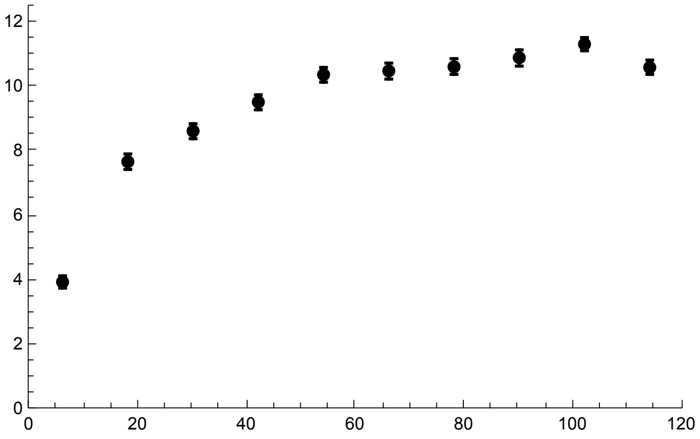


MSMS simulation plot

```
SPh05p005MSMSS = ErrorListPlot[SSh05p005MSMSS,
  PlotStyle → {Gray, PointSize[0.02]}, PlotRange → {{0, 120}, {0, 12}}]
```

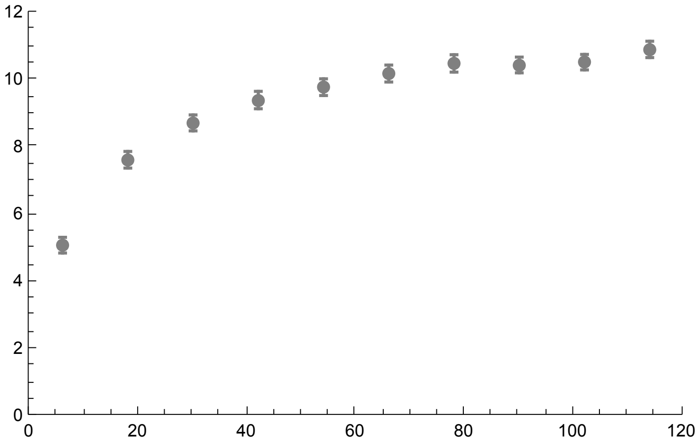


All together

```
h05MSMS5pcSegSites = Show[SSPh05p005, SPh05p005S, SPh05p005MSMSS, ImageSize → 200]
```

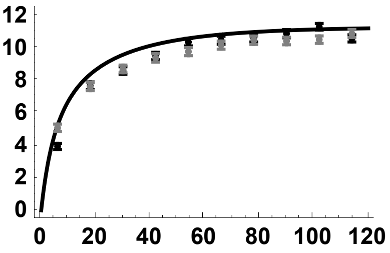


h = 0.1

SLiM simulation data

```
SSh01p005 = Import[
    "SLIM_F0/StatsProc_R_120_h_0.1_self_0_f_0.05_10b_SLIM.dat", "Table"][[1]];
SSh01p005CIB = Import[
    "SLIM_F0/StatsProc_R_120_h_0.1_self_0_f_0.05_10b_SLIM.dat", "Table"][[2]];
SSh01p005CIT = Import[
    "SLIM_F0/StatsProc_R_120_h_0.1_self_0_f_0.05_10b_SLIM.dat", "Table"][[3]];
SSh01p005S = Partition[Riffle[Partition[Riffle[Rin, SSh01p005], 2],
    Map[ErrorBar, Partition[Riffle[-SSh01p005CIB, SSh01p005CIT], 2]]], 2];
```

MSMS simulation data

```
SSh01p005MSMS =
  Import["MSMS_SV_5K/StatsProc_R_240_h_0.1_self_0_f_0.05_10b_MSMS.dat",
    "Table"][[1]];
SSh01p005MSMSCIB = Import[
    "MSMS_SV_5K/StatsProc_R_240_h_0.1_self_0_f_0.05_10b_MSMS.dat",
    "Table"][[2]];
SSh01p005MSMSCIT = Import[
    "MSMS_SV_5K/StatsProc_R_240_h_0.1_self_0_f_0.05_10b_MSMS.dat",
    "Table"][[3]];
SSh01p005MSMSS = Partition[Riffle[Partition[Riffle[Rin, SSh01p005MSMS], 2], Map[
     ErrorBar, Partition[Riffle[-SSh01p005MSMSCIB, SSh01p005MSMSCIT], 2]]], 2];
```

Analytical result

```
SSPh01p005 = Plot[ESS2[5000, 0, 0.05, 0.1, R, 4, 0.05, 10],
  {R, 0, 120}, PlotStyle → {{Red, Thick}},
  PlotRange → {All, {-0.5, 12.5}}, Frame → {{True, False}, {True, False}},
  BaseStyle → {FontWeight → "Bold", FontColor → Black, FontSize → 12}]
```

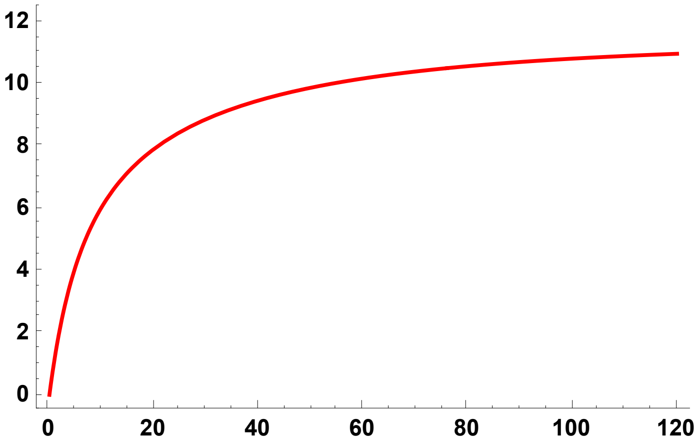


SLiM simulation plot

```
SPh01p005S = ErrorListPlot[SSh01p005S,
  PlotStyle → {Red, PointSize[0.02]}, PlotRange → {{0, 120}, {0, 12}}]
```

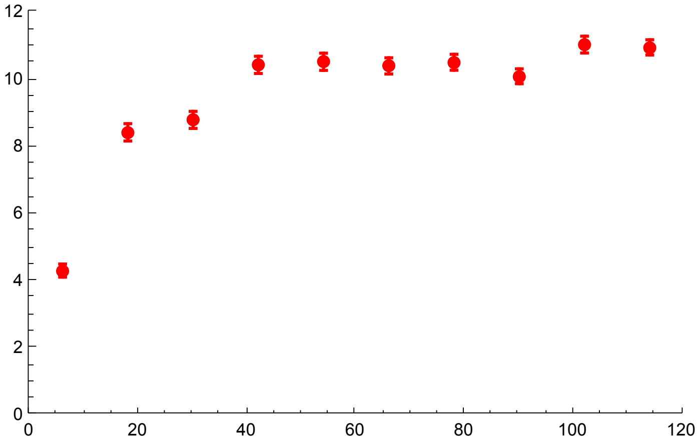


MSMS simulation plot

```
SPh01p005MSMSS = ErrorListPlot[SSh01p005MSMSS,
  PlotStyle → {Pink, PointSize[0.02]}, PlotRange → {{0, 120}, {0, 12}}]
```

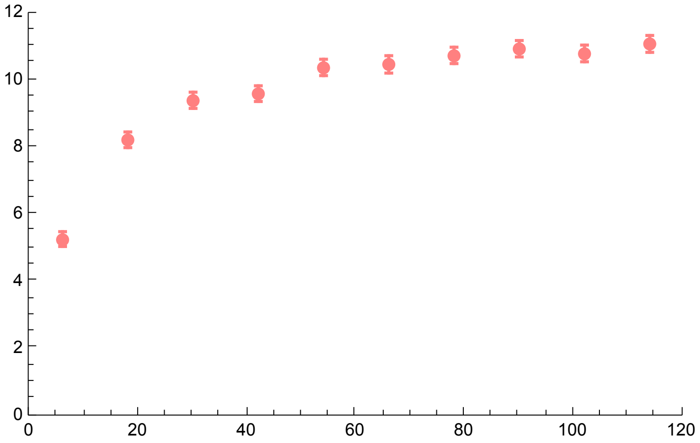


All together

```
h01MSMS5pcSegSites = Show[SSPh01p005, SPh01p005S, SPh01p005MSMSS, ImageSize → 200]
```

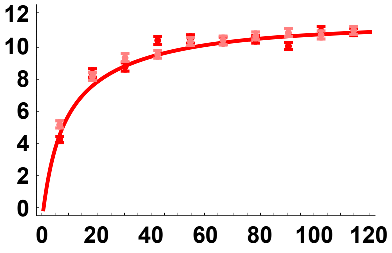


h = 0.9

SLiM simulation data

```
SSh09p005 = Import[
    "SLIM_F0/StatsProc_R_120_h_0.9_self_0_f_0.05_10b_SLIM.dat", "Table"][[1]];
SSh09p005CIB = Import[
    "SLIM_F0/StatsProc_R_120_h_0.9_self_0_f_0.05_10b_SLIM.dat", "Table"][[2]];
SSh09p005CIT = Import[
    "SLIM_F0/StatsProc_R_120_h_0.9_self_0_f_0.05_10b_SLIM.dat", "Table"][[3]];
SSh09p005S = Partition[Riffle[Partition[Riffle[Rin, SSh09p005], 2],
    Map[ErrorBar, Partition[Riffle[-SSh09p005CIB, SSh09p005CIT], 2]]], 2];
```

MSMS simulation data

```
SSh09p005MSMS =
  Import["MSMS_SV_5K/StatsProc_R_240_h_0.9_self_0_f_0.05_10b_MSMS.dat",
    "Table"][[1]];
SSh09p005MSMSCIB = Import[
    "MSMS_SV_5K/StatsProc_R_240_h_0.9_self_0_f_0.05_10b_MSMS.dat",
    "Table"][[2]];
SSh09p005MSMSCIT = Import[
    "MSMS_SV_5K/StatsProc_R_240_h_0.9_self_0_f_0.05_10b_MSMS.dat",
    "Table"][[3]];
SSh09p005MSMSS = Partition[Riffle[Partition[Riffle[Rin, SSh09p005MSMS], 2], Map[
     ErrorBar, Partition[Riffle[-SSh09p005MSMSCIB, SSh09p005MSMSCIT], 2]]], 2];
```

Analytical result

```
SSPh09p005 = Plot[ESS2[5000, 0, 0.05, 0.9, R, 4, 0.05, 10],
  {R, 0, 120}, PlotStyle → {{Blue, Thick}},
  PlotRange → {All, {-0.5, 12}}, Frame → {{True, False}, {True, False}},
  BaseStyle → {FontWeight → "Bold", FontColor → Black, FontSize → 12}]
```

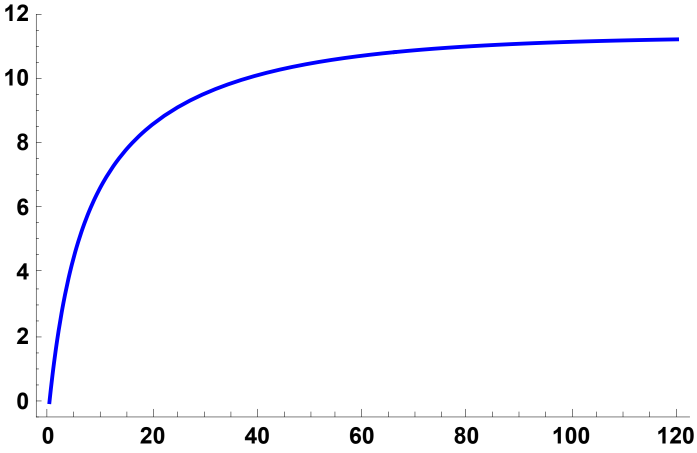


SLiM simulation plot

```
SPh09p005S = ErrorListPlot[SSh09p005S,
  PlotStyle → {Blue, PointSize[0.02]}, PlotRange → {{0, 120}, {0, 12}}]
```

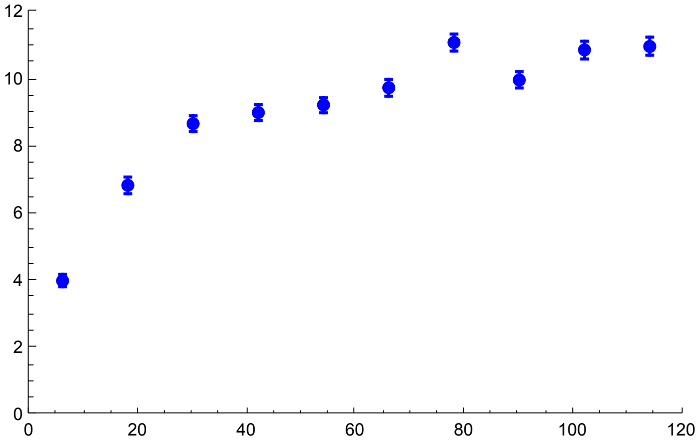


MSMS simulation plot

```
SPh09p005MSMSS = ErrorListPlot[SSh09p005MSMSS,
  PlotStyle → {Cyan, PointSize[0.02]}, PlotRange → {{0, 120}, {0, 12}}]
```

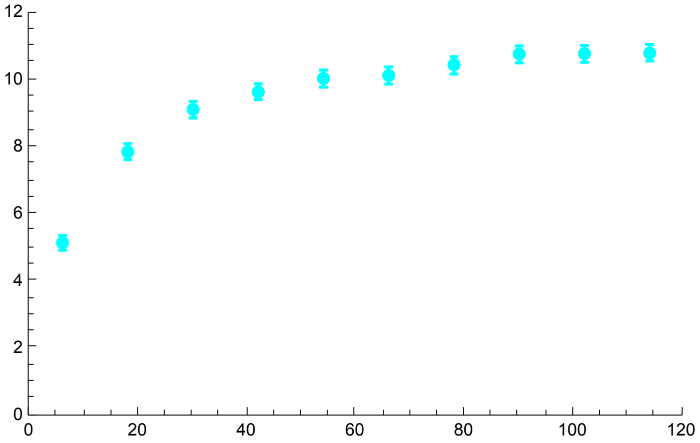


```
h09MSMS5pcSegSites = Show[SSPh09p005, SPh09p005S, SPh09p005MSMSS, ImageSize → 200]
```

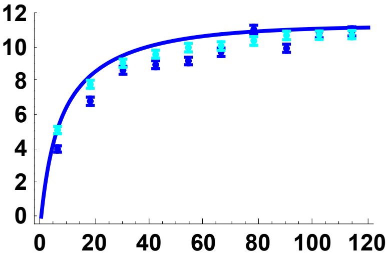


All *h* results, analytical solution and SLiM results

```
SimSegSitess05pc = Show[SSPh01p005, SPh01p005S,
  SSPh05p005, SPh05p005S, SSPh09p005, SPh09p005S, ImageSize → 200]
```

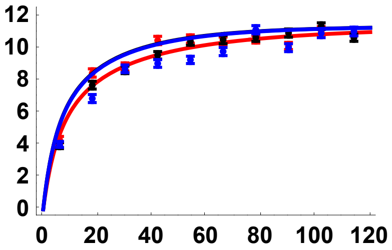


#### $\sigma = 1/2$ ($F = 1/3$)

All simulation data from here on is from SLiM.

Preliminaries

```
SetDirectory[NotebookDirectory[]];
Rin = Table[11 + 22 * i, {i, 0, 9}];
```

##### Simulation comparisons, from initial frequency $p_0 = 1/2N$

h = 0.5

```
SSh05s05 =
  Import["SLIM_F033/StatsProc_R_220_h_0.5_self_0.5_f_1e-04_10b_SLIM.dat",
    "Table"][[1]];
SSh05s05CIB = Import[
    "SLIM_F033/StatsProc_R_220_h_0.5_self_0.5_f_1e-04_10b_SLIM.dat",
    "Table"][[2]];
SSh05s05CIT = Import[
    "SLIM_F033/StatsProc_R_220_h_0.5_self_0.5_f_1e-04_10b_SLIM.dat",
    "Table"][[3]];
SSh05s05S = Partition[Riffle[Partition[Riffle[Rin, SSh05s05], 2],
    Map[ErrorBar, Partition[Riffle[-SSh05s05CIB, SSh05s05CIT], 2]]], 2];
```

```
Sh05s05 =
 Plot[{ESS2[5000, 0.5, 0.05, 0.5, R, 4/(1 + 1/3), Boostp0[5000, 0.05, 0.5, 0.5], 10]},
  {R, 0, 220}, PlotStyle → {{Black, Thick}},
  PlotRange → {All, {-0.5, 12}}, Frame → {{True, False}, {True, False}},
  BaseStyle → {FontWeight → "Bold", FontColor → Black, FontSize → 12}]
```

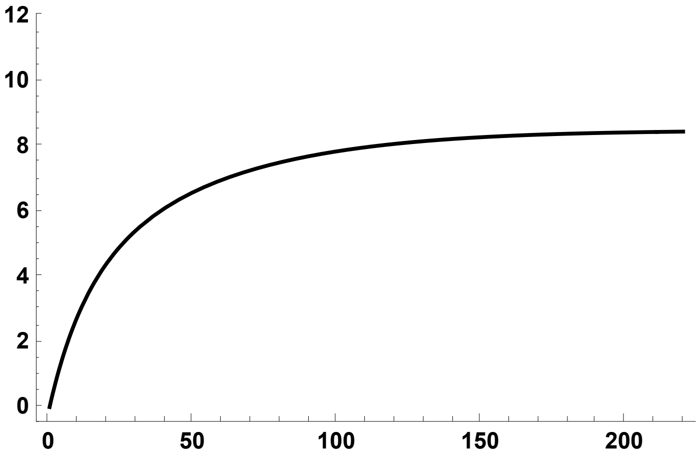


```
Sh05s05S = ErrorListPlot[SSh05s05S,
  PlotStyle → {Black, PointSize[0.02]}, PlotRange → {{0, 220}, {0, 12}}]
```

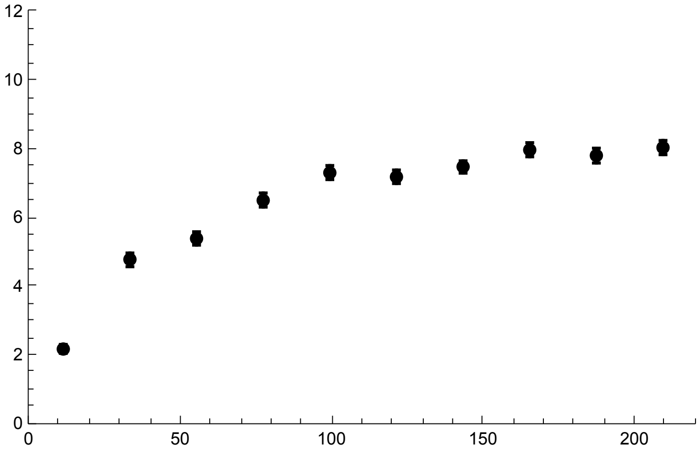


```
Show[Sh05s05, Sh05s05S]
```

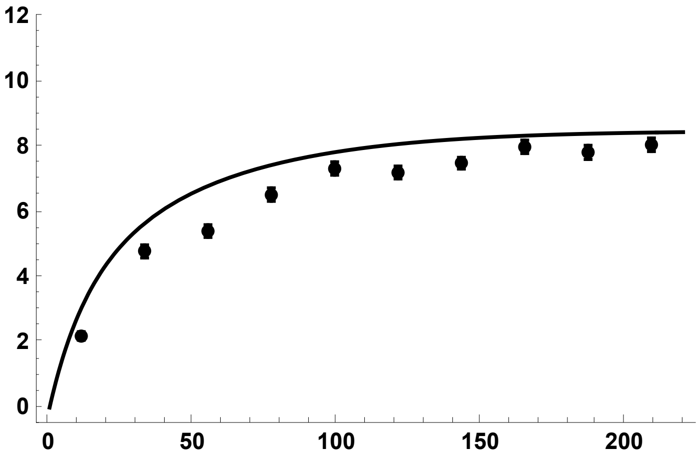

h = 0.1

```
SSh01s05 =
  Import["SLIM_F033/StatsProc_R_220_h_0.1_self_0.5_f_1e-04_10b_SLIM.dat",
    "Table"][[1]];
SSh01s05CIB = Import[
    "SLIM_F033/StatsProc_R_220_h_0.1_self_0.5_f_1e-04_10b_SLIM.dat",
    "Table"][[2]];
SSh01s05CIT = Import[
    "SLIM_F033/StatsProc_R_220_h_0.1_self_0.5_f_1e-04_10b_SLIM.dat",
    "Table"][[3]];
SSh01s05S = Partition[Riffle[Partition[Riffle[Rin, SSh01s05], 2],
    Map[ErrorBar, Partition[Riffle[-SSh01s05CIB, SSh01s05CIT], 2]]], 2];
```

```
Sh01s05 =
 Plot[{ESS2[5000, 0.5, 0.05, 0.1, R, 4/(1 + 1/3), Boostp0[5000, 0.05, 0.1, 0.5], 10]},
  {R, 0, 220}, PlotStyle → {{Red, Thick}},
  PlotRange → {All, {-0.5, 12}}, Frame → {{True, False}, {True, False}},
  BaseStyle → {FontWeight → "Bold", FontColor → Black, FontSize → 12}]
```

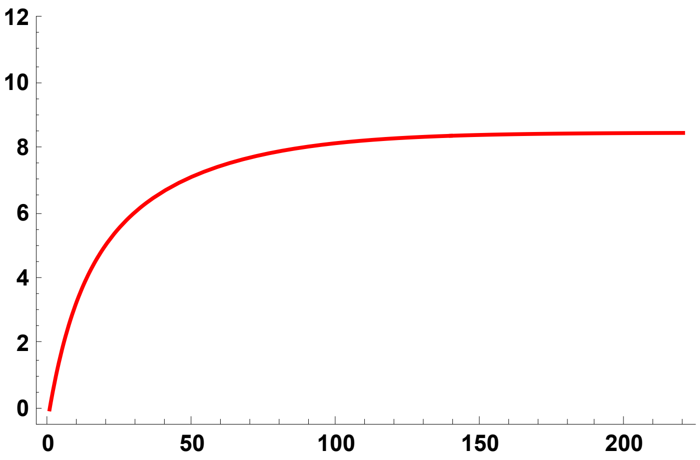


```
Sh01s05S = ErrorListPlot[SSh01s05S,
  PlotStyle → {Red, PointSize[0.02]}, PlotRange → {{0, 220}, {0, 12}}]
```

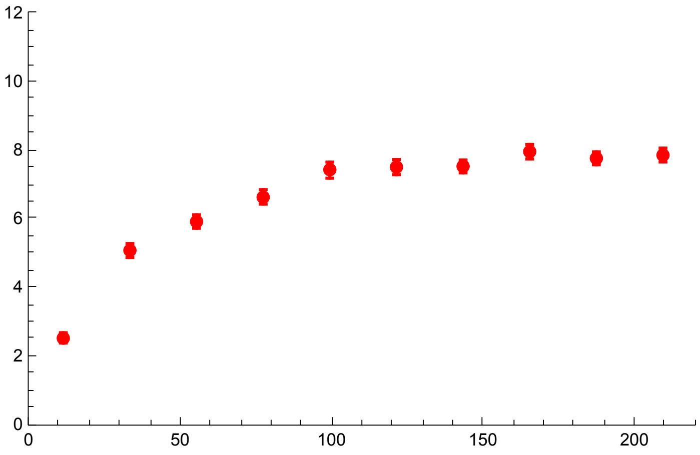

```
Show[Sh01s05, Sh01s05S]
```

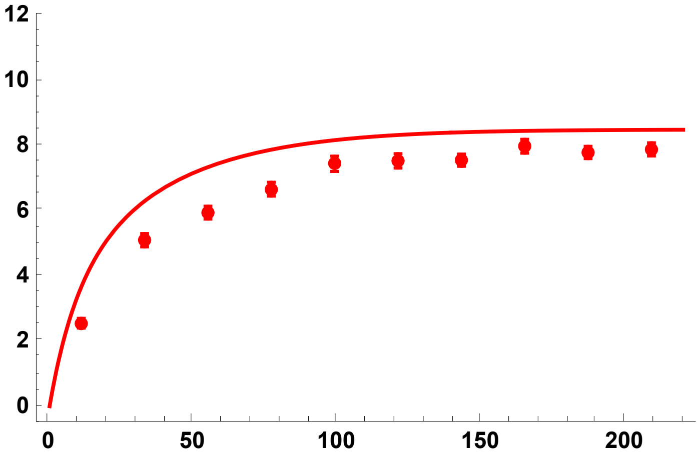


h = 0.9

```
SSh09s05 =
  Import["SLIM_F033/StatsProc_R_220_h_0.9_self_0.5_f_1e-04_10b_SLIM.dat",
    "Table"][[1]];
SSh09s05CIB = Import[
    "SLIM_F033/StatsProc_R_220_h_0.9_self_0.5_f_1e-04_10b_SLIM.dat",
    "Table"][[2]];
SSh09s05CIT = Import[
    "SLIM_F033/StatsProc_R_220_h_0.9_self_0.5_f_1e-04_10b_SLIM.dat",
    "Table"][[3]];
SSh09s05S = Partition[Riffle[Partition[Riffle[Rin, SSh09s05], 2],
    Map[ErrorBar, Partition[Riffle[-SSh09s05CIB, SSh09s05CIT], 2]]], 2];
```

```
Sh09s05 =
```

$$\text{Plot}\left[\left\{\text{ESS2}\left[5000, 0.5, 0.05, 0.9, R, \frac{4}{1+\frac{1}{3}}, \text{Boostp0}[5000, 0.05, 0.9, 0.5], 10\right]\right\},\right.$$

```
  {R, 0, 220}, PlotStyle → {{Blue, Thick}},
  PlotRange → {All, {-0.5, 12}}, Frame → {{True, False}, {True, False}},
  BaseStyle → {FontWeight → "Bold", FontColor → Black, FontSize → 12}]
```

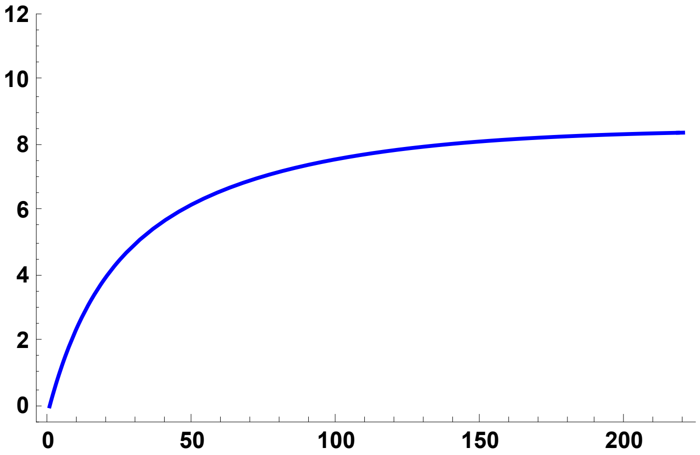

```
Sh09s05S = ErrorListPlot[SSh09s05S,
  PlotStyle → {Blue, PointSize[0.02]}, PlotRange → {{0, 220}, {0, 12}}]
```

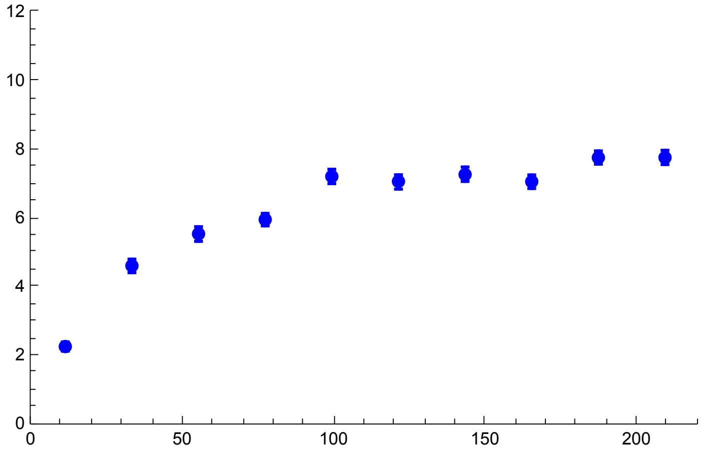

```
Show[Sh09s05, Sh09s05S]
```

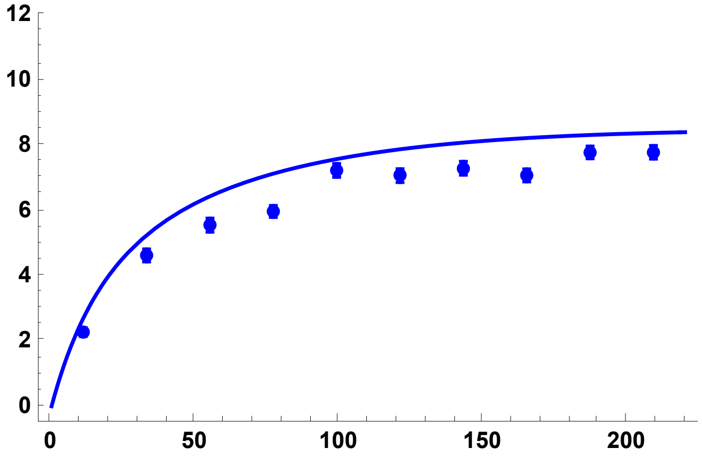

All results together

```
SimSegSitess05DN =
 Show[Sh01s05, Sh01s05S, Sh05s05, Sh05s05S, Sh09s05, Sh09s05S, ImageSize → 200]
```

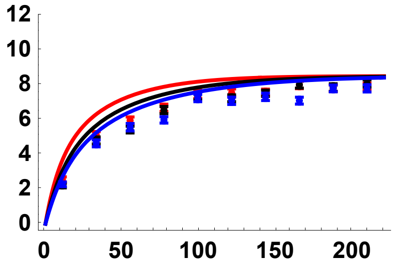

#### Simulation comparisons, $p_0$ = 0.02

h = 0.5

```
SSh05p002s05 =
  Import["SLIM_F033/StatsProc_R_220_h_0.9_self_0.5_f_0.02_10b_SLIM.dat",
    "Table"][[1]];
SSh05p002s05CIB = Import[
    "SLIM_F033/StatsProc_R_220_h_0.9_self_0.5_f_0.02_10b_SLIM.dat",
    "Table"][[2]];
SSh05p002s05CIT = Import[
    "SLIM_F033/StatsProc_R_220_h_0.9_self_0.5_f_0.02_10b_SLIM.dat",
    "Table"][[3]];
SSh05p002s05S = Partition[Riffle[Partition[Riffle[Rin, SSh05p002s05], 2],
    Map[ErrorBar, Partition[Riffle[-SSh05p002s05CIB, SSh05p002s05CIT], 2]]], 2];
```

SSPh05p002s05 = Plot[{ESS2[5000, 0.5, 0.05, 0.5, R, $\frac{4}{1+\frac{1}{3}}$, 0.02, 10]},

```
  {R, 0, 220}, PlotStyle → {{Black, Thick}},
  PlotRange → {All, {-0.5, 12}}, Frame → {{True, False}, {True, False}},
  BaseStyle → {FontWeight → "Bold", FontColor → Black, FontSize → 12}]
```

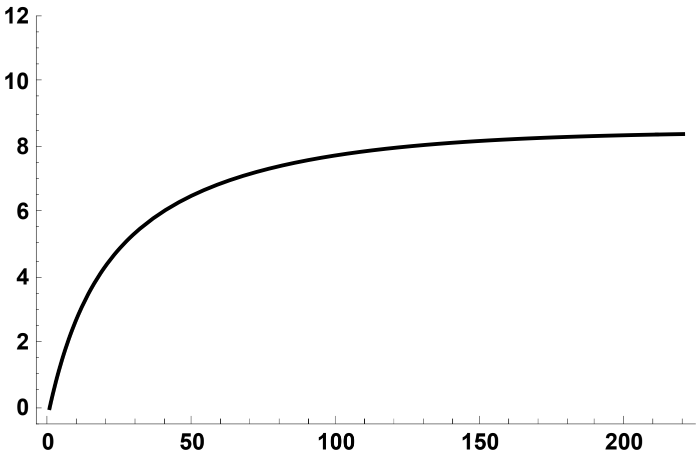

```
SPh05p002s05S = ErrorListPlot[SSh05p002s05S,
  PlotStyle → {Black, PointSize[0.02]}, PlotRange → {{0, 220}, {0, 12}}]
```

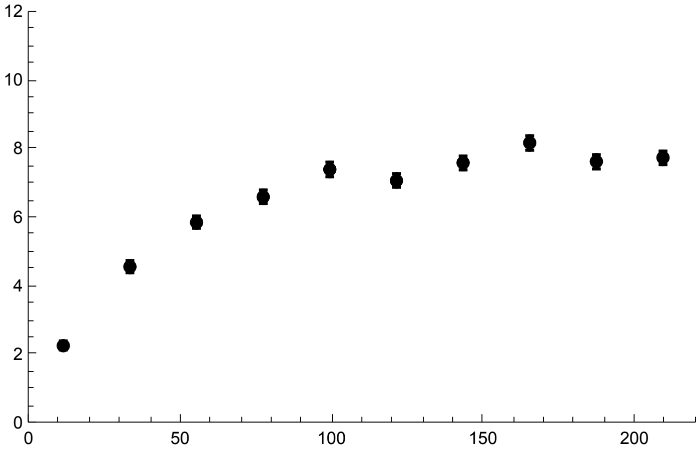

```
Show[SSPh05p002s05, SPh05p002s05S]
```

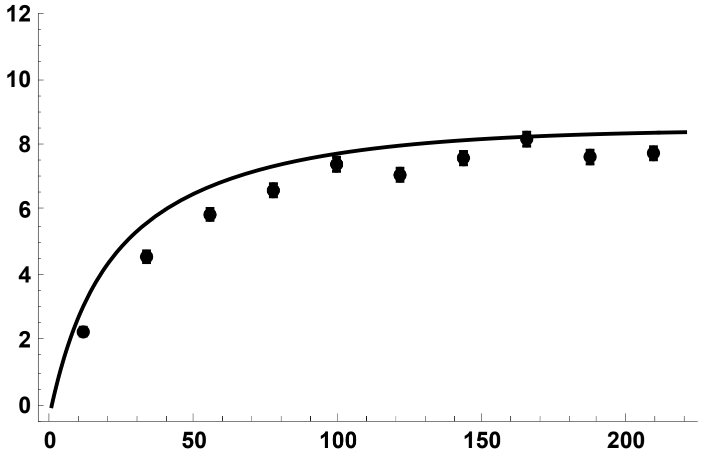


h = 0.1

```
SSh01p002s05 =
  Import["SLIM_F033/StatsProc_R_220_h_0.1_self_0.5_f_0.02_10b_SLIM.dat",
    "Table"][[1]];
SSh01p002s05CIB = Import[
    "SLIM_F033/StatsProc_R_220_h_0.1_self_0.5_f_0.02_10b_SLIM.dat",
    "Table"][[2]];
SSh01p002s05CIT = Import[
    "SLIM_F033/StatsProc_R_220_h_0.1_self_0.5_f_0.02_10b_SLIM.dat",
    "Table"][[3]];
SSh01p002s05S = Partition[Riffle[Partition[Riffle[Rin, SSh01p002s05], 2],
    Map[ErrorBar, Partition[Riffle[-SSh01p002s05CIB, SSh01p002s05CIT], 2]]], 2];
```

SSPh01p002s05 = Plot$\left[\left\{\text{ESS2}\left[5000, 0.5, 0.05, 0.1, R, \frac{4}{1+\frac{1}{3}}, 0.02, 10\right]\right\},\right.$

```
  {R, 0, 220}, PlotStyle → {{Red, Thick}},
  PlotRange → {All, {-0.5, 12}}, Frame → {{True, False}, {True, False}},
  BaseStyle → {FontWeight → "Bold", FontColor → Black, FontSize → 12}]
```

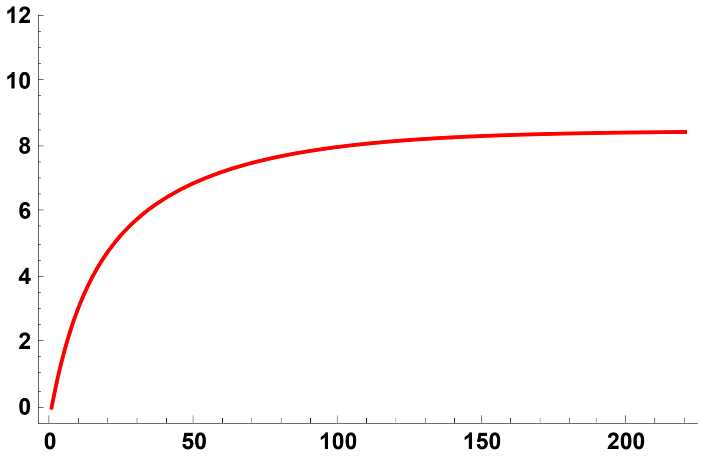

```
SPh01p002s05S = ErrorListPlot[SSh01p002s05S,
  PlotStyle → {Red, PointSize[0.02]}, PlotRange → {{0, 220}, {0, 12}}]
```

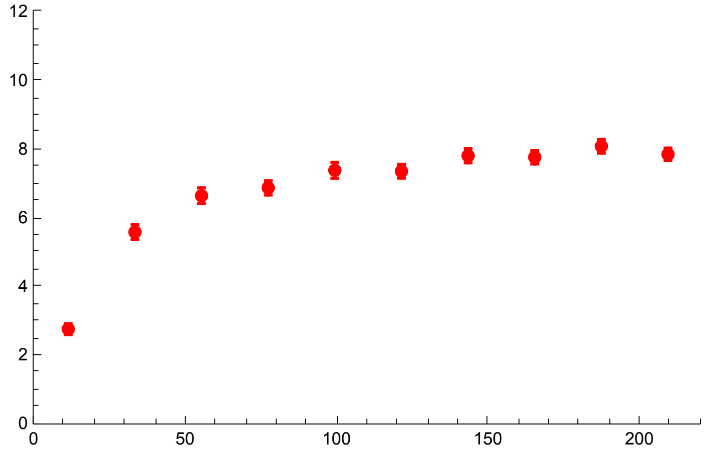


```
Show[SSPh01p002s05, SPh01p002s05S]
```

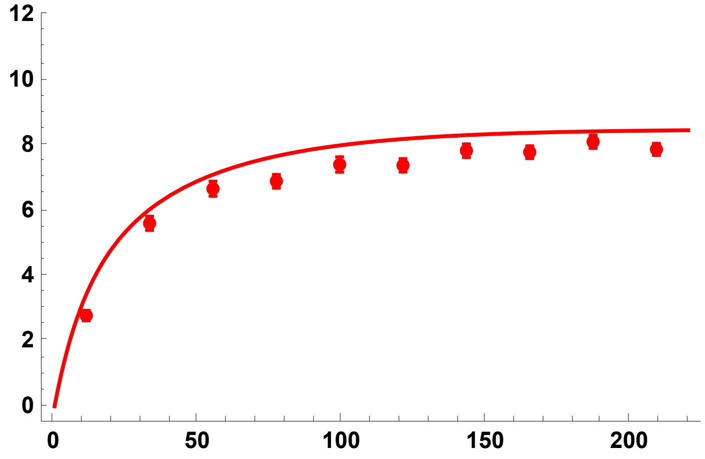


h = 0.9

```
SSh09p002s05 =
  Import["SLIM_F033/StatsProc_R_220_h_0.9_self_0.5_f_0.02_10b_SLIM.dat",
    "Table"][[1]];
SSh09p002s05CIB = Import[
    "SLIM_F033/StatsProc_R_220_h_0.9_self_0.5_f_0.02_10b_SLIM.dat",
    "Table"][[2]];
SSh09p002s05CIT = Import[
    "SLIM_F033/StatsProc_R_220_h_0.9_self_0.5_f_0.02_10b_SLIM.dat",
    "Table"][[3]];
SSh09p002s05S = Partition[Riffle[Partition[Riffle[Rin, SSh09p002s05], 2],
    Map[ErrorBar, Partition[Riffle[-SSh09p002s05CIB, SSh09p002s05CIT], 2]]], 2];
```

```
SSPh09p002s05 = Plot[{ESS2[5000, 0.5, 0.05, 0.9, R, 4/(1 + 1/3), 0.02, 10]},
  {R, 0, 220}, PlotStyle → {{Blue, Thick}},
  PlotRange → {All, {-0.5, 12}}, Frame → {{True, False}, {True, False}},
  BaseStyle → {FontWeight → "Bold", FontColor → Black, FontSize → 12}]
```

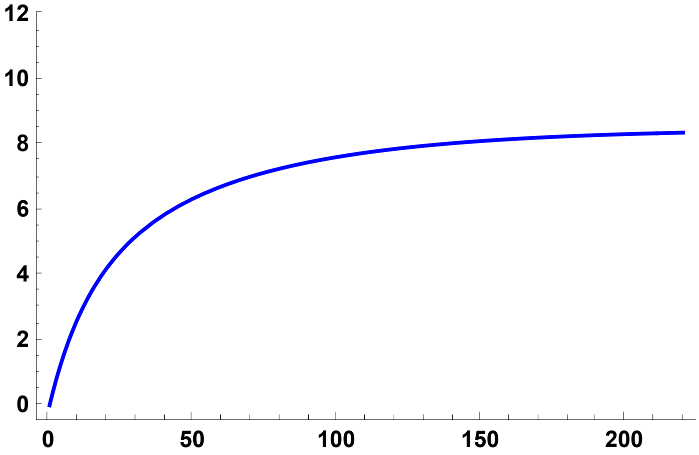


```
SPh09p002s05S = ErrorListPlot[SSh09p002s05S,
  PlotStyle → {Blue, PointSize[0.02]}, PlotRange → {{0, 220}, {0, 12}}]
```

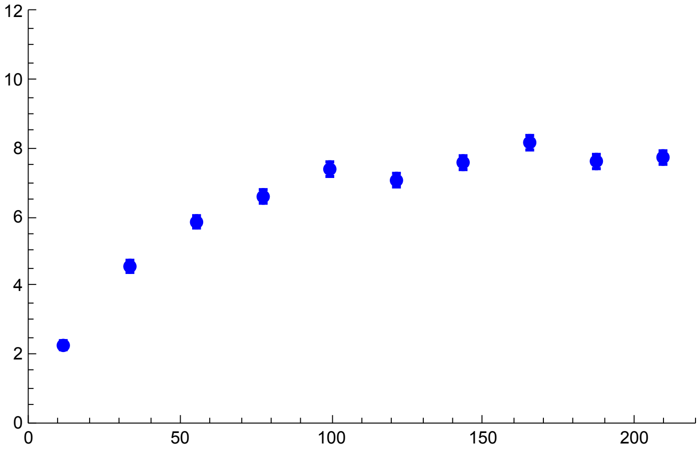


```
Show[SSPh09p002s05, SPh09p002s05S]
```

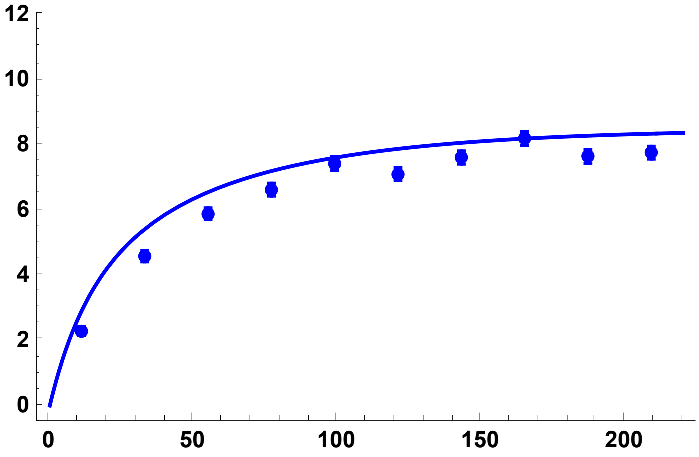

All results together

```
SimSegSitess052pc = Show[SSPh01p002s05, SPh01p002s05S, SSPh05p002s05,
  SPh05p002s05S, SSPh09p002s05, SPh09p002s05S, ImageSize → 200]
```

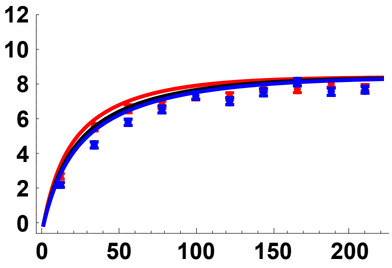


#### Simulation comparisons, $p_0$ = 0.05

h = 0.5

```
SSh05p005s05 =
  Import["SLIM_F033/StatsProc_R_220_h_0.5_self_0.5_f_0.05_10b_SLIM.dat",
    "Table"][[1]];
SSh05p005s05CIB = Import[
    "SLIM_F033/StatsProc_R_220_h_0.5_self_0.5_f_0.05_10b_SLIM.dat",
    "Table"][[2]];
SSh05p005s05CIT = Import[
    "SLIM_F033/StatsProc_R_220_h_0.5_self_0.5_f_0.05_10b_SLIM.dat",
    "Table"][[3]];
SSh05p005s05S = Partition[Riffle[Partition[Riffle[Rin, SSh05p005s05], 2],
    Map[ErrorBar, Partition[Riffle[-SSh05p005s05CIB, SSh05p005s05CIT], 2]]], 2];
```

SSPh05p005s05 = Plot$\left[\left\{ESS2\left[5000, 0.5, 0.05, 0.5, R, \frac{4}{1+\frac{1}{3}}, 0.05, 10\right]\right\},\right.$

```
  {R, 0, 220}, PlotStyle → {{Black, Thick}},
  PlotRange → {All, {-0.5, 12}}, Frame → {{True, False}, {True, False}},
  BaseStyle → {FontWeight → "Bold", FontColor → Black, FontSize → 12}]
```

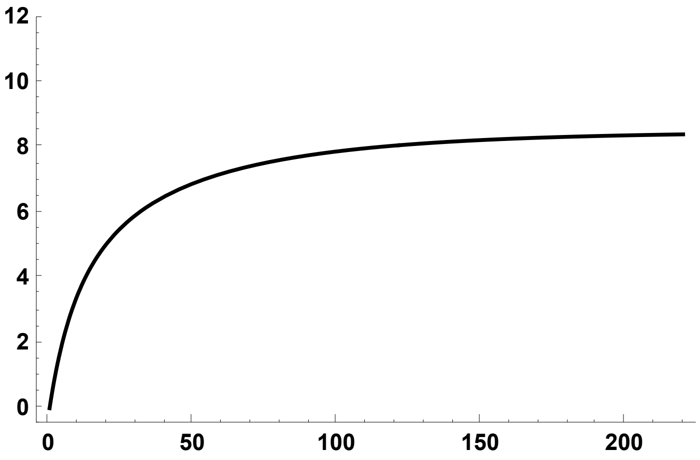

```
SPh05p005s05S = ErrorListPlot[SSh05p005s05S,
  PlotStyle → {Black, PointSize[0.02]}, PlotRange → {{0, 220}, {0, 12}}]
```

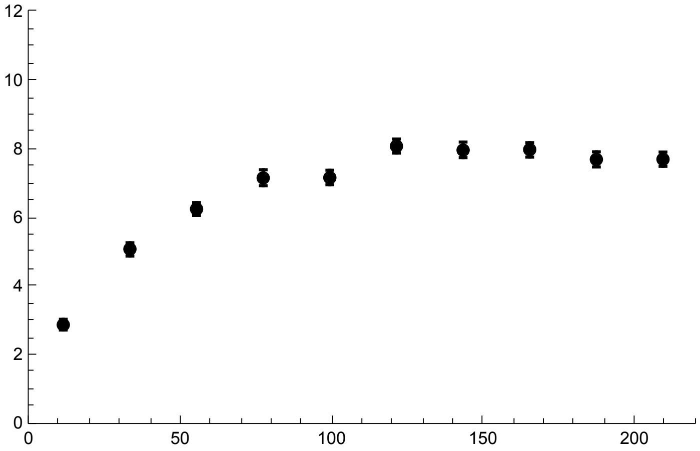


```
Show[SSPh05p005s05, SPh05p005s05S]
```

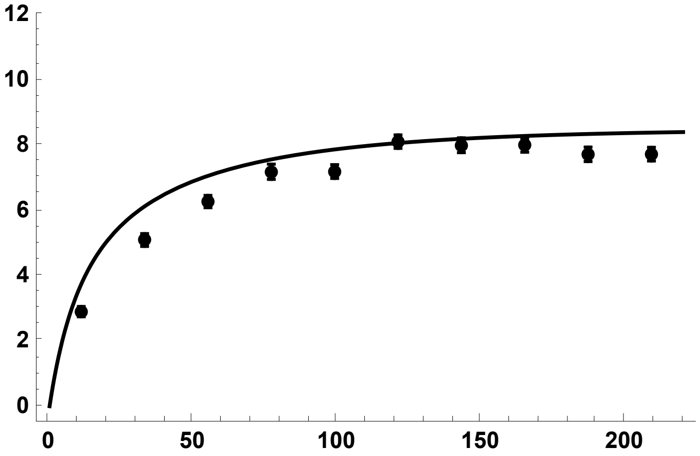


h = 0.1

```
SSh01p005s05 =
  Import["SLIM_F033/StatsProc_R_220_h_0.1_self_0.5_f_0.05_10b_SLIM.dat",
    "Table"][[1]];
SSh01p005s05CIB = Import[
    "SLIM_F033/StatsProc_R_220_h_0.1_self_0.5_f_0.05_10b_SLIM.dat",
    "Table"][[2]];
SSh01p005s05CIT = Import[
    "SLIM_F033/StatsProc_R_220_h_0.1_self_0.5_f_0.05_10b_SLIM.dat",
    "Table"][[3]];
SSh01p005s05S = Partition[Riffle[Partition[Riffle[Rin, SSh01p005s05], 2],
    Map[ErrorBar, Partition[Riffle[-SSh01p005s05CIB, SSh01p005s05CIT], 2]]], 2];
```

```
SSPh01p005s05 = Plot[{ESS2[5000, 0.5, 0.05, 0.1, R, 4/(1 + 1/3), 0.05, 10]},
  {R, 0, 220}, PlotStyle → {{Red, Thick}},
  PlotRange → {All, {-0.5, 12}}, Frame → {{True, False}, {True, False}},
  BaseStyle → {FontWeight → "Bold", FontColor → Black, FontSize → 12}]
```

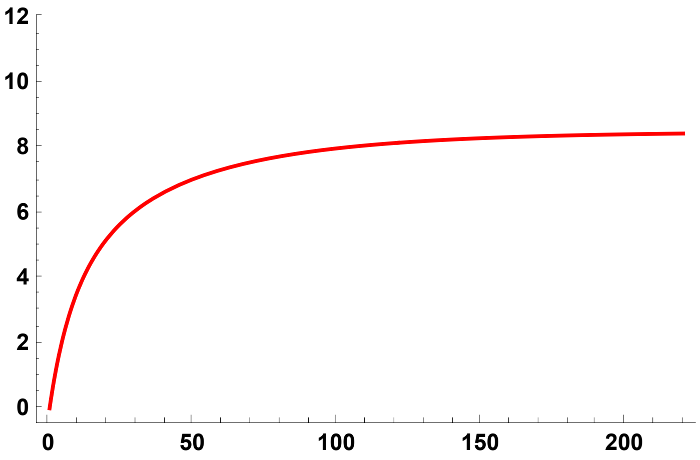


```
SPh01p005s05S = ErrorListPlot[SSh01p005s05S,
  PlotStyle → {Red, PointSize[0.02]}, PlotRange → {{0, 220}, {0, 12}}]
```

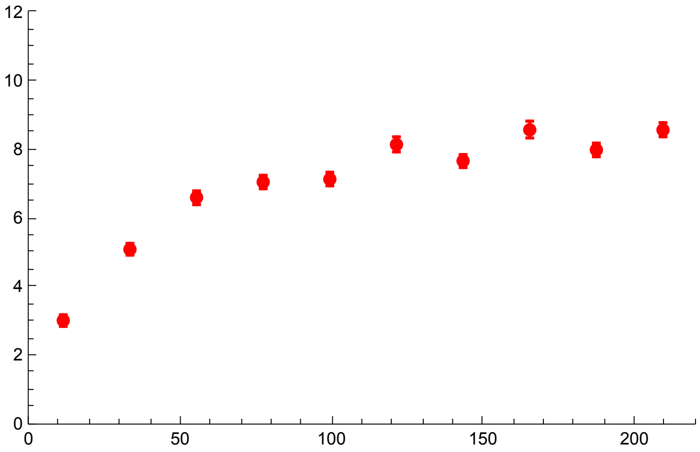


```
Show[SSPh01p005s05, SPh01p005s05S]
```

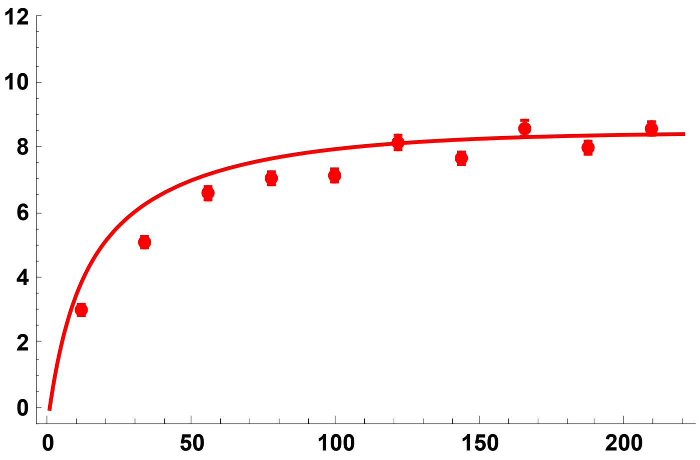

h = 0.9

```
SSh09p005s05 =
  Import["SLIM_F033/StatsProc_R_220_h_0.9_self_0.5_f_0.05_10b_SLIM.dat",
    "Table"][[1]];
SSh09p005s05CIB = Import[
    "SLIM_F033/StatsProc_R_220_h_0.9_self_0.5_f_0.05_10b_SLIM.dat",
    "Table"][[2]];
SSh09p005s05CIT = Import[
    "SLIM_F033/StatsProc_R_220_h_0.9_self_0.5_f_0.05_10b_SLIM.dat",
    "Table"][[3]];
SSh09p005s05S = Partition[Riffle[Partition[Riffle[Rin, SSh09p005s05], 2],
    Map[ErrorBar, Partition[Riffle[-SSh09p005s05CIB, SSh09p005s05CIT], 2]]], 2];
```

```
SSPh09p005s05 = Plot[{ESS2[5000, 0.5, 0.05, 0.9, R, 4/(1 + 1/3), 0.05, 10]},
  {R, 0, 220}, PlotStyle → {{Blue, Thick}},
  PlotRange → {All, {0, 12}}, Frame → {{True, False}, {True, False}},
  BaseStyle → {FontWeight → "Bold", FontColor → Black, FontSize → 12}]
```

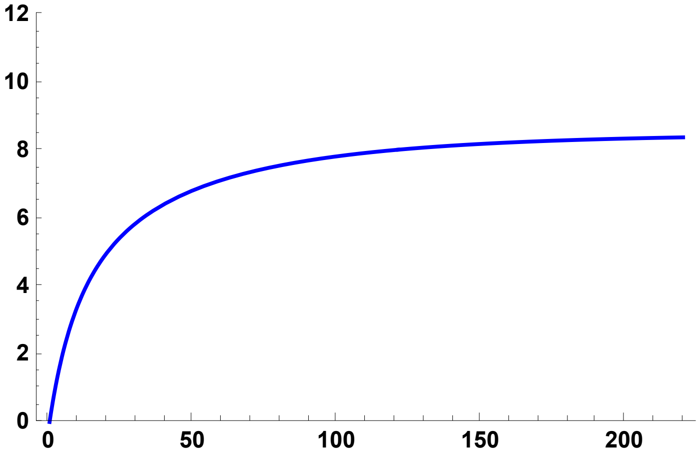


```
SPh09p005s05S = ErrorListPlot[SSh09p005s05S,
  PlotStyle → {Blue, PointSize[0.02]}, PlotRange → {{0, 220}, {0, 12}}]
```

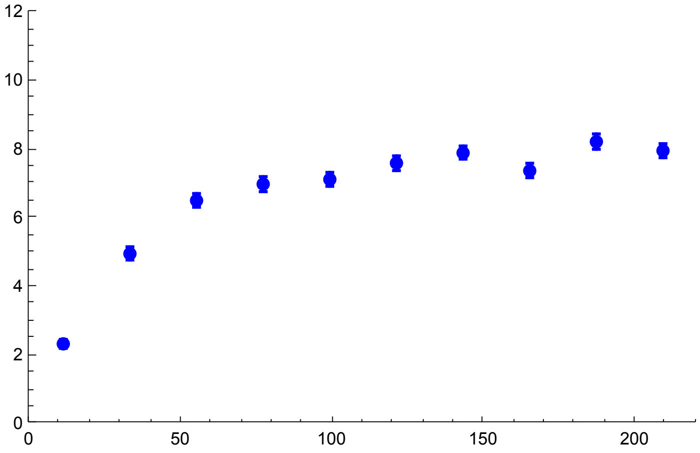

```
Show[SSPh09p005s05, SPh09p005s05S]
```

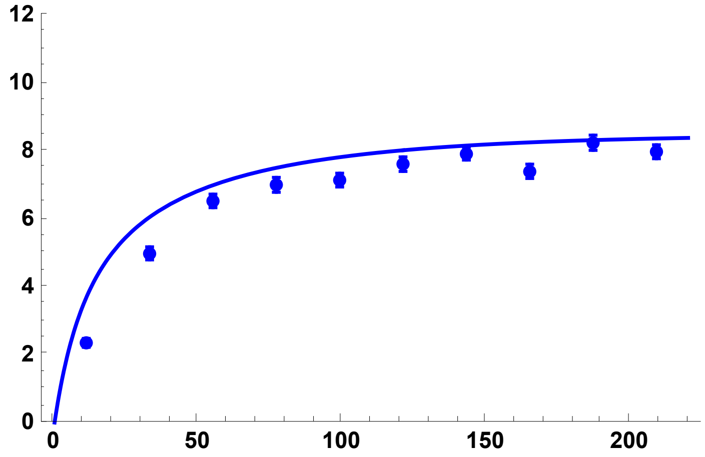


All results together

```
SimSegSitess055pc = Show[SSPh01p005s05, SPh01p005s05S, SSPh05p005s05,
  SPh05p005s05S, SSPh09p005s05, SPh09p005s05S, ImageSize → 200]
```

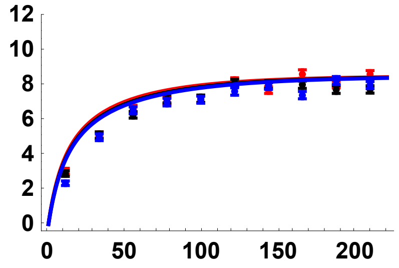


#### $\sigma = 0.95$ ($F \simeq 0.904$)

Preliminaries

```
SetDirectory[NotebookDirectory[]];
Rin = Table[100 + 200 * i, {i, 0, 9}];
```

##### Simulation comparisons, from initial frequency $p_0 = 1/2N$

h = 0.5

```
SSh05s095 =
  Import["SLIM_F09/StatsProc_R_2000_h_0.5_self_0.95_f_1e-04_10b_SLIM.dat",
    "Table"][[1]];
SSh05s095CIB = Import[
    "SLIM_F09/StatsProc_R_2000_h_0.5_self_0.95_f_1e-04_10b_SLIM.dat",
    "Table"][[2]];
SSh05s095CIT = Import[
    "SLIM_F09/StatsProc_R_2000_h_0.5_self_0.95_f_1e-04_10b_SLIM.dat",
    "Table"][[3]];
SSh05s095S = Partition[Riffle[Partition[Riffle[Rin, SSh05s095], 2],
    Map[ErrorBar, Partition[Riffle[-SSh05s095CIB, SSh05s095CIT], 2]]], 2];
```

```
Sh05s095 = Plot[
  {ESS2[5000, 0.95, 0.05, 0.5, R, 4/(1 + 0.95/(2-0.95)), Boostp0[5000, 0.05, 0.5, 0.95], 10]},
  {R, 0, 2000}, PlotStyle → {{Black, Thick}},
  PlotRange → {All, {-0.5, 12}}, Frame → {{True, False}, {True, False}},
  BaseStyle → {FontWeight → "Bold", FontColor → Black, FontSize → 12}]
```

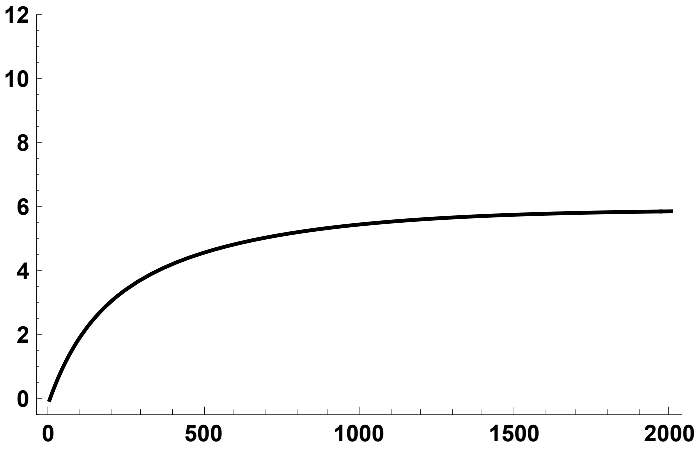

```
Sh05s095S = ErrorListPlot[SSh05s095S,
  PlotStyle → {Black, PointSize[0.02]}, PlotRange → {{0, 2000}, {0, 12}}]
```

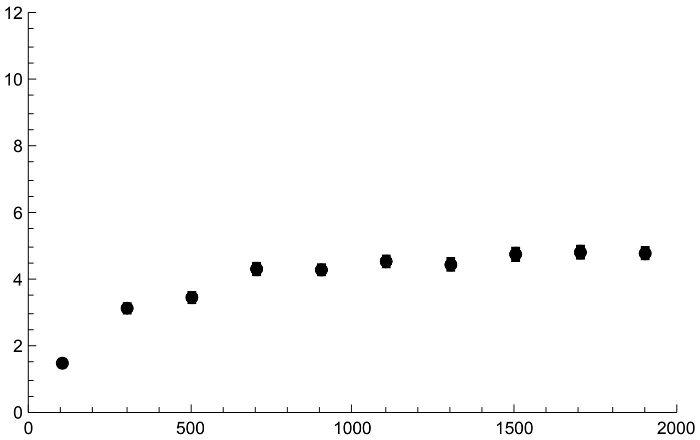

```
Show[Sh05s095, Sh05s095S]
```

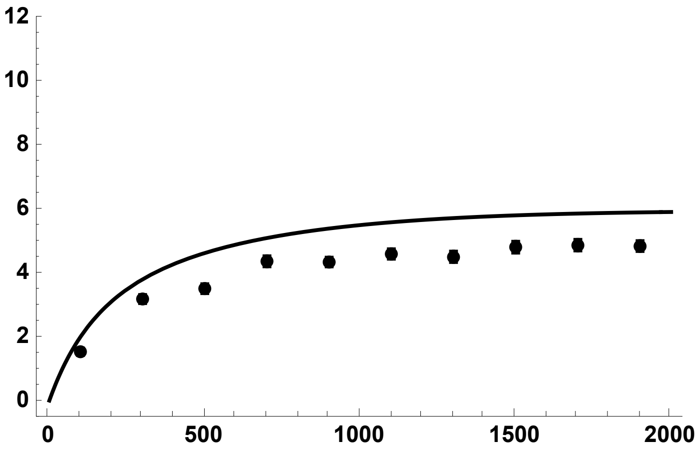

h = 0.1

```
SSh01s095 =
  Import["SLIM_F09/StatsProc_R_2000_h_0.1_self_0.95_f_1e-04_10b_SLIM.dat",
    "Table"][[1]];
SSh01s095CIB = Import[
    "SLIM_F09/StatsProc_R_2000_h_0.1_self_0.95_f_1e-04_10b_SLIM.dat",
    "Table"][[2]];
SSh01s095CIT = Import[
    "SLIM_F09/StatsProc_R_2000_h_0.1_self_0.95_f_1e-04_10b_SLIM.dat",
    "Table"][[3]];
SSh01s095S = Partition[Riffle[Partition[Riffle[Rin, SSh01s095], 2],
    Map[ErrorBar, Partition[Riffle[-SSh01s095CIB, SSh01s095CIT], 2]]], 2];
```

```
Sh01s095 = Plot[
  {ESS2[5000, 0.95, 0.05, 0.1, R, 4/(1 + 0.95/(2-0.95)), Boostp0[5000, 0.05, 0.1, 0.95], 10]},
  {R, 0, 2000},
  PlotStyle → {{Red, Thick}, {Red, Thick, Dashed}, {Red, Thick, Dashing[Large]}},
  PlotRange → {All, {-0.5, 12}}, Frame → {{True, False}, {True, False}},
  Frame → {{True, False}, {True, False}},
  BaseStyle → {FontWeight → "Bold", FontColor → Black, FontSize → 12}]
```

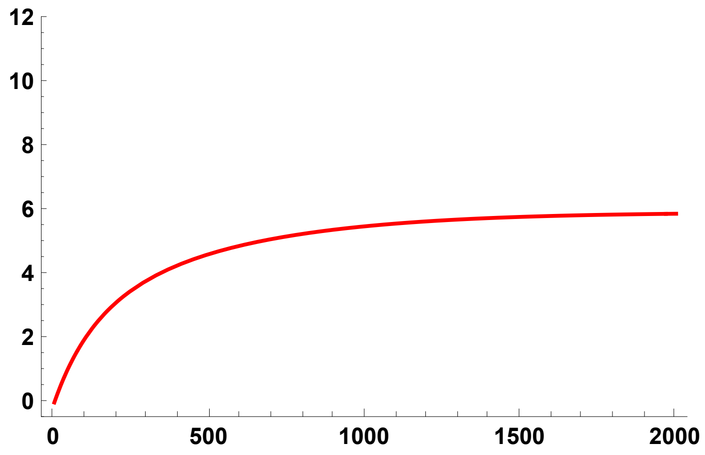

```
Sh01s095S = ErrorListPlot[SSh01s095S,
  PlotStyle → {Red, PointSize[0.02]}, PlotRange → {{0, 2000}, {0, 12}}]
```

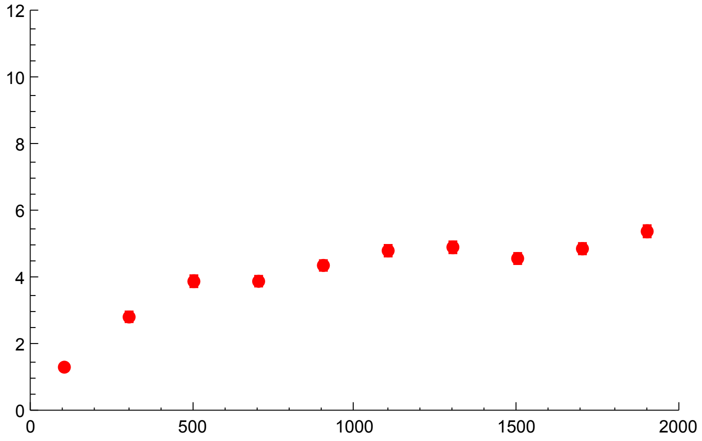


```
Show[Sh01s095, Sh01s095S]
```

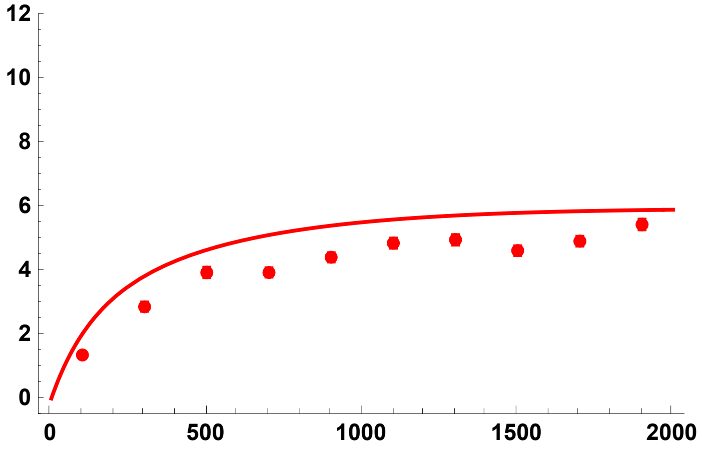


h = 0.9

```
SSh09s095 =
  Import["SLIM_F09/StatsProc_R_2000_h_0.9_self_0.95_f_1e-04_10b_SLIM.dat",
    "Table"][[1]];
SSh09s095CIB = Import[
    "SLIM_F09/StatsProc_R_2000_h_0.9_self_0.95_f_1e-04_10b_SLIM.dat",
    "Table"][[2]];
SSh09s095CIT = Import[
    "SLIM_F09/StatsProc_R_2000_h_0.9_self_0.95_f_1e-04_10b_SLIM.dat",
    "Table"][[3]];
SSh09s095S = Partition[Riffle[Partition[Riffle[Rin, SSh09s095], 2],
    Map[ErrorBar, Partition[Riffle[-SSh09s095CIB, SSh09s095CIT], 2]]], 2];
```

```
Sh09s095 = Plot[
  {ESS2[5000, 0.95, 0.05, 0.9, R, 4/(1 + 0.95/(2-0.95)), Boostp0[5000, 0.05, 0.9, 0.95], 10]},
  {R, 0, 2000}, PlotStyle →
   {{Blue, Thick}, {Blue, Thick, Dashed}, {Blue, Thick, Dashing[Large]}},
  PlotRange → {All, {-0.5, 12}}, Frame → {{True, False}, {True, False}},
  BaseStyle → {FontWeight → "Bold", FontColor → Black, FontSize → 12}]
```

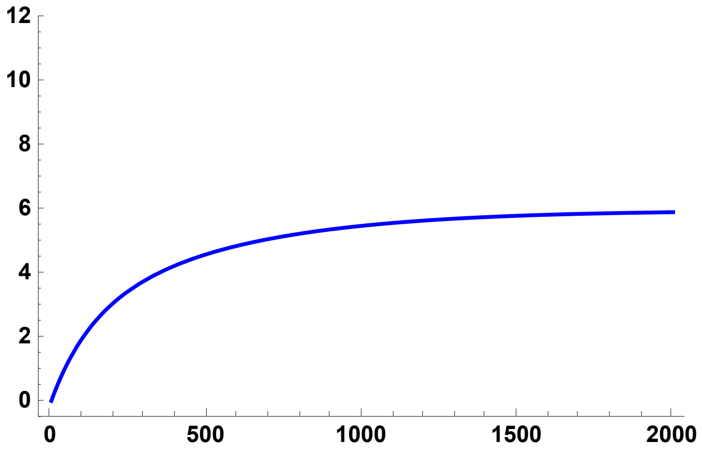

```
Sh09s095S = ErrorListPlot[SSh09s095S,
  PlotStyle → {Blue, PointSize[0.02]}, PlotRange → {{0, 2000}, {0, 12}}]
```

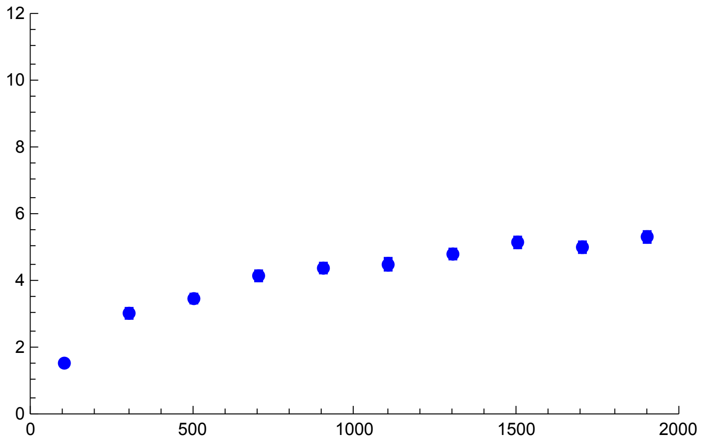

```
Show[Sh09s095, Sh09s095S]
```

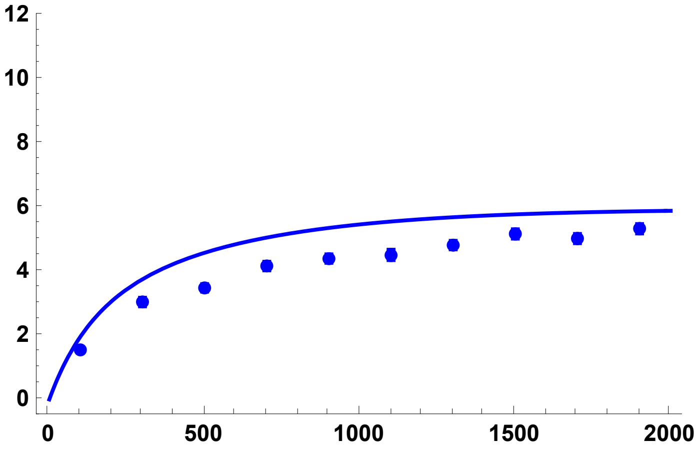


All results together

```
SimSegSitess095DN = Show[Sh01s095, Sh01s095S,
  Sh05s095, Sh05s095S, Sh09s095, Sh09s095S, ImageSize → 200]
```

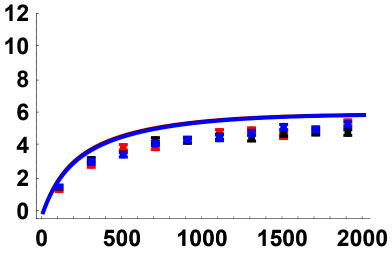


#### Simulation comparisons, $p_0$ = 0.02

h = 0.5

```
SSh05p002s095 =
  Import["SLIM_F09/StatsProc_R_2000_h_0.9_self_0.95_f_0.02_10b_SLIM.dat",
    "Table"][[1]];
SSh05p002s095CIB = Import[
    "SLIM_F09/StatsProc_R_2000_h_0.9_self_0.95_f_0.02_10b_SLIM.dat",
    "Table"][[2]];
SSh05p002s095CIT = Import[
    "SLIM_F09/StatsProc_R_2000_h_0.9_self_0.95_f_0.02_10b_SLIM.dat",
    "Table"][[3]];
SSh05p002s095S = Partition[Riffle[Partition[Riffle[Rin, SSh05p002s095], 2], Map[
     ErrorBar, Partition[Riffle[-SSh05p002s095CIB, SSh05p002s095CIT], 2]]], 2];
```

```
SSPh05p002s095 = Plot[{ESS2[5000, 0.95, 0.05, 0.5, R, 4/(1 + 0.95/(2-0.95)), 0.02, 10]},
  {R, 0, 2000}, PlotStyle → {{Black, Thick}},
  PlotRange → {All, {-0.5, 12}}, Frame → {{True, False}, {True, False}},
  BaseStyle → {FontWeight → "Bold", FontColor → Black, FontSize → 12}]
```

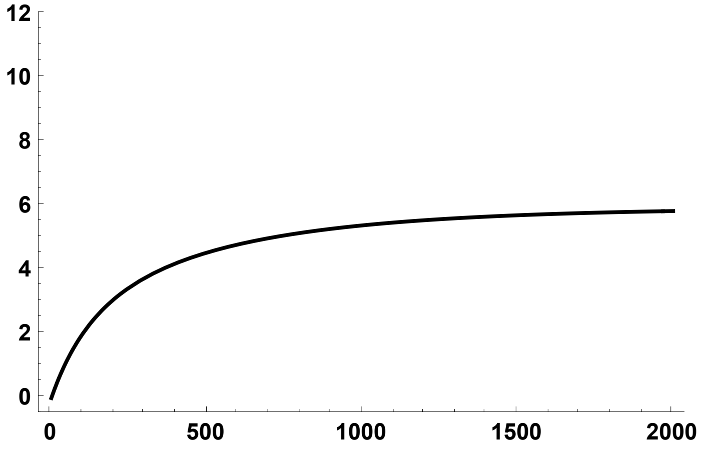


```
SPh05p002s095S = ErrorListPlot[SSh05p002s095S,
  PlotStyle → {Black, PointSize[0.02]}, PlotRange → {{0, 2000}, {0, 12}}]
```

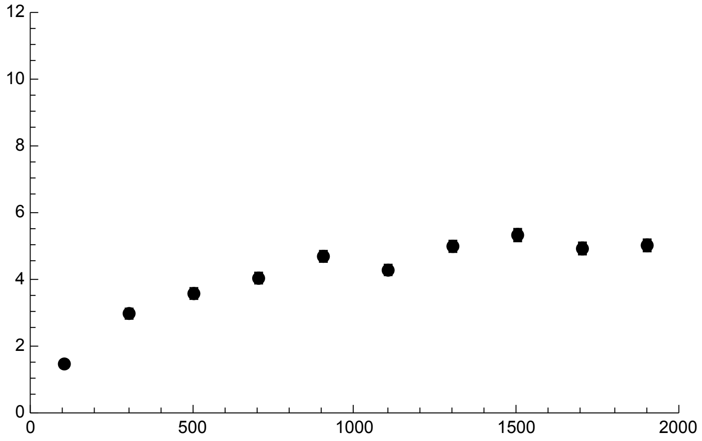


```
Show[SSPh05p002s095, SPh05p002s095S]
```

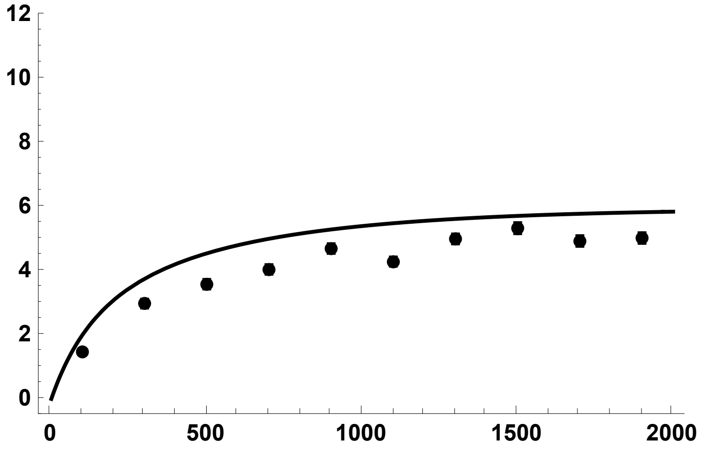


h = 0.1

```
SSh01p002s095 =
  Import["SLIM_F09/StatsProc_R_2000_h_0.1_self_0.95_f_0.02_10b_SLIM.dat",
    "Table"][[1]];
SSh01p002s095CIB = Import[
    "SLIM_F09/StatsProc_R_2000_h_0.1_self_0.95_f_0.02_10b_SLIM.dat",
    "Table"][[2]];
SSh01p002s095CIT = Import[
    "SLIM_F09/StatsProc_R_2000_h_0.1_self_0.95_f_0.02_10b_SLIM.dat",
    "Table"][[3]];
SSh01p002s095S = Partition[Riffle[Partition[Riffle[Rin, SSh01p002s095], 2], Map[
     ErrorBar, Partition[Riffle[-SSh01p002s095CIB, SSh01p002s095CIT], 2]]], 2];
```

$$SSPh01p002s095 = Plot\left[\left\{ESS2\left[5000, 0.95, 0.05, 0.1, R, \frac{4}{1+\frac{0.95}{2-0.95}}, 0.02, 10\right]\right\},\right.$$

```
  {R, 0, 2000}, PlotStyle → {{Red, Thick}},
  PlotRange → {All, {-0.5, 12}}, Frame → {{True, False}, {True, False}},
  BaseStyle → {FontWeight → "Bold", FontColor → Black, FontSize → 12}]
```

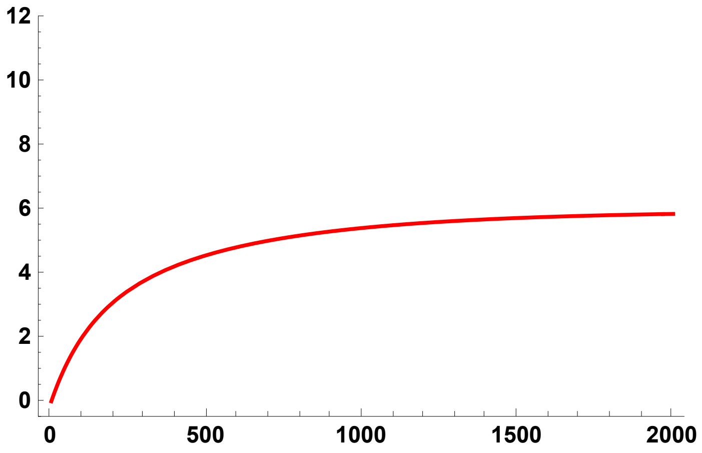


```
SPh01p002s095S = ErrorListPlot[SSh01p002s095S,
  PlotStyle → {Red, PointSize[0.02]}, PlotRange → {{0, 2000}, {0, 12}}]
```

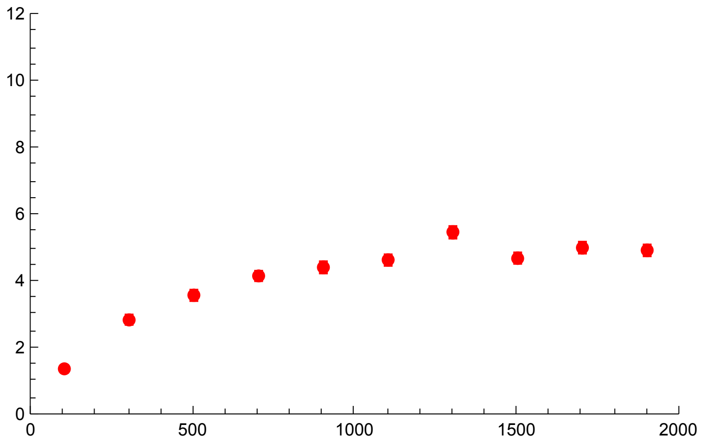

```
Show[SSPh01p002s095, SPh01p002s095S]
```

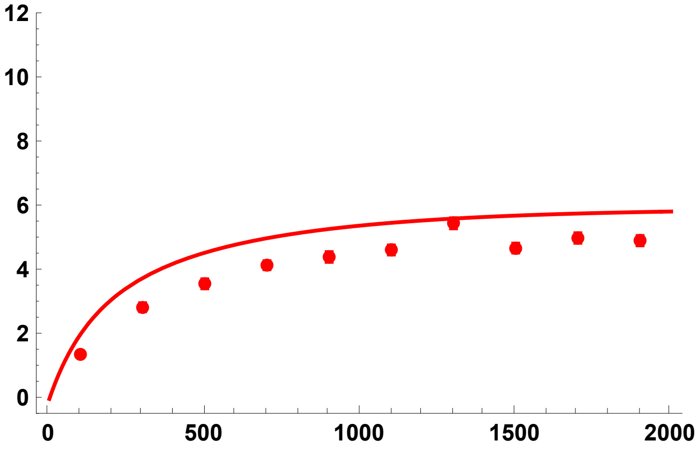


h = 0.9

```
SSh09p002s095 =
  Import["SLIM_F09/StatsProc_R_2000_h_0.9_self_0.95_f_0.02_10b_SLIM.dat",
    "Table"][[1]];
SSh09p002s095CIB = Import[
    "SLIM_F09/StatsProc_R_2000_h_0.9_self_0.95_f_0.02_10b_SLIM.dat",
    "Table"][[2]];
SSh09p002s095CIT = Import[
    "SLIM_F09/StatsProc_R_2000_h_0.9_self_0.95_f_0.02_10b_SLIM.dat",
    "Table"][[3]];
SSh09p002s095S = Partition[Riffle[Partition[Riffle[Rin, SSh09p002s095], 2], Map[
     ErrorBar, Partition[Riffle[-SSh09p002s095CIB, SSh09p002s095CIT], 2]]], 2];
```

SSPh09p002s095 = Plot[{ESS2[5000, 0.95, 0.05, 0.9, R, $\frac{4}{1+\frac{0.95}{2-0.95}}$, 0.02, 10]},

```
  {R, 0, 2000}, PlotStyle → {{Blue, Thick}},
  PlotRange → {All, {-0.5, 12}}, Frame → {{True, False}, {True, False}},
  BaseStyle → {FontWeight → "Bold", FontColor → Black, FontSize → 12}]
```

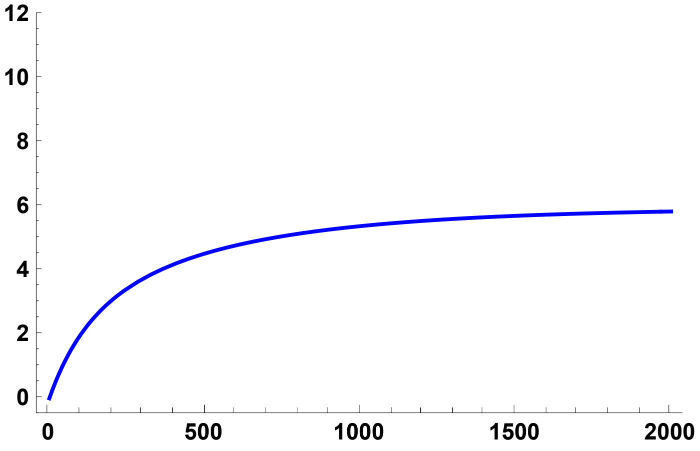

```
SPh09p002s095S = ErrorListPlot[SSh09p002s095S,
  PlotStyle → {Blue, PointSize[0.02]}, PlotRange → {{0, 2000}, {0, 12}}]
```

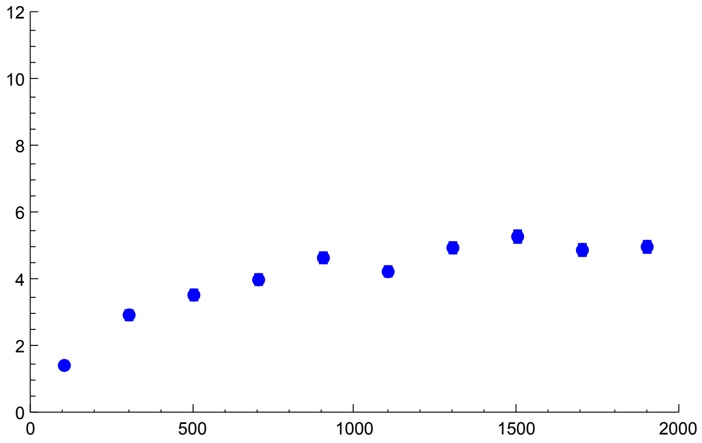


```
Show[SSPh09p002s095, SPh09p002s095S]
```

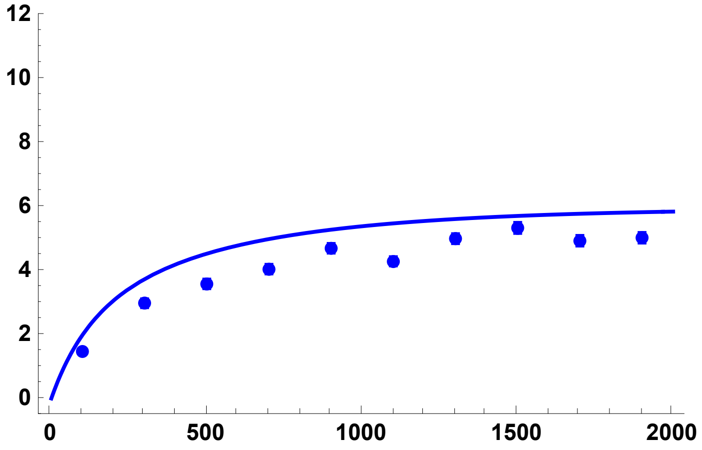


All results together

```
SimSegSitess0952pc = Show[SSPh01p002s095, SPh01p002s095S, SSPh05p002s095,
  SPh05p002s095S, SSPh09p002s095, SPh09p002s095S, ImageSize → 200]
```

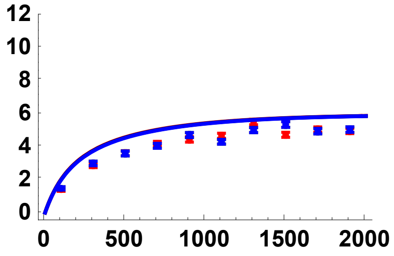


#### Simulation comparisons, $p_0 = 0.05$

h = 0.5

```
SSh05p005s095 =
  Import["SLIM_F09/StatsProc_R_2000_h_0.5_self_0.95_f_0.05_10b_SLIM.dat",
    "Table"][[1]];
SSh05p005s095CIB = Import[
    "SLIM_F09/StatsProc_R_2000_h_0.5_self_0.95_f_0.05_10b_SLIM.dat",
    "Table"][[2]];
SSh05p005s095CIT = Import[
    "SLIM_F09/StatsProc_R_2000_h_0.5_self_0.95_f_0.05_10b_SLIM.dat",
    "Table"][[3]];
SSh05p005s095S = Partition[Riffle[Partition[Riffle[Rin, SSh05p005s095], 2], Map[
     ErrorBar, Partition[Riffle[-SSh05p005s095CIB, SSh05p005s095CIT], 2]]], 2];
```

SSPh05p005s095 = Plot[{ESS2[5000, 0.95, 0.05, 0.5, R, $\frac{4}{1+\frac{0.95}{2-0.95}}$, 0.05, 10]},

```
  {R, 0, 2000}, PlotStyle → {{Black, Thick}},
  PlotRange → {All, {-0.5, 12}}, Frame → {{True, False}, {True, False}},
  BaseStyle → {FontWeight → "Bold", FontColor → Black, FontSize → 12}]
```

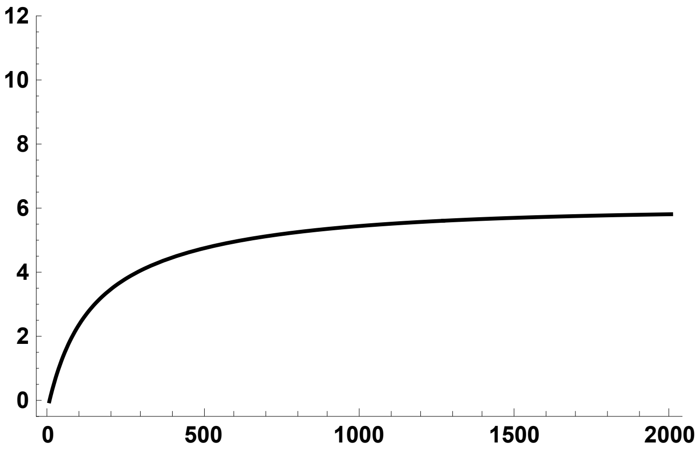

```
SPh05p005s095S = ErrorListPlot[SSh05p005s095S,
  PlotStyle → {Black, PointSize[0.02]}, PlotRange → {{0, 2000}, {0, 12}}]
```

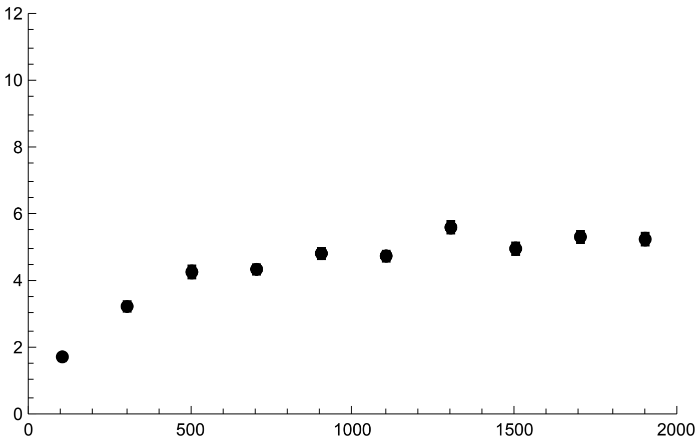

```
Show[SSPh05p005s095, SPh05p005s095S]
```

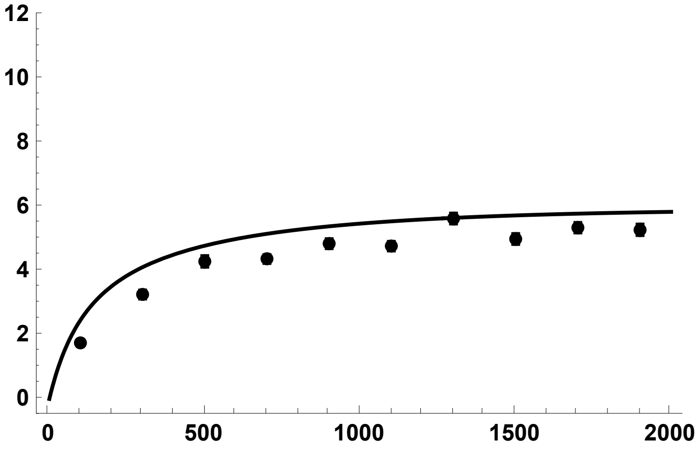


h = 0.1

```
SSh01p005s095 =
  Import["SLIM_F09/StatsProc_R_2000_h_0.1_self_0.95_f_0.05_10b_SLIM.dat",
    "Table"][[1]];
SSh01p005s095CIB = Import[
    "SLIM_F09/StatsProc_R_2000_h_0.1_self_0.95_f_0.05_10b_SLIM.dat",
    "Table"][[2]];
SSh01p005s095CIT = Import[
    "SLIM_F09/StatsProc_R_2000_h_0.1_self_0.95_f_0.05_10b_SLIM.dat",
    "Table"][[3]];
SSh01p005s095S = Partition[Riffle[Partition[Riffle[Rin, SSh01p005s095], 2], Map[
     ErrorBar, Partition[Riffle[-SSh01p005s095CIB, SSh01p005s095CIT], 2]]], 2];
```

SSPh01p005s095 = Plot[{ESS2[5000, 0.95, 0.05, 0.1, R, $\frac{4}{1+\frac{0.95}{2-0.95}}$, 0.05, 10]},

```
  {R, 0, 2000}, PlotStyle → {{Red, Thick}},
  PlotRange → {All, {-0.5, 12}}, Frame → {{True, False}, {True, False}},
  BaseStyle → {FontWeight → "Bold", FontColor → Black, FontSize → 12}]
```

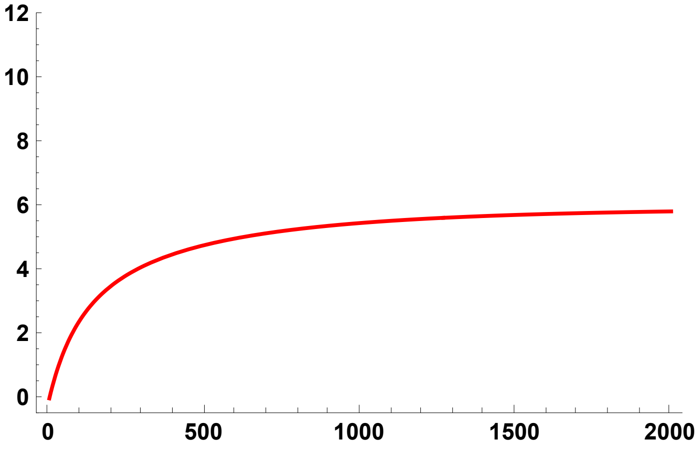

```
SPh01p005s095S = ErrorListPlot[SSh01p005s095S,
  PlotStyle → {Red, PointSize[0.02]}, PlotRange → {{0, 2000}, {0, 12}}]
```

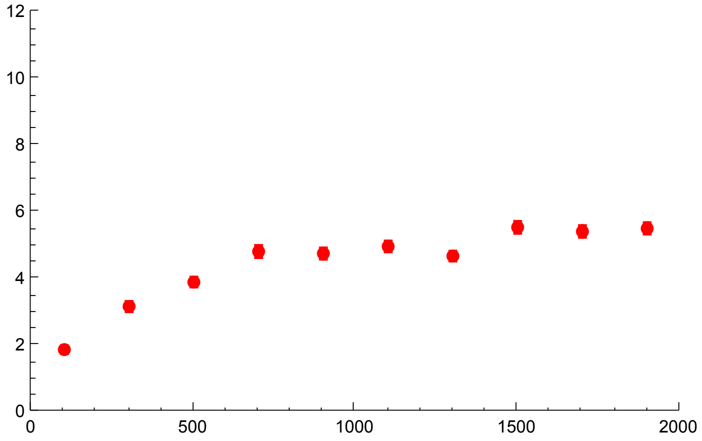


```
Show[SSPh01p005s095, SPh01p005s095S]
```

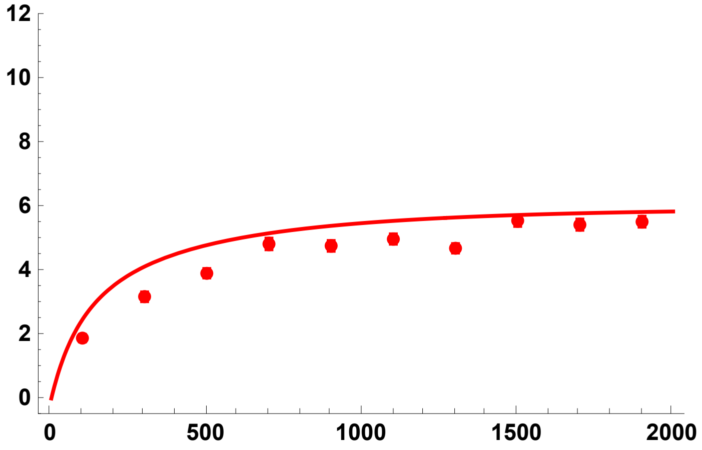


h = 0.9

```
SSh09p005s095 =
  Import["SLIM_F09/StatsProc_R_2000_h_0.9_self_0.95_f_0.05_10b_SLIM.dat",
    "Table"][[1]];
SSh09p005s095CIB = Import[
    "SLIM_F09/StatsProc_R_2000_h_0.9_self_0.95_f_0.05_10b_SLIM.dat",
    "Table"][[2]];
SSh09p005s095CIT = Import[
    "SLIM_F09/StatsProc_R_2000_h_0.9_self_0.95_f_0.05_10b_SLIM.dat",
    "Table"][[3]];
SSh09p005s095S = Partition[Riffle[Partition[Riffle[Rin, SSh09p005s095], 2], Map[
      ErrorBar, Partition[Riffle[-SSh09p005s095CIB, SSh09p005s095CIT], 2]]], 2];
```

```
SSPh09p005s095 = Plot[{ESS2[5000, 0.95, 0.05, 0.9, R, 4/(1 + 0.95/(2-0.95)), 0.05, 10]},
  {R, 0, 2000}, PlotStyle → {{Blue, Thick}},
  PlotRange → {All, {-0.5, 12}}, Frame → {{True, False}, {True, False}},
  BaseStyle → {FontWeight → "Bold", FontColor → Black, FontSize → 12}]
```

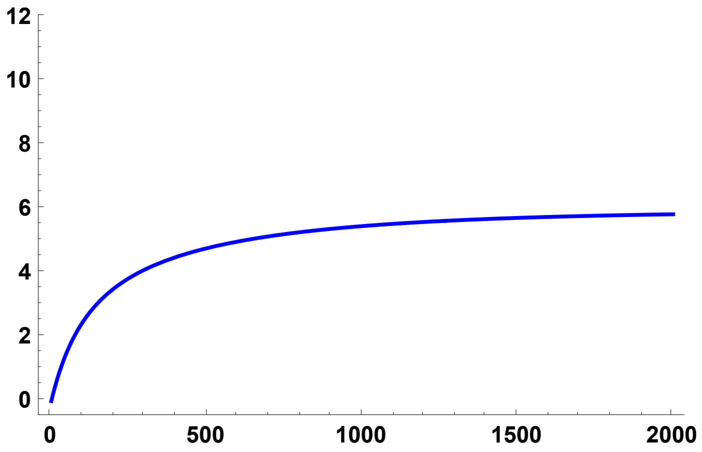

```
SPh09p005s095S = ErrorListPlot[SSh09p005s095S,
  PlotStyle → {Blue, PointSize[0.02]}, PlotRange → {{0, 2000}, {0, 12}}]
```

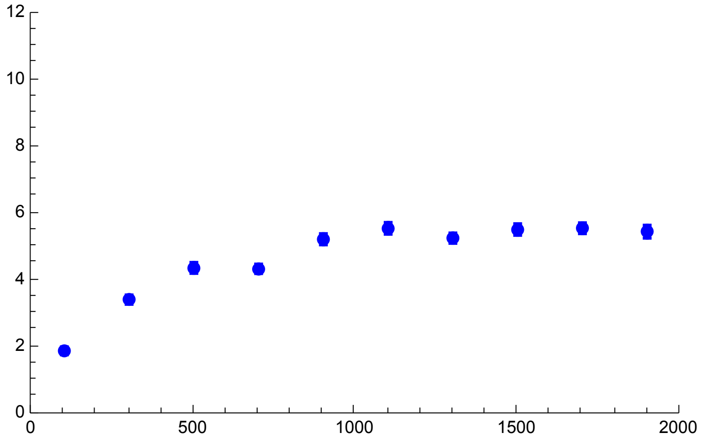

```
Show[SSPh09p005s095, SPh09p005s095S]
```

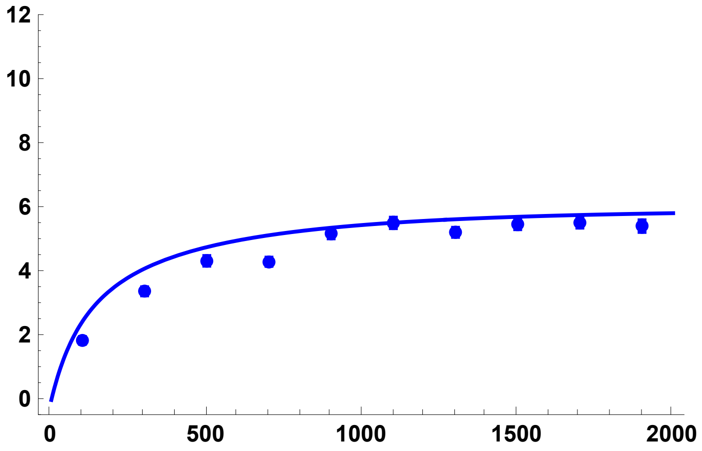

All results together

```
SimSegSitess0955pc = Show[SSPh01p005s095, SPh01p005s095S, SSPh05p005s095,
  SPh05p005s095S, SSPh09p005s095, SPh09p005s095S, ImageSize → 200]
```

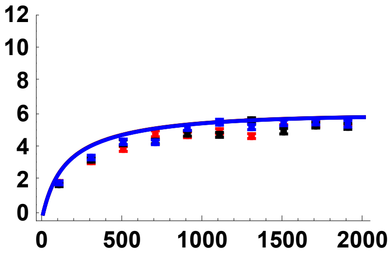

### Grid of key results

#### Comparing analytical results and SLiM simulations

```
SimCompSegSites = Labeled[
  Grid[{{Text@TraditionalForm@Style["p0 = 1/2N", 24], Text@TraditionalForm@
       Style["p0 = 0.02", 24], Text@TraditionalForm@Style["p0 = 0.05", 24],},
    {SimSegSitess0DN, SimSegSitess02pc, SimSegSitess05pc,
     Text@TraditionalForm@Style["σ = 0.00\n(F = 0.00)", 24]},
    {SimSegSitess05DN, SimSegSitess052pc, SimSegSitess055pc,
     Text@TraditionalForm@Style["σ = 0.50\n(F ≃ 0.33)", 24]},
    {SimSegSitess095DN, SimSegSitess0952pc, SimSegSitess0955pc, Text@
      TraditionalForm@Style["σ = 0.95\n(F ≃ 0.90)", 24]}}, Spacings → {2, 1}],
  {Text@TraditionalForm@Style["Number of\nSegregating\nSites", 24],
   Text@TraditionalForm@Style["Scaled Recombination Rate, 2Nr", 24]}, {Left,
   Bottom}]
```

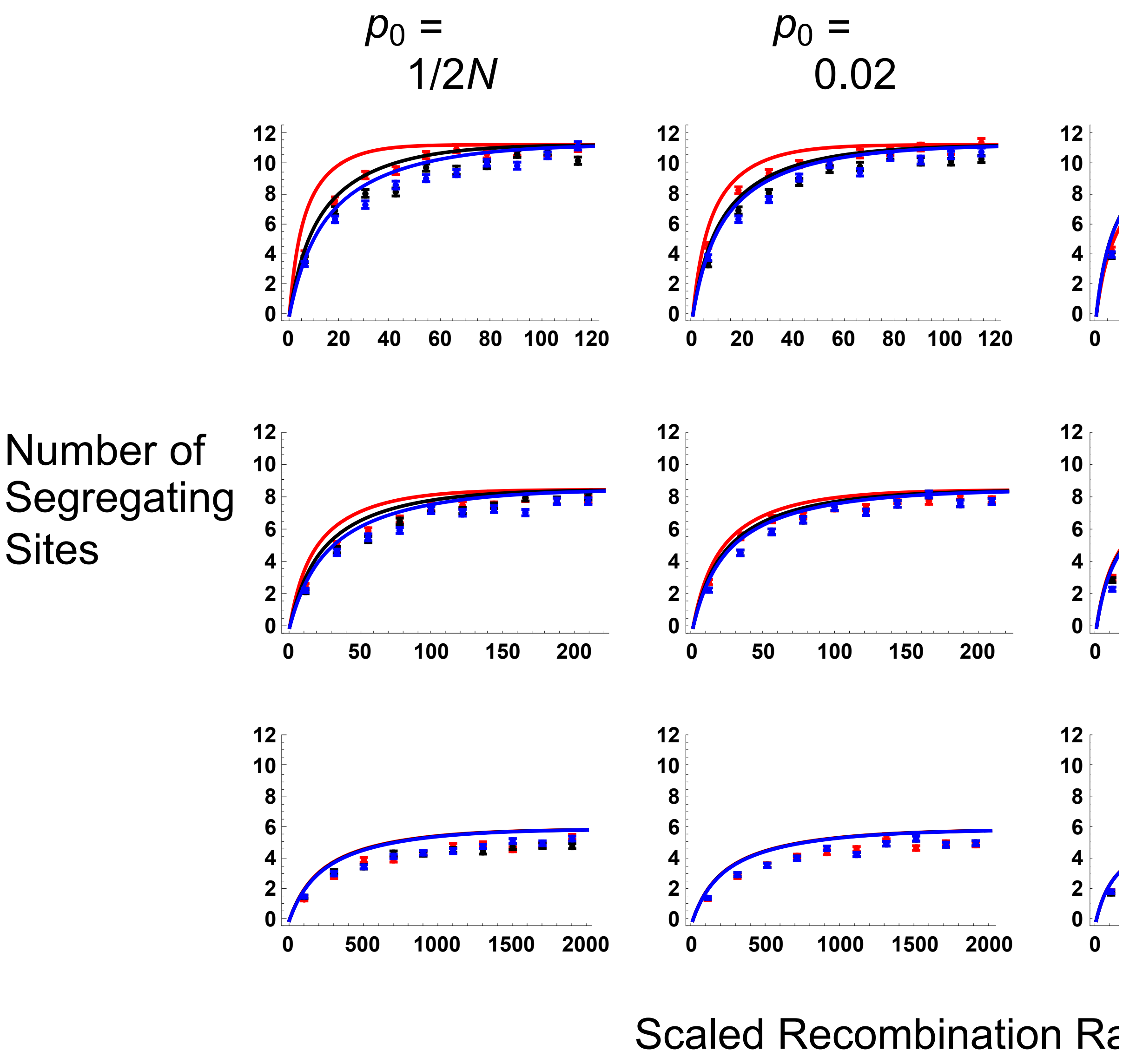

#### Comparing SLiM and MSMS simulations ($\sigma = 0$)

```
SLiMandMSMSResSegSites = Labeled[
  Grid[{{Text@TraditionalForm@Style["h = 0.1", 24, TextAlignment → Center],
     Text@TraditionalForm@Style["h = 0.5", 24], Text@
      TraditionalForm@Style["h = 0.9", 24],}, {Text@TraditionalForm@Style[
        "Red points: forward-in-time\nPink points: coalescent simulations",
        14, TextAlignment → Center], Text@TraditionalForm@Style[
        "Black points: forward-in-time\nGrey points: coalescent simulations",
        14, TextAlignment → Center], Text@TraditionalForm@Style[
        "Blue points: forward-in-time\nCyan points: coalescent simulations",
        14, TextAlignment → Center],},
    {h01MSMSSegsites, h05MSMSSegsites, h09MSMSSegsites,
     Text@TraditionalForm@Style["p0 = 1/2N", 24]},
    {h01MSMS2pcSegsites, h05MSMS2pcSegsites, h09MSMS2pcSegsites,
     Text@TraditionalForm@Style["p0 = 0.02", 24]},
    {h01MSMS5pcSegSites, h05MSMS5pcSegSites, h09MSMS5pcSegSites,
     Text@TraditionalForm@Style["p0 = 0.05", 24]}}, Spacings → {2, 1}],
  {Text@TraditionalForm@Style["Number of\nSegregating\nSites", 24],
   Text@TraditionalForm@Style["Scaled Recombination Rate, 2Nr", 24]}, {Left,
   Bottom}]
```

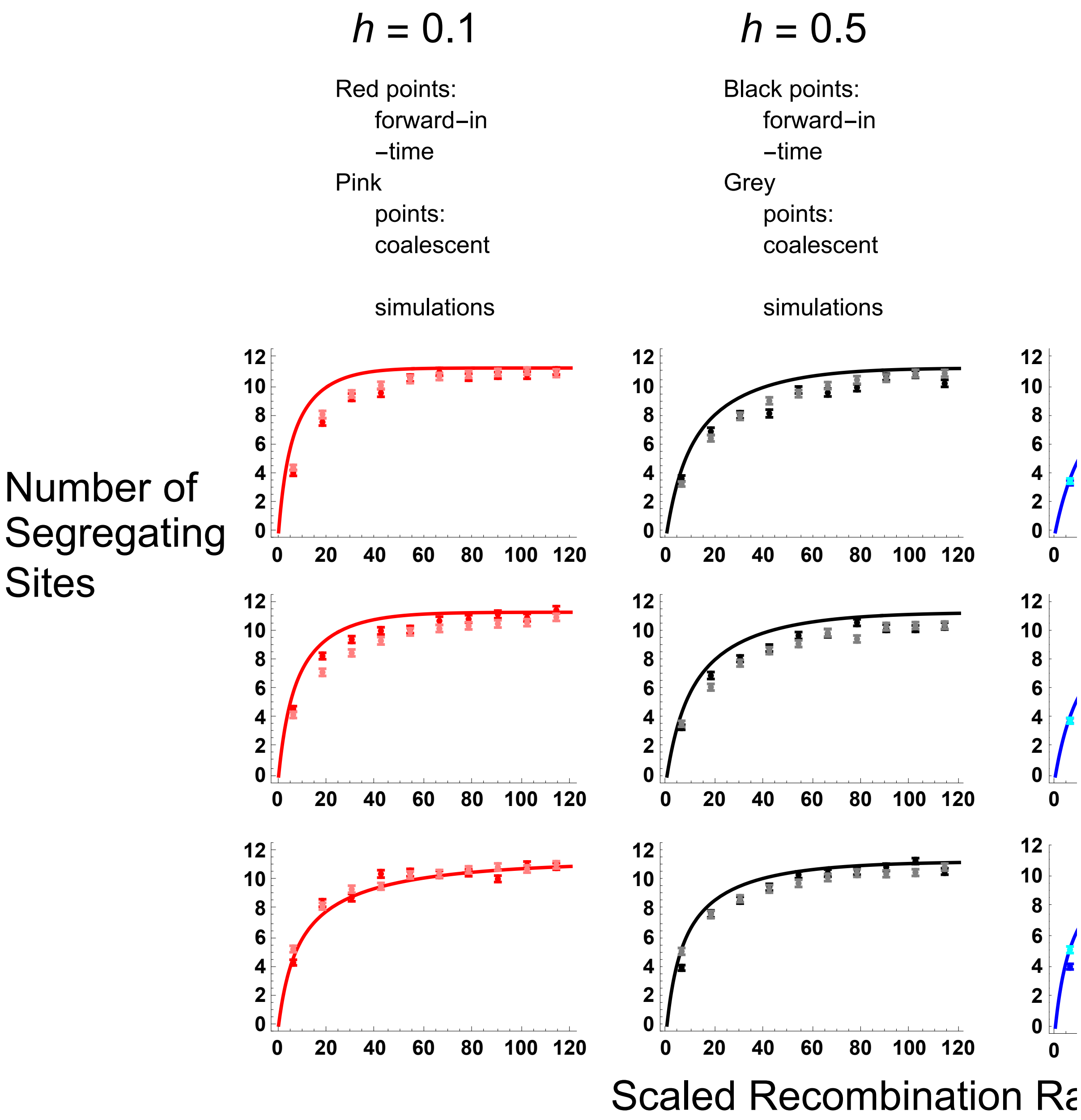


### Section E: Simulation comparisons, Site Frequency Spectrum

Setting working directory (to enable reading in of files)

```
SetDirectory[NotebookDirectory[]];
```

#### Equations

The complete function, accounting for the different outcomes depending on *i:*

$$F[\sigma\_] := \frac{\sigma}{2-\sigma};$$

$$\Phi[r\_, \sigma\_] := \frac{\sigma\,(2-\sigma-2\,(1-r)\,r\,(2-3\,\sigma))}{(2-\sigma)\,(2-(1-2\,(1-r)\,r)\,\sigma)};$$

$$PNR[Na\_, F\_, \Phi\_, s\_, h\_, R\_, p0\_] := \left(\left(\frac{(F+h-F\,h)}{(1-h+F\,h)}\left(\frac{1}{p0}+1\right)-1\right)^{-\frac{R\,(1-F)}{2\,Na\,(F+h-F\,h)\,s}}\right);$$

$$PrKR[k\_, n\_, R\_, F\_, \Phi\_, p0\_] := \left(\left(2\,\frac{R}{1+F}\,(1-2\,F+\Phi)\,p0\,(1-p0)\right)^{k} Abs[StirlingS1[n, k]]\right)\Big/ Product\left[\left(2\,\frac{R}{1+F}\,(1-2\,F+\Phi)\,p0\,(1-p0)+a\right), \{a, 0, n-1\}\right];$$

$$PrJ[j\_, k\_] := \frac{1}{j\,Sum\left[\frac{1}{a}, \{a, k-1\}\right]};$$

```
H[g_, j_, k_, n_, i_] :=
  (Binomial[n - i, g] * Binomial[k, j - g]) / Binomial[k + n - i, j];
PrL[n_, k_, j_, l_] :=
  ((Binomial[n, l] * Abs[StirlingS1[l, j]] * Abs[StirlingS1[n - l, k - j]]) /
    (Binomial[k, j] * Abs[StirlingS1[n, k]]));
```

$$TFixI[Na\_, s\_, h\_, F\_, p0\_] := \frac{EulerGamma + Log\left[\frac{4\,Na\,s\,(1-(1-F)\,h)\,(1-p0)}{1+F}\right]}{s\,(1-(1-F)\,h)} - \frac{Log[p0]}{s\,(h+F-h\,F)} + \frac{(1-F)\,(1-2\,h)}{s\,(h+F-h\,F)\,(1-(1-F)\,h)}\,Log\left[\frac{h+F-h\,F+(1-F)\,(1-2\,h)\,p0}{1-(1-F)\,h}\right]$$

$$TFixIC[Na\_, s\_, h\_, F\_, p0\_] := \frac{(1+F)\,TFixI[Na, s, h, F, p0]}{2\,Na}$$

$$Boostp0[Na\_, s\_, h\_, \sigma\_] := \frac{1+F[\sigma]}{4\,Na\,s\,(F[\sigma]+h-F[\sigma]\,h)}$$

```
PLNH[Na_, s_, h_, σ_, R_, n_, l_, p0_, θ_] :=
 Sum[PDF[BinomialDistribution[n, PNR[Na, F[σ], Φ[R/(2 Na), σ], s, h, R, p0]], i] *
   (Piecewise[{{Sum[PrKR[k, i, R, F[σ], Φ[R/(2 Na), σ], p0] *
          Sum[PrJ[j, k + n - i] * Sum[H[g, j, k, n, i] * PrL[i, k, j - g, l - g],
             {g, Max[{j - k, l - i}], Min[{j, l, n - i}]}], {j, 1,
            Min[{k + n - i - 1, l}]}], {k, 1, i}], 0 < i < n}, {0, i == 0 || i == n}}] +
     Piecewise[{{1/(l Sum[1/a, {a, n - 1}]), i == 0}, {0, i ≠ 0}}] +
     Piecewise[{{PrKR[1, i, R, F[σ], Φ[R/(2 Na), σ], p0] * Piecewise[
           {{θ * p0 + (θ n)/2 TFixIC[Na, s, h, F[σ], p0], l == 1}, {(θ * p0)/l, l ≠ 1}}] +
          Sum[PrKR[k, i, R, F[σ], Φ[R/(2 Na), σ], p0] * Sum[PrJ[j, k] * PrL[n, k, j, l],
             {j, 1, Min[{k - 1, l}]}], {k, 2, n}], i == n}, {0, i ≠ n}}]), {i, 0, n}]

PLNHS[Na_, s_, h_, σ_, R_, n_, p0_, θ_] :=
 Sum[PLNH[Na, s, h, σ, R, n, L, p0, θ], {L, 1, n - 1}]
PLNH2[Na_, s_, h_, σ_, R_, n_, l_, p0_, θ_] := PLNH[Na, s, h, σ, R, n, l, p0, θ] / PLNHS[Na, s, h, σ, R, n, p0, θ]
```

### Outcrossing results ($\sigma = F = 0$)

#### Results with $p_0 = 1/2N$

R = 6

SLiM simulation results

```
datfDNR6h01 = Import[
    "SLIM_SFS_F0/SFSTab_R_6_h_0.1_self_0_f_1e-04_10b_SLIM.dat", "Table"][[1]];
datfDNR6h01CI = Import[
    "SLIM_SFS_F0/SFSTab_R_6_h_0.1_self_0_f_1e-04_10b_SLIM.dat", "Table"][[2]];
datfDNR6h05 = Import["SLIM_SFS_F0/SFSTab_R_6_h_0.5_self_0_f_1e-04_10b_SLIM.dat",
    "Table"][[1]];
datfDNR6h05CI = Import[
    "SLIM_SFS_F0/SFSTab_R_6_h_0.5_self_0_f_1e-04_10b_SLIM.dat", "Table"][[2]];
datfDNR6h09 = Import["SLIM_SFS_F0/SFSTab_R_6_h_0.9_self_0_f_1e-04_10b_SLIM.dat",
    "Table"][[1]];
datfDNR6h09CI = Import[
    "SLIM_SFS_F0/SFSTab_R_6_h_0.9_self_0_f_1e-04_10b_SLIM.dat", "Table"][[2]];
datfDNR6h01T = Partition[Riffle[Partition[Riffle[Range[9], datfDNR6h01], 2],
    Map[ErrorBar, datfDNR6h01CI]], 2];
datfDNR6h05T = Partition[Riffle[Partition[Riffle[Range[9], datfDNR6h05], 2],
    Map[ErrorBar, datfDNR6h05CI]], 2];
datfDNR6h09T = Partition[Riffle[Partition[Riffle[Range[9], datfDNR6h09], 2],
    Map[ErrorBar, datfDNR6h09CI]], 2];
```

MSMS simulation results

```
datfDNR6h01MSMS =
  Import["SFS_MSMS_SV_5K/SFSTab_R_12_h_0.1_self_0_f_1e-04_10b_MSMS.dat",
    "Table"][[1]];
datfDNR6h01MSMSCI = Import[
    "SFS_MSMS_SV_5K/SFSTab_R_12_h_0.1_self_0_f_1e-04_10b_MSMS.dat",
    "Table"][[2]];
datfDNR6h05MSMS = Import[
    "SFS_MSMS_SV_5K/SFSTab_R_12_h_0.5_self_0_f_1e-04_10b_MSMS.dat",
    "Table"][[1]];
datfDNR6h05MSMSCI = Import[
    "SFS_MSMS_SV_5K/SFSTab_R_12_h_0.5_self_0_f_1e-04_10b_MSMS.dat",
    "Table"][[2]];
datfDNR6h09MSMS = Import[
    "SFS_MSMS_SV_5K/SFSTab_R_12_h_0.9_self_0_f_1e-04_10b_MSMS.dat",
    "Table"][[1]];
datfDNR6h09MSMSCI = Import[
    "SFS_MSMS_SV_5K/SFSTab_R_12_h_0.9_self_0_f_1e-04_10b_MSMS.dat",
    "Table"][[2]];
datfDNR6h01MSMST = Partition[Riffle[Partition[Riffle[Range[9], datfDNR6h01MSMS],
     2], Map[ErrorBar, datfDNR6h01MSMSCI]], 2];
datfDNR6h05MSMST = Partition[Riffle[Partition[Riffle[Range[9], datfDNR6h05MSMS],
     2], Map[ErrorBar, datfDNR6h05MSMSCI]], 2];
datfDNR6h09MSMST = Partition[Riffle[Partition[Riffle[Range[9], datfDNR6h09MSMS],
     2], Map[ErrorBar, datfDNR6h09MSMSCI]], 2];
```

Analytical solutions

```
p1R6 = ListPlot[
  {Table[{l, PLNH2[5000, 0.05, 0.1, 0, 6, 10, l, Boostp0[5000, 0.05, 0.1, 0], 4]},
    {l, 1, 9}], Table[{l, PLNH2[5000, 0.05, 0.5, 0, 6,
      10, l, Boostp0[5000, 0.05, 0.5, 0], 4]}, {l, 1, 9}],
   Table[{l, PLNH2[5000, 0.05, 0.9, 0, 6, 10, l, Boostp0[5000, 0.05, 0.9, 0], 4]},
    {l, 1, 9}], Table[{l, PrJ[l, 10]}, {l, 1, 9}]}, PlotRange → All,
  PlotStyle → {Red, Black, Blue, {Gray, Dashed}}, Joined → True,
  Ticks → {{1, 2, 3, 4, 5, 6, 7, 8, 9}, Automatic},
  BaseStyle → {FontWeight → "Bold", FontColor → Black, FontSize → 12}]
```

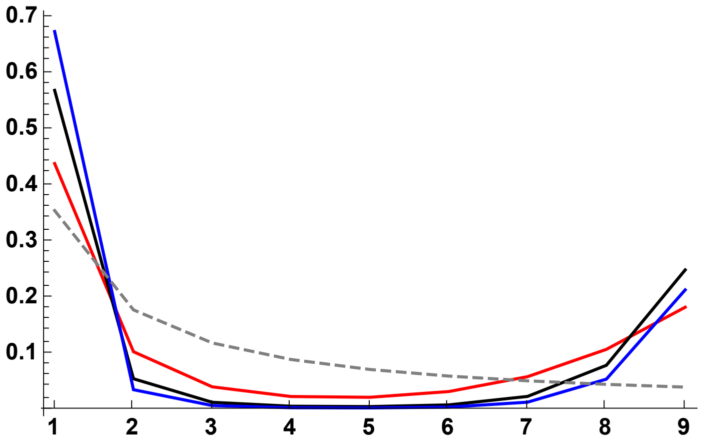

SLiM simulations plot

```
p2R6 = ErrorListPlot[
  {datfDNR6h01T, datfDNR6h05T, datfDNR6h09T}, PlotRange → All, PlotStyle →
   {{Red, PointSize[0.02]}, {Black, PointSize[0.02]}, {Blue, PointSize[0.02]}}]
```

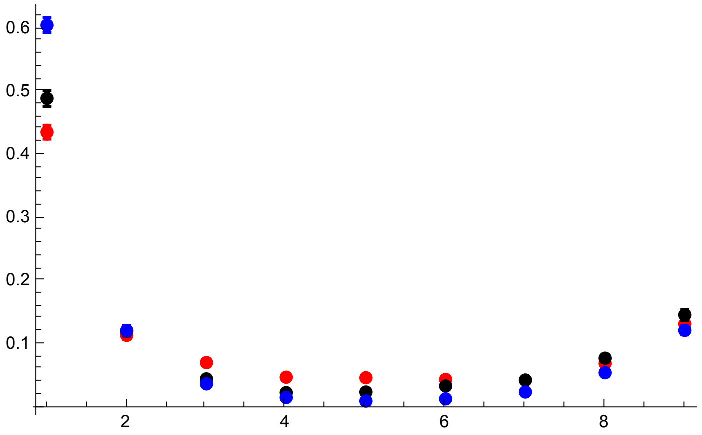

MSMS simulations plot

```
p2AR6 = ErrorListPlot[{datfDNR6h01MSMST, datfDNR6h05MSMST, datfDNR6h09MSMST},
  PlotRange → All, PlotStyle →
   {{Pink, PointSize[0.02]}, {Gray, PointSize[0.02]}, {Cyan, PointSize[0.02]}}]
```

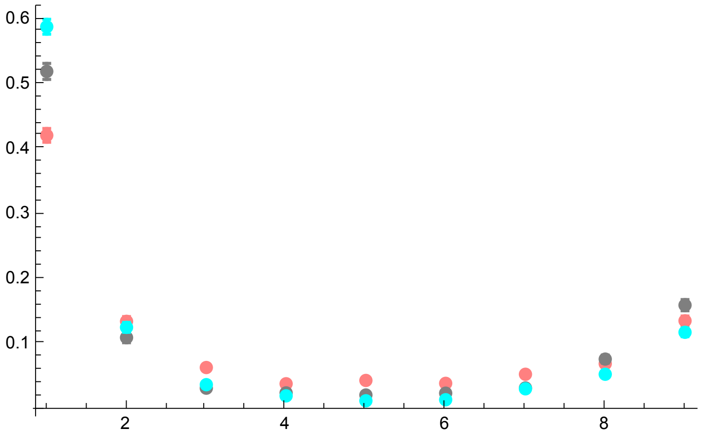


All together

```
p3R6s0DN = Show[p1R6, p2R6, p2AR6, PlotRange → All, ImageSize → 325]
```

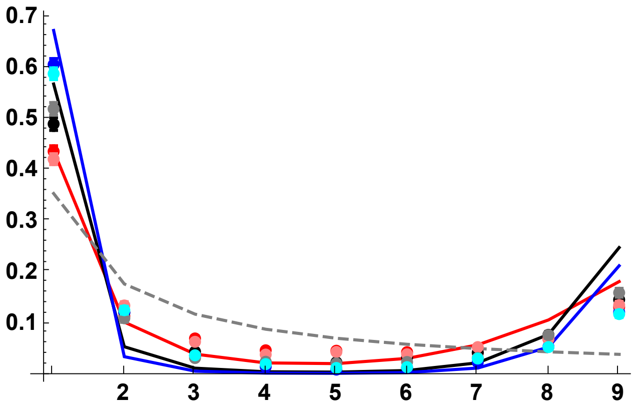


Analytical results and SLiM simulations

```
p3AR6s0DN = Show[p1R6, p2R6, PlotRange → All, ImageSize → 325]
```

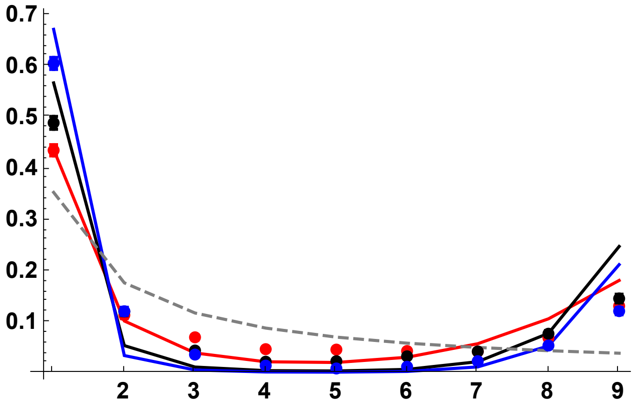


Simulation results only (SLiM and MSMS)

```
p3BR6s0DN = Show[p2R6, p2AR6, PlotRange → All]
```

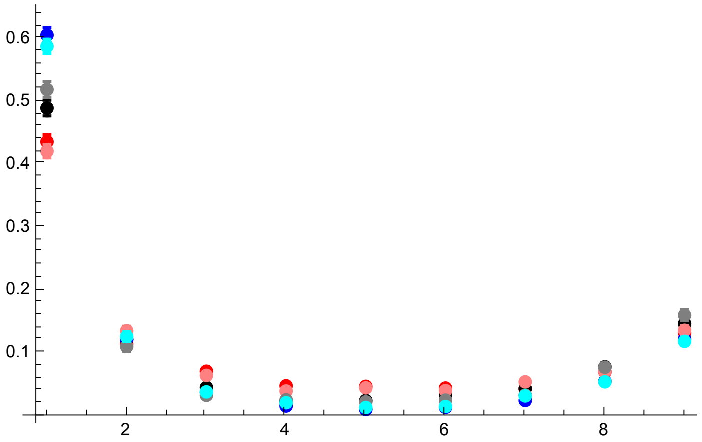


R = 18

SLiM simulation results

```
datfDNR18h01 =
  Import["SLIM_SFS_F0/SFSTab_R_18_h_0.1_self_0_f_1e-04_10b_SLIM.dat",
    "Table"][[1]];
datfDNR18h01CI = Import[
    "SLIM_SFS_F0/SFSTab_R_18_h_0.1_self_0_f_1e-04_10b_SLIM.dat",
    "Table"][[2]];
datfDNR18h05 = Import[
    "SLIM_SFS_F0/SFSTab_R_18_h_0.5_self_0_f_1e-04_10b_SLIM.dat",
    "Table"][[1]];
datfDNR18h05CI = Import[
    "SLIM_SFS_F0/SFSTab_R_18_h_0.5_self_0_f_1e-04_10b_SLIM.dat",
    "Table"][[2]];
datfDNR18h09 = Import[
    "SLIM_SFS_F0/SFSTab_R_18_h_0.9_self_0_f_1e-04_10b_SLIM.dat",
    "Table"][[1]];
datfDNR18h09CI = Import[
    "SLIM_SFS_F0/SFSTab_R_18_h_0.9_self_0_f_1e-04_10b_SLIM.dat",
    "Table"][[2]];
datfDNR18h01T = Partition[Riffle[Partition[Riffle[Range[9], datfDNR18h01], 2],
    Map[ErrorBar, datfDNR18h01CI]], 2];
datfDNR18h05T = Partition[Riffle[Partition[Riffle[Range[9], datfDNR18h05], 2],
    Map[ErrorBar, datfDNR18h05CI]], 2];
datfDNR18h09T = Partition[Riffle[Partition[Riffle[Range[9], datfDNR18h09], 2],
    Map[ErrorBar, datfDNR18h09CI]], 2];
```

MSMS simulation results

```
datfDNR18h01MSMS =
  Import["SFS_MSMS_SV_5K/SFSTab_R_36_h_0.1_self_0_f_1e-04_10b_MSMS.dat",
    "Table"][[1]];
datfDNR18h01MSMSCI = Import[
    "SFS_MSMS_SV_5K/SFSTab_R_36_h_0.1_self_0_f_1e-04_10b_MSMS.dat",
    "Table"][[2]];
datfDNR18h05MSMS = Import[
    "SFS_MSMS_SV_5K/SFSTab_R_36_h_0.5_self_0_f_1e-04_10b_MSMS.dat",
    "Table"][[1]];
datfDNR18h05MSMSCI = Import[
    "SFS_MSMS_SV_5K/SFSTab_R_36_h_0.5_self_0_f_1e-04_10b_MSMS.dat",
    "Table"][[2]];
datfDNR18h09MSMS = Import[
    "SFS_MSMS_SV_5K/SFSTab_R_36_h_0.9_self_0_f_1e-04_10b_MSMS.dat",
    "Table"][[1]];
datfDNR18h09MSMSCI = Import[
    "SFS_MSMS_SV_5K/SFSTab_R_36_h_0.9_self_0_f_1e-04_10b_MSMS.dat",
    "Table"][[2]];
datfDNR18h01MSMST = Partition[Riffle[Partition[Riffle[Range[9],
      datfDNR18h01MSMS], 2], Map[ErrorBar, datfDNR18h01MSMSCI]], 2];
datfDNR18h05MSMST = Partition[Riffle[Partition[Riffle[Range[9],
      datfDNR18h05MSMS], 2], Map[ErrorBar, datfDNR18h05MSMSCI]], 2];
datfDNR18h09MSMST = Partition[Riffle[Partition[Riffle[Range[9],
      datfDNR18h09MSMS], 2], Map[ErrorBar, datfDNR18h09MSMSCI]], 2];
```

Analytical solutions

```
p1R18 = ListPlot[
  {Table[{l, PLNH2[5000, 0.05, 0.1, 0, 18, 10, l, Boostp0[5000, 0.05, 0.1, 0], 4]},
    {l, 1, 9}], Table[{l, PLNH2[5000, 0.05, 0.5, 0, 18,
      10, l, Boostp0[5000, 0.05, 0.5, 0], 4]}, {l, 1, 9}],
   Table[{l, PLNH2[5000, 0.05, 0.9, 0, 18, 10, l, Boostp0[5000, 0.05, 0.9, 0], 4]},
    {l, 1, 9}], Table[{l, PrJ[l, 10]}, {l, 1, 9}]}, PlotRange → All,
  PlotStyle → {Red, Black, Blue, {Gray, Dashed}}, Joined → True,
  Ticks → {{1, 2, 3, 4, 5, 6, 7, 8, 9}, Automatic},
  BaseStyle → {FontWeight → "Bold", FontColor → Black, FontSize → 12}]
```

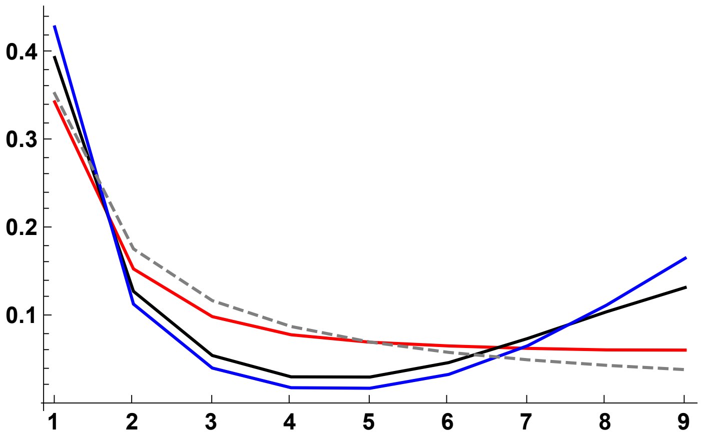

SLiM simulations plot

```
p2R18 = ErrorListPlot[
  {datfDNR18h01T, datfDNR18h05T, datfDNR18h09T}, PlotRange → All, PlotStyle →
   {{Red, PointSize[0.02]}, {Black, PointSize[0.02]}, {Blue, PointSize[0.02]}}]
```

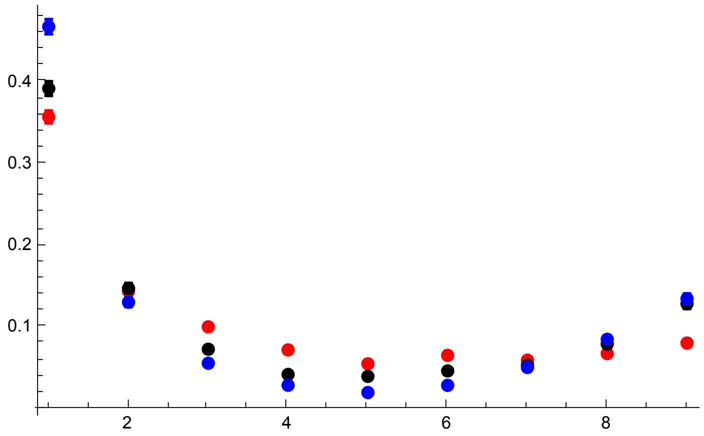


MSMS simulation results

```
p2AR18 = ErrorListPlot[{datfDNR18h01MSMST, datfDNR18h05MSMST, datfDNR18h09MSMST},
  PlotRange → All, PlotStyle →
   {{Pink, PointSize[0.02]}, {Gray, PointSize[0.02]}, {Cyan, PointSize[0.02]}}]
```

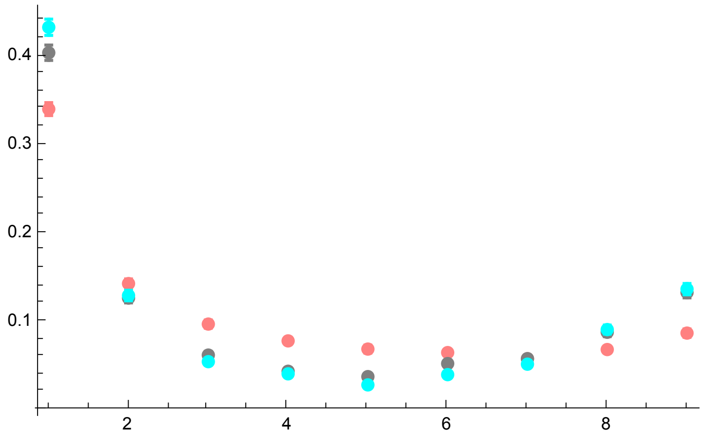


All together

```
p3R18s0DN = Show[p1R18, p2R18, p2AR18, PlotRange → All, ImageSize → 325]
```

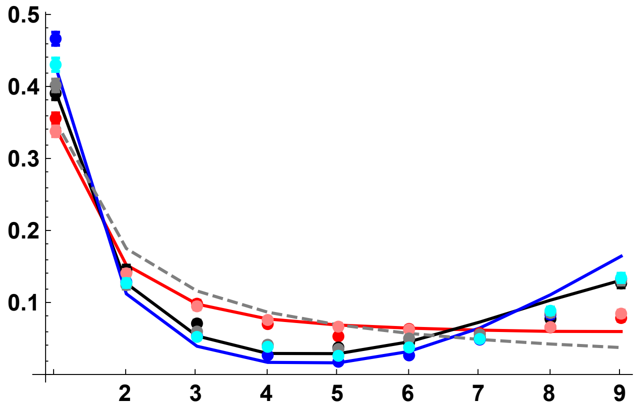

Analytical results and SLiM simulations

```
p3AR18 = Show[p1R18, p2R18, PlotRange → All]
```

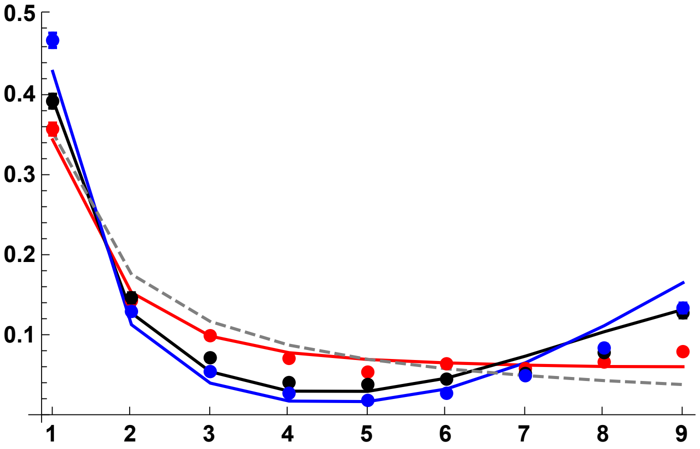


Simulation results only (SLiM and MSMS)

```
p3BR18 = Show[p2R18, p2AR18, PlotRange → All]
```

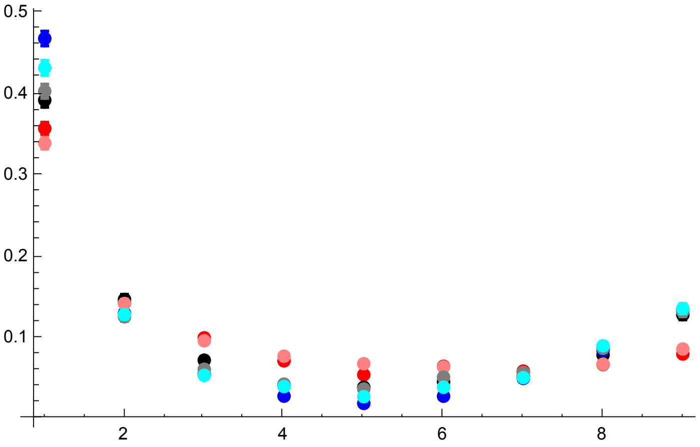


R = 30

SLiM simulations plot

```
datfDNR30h01 =
  Import["SLIM_SFS_F0/SFSTab_R_30_h_0.1_self_0_f_1e-04_10b_SLIM.dat",
    "Table"][[1]];
datfDNR30h01CI = Import[
    "SLIM_SFS_F0/SFSTab_R_30_h_0.1_self_0_f_1e-04_10b_SLIM.dat",
    "Table"][[2]];
datfDNR30h05 = Import[
    "SLIM_SFS_F0/SFSTab_R_30_h_0.5_self_0_f_1e-04_10b_SLIM.dat",
    "Table"][[1]];
datfDNR30h05CI = Import[
    "SLIM_SFS_F0/SFSTab_R_30_h_0.5_self_0_f_1e-04_10b_SLIM.dat",
    "Table"][[2]];
datfDNR30h09 = Import[
    "SLIM_SFS_F0/SFSTab_R_30_h_0.9_self_0_f_1e-04_10b_SLIM.dat",
    "Table"][[1]];
datfDNR30h09CI = Import[
    "SLIM_SFS_F0/SFSTab_R_30_h_0.9_self_0_f_1e-04_10b_SLIM.dat",
    "Table"][[2]];
datfDNR30h01T = Partition[Riffle[Partition[Riffle[Range[9], datfDNR30h01], 2],
    Map[ErrorBar, datfDNR30h01CI]], 2];
datfDNR30h05T = Partition[Riffle[Partition[Riffle[Range[9], datfDNR30h05], 2],
    Map[ErrorBar, datfDNR30h05CI]], 2];
datfDNR30h09T = Partition[Riffle[Partition[Riffle[Range[9], datfDNR30h09], 2],
    Map[ErrorBar, datfDNR30h09CI]], 2];
```

MSMS simulation results

```
datfDNR30h01MSMS =
  Import["SFS_MSMS_SV_5K/SFSTab_R_60_h_0.1_self_0_f_1e-04_10b_MSMS.dat",
    "Table"][[1]];
datfDNR30h01MSMSCI = Import[
    "SFS_MSMS_SV_5K/SFSTab_R_60_h_0.1_self_0_f_1e-04_10b_MSMS.dat",
    "Table"][[2]];
datfDNR30h05MSMS = Import[
    "SFS_MSMS_SV_5K/SFSTab_R_60_h_0.5_self_0_f_1e-04_10b_MSMS.dat",
    "Table"][[1]];
datfDNR30h05MSMSCI = Import[
    "SFS_MSMS_SV_5K/SFSTab_R_60_h_0.5_self_0_f_1e-04_10b_MSMS.dat",
    "Table"][[2]];
datfDNR30h09MSMS = Import[
    "SFS_MSMS_SV_5K/SFSTab_R_60_h_0.9_self_0_f_1e-04_10b_MSMS.dat",
    "Table"][[1]];
datfDNR30h09MSMSCI = Import[
    "SFS_MSMS_SV_5K/SFSTab_R_60_h_0.9_self_0_f_1e-04_10b_MSMS.dat",
    "Table"][[2]];
datfDNR30h01MSMST = Partition[Riffle[Partition[Riffle[Range[9],
      datfDNR30h01MSMS], 2], Map[ErrorBar, datfDNR30h01MSMSCI]], 2];
datfDNR30h05MSMST = Partition[Riffle[Partition[Riffle[Range[9],
      datfDNR30h05MSMS], 2], Map[ErrorBar, datfDNR30h05MSMSCI]], 2];
datfDNR30h09MSMST = Partition[Riffle[Partition[Riffle[Range[9],
      datfDNR30h09MSMS], 2], Map[ErrorBar, datfDNR30h09MSMSCI]], 2];
```

Analytical solutions

```
p1R30 = ListPlot[
  {Table[{l, PLNH2[5000, 0.05, 0.1, 0, 30, 10, l, Boostp0[5000, 0.05, 0.1, 0], 4]},
    {l, 1, 9}], Table[{l, PLNH2[5000, 0.05, 0.5, 0, 30,
      10, l, Boostp0[5000, 0.05, 0.5, 0], 4]}, {l, 1, 9}],
   Table[{l, PLNH2[5000, 0.05, 0.9, 0, 30, 10, l, Boostp0[5000, 0.05, 0.9, 0], 4]},
    {l, 1, 9}], Table[{l, PrJ[l, 10]}, {l, 1, 9}]}, PlotRange → All,
  PlotStyle → {Red, Black, Blue, {Gray, Dashed}}, Joined → True]
```

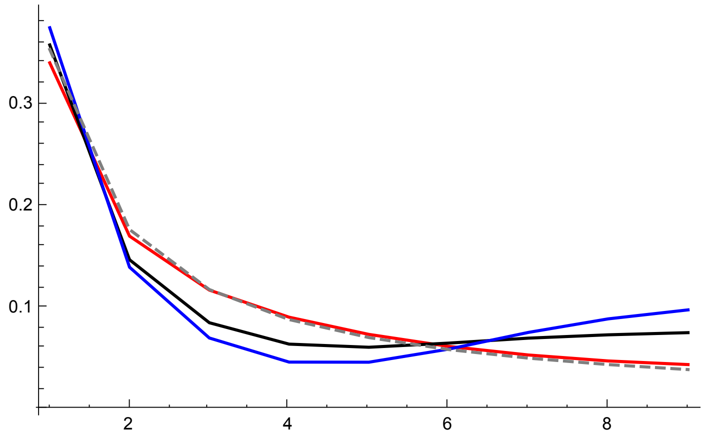


SLiM simulations plot

```
p2R30 = ErrorListPlot[
  {datfDNR30h01T, datfDNR30h05T, datfDNR30h09T}, PlotRange → All, PlotStyle →
   {{Red, PointSize[0.02]}, {Black, PointSize[0.02]}, {Blue, PointSize[0.02]}}]
```

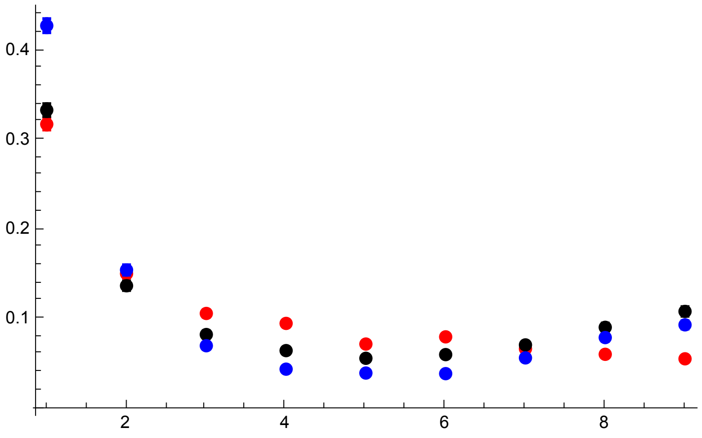


MSMS simulation results

```
p2AR30 = ErrorListPlot[{datfDNR30h01MSMST, datfDNR30h05MSMST, datfDNR30h09MSMST},
  PlotRange → All, PlotStyle →
   {{Pink, PointSize[0.02]}, {Gray, PointSize[0.02]}, {Cyan, PointSize[0.02]}}]
```

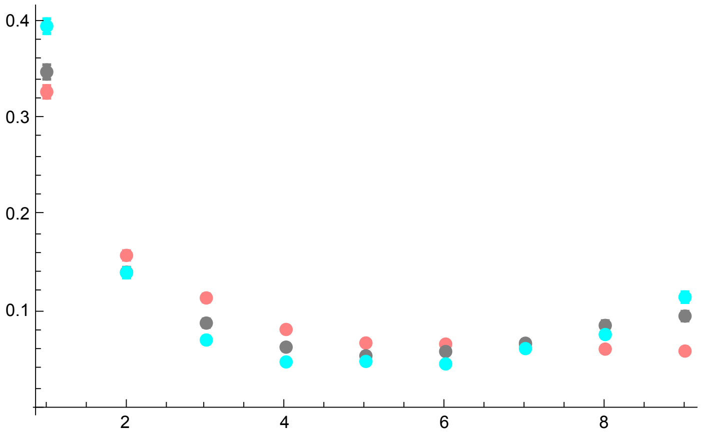


All together

```
p3R30 = Show[p1R30, p2R30, p2AR30, PlotRange → All]
```

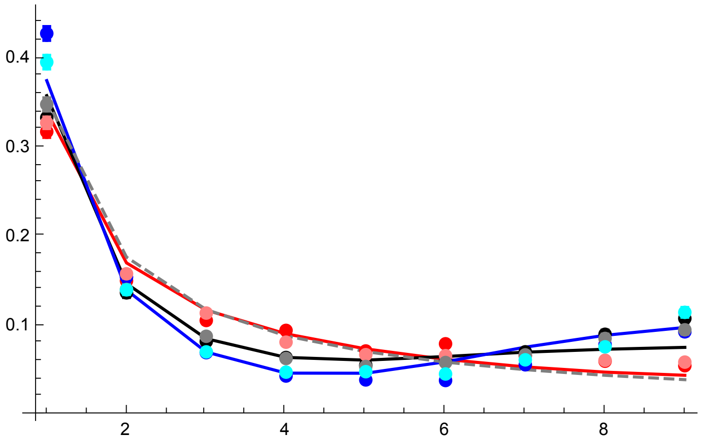


Analytical results and SLiM simulations

```
p3AR30 = Show[p1R30, p2R30, PlotRange → All]
```

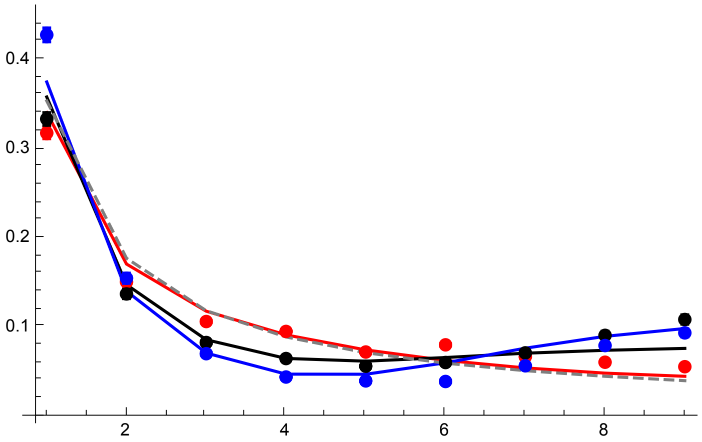


Simulation results only (SLiM and MSMS)

```
p3BR30 = Show[p2R30, p2AR30, PlotRange → All]
```

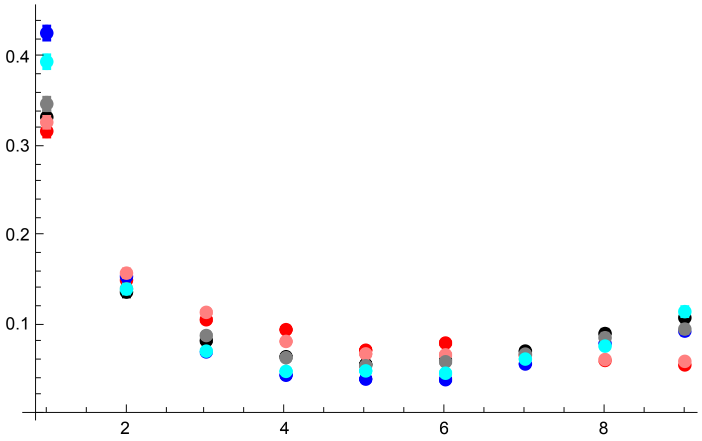


R = 42

SLiM simulations results

```
datfDNR42h01 =
  Import["SLIM_SFS_F0/SFSTab_R_42_h_0.1_self_0_f_1e-04_10b_SLIM.dat",
    "Table"][[1]];
datfDNR42h01CI = Import[
    "SLIM_SFS_F0/SFSTab_R_42_h_0.1_self_0_f_1e-04_10b_SLIM.dat",
    "Table"][[2]];
datfDNR42h05 = Import[
    "SLIM_SFS_F0/SFSTab_R_42_h_0.5_self_0_f_1e-04_10b_SLIM.dat",
    "Table"][[1]];
datfDNR42h05CI = Import[
    "SLIM_SFS_F0/SFSTab_R_42_h_0.5_self_0_f_1e-04_10b_SLIM.dat",
    "Table"][[2]];
datfDNR42h09 = Import[
    "SLIM_SFS_F0/SFSTab_R_42_h_0.9_self_0_f_1e-04_10b_SLIM.dat",
    "Table"][[1]];
datfDNR42h09CI = Import[
    "SLIM_SFS_F0/SFSTab_R_42_h_0.9_self_0_f_1e-04_10b_SLIM.dat",
    "Table"][[2]];
datfDNR42h01T = Partition[Riffle[Partition[Riffle[Range[9], datfDNR42h01], 2],
    Map[ErrorBar, datfDNR42h01CI]], 2];
datfDNR42h05T = Partition[Riffle[Partition[Riffle[Range[9], datfDNR42h05], 2],
    Map[ErrorBar, datfDNR42h05CI]], 2];
datfDNR42h09T = Partition[Riffle[Partition[Riffle[Range[9], datfDNR42h09], 2],
    Map[ErrorBar, datfDNR42h09CI]], 2];
```

MSMS simulation results

```
datfDNR42h01MSMS =
  Import["SFS_MSMS_SV_5K/SFSTab_R_84_h_0.1_self_0_f_1e-04_10b_MSMS.dat",
    "Table"][[1]];
datfDNR42h01MSMSCI = Import[
    "SFS_MSMS_SV_5K/SFSTab_R_84_h_0.1_self_0_f_1e-04_10b_MSMS.dat",
    "Table"][[2]];
datfDNR42h05MSMS = Import[
    "SFS_MSMS_SV_5K/SFSTab_R_84_h_0.5_self_0_f_1e-04_10b_MSMS.dat",
    "Table"][[1]];
datfDNR42h05MSMSCI = Import[
    "SFS_MSMS_SV_5K/SFSTab_R_84_h_0.5_self_0_f_1e-04_10b_MSMS.dat",
    "Table"][[2]];
datfDNR42h09MSMS = Import[
    "SFS_MSMS_SV_5K/SFSTab_R_84_h_0.9_self_0_f_1e-04_10b_MSMS.dat",
    "Table"][[1]];
datfDNR42h09MSMSCI = Import[
    "SFS_MSMS_SV_5K/SFSTab_R_84_h_0.9_self_0_f_1e-04_10b_MSMS.dat",
    "Table"][[2]];
datfDNR42h01MSMST = Partition[Riffle[Partition[Riffle[Range[9],
      datfDNR42h01MSMS], 2], Map[ErrorBar, datfDNR42h01MSMSCI]], 2];
datfDNR42h05MSMST = Partition[Riffle[Partition[Riffle[Range[9],
      datfDNR42h05MSMS], 2], Map[ErrorBar, datfDNR42h05MSMSCI]], 2];
datfDNR42h09MSMST = Partition[Riffle[Partition[Riffle[Range[9],
      datfDNR42h09MSMS], 2], Map[ErrorBar, datfDNR42h09MSMSCI]], 2];
```

Analytic solutions

```
p1R42 = ListPlot[
  {Table[{l, PLNH2[5000, 0.05, 0.1, 0, 42, 10, l, Boostp0[5000, 0.05, 0.1, 0], 4]},
    {l, 1, 9}], Table[{l, PLNH2[5000, 0.05, 0.5, 0, 42,
      10, l, Boostp0[5000, 0.05, 0.5, 0], 4]}, {l, 1, 9}],
   Table[{l, PLNH2[5000, 0.05, 0.9, 0, 42, 10, l, Boostp0[5000, 0.05, 0.9, 0], 4]},
    {l, 1, 9}], Table[{l, PrJ[l, 10]}, {l, 1, 9}]}, PlotRange → All,
  PlotStyle → {Red, Black, Blue, {Gray, Dashed}}, Joined → True]
```

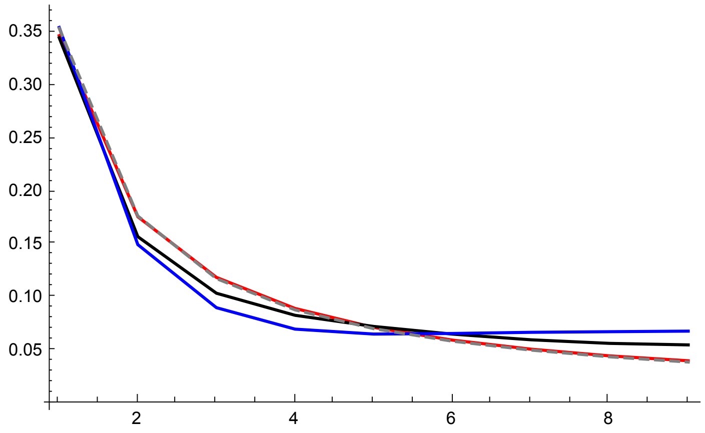


SLiM simulations plot

```
p2R42 = ErrorListPlot[
  {datfDNR42h01T, datfDNR42h05T, datfDNR42h09T}, PlotRange → All, PlotStyle →
   {{Red, PointSize[0.02]}, {Black, PointSize[0.02]}, {Blue, PointSize[0.02]}}]
```

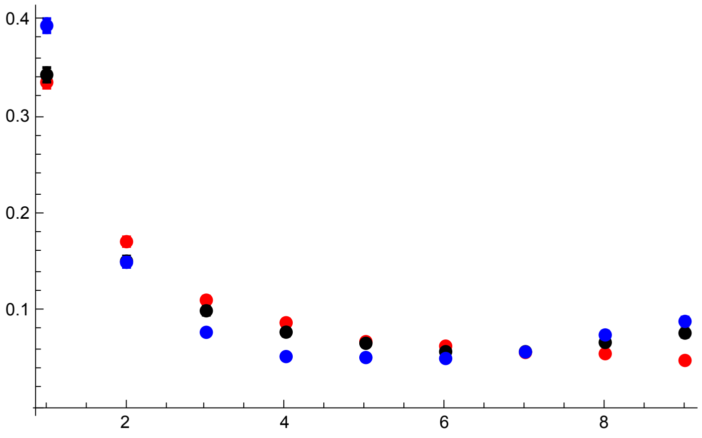


MSMS simulation results

```
p2AR42 = ErrorListPlot[{datfDNR42h01MSMST, datfDNR42h05MSMST, datfDNR42h09MSMST},
  PlotRange → All, PlotStyle →
   {{Pink, PointSize[0.02]}, {Gray, PointSize[0.02]}, {Cyan, PointSize[0.02]}}]
```

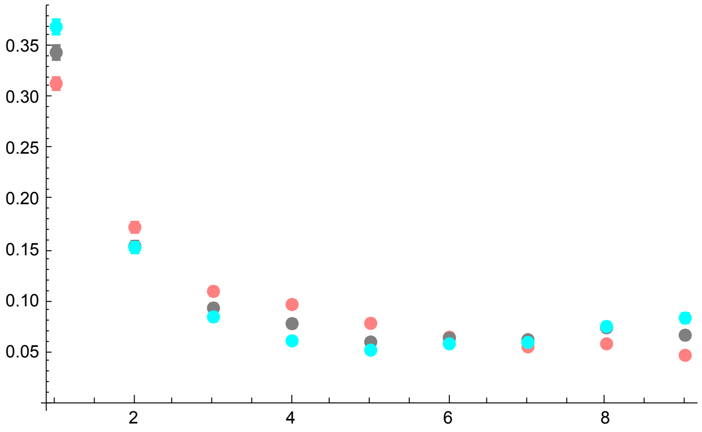


All together

```
p3R42 = Show[p1R42, p2R42, p2AR42, PlotRange → All]
```

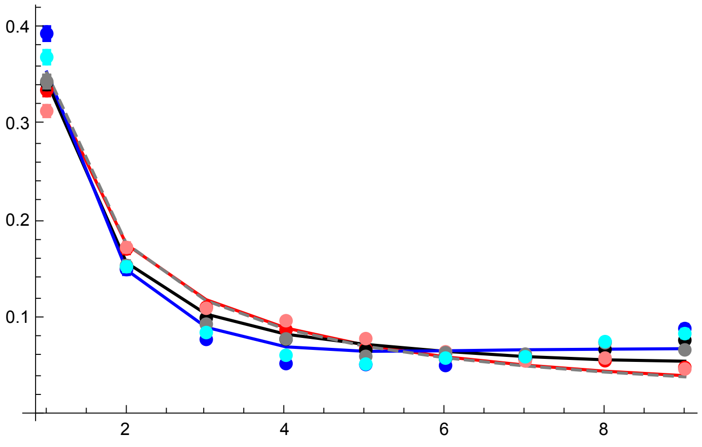


Analytical solutions and SLIM results

```
p3AR42 = Show[p1R42, p2R42, PlotRange → All]
```

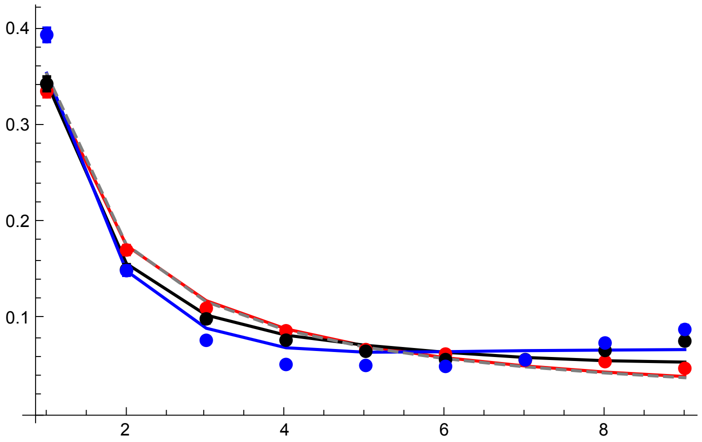


Simulation results only (SLiM and MSMS)

```
p3BR42 = Show[p2R42, p2AR42, PlotRange → All]
```

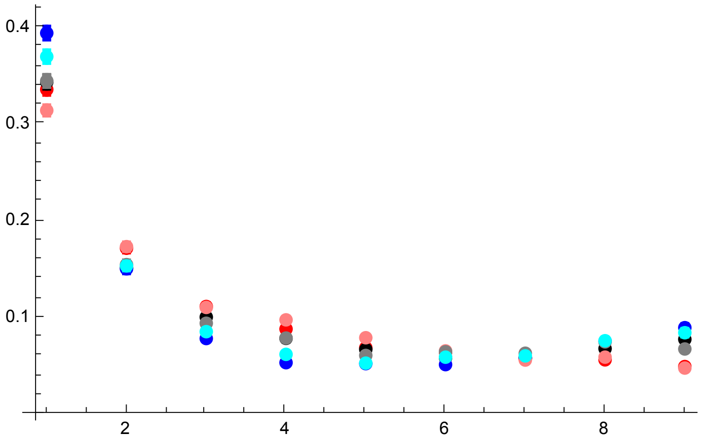


All plots compared

```
GraphicsGrid[{{p3AR6s0DN, p3AR18}, {p3AR30, p3AR42}}]
```

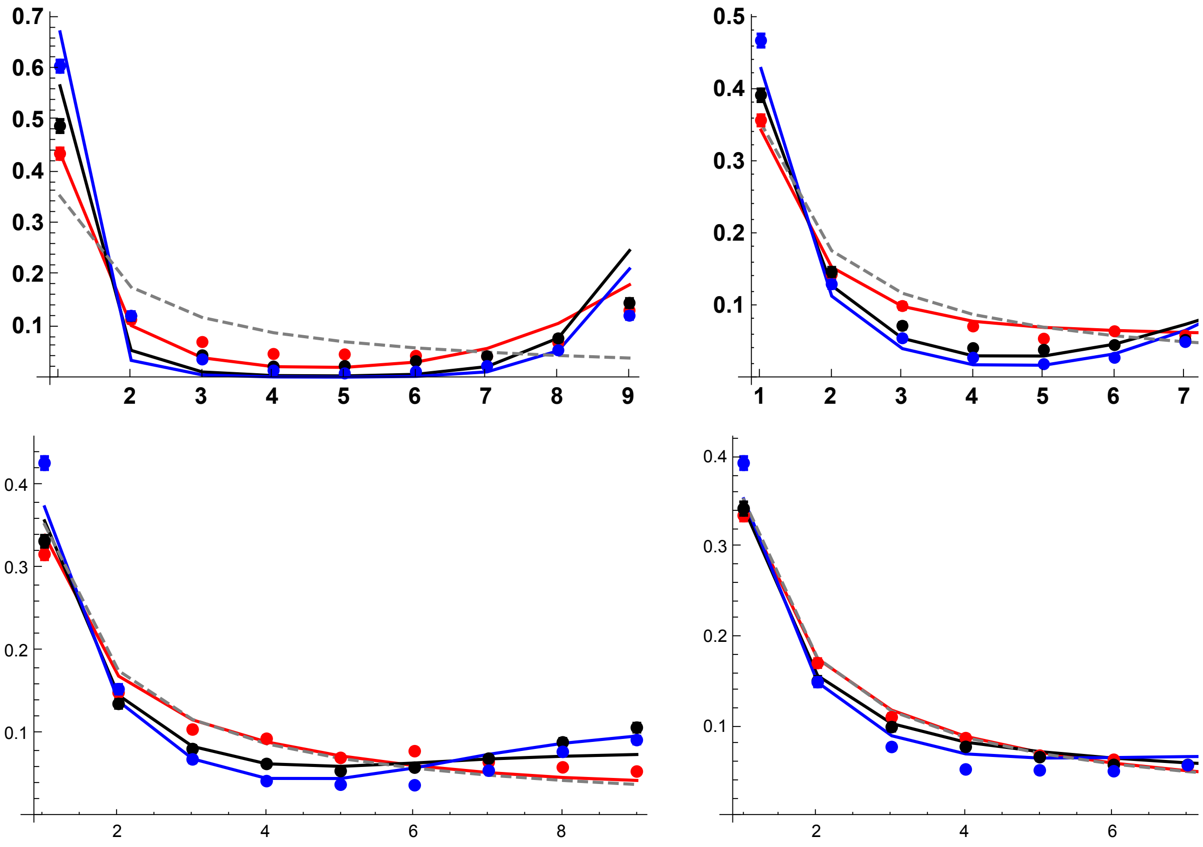


#### Results with $p_0 = 0.02$

R = 6

SLiM simulations results

```
datp002R6h01 = Import[
    "SLIM_SFS_F0/SFSTab_R_6_h_0.1_self_0_f_0.02_10b_SLIM.dat", "Table"][[1]];
datp002R6h01CI = Import[
    "SLIM_SFS_F0/SFSTab_R_6_h_0.1_self_0_f_0.02_10b_SLIM.dat", "Table"][[2]];
datp002R6h05 = Import["SLIM_SFS_F0/SFSTab_R_6_h_0.5_self_0_f_0.02_10b_SLIM.dat",
    "Table"][[1]];
datp002R6h05CI = Import[
    "SLIM_SFS_F0/SFSTab_R_6_h_0.5_self_0_f_0.02_10b_SLIM.dat", "Table"][[2]];
datp002R6h09 = Import["SLIM_SFS_F0/SFSTab_R_6_h_0.9_self_0_f_0.02_10b_SLIM.dat",
    "Table"][[1]];
datp002R6h09CI = Import[
    "SLIM_SFS_F0/SFSTab_R_6_h_0.9_self_0_f_0.02_10b_SLIM.dat", "Table"][[2]];
datp002R6h01T = Partition[Riffle[Partition[Riffle[Range[9], datp002R6h01], 2],
    Map[ErrorBar, datp002R6h01CI]], 2];
datp002R6h05T = Partition[Riffle[Partition[Riffle[Range[9], datp002R6h05], 2],
    Map[ErrorBar, datp002R6h05CI]], 2];
datp002R6h09T = Partition[Riffle[Partition[Riffle[Range[9], datp002R6h09], 2],
    Map[ErrorBar, datp002R6h09CI]], 2];
```

MSMS simulation results

```
datp002R6h01MSMS =
  Import["SFS_MSMS_SV_5K/SFSTab_R_12_h_0.1_self_0_f_0.02_10b_MSMS.dat",
    "Table"][[1]];
datp002R6h01MSMSCI = Import[
    "SFS_MSMS_SV_5K/SFSTab_R_12_h_0.1_self_0_f_0.02_10b_MSMS.dat",
    "Table"][[2]];
datp002R6h05MSMS = Import[
    "SFS_MSMS_SV_5K/SFSTab_R_12_h_0.5_self_0_f_0.02_10b_MSMS.dat",
    "Table"][[1]];
datp002R6h05MSMSCI = Import[
    "SFS_MSMS_SV_5K/SFSTab_R_12_h_0.5_self_0_f_0.02_10b_MSMS.dat",
    "Table"][[2]];
datp002R6h09MSMS = Import[
    "SFS_MSMS_SV_5K/SFSTab_R_12_h_0.9_self_0_f_0.02_10b_MSMS.dat",
    "Table"][[1]];
datp002R6h09MSMSCI = Import[
    "SFS_MSMS_SV_5K/SFSTab_R_12_h_0.9_self_0_f_0.02_10b_MSMS.dat",
    "Table"][[2]];
datp002R6h01MSMST = Partition[Riffle[Partition[Riffle[Range[9],
      datp002R6h01MSMS], 2], Map[ErrorBar, datp002R6h01MSMSCI]], 2];
datp002R6h05MSMST = Partition[Riffle[Partition[Riffle[Range[9],
      datp002R6h05MSMS], 2], Map[ErrorBar, datp002R6h05MSMSCI]], 2];
datp002R6h09MSMST = Partition[Riffle[Partition[Riffle[Range[9],
      datp002R6h09MSMS], 2], Map[ErrorBar, datp002R6h09MSMSCI]], 2];
```

Analytical solutions

```
p1R6 = ListPlot[{Table[{l, PLNH2[5000, 0.05, 0.1, 0, 6, 10, l, 0.02, 4]}, {l, 1, 9}],
   Table[{l, PLNH2[5000, 0.05, 0.5, 0, 6, 10, l, 0.02, 4]}, {l, 1, 9}],
   Table[{l, PLNH2[5000, 0.05, 0.9, 0, 6, 10, l, 0.02, 4]}, {l, 1, 9}],
   Table[{l, PrJ[l, 10]}, {l, 1, 9}]}, PlotRange → All,
  PlotStyle → {Red, Black, Blue, {Gray, Dashed}}, Joined → True,
  Ticks → {{1, 2, 3, 4, 5, 6, 7, 8, 9}, Automatic},
  BaseStyle → {FontWeight → "Bold", FontColor → Black, FontSize → 12}]
```

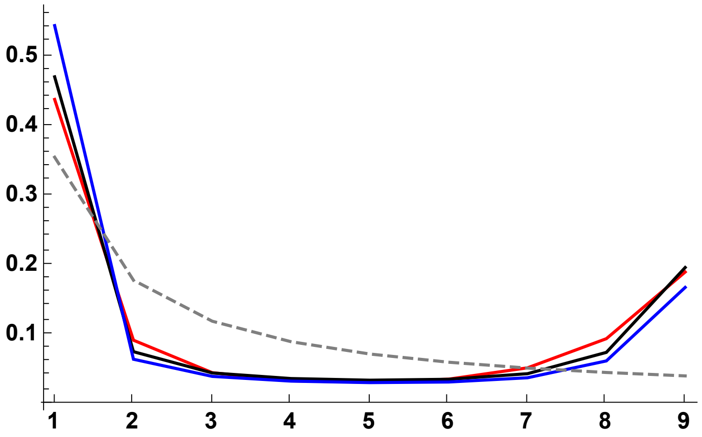


SLiM simulations plot

```
p2R6 = ErrorListPlot[
  {datp002R6h01T, datp002R6h05T, datp002R6h09T}, PlotRange → All, PlotStyle →
   {{Red, PointSize[0.02]}, {Black, PointSize[0.02]}, {Blue, PointSize[0.02]}}]
```

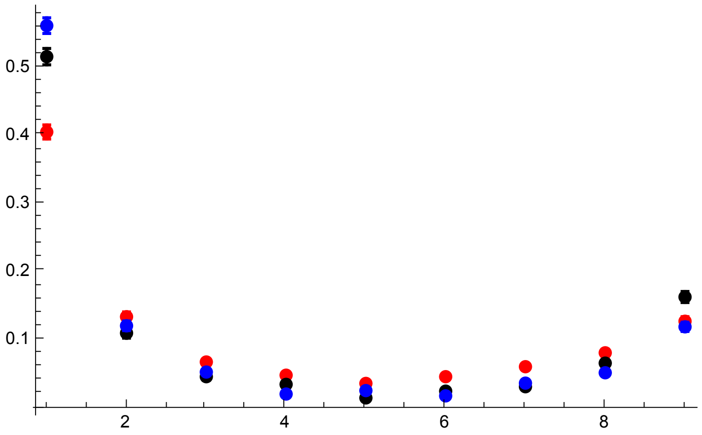


MSMS simulation results

```
p2AR6 = ErrorListPlot[{datp002R6h01MSMST, datp002R6h05MSMST, datp002R6h09MSMST},
  PlotRange → All, PlotStyle →
   {{Pink, PointSize[0.02]}, {Gray, PointSize[0.02]}, {Cyan, PointSize[0.02]}}]
```

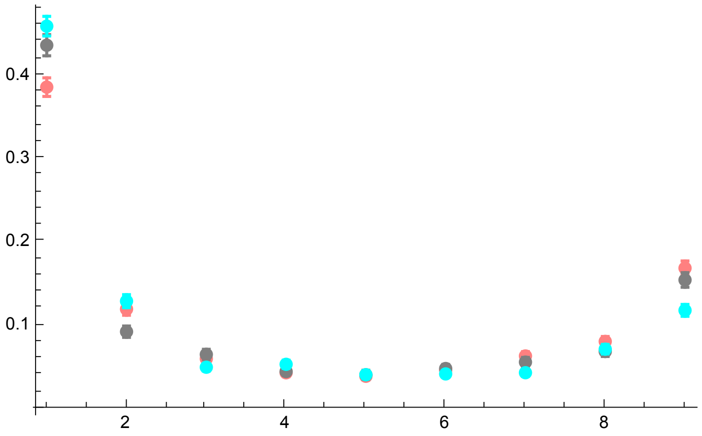


All together:

```
p3R6s0p02 = Show[p1R6, p2R6, p2AR6, PlotRange → All, ImageSize → 325]
```

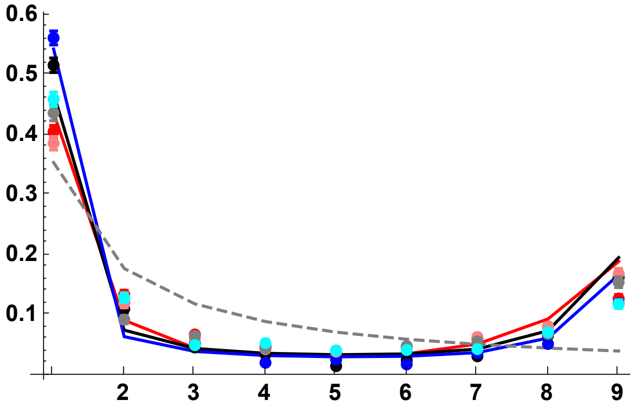


SLiM simulations and analytic results:

```
p3AR6s0p02 = Show[p1R6, p2R6, PlotRange → All, ImageSize → 325]
```

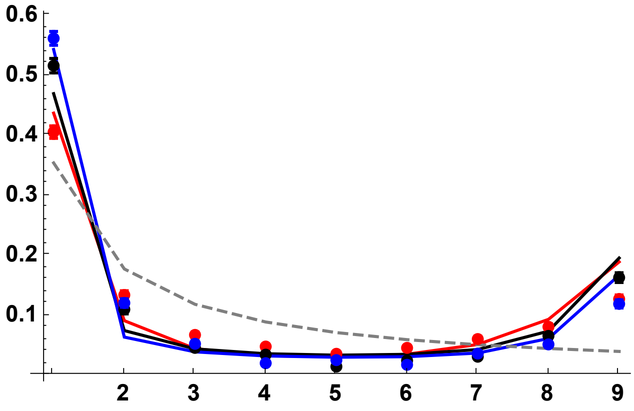


Simulation results only (SLiM and MSMS)

```
p3BR6 = Show[p2R6, p2AR6, PlotRange → All]
```

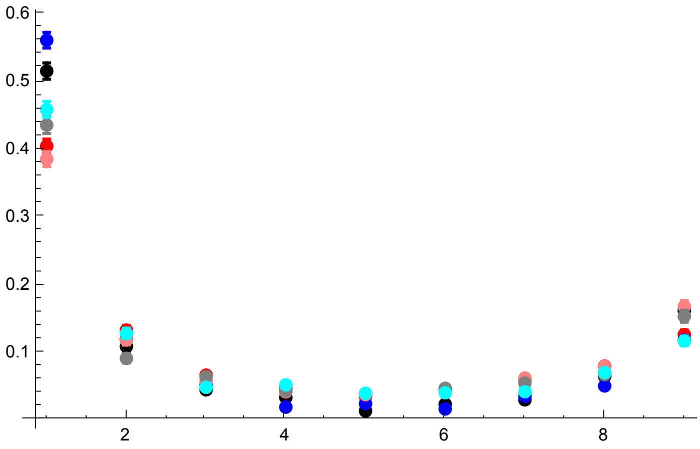


R = 18

SLiM simulations plot

```
datp002R18h01 = Import[
    "SLIM_SFS_F0/SFSTab_R_18_h_0.1_self_0_f_0.02_10b_SLIM.dat", "Table"][[1]];
datp002R18h01CI = Import[
    "SLIM_SFS_F0/SFSTab_R_18_h_0.1_self_0_f_0.02_10b_SLIM.dat", "Table"][[2]];
datp002R18h05 = Import[
    "SLIM_SFS_F0/SFSTab_R_18_h_0.5_self_0_f_0.02_10b_SLIM.dat", "Table"][[1]];
datp002R18h05CI = Import[
    "SLIM_SFS_F0/SFSTab_R_18_h_0.5_self_0_f_0.02_10b_SLIM.dat", "Table"][[2]];
datp002R18h09 = Import[
    "SLIM_SFS_F0/SFSTab_R_18_h_0.9_self_0_f_0.02_10b_SLIM.dat", "Table"][[1]];
datp002R18h09CI = Import[
    "SLIM_SFS_F0/SFSTab_R_18_h_0.9_self_0_f_0.02_10b_SLIM.dat", "Table"][[2]];
datp002R18h01T = Partition[Riffle[Partition[Riffle[Range[9], datp002R18h01], 2],
    Map[ErrorBar, datp002R18h01CI]], 2];
datp002R18h05T = Partition[Riffle[Partition[Riffle[Range[9], datp002R18h05], 2],
    Map[ErrorBar, datp002R18h05CI]], 2];
datp002R18h09T = Partition[Riffle[Partition[Riffle[Range[9], datp002R18h09], 2],
    Map[ErrorBar, datp002R18h09CI]], 2];
```

MSMS simulation results

```
datp002R18h01MSMS =
  Import["SFS_MSMS_SV_5K/SFSTab_R_36_h_0.1_self_0_f_0.02_10b_MSMS.dat",
    "Table"][[1]];
datp002R18h01MSMSCI = Import[
    "SFS_MSMS_SV_5K/SFSTab_R_36_h_0.1_self_0_f_0.02_10b_MSMS.dat",
    "Table"][[2]];
datp002R18h05MSMS = Import[
    "SFS_MSMS_SV_5K/SFSTab_R_36_h_0.5_self_0_f_0.02_10b_MSMS.dat",
    "Table"][[1]];
datp002R18h05MSMSCI = Import[
    "SFS_MSMS_SV_5K/SFSTab_R_36_h_0.5_self_0_f_0.02_10b_MSMS.dat",
    "Table"][[2]];
datp002R18h09MSMS = Import[
    "SFS_MSMS_SV_5K/SFSTab_R_36_h_0.9_self_0_f_0.02_10b_MSMS.dat",
    "Table"][[1]];
datp002R18h09MSMSCI = Import[
    "SFS_MSMS_SV_5K/SFSTab_R_36_h_0.9_self_0_f_0.02_10b_MSMS.dat",
    "Table"][[2]];
datp002R18h01MSMST = Partition[Riffle[Partition[Riffle[Range[9],
      datp002R18h01MSMS], 2], Map[ErrorBar, datp002R18h01MSMSCI]], 2];
datp002R18h05MSMST = Partition[Riffle[Partition[Riffle[Range[9],
      datp002R18h05MSMS], 2], Map[ErrorBar, datp002R18h05MSMSCI]], 2];
datp002R18h09MSMST = Partition[Riffle[Partition[Riffle[Range[9],
      datp002R18h09MSMS], 2], Map[ErrorBar, datp002R18h09MSMSCI]], 2];
```

Analytic plot

```
p1R18 =
 ListPlot[{Table[{l, PLNH2[5000, 0.05, 0.1, 0, 18, 10, l, 0.02, 4]}, {l, 1, 9}],
   Table[{l, PLNH2[5000, 0.05, 0.5, 0, 18, 10, l, 0.02, 4]}, {l, 1, 9}],
   Table[{l, PLNH2[5000, 0.05, 0.9, 0, 18, 10, l, 0.02, 4]}, {l, 1, 9}],
   Table[{l, PrJ[l, 10]}, {l, 1, 9}]}, PlotRange → All,
  PlotStyle → {Red, Black, Blue, {Gray, Dashed}}, Joined → True,
  Ticks → {{1, 2, 3, 4, 5, 6, 7, 8, 9}, Automatic},
  BaseStyle → {FontWeight → "Bold", FontColor → Black, FontSize → 12}]
```

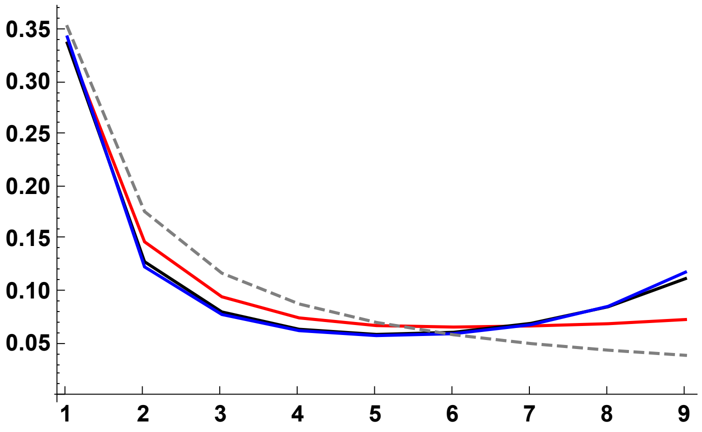


SLiM simulations plot

```
p2R18 = ErrorListPlot[
  {datp002R18h01T, datp002R18h05T, datp002R18h09T}, PlotRange → All, PlotStyle →
   {{Red, PointSize[0.02]}, {Black, PointSize[0.02]}, {Blue, PointSize[0.02]}}]
```

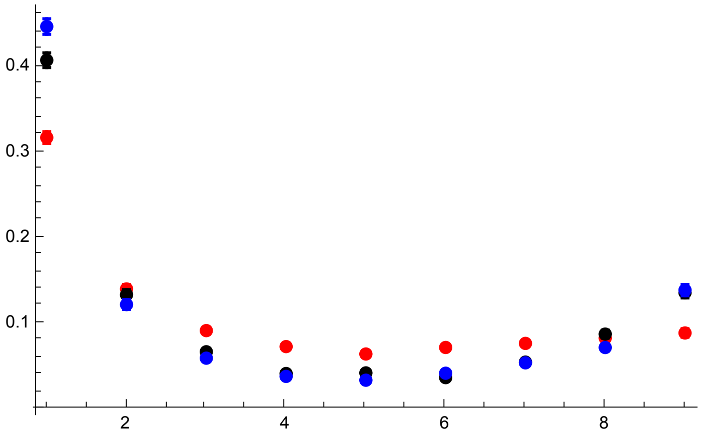


MSMS simulation results

```
p2AR18 = ErrorListPlot[{datp002R18h01MSMST,
   datp002R18h05MSMST, datp002R18h09MSMST}, PlotRange → All, PlotStyle →
   {{Pink, PointSize[0.02]}, {Gray, PointSize[0.02]}, {Cyan, PointSize[0.02]}}]
```

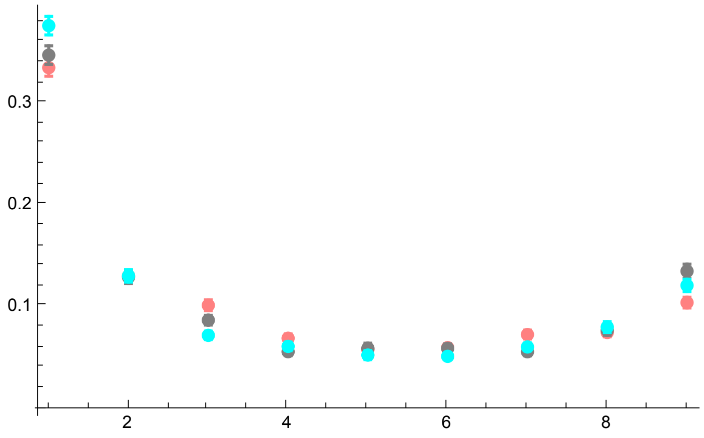


All together

```
p3R18s0p02 = Show[p1R18, p2R18, p2AR18, PlotRange → All, ImageSize → 325]
```

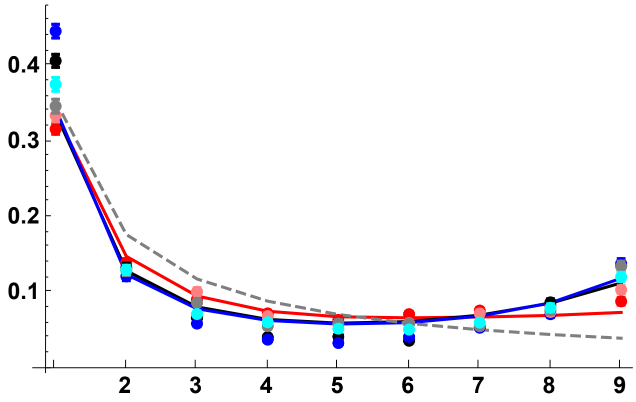


Simulation results only (SLiM and MSMS)

```
p3BR18 = Show[p2R18, p2AR18, PlotRange → All]
```

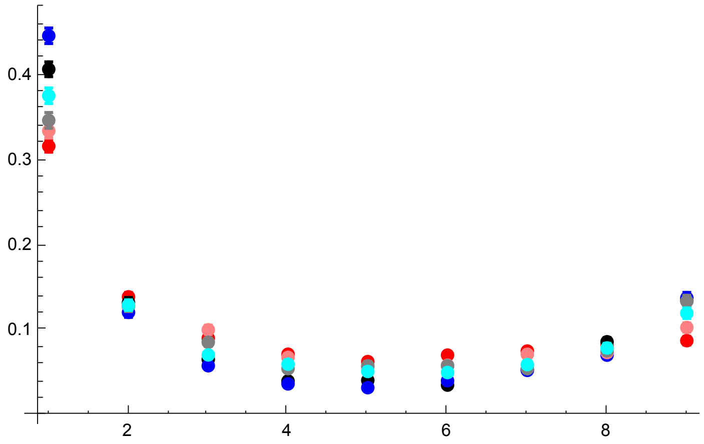


R = 30

SLiM simulations plot

```
datp002R30h01 = Import[
    "SLIM_SFS_F0/SFSTab_R_30_h_0.1_self_0_f_0.02_10b_SLIM.dat", "Table"][[1]];
datp002R30h01CI = Import[
    "SLIM_SFS_F0/SFSTab_R_30_h_0.1_self_0_f_0.02_10b_SLIM.dat", "Table"][[2]];
datp002R30h05 = Import[
    "SLIM_SFS_F0/SFSTab_R_30_h_0.5_self_0_f_0.02_10b_SLIM.dat", "Table"][[1]];
datp002R30h05CI = Import[
    "SLIM_SFS_F0/SFSTab_R_30_h_0.5_self_0_f_0.02_10b_SLIM.dat", "Table"][[2]];
datp002R30h09 = Import[
    "SLIM_SFS_F0/SFSTab_R_30_h_0.9_self_0_f_0.02_10b_SLIM.dat", "Table"][[1]];
datp002R30h09CI = Import[
    "SLIM_SFS_F0/SFSTab_R_30_h_0.9_self_0_f_0.02_10b_SLIM.dat", "Table"][[2]];
datp002R30h01T = Partition[Riffle[Partition[Riffle[Range[9], datp002R30h01], 2],
    Map[ErrorBar, datp002R30h01CI]], 2];
datp002R30h05T = Partition[Riffle[Partition[Riffle[Range[9], datp002R30h05], 2],
    Map[ErrorBar, datp002R30h05CI]], 2];
datp002R30h09T = Partition[Riffle[Partition[Riffle[Range[9], datp002R30h09], 2],
    Map[ErrorBar, datp002R30h09CI]], 2];
```

MSMS simulation results

```
datp002R30h01MSMS =
  Import["SFS_MSMS_SV_5K/SFSTab_R_60_h_0.1_self_0_f_0.02_10b_MSMS.dat",
    "Table"][[1]];
datp002R30h01MSMSCI = Import[
    "SFS_MSMS_SV_5K/SFSTab_R_60_h_0.1_self_0_f_0.02_10b_MSMS.dat",
    "Table"][[2]];
datp002R30h05MSMS = Import[
    "SFS_MSMS_SV_5K/SFSTab_R_60_h_0.5_self_0_f_0.02_10b_MSMS.dat",
    "Table"][[1]];
datp002R30h05MSMSCI = Import[
    "SFS_MSMS_SV_5K/SFSTab_R_60_h_0.5_self_0_f_0.02_10b_MSMS.dat",
    "Table"][[2]];
datp002R30h09MSMS = Import[
    "SFS_MSMS_SV_5K/SFSTab_R_60_h_0.9_self_0_f_0.02_10b_MSMS.dat",
    "Table"][[1]];
datp002R30h09MSMSCI = Import[
    "SFS_MSMS_SV_5K/SFSTab_R_60_h_0.9_self_0_f_0.02_10b_MSMS.dat",
    "Table"][[2]];
datp002R30h01MSMST = Partition[Riffle[Partition[Riffle[Range[9],
      datp002R30h01MSMS], 2], Map[ErrorBar, datp002R30h01MSMSCI]], 2];
datp002R30h05MSMST = Partition[Riffle[Partition[Riffle[Range[9],
      datp002R30h05MSMS], 2], Map[ErrorBar, datp002R30h05MSMSCI]], 2];
datp002R30h09MSMST = Partition[Riffle[Partition[Riffle[Range[9],
      datp002R30h09MSMS], 2], Map[ErrorBar, datp002R30h09MSMSCI]], 2];
```

Analytic res

```
p1R30 =
 ListPlot[{Table[{l, PLNH2[5000, 0.05, 0.1, 0, 30, 10, l, 0.02, 4]}, {l, 1, 9}],
   Table[{l, PLNH2[5000, 0.05, 0.5, 0, 30, 10, l, 0.02, 4]}, {l, 1, 9}],
   Table[{l, PLNH2[5000, 0.05, 0.9, 0, 30, 10, l, 0.02, 4]}, {l, 1, 9}],
   Table[{l, PrJ[l, 10]}, {l, 1, 9}]}, PlotRange → All,
  PlotStyle → {Red, Black, Blue, {Gray, Dashed}}, Joined → True]
```

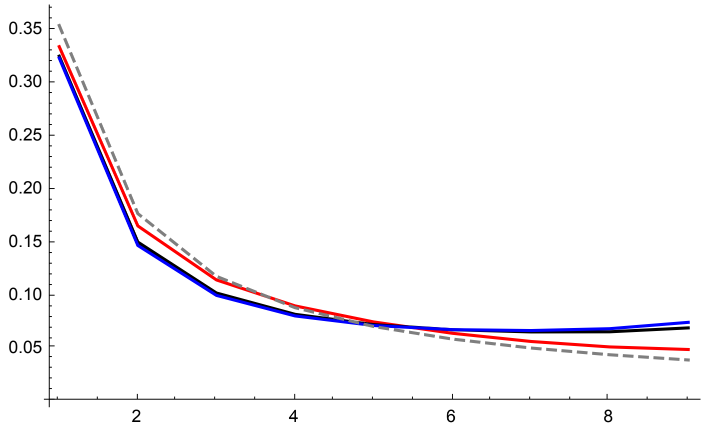


SLiM simulations plot

```
p2R30 = ErrorListPlot[
  {datp002R30h01T, datp002R30h05T, datp002R30h09T}, PlotRange → All, PlotStyle →
   {{Red, PointSize[0.02]}, {Black, PointSize[0.02]}, {Blue, PointSize[0.02]}}]
```

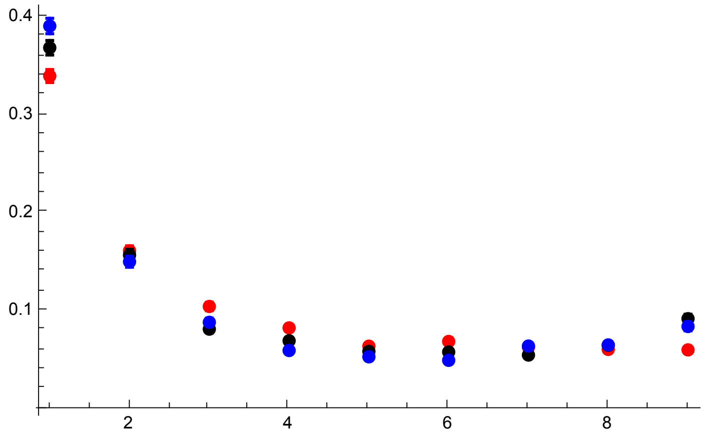


MSMS simulation results

```
p2AR30 = ErrorListPlot[{datp002R30h01MSMST,
    datp002R30h05MSMST, datp002R30h09MSMST}, PlotRange → All, PlotStyle →
    {{Pink, PointSize[0.02]}, {Gray, PointSize[0.02]}, {Cyan, PointSize[0.02]}}]
```

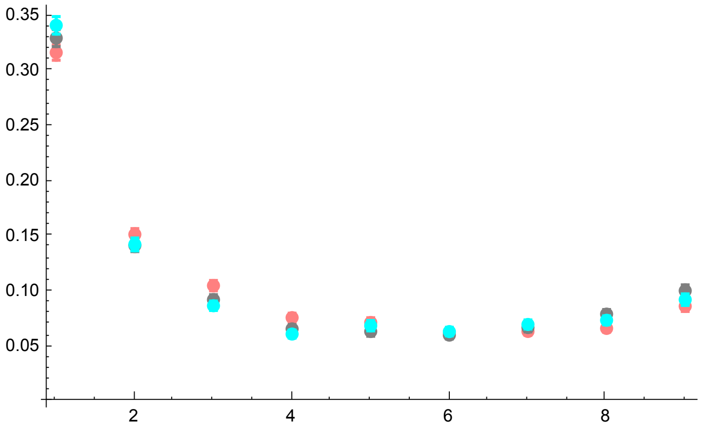


All together

```
p3R30 = Show[p1R30, p2R30, p2AR30, PlotRange → All]
```

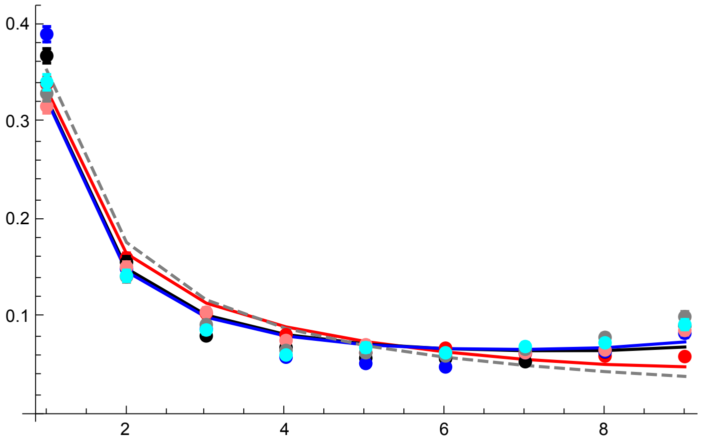


Simulation results only (SLiM and MSMS)

```
p3BR30 = Show[p2R30, p2AR30, PlotRange → All]
```

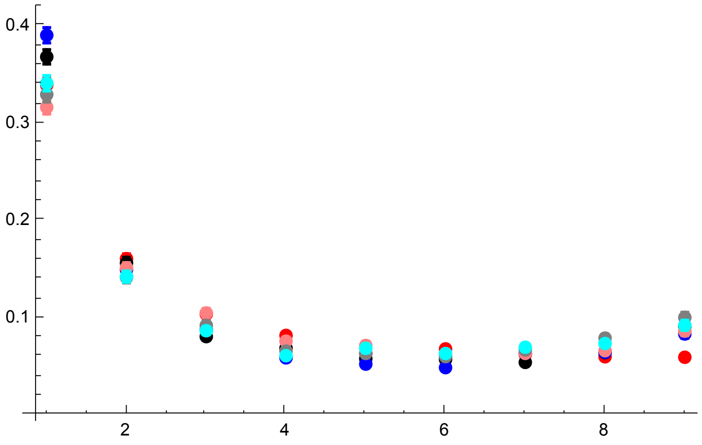


R = 42

Original sim data

```
datp002R42h01 = Import[
    "SLIM_SFS_F0/SFSTab_R_42_h_0.1_self_0_f_0.02_10b_SLIM.dat", "Table"][[1]];
datp002R42h01CI = Import[
    "SLIM_SFS_F0/SFSTab_R_42_h_0.1_self_0_f_0.02_10b_SLIM.dat", "Table"][[2]];
datp002R42h05 = Import[
    "SLIM_SFS_F0/SFSTab_R_42_h_0.5_self_0_f_0.02_10b_SLIM.dat", "Table"][[1]];
datp002R42h05CI = Import[
    "SLIM_SFS_F0/SFSTab_R_42_h_0.5_self_0_f_0.02_10b_SLIM.dat", "Table"][[2]];
datp002R42h09 = Import[
    "SLIM_SFS_F0/SFSTab_R_42_h_0.9_self_0_f_0.02_10b_SLIM.dat", "Table"][[1]];
datp002R42h09CI = Import[
    "SLIM_SFS_F0/SFSTab_R_42_h_0.9_self_0_f_0.02_10b_SLIM.dat", "Table"][[2]];
datp002R42h01T = Partition[Riffle[Partition[Riffle[Range[9], datp002R42h01], 2],
    Map[ErrorBar, datp002R42h01CI]], 2];
datp002R42h05T = Partition[Riffle[Partition[Riffle[Range[9], datp002R42h05], 2],
    Map[ErrorBar, datp002R42h05CI]], 2];
datp002R42h09T = Partition[Riffle[Partition[Riffle[Range[9], datp002R42h09], 2],
    Map[ErrorBar, datp002R42h09CI]], 2];
```

MSMS Sim data

```
datp002R42h01MSMS =
  Import["SFS_MSMS_SV_5K/SFSTab_R_84_h_0.1_self_0_f_0.02_10b_MSMS.dat",
    "Table"][[1]];
datp002R42h01MSMSCI = Import[
    "SFS_MSMS_SV_5K/SFSTab_R_84_h_0.1_self_0_f_0.02_10b_MSMS.dat",
    "Table"][[2]];
datp002R42h05MSMS = Import[
    "SFS_MSMS_SV_5K/SFSTab_R_84_h_0.5_self_0_f_0.02_10b_MSMS.dat",
    "Table"][[1]];
datp002R42h05MSMSCI = Import[
    "SFS_MSMS_SV_5K/SFSTab_R_84_h_0.5_self_0_f_0.02_10b_MSMS.dat",
    "Table"][[2]];
datp002R42h09MSMS = Import[
    "SFS_MSMS_SV_5K/SFSTab_R_84_h_0.9_self_0_f_0.02_10b_MSMS.dat",
    "Table"][[1]];
datp002R42h09MSMSCI = Import[
    "SFS_MSMS_SV_5K/SFSTab_R_84_h_0.9_self_0_f_0.02_10b_MSMS.dat",
    "Table"][[2]];
datp002R42h01MSMST = Partition[Riffle[Partition[Riffle[Range[9],
      datp002R42h01MSMS], 2], Map[ErrorBar, datp002R42h01MSMSCI]], 2];
datp002R42h05MSMST = Partition[Riffle[Partition[Riffle[Range[9],
      datp002R42h05MSMS], 2], Map[ErrorBar, datp002R42h05MSMSCI]], 2];
datp002R42h09MSMST = Partition[Riffle[Partition[Riffle[Range[9],
      datp002R42h09MSMS], 2], Map[ErrorBar, datp002R42h09MSMSCI]], 2];
```

Analytic res

```
p1R42 =
 ListPlot[{Table[{l, PLNH2[5000, 0.05, 0.1, 0, 42, 10, l, 0.02, 4]}, {l, 1, 9}],
   Table[{l, PLNH2[5000, 0.05, 0.5, 0, 42, 10, l, 0.02, 4]}, {l, 1, 9}],
   Table[{l, PLNH2[5000, 0.05, 0.9, 0, 42, 10, l, 0.02, 4]}, {l, 1, 9}],
   Table[{l, PrJ[l, 10]}, {l, 1, 9}]}, PlotRange → All,
  PlotStyle → {Red, Black, Blue, {Gray, Dashed}}, Joined → True]
```

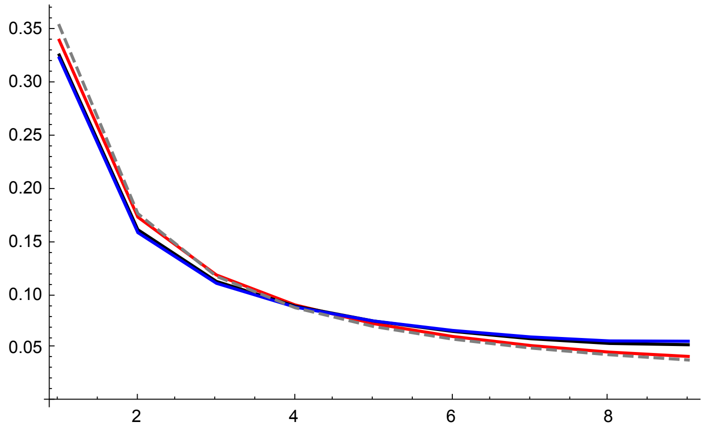


SLiM simulations plot

```
p2R42 = ErrorListPlot[
  {datp002R42h01T, datp002R42h05T, datp002R42h09T}, PlotRange → All, PlotStyle →
   {{Red, PointSize[0.02]}, {Black, PointSize[0.02]}, {Blue, PointSize[0.02]}}]
```

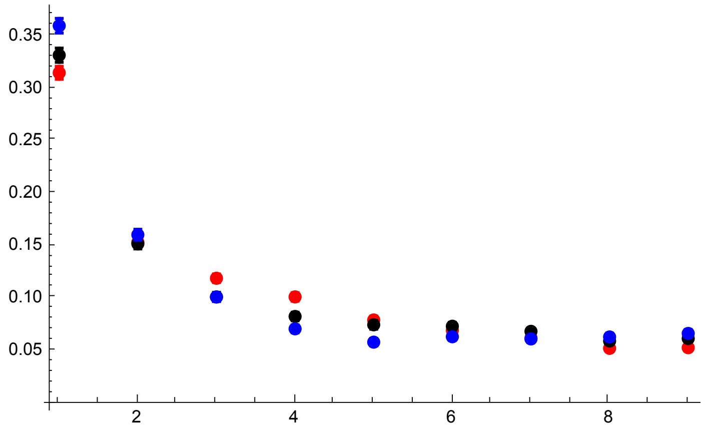


MSMS simulation results

```
p2AR42 = ErrorListPlot[{datp002R42h01MSMST,
    datp002R42h05MSMST, datp002R42h09MSMST}, PlotRange → All, PlotStyle →
    {{Pink, PointSize[0.02]}, {Gray, PointSize[0.02]}, {Cyan, PointSize[0.02]}}]
```

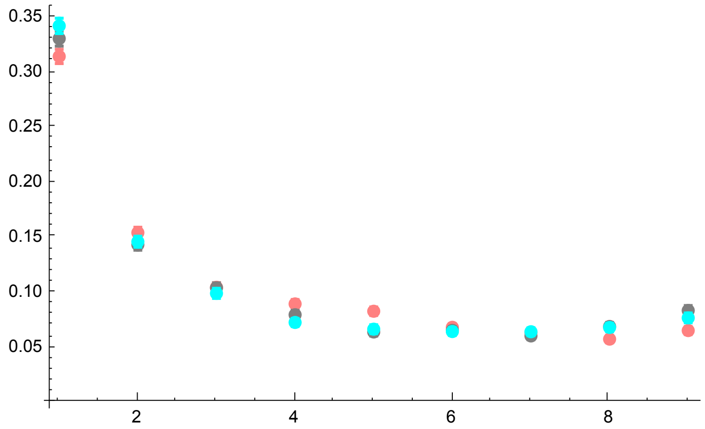


All together

```
p3R42 = Show[p1R42, p2R42, p2AR42, PlotRange → All]
```

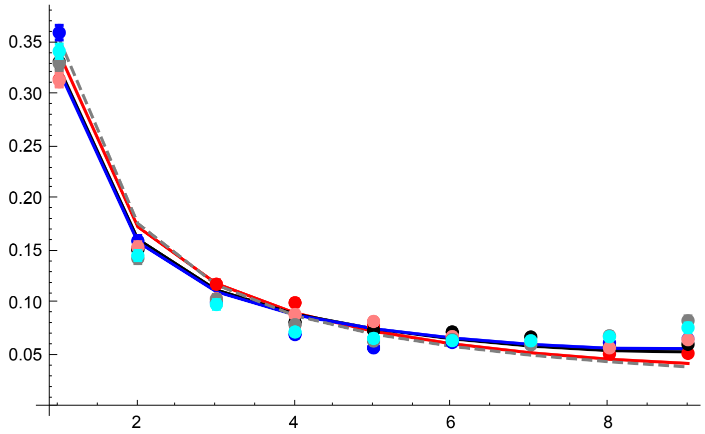


Simulation results only (SLiM and MSMS)

```
p3BR42 = Show[p2R42, p2AR42, PlotRange → All]
```

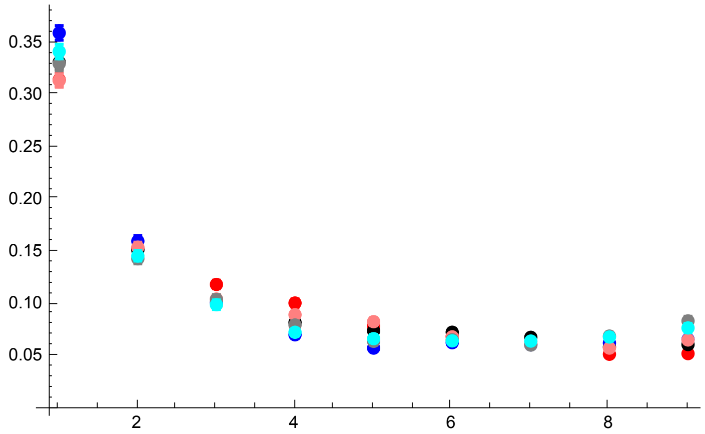


All plots compared

```
GraphicsGrid[{{p3R6s0p02, p3R18s0p02}, {p3R30, p3R42}}]
```

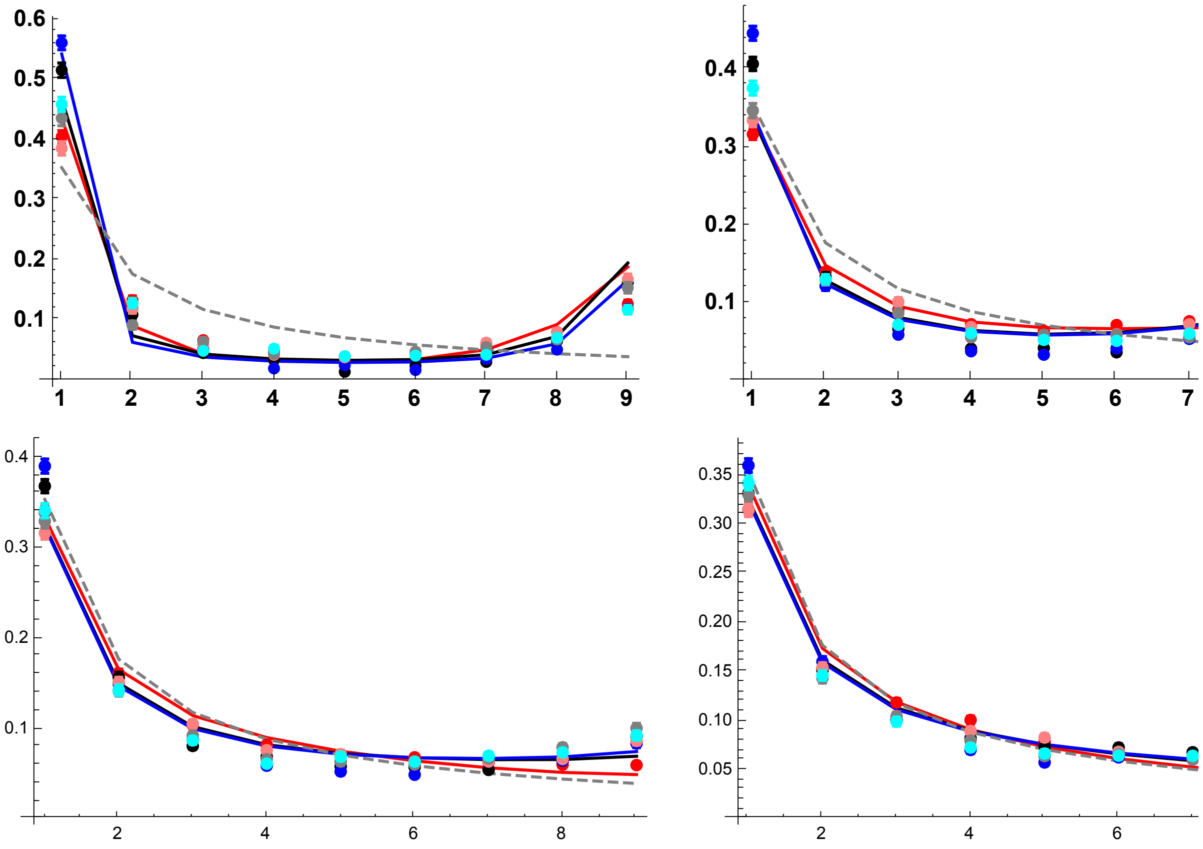


#### Results with $p_0 = 0.05$

R = 6

Original sim results

```
datp005R6h01 = Import[
    "SLIM_SFS_F0/SFSTab_R_6_h_0.1_self_0_f_0.05_10b_SLIM.dat", "Table"][[1]];
datp005R6h01CI = Import[
    "SLIM_SFS_F0/SFSTab_R_6_h_0.1_self_0_f_0.05_10b_SLIM.dat", "Table"][[2]];
datp005R6h05 = Import["SLIM_SFS_F0/SFSTab_R_6_h_0.5_self_0_f_0.05_10b_SLIM.dat",
    "Table"][[1]];
datp005R6h05CI = Import[
    "SLIM_SFS_F0/SFSTab_R_6_h_0.5_self_0_f_0.05_10b_SLIM.dat", "Table"][[2]];
datp005R6h09 = Import["SLIM_SFS_F0/SFSTab_R_6_h_0.9_self_0_f_0.05_10b_SLIM.dat",
    "Table"][[1]];
datp005R6h09CI = Import[
    "SLIM_SFS_F0/SFSTab_R_6_h_0.9_self_0_f_0.05_10b_SLIM.dat", "Table"][[2]];
datp005R6h01T = Partition[Riffle[Partition[Riffle[Range[9], datp005R6h01], 2],
    Map[ErrorBar, datp005R6h01CI]], 2];
datp005R6h05T = Partition[Riffle[Partition[Riffle[Range[9], datp005R6h05], 2],
    Map[ErrorBar, datp005R6h05CI]], 2];
datp005R6h09T = Partition[Riffle[Partition[Riffle[Range[9], datp005R6h09], 2],
    Map[ErrorBar, datp005R6h09CI]], 2];
```

MSMS Sim results

```
datp005R6h01MSMS =
  Import["SFS_MSMS_SV_5K/SFSTab_R_12_h_0.1_self_0_f_0.05_10b_MSMS.dat",
    "Table"][[1]];
datp005R6h01MSMSCI = Import[
    "SFS_MSMS_SV_5K/SFSTab_R_12_h_0.1_self_0_f_0.05_10b_MSMS.dat",
    "Table"][[2]];
datp005R6h05MSMS = Import[
    "SFS_MSMS_SV_5K/SFSTab_R_12_h_0.5_self_0_f_0.05_10b_MSMS.dat",
    "Table"][[1]];
datp005R6h05MSMSCI = Import[
    "SFS_MSMS_SV_5K/SFSTab_R_12_h_0.5_self_0_f_0.05_10b_MSMS.dat",
    "Table"][[2]];
datp005R6h09MSMS = Import[
    "SFS_MSMS_SV_5K/SFSTab_R_12_h_0.9_self_0_f_0.05_10b_MSMS.dat",
    "Table"][[1]];
datp005R6h09MSMSCI = Import[
    "SFS_MSMS_SV_5K/SFSTab_R_12_h_0.9_self_0_f_0.05_10b_MSMS.dat",
    "Table"][[2]];
datp005R6h01MSMST = Partition[Riffle[Partition[Riffle[Range[9],
      datp005R6h01MSMS], 2], Map[ErrorBar, datp005R6h01MSMSCI]], 2];
datp005R6h05MSMST = Partition[Riffle[Partition[Riffle[Range[9],
      datp005R6h05MSMS], 2], Map[ErrorBar, datp005R6h05MSMSCI]], 2];
datp005R6h09MSMST = Partition[Riffle[Partition[Riffle[Range[9],
      datp005R6h09MSMS], 2], Map[ErrorBar, datp005R6h09MSMSCI]], 2];
```

```
p1R6 = ListPlot[{Table[{l, PLNH2[5000, 0.05, 0.1, 0, 6, 10, l, 0.05, 4]}, {l, 1, 9}],
   Table[{l, PLNH2[5000, 0.05, 0.5, 0, 6, 10, l, 0.05, 4]}, {l, 1, 9}],
   Table[{l, PLNH2[5000, 0.05, 0.9, 0, 6, 10, l, 0.05, 4]}, {l, 1, 9}],
   Table[{l, PrJ[l, 10]}, {l, 1, 9}]}, PlotRange → All,
  PlotStyle → {Red, Black, Blue, {Gray, Dashed}}, Joined → True,
  Ticks → {{1, 2, 3, 4, 5, 6, 7, 8, 9}, Automatic},
  BaseStyle → {FontWeight → "Bold", FontColor → Black, FontSize → 12}]
```

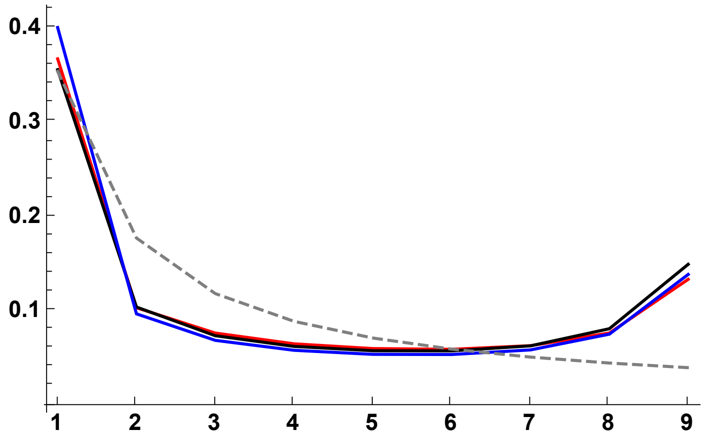


SLiM simulations plot

```
p2R6 = ErrorListPlot[
  {datp005R6h01T, datp005R6h05T, datp005R6h09T}, PlotRange → All, PlotStyle →
   {{Red, PointSize[0.02]}, {Black, PointSize[0.02]}, {Blue, PointSize[0.02]}}]
```

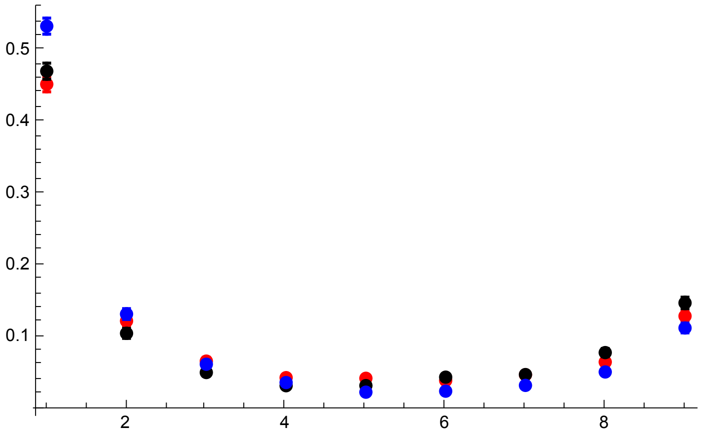


MSMS simulation results

```
p2AR6 = ErrorListPlot[{datp005R6h01MSMST, datp005R6h05MSMST, datp005R6h09MSMST},
  PlotRange → All, PlotStyle →
   {{Pink, PointSize[0.02]}, {Gray, PointSize[0.02]}, {Cyan, PointSize[0.02]}}]
```

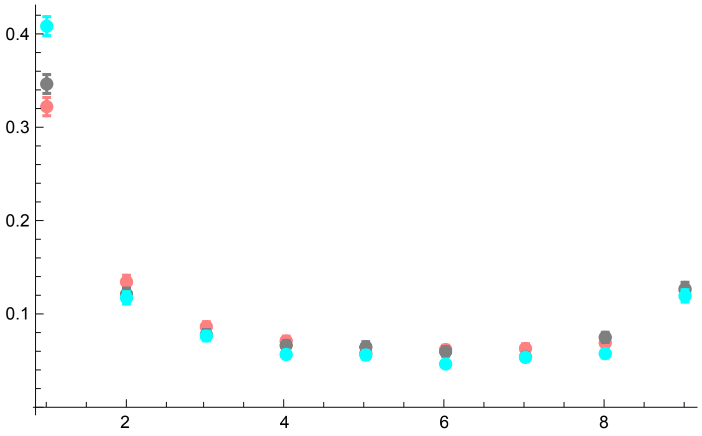


All together

```
p3R6s0p05 = Show[p1R6, p2R6, p2AR6, PlotRange → All, ImageSize → 325]
```

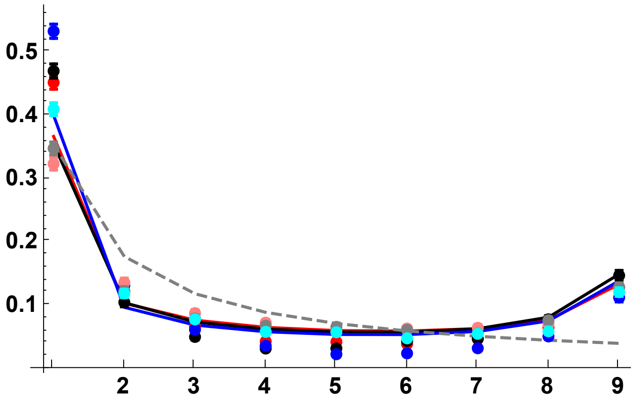


SLiM Simulations and analytical results

```
p3AR6s0p05 = Show[p1R6, p2R6, PlotRange → All, ImageSize → 325]
```

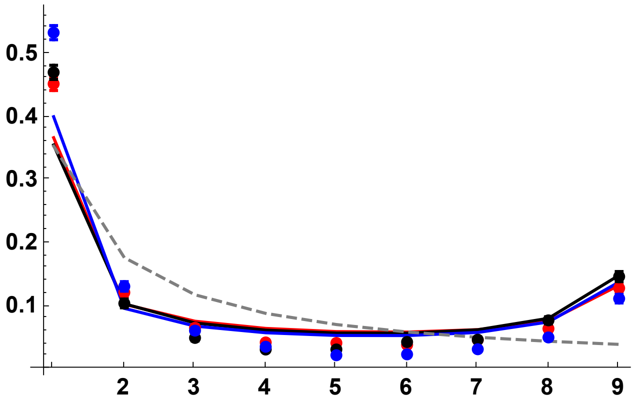


Simulation results only (SLiM and MSMS)

```
p3BR6 = Show[p2R6, p2AR6, PlotRange → All]
```

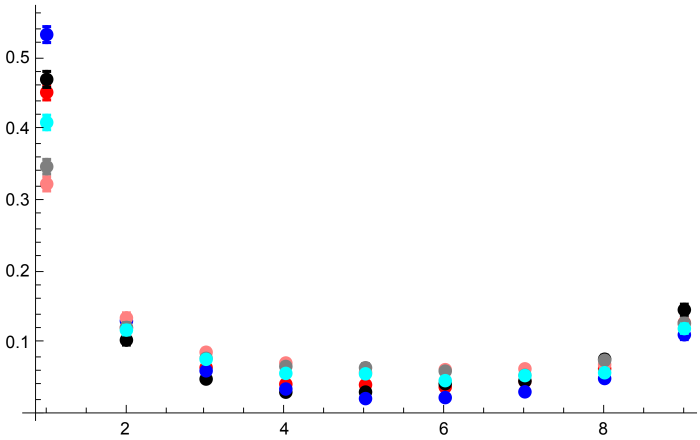


R = 18

SLiM simulations plot

```
datp005R18h01 = Import[
    "SLIM_SFS_F0/SFSTab_R_18_h_0.1_self_0_f_0.05_10b_SLIM.dat", "Table"][[1]];
datp005R18h01CI = Import[
    "SLIM_SFS_F0/SFSTab_R_18_h_0.1_self_0_f_0.05_10b_SLIM.dat", "Table"][[2]];
datp005R18h05 = Import[
    "SLIM_SFS_F0/SFSTab_R_18_h_0.5_self_0_f_0.05_10b_SLIM.dat", "Table"][[1]];
datp005R18h05CI = Import[
    "SLIM_SFS_F0/SFSTab_R_18_h_0.5_self_0_f_0.05_10b_SLIM.dat", "Table"][[2]];
datp005R18h09 = Import[
    "SLIM_SFS_F0/SFSTab_R_18_h_0.9_self_0_f_0.05_10b_SLIM.dat", "Table"][[1]];
datp005R18h09CI = Import[
    "SLIM_SFS_F0/SFSTab_R_18_h_0.9_self_0_f_0.05_10b_SLIM.dat", "Table"][[2]];
datp005R18h01T = Partition[Riffle[Partition[Riffle[Range[9], datp005R18h01], 2],
    Map[ErrorBar, datp005R18h01CI]], 2];
datp005R18h05T = Partition[Riffle[Partition[Riffle[Range[9], datp005R18h05], 2],
    Map[ErrorBar, datp005R18h05CI]], 2];
datp005R18h09T = Partition[Riffle[Partition[Riffle[Range[9], datp005R18h09], 2],
    Map[ErrorBar, datp005R18h09CI]], 2];
```

MSMS simulation results

```
datp005R18h01MSMS =
  Import["SFS_MSMS_SV_5K/SFSTab_R_36_h_0.1_self_0_f_0.05_10b_MSMS.dat",
    "Table"][[1]];
datp005R18h01MSMSCI = Import[
    "SFS_MSMS_SV_5K/SFSTab_R_36_h_0.1_self_0_f_0.05_10b_MSMS.dat",
    "Table"][[2]];
datp005R18h05MSMS = Import[
    "SFS_MSMS_SV_5K/SFSTab_R_36_h_0.5_self_0_f_0.05_10b_MSMS.dat",
    "Table"][[1]];
datp005R18h05MSMSCI = Import[
    "SFS_MSMS_SV_5K/SFSTab_R_36_h_0.5_self_0_f_0.05_10b_MSMS.dat",
    "Table"][[2]];
datp005R18h09MSMS = Import[
    "SFS_MSMS_SV_5K/SFSTab_R_36_h_0.9_self_0_f_0.05_10b_MSMS.dat",
    "Table"][[1]];
datp005R18h09MSMSCI = Import[
    "SFS_MSMS_SV_5K/SFSTab_R_36_h_0.9_self_0_f_0.05_10b_MSMS.dat",
    "Table"][[2]];
datp005R18h01MSMST = Partition[Riffle[Partition[Riffle[Range[9],
      datp005R18h01MSMS], 2], Map[ErrorBar, datp005R18h01MSMSCI]], 2];
datp005R18h05MSMST = Partition[Riffle[Partition[Riffle[Range[9],
      datp005R18h05MSMS], 2], Map[ErrorBar, datp005R18h05MSMSCI]], 2];
datp005R18h09MSMST = Partition[Riffle[Partition[Riffle[Range[9],
      datp005R18h09MSMS], 2], Map[ErrorBar, datp005R18h09MSMSCI]], 2];
```

Analytic results

```
p1R18 =
 ListPlot[{Table[{l, PLNH2[5000, 0.05, 0.1, 0, 18, 10, l, 0.05, 4]}, {l, 1, 9}],
    Table[{l, PLNH2[5000, 0.05, 0.5, 0, 18, 10, l, 0.05, 4]}, {l, 1, 9}],
    Table[{l, PLNH2[5000, 0.05, 0.9, 0, 18, 10, l, 0.05, 4]}, {l, 1, 9}],
    Table[{l, PrJ[l, 10]}, {l, 1, 9}]}, PlotRange → All,
  PlotStyle → {Red, Black, Blue, {Gray, Dashed}}, Joined → True,
  Ticks → {{1, 2, 3, 4, 5, 6, 7, 8, 9}, Automatic},
  BaseStyle → {FontWeight → "Bold", FontColor → Black, FontSize → 12}]
```

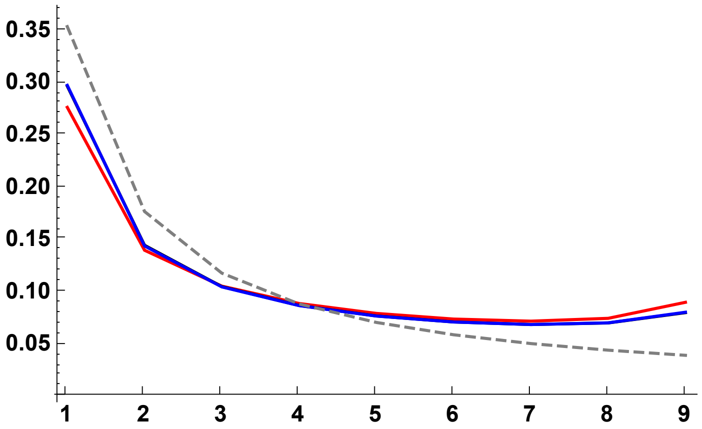


SLiM simulations plot

```
p2R18 = ErrorListPlot[
  {datp005R18h01T, datp005R18h05T, datp005R18h09T}, PlotRange → All, PlotStyle →
   {{Red, PointSize[0.02]}, {Black, PointSize[0.02]}, {Blue, PointSize[0.02]}}]
```

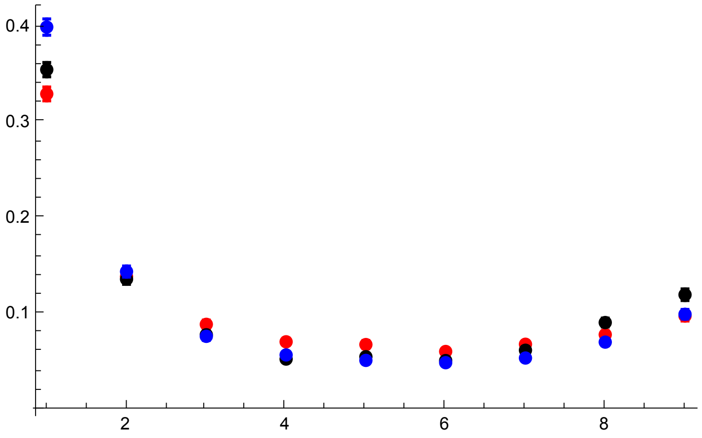


MSMS simulation results

```
p2AR18 = ErrorListPlot[{datp005R18h01MSMST,
   datp005R18h05MSMST, datp005R18h09MSMST}, PlotRange → All, PlotStyle →
   {{Pink, PointSize[0.02]}, {Gray, PointSize[0.02]}, {Cyan, PointSize[0.02]}}]
```

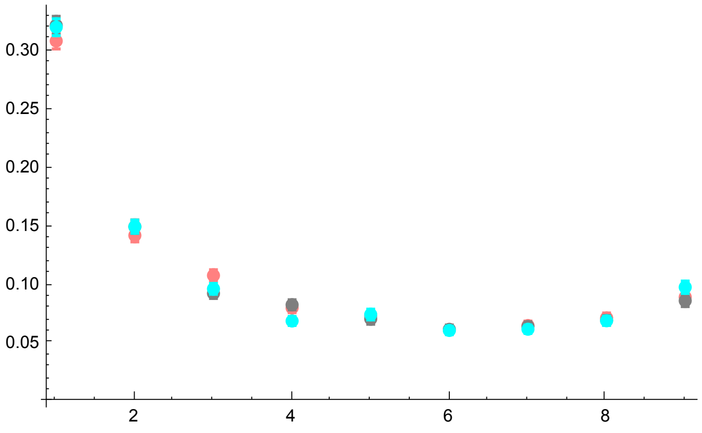


All together

```
p3R18s0p05 = Show[p1R18, p2R18, p2AR18, PlotRange → All, ImageSize → 325]
```

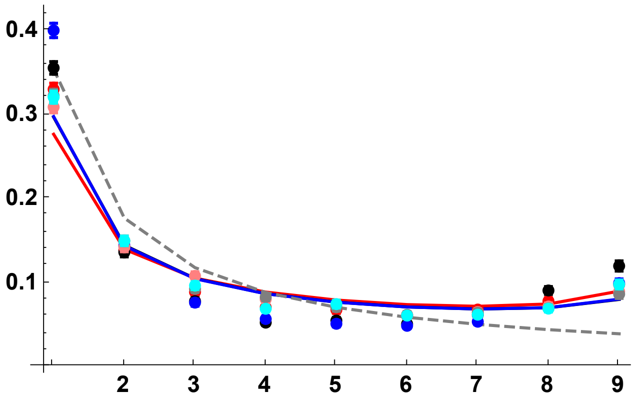


Simulation results only (SLiM and MSMS)

```
p3BR18 = Show[p2R18, p2AR18, PlotRange → All]
```

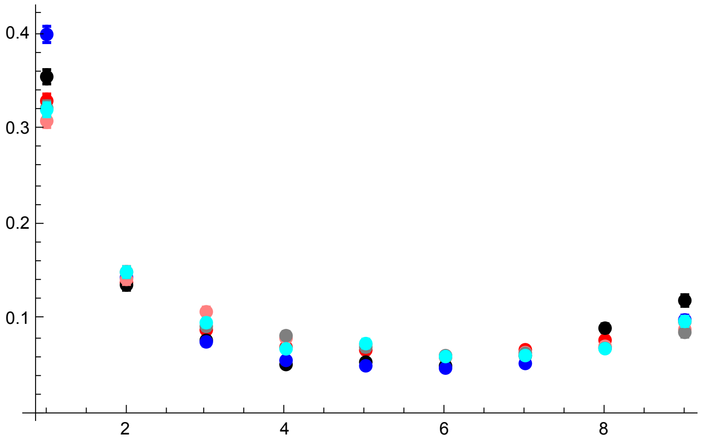


R = 30

SLiM simulations plot

```
datp005R30h01 = Import[
    "SLIM_SFS_F0/SFSTab_R_30_h_0.1_self_0_f_0.05_10b_SLIM.dat", "Table"][[1]];
datp005R30h01CI = Import[
    "SLIM_SFS_F0/SFSTab_R_30_h_0.1_self_0_f_0.05_10b_SLIM.dat", "Table"][[2]];
datp005R30h05 = Import[
    "SLIM_SFS_F0/SFSTab_R_30_h_0.5_self_0_f_0.05_10b_SLIM.dat", "Table"][[1]];
datp005R30h05CI = Import[
    "SLIM_SFS_F0/SFSTab_R_30_h_0.5_self_0_f_0.05_10b_SLIM.dat", "Table"][[2]];
datp005R30h09 = Import[
    "SLIM_SFS_F0/SFSTab_R_30_h_0.9_self_0_f_0.05_10b_SLIM.dat", "Table"][[1]];
datp005R30h09CI = Import[
    "SLIM_SFS_F0/SFSTab_R_30_h_0.9_self_0_f_0.05_10b_SLIM.dat", "Table"][[2]];
datp005R30h01T = Partition[Riffle[Partition[Riffle[Range[9], datp005R30h01], 2],
    Map[ErrorBar, datp005R30h01CI]], 2];
datp005R30h05T = Partition[Riffle[Partition[Riffle[Range[9], datp005R30h05], 2],
    Map[ErrorBar, datp005R30h05CI]], 2];
datp005R30h09T = Partition[Riffle[Partition[Riffle[Range[9], datp005R30h09], 2],
    Map[ErrorBar, datp005R30h09CI]], 2];
```

MSMS simulation results

```
datp005R30h01MSMS =
  Import["SFS_MSMS_SV_5K/SFSTab_R_60_h_0.1_self_0_f_0.05_10b_MSMS.dat",
    "Table"][[1]];
datp005R30h01MSMSCI = Import[
    "SFS_MSMS_SV_5K/SFSTab_R_60_h_0.1_self_0_f_0.05_10b_MSMS.dat",
    "Table"][[2]];
datp005R30h05MSMS = Import[
    "SFS_MSMS_SV_5K/SFSTab_R_60_h_0.5_self_0_f_0.05_10b_MSMS.dat",
    "Table"][[1]];
datp005R30h05MSMSCI = Import[
    "SFS_MSMS_SV_5K/SFSTab_R_60_h_0.5_self_0_f_0.05_10b_MSMS.dat",
    "Table"][[2]];
datp005R30h09MSMS = Import[
    "SFS_MSMS_SV_5K/SFSTab_R_60_h_0.9_self_0_f_0.05_10b_MSMS.dat",
    "Table"][[1]];
datp005R30h09MSMSCI = Import[
    "SFS_MSMS_SV_5K/SFSTab_R_60_h_0.9_self_0_f_0.05_10b_MSMS.dat",
    "Table"][[2]];
datp005R30h01MSMST = Partition[Riffle[Partition[Riffle[Range[9],
      datp005R30h01MSMS], 2], Map[ErrorBar, datp005R30h01MSMSCI]], 2];
datp005R30h05MSMST = Partition[Riffle[Partition[Riffle[Range[9],
      datp005R30h05MSMS], 2], Map[ErrorBar, datp005R30h05MSMSCI]], 2];
datp005R30h09MSMST = Partition[Riffle[Partition[Riffle[Range[9],
      datp005R30h09MSMS], 2], Map[ErrorBar, datp005R30h09MSMSCI]], 2];
```

```
p1R30 =
 ListPlot[{Table[{l, PLNH2[5000, 0.05, 0.1, 0, 30, 10, l, 0.05, 4]}, {l, 1, 9}],
   Table[{l, PLNH2[5000, 0.05, 0.5, 0, 30, 10, l, 0.05, 4]}, {l, 1, 9}],
   Table[{l, PLNH2[5000, 0.05, 0.9, 0, 30, 10, l, 0.05, 4]}, {l, 1, 9}],
   Table[{l, PrJ[l, 10]}, {l, 1, 9}]}, PlotRange → All,
  PlotStyle → {Red, Black, Blue, {Gray, Dashed}}, Joined → True]
```

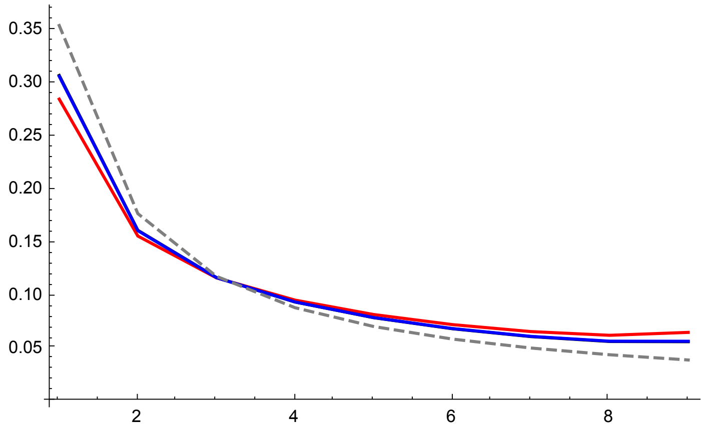


SLiM simulations plot

```
p2R30 = ErrorListPlot[
  {datp005R30h01T, datp005R30h05T, datp005R30h09T}, PlotRange → All, PlotStyle →
   {{Red, PointSize[0.02]}, {Black, PointSize[0.02]}, {Blue, PointSize[0.02]}}]
```

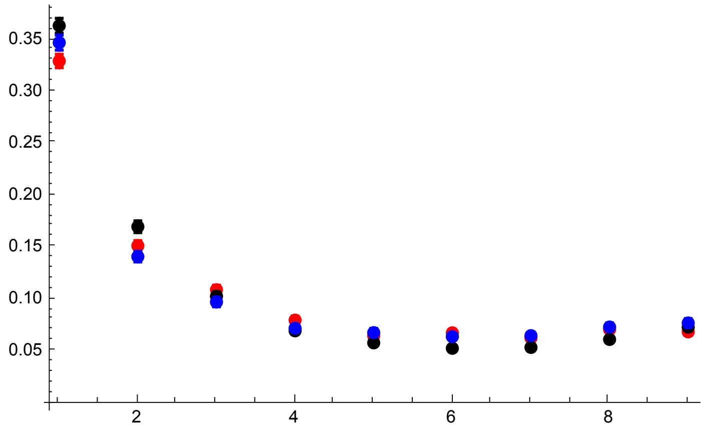


MSMS simulation results

```
p2AR30 = ErrorListPlot[{datp005R30h01MSMST,
   datp005R30h05MSMST, datp005R30h09MSMST}, PlotRange → All, PlotStyle →
   {{Pink, PointSize[0.02]}, {Gray, PointSize[0.02]}, {Cyan, PointSize[0.02]}}]
```

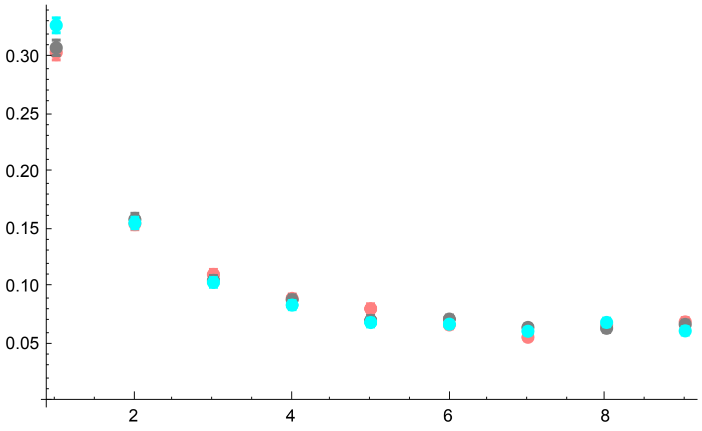


All together

```
p3R30 = Show[p1R30, p2R30, p2AR30, PlotRange → All]
```

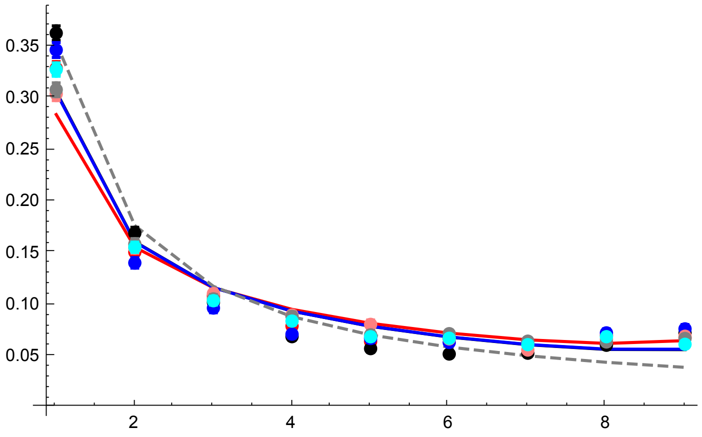


Simulation results only (SLiM and MSMS)

```
p3BR30 = Show[p2R30, p2AR30, PlotRange → All]
```

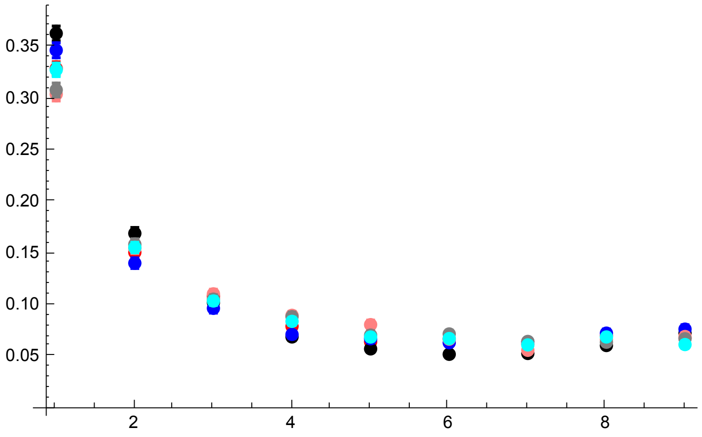


R = 42

Original sim data

```
datp005R42h01 = Import[
    "SLIM_SFS_F0/SFSTab_R_42_h_0.1_self_0_f_0.05_10b_SLIM.dat", "Table"][[1]];
datp005R42h01CI = Import[
    "SLIM_SFS_F0/SFSTab_R_42_h_0.1_self_0_f_0.05_10b_SLIM.dat", "Table"][[2]];
datp005R42h05 = Import[
    "SLIM_SFS_F0/SFSTab_R_42_h_0.5_self_0_f_0.05_10b_SLIM.dat", "Table"][[1]];
datp005R42h05CI = Import[
    "SLIM_SFS_F0/SFSTab_R_42_h_0.5_self_0_f_0.05_10b_SLIM.dat", "Table"][[2]];
datp005R42h09 = Import[
    "SLIM_SFS_F0/SFSTab_R_42_h_0.9_self_0_f_0.05_10b_SLIM.dat", "Table"][[1]];
datp005R42h09CI = Import[
    "SLIM_SFS_F0/SFSTab_R_42_h_0.9_self_0_f_0.05_10b_SLIM.dat", "Table"][[2]];
datp005R42h01T = Partition[Riffle[Partition[Riffle[Range[9], datp005R42h01], 2],
    Map[ErrorBar, datp005R42h01CI]], 2];
datp005R42h05T = Partition[Riffle[Partition[Riffle[Range[9], datp005R42h05], 2],
    Map[ErrorBar, datp005R42h05CI]], 2];
datp005R42h09T = Partition[Riffle[Partition[Riffle[Range[9], datp005R42h09], 2],
    Map[ErrorBar, datp005R42h09CI]], 2];
```

MSMS simulation results

```
datp005R42h01MSMS =
  Import["SFS_MSMS_SV_5K/SFSTab_R_84_h_0.1_self_0_f_0.05_10b_MSMS.dat",
    "Table"][[1]];
datp005R42h01MSMSCI = Import[
    "SFS_MSMS_SV_5K/SFSTab_R_84_h_0.1_self_0_f_0.05_10b_MSMS.dat",
    "Table"][[2]];
datp005R42h05MSMS = Import[
    "SFS_MSMS_SV_5K/SFSTab_R_84_h_0.5_self_0_f_0.05_10b_MSMS.dat",
    "Table"][[1]];
datp005R42h05MSMSCI = Import[
    "SFS_MSMS_SV_5K/SFSTab_R_84_h_0.5_self_0_f_0.05_10b_MSMS.dat",
    "Table"][[2]];
datp005R42h09MSMS = Import[
    "SFS_MSMS_SV_5K/SFSTab_R_84_h_0.9_self_0_f_0.05_10b_MSMS.dat",
    "Table"][[1]];
datp005R42h09MSMSCI = Import[
    "SFS_MSMS_SV_5K/SFSTab_R_84_h_0.9_self_0_f_0.05_10b_MSMS.dat",
    "Table"][[2]];
datp005R42h01MSMST = Partition[Riffle[Partition[Riffle[Range[9],
      datp005R42h01MSMS], 2], Map[ErrorBar, datp005R42h01MSMSCI]], 2];
datp005R42h05MSMST = Partition[Riffle[Partition[Riffle[Range[9],
      datp005R42h05MSMS], 2], Map[ErrorBar, datp005R42h05MSMSCI]], 2];
datp005R42h09MSMST = Partition[Riffle[Partition[Riffle[Range[9],
      datp005R42h09MSMS], 2], Map[ErrorBar, datp005R42h09MSMSCI]], 2];
```

Analytical solutions

```
p1R42 =
 ListPlot[{Table[{l, PLNH2[5000, 0.05, 0.1, 0, 42, 10, l, 0.05, 4]}, {l, 1, 9}],
   Table[{l, PLNH2[5000, 0.05, 0.5, 0, 42, 10, l, 0.05, 4]}, {l, 1, 9}],
   Table[{l, PLNH2[5000, 0.05, 0.9, 0, 42, 10, l, 0.05, 4]}, {l, 1, 9}],
   Table[{l, PrJ[l, 10]}, {l, 1, 9}]}, PlotRange → All,
  PlotStyle → {Red, Black, Blue, {Gray, Dashed}}, Joined → True]
```

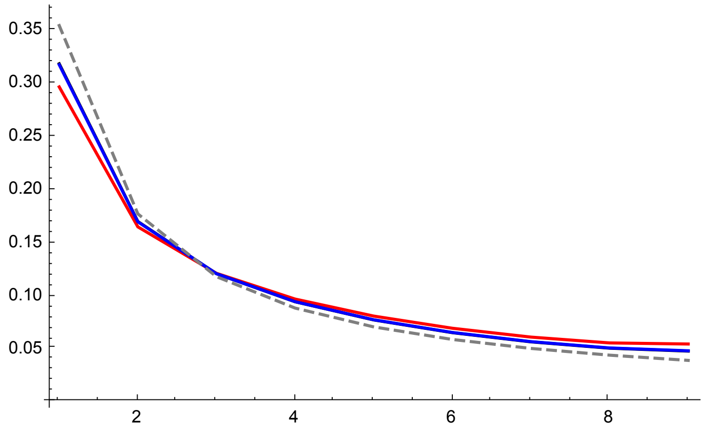


SLiM simulations plot

```
p2R42 = ErrorListPlot[
  {datp005R42h01T, datp005R42h05T, datp005R42h09T}, PlotRange → All, PlotStyle →
   {{Red, PointSize[0.02]}, {Black, PointSize[0.02]}, {Blue, PointSize[0.02]}}]
```

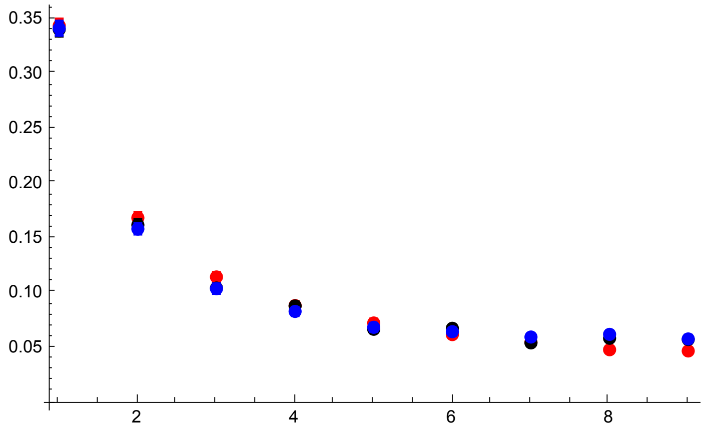


MSMS simulation results

```
p2AR42 = ErrorListPlot[{datp005R42h01MSMST,
    datp005R42h05MSMST, datp005R42h09MSMST}, PlotRange → All, PlotStyle →
    {{Pink, PointSize[0.02]}, {Gray, PointSize[0.02]}, {Cyan, PointSize[0.02]}}]
```

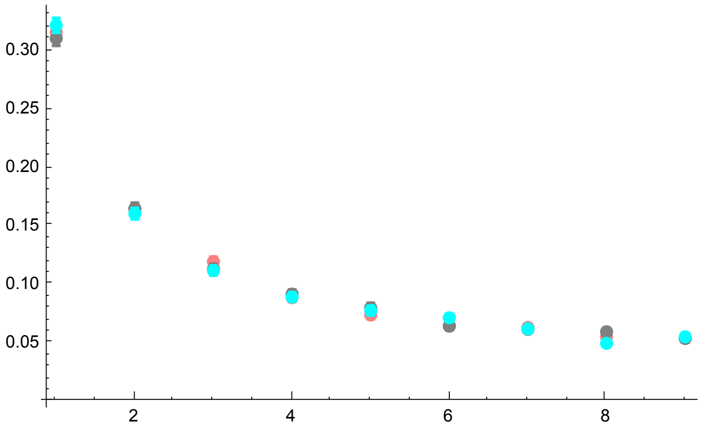


All together

```
p3R42 = Show[p1R42, p2R42, p2AR42, PlotRange → All]
```

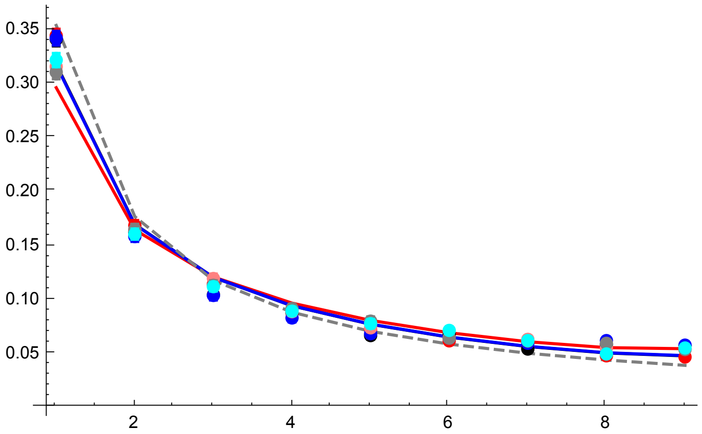


Simulation results only (SLiM and MSMS)

```
p3BR42 = Show[p2R42, p2AR42, PlotRange → All]
```

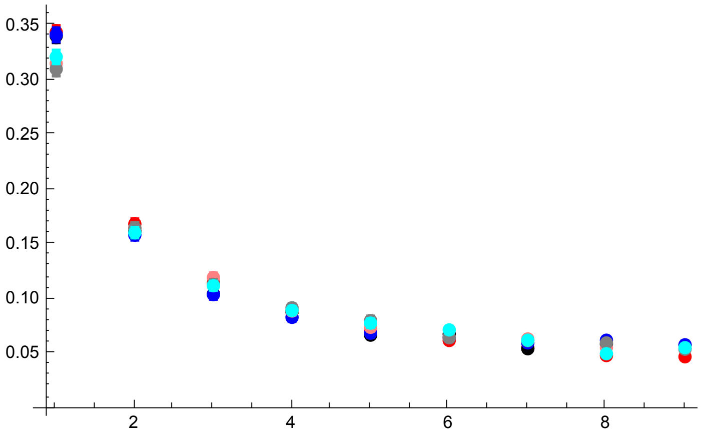


All plots compared

```
GraphicsGrid[{{p3R6s0p05, p3R18s0p05}, {p3R30, p3R42}}]
```

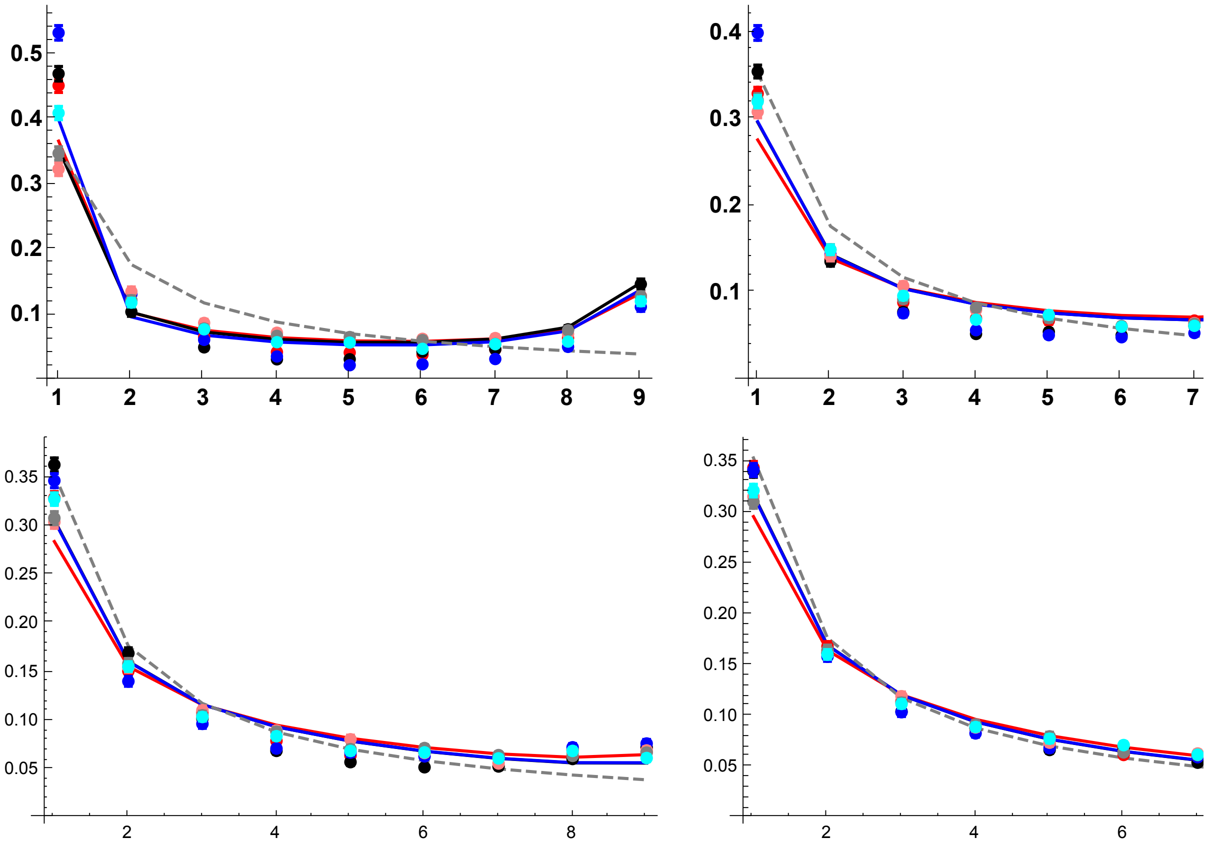


#### $\sigma = 1/2$, $F = 1/3$ case

##### Results with $p_0 = 1/2N$

R = 11

```
datfDNR11h01s05 =
  Import["SLIM_SFS_F033/SFSTab_R_11_h_0.1_self_0.5_f_1e-04_10b_SLIM.dat",
    "Table"][[1]];
datfDNR11h01s05CI = Import[
    "SLIM_SFS_F033/SFSTab_R_11_h_0.1_self_0.5_f_1e-04_10b_SLIM.dat",
    "Table"][[2]];
datfDNR11h05s05 = Import[
    "SLIM_SFS_F033/SFSTab_R_11_h_0.5_self_0.5_f_1e-04_10b_SLIM.dat",
    "Table"][[1]];
datfDNR11h05s05CI = Import[
    "SLIM_SFS_F033/SFSTab_R_11_h_0.5_self_0.5_f_1e-04_10b_SLIM.dat",
    "Table"][[2]];
datfDNR11h09s05 = Import[
    "SLIM_SFS_F033/SFSTab_R_11_h_0.9_self_0.5_f_1e-04_10b_SLIM.dat",
    "Table"][[1]];
datfDNR11h09s05CI = Import[
    "SLIM_SFS_F033/SFSTab_R_11_h_0.9_self_0.5_f_1e-04_10b_SLIM.dat",
    "Table"][[2]];
datfDNR11h01s05T = Partition[Riffle[Partition[Riffle[Range[9], datfDNR11h01s05],
     2], Map[ErrorBar, datfDNR11h01s05CI]], 2];
datfDNR11h05s05T = Partition[Riffle[Partition[Riffle[Range[9], datfDNR11h05s05],
     2], Map[ErrorBar, datfDNR11h05s05CI]], 2];
datfDNR11h09s05T = Partition[Riffle[Partition[Riffle[Range[9], datfDNR11h09s05],
     2], Map[ErrorBar, datfDNR11h09s05CI]], 2];
```

```
p1R11s05 = ListPlot[
  {Table[{l, PLNH2[5000, 0.05, 0.1, 0.5, 11, 10, l, Boostp0[5000, 0.05, 0.1, 0.5],
        4/(1 + 0.5/(2-0.5))]}, {l, 1, 9}], Table[{l, PLNH2[5000, 0.05, 0.5, 0.5,
        11, 10, l, Boostp0[5000, 0.05, 0.5, 0.5], 4/(1 + 0.5/(2-0.5))]}, {l, 1, 9}],
   Table[{l, PLNH2[5000, 0.05, 0.9, 0.5, 11, 10, l, Boostp0[5000, 0.05, 0.9, 0.5],
        4/(1 + 0.5/(2-0.5))]}, {l, 1, 9}], Table[{l, PrJ[l, 10]}, {l, 1, 9}]},
  PlotRange → All, PlotStyle → {Red, Black, Blue, {Gray, Dashed}},
  Joined → True, Ticks → {{1, 2, 3, 4, 5, 6, 7, 8, 9}, Automatic},
  BaseStyle → {FontWeight → "Bold", FontColor → Black, FontSize → 12}]
```

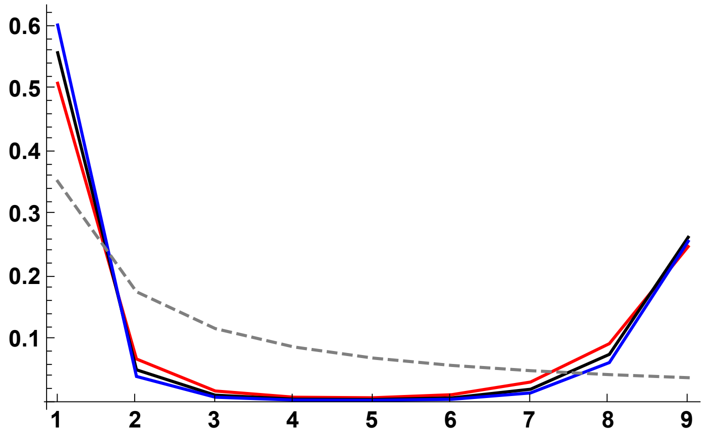


```
p2R11s05 = ErrorListPlot[{datfDNR11h01s05T, datfDNR11h05s05T, datfDNR11h09s05T},
  PlotRange → All, PlotStyle →
   {{Red, PointSize[0.02]}, {Black, PointSize[0.02]}, {Blue, PointSize[0.02]}}]
```

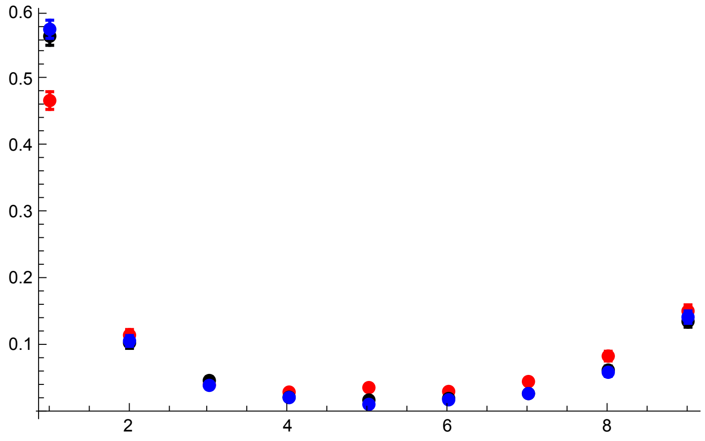

```
p3R11s05DN = Show[p1R11s05, p2R11s05, PlotRange → All, ImageSize → 325]
```

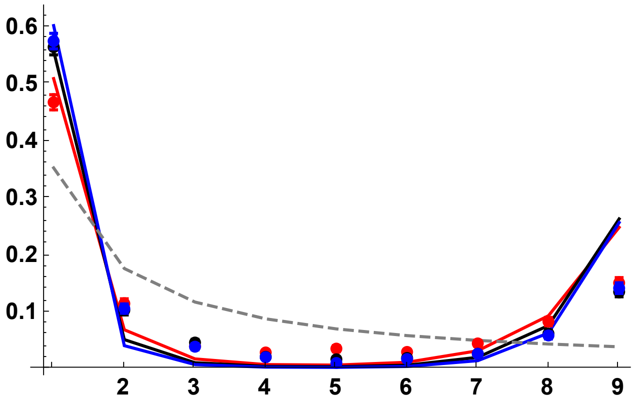


R = 33

```
datfDNR33h01s05 =
  Import["SLIM_SFS_F033/SFSTab_R_33_h_0.1_self_0.5_f_1e-04_10b_SLIM.dat",
    "Table"][[1]];
datfDNR33h01s05CI = Import[
    "SLIM_SFS_F033/SFSTab_R_33_h_0.1_self_0.5_f_1e-04_10b_SLIM.dat",
    "Table"][[2]];
datfDNR33h05s05 = Import[
    "SLIM_SFS_F033/SFSTab_R_33_h_0.5_self_0.5_f_1e-04_10b_SLIM.dat",
    "Table"][[1]];
datfDNR33h05s05CI = Import[
    "SLIM_SFS_F033/SFSTab_R_33_h_0.5_self_0.5_f_1e-04_10b_SLIM.dat",
    "Table"][[2]];
datfDNR33h09s05 = Import[
    "SLIM_SFS_F033/SFSTab_R_33_h_0.9_self_0.5_f_1e-04_10b_SLIM.dat",
    "Table"][[1]];
datfDNR33h09s05CI = Import[
    "SLIM_SFS_F033/SFSTab_R_33_h_0.9_self_0.5_f_1e-04_10b_SLIM.dat",
    "Table"][[2]];
datfDNR33h01s05T = Partition[Riffle[Partition[Riffle[Range[9], datfDNR33h01s05],
     2], Map[ErrorBar, datfDNR33h01s05CI]], 2];
datfDNR33h05s05T = Partition[Riffle[Partition[Riffle[Range[9], datfDNR33h05s05],
     2], Map[ErrorBar, datfDNR33h05s05CI]], 2];
datfDNR33h09s05T = Partition[Riffle[Partition[Riffle[Range[9], datfDNR33h09s05],
     2], Map[ErrorBar, datfDNR33h09s05CI]], 2];
```

```
p1R33s05 = ListPlot[
  {Table[{l, PLNH2[5000, 0.05, 0.1, 0.5, 33, 10, l, Boostp0[5000, 0.05, 0.1, 0.5],
       4/(1 + 0.5/(2-0.5))]}, {l, 1, 9}], Table[{l, PLNH2[5000, 0.05, 0.5, 0.5,
       33, 10, l, Boostp0[5000, 0.05, 0.5, 0.5], 4/(1 + 0.5/(2-0.5))]}, {l, 1, 9}],
   Table[{l, PLNH2[5000, 0.05, 0.9, 0.5, 33, 10, l, Boostp0[5000, 0.05, 0.9, 0.5],
       4/(1 + 0.5/(2-0.5))]}, {l, 1, 9}], Table[{l, PrJ[l, 10]}, {l, 1, 9}]},
  PlotRange → All, PlotStyle → {Red, Black, Blue, {Gray, Dashed}},
  Joined → True]
```

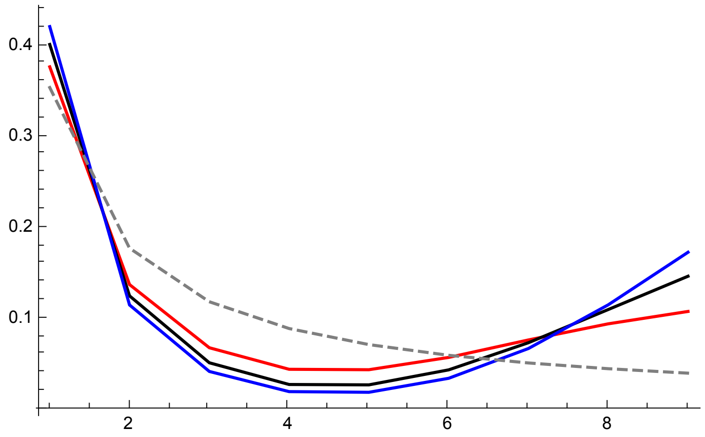


```
p2R33s05 = ErrorListPlot[{datfDNR33h01s05T, datfDNR33h05s05T, datfDNR33h09s05T},
  PlotRange → All, PlotStyle →
   {{Red, PointSize[0.02]}, {Black, PointSize[0.02]}, {Blue, PointSize[0.02]}}]
```

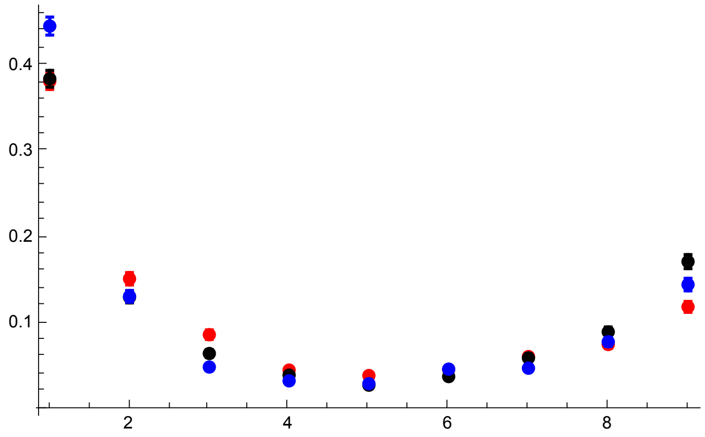

```
p3R33s05 = Show[p1R33s05, p2R33s05, PlotRange → All]
```

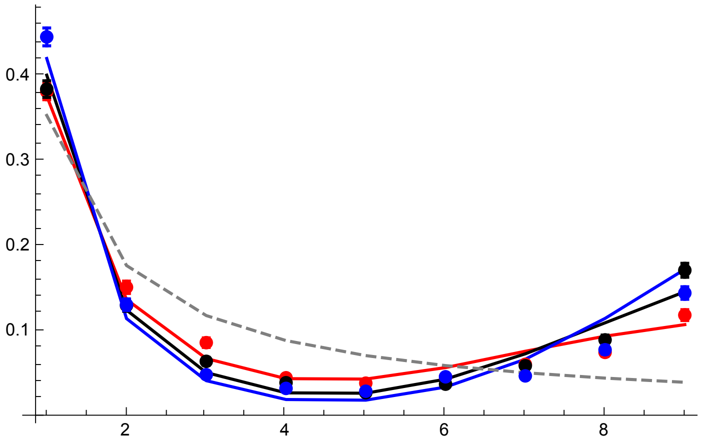


R = 55

```
datfDNR55h01s05 =
  Import["SLIM_SFS_F033/SFSTab_R_55_h_0.1_self_0.5_f_1e-04_10b_SLIM.dat",
    "Table"][[1]];
datfDNR55h01s05CI = Import[
    "SLIM_SFS_F033/SFSTab_R_55_h_0.1_self_0.5_f_1e-04_10b_SLIM.dat",
    "Table"][[2]];
datfDNR55h05s05 = Import[
    "SLIM_SFS_F033/SFSTab_R_55_h_0.5_self_0.5_f_1e-04_10b_SLIM.dat",
    "Table"][[1]];
datfDNR55h05s05CI = Import[
    "SLIM_SFS_F033/SFSTab_R_55_h_0.5_self_0.5_f_1e-04_10b_SLIM.dat",
    "Table"][[2]];
datfDNR55h09s05 = Import[
    "SLIM_SFS_F033/SFSTab_R_55_h_0.9_self_0.5_f_1e-04_10b_SLIM.dat",
    "Table"][[1]];
datfDNR55h09s05CI = Import[
    "SLIM_SFS_F033/SFSTab_R_55_h_0.9_self_0.5_f_1e-04_10b_SLIM.dat",
    "Table"][[2]];
datfDNR55h01s05T = Partition[Riffle[Partition[Riffle[Range[9], datfDNR55h01s05],
     2], Map[ErrorBar, datfDNR55h01s05CI]], 2];
datfDNR55h05s05T = Partition[Riffle[Partition[Riffle[Range[9], datfDNR55h05s05],
     2], Map[ErrorBar, datfDNR55h05s05CI]], 2];
datfDNR55h09s05T = Partition[Riffle[Partition[Riffle[Range[9], datfDNR55h09s05],
     2], Map[ErrorBar, datfDNR55h09s05CI]], 2];
```

```
p1R55s05 = ListPlot[
  {Table[{l, PLNH2[5000, 0.05, 0.1, 0.5, 55, 10, l, Boostp0[5000, 0.05, 0.1, 0.5],
        4/(1 + 0.5/(2-0.5))]}, {l, 1, 9}], Table[{l, PLNH2[5000, 0.05, 0.5, 0.5,
        55, 10, l, Boostp0[5000, 0.05, 0.5, 0.5], 4/(1 + 0.5/(2-0.5))]}, {l, 1, 9}],
   Table[{l, PLNH2[5000, 0.05, 0.9, 0.5, 55, 10, l, Boostp0[5000, 0.05, 0.9, 0.5],
        4/(1 + 0.5/(2-0.5))]}, {l, 1, 9}], Table[{l, PrJ[l, 10]}, {l, 1, 9}]},
  PlotRange → All, PlotStyle → {Red, Black, Blue, {Gray, Dashed}},
  Joined → True]
```

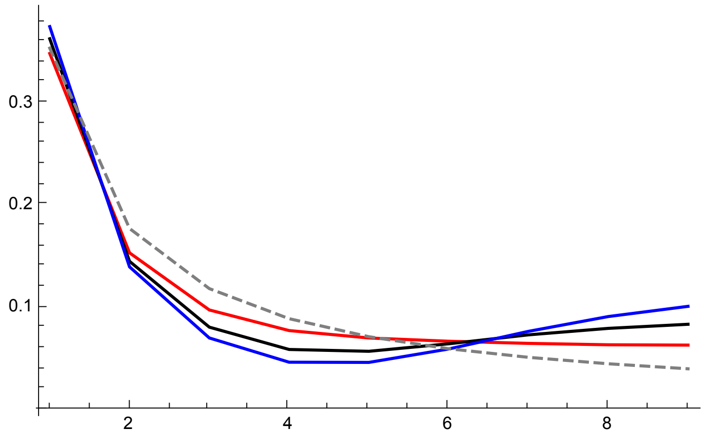


```
p2R55s05 = ErrorListPlot[{datfDNR55h01s05T, datfDNR55h05s05T, datfDNR55h09s05T},
  PlotRange → All, PlotStyle →
   {{Red, PointSize[0.02]}, {Black, PointSize[0.02]}, {Blue, PointSize[0.02]}}]
```

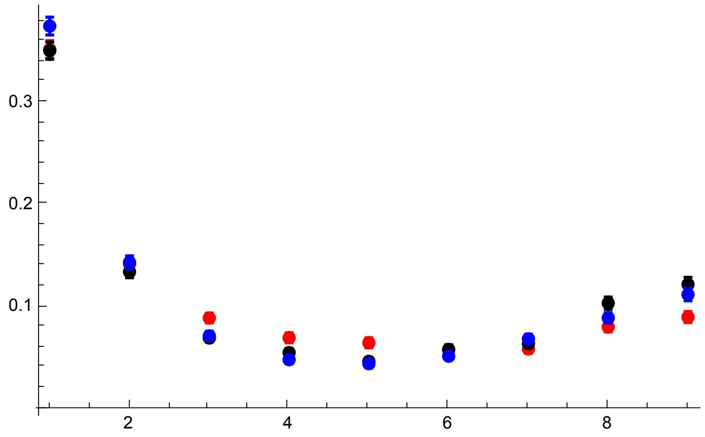

```
p3R55s05 = Show[p1R55s05, p2R55s05, PlotRange → All]
```

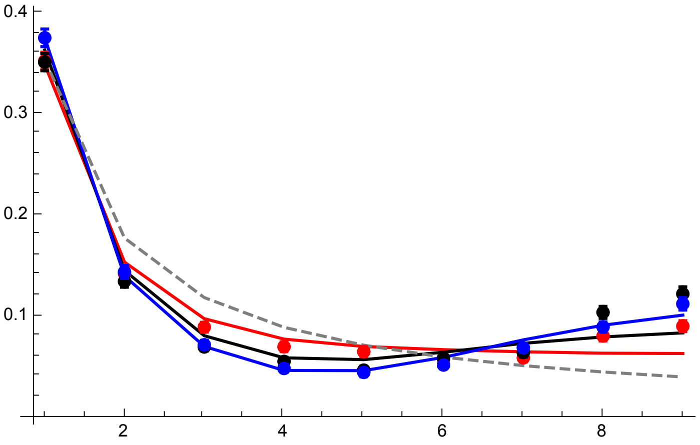


All plots compared

```
GraphicsGrid[{{p3R11s05DN, p3R33s05}, {p3R55s05,}}]
```

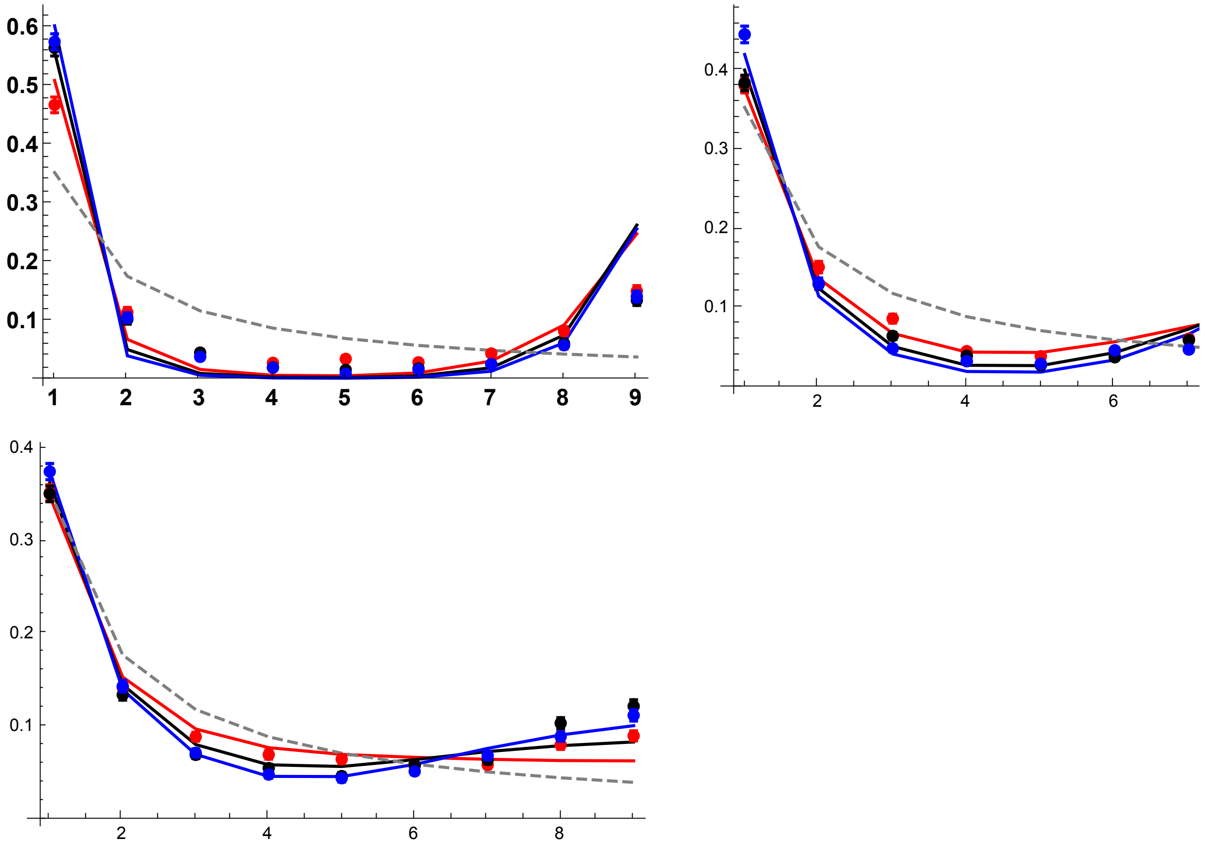


#### Results with $p_0 = 0.02$

R = 11

```
datp002R11h01s05 =
  Import["SLIM_SFS_F033/SFSTab_R_11_h_0.1_self_0.5_f_0.02_10b_SLIM.dat",
    "Table"][[1]];
datp002R11h01s05CI = Import[
    "SLIM_SFS_F033/SFSTab_R_11_h_0.1_self_0.5_f_0.02_10b_SLIM.dat",
    "Table"][[2]];
datp002R11h05s05 = Import[
    "SLIM_SFS_F033/SFSTab_R_11_h_0.5_self_0.5_f_0.02_10b_SLIM.dat",
    "Table"][[1]];
datp002R11h05s05CI = Import[
    "SLIM_SFS_F033/SFSTab_R_11_h_0.5_self_0.5_f_0.02_10b_SLIM.dat",
    "Table"][[2]];
datp002R11h09s05 = Import[
    "SLIM_SFS_F033/SFSTab_R_11_h_0.9_self_0.5_f_0.02_10b_SLIM.dat",
    "Table"][[1]];
datp002R11h09s05CI = Import[
    "SLIM_SFS_F033/SFSTab_R_11_h_0.9_self_0.5_f_0.02_10b_SLIM.dat",
    "Table"][[2]];
datp002R11h01s05T = Partition[Riffle[Partition[Riffle[Range[9],
      datp002R11h01s05], 2], Map[ErrorBar, datp002R11h01s05CI]], 2];
datp002R11h05s05T = Partition[Riffle[Partition[Riffle[Range[9],
      datp002R11h05s05], 2], Map[ErrorBar, datp002R11h05s05CI]], 2];
datp002R11h09s05T = Partition[Riffle[Partition[Riffle[Range[9],
      datp002R11h09s05], 2], Map[ErrorBar, datp002R11h09s05CI]], 2];
```

```
p1R11s05 = ListPlot[
  {Table[{l, PLNH2[5000, 0.05, 0.1, 0.5, 11, 10, l, 0.02, 4/(1 + 0.5/(2-0.5))]}, {l, 1, 9}],
   Table[{l, PLNH2[5000, 0.05, 0.5, 0.5, 11, 10, l, 0.02, 4/(1 + 0.5/(2-0.5))]}, {l, 1, 9}],
   Table[{l, PLNH2[5000, 0.05, 0.9, 0.5, 11, 10, l, 0.02, 4/(1 + 0.5/(2-0.5))]}, {l, 1, 9}],
   Table[{l, PrJ[l, 10]}, {l, 1, 9}]}, PlotRange → All,
  PlotStyle → {Red, Black, Blue, {Gray, Dashed}}, Joined → True,
  Ticks → {{1, 2, 3, 4, 5, 6, 7, 8, 9}, Automatic},
  BaseStyle → {FontWeight → "Bold", FontColor → Black, FontSize → 12}]
```

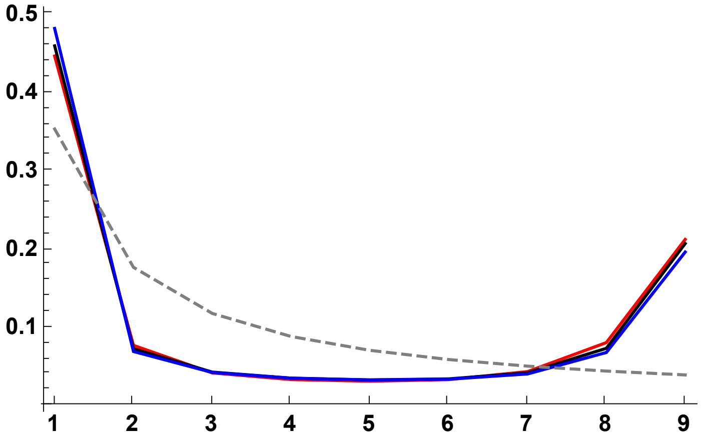


```
p2R11s05 = ErrorListPlot[{datp002R11h01s05T,
   datp002R11h05s05T, datp002R11h09s05T}, PlotRange → All, PlotStyle →
   {{Red, PointSize[0.02]}, {Black, PointSize[0.02]}, {Blue, PointSize[0.02]}}]
```

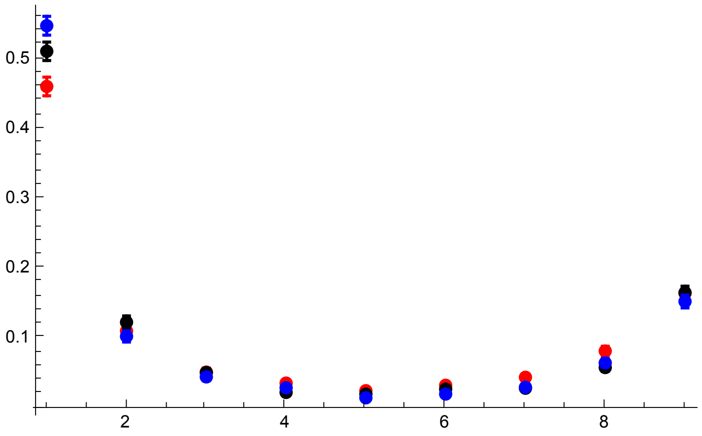

```
p3R11s05p02 = Show[p1R11s05, p2R11s05, PlotRange → All, ImageSize → 325]
```

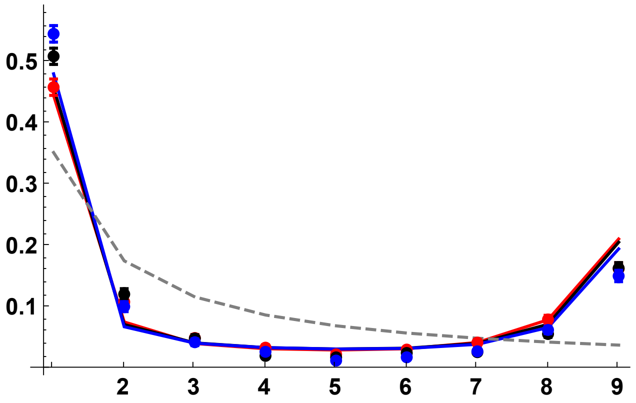


R = 33

```
datp002R33h01s05 =
  Import["SLIM_SFS_F033/SFSTab_R_33_h_0.1_self_0.5_f_0.02_10b_SLIM.dat",
    "Table"][[1]];
datp002R33h01s05CI = Import[
    "SLIM_SFS_F033/SFSTab_R_33_h_0.1_self_0.5_f_0.02_10b_SLIM.dat",
    "Table"][[2]];
datp002R33h05s05 = Import[
    "SLIM_SFS_F033/SFSTab_R_33_h_0.5_self_0.5_f_0.02_10b_SLIM.dat",
    "Table"][[1]];
datp002R33h05s05CI = Import[
    "SLIM_SFS_F033/SFSTab_R_33_h_0.5_self_0.5_f_0.02_10b_SLIM.dat",
    "Table"][[2]];
datp002R33h09s05 = Import[
    "SLIM_SFS_F033/SFSTab_R_33_h_0.9_self_0.5_f_0.02_10b_SLIM.dat",
    "Table"][[1]];
datp002R33h09s05CI = Import[
    "SLIM_SFS_F033/SFSTab_R_33_h_0.9_self_0.5_f_0.02_10b_SLIM.dat",
    "Table"][[2]];
datp002R33h01s05T = Partition[Riffle[Partition[Riffle[Range[9],
      datp002R33h01s05], 2], Map[ErrorBar, datp002R33h01s05CI]], 2];
datp002R33h05s05T = Partition[Riffle[Partition[Riffle[Range[9],
      datp002R33h05s05], 2], Map[ErrorBar, datp002R33h05s05CI]], 2];
datp002R33h09s05T = Partition[Riffle[Partition[Riffle[Range[9],
      datp002R33h09s05], 2], Map[ErrorBar, datp002R33h09s05CI]], 2];
```

```
p1R33s05 = ListPlot[
  {Table[{l, PLNH2[5000, 0.05, 0.1, 0.5, 33, 10, l, 0.02, 4/(1 + 0.5/(2-0.5))]}, {l, 1, 9}],
   Table[{l, PLNH2[5000, 0.05, 0.5, 0.5, 33, 10, l, 0.02, 4/(1 + 0.5/(2-0.5))]}, {l, 1, 9}],
   Table[{l, PLNH2[5000, 0.05, 0.9, 0.5, 33, 10, l, 0.02, 4/(1 + 0.5/(2-0.5))]}, {l, 1, 9}],
   Table[{l, PrJ[l, 10]}, {l, 1, 9}]}, PlotRange → All,
  PlotStyle → {Red, Black, Blue, {Gray, Dashed}}, Joined → True]
```

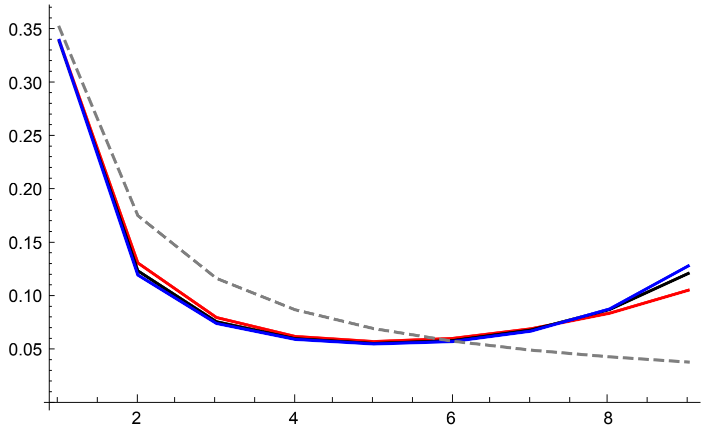


```
p2R33s05 = ErrorListPlot[{datp002R33h01s05T,
   datp002R33h05s05T, datp002R33h09s05T}, PlotRange → All, PlotStyle →
   {{Red, PointSize[0.02]}, {Black, PointSize[0.02]}, {Blue, PointSize[0.02]}}]
```

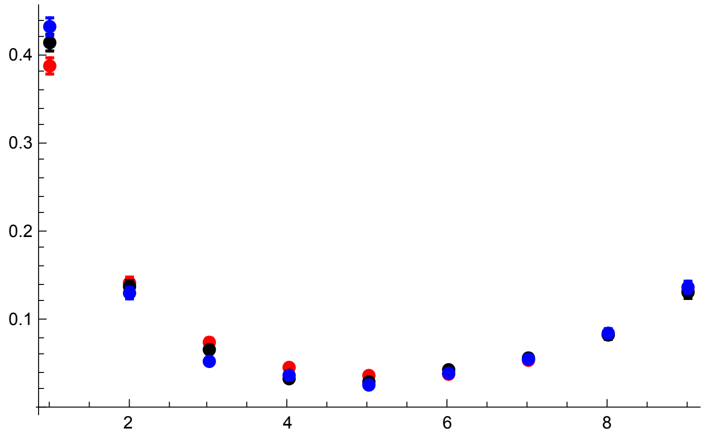

```
p3R33s05 = Show[p1R33s05, p2R33s05, PlotRange → All]
```

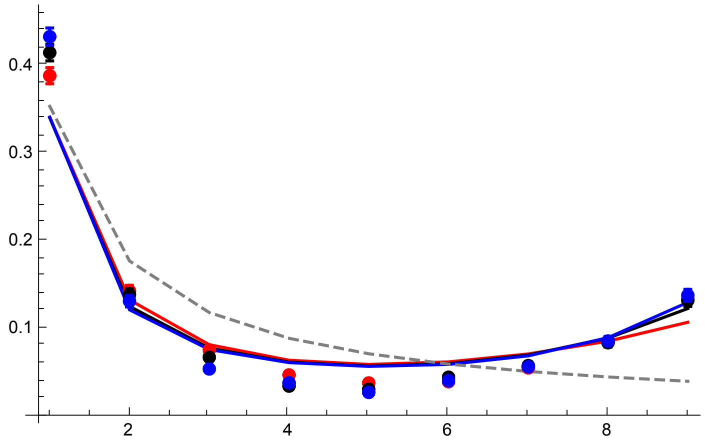


R = 55

```
datp002R55h01s05 =
  Import["SLIM_SFS_F033/SFSTab_R_55_h_0.1_self_0.5_f_0.02_10b_SLIM.dat",
    "Table"][[1]];
datp002R55h01s05CI = Import[
    "SLIM_SFS_F033/SFSTab_R_55_h_0.1_self_0.5_f_0.02_10b_SLIM.dat",
    "Table"][[2]];
datp002R55h05s05 = Import[
    "SLIM_SFS_F033/SFSTab_R_55_h_0.5_self_0.5_f_0.02_10b_SLIM.dat",
    "Table"][[1]];
datp002R55h05s05CI = Import[
    "SLIM_SFS_F033/SFSTab_R_55_h_0.5_self_0.5_f_0.02_10b_SLIM.dat",
    "Table"][[2]];
datp002R55h09s05 = Import[
    "SLIM_SFS_F033/SFSTab_R_55_h_0.9_self_0.5_f_0.02_10b_SLIM.dat",
    "Table"][[1]];
datp002R55h09s05CI = Import[
    "SLIM_SFS_F033/SFSTab_R_55_h_0.9_self_0.5_f_0.02_10b_SLIM.dat",
    "Table"][[2]];
datp002R55h01s05T = Partition[Riffle[Partition[Riffle[Range[9],
      datp002R55h01s05], 2], Map[ErrorBar, datp002R55h01s05CI]], 2];
datp002R55h05s05T = Partition[Riffle[Partition[Riffle[Range[9],
      datp002R55h05s05], 2], Map[ErrorBar, datp002R55h05s05CI]], 2];
datp002R55h09s05T = Partition[Riffle[Partition[Riffle[Range[9],
      datp002R55h09s05], 2], Map[ErrorBar, datp002R55h09s05CI]], 2];
```

```
p1R55s05 = ListPlot[
  {Table[{l, PLNH2[5000, 0.05, 0.1, 0.5, 55, 10, l, 0.02, 4/(1 + 0.5/(2-0.5))]}, {l, 1, 9}],
   Table[{l, PLNH2[5000, 0.05, 0.5, 0.5, 55, 10, l, 0.02, 4/(1 + 0.5/(2-0.5))]}, {l, 1, 9}],
   Table[{l, PLNH2[5000, 0.05, 0.9, 0.5, 55, 10, l, 0.02, 4/(1 + 0.5/(2-0.5))]}, {l, 1, 9}],
   Table[{l, PrJ[l, 10]}, {l, 1, 9}]}, PlotRange → All,
  PlotStyle → {Red, Black, Blue, {Gray, Dashed}}, Joined → True]
```

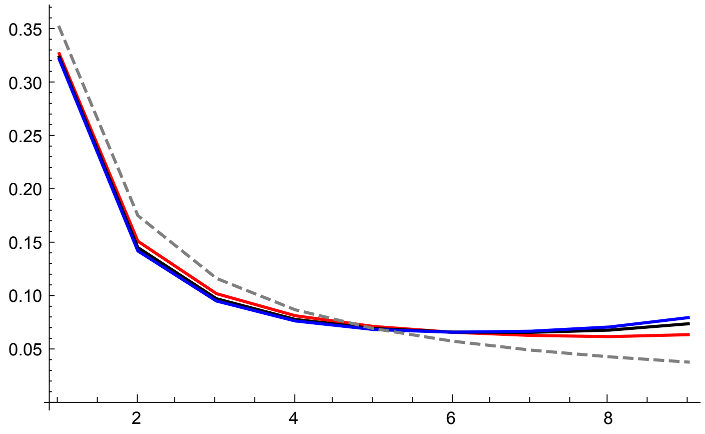


```
p2R55s05 = ErrorListPlot[{datp002R55h01s05T,
   datp002R55h05s05T, datp002R55h09s05T}, PlotRange → All, PlotStyle →
   {{Red, PointSize[0.02]}, {Black, PointSize[0.02]}, {Blue, PointSize[0.02]}}]
```

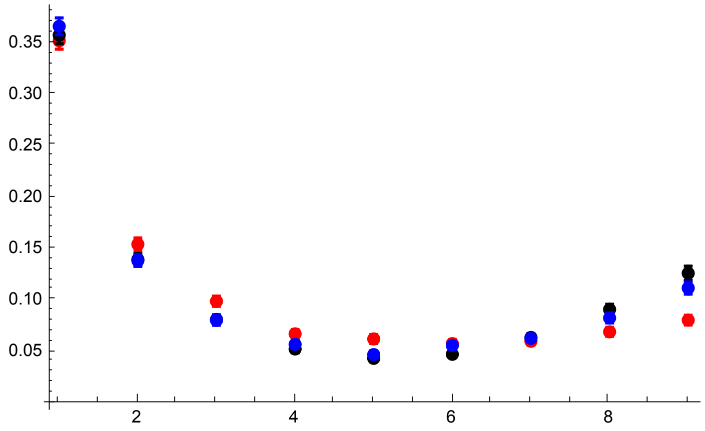

```
p3R55s05 = Show[p1R55s05, p2R55s05, PlotRange → All]
```

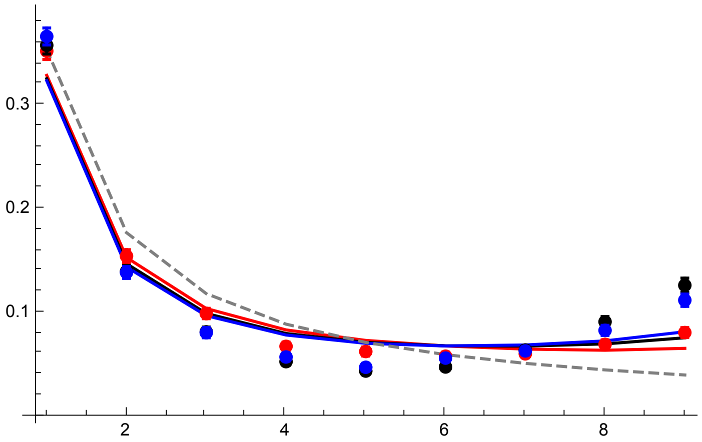


All plots compared

```
GraphicsGrid[{{p3R11s05p02, p3R33s05}, {p3R55s05,}}]
```

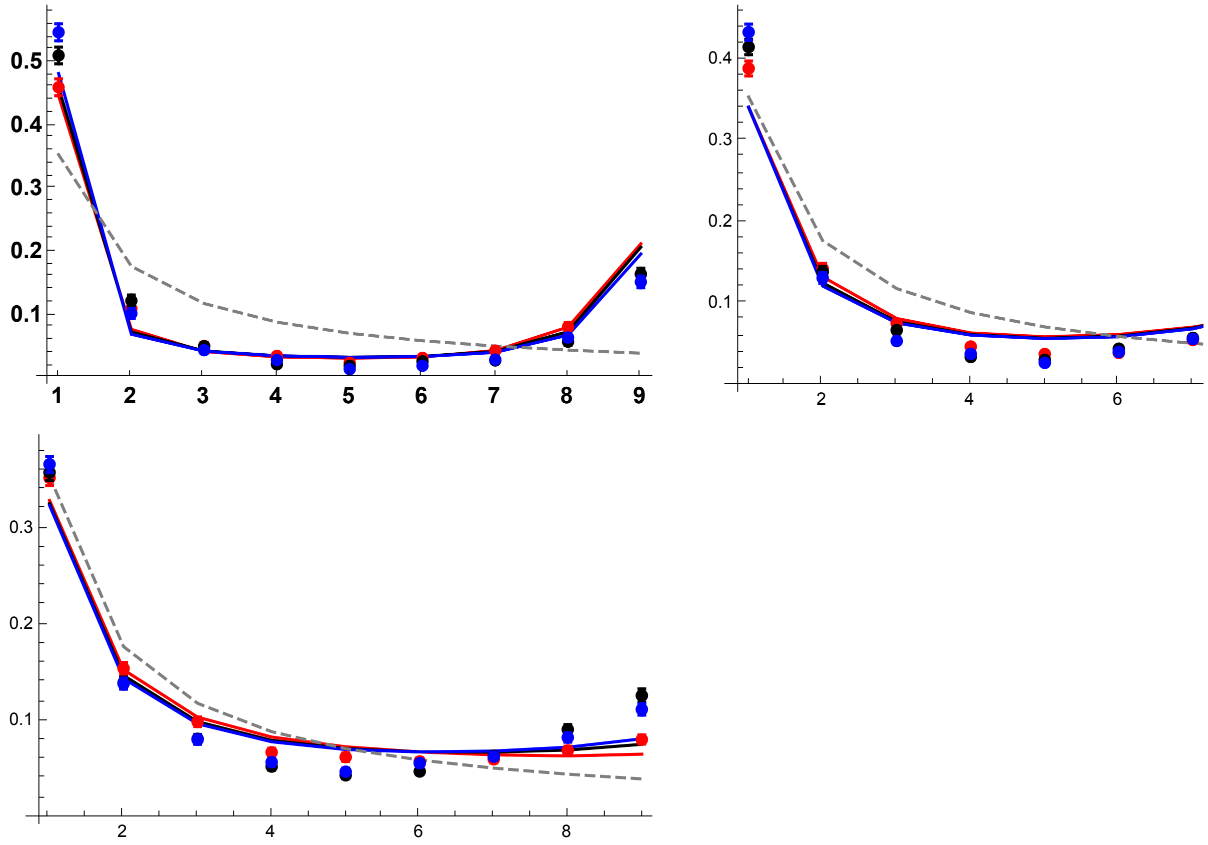


#### Results with $p_0 = 0.05$

R = 11

```
datp005R11h01s05 =
  Import["SLIM_SFS_F033/SFSTab_R_11_h_0.1_self_0.5_f_0.05_10b_SLIM.dat",
    "Table"][[1]];
datp005R11h01s05CI = Import[
    "SLIM_SFS_F033/SFSTab_R_11_h_0.1_self_0.5_f_0.05_10b_SLIM.dat",
    "Table"][[2]];
datp005R11h05s05 = Import[
    "SLIM_SFS_F033/SFSTab_R_11_h_0.5_self_0.5_f_0.05_10b_SLIM.dat",
    "Table"][[1]];
datp005R11h05s05CI = Import[
    "SLIM_SFS_F033/SFSTab_R_11_h_0.5_self_0.5_f_0.05_10b_SLIM.dat",
    "Table"][[2]];
datp005R11h09s05 = Import[
    "SLIM_SFS_F033/SFSTab_R_11_h_0.9_self_0.5_f_0.05_10b_SLIM.dat",
    "Table"][[1]];
datp005R11h09s05CI = Import[
    "SLIM_SFS_F033/SFSTab_R_11_h_0.9_self_0.5_f_0.05_10b_SLIM.dat",
    "Table"][[2]];
datp005R11h01s05T = Partition[Riffle[Partition[Riffle[Range[9],
      datp005R11h01s05], 2], Map[ErrorBar, datp005R11h01s05CI]], 2];
datp005R11h05s05T = Partition[Riffle[Partition[Riffle[Range[9],
      datp005R11h05s05], 2], Map[ErrorBar, datp005R11h05s05CI]], 2];
datp005R11h09s05T = Partition[Riffle[Partition[Riffle[Range[9],
      datp005R11h09s05], 2], Map[ErrorBar, datp005R11h09s05CI]], 2];
```

```
p1R11s05 = ListPlot[
  {Table[{l, PLNH2[5000, 0.05, 0.1, 0.5, 11, 10, l, 0.05, 4/(1 + 0.5/(2-0.5))]}, {l, 1, 9}],
   Table[{l, PLNH2[5000, 0.05, 0.5, 0.5, 11, 10, l, 0.05, 4/(1 + 0.5/(2-0.5))]}, {l, 1, 9}],
   Table[{l, PLNH2[5000, 0.05, 0.9, 0.5, 11, 10, l, 0.05, 4/(1 + 0.5/(2-0.5))]}, {l, 1, 9}],
   Table[{l, PrJ[l, 10]}, {l, 1, 9}]}, PlotRange → All,
  PlotStyle → {Red, Black, Blue, {Gray, Dashed}}, Joined → True,
  Ticks → {{1, 2, 3, 4, 5, 6, 7, 8, 9}, Automatic},
  BaseStyle → {FontWeight → "Bold", FontColor → Black, FontSize → 12}]
```

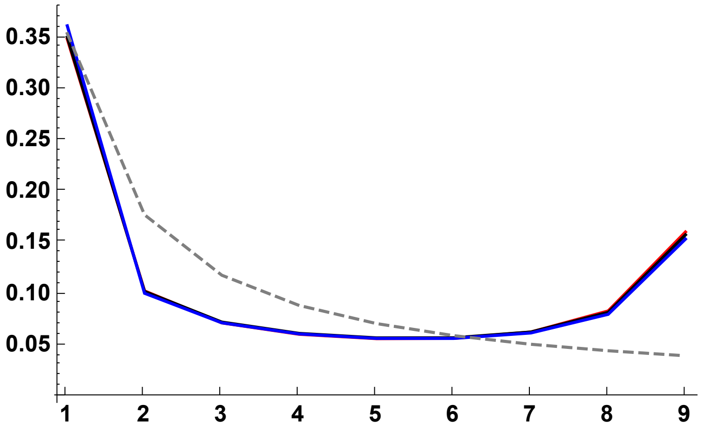


```
p2R11s05 = ErrorListPlot[{datp005R11h01s05T,
   datp005R11h05s05T, datp005R11h09s05T}, PlotRange → All, PlotStyle →
   {{Red, PointSize[0.02]}, {Black, PointSize[0.02]}, {Blue, PointSize[0.02]}}]
```

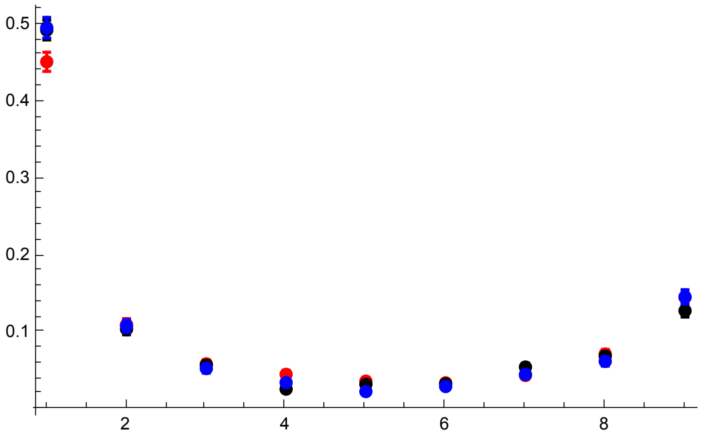

```
p3R11s05p05 = Show[p1R11s05, p2R11s05, PlotRange → All, ImageSize → 325]
```

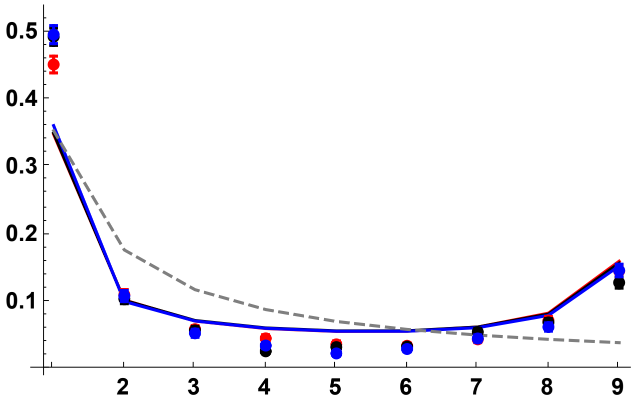


R = 33

```
datp005R33h01s05 =
  Import["SLIM_SFS_F033/SFSTab_R_33_h_0.1_self_0.5_f_0.05_10b_SLIM.dat",
    "Table"][[1]];
datp005R33h01s05CI = Import[
    "SLIM_SFS_F033/SFSTab_R_33_h_0.1_self_0.5_f_0.05_10b_SLIM.dat",
    "Table"][[2]];
datp005R33h05s05 = Import[
    "SLIM_SFS_F033/SFSTab_R_33_h_0.5_self_0.5_f_0.05_10b_SLIM.dat",
    "Table"][[1]];
datp005R33h05s05CI = Import[
    "SLIM_SFS_F033/SFSTab_R_33_h_0.5_self_0.5_f_0.05_10b_SLIM.dat",
    "Table"][[2]];
datp005R33h09s05 = Import[
    "SLIM_SFS_F033/SFSTab_R_33_h_0.9_self_0.5_f_0.05_10b_SLIM.dat",
    "Table"][[1]];
datp005R33h09s05CI = Import[
    "SLIM_SFS_F033/SFSTab_R_33_h_0.9_self_0.5_f_0.05_10b_SLIM.dat",
    "Table"][[2]];
datp005R33h01s05T = Partition[Riffle[Partition[Riffle[Range[9],
      datp005R33h01s05], 2], Map[ErrorBar, datp005R33h01s05CI]], 2];
datp005R33h05s05T = Partition[Riffle[Partition[Riffle[Range[9],
      datp005R33h05s05], 2], Map[ErrorBar, datp005R33h05s05CI]], 2];
datp005R33h09s05T = Partition[Riffle[Partition[Riffle[Range[9],
      datp005R33h09s05], 2], Map[ErrorBar, datp005R33h09s05CI]], 2];
```

```
p1R33s05 = ListPlot[
  {Table[{l, PLNH2[5000, 0.05, 0.1, 0.5, 33, 10, l, 0.05, 4/(1 + 0.5/(2-0.5))]}, {l, 1, 9}],
   Table[{l, PLNH2[5000, 0.05, 0.5, 0.5, 33, 10, l, 0.05, 4/(1 + 0.5/(2-0.5))]}, {l, 1, 9}],
   Table[{l, PLNH2[5000, 0.05, 0.9, 0.5, 33, 10, l, 0.05, 4/(1 + 0.5/(2-0.5))]}, {l, 1, 9}],
   Table[{l, PrJ[l, 10]}, {l, 1, 9}]}, PlotRange → All,
  PlotStyle → {Red, Black, Blue, {Gray, Dashed}}, Joined → True]
```

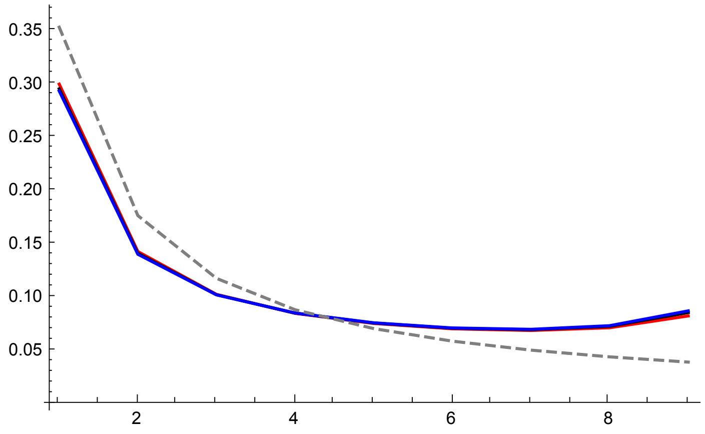


```
p2R33s05 = ErrorListPlot[{datp005R33h01s05T,
   datp005R33h05s05T, datp005R33h09s05T}, PlotRange → All, PlotStyle →
   {{Red, PointSize[0.02]}, {Black, PointSize[0.02]}, {Blue, PointSize[0.02]}}]
```

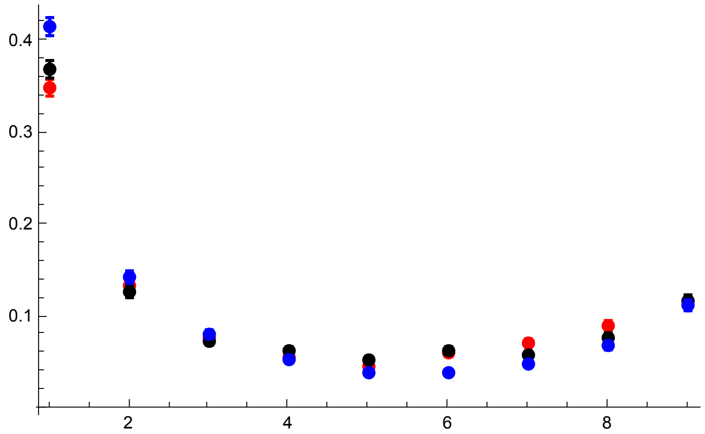

```
p3R33s05 = Show[p1R33s05, p2R33s05, PlotRange → All]
```

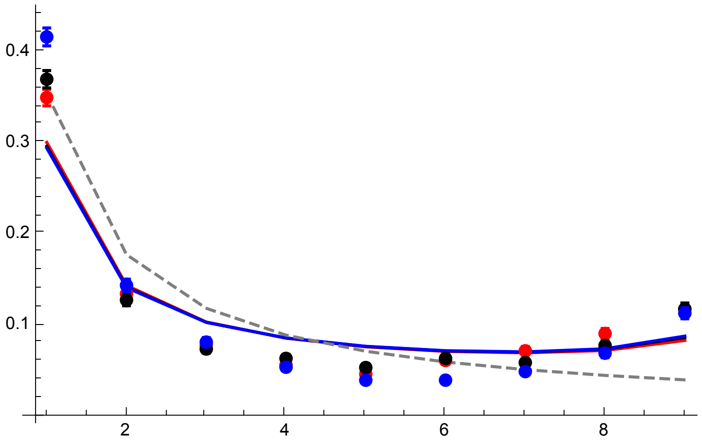


R = 55

```
datp005R55h01s05 =
  Import["SLIM_SFS_F033/SFSTab_R_55_h_0.1_self_0.5_f_0.05_10b_SLIM.dat",
    "Table"][[1]];
datp005R55h01s05CI = Import[
    "SLIM_SFS_F033/SFSTab_R_55_h_0.1_self_0.5_f_0.05_10b_SLIM.dat",
    "Table"][[2]];
datp005R55h05s05 = Import[
    "SLIM_SFS_F033/SFSTab_R_55_h_0.5_self_0.5_f_0.05_10b_SLIM.dat",
    "Table"][[1]];
datp005R55h05s05CI = Import[
    "SLIM_SFS_F033/SFSTab_R_55_h_0.5_self_0.5_f_0.05_10b_SLIM.dat",
    "Table"][[2]];
datp005R55h09s05 = Import[
    "SLIM_SFS_F033/SFSTab_R_55_h_0.9_self_0.5_f_0.05_10b_SLIM.dat",
    "Table"][[1]];
datp005R55h09s05CI = Import[
    "SLIM_SFS_F033/SFSTab_R_55_h_0.9_self_0.5_f_0.05_10b_SLIM.dat",
    "Table"][[2]];
datp005R55h01s05T = Partition[Riffle[Partition[Riffle[Range[9],
      datp005R55h01s05], 2], Map[ErrorBar, datp005R55h01s05CI]], 2];
datp005R55h05s05T = Partition[Riffle[Partition[Riffle[Range[9],
      datp005R55h05s05], 2], Map[ErrorBar, datp005R55h05s05CI]], 2];
datp005R55h09s05T = Partition[Riffle[Partition[Riffle[Range[9],
      datp005R55h09s05], 2], Map[ErrorBar, datp005R55h09s05CI]], 2];
```

```
p1R55s05 = ListPlot[
  {Table[{l, PLNH2[5000, 0.05, 0.1, 0.5, 55, 10, l, 0.05, 4/(1 + 0.5/(2-0.5))]}, {l, 1, 9}],
   Table[{l, PLNH2[5000, 0.05, 0.5, 0.5, 55, 10, l, 0.05, 4/(1 + 0.5/(2-0.5))]}, {l, 1, 9}],
   Table[{l, PLNH2[5000, 0.05, 0.9, 0.5, 55, 10, l, 0.05, 4/(1 + 0.5/(2-0.5))]}, {l, 1, 9}],
   Table[{l, PrJ[l, 10]}, {l, 1, 9}]}, PlotRange → All,
  PlotStyle → {Red, Black, Blue, {Gray, Dashed}}, Joined → True]
```

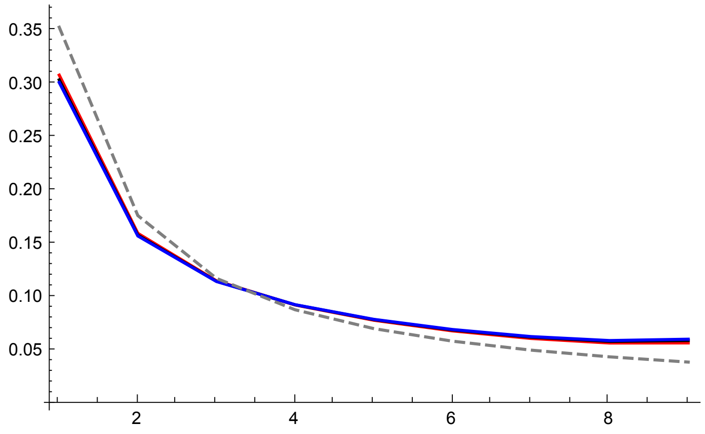


```
p2R55s05 = ErrorListPlot[{datp005R55h01s05T,
   datp005R55h05s05T, datp005R55h09s05T}, PlotRange → All, PlotStyle →
   {{Red, PointSize[0.02]}, {Black, PointSize[0.02]}, {Blue, PointSize[0.02]}}]
```

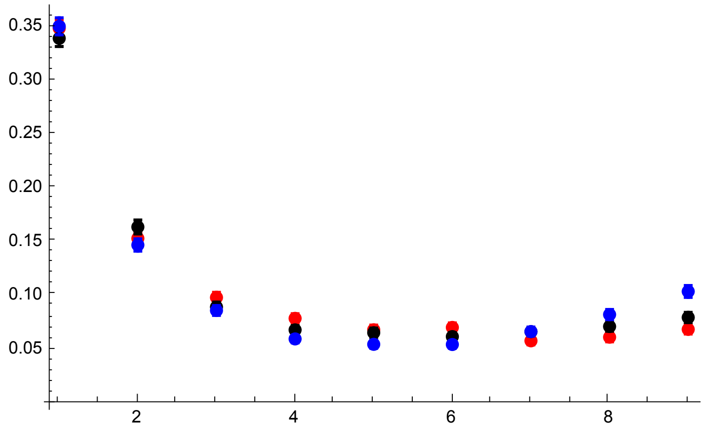

```
p3R55s05 = Show[p1R55s05, p2R55s05, PlotRange → All]
```

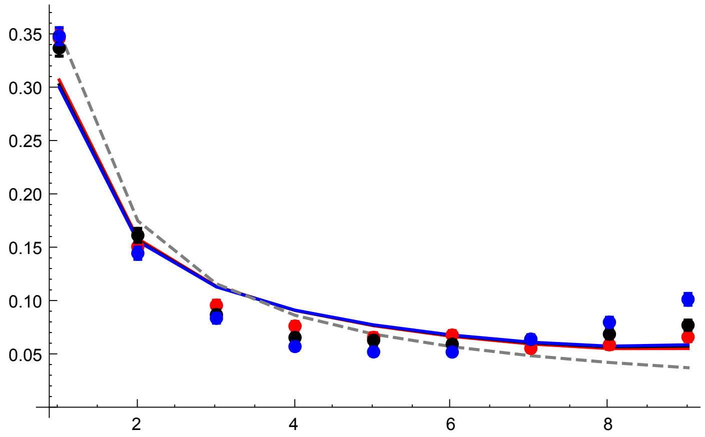


All plots compared

```
GraphicsGrid[{{p3R11s05p05, p3R33s05}, {p3R55s05,}}]
```

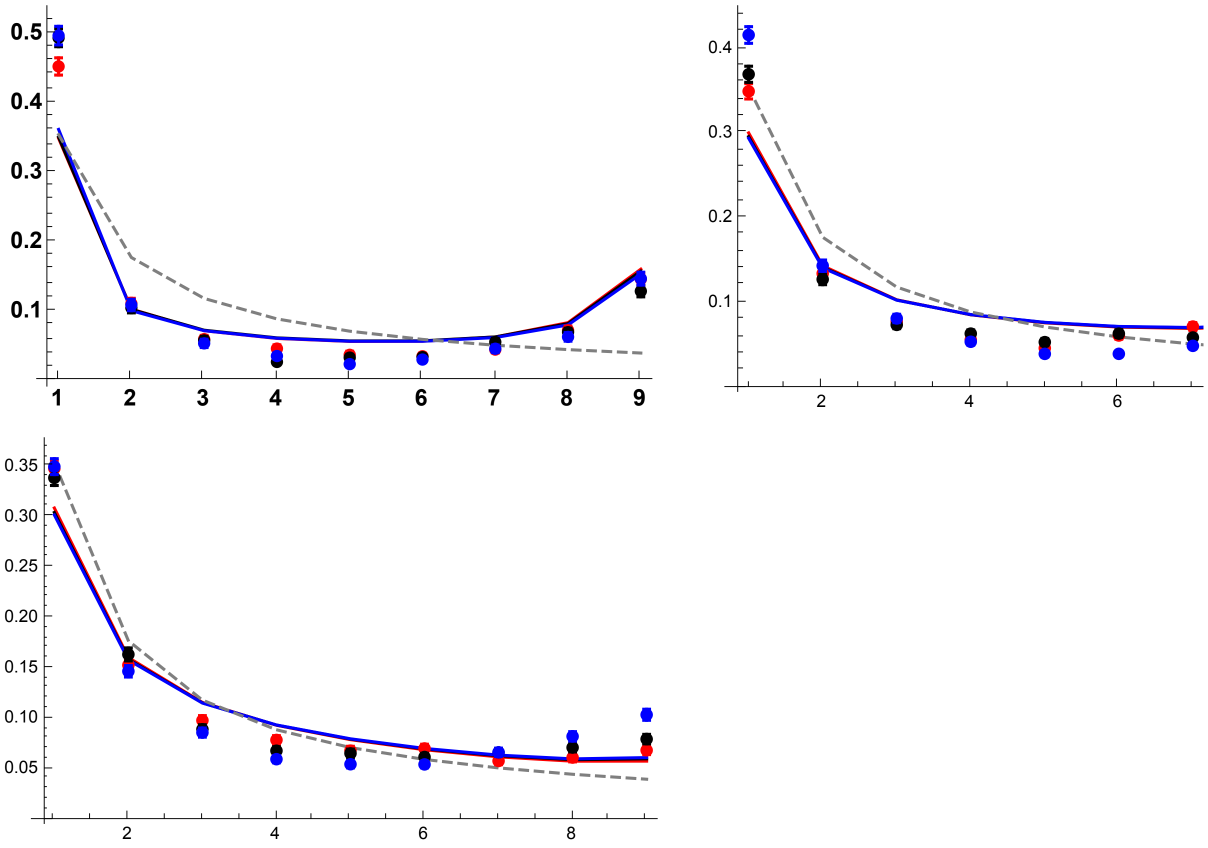


### $\sigma = 0.95$, $F \simeq 0.904$ case

#### Results with $p_0 = 1/2N$

R = 100

```
datfDNR100h01s095 =
  Import["SLIM_SFS_F09/SFSTab_R_100_h_0.1_self_0.95_f_1e-04_10b_SLIM.dat",
    "Table"][[1]];
datfDNR100h01s095CI = Import[
    "SLIM_SFS_F09/SFSTab_R_100_h_0.1_self_0.95_f_1e-04_10b_SLIM.dat",
    "Table"][[2]];
datfDNR100h05s095 = Import[
    "SLIM_SFS_F09/SFSTab_R_100_h_0.5_self_0.95_f_1e-04_10b_SLIM.dat",
    "Table"][[1]];
datfDNR100h05s095CI = Import[
    "SLIM_SFS_F09/SFSTab_R_100_h_0.5_self_0.95_f_1e-04_10b_SLIM.dat",
    "Table"][[2]];
datfDNR100h09s095 = Import[
    "SLIM_SFS_F09/SFSTab_R_100_h_0.9_self_0.95_f_1e-04_10b_SLIM.dat",
    "Table"][[1]];
datfDNR100h09s095CI = Import[
    "SLIM_SFS_F09/SFSTab_R_100_h_0.9_self_0.95_f_1e-04_10b_SLIM.dat",
    "Table"][[2]];
datfDNR100h01s095T = Partition[Riffle[Partition[Riffle[Range[9],
      datfDNR100h01s095], 2], Map[ErrorBar, datfDNR100h01s095CI]], 2];
datfDNR100h05s095T = Partition[Riffle[Partition[Riffle[Range[9],
      datfDNR100h05s095], 2], Map[ErrorBar, datfDNR100h05s095CI]], 2];
datfDNR100h09s095T = Partition[Riffle[Partition[Riffle[Range[9],
      datfDNR100h09s095], 2], Map[ErrorBar, datfDNR100h09s095CI]], 2];
```

```
p1R100s095 =
 ListPlot[{Table[{l, PLNH2[5000, 0.05, 0.1, 0.95, 100, 10, l, Boostp0[5000, 0.05,
        0.1, 0.95], 4/(1 + 0.95/(2-0.95))]}, {l, 1, 9}], Table[{l, PLNH2[5000, 0.05, 0.5, 0.95,
       100, 10, l, Boostp0[5000, 0.05, 0.5, 0.95], 4/(1 + 0.95/(2-0.95))]}, {l, 1, 9}],
    Table[{l, PLNH2[5000, 0.05, 0.9, 0.95, 100, 10, l,
       Boostp0[5000, 0.05, 0.9, 0.95], 4/(1 + 0.95/(2-0.95))]}, {l, 1, 9}],
    Table[{l, PrJ[l, 10]}, {l, 1, 9}]}, PlotRange → All,
  PlotStyle → {Red, Black, Blue, {Gray, Dashed}}, Joined → True]
```

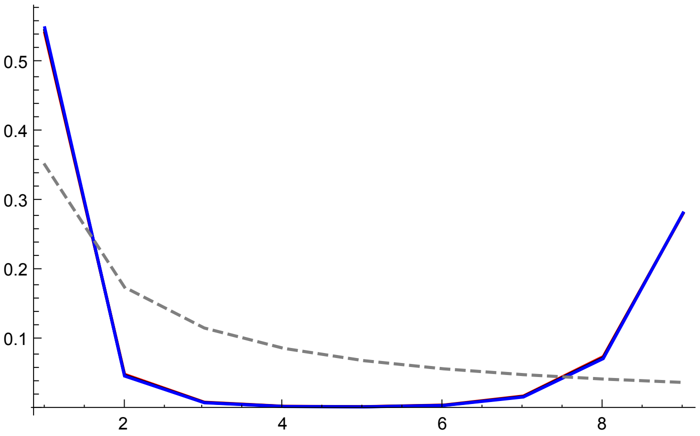


```
p2R100s095 = ErrorListPlot[{datfDNR100h01s095T,
   datfDNR100h05s095T, datfDNR100h09s095T}, PlotRange → All, PlotStyle →
   {{Red, PointSize[0.02]}, {Black, PointSize[0.02]}, {Blue, PointSize[0.02]}}]
```

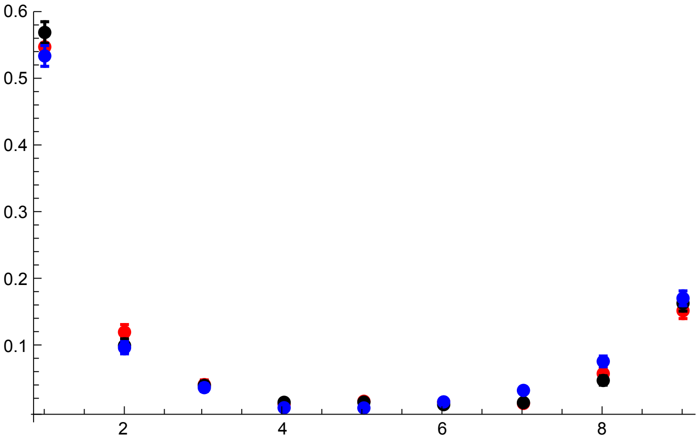

```
p3R100s095 = Show[p1R100s095, p2R100s095, PlotRange → All]
```

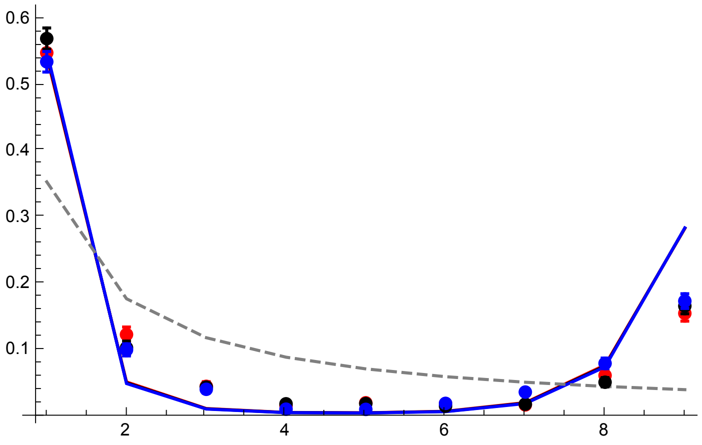


R = 300

```
datfDNR300h01s095 =
  Import["SLIM_SFS_F09/SFSTab_R_300_h_0.1_self_0.95_f_1e-04_10b_SLIM.dat",
    "Table"][[1]];
datfDNR300h01s095CI = Import[
    "SLIM_SFS_F09/SFSTab_R_300_h_0.1_self_0.95_f_1e-04_10b_SLIM.dat",
    "Table"][[2]];
datfDNR300h05s095 = Import[
    "SLIM_SFS_F09/SFSTab_R_300_h_0.5_self_0.95_f_1e-04_10b_SLIM.dat",
    "Table"][[1]];
datfDNR300h05s095CI = Import[
    "SLIM_SFS_F09/SFSTab_R_300_h_0.5_self_0.95_f_1e-04_10b_SLIM.dat",
    "Table"][[2]];
datfDNR300h09s095 = Import[
    "SLIM_SFS_F09/SFSTab_R_300_h_0.9_self_0.95_f_1e-04_10b_SLIM.dat",
    "Table"][[1]];
datfDNR300h09s095CI = Import[
    "SLIM_SFS_F09/SFSTab_R_300_h_0.9_self_0.95_f_1e-04_10b_SLIM.dat",
    "Table"][[2]];
datfDNR300h01s095T = Partition[Riffle[Partition[Riffle[Range[9],
      datfDNR300h01s095], 2], Map[ErrorBar, datfDNR300h01s095CI]], 2];
datfDNR300h05s095T = Partition[Riffle[Partition[Riffle[Range[9],
      datfDNR300h05s095], 2], Map[ErrorBar, datfDNR300h05s095CI]], 2];
datfDNR300h09s095T = Partition[Riffle[Partition[Riffle[Range[9],
      datfDNR300h09s095], 2], Map[ErrorBar, datfDNR300h09s095CI]], 2];
```

```
p1R300s095 =
 ListPlot[{Table[{l, PLNH2[5000, 0.05, 0.1, 0.95, 300, 10, l, Boostp0[5000, 0.05,
        0.1, 0.95], 4/(1 + 0.95/(2-0.95))]}, {l, 1, 9}], Table[{l, PLNH2[5000, 0.05, 0.5, 0.95,
       300, 10, l, Boostp0[5000, 0.05, 0.5, 0.95], 4/(1 + 0.95/(2-0.95))]}, {l, 1, 9}],
   Table[{l, PLNH2[5000, 0.05, 0.9, 0.95, 300, 10, l,
       Boostp0[5000, 0.05, 0.9, 0.95], 4/(1 + 0.95/(2-0.95))]}, {l, 1, 9}],
   Table[{l, PrJ[l, 10]}, {l, 1, 9}]}, PlotRange → All,
  PlotStyle → {Red, Black, Blue, {Gray, Dashed}}, Joined → True]
```

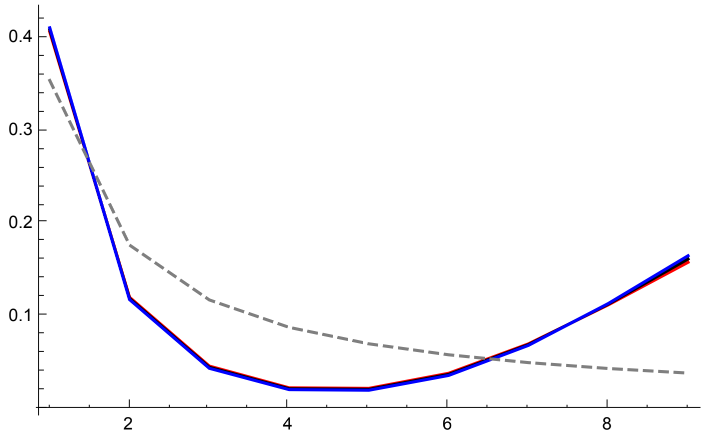


```
p2R300s095 = ErrorListPlot[{datfDNR300h01s095T,
   datfDNR300h05s095T, datfDNR300h09s095T}, PlotRange → All, PlotStyle →
   {{Red, PointSize[0.02]}, {Black, PointSize[0.02]}, {Blue, PointSize[0.02]}}]
```

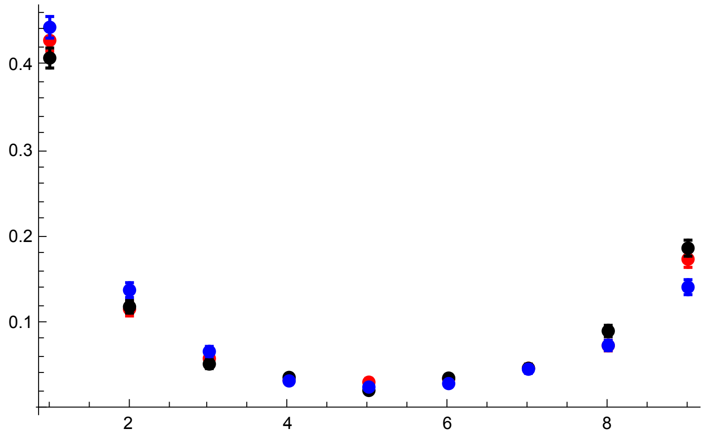

```
p3R300s095 = Show[p1R300s095, p2R300s095, PlotRange → All]
```

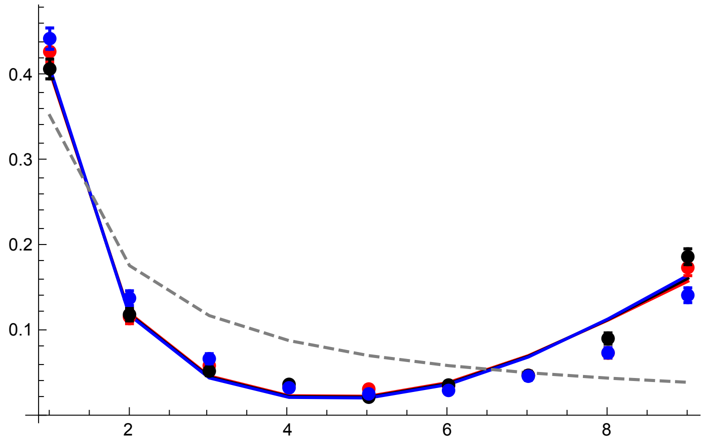


R = 500

```
datfDNR500h01s095 =
  Import["SLIM_SFS_F09/SFSTab_R_500_h_0.1_self_0.95_f_1e-04_10b_SLIM.dat",
    "Table"][[1]];
datfDNR500h01s095CI = Import[
    "SLIM_SFS_F09/SFSTab_R_500_h_0.1_self_0.95_f_1e-04_10b_SLIM.dat",
    "Table"][[2]];
datfDNR500h05s095 = Import[
    "SLIM_SFS_F09/SFSTab_R_500_h_0.5_self_0.95_f_1e-04_10b_SLIM.dat",
    "Table"][[1]];
datfDNR500h05s095CI = Import[
    "SLIM_SFS_F09/SFSTab_R_500_h_0.5_self_0.95_f_1e-04_10b_SLIM.dat",
    "Table"][[2]];
datfDNR500h09s095 = Import[
    "SLIM_SFS_F09/SFSTab_R_500_h_0.9_self_0.95_f_1e-04_10b_SLIM.dat",
    "Table"][[1]];
datfDNR500h09s095CI = Import[
    "SLIM_SFS_F09/SFSTab_R_500_h_0.9_self_0.95_f_1e-04_10b_SLIM.dat",
    "Table"][[2]];
datfDNR500h01s095T = Partition[Riffle[Partition[Riffle[Range[9],
      datfDNR500h01s095], 2], Map[ErrorBar, datfDNR500h01s095CI]], 2];
datfDNR500h05s095T = Partition[Riffle[Partition[Riffle[Range[9],
      datfDNR500h05s095], 2], Map[ErrorBar, datfDNR500h05s095CI]], 2];
datfDNR500h09s095T = Partition[Riffle[Partition[Riffle[Range[9],
      datfDNR500h09s095], 2], Map[ErrorBar, datfDNR500h09s095CI]], 2];
```

```
p1R500s095 =
 ListPlot[{Table[{l, PLNH2[5000, 0.05, 0.1, 0.95, 500, 10, l, Boostp0[5000, 0.05,
        0.1, 0.95], 4/(1 + 0.95/(2-0.95))]}, {l, 1, 9}], Table[{l, PLNH2[5000, 0.05, 0.5, 0.95,
       500, 10, l, Boostp0[5000, 0.05, 0.5, 0.95], 4/(1 + 0.95/(2-0.95))]}, {l, 1, 9}],
   Table[{l, PLNH2[5000, 0.05, 0.9, 0.95, 500, 10, l,
       Boostp0[5000, 0.05, 0.9, 0.95], 4/(1 + 0.95/(2-0.95))]}, {l, 1, 9}],
   Table[{l, PrJ[l, 10]}, {l, 1, 9}]}, PlotRange → All,
  PlotStyle → {Red, Black, Blue, {Gray, Dashed}}, Joined → True]
```

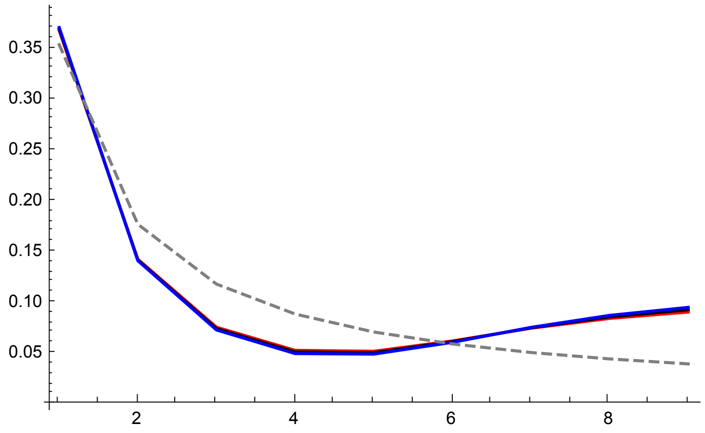


```
p2R500s095 = ErrorListPlot[{datfDNR500h01s095T,
   datfDNR500h05s095T, datfDNR500h09s095T}, PlotRange → All, PlotStyle →
   {{Red, PointSize[0.02]}, {Black, PointSize[0.02]}, {Blue, PointSize[0.02]}}]
```

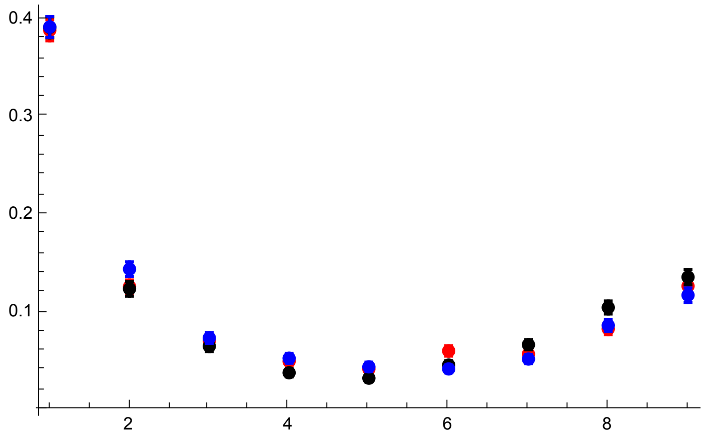

```
p3R500s095 = Show[p1R500s095, p2R500s095, PlotRange → All]
```

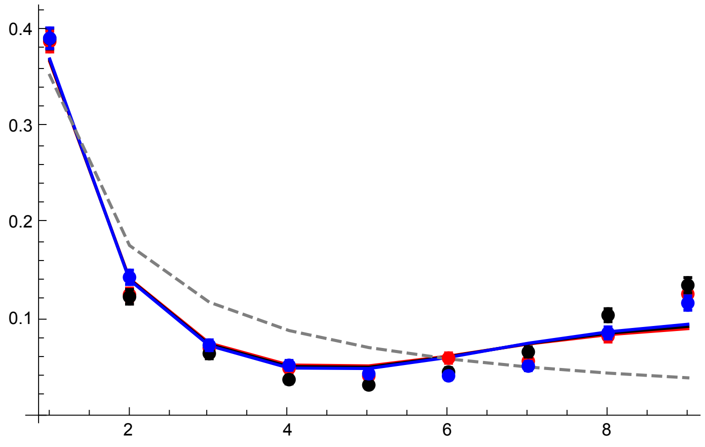


All plots compared

```
GraphicsGrid[{{p3R100s095, p3R300s095}, {p3R500s095,}}]
```

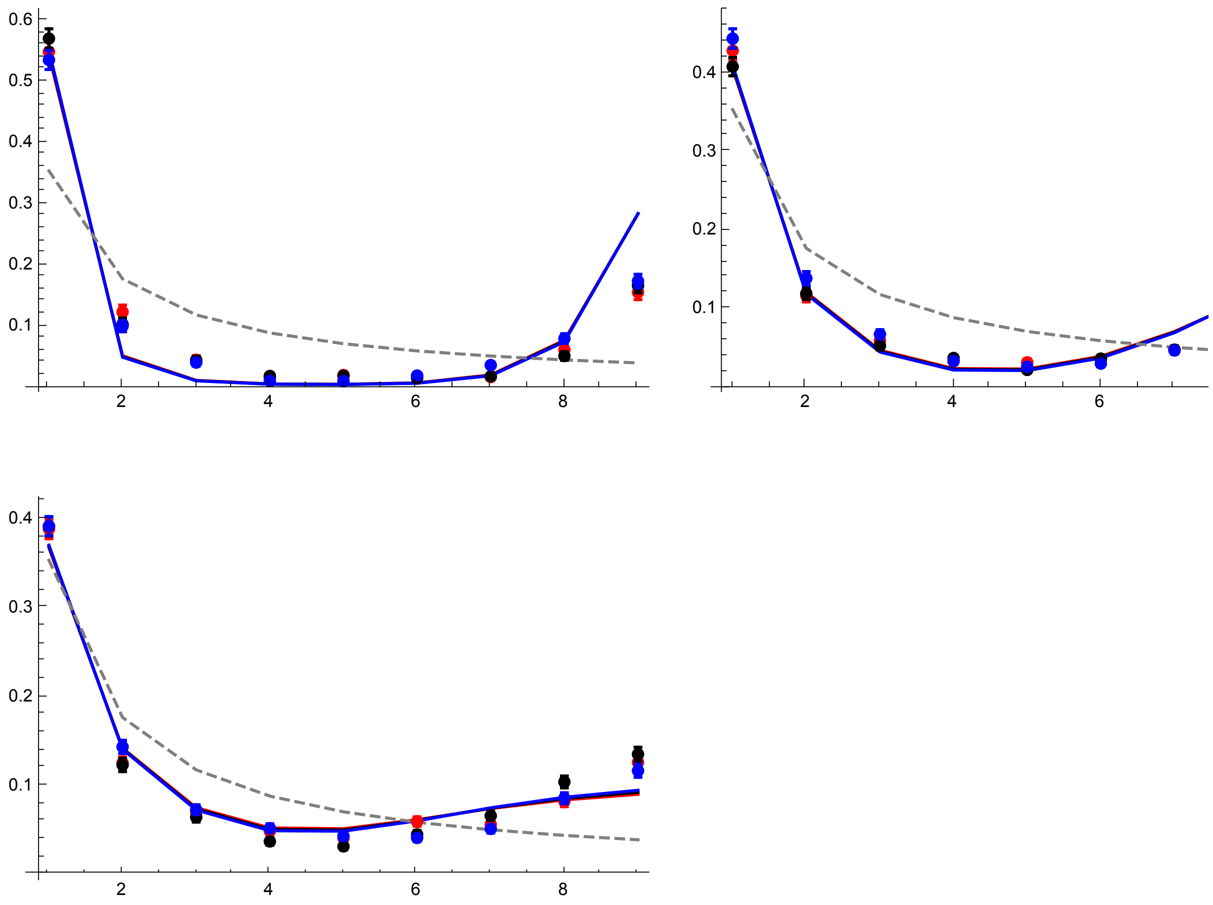


#### Results with $p_0 = 0.02$

R = 100

```
datp002R100h01s095 =
  Import["SLIM_SFS_F09/SFSTab_R_100_h_0.1_self_0.95_f_0.02_10b_SLIM.dat",
    "Table"][[1]];
datp002R100h01s095CI = Import[
    "SLIM_SFS_F09/SFSTab_R_100_h_0.1_self_0.95_f_0.02_10b_SLIM.dat",
    "Table"][[2]];
datp002R100h05s095 = Import[
    "SLIM_SFS_F09/SFSTab_R_100_h_0.5_self_0.95_f_0.02_10b_SLIM.dat",
    "Table"][[1]];
datp002R100h05s095CI = Import[
    "SLIM_SFS_F09/SFSTab_R_100_h_0.5_self_0.95_f_0.02_10b_SLIM.dat",
    "Table"][[2]];
datp002R100h09s095 = Import[
    "SLIM_SFS_F09/SFSTab_R_100_h_0.9_self_0.95_f_0.02_10b_SLIM.dat",
    "Table"][[1]];
datp002R100h09s095CI = Import[
    "SLIM_SFS_F09/SFSTab_R_100_h_0.9_self_0.95_f_0.02_10b_SLIM.dat",
    "Table"][[2]];
datp002R100h01s095T = Partition[Riffle[Partition[Riffle[Range[9],
      datp002R100h01s095], 2], Map[ErrorBar, datp002R100h01s095CI]], 2];
datp002R100h05s095T = Partition[Riffle[Partition[Riffle[Range[9],
      datp002R100h05s095], 2], Map[ErrorBar, datp002R100h05s095CI]], 2];
datp002R100h09s095T = Partition[Riffle[Partition[Riffle[Range[9],
      datp002R100h09s095], 2], Map[ErrorBar, datp002R100h09s095CI]], 2];
```

```
p1R100s095 = ListPlot[
  {Table[{l, PLNH2[5000, 0.05, 0.1, 0.95, 100, 10, l, 0.02, 4/(1 + 0.95/(2-0.95))]}, {l, 1, 9}],
   Table[{l, PLNH2[5000, 0.05, 0.5, 0.95, 100, 10, l, 0.02, 4/(1 + 0.95/(2-0.95))]}, {l, 1, 9}],
   Table[{l, PLNH2[5000, 0.05, 0.9, 0.95, 100, 10, l, 0.02, 4/(1 + 0.95/(2-0.95))]}, {l, 1, 9}],
   Table[{l, PrJ[l, 10]}, {l, 1, 9}]}, PlotRange → All,
  PlotStyle → {Red, Black, Blue, {Gray, Dashed}}, Joined → True]
```

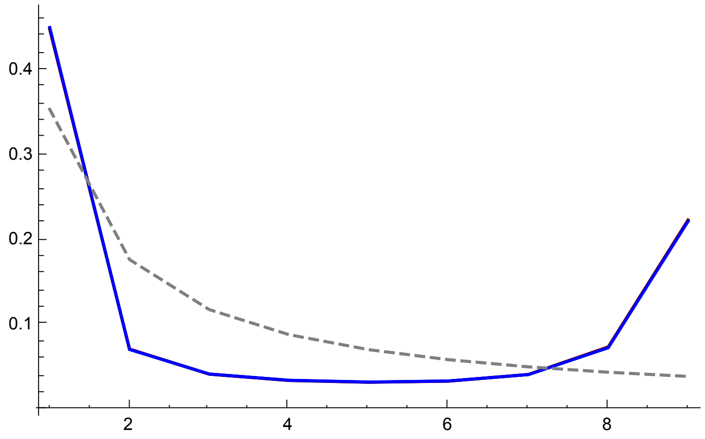


```
p2R100s095 = ErrorListPlot[{datp002R100h01s095T,
   datp002R100h05s095T, datp002R100h09s095T}, PlotRange → All, PlotStyle →
   {{Red, PointSize[0.02]}, {Black, PointSize[0.02]}, {Blue, PointSize[0.02]}}]
```

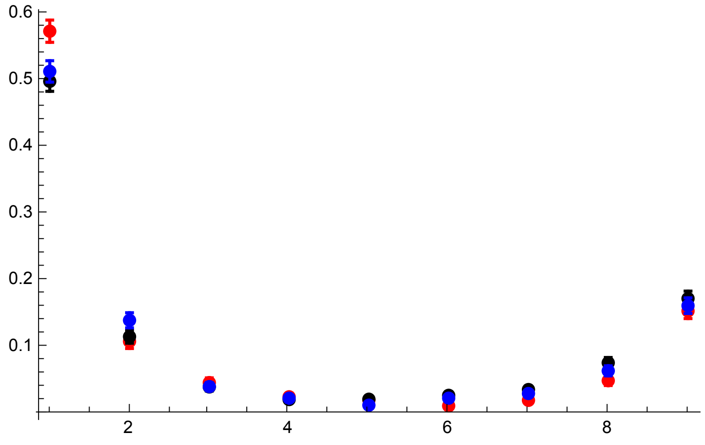

```
p3R100s095 = Show[p1R100s095, p2R100s095, PlotRange → All]
```

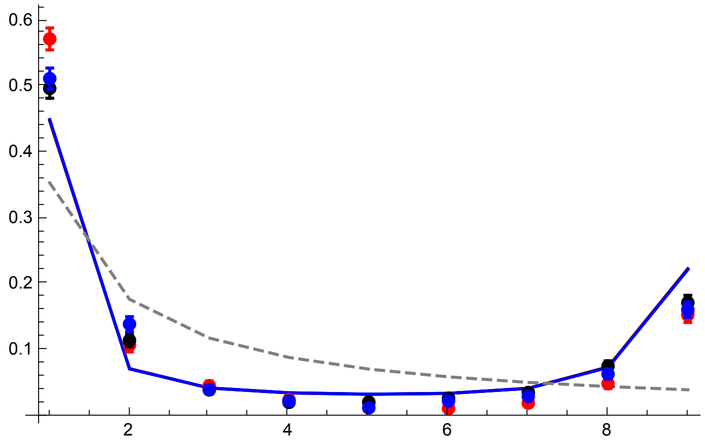


R = 300

```
datp002R300h01s095 =
  Import["SLIM_SFS_F09/SFSTab_R_300_h_0.1_self_0.95_f_0.02_10b_SLIM.dat",
    "Table"][[1]];
datp002R300h01s095CI = Import[
    "SLIM_SFS_F09/SFSTab_R_300_h_0.1_self_0.95_f_0.02_10b_SLIM.dat",
    "Table"][[2]];
datp002R300h05s095 = Import[
    "SLIM_SFS_F09/SFSTab_R_300_h_0.5_self_0.95_f_0.02_10b_SLIM.dat",
    "Table"][[1]];
datp002R300h05s095CI = Import[
    "SLIM_SFS_F09/SFSTab_R_300_h_0.5_self_0.95_f_0.02_10b_SLIM.dat",
    "Table"][[2]];
datp002R300h09s095 = Import[
    "SLIM_SFS_F09/SFSTab_R_300_h_0.9_self_0.95_f_0.02_10b_SLIM.dat",
    "Table"][[1]];
datp002R300h09s095CI = Import[
    "SLIM_SFS_F09/SFSTab_R_300_h_0.9_self_0.95_f_0.02_10b_SLIM.dat",
    "Table"][[2]];
datp002R300h01s095T = Partition[Riffle[Partition[Riffle[Range[9],
      datp002R300h01s095], 2], Map[ErrorBar, datp002R300h01s095CI]], 2];
datp002R300h05s095T = Partition[Riffle[Partition[Riffle[Range[9],
      datp002R300h05s095], 2], Map[ErrorBar, datp002R300h05s095CI]], 2];
datp002R300h09s095T = Partition[Riffle[Partition[Riffle[Range[9],
      datp002R300h09s095], 2], Map[ErrorBar, datp002R300h09s095CI]], 2];
```

```
p1R300s095 = ListPlot[
  {Table[{l, PLNH2[5000, 0.05, 0.1, 0.95, 300, 10, l, 0.02, 4/(1 + 0.95/(2-0.95))]}, {l, 1, 9}],
   Table[{l, PLNH2[5000, 0.05, 0.5, 0.95, 300, 10, l, 0.02, 4/(1 + 0.95/(2-0.95))]}, {l, 1, 9}],
   Table[{l, PLNH2[5000, 0.05, 0.9, 0.95, 300, 10, l, 0.02, 4/(1 + 0.95/(2-0.95))]}, {l, 1, 9}],
   Table[{l, PrJ[l, 10]}, {l, 1, 9}]}, PlotRange → All,
  PlotStyle → {Red, Black, Blue, {Gray, Dashed}}, Joined → True]
```

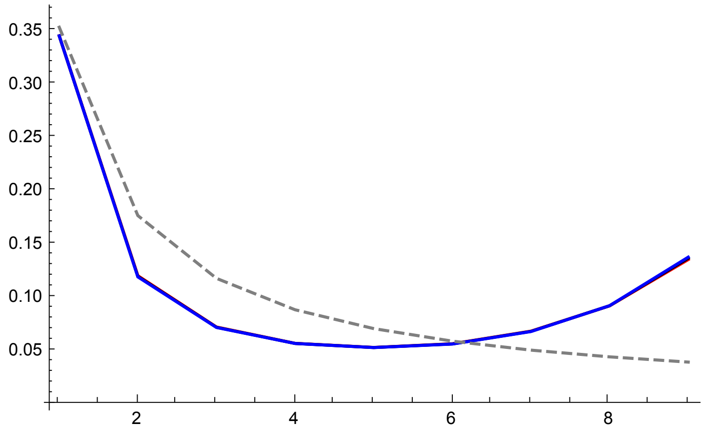

```
p2R300s095 = ErrorListPlot[{datp002R300h01s095T,
   datp002R300h05s095T, datp002R300h09s095T}, PlotRange → All, PlotStyle →
   {{Red, PointSize[0.02]}, {Black, PointSize[0.02]}, {Blue, PointSize[0.02]}}]
```

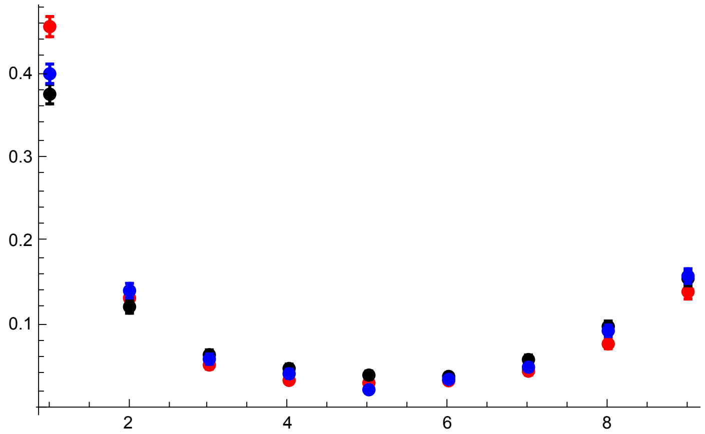

```
p3R300s095 = Show[p1R300s095, p2R300s095, PlotRange → All]
```

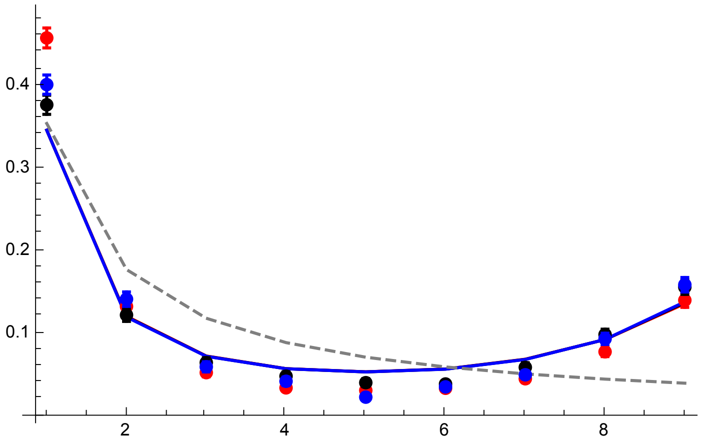


R = 500

```
datp002R500h01s095 =
  Import["SLIM_SFS_F09/SFSTab_R_500_h_0.1_self_0.95_f_0.02_10b_SLIM.dat",
    "Table"][[1]];
datp002R500h01s095CI = Import[
    "SLIM_SFS_F09/SFSTab_R_500_h_0.1_self_0.95_f_0.02_10b_SLIM.dat",
    "Table"][[2]];
datp002R500h05s095 = Import[
    "SLIM_SFS_F09/SFSTab_R_500_h_0.5_self_0.95_f_0.02_10b_SLIM.dat",
    "Table"][[1]];
datp002R500h05s095CI = Import[
    "SLIM_SFS_F09/SFSTab_R_500_h_0.5_self_0.95_f_0.02_10b_SLIM.dat",
    "Table"][[2]];
datp002R500h09s095 = Import[
    "SLIM_SFS_F09/SFSTab_R_500_h_0.9_self_0.95_f_0.02_10b_SLIM.dat",
    "Table"][[1]];
datp002R500h09s095CI = Import[
    "SLIM_SFS_F09/SFSTab_R_500_h_0.9_self_0.95_f_0.02_10b_SLIM.dat",
    "Table"][[2]];
datp002R500h01s095T = Partition[Riffle[Partition[Riffle[Range[9],
      datp002R500h01s095], 2], Map[ErrorBar, datp002R500h01s095CI]], 2];
datp002R500h05s095T = Partition[Riffle[Partition[Riffle[Range[9],
      datp002R500h05s095], 2], Map[ErrorBar, datp002R500h05s095CI]], 2];
datp002R500h09s095T = Partition[Riffle[Partition[Riffle[Range[9],
      datp002R500h09s095], 2], Map[ErrorBar, datp002R500h09s095CI]], 2];
```

```
p1R500s095 = ListPlot[
  {Table[{l, PLNH2[5000, 0.05, 0.1, 0.95, 500, 10, l, 0.02, 4/(1 + 0.95/(2-0.95))]}, {l, 1, 9}],
   Table[{l, PLNH2[5000, 0.05, 0.5, 0.95, 500, 10, l, 0.02, 4/(1 + 0.95/(2-0.95))]}, {l, 1, 9}],
   Table[{l, PLNH2[5000, 0.05, 0.9, 0.95, 500, 10, l, 0.02, 4/(1 + 0.95/(2-0.95))]}, {l, 1, 9}],
   Table[{l, PrJ[l, 10]}, {l, 1, 9}]}, PlotRange → All,
  PlotStyle → {Red, Black, Blue, {Gray, Dashed}}, Joined → True]
```

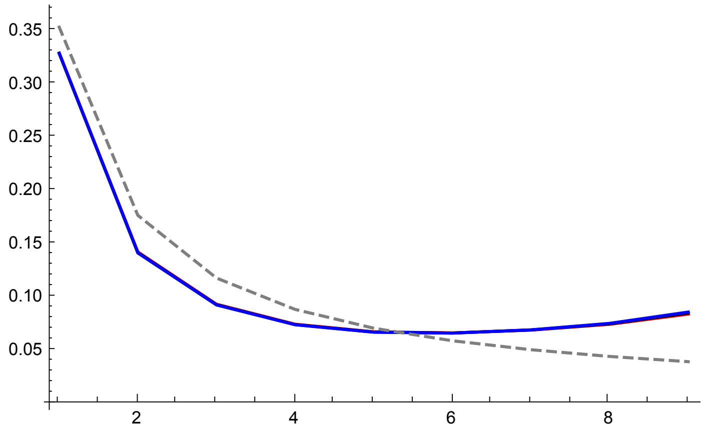


```
p2R500s095 = ErrorListPlot[{datp002R500h01s095T,
   datp002R500h05s095T, datp002R500h09s095T}, PlotRange → All, PlotStyle →
   {{Red, PointSize[0.02]}, {Black, PointSize[0.02]}, {Blue, PointSize[0.02]}}]
```

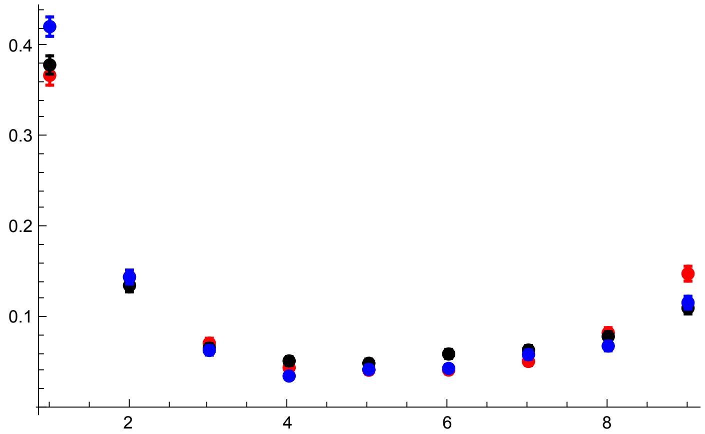

```
p3R500s095 = Show[p1R500s095, p2R500s095, PlotRange → All]
```

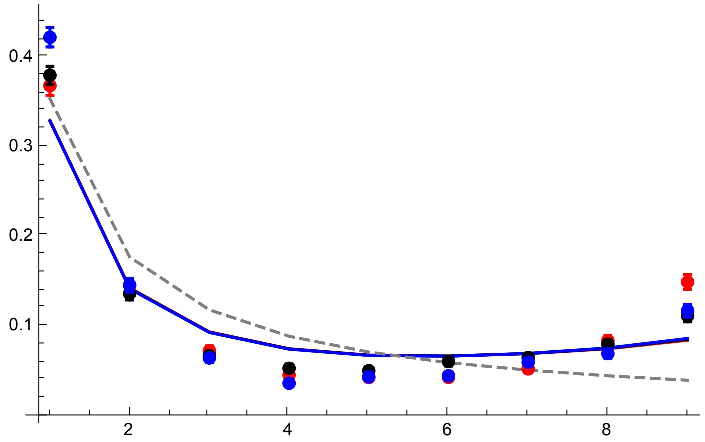


All plots compared

```
GraphicsGrid[{{p3R100s095, p3R300s095}, {p3R500s095,}}]
```

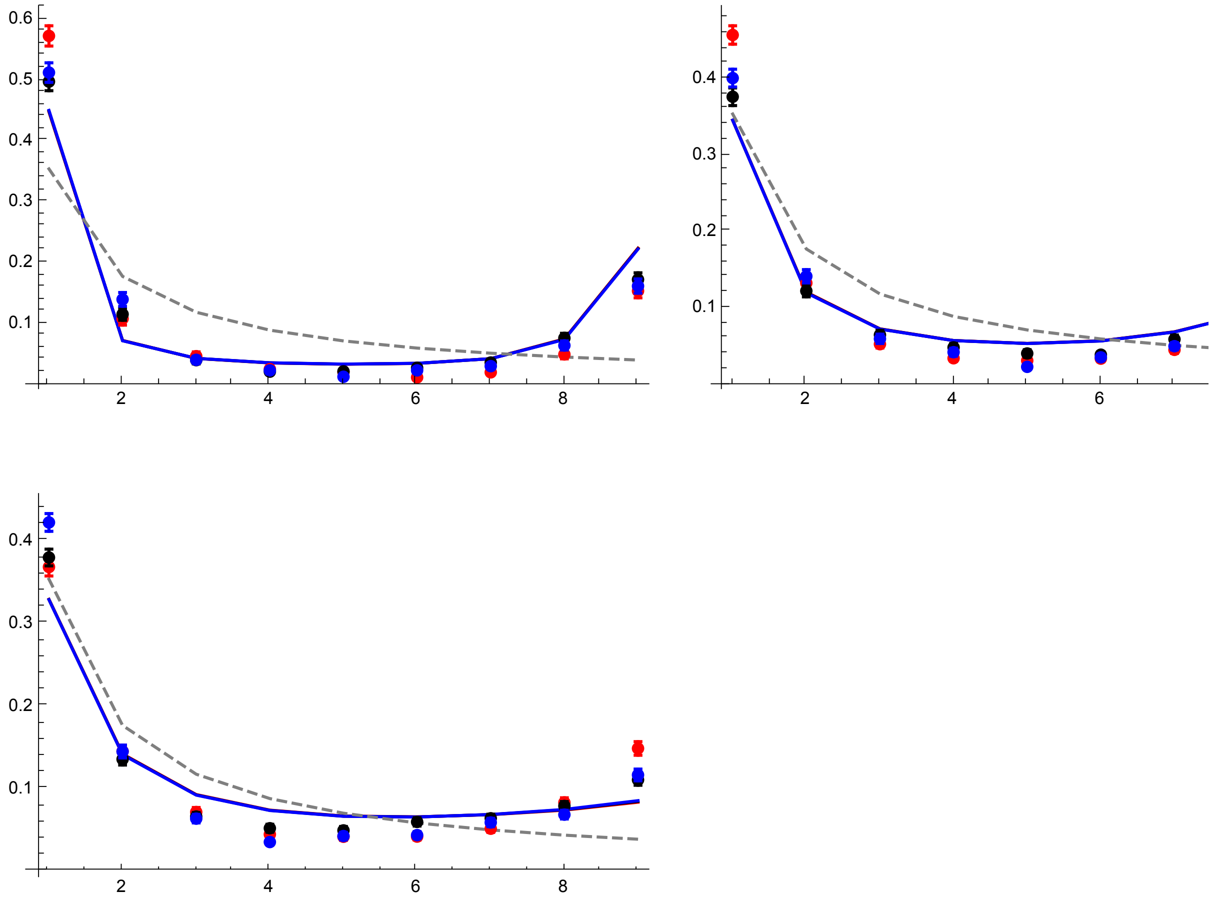

#### Results with $p_0 = 0.05$

R = 100

```
datp005R100h01s095 =
  Import["SLIM_SFS_F09/SFSTab_R_100_h_0.1_self_0.95_f_0.05_10b_SLIM.dat",
    "Table"][[1]];
datp005R100h01s095CI = Import[
    "SLIM_SFS_F09/SFSTab_R_100_h_0.1_self_0.95_f_0.05_10b_SLIM.dat",
    "Table"][[2]];
datp005R100h05s095 = Import[
    "SLIM_SFS_F09/SFSTab_R_100_h_0.5_self_0.95_f_0.05_10b_SLIM.dat",
    "Table"][[1]];
datp005R100h05s095CI = Import[
    "SLIM_SFS_F09/SFSTab_R_100_h_0.5_self_0.95_f_0.05_10b_SLIM.dat",
    "Table"][[2]];
datp005R100h09s095 = Import[
    "SLIM_SFS_F09/SFSTab_R_100_h_0.9_self_0.95_f_0.05_10b_SLIM.dat",
    "Table"][[1]];
datp005R100h09s095CI = Import[
    "SLIM_SFS_F09/SFSTab_R_100_h_0.9_self_0.95_f_0.05_10b_SLIM.dat",
    "Table"][[2]];
datp005R100h01s095T = Partition[Riffle[Partition[Riffle[Range[9],
      datp005R100h01s095], 2], Map[ErrorBar, datp005R100h01s095CI]], 2];
datp005R100h05s095T = Partition[Riffle[Partition[Riffle[Range[9],
      datp005R100h05s095], 2], Map[ErrorBar, datp005R100h05s095CI]], 2];
datp005R100h09s095T = Partition[Riffle[Partition[Riffle[Range[9],
      datp005R100h09s095], 2], Map[ErrorBar, datp005R100h09s095CI]], 2];
```

```
p1R100s095 = ListPlot[
  {Table[{l, PLNH2[5000, 0.05, 0.1, 0.95, 100, 10, l, 0.05, 4/(1 + 0.95/(2-0.95))]}, {l, 1, 9}],
   Table[{l, PLNH2[5000, 0.05, 0.5, 0.95, 100, 10, l, 0.05, 4/(1 + 0.95/(2-0.95))]}, {l, 1, 9}],
   Table[{l, PLNH2[5000, 0.05, 0.9, 0.95, 100, 10, l, 0.05, 4/(1 + 0.95/(2-0.95))]}, {l, 1, 9}],
   Table[{l, PrJ[l, 10]}, {l, 1, 9}]}, PlotRange → All,
  PlotStyle → {Red, Black, Blue, {Gray, Dashed}}, Joined → True]
```

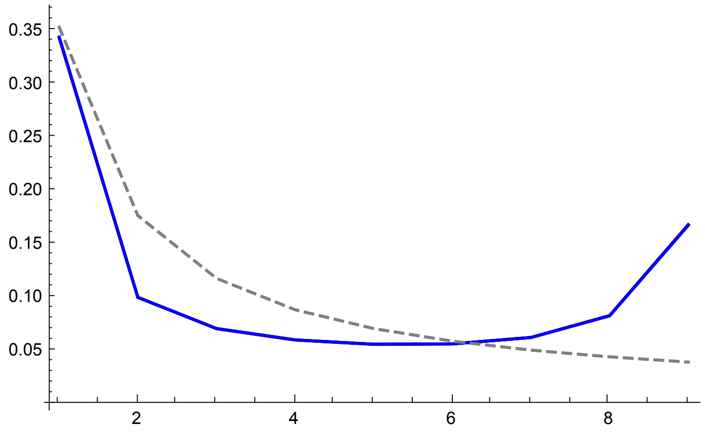


```
p2R100s095 = ErrorListPlot[{datp005R100h01s095T,
   datp005R100h05s095T, datp005R100h09s095T}, PlotRange → All, PlotStyle →
   {{Red, PointSize[0.02]}, {Black, PointSize[0.02]}, {Blue, PointSize[0.02]}}]
```

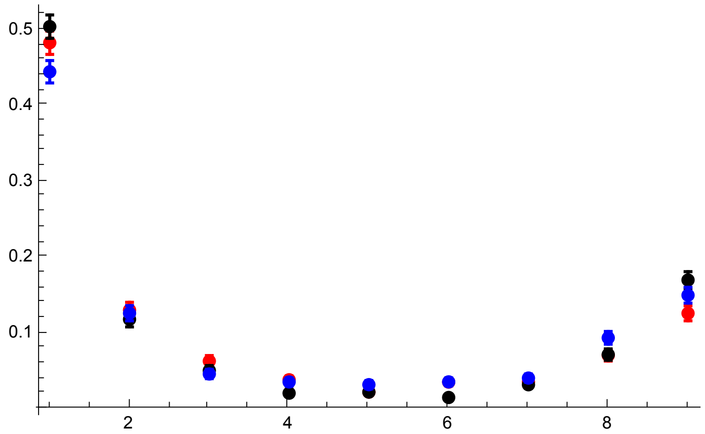

```
p3R100s095 = Show[p1R100s095, p2R100s095, PlotRange → All]
```

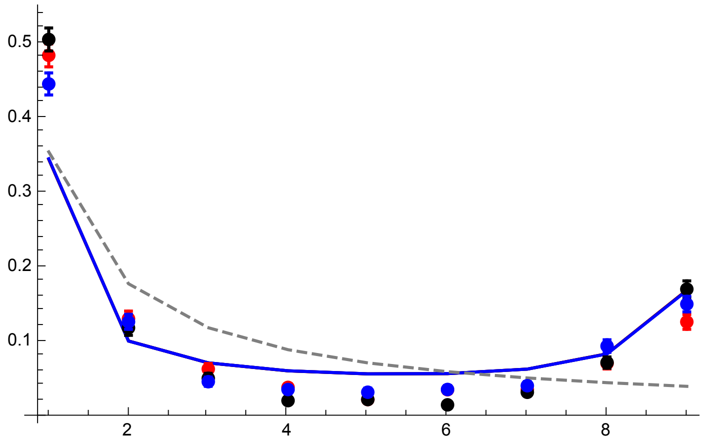


R = 300

```
datp005R300h01s095 =
  Import["SLIM_SFS_F09/SFSTab_R_300_h_0.1_self_0.95_f_0.05_10b_SLIM.dat",
    "Table"][[1]];
datp005R300h01s095CI = Import[
    "SLIM_SFS_F09/SFSTab_R_300_h_0.1_self_0.95_f_0.05_10b_SLIM.dat",
    "Table"][[2]];
datp005R300h05s095 = Import[
    "SLIM_SFS_F09/SFSTab_R_300_h_0.5_self_0.95_f_0.05_10b_SLIM.dat",
    "Table"][[1]];
datp005R300h05s095CI = Import[
    "SLIM_SFS_F09/SFSTab_R_300_h_0.5_self_0.95_f_0.05_10b_SLIM.dat",
    "Table"][[2]];
datp005R300h09s095 = Import[
    "SLIM_SFS_F09/SFSTab_R_300_h_0.9_self_0.95_f_0.05_10b_SLIM.dat",
    "Table"][[1]];
datp005R300h09s095CI = Import[
    "SLIM_SFS_F09/SFSTab_R_300_h_0.9_self_0.95_f_0.05_10b_SLIM.dat",
    "Table"][[2]];
datp005R300h01s095T = Partition[Riffle[Partition[Riffle[Range[9],
      datp005R300h01s095], 2], Map[ErrorBar, datp005R300h01s095CI]], 2];
datp005R300h05s095T = Partition[Riffle[Partition[Riffle[Range[9],
      datp005R300h05s095], 2], Map[ErrorBar, datp005R300h05s095CI]], 2];
datp005R300h09s095T = Partition[Riffle[Partition[Riffle[Range[9],
      datp005R300h09s095], 2], Map[ErrorBar, datp005R300h09s095CI]], 2];
```

```
p1R300s095 = ListPlot[
  {Table[{l, PLNH2[5000, 0.05, 0.1, 0.95, 300, 10, l, 0.05, 4/(1 + 0.95/(2-0.95))]}, {l, 1, 9}],
   Table[{l, PLNH2[5000, 0.05, 0.5, 0.95, 300, 10, l, 0.05, 4/(1 + 0.95/(2-0.95))]}, {l, 1, 9}],
   Table[{l, PLNH2[5000, 0.05, 0.9, 0.95, 300, 10, l, 0.05, 4/(1 + 0.95/(2-0.95))]}, {l, 1, 9}],
   Table[{l, PrJ[l, 10]}, {l, 1, 9}]}, PlotRange → All,
  PlotStyle → {Red, Black, Blue, {Gray, Dashed}}, Joined → True]
```

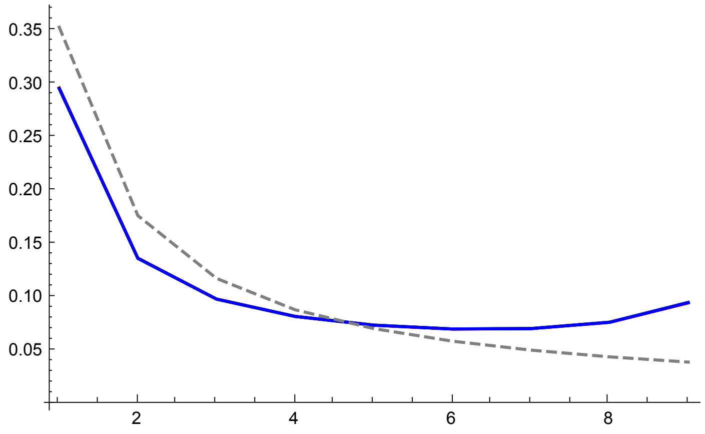


```
p2R300s095 = ErrorListPlot[{datp005R300h01s095T,
   datp005R300h05s095T, datp005R300h09s095T}, PlotRange → All, PlotStyle →
   {{Red, PointSize[0.02]}, {Black, PointSize[0.02]}, {Blue, PointSize[0.02]}}]
```

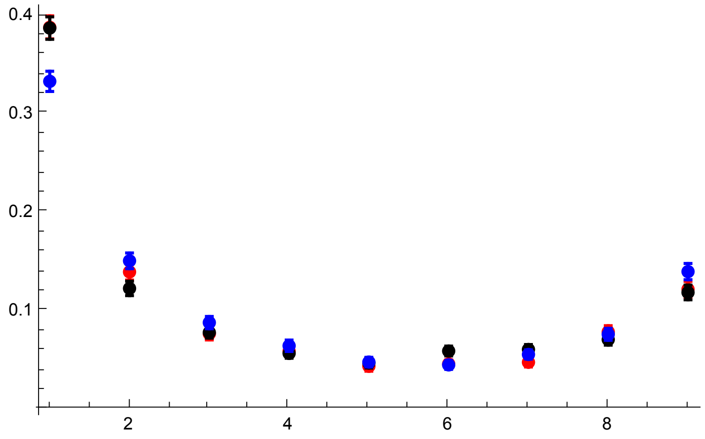

```
p3R300s095 = Show[p1R300s095, p2R300s095, PlotRange → All]
```

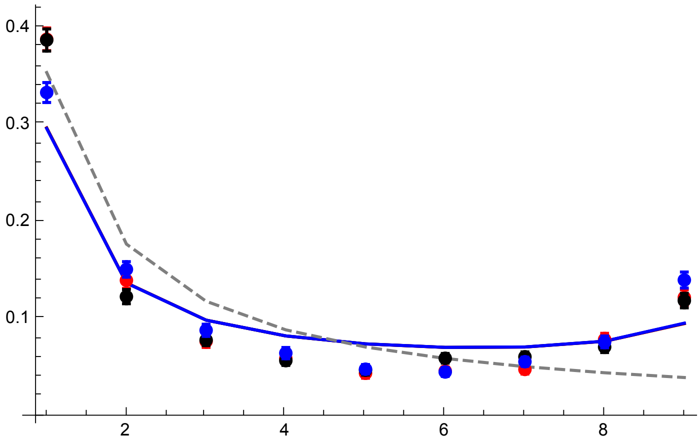


R = 500

```
datp005R500h01s095 =
  Import["SLIM_SFS_F09/SFSTab_R_500_h_0.1_self_0.95_f_0.05_10b_SLIM.dat",
    "Table"][[1]];
datp005R500h01s095CI = Import[
    "SLIM_SFS_F09/SFSTab_R_500_h_0.1_self_0.95_f_0.05_10b_SLIM.dat",
    "Table"][[2]];
datp005R500h05s095 = Import[
    "SLIM_SFS_F09/SFSTab_R_500_h_0.5_self_0.95_f_0.05_10b_SLIM.dat",
    "Table"][[1]];
datp005R500h05s095CI = Import[
    "SLIM_SFS_F09/SFSTab_R_500_h_0.5_self_0.95_f_0.05_10b_SLIM.dat",
    "Table"][[2]];
datp005R500h09s095 = Import[
    "SLIM_SFS_F09/SFSTab_R_500_h_0.9_self_0.95_f_0.05_10b_SLIM.dat",
    "Table"][[1]];
datp005R500h09s095CI = Import[
    "SLIM_SFS_F09/SFSTab_R_500_h_0.9_self_0.95_f_0.05_10b_SLIM.dat",
    "Table"][[2]];
datp005R500h01s095T = Partition[Riffle[Partition[Riffle[Range[9],
      datp005R500h01s095], 2], Map[ErrorBar, datp005R500h01s095CI]], 2];
datp005R500h05s095T = Partition[Riffle[Partition[Riffle[Range[9],
      datp005R500h05s095], 2], Map[ErrorBar, datp005R500h05s095CI]], 2];
datp005R500h09s095T = Partition[Riffle[Partition[Riffle[Range[9],
      datp005R500h09s095], 2], Map[ErrorBar, datp005R500h09s095CI]], 2];
```

```
p1R500s095 = ListPlot[
  {Table[{l, PLNH2[5000, 0.05, 0.1, 0.95, 500, 10, l, 0.05, 4/(1 + 0.95/(2-0.95))]}, {l, 1, 9}],
   Table[{l, PLNH2[5000, 0.05, 0.5, 0.95, 500, 10, l, 0.05, 4/(1 + 0.95/(2-0.95))]}, {l, 1, 9}],
   Table[{l, PLNH2[5000, 0.05, 0.9, 0.95, 500, 10, l, 0.05, 4/(1 + 0.95/(2-0.95))]}, {l, 1, 9}],
   Table[{l, PrJ[l, 10]}, {l, 1, 9}]}, PlotRange → All,
  PlotStyle → {Red, Black, Blue, {Gray, Dashed}}, Joined → True]
```

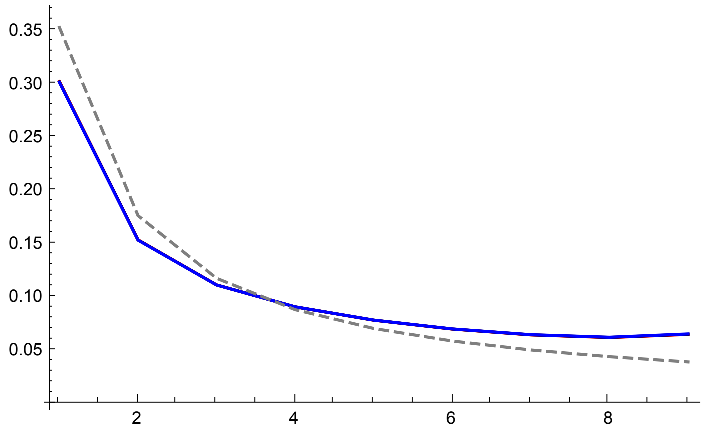


```
p2R500s095 = ErrorListPlot[{datp005R500h01s095T,
   datp005R500h05s095T, datp005R500h09s095T}, PlotRange → All, PlotStyle →
   {{Red, PointSize[0.02]}, {Black, PointSize[0.02]}, {Blue, PointSize[0.02]}}]
```

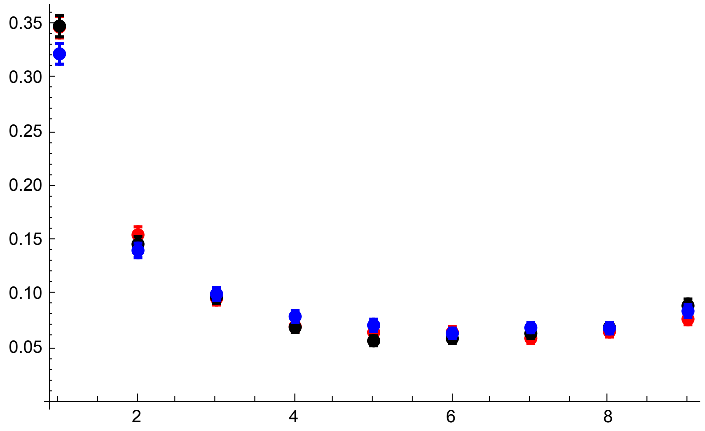

```
p3R500s095 = Show[p1R500s095, p2R500s095, PlotRange → All]
```

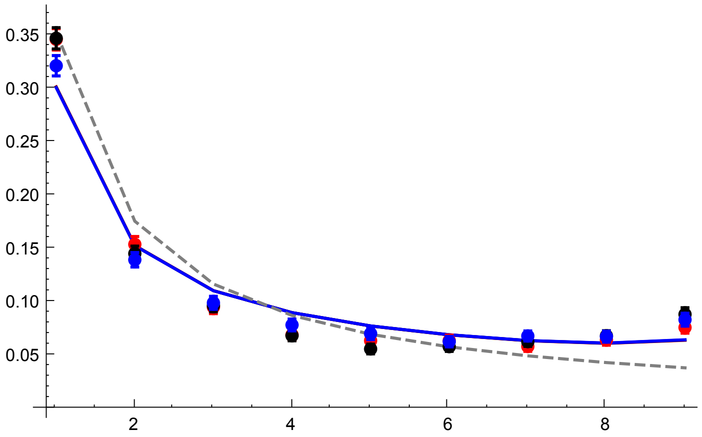


All plots compared

```
GraphicsGrid[{{p3R100s095, p3R300s095}, {p3R500s095,}}]
```

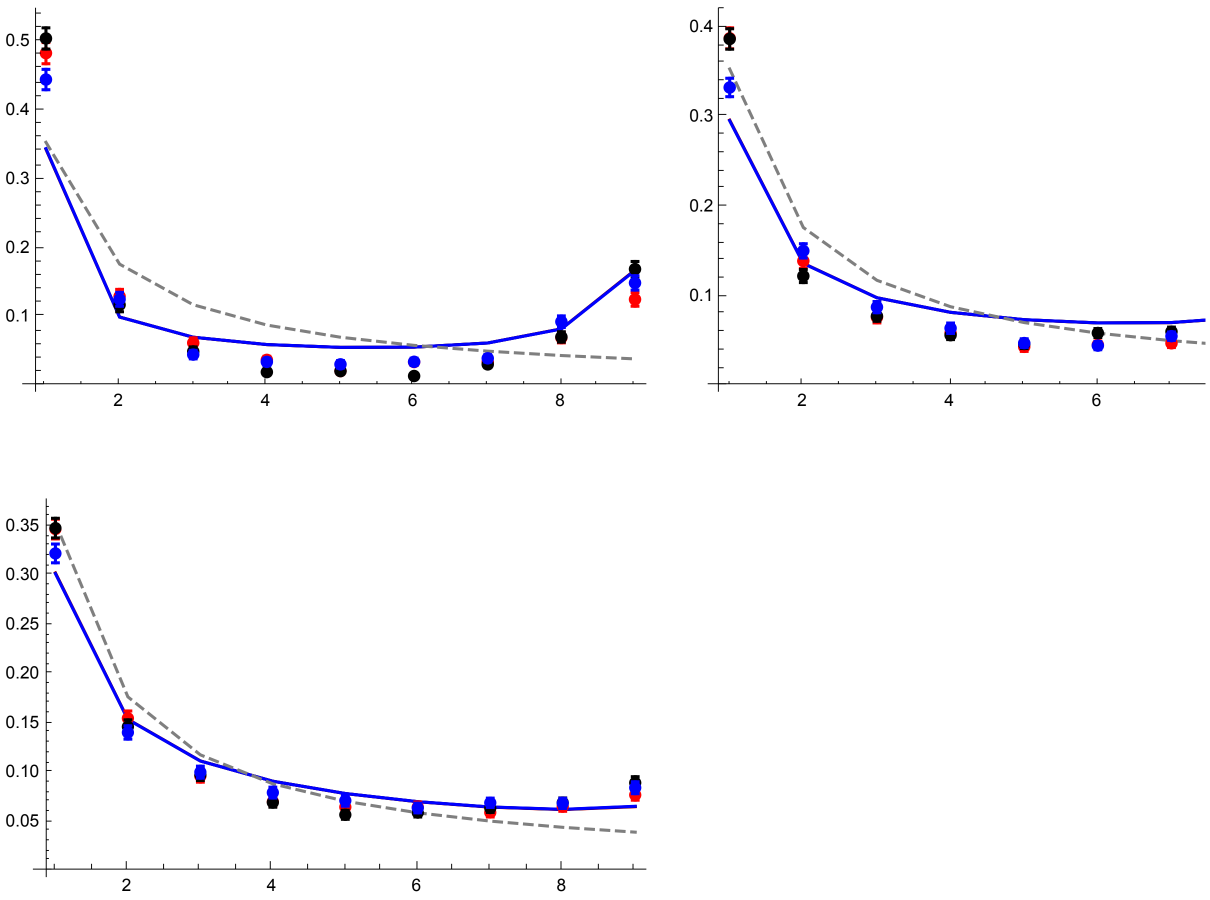


### Grid of key results

#### Comparing analytical results and SLiM simulations

Full list:

```
SimCompSFS = Labeled[
  Grid[{{Text@TraditionalForm@Style["p₀ = 1/2N", 24], Text@TraditionalForm@
       Style["p₀ = 0.02", 24], Text@TraditionalForm@Style["p₀ = 0.05", 24],},
    {p3AR6s0DN, p3AR6s0p02, p3AR6s0p05, Text@TraditionalForm@
       Style["σ = 0.00\n(F = 0.00,\nR = 6)", 24]},
    {p3R11s05DN, p3R11s05p02, p3R11s05p05,
     Text@TraditionalForm@Style["σ = 0.50\n(F = 0.33,\nR = 11)", 24]}},
   Spacings → {2, 1}], {Text@TraditionalForm@Style["Frequency", 24],
   Text@TraditionalForm@Style["Derived Allele Count", 24]}, {Left, Bottom}]
```

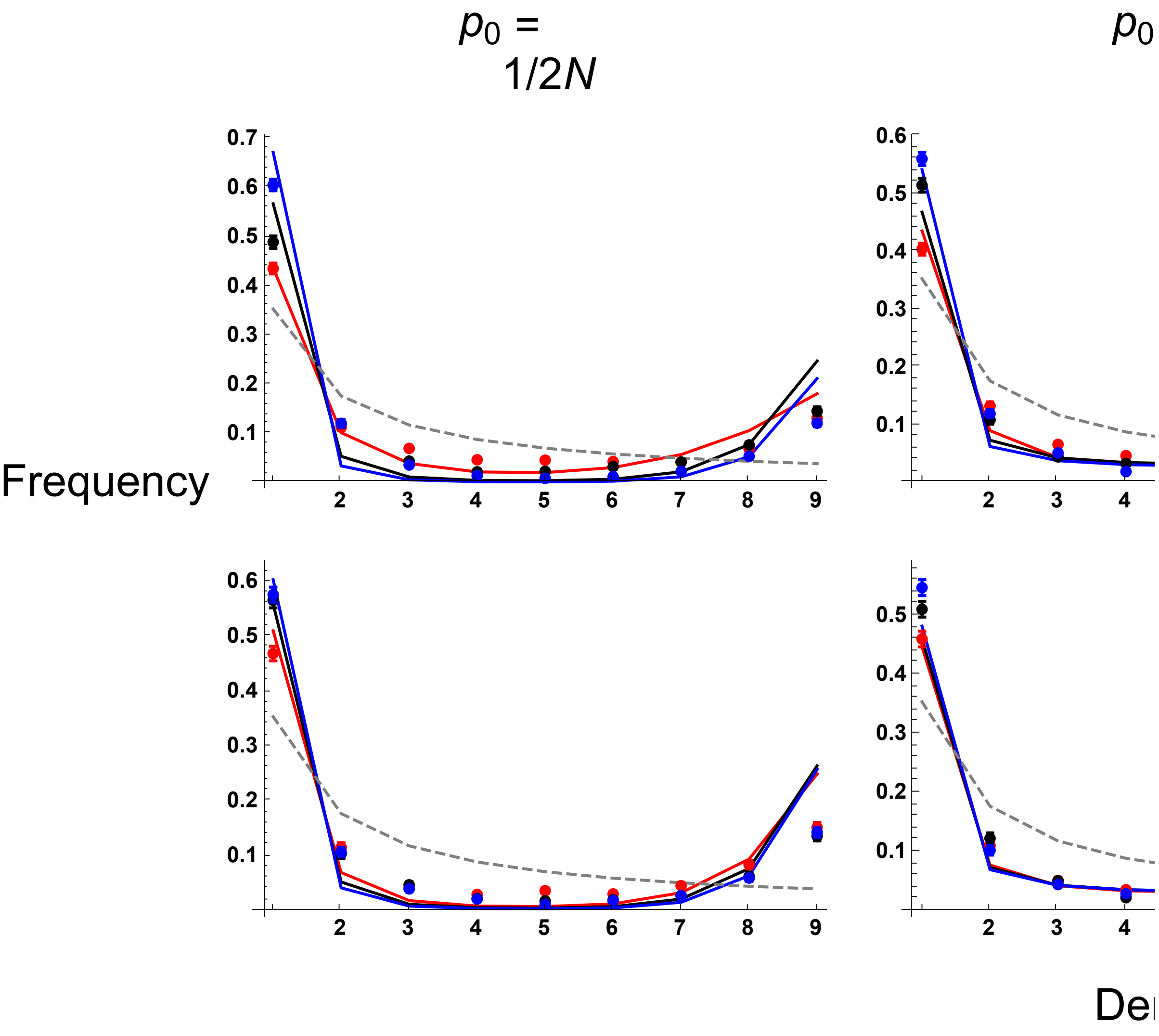


Reduced grid:

```
SimCompSFS2 = Labeled[Grid[{{Text@TraditionalForm@Style["p0 = 1/2N", 24],
    Text@TraditionalForm@Style["p0 = 0.05", 24],}, {p3AR6s0DN, p3AR6s0p05,
    Text@TraditionalForm@Style["σ = 0.00\n(F = 0.00,\nR = 6)", 24]},
   {p3R11s05DN, p3R11s05p05,
    Text@TraditionalForm@Style["σ = 0.50\n(F = 0.33,\nR = 11)", 24]}},
  Spacings → {2, 1}], {Text@TraditionalForm@Style["Frequency", 24],
  Text@TraditionalForm@Style["Derived Allele Count", 24]}, {Left, Bottom}]
```

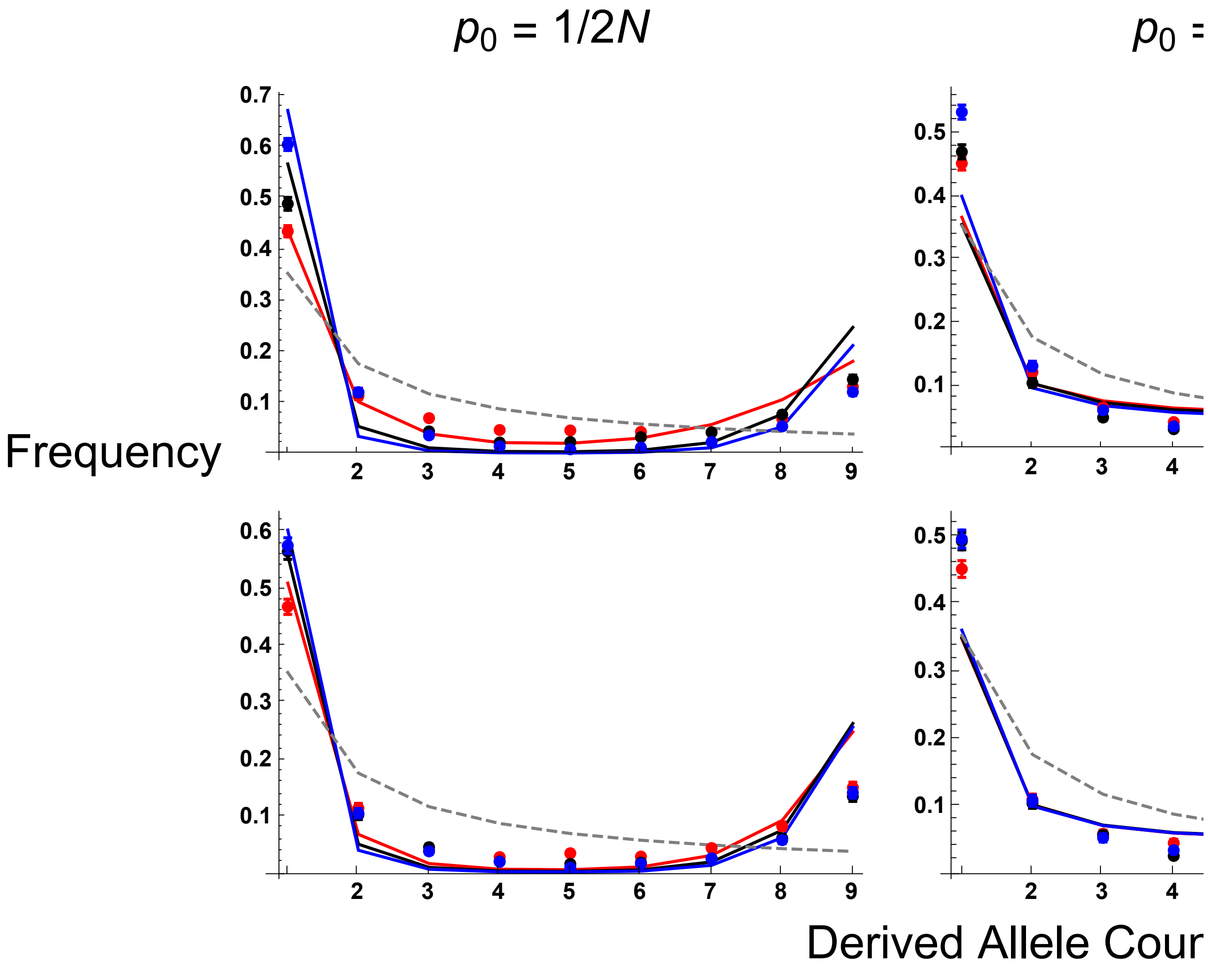

#### Comparing SLiM and MSMS simulations ($\sigma = 0$)

```
SLIMandMSMSCompSFS =
 Labeled[Grid[{{Text@TraditionalForm@Style["p0 = 1/2N", 24], Text@
      TraditionalForm@Style["p0 = 0.02", 24], Text@TraditionalForm@
       Style["p0 = 0.05", 24],}, {Text@TraditionalForm@Style[
        "Red points: forward-in-time\nPink points: coalescent simulations",
        14, TextAlignment → Center], Text@TraditionalForm@Style[
        "Black points: forward-in-time\nGrey points: coalescent simulations",
        14, TextAlignment → Center], Text@TraditionalForm@Style[
        "Blue points: forward-in-time\nCyan points: coalescent simulations",
        14, TextAlignment → Center],},
    {p3R6s0DN, p3R6s0p02, p3R6s0p05, Text@TraditionalForm@Style["R = 6", 24]},
    {p3R18s0DN, p3R18s0p02, p3R18s0p05,
     Text@TraditionalForm@Style["R = 18", 24]}}, Spacings → {2, 1}],
  {Text@TraditionalForm@Style["Frequency", 24],
   Text@TraditionalForm@Style["Derived Allele Count", 24]}, {Left,
   Bottom}]
```

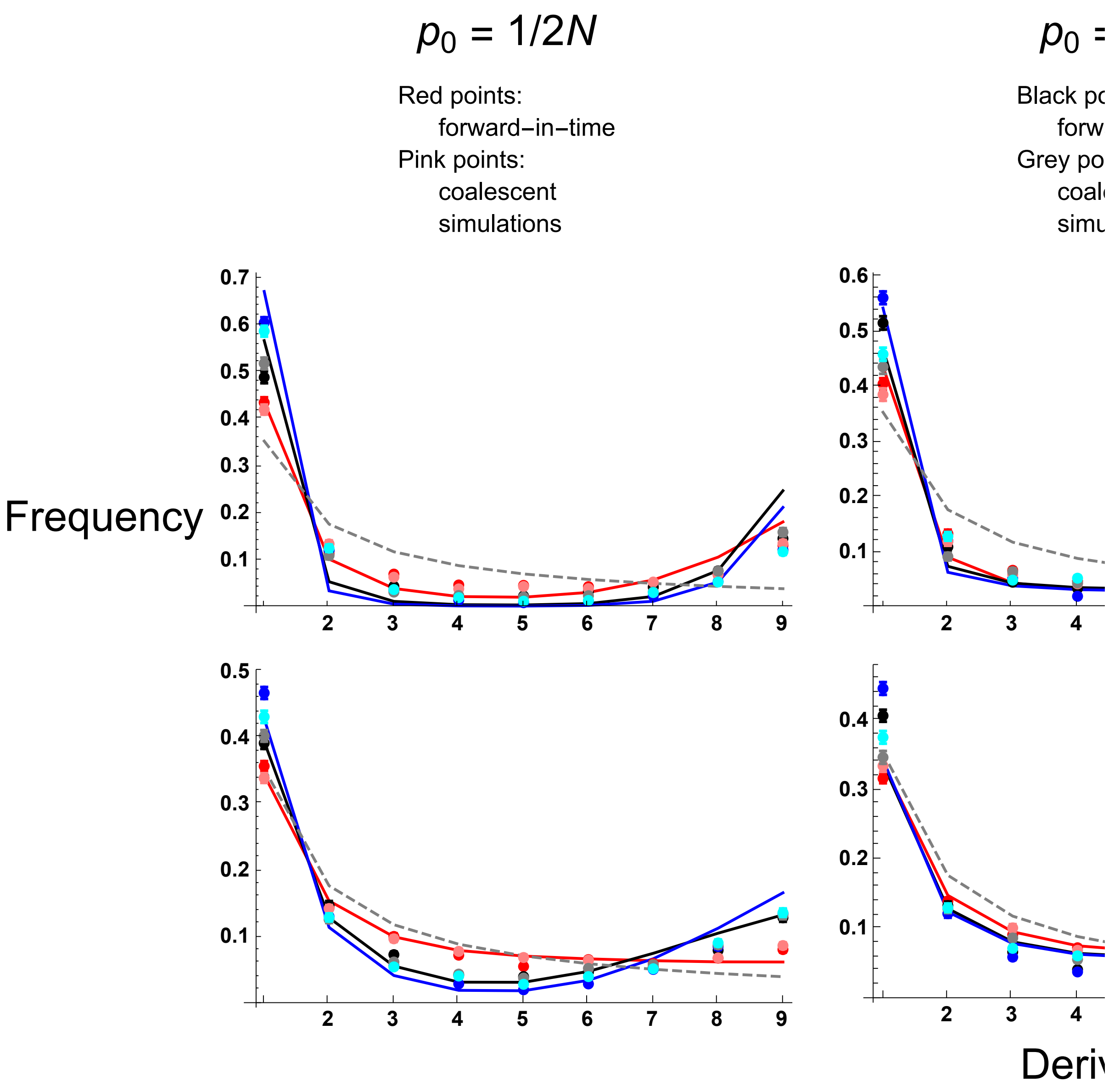


### Section F: Comparing sweeps from standing variation to sweeps from recurrent mutation

Clearing memory before running this section to avoid conflicts with previous sections

```
ClearAll["`*"];
Needs["ErrorBarPlots`"]
```

#### Model description

In this scenario, there are five possible outcomes:

(1) Coalescence during the sweep phase;

(2) Recombination during the sweep phase;

(3) Mutation during the sweep phase;

(4) Coalescence at the allele origin;
(5) Mutation at the allele origin.

Note that we do not consider recombination during the ‘standing phase’ since it does not exist in this model.

If events (1), (4) occur then $\pi \simeq 0$.
If events (2), (3), (5) occur then $\pi \simeq \pi_0$, the background levels of diversity.
Hence $\mathbb{E}(\pi/\pi_0)$ = P(Event 2) + P(Event 3) + P(Event 5).

Going through these in turn;
To calculate P(Event 2), we need to determine the probabilities that
(i) no event (coalescence, recombination, mutation) occurs in the sweep phase when the derived allele is between a frequency of *p* to 1;
(ii) recombination occurs at frequency *p*;
(iii) integrating this solution over all frequencies *p0* to 1.
P(Event 3) is calculated in a similar way, but instead at point (ii) we calculate the probability that mutation occurs at frequency *p*.

To calculate P(Event 5), we need to determine the probabilities that
(i) no event occurs in the sweep phase when the derived allele is between a frequency of *p0,A* to 1;
(ii) Mutation occurs when the derived allele first appears.

Let’s look at the relative probability of each event over each timestep.
If the frequency of the beneficial allele is *p* at a certain time, then the probability of coalescence is $\frac{1}{2\,\mathrm{Ne}\,p} = \frac{1+F}{2\,N\,p}$.
The probability of one of the two samples recombining out is 2r(1-2F+Φ)(1-p).
The probability of one of the two samples mutating is 2$\mu$(1-p)/p.

The probability of no action occurring at any timepoint is $1 - \frac{1}{2\,\mathrm{Ne}\,p} - 2r(1\text{-}2F\text{+}\Phi)(1\text{-}p) - 2\mu\frac{(1-p)}{p}$. The total probability of neither action over the entire sweep phase is
$\simeq \mathrm{Exp}\left[-\int_{t=0}^{\tau f}\left(\frac{1+F}{2\,N\,p} + 2\,r(1-2\,F+\Phi)\,(1-p) + \frac{2\,\mu(1-p)}{p}\right) dt\right]$ for fixation time $\tau$f, or
$$\int_{p=1-\epsilon 1}^{p0} \frac{\left(\frac{1+F}{2\,N\,p} + 2\,r\,(1-2\,F+\Phi)\,(1-p) + \frac{2\,\mu\,(1-p)}{p}\right)}{dp/dt}\,dp.$$

The sum of the coalescence and recombination probabilities are:

$$2\,r\,(1-2\,F+\Phi)\,(1-p) + \frac{1+F}{2\,\mathrm{Na}\,p} + \frac{2\,\mu\,(1-p)}{p} \text{ // Together}$$

$$\frac{1}{2\,\mathrm{Na}\,p}\left(1 + F + 4\,\mathrm{Na}\,p\,r - 8\,F\,\mathrm{Na}\,p\,r - 4\,\mathrm{Na}\,p^2\,r + 8\,F\,\mathrm{Na}\,p^2\,r + 4\,\mathrm{Na}\,\mu - 4\,\mathrm{Na}\,p\,\mu + 4\,\mathrm{Na}\,p\,r\,\Phi - 4\,\mathrm{Na}\,p^2\,r\,\Phi\right)$$

Tidying up:

```
1/(2 Na p) (1 + F + 4 Na p (1 - p) (1 - 2 F + Φ) r + 4 Na (1 - p) μ) -
  {1/(2 Na p) (1 + F + 4 Na p r - 8 F Na p r - 4 Na p^2 r + 8 F Na p^2 r +
      4 Na μ - 4 Na p μ + 4 Na p r Φ - 4 Na p^2 r Φ)} // Simplify
```

{0}

Dividing by $-s\,(1-p)\,p\,(F+h-F\,h+(1-F)\,(1-2\,h)\,p)$ and taking the indefinite integral;

```
Integrate[-((1 + F + 4 Na p (1 - p) (1 - 2 F + Φ) r + 4 Na (1 - p) μ) /
    ((2 Na p) (s (1 - p) p (F + h - F h + (1 - F) (1 - 2 h) p)))), p] // Simplify
```

$$\frac{1}{2\,\text{Na}\,s}\left(\frac{1+F+4\,\text{Na}\,\mu}{(F+h-F\,h)\,p}+\frac{(1+F)\,\text{Log}[1-p]}{1+(-1+F)\,h}-\left(\left(-1+F+F^2\,(2-8\,\text{Na}\,r+h\,(-3+8\,\text{Na}\,r))-4\,\text{Na}\,\mu+h\,(3+4\,\text{Na}\,(r+2\,\mu+r\,\Phi))-4\,F\,\text{Na}\,((-1+2\,h)\,\mu+r\,(-1-\Phi+h\,(3+\Phi)))\right)\text{Log}[p]\right)\Big/(F+h-F\,h)^2+\frac{1}{(1+(-1+F)\,h)\,(F+h-F\,h)^2}\left(-1+F^3\left(-1+h\,(4-8\,\text{Na}\,r)+h^2\,(-4+8\,\text{Na}\,r)\right)-4\,\text{Na}\,\mu-4\,h^2\,(1+\text{Na}\,(r+2\,\mu+r\,\Phi))+4\,h\,(1+\text{Na}\,(r+3\,\mu+r\,\Phi))+F\left(1+4\,\text{Na}\,(r+\mu+r\,\Phi)-4\,h\,(1+4\,\text{Na}\,\mu+2\,\text{Na}\,r\,(2+\Phi))+4\,h^2\,(1+4\,\text{Na}\,\mu+2\,\text{Na}\,r\,(2+\Phi))\right)+F^2\left(1-8\,\text{Na}\,r-4\,h^2\,(-1+2\,\text{Na}\,\mu+\text{Na}\,r\,(5+\Phi))+4\,h\,(-1+\text{Na}\,(\mu+r\,(5+\Phi)))\right)\right)\text{Log}[F+h+p-F\,p-2\,h\,p+F\,h\,(-1+2\,p)]\right)$$

```
SweepIntFHM[Na_, s_, h_, F_, Φ_, r_, μ_, p_] :=
  1/(2 Na s) ((1 + F + 4 Na μ)/((F + h - F h) p) + ((1 + F) Log[1 - p])/(1 + (-1 + F) h) -
     ((-1 + F + F^2 (2 - 8 Na r + h (-3 + 8 Na r)) - 4 Na μ + h (3 + 4 Na (r + 2 μ + r Φ)) -
         4 F Na ((-1 + 2 h) μ + r (-1 - Φ + h (3 + Φ)))) Log[p]) / (F + h - F h)^2 +
     1/((1 + (-1 + F) h) (F + h - F h)^2) (-1 + F^3 (-1 + h (4 - 8 Na r) + h^2 (-4 + 8 Na r)) - 4 Na μ -
        4 h^2 (1 + Na (r + 2 μ + r Φ)) + 4 h (1 + Na (r + 3 μ + r Φ)) + F (1 + 4 Na (r + μ + r Φ) -
           4 h (1 + 4 Na μ + 2 Na r (2 + Φ)) + 4 h^2 (1 + 4 Na μ + 2 Na r (2 + Φ))) +
        F^2 (1 - 8 Na r - 4 h^2 (-1 + 2 Na μ + Na r (5 + Φ)) + 4 h (-1 + Na (μ + r (5 + Φ)))))
      Log[F + h + p - F p - 2 h p + F h (-1 + 2 p)])
```

We can use this integral to calculate (A) the probability of recombination (mutation) during the sweep phase; (B) the probability of coalescence or mutation at the sweep origin, given no actions during the sweep phase.

The total probability of no event occurring for frequency *p* is the difference of this integral between p and 1 - $p_{0,A}$(1-h) (where $p_{0,A}\,(1-h)=\frac{1+F}{4\,\text{Na}\,s\,(1-h)}$ is the elevated ending frequency for a selected allele going to fixation with dominance *h*). This term is then multiplied by (2 r (1-2F+Φ)(1-

p)) to determine the probability that recombination acts at frequency *p;* or $\frac{2\mu(1-p)}{p}$ for the probability that mutations acts at frequency *p*.

```
Boostp0[Na_, s_, h_, F_] := (1 + F) / (4 Na s (F + h - F h))
```

```
PRecPFHM[Na_, s_, h_, F_, Φ_, r_, μ_, p_] :=
  (2 r (1 - 2 F + Φ) (1 - p)) Exp[- (SweepIntFHM[Na, s, h, F, Φ, r, μ, p] -
      SweepIntFHM[Na, s, h, F, Φ, r, μ, 1 - Boostp0[Na, s, 1 - h, F]])]
```

```
PMutPFHM[Na_, s_, h_, F_, Φ_, r_, μ_, p_] :=
  (2 μ (1 - p)) / p Exp[- (SweepIntFHM[Na, s, h, F, Φ, r, μ, p] -
      SweepIntFHM[Na, s, h, F, Φ, r, μ, 1 - Boostp0[Na, s, 1 - h, F]])]
```

(A) can be calculated by integrating the total recombination probability between $p_{0,A}$(h) and 1 - $p_{0,A}$(1-h), again after diving by *d*p/*d*t to convert from a time integral into a frequency integral. It does not appear that this integral has an analytical solution, so this integral is instead integrated numerically:

```
PRecFHO[Na_, s_, h_, F_, Φ_, r_, μ_] := NIntegrate[
   PRecPFHM[Na, s, h, F, Φ, r, μ, p] / (s (1 - p) p (F + h - F h + (1 - F) (1 - 2 h) p)),
   {p, Boostp0[Na, s, h, F], 1 - Boostp0[Na, s, 1 - h, F]}]
```

```
PMutFHO[Na_, s_, h_, F_, Φ_, r_, μ_] := NIntegrate[
   PMutPFHM[Na, s, h, F, Φ, r, μ, p] / (s (1 - p) p (F + h - F h + (1 - F) (1 - 2 h) p)),
   {p, Boostp0[Na, s, h, F], 1 - Boostp0[Na, s, 1 - h, F]}]
```

(B) can be calculated by working out (i) the probability of no event acting between frequency $p_{0,A}$(h) and $1 - p_{0,A}(1 - h)$, and (ii) multiplying it by the probability that a mutation event takes precedence over coalescence at the sweep origin.

(i) is obtained by taking the integral solution (as defined in ‘SweepIntFHM’). For (ii), the probability is equal to:

```
(2 μ (1-p0) / p0) / ((1+F) / (2 Na p0) + 2 μ (1-p0) / p0) // FullSimplify
```

$$-\frac{4\,\text{Na}\,(-1+\text{p0})\,\mu}{1+\text{F}-4\,\text{Na}\,(-1+\text{p0})\,\mu}$$

Which can be rewritten as:

```
(4 Na (1 - p0) μ) / (1 + F + 4 Na (1 - p0) μ) - {- (4 Na (-1 + p0) μ) / (1 + F - 4 Na (-1 + p0) μ)} // Simplify
```

{0}

This result can be further simplified by assuming 1-p0 ≃ 1 and substituting in $\Theta_b = 2\,N\mu$:

```
4 Na (1 - p0) μ / (1 + F + 4 Na (1 - p0) μ) /. μ → Θb / (2 Na) // Simplify
```

$$\frac{2\,(1-p0)\,\Theta b}{1+F-2\,(-1+p0)\,\Theta b}$$

```
2 (1 - p0) (Θb/(1+F)) / (1 + 2 (1 - p0) (Θb/(1+F))) - {2 (1 - p0) Θb / (1 + F - 2 (-1 + p0) Θb)} // Simplify
```

{0}

The recombination rate ($R_{\text{lim}}$) at which the coalescence probability for the std var case equates that for the mutation case:

```
Solve[1 / (1 + (2 R/(1+F)) (1 - 2 F + Φ) p0 (1 - p0)) == 1 / (1 + 2 (1 - p0) (Θb/(1+F))), R]
```

$$\left\{\left\{R \to -\frac{\Theta b}{p0\,(-1+2\,F-\Phi)}\right\}\right\}$$

Below we define (i) the probability of no event between $p_{0,A}(h)$ and $1 - p_{0,A}(1-h)$, and the probability of mutation during the standing phase if no event occurs in the sweep phase.

```
PNoActFHM[Na_, s_, h_, F_, Φ_, r_, μ_] :=
  Exp[-(SweepIntFHM[Na, s, h, F, Φ, r, μ, Boostp0[Na, s, h, F]] -
      SweepIntFHM[Na, s, h, F, Φ, r, μ, 1 - Boostp0[Na, s, 1 - h, F]])]

PMutFHM[Na_, s_, h_, F_, Φ_, r_, μ_] :=
  PNoActFHM[Na, s, h, F, Φ, r, μ] * (4 Na (1 - Boostp0[Na, s, h, F]) μ / (1 + F + 4 Na (1 - Boostp0[Na, s, h, F]) μ))
```

$\mathbb{E}[\pi/\pi_0]$ is the sum of the probabilities that recombination occurs during the sweep phase and standing phase respectively:

```
F[σ_] := σ / (2 - σ)
Φ[r_, σ_] := σ (2 - σ - 2 (1 - r) r (2 - 3 σ)) / ((2 - σ) (2 - (1 - 2 (1 - r) r) σ))
ExPiMut[Na_, s_, h_, σ_, r_, μ_] := PRecFHO[Na, s, h, F[σ], Φ[r, σ], r, μ] +
   PMutFHO[Na, s, h, F[σ], Φ[r, σ], r, μ] + PMutFHM[Na, s, h, F[σ], Φ[r, σ], r, μ]
```

#### Plots - reduction in pairwise diversity

Below are the equations for $\pi/\pi_0$ for std var sweep

```
SweepIntFH[Na_, s_, h_, F_, Φ_, r_, p_] :=
 - 1/(2 Na s) (- (1 + F)/((F + h - F h) p) - ((1 + F) Log[1 - p])/(1 + (-1 + F) h) + ((-1 + F + F^2 (2 - 8 Na r + h (-3 + 8 Na r)) +
        h (3 + 4 Na r (1 + Φ)) - 4 F Na r (-1 - Φ + h (3 + Φ))) Log[p]) / (F + h - F h)^2 -
    1/((1 + (-1 + F) h) (F + h - F h)^2) (-1 + F^3 (-1 + h (4 - 8 Na r) + h^2 (-4 + 8 Na r)) +
       4 h (1 + Na r (1 + Φ)) - 4 h^2 (1 + Na r (1 + Φ)) +
       F (1 + 4 Na r (1 + Φ) - 4 h (1 + 2 Na r (2 + Φ)) + 4 h^2 (1 + 2 Na r (2 + Φ))) +
       F^2 (1 - 8 Na r + 4 h (-1 + Na r (5 + Φ)) - 4 h^2 (-1 + Na r (5 + Φ))))
     Log[F + h + p - F p - 2 h p + F h (-1 + 2 p)])
PRecPFH[Na_, s_, h_, F_, Φ_, r_, p_] := (2 r (1 - 2 F + Φ) (1 - p))
  Exp[- (SweepIntFH[Na, s, h, F, Φ, r, p] -
      SweepIntFH[Na, s, h, F, Φ, r, 1 - Boostp0[Na, s, 1 - h, F]])]
PRecFHp0[Na_, s_, h_, F_, Φ_, r_, p0_] := NIntegrate[PRecPFH[Na, s, h, F, Φ, r, p] /
   (s (1 - p) p (F + h - F h + (1 - F) (1 - 2 h) p)), {p, p0, 1 - Boostp0[Na, s, 1 - h, F]}]
PNoActFH[Na_, s_, h_, F_, Φ_, r_, p0_] := Exp[- (SweepIntFH[Na, s, h, F, Φ, r, p0] -
     SweepIntFH[Na, s, h, F, Φ, r, 1 - Boostp0[Na, s, 1 - h, F]])]
PRecp0FH[Na_, s_, h_, F_, Φ_, r_, p0_] := PNoActFH[Na, s, h, F, Φ, r, p0] *
  ((4 Na r (1 - 2 F + Φ) (1 - p0) p0)/(1 + F + 4 Na r (1 - 2 F + Φ) (1 - p0) p0))
ExPiSV[Na_, s_, h_, σ_, r_, p0_] := PRecp0FH[Na, s, h, F[σ], Φ[r, σ], r, p0] +
  PRecFHp0[Na, s, h, F[σ], Φ[r, σ], r, p0]
```

‘PiSw’ is the recombination value, above which the standing variation model shows higher diversity than the mutation model. Note that we use the approximate recombination scaling (1–F) instead of (1–2F+Φ):

```
PiSw[Θ_, F_, p0_] := Θ/((1 - F) p0)
```

Comparing std var (solid lines) to mutation case (dashed lines). f0 = 5%, F = 0.
Here, the recurrent mutation case gives different outcomes with varying dominance coefficients, while standing variation case does not.

```
ExPis0 = Plot[{ExPiSV[5000, 0.05, 0.5, 0, R/10000, 0.05],
    ExPiMut[5000, 0.05, 0.5, 0, R/10000, 0.2/10000],
    ExPiSV[5000, 0.05, 0.1, 0, R/10000, 0.05], ExPiMut[5000, 0.05,
     0.1, 0, R/10000, 0.2/10000], ExPiSV[5000, 0.05, 0.9, 0, R/10000, 0.05],
    ExPiMut[5000, 0.05, 0.95, 0, R/10000, 0.2/10000]}, {R, 0, 120},
  PlotRange → All, PlotStyle → {{Black, Thick}, {Black, Dashed, Thick},
    {Red, Thick}, {Red, Dashed, Thick}, {Blue}, {Blue, Dashed, Thick}},
  GridLines → {{PiSw[0.2, 0, 0.05]}, {}}, Frame → {{True, False}, {True, False}},
  BaseStyle → {FontWeight → "Bold", FontColor → Black, FontSize → 16},
  GridLinesStyle → Directive[Black, Thick, Dashed], ImageSize → 375]
```

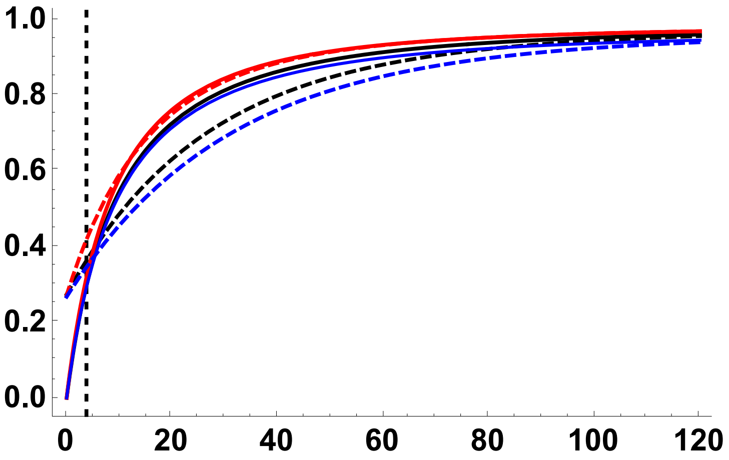


Ratio of the std var to mutation case

```
RatPis0 = Plot[{ExPiSV[5000, 0.05, 0.5, 0, R/10000, 0.05]/
    ExPiMut[5000, 0.05, 0.5, 0, R/10000, 0.2/10000],
   ExPiSV[5000, 0.05, 0.1, 0, R/10000, 0.05]/ ExPiMut[5000, 0.05, 0.1,
    0, R/10000, 0.2/10000], ExPiSV[5000, 0.05, 0.9, 0, R/10000, 0.05]/
    ExPiMut[5000, 0.05, 0.9, 0, R/10000, 0.2/10000]}, {R, 0, 120},
  PlotRange → All, PlotStyle → {{Black, Thick}, {Red, Thick}, {Blue, Thick}},
  GridLines → {{PiSw[0.2, 0, 0.05]}, {1}},
  Frame → {{True, False}, {True, False}},
  BaseStyle → {FontWeight → "Bold", FontColor → Black, FontSize → 16},
  GridLinesStyle → Directive[Black, Thick, Dashed], ImageSize → 375]
```

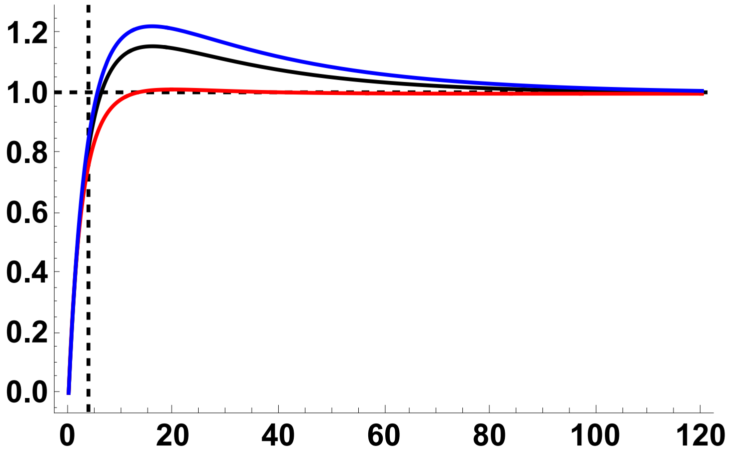


Comparing std var to mutation case. f0 = 0.05, $\sigma$ = 0.95.

Interestingly, sweep signatures are stronger for recurrent mutation as opposed to standing variation. In the sense that they are stretched out over longer regions of the genome.

```
ExPis095 = Plot[{ExPiSV[5000, 0.05, 0.5, 0.95, R/10000, 0.05],
   ExPiMut[5000, 0.05, 0.5, 0.95, R/10000, 0.2/10000],
   ExPiSV[5000, 0.05, 0.1, 0.95, R/10000, 0.05], ExPiMut[5000, 0.05, 0.1,
    0.95, R/10000, 0.2/10000], ExPiSV[5000, 0.05, 0.9, 0.95, R/10000, 0.05],
   ExPiMut[5000, 0.05, 0.95, 0.95, R/10000, 0.2/10000]}, {R, 0, 2000},
  PlotRange → {All, {0, 1}}, PlotStyle → {{Black, Thick}, {Black, Dashed, Thick},
    {Red, Thick}, {Red, Dashed, Thick}, {Blue, Thick}, {Blue, Dashed, Thick}},
  GridLines → {{PiSw[0.2, 0.95/(2 - 0.95), 0.05]}, {}},
  Frame → {{True, False}, {True, False}},
  BaseStyle → {FontWeight → "Bold", FontColor → Black, FontSize → 16},
  GridLinesStyle → Directive[Black, Thick, Dashed], ImageSize → 375]
```

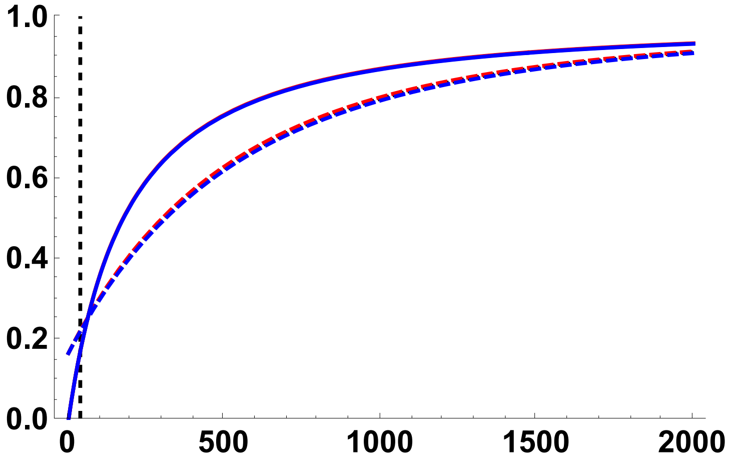

```
RatPis095 =
 Plot[{ExPiSV[5000, 0.05, 0.5, 0.95, R/10 000, 0.05] / ExPiMut[5000, 0.05, 0.5,
     0.95, R/10 000, 0.2/10 000], ExPiSV[5000, 0.05, 0.1, 0.95, R/10 000, 0.05] /
    ExPiMut[5000, 0.05, 0.1, 0.95, R/10 000, 0.2/10 000],
   ExPiSV[5000, 0.05, 0.9, 0.95, R/10 000, 0.05] /
    ExPiMut[5000, 0.05, 0.9, 0.95, R/10 000, 0.2/10 000]}, {R, 0, 2000},
  PlotRange → All, PlotStyle → {{Black, Thick}, {Red, Thick}, {Blue, Thick}},
  GridLines → {{PiSw[0.2, 0.95/(2 - 0.95), 0.05]}, {1}},
  Frame → {{True, False}, {True, False}},
  BaseStyle → {FontWeight → "Bold", FontColor → Black, FontSize → 16},
  GridLinesStyle → Directive[Black, Thick, Dashed], ImageSize → 375]
```

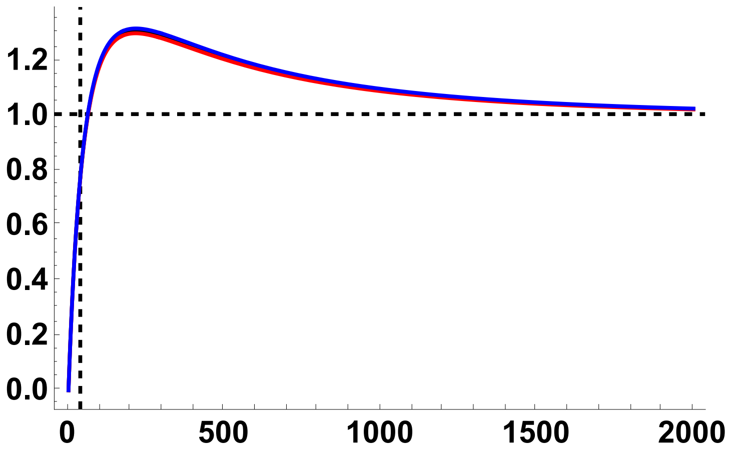


Graphics grid:

```
SVandMut =
  Labeled[Grid[{{, Text@TraditionalForm@Style["σ = 0.00\n(F = 0.00)", 24],
      Text@TraditionalForm@Style["σ = 0.95\n(F ≃ 0.90)", 24]},
     {Text@TraditionalForm@Style["𝔼[πSV/π0],\n𝔼[πM/π0]", 24], ExPis0, ExPis095},
     {Text@TraditionalForm@Style["𝔼[πSV/π0]\n————\n𝔼[πM/π0]", 24],
      RatPis0, RatPis095}}, Spacings → {2, 1}],
   {Text@TraditionalForm@Style["Scaled Recombination Rate, 2Nr", 24]}, {Bottom}]
```

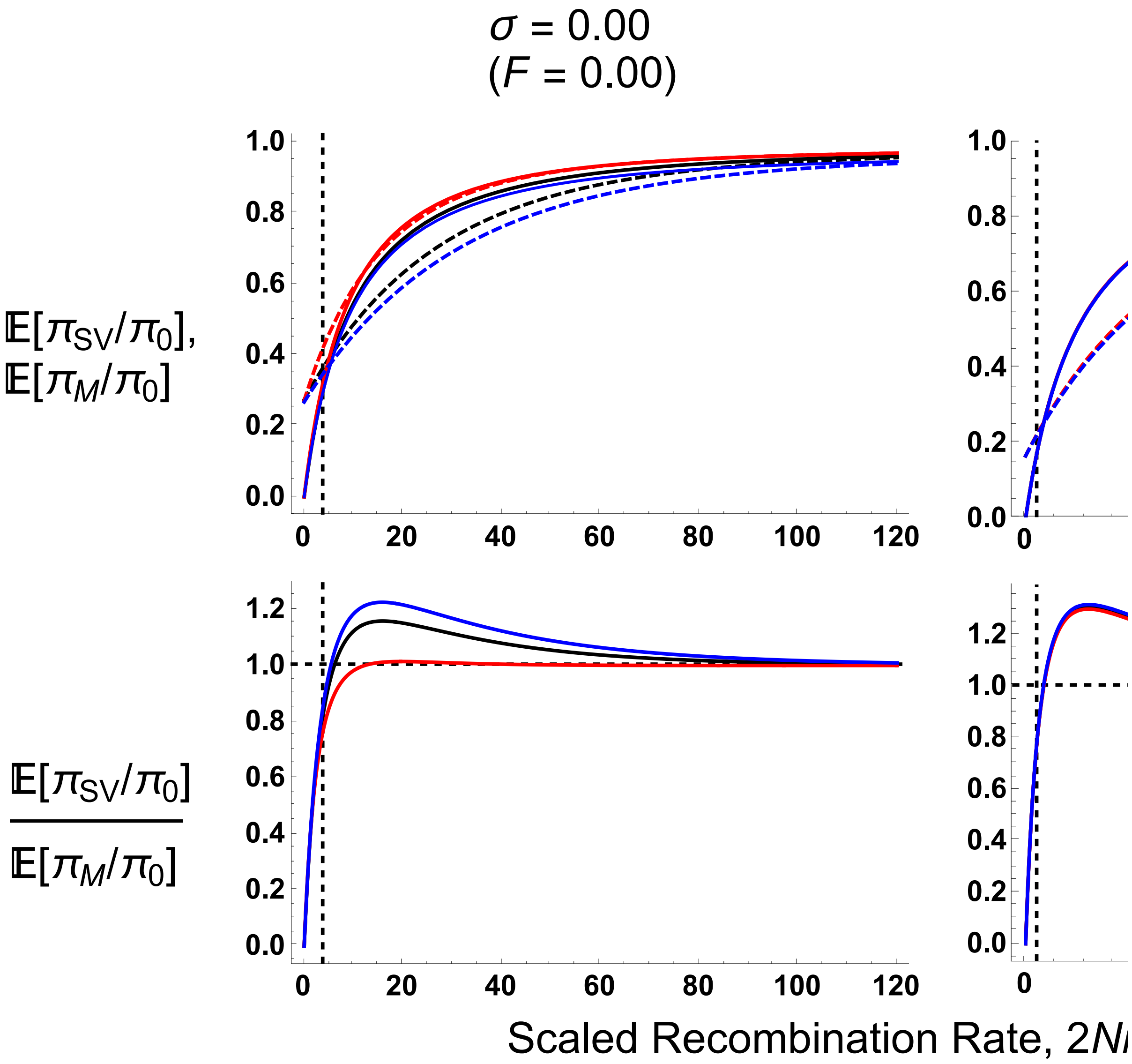


#### Plots – Site Frequency Spectrum (SFS)

##### Equations

For SFS from standing variation:

$$F[\sigma\_] := \frac{\sigma}{2-\sigma};$$

$$\Phi[r\_, \sigma\_] := \frac{\sigma\,(2-\sigma-2\,(1-r)\,r\,(2-3\,\sigma))}{(2-\sigma)\,(2-(1-2\,(1-r)\,r)\,\sigma)};$$

$$\text{PNR}[\text{Na}\_, F\_, \Phi\_, s\_, h\_, R\_, \text{p0}\_] := \left(\left(\frac{(F+h-F\,h)}{(1-h+F\,h)}\left(\frac{1}{\text{p0}}+1\right)-1\right)^{-\frac{R\,(1-F)}{2\,\text{Na}\,(F+h-F\,h)\,s}}\right);$$

$$\text{PrKR}[k\_, n\_, R\_, F\_, \Phi\_, \text{p0}\_] := \left(\left(2\,\frac{R}{1+F}\,(1-2\,F+\Phi)\,\text{p0}\,(1-\text{p0})\right)^{k}\text{Abs}[\text{StirlingS1}[n, k]]\right)\Big/ \text{Product}\left[\left(2\,\frac{R}{1+F}\,(1-2\,F+\Phi)\,\text{p0}\,(1-\text{p0})+a\right), \{a, 0, n-1\}\right];$$

$$\text{PrJ}[j\_, k\_] := \frac{1}{j\,\text{Sum}\left[\frac{1}{a}, \{a, k-1\}\right]};$$

```
H[g_, j_, k_, n_, i_] :=
  (Binomial[n - i, g] * Binomial[k, j - g]) / Binomial[k + n - i, j];
PrL[n_, k_, j_, l_] :=
  ((Binomial[n, l] * Abs[StirlingS1[l, j]] * Abs[StirlingS1[n - l, k - j]]) /
    (Binomial[k, j] * Abs[StirlingS1[n, k]]));
```

$$\text{TFixI}[\text{Na}\_, s\_, h\_, F\_, \text{p0}\_] := \frac{\text{EulerGamma}+\text{Log}\left[\frac{4\,\text{Na}\,s\,(1-(1-F)\,h)\,(1-\text{p0})}{1+F}\right]}{s\,(1-(1-F)\,h)} - \frac{\text{Log}[\text{p0}]}{s\,(h+F-h\,F)} + \frac{(1-F)\,(1-2\,h)}{s\,(h+F-h\,F)\,(1-(1-F)\,h)}\,\text{Log}\left[\frac{h+F-h\,F+(1-F)\,(1-2\,h)\,\text{p0}}{1-(1-F)\,h}\right]$$

$$\text{TFixIC}[\text{Na}\_, s\_, h\_, F\_, \text{p0}\_] := \frac{(1+F)\,\text{TFixI}[\text{Na}, s, h, F, \text{p0}]}{2\,\text{Na}}$$

$$\text{Boostp0}[\text{Na}\_, s\_, h\_, \sigma\_] := \frac{1+F[\sigma]}{4\,\text{Na}\,s\,(F[\sigma]+h-F[\sigma]\,h)}$$

```
PLNH[Na_, s_, h_, σ_, R_, n_, l_, p0_, θ_] :=
 Sum[PDF[BinomialDistribution[n, PNR[Na, F[σ], Φ[R/(2 Na), σ], s, h, R, p0]], i] *
   (Piecewise[{{Sum[PrKR[k, i, R, F[σ], Φ[R/(2 Na), σ], p0] *
            Sum[PrJ[j, k + n - i] * Sum[H[g, j, k, n, i] * PrL[i, k, j - g, l - g],
               {g, Max[{j - k, l - i}], Min[{j, l, n - i}]}], {j, 1,
              Min[{k + n - i - 1, l}]}], {k, 1, i}], 0 < i < n}, {0, i == 0 || i == n}}] +
     Piecewise[{{1/(l Sum[1/a, {a, n - 1}]), i == 0}, {0, i ≠ 0}}] +
     Piecewise[{{PrKR[1, i, R, F[σ], Φ[R/(2 Na), σ], p0] * Piecewise[
            {{θ * p0 + (θ n)/2 TFixIC[Na, s, h, F[σ], p0], l == 1}, {(θ * p0)/l, l ≠ 1}}] +
          Sum[PrKR[k, i, R, F[σ], Φ[R/(2 Na), σ], p0] * Sum[PrJ[j, k] * PrL[n, k, j, l],
             {j, 1, Min[{k - 1, l}]}], {k, 2, n}], i == n}, {0, i ≠ n}}]), {i, 0, n}]

PLNHS[Na_, s_, h_, σ_, R_, n_, p0_, θ_] :=
 Sum[PLNH[Na, s, h, σ, R, n, L, p0, θ], {L, 1, n - 1}]
PLNH2[Na_, s_, h_, σ_, R_, n_, l_, p0_, θ_] := PLNH[Na, s, h, σ, R, n, l, p0, θ] / PLNHS[Na, s, h, σ, R, n, p0, θ]
```

Modifying equations for the SFS from standing variation to account for different Ewens' Sampling Formula under the recurrent mutation case:

$$F[\sigma\_] := \frac{\sigma}{2-\sigma};$$

$$\Phi[r\_, \sigma\_] := \frac{\sigma\,(2-\sigma-2\,(1-r)\,r\,(2-3\,\sigma))}{(2-\sigma)\,(2-(1-2\,(1-r)\,r)\,\sigma)};$$

$$PNR[Na\_, F\_, \Phi\_, s\_, h\_, R\_, p0\_] := \left(\left(\frac{(F+h-F\,h)}{(1-h+F\,h)}\left(\frac{1}{p0}+1\right)-1\right)^{-\frac{R\,(1-F)}{2\,Na\,(F+h-F\,h)\,s}}\right);$$

$$PrKRM[k\_, n\_, F\_, \Theta\_] := \frac{\left(2\,\frac{\Theta}{1+F}\right)^{k} Abs[StirlingS1[n, k]]}{Product\left[\left(2\,\frac{\Theta}{1+F}+a\right), \{a, 0, n-1\}\right]}$$

$$PrJ[j\_, k\_] := \frac{1}{j\,Sum\left[\frac{1}{a}, \{a, k-1\}\right]};$$

```
H[g_, j_, k_, n_, i_] :=
  (Binomial[n - i, g] * Binomial[k, j - g]) / Binomial[k + n - i, j];
PrL[n_, k_, j_, l_] :=
  ((Binomial[n, l] * Abs[StirlingS1[l, j]] * Abs[StirlingS1[n - l, k - j]]) /
    (Binomial[k, j] * Abs[StirlingS1[n, k]]));
TFixI[Na_, s_, h_, F_, p0_] :=
```

$$\frac{EulerGamma + Log\left[\frac{4\,Na\,s\,(1-(1-F)\,h)\,(1-p0)}{1+F}\right]}{s\,(1-(1-F)\,h)} - \frac{Log[p0]}{s\,(h+F-h\,F)} + \frac{(1-F)\,(1-2\,h)}{s\,(h+F-h\,F)\,(1-(1-F)\,h)}\,Log\left[\frac{h+F-h\,F+(1-F)\,(1-2\,h)\,p0}{1-(1-F)\,h}\right]$$

$$TFixIC[Na\_, s\_, h\_, F\_, p0\_] := \frac{(1+F)\,TFixI[Na, s, h, F, p0]}{2\,Na}$$

$$Boostp0[Na\_, s\_, h\_, F\_] := \frac{1+F}{4\,Na\,s\,(F+h-F\,h)}$$

```
PLNHM[Na_, s_, h_, σ_, R_, n_, l_, Θ_, θ_] :=
 Sum[PDF[BinomialDistribution[n, PNR[Na, F[σ],
      Φ[R/(2 Na), σ], s, h, R, Boostp0[Na, s, h, F[σ]]]], i] * (Piecewise[
      {{Sum[PrKRM[k, i, F[σ], Θ] * Sum[PrJ[j, k + n - i] * Sum[H[g, j, k, n, i] *
               PrL[i, k, j - g, l - g], {g, Max[{j - k, l - i}], Min[{j, l, n - i}]}], {j,
             1, Min[{k + n - i - 1, l}]}], {k, 1, i}], 0 < i < n}, {0, i == 0 || i == n}}] +
     Piecewise[{{1/(l Sum[1/a, {a, n - 1}]), i == 0}, {0, i ≠ 0}}] +
     Piecewise[{{PrKRM[1, i, F[σ], Θ] * Piecewise[
            {{θ * Boostp0[Na, s, h, F[σ]] + (θ n)/2 TFixIC[Na, s, h, F[σ], Boostp0[Na,
                   s, h, F[σ]]], l == 1}, {1/l θ * Boostp0[Na, s, h, F[σ]], l ≠ 1}}] +
          Sum[PrKRM[k, i, F[σ], Θ] * Sum[PrJ[j, k] * PrL[n, k, j, l], {j, 1, k - 1}],
           {k, 2, n}], i == n}, {0, i ≠ n}}]), {i, 0, n}]

PLNHMS[Na_, s_, h_, σ_, R_, n_, Θ_, θ_] :=
 Sum[PLNHM[Na, s, h, σ, R, n, L, Θ, θ], {L, 1, n - 1}]
PLNHM2[Na_, s_, h_, σ_, R_, n_, l_, Θ_, θ_] := PLNHM[Na, s, h, σ, R, n, l, Θ, θ] / PLNHMS[Na, s, h, σ, R, n, Θ, θ]
```

#### Figures

Standing variation case (5%, black) vs recurrent mutation ($\Theta = 0.2$, red), $\sigma = 0$, $R = 2$

```
SFSM1 =
 ListPlot[{Table[{l, PLNH2[5000, 0.05, 0.5, 0, 2, 10, l, 0.05, 4]}, {l, 1, 9}],
   Table[{l, PLNHM2[5000, 0.05, 0.5, 0, 2, 10, l, 0.2, 4]}, {l, 1, 9}],
   Table[{l, PrJ[l, 10]}, {l, 1, 9}]}, PlotRange → All,
  PlotStyle → {Black, Red, {Gray, Dashed}}, Joined → True,
  BaseStyle → {FontWeight → "Bold", FontColor → Black, FontSize → 16},
  Ticks → {{1, 2, 3, 4, 5, 6, 7, 8, 9}, Automatic}, ImageSize → 375]
```

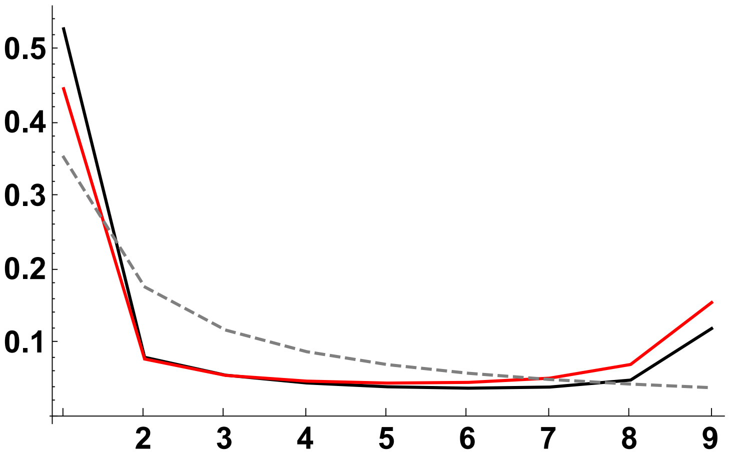


Standing variation case (5%, black) vs recurrent mutation (Θ = 0.2, red), $\sigma = 0$, R = 10

```
SFSM2 =
 ListPlot[{Table[{l, PLNH2[5000, 0.05, 0.5, 0, 10, 10, l, 0.05, 4]}, {l, 1, 9}],
   Table[{l, PLNHM2[5000, 0.05, 0.5, 0, 10, 10, l, 0.2, 4]}, {l, 1, 9}],
   Table[{l, PrJ[l, 10]}, {l, 1, 9}]}, PlotRange → All,
  PlotStyle → {Black, Red, {Gray, Dashed}}, Joined → True,
  BaseStyle → {FontWeight → "Bold", FontColor → Black, FontSize → 16},
  Ticks → {{1, 2, 3, 4, 5, 6, 7, 8, 9}, Automatic}, ImageSize → 375]
```

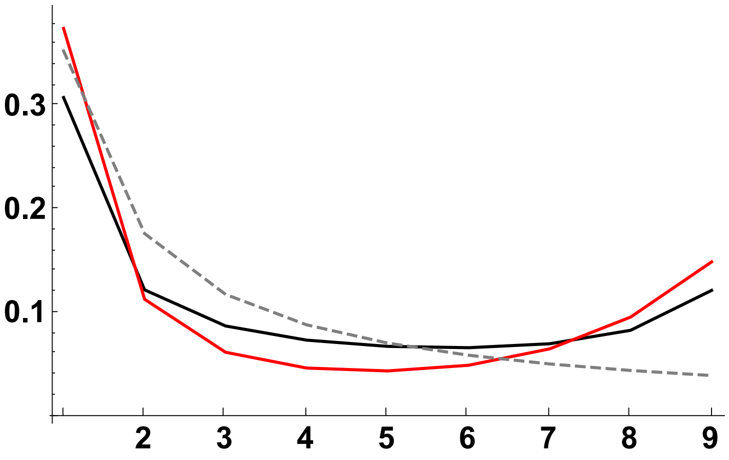


Standing variation case (5%, black) vs recurrent mutation (Θ = 0.2, red), $\sigma = 0.95$, R = 10

```
SFSM3 = ListPlot[
  {Table[{l, PLNH2[5000, 0.05, 0.5, 0.95, 10, 10, l, 0.05, 4/(1 + 0.95/(2-0.95))]}, {l, 1, 9}],
   Table[{l, PLNHM2[5000, 0.05, 0.5, 0.95, 10, 10, l, 0.2, 4/(1 + 0.95/(2-0.95))]}, {l, 1, 9}],
   Table[{l, PrJ[l, 10]}, {l, 1, 9}]}, PlotRange → All,
  PlotStyle → {Black, Red, {Gray, Dashed}}, Joined → True,
  BaseStyle → {FontWeight → "Bold", FontColor → Black, FontSize → 16},
  Ticks → {{1, 2, 3, 4, 5, 6, 7, 8, 9}, Automatic}, ImageSize → 375]
```

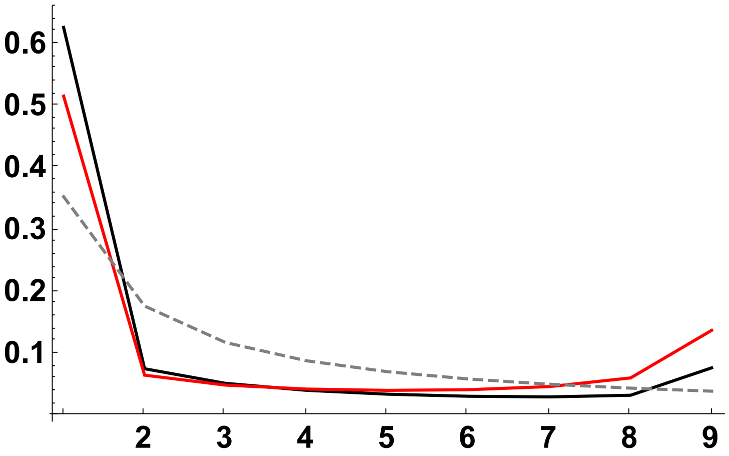


Standing variation case (5%, black) vs recurrent mutation (Θ = 0.2, red), $\sigma$ = 0.95, R = 100

```
SFSM4 = ListPlot[
  {Table[{l, PLNH2[5000, 0.05, 0.5, 0.95, 100, 10, l, 0.05, 4/(1 + 0.95/(2-0.95))]}, {l, 1, 9}],
   Table[{l, PLNHM2[5000, 0.05, 0.5, 0.95, 100, 10, l, 0.2, 4/(1 + 0.95/(2-0.95))]}, {l, 1, 9}],
   Table[{l, PrJ[l, 10]}, {l, 1, 9}]}, PlotRange → All,
  PlotStyle → {Black, Red, {Gray, Dashed}}, Joined → True,
  BaseStyle → {FontWeight → "Bold", FontColor → Black, FontSize → 16},
  Ticks → {{1, 2, 3, 4, 5, 6, 7, 8, 9}, Automatic}, ImageSize → 375]
```

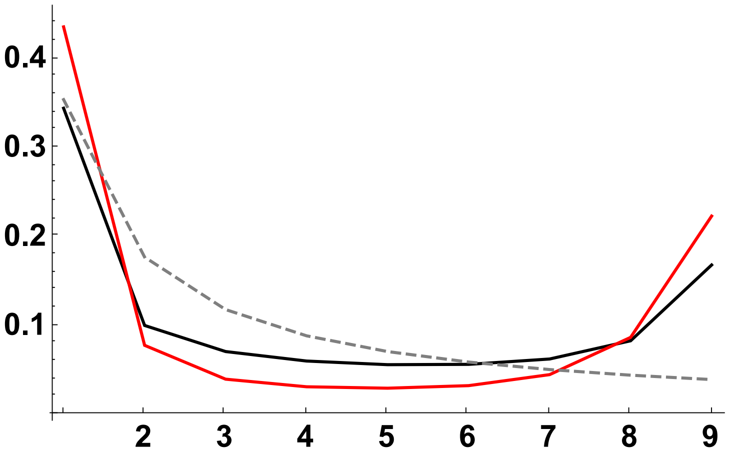


Graphics grid:

```
SVandMutSFS = Labeled[Grid[{{Text@TraditionalForm@Style["R = 2", 24],
      Text@TraditionalForm@Style["R = 10", 24],},
     {SFSM1, SFSM2, Text@TraditionalForm@Style["σ = 0.00\n(F = 0.00)", 24]},
     {Text@TraditionalForm@Style["R = 10", 24],
      Text@TraditionalForm@Style["R = 100", 24],},
     {SFSM3, SFSM4, Text@TraditionalForm@Style["σ = 0.95\n(F ≃ 0.90)", 24]}},
    Spacings → {2, 1}], {Text@TraditionalForm@Style["Frequency", 24],
    Text@TraditionalForm@Style["Derived Allele Count", 24]}, {Left, Bottom}]
```

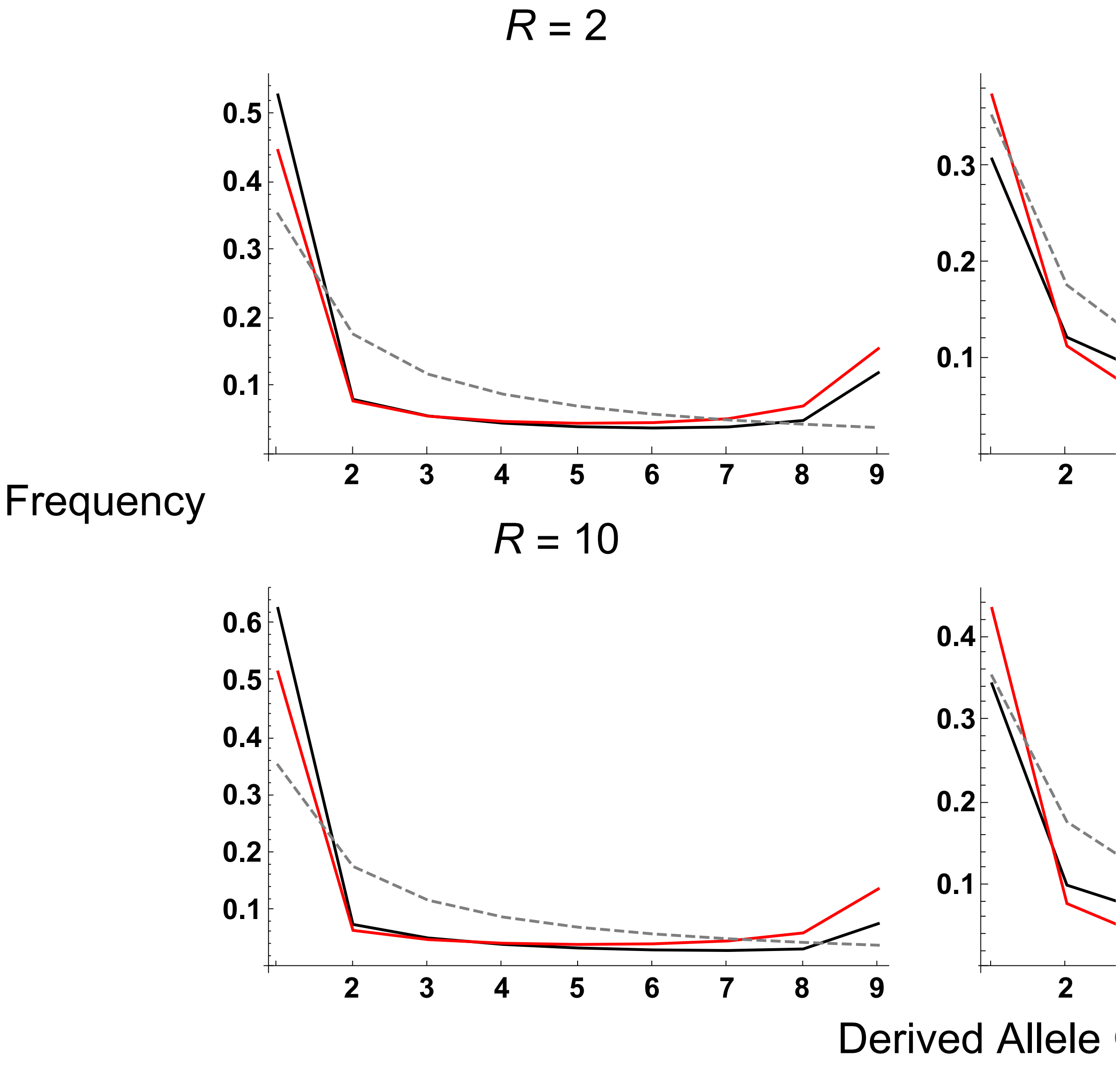
