## Supplemental File S2 for "Selective sweeps under dominance and inbreeding"

### Simulation Procedures

#### Forward–in–time simulations

Forward–in–time simulations were ran using SLiM version 3.3 (Haller and Messer 2019). There exists $N$ diploid individuals, each carrying a chromosome of length 100,000 nucleotides. The target locus carrying the derived allele is present at the left-hand end of the haplotype, while the remaining loci can carry neutral mutations only. Selected alleles have a homozygous selective advantage $s$ and dominance coefficient $h$. Mutation and recombination parameters are input as population-level rates, which are subsequently scaled down to obtain per-locus mutation rates, or a per-inter-base-pair recombination probability.

A 'burn-in' phase is first run to generate background neutral diversity, where the population evolves without any beneficial alleles present for $20N$ generations; this population was subsequently saved. The second phase acts differently depending on whether the beneficial allele is instantly selected for (a hard sweep), or whether it went through a neutral phase. If a hard sweep was simulated, then the beneficial allele was introduced into a single individual as a heterozygote, and tracked until it is fixed or lost. If the latter, the burn-in population is reloaded, the random seed changed and the beneficial mutation reintroduced. The procedure is repeated until the mutation has fixed.

If the derived mutation was initially neutral, then following the burn-in a neutral allele is introduced into a random individual as a heterozygote, and tracked until it is lost or it reaches a frequency $p_0$. If it is lost then the burn-in population is reloaded, the random seed changed and the derived allele is reintroduced. If the mutation reaches the target frequency $p_0$ then it is then converted into a selected mutation, and is tracked until fixation or loss. If the beneficial mutation is subsequently lost then the simulation is stopped and restarted from scratch (i.e., the burn-in population is also regenerated).

100 burn-in populations were generated for each parameter set. After the beneficial allele has gone to fixation, we sampled 10 haplotypes 10 times from each burn-in population to create 1,000 replicate simulations. Mutations are placed in one of 10 bins depending on the distance from the sweep. Relevant statistics (pairwise diversity, relative to neutral expectations; number of segregating sites; site frequency spectrum) were calculated per bin. Mean values are calculated over all 1,000 outputs. 95% confidence intervals were calculated by bootstrapping the simulation results; specifically, simulation results were resampled with replacement 1,000 times, and the per–bin means were calculated from these values. The 2.5% – 97.5% quantile of these resampled means was used to estimate the confidence interval.

#### Coalescent simulations for outcrossing populations

In the forward-in-time simulations, the neutral population was tracked until the segregating variant reached a frequency $p_0$, then it became beneficial. However, this procedure may not fully reflect the genetic history of a sweep from standing variation, as it could have reached a frequency $p_0$ several times during the standing phase before it became selected for. To test whether this assumption introduces any substantial inaccuracies, we also ran coalescent simulations for outcrossing cases using *msms* (Ewing and Hermisson 2010), which is based on the exact history of the derived allele.

The gene genealogies of 10 alleles were simulated, where each haplotype consisted of the derived allele at the left-hand of the genome, and neutral polymorphisms added using an infinite sites model (although we define a finite number of sites). The same parameters were used, and summary statistics were calculated as for the forward-in-time simulations. *msms* does not have an explicit model for simulating sweeps from standing variation, so the derived allele trajectory was generated first before being used as an input for coalescent simulations (using the `--Strace` option). Given the derived allele at a frequency $p_0$ when it started to become selected for, initial genotypes were formed assuming Hardy-Weinberg equilibrium. To form the selected allele trajectory, each generation the genotypes frequencies were first changed by selection by $w_g/\overline{w}$, for $w_g$ the fitness of genotype $g$ (either the ancestral or derived homozygote, or the heterozygote genotype). Reproduction then occurred by creating genotypes according to Hardy-Weinburg expectations. $2N$ haplotypes were then sampled (equivalent to a diploid population of size $N$ under obligate outcrossing) using these new genotypes frequencies to account for random drift. This procedure was repeated until the derived allele reached fixation or was lost; if the latter then the entire selected allele trajectory was recreated until the allele fixed. The neutral trajectory was then created by randomly sampling genotypes from a multinomial distribution back in time from a frequency $p_0$ until the mutation was fixed or lost. If it fixed then the process was repeated until loss occurred. The two trajectories were then pasted together to create the entire allele trajectory. 1,000 trajectories were simulated, with each one used as a basis for individual *msms* simulation runs.

### Additional analytical results

#### Deriving the 'star-like' approximation for $\mathbb{E}(\pi_{SV}/\pi_0)$; comparing against the full solution

It is possible to obtain more tractable analytical solutions by using a 'separation-of-timescales' approximation, and assume that no coalescence occurs during the sweep phase (Pennings and Hermisson 2006b; Berg and Coop 2015). Here we only have to calculate the probability that no recombination occurs during the sweep phase, which for a single lineage equals:

$$\begin{aligned}
P_{NE,SL} &= \exp\left(-\int_{p=1}^{p_0} \frac{(P_r(p)/2)}{\mathrm{d}p/\mathrm{d}t}\mathrm{d}p\right) \\
&= \exp\left(-\int_{p=1}^{p_0} \frac{r_{eff}(1-p)}{\mathrm{d}p/\mathrm{d}t}\mathrm{d}p\right) \\
&= \exp\left(-\frac{r_{eff}}{H_l s}\log\left[\frac{H_l}{H_h}\left(\frac{1}{p_0}+1\right)-1\right]\right) \\
&= \left[\frac{H_l}{H_h}\left(\frac{1}{p_0}+1\right)-1\right]^{-r_{eff}/(H_l s)} \qquad (1)
\end{aligned}$$

$P_r$ is divided by 2 in Equation 1 as it only considers a single lineage. For the special case $\sigma = 0$ and $h = 1/2$, $H_l = H_h = 1/2$ and Equation 1 reduces to $(1/p_0)^{-(2r/s)}$, which is equivalent to Equation 2 of Berg and Coop (2015) after scaling the selection coefficient by 1/2 to include semidominance. Note that $p_0$ has to generally be set to a higher value than $1/2N$ for a hard sweep, to condition on the beneficial allele going to fixation. This effect is considered in the section 'Effective starting frequency for a *de novo* mutation' in the main text. Equation 1 can be used to derived an analytical solution for the relative diversity (Equation 10 in the main text).

Figure A demonstrates how the 'star-like' approximation for $\mathbb{E}(\pi_{SV}/\pi_0)$ overestimates the diversity following a sweep, when compared to the solution that considers coalescence during the sweep phase (Equation 9 in the main text). The differences can be substantial for hard sweeps in outcrossers, especially where the underlying adaptive mutation is recessive.

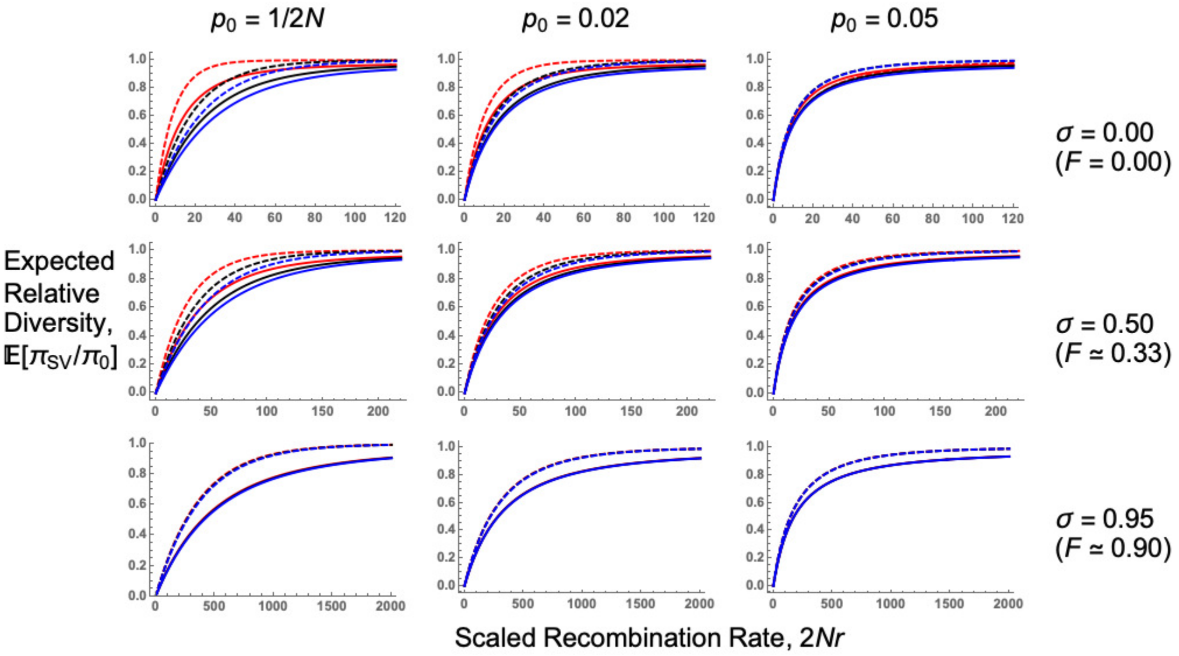


Figure A: **Comparing the full solution against the star-like approximation.** Plots of $\mathbb{E}(\pi_{SV}/\pi_0)$ as a function of the recombination rate scaled to population size $2Nr$. Solid lines are the full analytical solutions (Equation 9 in the main text), while dashed lines are the star-like approximation (Equation 10 in the main text). $N = 5,000$, $s = 0.05$, $4N\mu = 40$, and dominance coefficient $h = 0.1$ (red lines), 0.5 (black lines), or 0.9 (blue lines).

#### Number of Segregating Sites

We next calculate the number of segregating sites, and the site frequency spectrum. Both derivations are based on determining the number of lineages that are present at the end of the sweep phase. Approximations can be more readily obtained if using the star-like approximation, so the following results are likely to be most accurate if applied to strongly-selected beneficial mutations.

We calculate the total time underlying the genealogy, $\mathbb{E}(T_{tot})$, and therefore the expected number of segregating sites $\mathbb{E}(S)$. We consider $n$ samples of the derived allele; looking back in time, $i$ of these alleles fail to recombine away from the derived background during the sweep. The probability of this event can be drawn from a binomial distribution with probability $P_{NE,SL}$ (Equation 1). We denote this value $P_{NE}(i|n) \sim Bin(n, P_{NE,SL})$. Out of these $i$ alleles, let $k$ of them recombine during the sweep phase to create different ancestral backgrounds of the derived allele. Berg and Coop (2015) demonstrated how the number of lineages that recombine away from the derived background can be determined using Ewens' Sampling Formula:

$$P_{ESF}(k|i) = S(i,k)\frac{R_{p_0}^k}{\prod_{l=1}^{i-1}(R_{p_0}+l)} \tag{2}$$

where $R_{p_0} = 4Nrp_0(1-p_0)$ is the scaled recombination rate acting on the ancestral background at frequency $p_0$, and $S(i,k)$ are non-negative Stirling numbers of the first kind (Abramowitz and Stegun 1970; Pennings and Hermisson 2006b; Berg and Coop 2015). Here, we use the rescaled version of $R_{p_0}$ accounting for the reduced effective recombination rate and effective population size caused by self-fertilisation:

$$P_{ESF}(k|i) = S(i,k)\frac{(2R(1-2F+\Phi)p_0(1-p_0)/(1+F))^k}{\prod_{l=1}^{i-1}((2R(1-2F+\Phi)p_0(1-p_0)/(1+F))+l)} \quad (3)$$

Finally, for the $k$ neutral lineages created in the standing phase, along with the $n-i$ neutral lineages created in the sweep phase, the expected total time for the genealogy for all of them, in units of $2N_e$ generations, equals $\sum_{j=1}^{k+n-i-1} 1/j$ (Watterson 1975). The total time covered by the genealogy is the product of these three terms, summed over all possible outcomes:

$$\mathbb{E}(T_{tot}) = \sum_{i=0}^{n} P_{NE}(i|n) \sum_{k=0}^{i} P_{ESF}(k|i) \sum_{j=1}^{k+n-i-1} 1/j \quad (4)$$

$\mathbb{E}(S)$ is $\theta\mathbb{E}(T_{tot})$ where $\theta = 4N_e\mu$ is the population level mutation rate (Hudson 1990). Equivalent results for outcrossing populations are given by Pennings and Hermisson (2006b, Equation 15) for adaptation from recurrent mutation, and Berg and Coop (2015, Equation 10) for adaptation from standing variation. Both these derivations assume $k > 1$ in the standing phase, as it was argued that $\mathbb{E}(T_{tot}) = 0$ so no segregating polymorphisms exist. Since simulation results show that this outcome is possible under low recombination rates, we do not include this conditioning in Equation 4.

Figure B plots $\mathbb{E}(S)$ alongside simulation results. The analytical solution provides a good fit but tends to overestimate simulations, especially for recessive mutations in outcrossing populations. Overestimation was also observed by Berg and Coop (2015), and likely arises as a consequence of not accounting for coalescence during the sweep phase. Also note that fewer segregating sites are present with partial selfing, due to a reduction in the net mutation rate $\theta = 4N_e\mu$

caused by lower $N_e$.

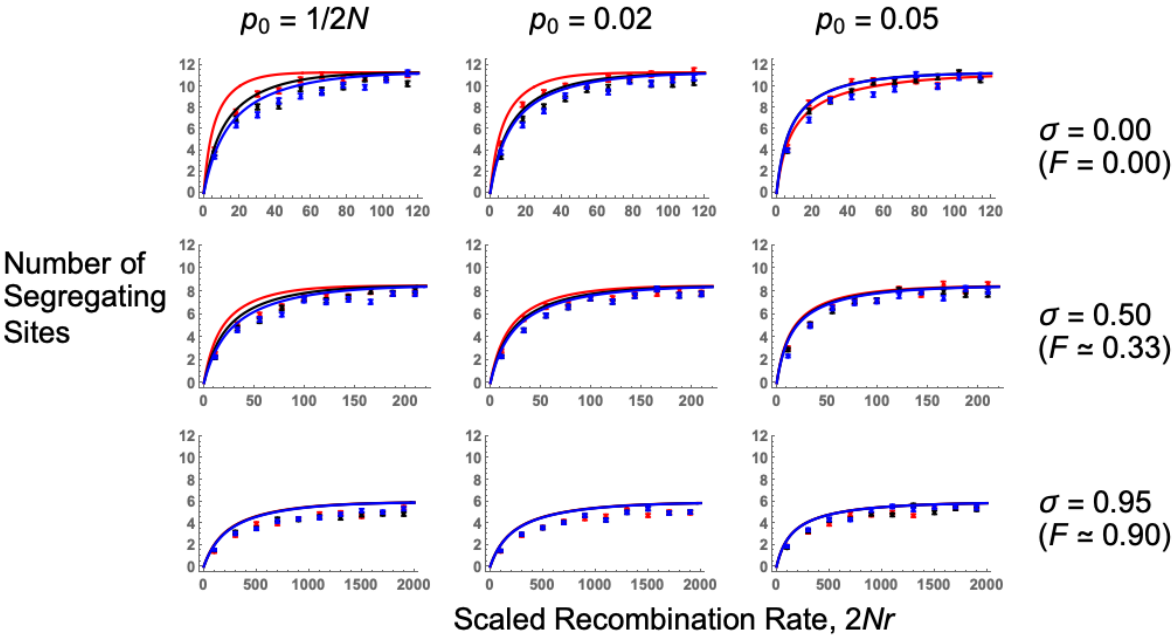


Figure B: **Expected number of segregating sites following a selective sweep.** A plot of $\mathbb{E}(S)$, as a function of the recombination rate scaled to population size $2Nr$. Lines are analytical solutions (Equation 4 multiplied by $\theta$), points are simulation results. $N = 5,000$, $s = 0.05$, $4N\mu = 40$ (so $\theta = 4N_e\mu$ per bin is 4 for $\sigma = 0$, 3 for $\sigma = 0.5$, and 2.1 for $\sigma = 0.95$), and dominance coefficient $h = 0.1$ (red lines, points), 0.5 (black lines, points), or 0.9 (blue lines, points). Further results are plotted in Section D of Supplementary File S1.

### Deriving the Site Frequency Spectrum

The calculations for $\mathbb{E}(S)$ can be extended to determine the full site-frequency spectrum (SFS) following a sweep; that is, the probability that out of $n$ sampled alleles, $l = 1, 2 \ldots n-1$ of them carry the derived mutation. Here we outline the derivation, based on that used by Berg and Coop (2015, Equation 15).

The form of the SFS depends on how many of the $n$ sampled lineages recombine away from the derived background during the sweep, which is determined by the probability $P_{NR}(i|n)$. First look at the special case of $i = 0$. Here, all lineages recombine onto the ancestral neutral background during the sweep. In this case the frequency of allele counts $l$ are determined by the neutral SFS:

$$p(l|n, i = 0) = q(l, n) \tag{5}$$
$$= \frac{1}{la_n} \tag{6}$$

where $a_n = \sum_{a=1}^{n-1}(1/a)$ (Wakeley 2009).

Now look at the opposite extreme where $i = n$, so none of the $n$ lineages recombine away from the derived background. In this case the probability that the $n$ lineages recombine into $k$ different ancestral backgrounds is given by $P_{ESF}$ (Equation 3). If $k > 1$ and $j$ of these $k$ backgrounds carry the derived allele, we can obtain the probability that $l$ of $n$ alleles carry the derived mutation by partitioning the $j$ derived background amongst the $l$ sampled backgrounds, as also used by Pennings and Hermisson (2006b, Equation 15) and Berg and Coop (2015, Equation 11):

$$p_{anc}(l|j, k, n) = \frac{\binom{n}{l}}{\binom{k}{j}} \frac{S(l, j)S(n - l, k - j)}{S(n, k)} \tag{7}$$

where $S(i, j)$ represents Stirling numbers of the first kind (Abramowitz and Stegun 1970). Of these $j$ derived backgrounds, the relative frequency of each is given by $q(j, k)$ as defined by Equation 5. By summing over $k > 1$ one obtains the expression for the frequency spectrum, following Berg and Coop (2015, Equation 14):

$$p(l|n, i = n, k > 1) = \sum_{k=2}^{i} P_{ESF}(k|R_{p0,F}, n) \sum_{j=1}^{k-1} p_{anc}(l|j, k, n)q(j, k) \tag{8}$$

$R_{p0,F} = 2R(1-2F+\Phi)p_0(1-p_0)/(1+F)$ is the rescaled Ewens' Sampling Formula parameter for the probability of recombination.

If $k = 1$ then Equation 7 becomes invalid since it evaluates to zero, so we have to consider this case separately. Simulation results (Figure 5 in the main text) suggest that a large number of low-frequency variants are created by private polymorphisms appearing on each haplotype during the sweep phase. We hence update Equation 8 to account for these new polymorphisms (Berg and Coop 2015):

$$p(l|n, i = n) = P_{ESF}(1|R_{p0,F}, n) \cdot q_{new}(l|i = n) + p(l|n, i = n, k > 1) \tag{9}$$

where $q_{new}(l)$ accounts for new mutations arising during the sweep:

$$q_{new}(l|i = n) = \theta \left( \frac{p_0}{l} + \mathbb{1}_1(l) \cdot \frac{n}{2}\tau_{p_0} \right) \tag{10}$$

where $\theta$ is the population-level neutral mutation rate for the region of interest, and $\mathbb{1}_1(l)$ is an indicator function, equal to 1 at $l = 1$ and zero for all other values. $\tau_{p_0}$ is the fixation time of the sweep from initial frequency $p_0$; we use Glémin (2012, Equation 21), scaling results by $2N_e = 2N/(1 + F)$ so they are on the coalescent timescale.

Finally, we consider the general case $0 < i < n$. Here we consider the probability that $n - i$ recombinant backgrounds were created during the sweep, with $k$ more created in the standing phase. $j$ of these $k + n - i$ recombinant backgrounds carry the derived allele; we then further partition which $g$ of these $j$ alleles lie on recombinant backgrounds created during the sweep. The full partition is given by Berg and Coop (2015, Equations 15–16):

$$\begin{aligned} p(l|n, 0 < i < n) = &\sum_{k=1}^{i} P_{ESF}(k|R_{p_0,F}, i) \\ &\times \sum_{j=1}^{Min(k+n-i-1,l)} q(j, k+n-i) \\ &\times \sum_{g=Max(j-k,l-i)}^{Min(j,l,n-i)} H(g|j,k,n-i) p_{anc}(l-g|j-g,k,i) \end{aligned} \tag{11}$$

where $H$ is the partitioning function:

$$H(g|j,k,n-i) = \frac{\binom{n-i}{g}\binom{k}{j-g}}{\binom{k+n-i}{j}} \tag{12}$$

Note that Equation 11 corrects a couple of errors present in Equation 15 of Berg and Coop (2015). First, $P_{ESF}$ takes $i$ as an input as opposed to $n-i$ that Berg and Coop (2015) use. This is because we wish to determine the probability that $k$ recombinant lineages form from $i$ lineages that did not recombine during the sweep phase. Second, the lower bound of the sum over $g$ depends on $Max(j-k, l-i)$, while Berg and Coop (2015) use $Max(j-k, 0)$. Since $j-g$ of the derived alleles are on the $k$ recombinant backgrounds created during the standing phase, $g \geq j-k$, hence the inclusion of this term in the sum's lower bound. However, $l-g$ of the derived alleles are on the $i$ recombinant backgrounds created during the sweep phase, implying that $g \geq l-i$ as well. Finally, if $i = 1$ then Equation 11 collapses to the same result as for $i = 0$.

Putting all this information together, the site frequency spectrum equals:

$$
\begin{aligned}
p(l|n) = &\sum_{i=0}^{n} P_{NR}(i|n) \\
&\times [p(l|n, i = 0)\mathbb{1}_0(i) \\
&+ p(l|n, 0 < i < n)\mathbb{1}_{0<i<n}(i) \\
&+ p(l|n, i = n)\mathbb{1}_n(i)]
\end{aligned} \tag{13}
$$

where $\mathbb{1}_A(i)$ equals 1 for $i \in A$ and is otherwise zero. Equation 13 needs to be normalised, due to the addition of the new polymorphism terms (Equation 9). Hence the normalised SFS is given by dividing each allele entry $p(l|n)$ by $\sum_l p(l|n)$.

#### The SFS with recurrent mutation

Equation 13 can be modified to account for sweeps from recurrent mutation. Pennings and Hermisson (2006a) demonstrated that after the sweep phase, the number of ancestral haplotypes created in the standing phase is given by Ewens' Sampling Formula; i.e., Equation 2 but with parameter $4N_e\mu_b = 2\Theta_b/(1+F)$, instead of $R_{p_0}$. We also set $p_0 = p_{0,A}$ in $P_{NE}$. Figure C compares SFS from standing variation or recurrent mutation. For $R < R_{Lim}$ (Equation 16 in the main text), sweeps from recurrent mutation show more intermediate-frequency variants, which is a typical soft sweep signature. Conversely sweeps from standing variation display more intermediate-frequency variants once $R > R_{Lim}$.

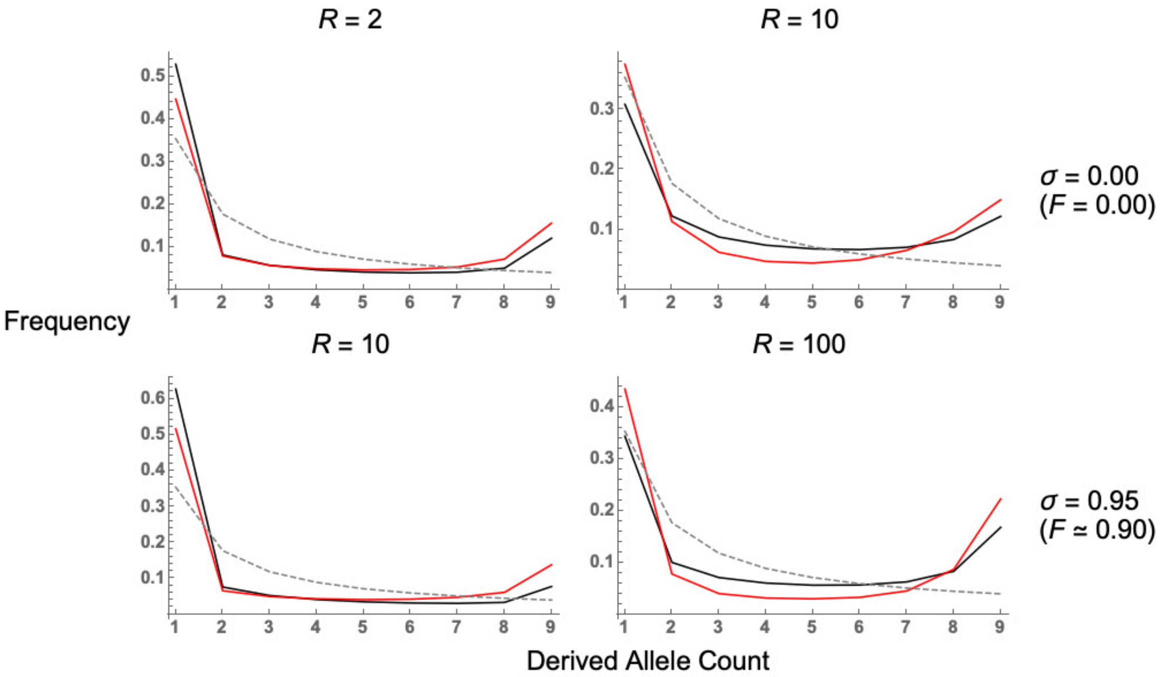


Figure C: Comparing SFS for soft sweeps from standing variation ($p_0 = 0.05$, black lines) against those from recurrent mutation ($\Theta = 0.2$, red lines). Neutral expectations are also shown for comparison (gray dashed lines). $N = 5,000$, $s = 0.05$, $h = 1/2$. Note that for these parameters, $R_{Lim} = 4$ under $\sigma = 0$, or $R_{Lim} = 42$ with $\sigma = 0.95$.

#### Comparing forward-in-time simulations against coalescent simulations

We compared the three main analytical results (for expected relative diversity in Figure D; for the number of segregating sites in Figure E; and the SFS in Figure F) to both sets of simulation results. As expected, for a hard sweep ($p_0 = 1/2N$) the two approaches give equivalent results. For higher $p_0$ the approaches are similar for higher recombination rates, however for lower recombination rates the forward-in-time simulation yields higher estimates of both $\mathbb{E}(\pi_{SV}/\pi_0)$ and $\mathbb{E}(S)$. In addition, forward-in-time simulations predict a visibly higher number of singletons in the SFS than the coalescent approaches. Overall it seems that forward-in-time simulations accurately captures the behaviour of a sweep from standing variation, but caution must be used when simulating results with low recombination rates, especially if investigating the site frequency spectrum.

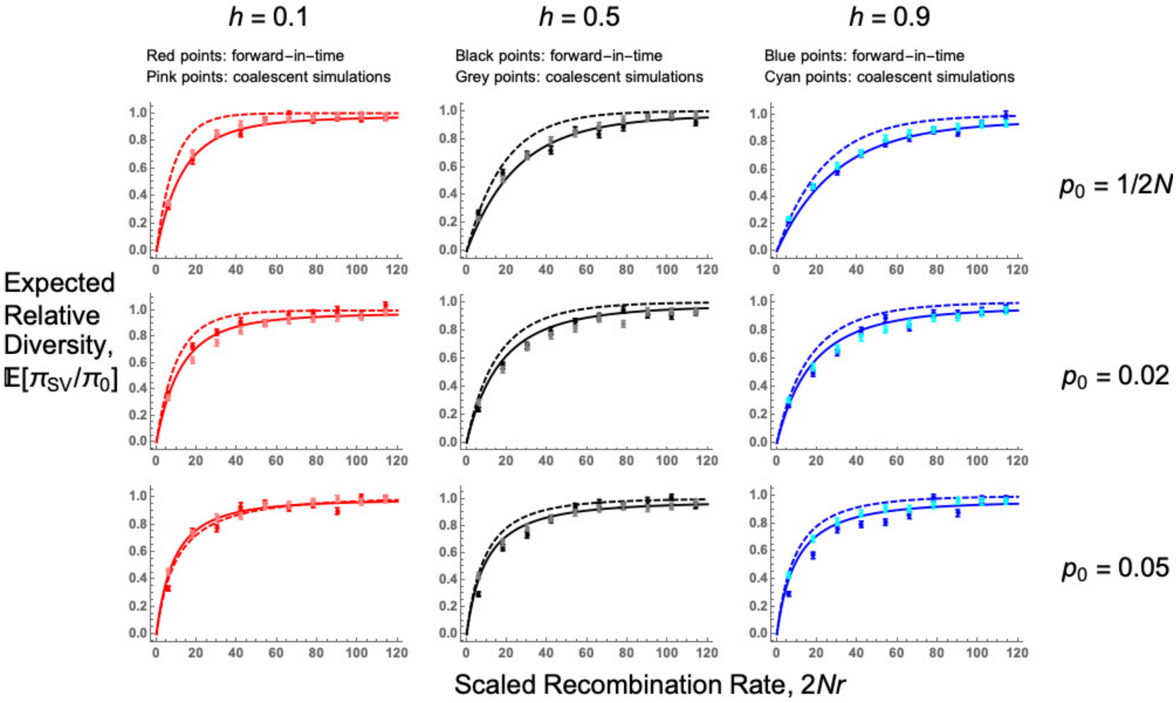


Figure D: **Comparing SLiM forward-in-time simulations with MSMS coalescent simulations, for $\mathbb{E}(\pi_{SV}/\pi_0)$.** Plots are as a function of the recombination rate scaled to population size $2Nr$. Solid lines are the full analytical solutions (Equation 9 in main text), while dashed lines are the star-like approximation (Equation 10 in the main text). Points are the simulation results, with different colours representing different simulation approaches, as denoted in the figure headings. $N = 5,000$, $s = 0.05$, $4N\mu = 40$. Rate of self-fertilisation equals 0. Values for the dominance coefficients and starting frequencies are listed in the figure.

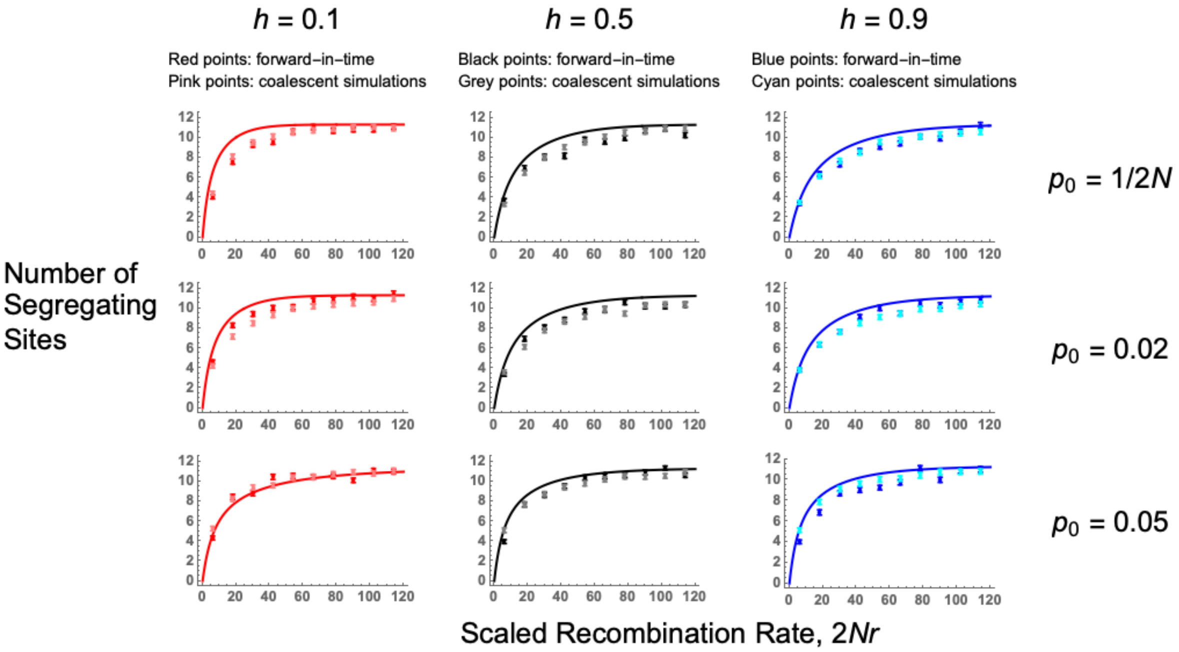


Figure E: **Comparing SLiM forward-in-time simulations with MSMS coalescent simulations, for number of segregating sites.** Plots are as a function of the recombination rate scaled to population size $2Nr$. Solid lines are the analytical solutions (Equation 4 multiplied by $\theta$). Points are the simulation results, with different colours representing different simulation approaches, as denoted in the figure headings. $N = 5,000$, $s = 0.05$, $4N\mu = 40$. Rate of self-fertilisation equals 0.Values for the dominance coefficients and starting frequencies are listed in the figure.

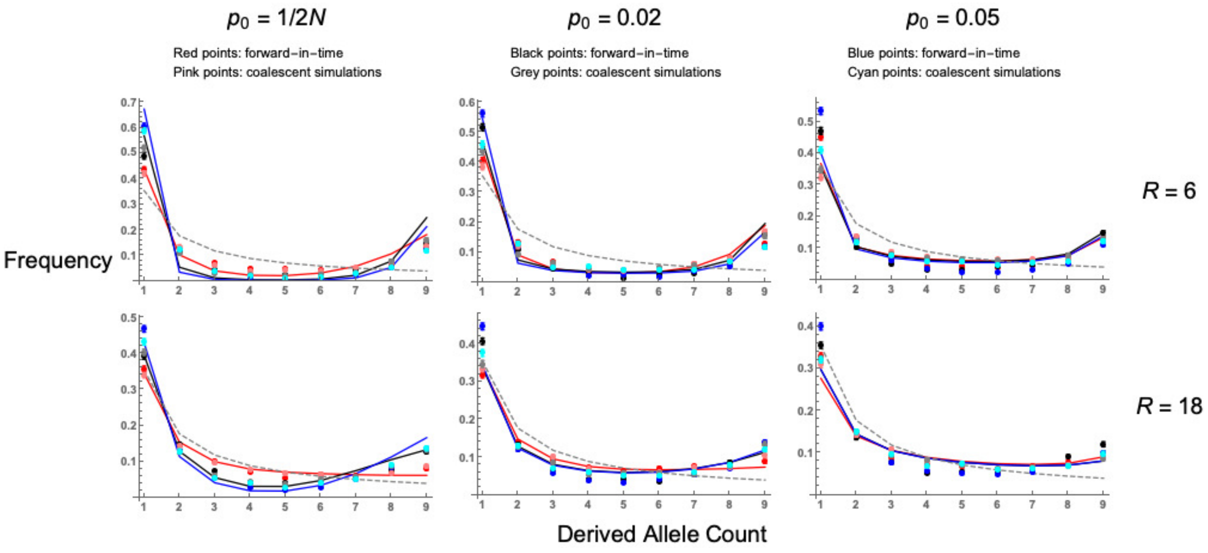


Figure F: **Comparing SLiM forward-in-time simulations with MSMS coalescent simulations, for SFS.** Lines are analytical solutions (Equation 13), points are different simulation results as indicated above the plots. $N = 5,000$, $s = 0.05$, $4N\mu = 40$, and dominance coefficient $h = 0.1$ (red lines, points), 0.5 (black lines, points), or 0.9 (blue lines, points). The neutral SFS is also included for comparisons (grey dashed line). Rate of self-fertilisation $\sigma = 0$.
