## Supplemental File S3 for "Selective sweeps under dominance and inbreeding": DataRun.sh

```
#!/bin/bash
#SBATCH -c 1
#SBATCH --mem-per-cpu 128
#SBATCH --partition short
#SBATCH --time=0:20:00

# Script for analysing SLiM data on GenomeDK server

# Inputs
POP=5000
IX=0.05
THETA=40
REC=2000
DOM=0.9
SELF=0.95
SAMPS=10
NPS=10
REPS=100
TOT=$(($NPS*$REPS))

source /com/extra/R/LATEST/load.sh
rm Sim*
rm Mutations/MT_* Mutations/Muts_*
rm *.out

# Preparing MS output files to input standard
for ((i=1; i<=TOT; i++));
do
	printf -v x "Mutations/RawMut%d.dat" $i
	awk -v fx=$i 'BEGIN {RS="//";}{ file = "Mutations/MT_" sprintf("%03d", fx) ".dat"}{for (k=1; k<=NF; k++) print $k," "> file; close(file)}' $x
	printf -v j "Mutations/MT_%03d.dat" $i
 	printf -v k "Mutations/Muts_%03d.dat" $i
 	tail -n +4 $j > $k
done

Rscript DataScriptBatch_Bins.R $TOT $SAMPS $THETA $REC $DOM $SELF $IX 0
```
