## Supplemental File S3 for "Selective sweeps under dominance and inbreeding": DataScriptBatch_Bins.R

```
# Scripts for reading in simulation data
# Then calculating key parameters
# Usage: Rscript DataScriptBatch_Bins.R l s th rec h self f0 sw
# l = Number of input files
# s = Number of haplotype samples
# th = 4Nu is the population-level mutation rate
# rec = 2Nr is the population-level recombination rate
# h = dominance coefficient of beneficial mutation
# self = Frequency of self-fertilisation
# f0 = Starting frequency of sweep when it started to be selected for
# sw = Switch to determine output file name (0 = 'SLIM'; 1 = 'MSMS')

rm(list = ls()) ## Clearing space
require(compiler)	## For speeding up execution
invisible(enableJIT(3))

args <- commandArgs(trailingOnly = TRUE)
l <- as.double(args[1])
s <- as.double(args[2])
th <- as.double(args[3])
rec <- as.double(args[4])
h <- as.double(args[5])
self <- as.double(args[6])
f0 <- as.double(args[7])
sw <- as.double(args[8])

bin <- 10
xa <- seq((0.5/bin),(1-0.5/bin),1/bin)
xa <- xa*rec
Fis <- self/(2-self)
tpb <- (th/(bin*(1 + Fis)))	#Theta per bin, normalised to expected value with given inbreeding level

pwi <- combn(1:s,2)		# List of pairwise individuals
sum2 <- function(x) {sum(x)%%2}	# Sum of entries mod 2 (for pairwise diff calculation)
meanna <- function(x) {mean(x,na.rm=T)}
varna <- function(x) {var(x,na.rm=T)}

SegSites <- matrix(data=NA,nrow=l,ncol=bin)		# Number of segregating sites
PairDiff <- matrix(data=NA,nrow=l,ncol=bin)		# Number of pairwise differences
PDScaled <- matrix(data=NA,nrow=l,ncol=bin)		# Number of pairwise differences scaled to neutral exp
Watt <- matrix(data=NA,nrow=l,ncol=bin)			# Watterson's estimator for theta
TD <- matrix(data=NA,nrow=l,ncol=bin)			# Tajima's D for each run
FWH <- matrix(data=NA,nrow=l,ncol=bin)			# Fay and Wu's H per run
NFWH <- matrix(data=NA,nrow=l,ncol=bin)			# Normalised Fay and Wu's H per run
UniqueH <- matrix(data=NA,nrow=l,ncol=bin)		# Number unique haplotypes in a run
H1 <- matrix(data=NA,nrow=l,ncol=bin)			# Sweep statistics (haplotype-based)
H2 <- matrix(data=NA,nrow=l,ncol=bin)			
H12 <- matrix(data=NA,nrow=l,ncol=bin)
H2H1 <- matrix(data=NA,nrow=l,ncol=bin)
SFST <- lapply(1:bin, function(x) matrix(NA, nrow=l, ncol=s-1))		# List of SFS per bin per sim
#SFSM <- matrix(data=NA,nrow=l,ncol=s-1)			# Matrix of mean SFS outcomes per bin
#SFSV <- matrix(data=NA,nrow=l,ncol=s-1)			# Matrix of mean SFS outcomes per bin

# Function to calculate Tajima's D from data
tajd <- function(pi,seg,n){
	a1 <- sum(1/(c(1:(n-1)))^1)
	a2 <- sum(1/(c(1:(n-1)))^2)
	b1 <- (n+1)/(3*(n-1))
	b2 <- (2*(n^2+n+3))/(9*n*(n-1))
	c1 <- b1 - (1/a1)
	c2 <- b2 - (n+2)/(a1*n) + (a2)/(a1^2)
	e1 <- c1/a1
	e2 <- c2/(a1^2 + a2)
	
	Dee <- (pi - seg/a1)/(sqrt(e1*seg + e2*seg*(seg - 1)))
	return(Dee)
}	# End of 'tajd' function

# Function to calculate Fay and Wu's H
faywu <- function(sf,n){
	# Standard FWH
	tp <- sum((c(1:(n-1))*((n-1)-c(1:(n-1))))*sf)/(choose(n,2))
	tH <- sum((c(1:(n-1))^2)*sf)/(choose(n,2))
	
	# 'Normalised' FWH
	a1 <- sum(1/(c(1:(n-1)))^1)
	a2 <- sum(1/(c(1:(n-1)))^2)
	a2a <- sum(1/(c(1:n))^2)
	tL <- sum(c(1:(n-1))*sf)/(n-1)
	tW <- sum(sf)/a1
	v <- ((n-2)/(6*(n-1)))*tW + ((18*(n^2)*((3*n) + 2)*a2a - (88*(n^3) + 9*(n^2) - 13*n + 6))/(9*n*((n - 1)^2)))*((sum(sf)*(sum(sf)-1))/(a1^2 + a2))
		
	return(c(tp-tH,(tp-tL)/sqrt(v)))
}	# End of 'faywu' function

a1 <- function(n) {sum(1/(c(1:(n-1)))^1)}

for(i in 1:l){
	# print(i)
	sze <- file.info(sprintf("Mutations/Muts_%03d.dat",i))$size
	if(sze==0){
		for(j in 1:bin){
			SegSites[i,j] <- 0
			UniqueH[i,j] <- 1
			PairDiff[i,j] <- 0
			PDScaled[i,j] <- 0
			Watt[i,j] <- 0
			TD[i,j] <- NA
			H1[i,j] <- 1
			H12[i,j] <- 1
			H2[i,j] <- 0
			H2H1[i,j] <- 0
			SFST[[j]][i,] <- NA
			FWH[i,j] <- NA
			NFWH[i,j] <- NA
		}
	}else{
		spot <- as.matrix(read.table(sprintf("Mutations/Muts_%03d.dat",i))$V1[1:(length(read.table(sprintf("Mutations/Muts_%03d.dat",i))$V1)-s)])
		SegS <- length(spot)
		# Then mutants themselves
		pie <- read.table(sprintf("Mutations/Muts_%03d.dat",i),colClasses=c(rep("character",1)))$V1[(SegS + 1):(length(read.table(sprintf("Mutations/Muts_%03d.dat",i))$V1))]
		pie2 <- gsub("(.)", "\\1 ", pie)
		elems <- as.numeric(unlist( strsplit( pie2 , " " ) ))
		MutsT <- as.data.frame(matrix(elems , ncol = s, nrow=SegS, byrow=F ))	
		levs <- as.numeric(cut(spot,seq(0,1,1/bin)))
		for(j in 1:bin){
			Muts <- MutsT[which(levs==j),1:dim(MutsT)[2]]
			if(dim(Muts)[1]==0){
				SegSites[i,j] <- 0
				UniqueH[i,j] <- 1
				PairDiff[i,j] <- 0
				PDScaled[i,j] <- 0
				Watt[i,j] <- 0
				TD[i,j] <- NA
				H1[i,j] <- 1
				H12[i,j] <- 1
				H2[i,j] <- 0
				H2H1[i,j] <- 0
				SFST[[j]][i,] <- NA
				FWH[i,j] <- NA
				NFWH[i,j] <- NA
			}else{
				SegSites[i,j] <- dim(Muts)[1]		# Number of seg. sites
				Watt[i,j] <- SegSites[i,j]/a1(s);		
				UniqueH[i,j] <- dim(unique(as.matrix(Muts),MARGIN=2))[2] # Number unique haplotypes
			
				# Haplotype Statistics
				HapFreq <- vector(mode="numeric",length=UniqueH[i,j])		# Frequency of haplotypes
				for(k in 1:s){
					HapFreq[which(colSums(Muts[,k]==unique(as.matrix(Muts),MARGIN=2)) == SegSites[i,j])] <- HapFreq[which(colSums(Muts[,k]==unique(as.matrix(Muts),MARGIN=2)) == SegSites[i,j])] + 1
				}
				HapFreq <- HapFreq/s
				HapFreq <- sort(HapFreq,decreasing=T)
				H1[i,j] <- sum(HapFreq^2)
				H12[i,j] <- H1[i,j] + 2*HapFreq[1]*HapFreq[2]
				H2[i,j] <- sum(HapFreq[-1]^2)
				H2H1[i,j] <- H2[i,j]/H1[i,j]
			
				# Calculating mean pairwise differences, then Tajima's D
				PairDiff[i,j] <- mean(apply(pwi,2,function(x) sum(apply(Muts[x],1,sum2))))
				PDScaled[i,j] <- PairDiff[i,j]/tpb
				TD[i,j] <- tajd(PairDiff[i,j],SegSites[i,j],s)
			
				# Calculating SFS, then Fay + Wu's H using it
				SFS <- vector(mode="numeric",length=s-1)	
				SFS[1:length(tabulate(apply(Muts,1,sum)))] <- tabulate(apply(Muts,1,sum))
				fwo <- faywu(SFS,s)		# Calculating Fay and Wu's H USING UNSCALED SFS
				FWH[i,j] <- fwo[1]
				NFWH[i,j] <- fwo[2]
				SFST[[j]][i,] <- SFS/(dim(Muts)[1])
			}
		}
	}
}

# After compiling results, calculating bootstraps
## matrices to store bootstrap calculations
SegSitesBS <- matrix(data=NA,nrow=1000,ncol=bin)		# Number of segregating sites
PairDiffBS <- matrix(data=NA,nrow=1000,ncol=bin)		# Number of pairwise differences
PDBSScal <- matrix(data=NA,nrow=1000,ncol=bin)			# Number of pairwise differences
WattBS <- matrix(data=NA,nrow=1000,ncol=bin)			# Watterson's estimator for theta
TDBS <- matrix(data=NA,nrow=1000,ncol=bin)			# Tajima's D for each run
FWHBS <- matrix(data=NA,nrow=1000,ncol=bin)			# Fay and Wu's H per run
NFWHBS <- matrix(data=NA,nrow=1000,ncol=bin)			# Normalised Fay and Wu's H per run
UniqueHBS <- matrix(data=NA,nrow=1000,ncol=bin)		# Number unique haplotypes in a run
H1BS <- matrix(data=NA,nrow=1000,ncol=bin)			# Sweep statistics (haplotype-based)
H2BS <- matrix(data=NA,nrow=1000,ncol=bin)			
H12BS <- matrix(data=NA,nrow=1000,ncol=bin)
H2H1BS <- matrix(data=NA,nrow=1000,ncol=bin)

for(a in 1:1000){
	BSsamp <- sample.int(l, replace = T)
	PairDiffBS[a,] <- colMeans(PairDiff[BSsamp,],na.rm=T)
	PDBSScal[a,] <- colMeans(PDScaled[BSsamp,],na.rm=T)
	SegSitesBS[a,] <- colMeans(SegSites[BSsamp,],na.rm=T)
	WattBS[a,] <- colMeans(Watt[BSsamp,],na.rm=T)
	TDBS[a,] <- colMeans(TD[BSsamp,],na.rm=T)
	FWHBS[a,] <- colMeans(FWH[BSsamp,],na.rm=T)
	NFWHBS[a,] <- colMeans(NFWH[BSsamp,],na.rm=T)
	UniqueHBS[a,] <- colMeans(UniqueH[BSsamp,],na.rm=T)
	H1BS[a,] <- colMeans(H1[BSsamp,],na.rm=T)
	H2BS[a,] <- colMeans(H2[BSsamp,],na.rm=T)
	H12BS[a,] <- colMeans(H12[BSsamp,],na.rm=T)
	H2H1BS[a,] <- colMeans(H2H1[BSsamp,],na.rm=T)	
}

CIcalc <- function(y){
	apply(y,2,function(x) quantile(x,c(0.025,0.975)))
}

## Creating output table
stattab <- 
rbind(colMeans(SegSites,na.rm=T),abs(CIcalc(SegSitesBS)[1,]-colMeans(SegSites,na.rm=T)),CIcalc(SegSitesBS)[2,]-colMeans(SegSites,na.rm=T),
colMeans(PairDiff,na.rm=T),abs(CIcalc(PairDiffBS)[1,]-colMeans(PairDiff,na.rm=T)),CIcalc(PairDiffBS)[2,]-colMeans(PairDiff,na.rm=T),
colMeans(Watt,na.rm=T),abs(CIcalc(WattBS)[1,]-colMeans(Watt,na.rm=T)),CIcalc(WattBS)[2,]-colMeans(Watt,na.rm=T),
colMeans(PDScaled,na.rm=T),abs(CIcalc(PDBSScal)[1,]-colMeans(PDScaled,na.rm=T)),CIcalc(PDBSScal)[2,]-colMeans(PDScaled,na.rm=T),
colMeans(TD,na.rm=T),abs(CIcalc(TDBS)[1,]-colMeans(TD,na.rm=T)),CIcalc(TDBS)[2,]-colMeans(TD,na.rm=T),
colMeans(NFWH,na.rm=T),abs(CIcalc(NFWHBS)[1,]-colMeans(NFWH,na.rm=T)),CIcalc(NFWHBS)[2,]-colMeans(NFWH,na.rm=T),
colMeans(UniqueH,na.rm=T),abs(CIcalc(UniqueHBS)[1,]-colMeans(UniqueH,na.rm=T)),CIcalc(UniqueHBS)[2,]-colMeans(UniqueH,na.rm=T),
colMeans(H1,na.rm=T),abs(CIcalc(H1BS)[1,]-colMeans(H1,na.rm=T)),CIcalc(H1BS)[2,]-colMeans(H1,na.rm=T),
colMeans(H2,na.rm=T),abs(CIcalc(H2BS)[1,]-colMeans(H2,na.rm=T)),CIcalc(H2BS)[2,]-colMeans(H2,na.rm=T),
colMeans(H12,na.rm=T),abs(CIcalc(H12BS)[1,]-colMeans(H12,na.rm=T)),CIcalc(H12BS)[2,]-colMeans(H12,na.rm=T),
colMeans(H2H1,na.rm=T),abs(CIcalc(H2H1BS)[1,]-colMeans(H2H1,na.rm=T)),CIcalc(H2H1BS)[2,]-colMeans(H2H1,na.rm=T)
)
if(sw==0){
	write.table(stattab,file=paste0('StatsProc_R_',rec,'_h_',h,'_self_',self,'_f_',f0,'_10b_SLIM.dat'),row.names=F,col.names=F,quote=F)
}else if(sw == 1){
	write.table(stattab,file=paste0('StatsProc_R_',rec,'_h_',h,'_self_',self,'_f_',f0,'_10b_MSMS.dat'),row.names=F,col.names=F,quote=F)
}

# Forming vector of 'mean' SFS (with CIs)
SFSM <- lapply(SFST,function(x) colMeans(x,na.rm=T))
SFSCI <- lapply(SFST,function(x) apply(x,2,function(y) sd(y,na.rm=T)/sqrt(sum(!is.na(y)))))
for(b in 1:bin){
	if(sw == 0){
		write.table(rbind(SFSM[[b]],SFSCI[[b]]),file=paste0('SFSRes/SFSTab_R_',xa[b],'_h_',h,'_self_',self,'_f_',f0,'_10b_SLIM.dat'),row.names=F,col.names=F,quote=F)
	}else if(sw == 1){
		write.table(rbind(SFSM[[b]],SFSCI[[b]]),file=paste0('SFSRes/SFSTab_R_',xa[b],'_h_',h,'_self_',self,'_f_',f0,'_10b_MSMS.dat'),row.names=F,col.names=F,quote=F)
	}
}

## EOF
```
