## Supplemental File S3 for "Selective sweeps under dominance and inbreeding": README.md

README FOR DOM SELF SWEEP FILES

Scripts used to simulate neutral diversity around a sweeping beneficial allele, while considering arbitrary levels of self-fertilisation and dominance. As used in the study "Selective sweeps under dominance and inbreeding". SLiM is available from https://messerlab.org/slim/.

***SLiM Files***

There are two scripts. "StdVarSweep_Hard.slim" is for simulating a 'hard' sweep, where a derived mutations has a selective advantage from it's appearance. "StdVarSweepFromNeut.slim" simulates sweeps arising from standing variation. These programs are meant to be run from the command line. For each program one first has to run 'mkdir Mutations/ BIPops/ Seeds/' to generate directories to store outputs. In each script the haplotype size is set to 100,000 basepairs, and the homozygote selection coefficient of the beneficial allele equals 0.05.

To run "StdVarSweep_Hard.slim", use:

slim -seed <SEED> -d N=<POP SIZE> -d Theta=<MUT RATE> -d R=<REC RATE> -d sps=<OUTPUT HAPLOTYPES> -d numsamp=<NUMBER OF SAMPLES> -d h=<DOMINANCE> -d sfrate=<SELFING FREQUENCY> -d simID=<ID> StdVarSweep_Hard.slim

Where:
- seed is the starting seed for the random number generator
- N is the population size
- Theta = 4Nu is the population-level mutation rate
- R = 2Nr is the population-level recombination rate across the entire haplotype
- sps is the number of haplotypes to sample from the final population
- numsamp is the number of times 'sps' haplotypes are sampled per simulation run
- h is the dominance value
- sfrate is the frequency of self-fertilisation
- simID is the simulation ID, used to label outputs

To run "StdVarSweepFromNeut.slim", use:

slim -seed <SEED> -d N=<POP SIZE> -d Theta=<MUT RATE> -d R=<REC RATE> -d x0=<STARTING FREQUNCY> -d sps=<OUTPUT HAPLOTYPES> -d numsamp=<NUMBER OF SAMPLES> -d h=<DOMINANCE> -d sfrate=<SELFING FREQUENCY> -d simID=<ID> ./StdVarSweepFromNeut_V6.slim

Parameters are the same as in "StdVarSweep_Hard.slim", except 'x0' is the frequency at which the derived allele becomes selected for.

The 'Seeds' folder contains a list of random number seeds used in each simulation. 'BIPops' outputs the state of the population after the burn-in. 'Mutations' outputs diversity (in MS format) for 'sps' haplotype samples after the sweep has fixed.

***C Code***

The program 'StdVarTrajs_Sampling.c' is used to simulate sweep trajectories to be used with MSMS. It uses routines found with the GNU Scientific Library (GSL) (http://www.gnu.org/software/gsl/) Since GSL is distributed under the GNU General Public License (http://www.gnu.org/copyleft/gpl.html), you must download it separately from this file.

Programs can be compiled with e.g. GCC using a command like: gcc StdVarTrajs_Sampling -lm -lgsl -lgslcblas -I/usr/local/include -L/usr/local/lib StdVarTrajs_Sampling.c.

The program is run by executing:
./StdVarTrajs_Sampling N s h x0 reps cutoff
Where:
- N is the (haploid) population size
- s is the selection coefficient of the beneficial allele
- h is dominance of the beneficial allele
- x0 is frequency of the allele when it started to be selected for
- reps is how many times the second allele should *fix* before simulation stops (the number of actual runs is greater due to stochastic loss)
- cutoff denotes how high the derived allele can be in the neutral phase ('0' for fixation, '1' for x0 + 1/2N; '0' was used for results in the manuscript)

Trajectories are output in the 'Traj' folder.

***R code***

The file 'DataScriptBatch_Bins.R' is an R script used to calculate summary statistics from simulation results. It can be run by:

Rscript DataScriptBatch_Bins.R l s th rec h self f0 sw

Where:
- l = Number of input files
- s = Number of haplotype samples
- th = 4Nu is the population-level mutation rate
- rec = 2Nr is the population-level recombination rate
- h = dominance coefficient of beneficial mutation
- self = Frequency of self-fertilisation
- f0 = Starting frequency of sweep when it started to be selected for
- sw = Switch to determine output file name (0 = 'SLIM'; 1 = 'MSMS')

Comments to.
