## Supplemental File S3 for "Selective sweeps under dominance and inbreeding": SlimRun.sh

```
#!/bin/bash
#SBATCH -c 1
#SBATCH --mem-per-cpu 128
#SBATCH --partition short
#SBATCH --time=0:20:00

# Script for running SLiM program on GenomeDK server

# Inputs
POP=5000
IX=0.05
THETA=40
REC=2000
DOM=0.9
SELF=0.95
SAMPS=10
NPS=10
REPS=100
TOT=$(($NPS*$REPS))
COND=$(expr 2*$POP*$IX/1 | bc)

rm Sim*
rm *.out
rm -r Mutations/ BIPops/ Seeds/ SFSRes/
mkdir Mutations/ BIPops/ Seeds/ SFSRes/

BSEED=90712784618784892
for ((i=0; i<REPS; i++));
do
	echo '#!/bin/bash' > Sim${i}.sh
	echo '#SBATCH -c 1' >> Sim${i}.sh
	echo '#SBATCH --mem-per-cpu 256' >> Sim${i}.sh
	echo '#SBATCH --partition short' >> Sim${i}.sh
	echo '#SBATCH --time=0:15:00' >> Sim${i}.sh
	echo 'source /com/extra/gcc/5.3.0/load.sh' >> Sim${i}.sh
	if (($COND == 1))
	then
		echo "~/SLiM/build/slim -seed $(($BSEED + $i)) -d N=$POP -d Theta=$THETA -d R=$REC -d sps=$SAMPS -d numsamp=$NPS -d h=$DOM -d sfrate=$SELF -d simID=$i ./StdVarSweep_Hard.slim &" >> Sim${i}.sh
	else
		echo "~/SLiM/build/slim -seed $(($BSEED + $i)) -d N=$POP -d Theta=$THETA -d R=$REC -d x0=$IX -d sps=$SAMPS -d numsamp=$NPS -d h=$DOM -d sfrate=$SELF -d simID=$i ./StdVarSweepFromNeut.slim &" >> Sim${i}.sh
	fi
	echo 'wait' >> Sim${i}.sh	
	echo 'exit 0' >> Sim${i}.sh
	chmod u+x Sim${i}.sh
	sbatch Sim${i}.sh
done
wait

exit 0
```
