## Supplemental File S3 for "Selective sweeps under dominance and inbreeding": StdVarTrajs_Sampling.c

```
/* StdVarTrajs_Sampling.c 

Simulation for calculating allele trajectories, adaptation from standing variation
To be fed into MSMS

Simulation uses routines found with the GNU Scientific Library (GSL)
(http://www.gnu.org/software/gsl/)
Since GSL is distributed under the GNU General Public License 
(http://www.gnu.org/copyleft/gpl.html), you must download it 
separately from this file.

This program can be compiled with e.g. GCC using a command like:
gcc StdVarTrajs_Sampling -lm -lgsl -lgslcblas -I/usr/local/include -L/usr/local/lib StdVarTrajs_Sampling.c

Then run by executing:
./StdVarTrajs_Sampling N s h x0 reps cutoff
Where:
- N is the (haploid) size
- s is the selection coefficient of the beneficial allele
- h is dominance of the beneficial allele
- x0 is frequency of the allele when it started to be selected for
- reps is how many times the second allele should FIX before simulation stops 
(the number of actual runs is greater due to stochastic loss)
- cutoff denotes how high the derived allele can be in the neutral phase ('0' for fixation, '1' for x0 + 1/2N)
*/

/* Preprocessor statements */
#include <stdio.h>
#include <stdlib.h>
#include <time.h>
#include <math.h>
#include <stddef.h>
#include <string.h>
#include <gsl/gsl_rng.h>
#include <gsl/gsl_randist.h>
#include <gsl/gsl_complex.h>
#include <gsl/gsl_complex_math.h>
#include <sys/stat.h>
#include <sys/types.h>

#define INITTS 310000

/* Function prototypes */
void geninit(double *geninit, double pee);
void selection(double *geninit, double h, double s);
void reproduction(double *geninit);
void bentraj(double *XFreq, unsigned int *btime, double *genotype, double x0, unsigned int rep, unsigned int N, double s, double h, const gsl_rng *r);
void neutraj(double *NFreq, unsigned int *ntime, double *genotype, double x0, unsigned int rep, unsigned int N, unsigned int cutoff, const gsl_rng *r);
void trajprint(double **AFreq, unsigned int *ttime, unsigned int rep);
void Wait();

/* Initialising genotypes */
void geninit(double *geninit, double pee){
	
	*(geninit + 0) = (1-pee)*(1-pee);
	*(geninit + 1) = 2*pee*(1-pee);
	*(geninit + 2) = pee*pee;

}	/* End of gen initiation routine */

/* Selection routine (single locus) */
void selection(double *geninit, double h, double s){
	/* Fitness of each genotype */
	double WAA, WAa, Waa;
	double Wmean;				/* Mean fitness */
	
	Waa = 1;
	WAa = 1 + h*s;
	WAA = 1 + s;
	
	/* Mean fitness calculation */
	Wmean = ((*(geninit + 0))*Waa) + ((*(geninit + 1))*WAa) + ((*(geninit + 2))*WAA);
	
	/* Changing frequencies by selection */
	*(geninit + 0) = ((*(geninit + 0))*Waa)/Wmean;
	*(geninit + 1) = ((*(geninit + 1))*WAa)/Wmean;
	*(geninit + 2) = ((*(geninit + 2))*WAA)/Wmean;
	
}	/* End of selection routine */

/* Reproduction routine */
void reproduction(double *geninit){

	/* Fed-in genotype frequencies (for ease of programming) */
	double gaas, gAas, gAAs;
	/* Haplotypes */
	double xA, xa;
	
	/* Initial definition of genotypes */
	gaas = *(geninit + 0);
	gAas = *(geninit + 1);
	gAAs = *(geninit + 2);
	
	/* Baseline change in haplotype frequencies with obligate outcrossing */
	xA = gAAs + (gAas)/2.0;
	xa = gaas + (gAas)/2.0;
	
	/* Change in frequencies (HWE) */
	*(geninit + 0) = pow(xa,2);
	*(geninit + 1) = 2.0*xA*xa;
	*(geninit + 2) = pow(xA,2);
		
}	/* End of reproduction routine */


/* Routine to simulate beneficial allele trajectory */
void bentraj(double *XFreq, unsigned int *btime, double *genotype, double x0, unsigned int rep, unsigned int N, double s, double h, const gsl_rng *r){

	unsigned int gt = 1;		/* Generation time */
	unsigned int done = 0;		/* Counter if routine done or not */
	unsigned int exitl = 0;		/* Decide if exit loop or not */
	unsigned int i;				/* Genotype counter */
	double x = 0;				/* Beneficial allele frequency */
	
	unsigned int *gensamp = calloc(3,sizeof(unsigned int));		/* New population samples */
	
	while(done == 0){
		exitl = 0;
		gt = 1;
		x = x0;
		*(XFreq + 0) = x0;
		/* Initiating genotypes */
		geninit(genotype,x0);
		while(exitl == 0){
		
			/* Selection routine */
    		selection(genotype,h,s);
    		
			/* Reproduction routine */
    		reproduction(genotype);
    		       		
    		/* Sampling based on new frequencies */
	       	gsl_ran_multinomial(r,3,N,genotype,gensamp);
       		for(i = 0; i < 3; i++){
    	   		*(genotype + i) = (*(gensamp + i))/(1.0*N);
	       	}
    		
	       	x = 0.5*(*(genotype + 1)) + (*(genotype + 2));			
			*(XFreq + gt) = x;
			gt += 1;
			if(gt > INITTS){
				fprintf(stderr,"Number of generations in selection trajectory exceed vector length (INITTS).\n");
				exit(1);
			}
			
			if( (x == 1) || (x == 0) ){
				exitl = 1;
			}
		}
	
		if(x == 1){
			done = 1;
			*(btime + rep) = (gt - 1);
		}
	}
	
	free(gensamp);
	
}

/* Routine to simulate neutral allele trajectory */
void neutraj(double *NFreq, unsigned int *ntime, double *genotype, double x0, unsigned int rep, unsigned int N, unsigned int cutoff, const gsl_rng *r){

	unsigned int gt = 1;		/* Generation time */
	unsigned int done = 0;		/* Counter if routine done or not */
	unsigned int exitl = 0;		/* Decide if exit loop or not */
	unsigned int i;				/* Genotype counter */
	double x = 0;				/* Neutral allele frequency */
	double xmax = 1;			/* Value of x to use as maximum cutoff */
	
	unsigned int *gensamp = calloc(3,sizeof(unsigned int));		/* New population samples */
	
	if(cutoff == 0){
		xmax = 1;
	}else if(cutoff == 1){
		xmax = x0 + (1.0/(2.0*N));
		if(xmax > 1){
			xmax = 1;
		}
	}
/*	printf("xmax is %lf\n",xmax);	*/
	
	while(done == 0){
		exitl = 0;
		gt = 0;		/* Note start from zero here since x0 case defined in selection trajectory */
		x = x0;
		
		/* Initiating genotypes */
		geninit(genotype,x0);
		
		while(exitl==0){
		
			/* Reproduction routine */
    		reproduction(genotype);
			
			/* Random sampling of allele frequencies (drift) */
	       	gsl_ran_multinomial(r,3,N,genotype,gensamp);
       		for(i = 0; i < 3; i++){
    	   		*(genotype + i) = (*(gensamp + i))/(1.0*N);
	       	}
	       	x = 0.5*(*(genotype + 1)) + (*(genotype + 2));			
			*(NFreq + gt) = x;

			gt++;
			if(gt > INITTS){
				printf("Number of generations in neutral trajectory exceed vector length (INITTS).\n");
				exit(1);
			}
			
			if( (x == 0) || (x >= xmax) ){
				exitl = 1;
			}
		}
	
		if(x == 0){
			done = 1;
			*(ntime + rep) = (gt - 1);
		}
	}
	
	free(gensamp);
	
}

/* Print out trajectories to file */
void trajprint(double **AFreq, unsigned int *ttime, unsigned int rep){
	unsigned int j;
	char filename[32];
	FILE *ofp_tr;				/* Pointer for file output */

	sprintf(filename,"Traj/ATraj%d.dat",(rep + 1));
	ofp_tr = fopen(filename,"w");
	
	for(j = 0; j < *(ttime + rep); j++){
		fprintf(ofp_tr,"%lf %lf %lf\n",*((*(AFreq + 0)) + j),*((*(AFreq + 1)) + j),*((*(AFreq + 2)) + j));
	}
	
	fclose(ofp_tr);
	
}

void Wait(){
	printf("Press Enter to Continue");
	while( getchar() != '\n' );
	printf("\n");	
}

/* Main program */
int main(int argc, char *argv[]){

	/* Declare variables here */
	unsigned int i, a;			/* Rep counter, memory counter */
	unsigned int N = 0;			/* Population Size */
	unsigned int Nreps = 0;		/* Number of repetitions */
	unsigned int acc = 0;		/* Accumulator (when merging two trajectories) */
	unsigned int es = 10;		/* How many extra steps to put in the trajectory files */
	unsigned int cutoff = 0;	/* What kind of back-in-time cutoff to use */
	int j;						/* Timestep counter */
	double s = 0;				/* Strength of selection */
	double h = 0;				/* Dominance level */
	double x0 = 0;				/* Initial allele frequency */
	double lim = 0;				/* Limit of acceptable allele frequencies */
	double dx = 0;				/* Rescaled timestep so can be used by MSMS */
	FILE *ofp_sd;				/* Pointer for seed output */

	/* GSL random number definitions */
	const gsl_rng_type * T;
	gsl_rng * r;
	
	/* Reading in data from command line */
	/* <Program> N s h x0 NReps cutoff */
	if(argc != 7){
		fprintf(stderr,"Six inputs are needed (N s h x0 Reps cutoff).\n");
		exit(1);
	}
	
	N = atoi(argv[1]);
	if(N <= 0){
		fprintf(stderr,"Total Population size N is zero or negative, not allowed.\n");
		exit(1);
	}
	
	/* Defining min frequency limit ** based on unscaled N ** */
	lim = (1.0/(1.0*N));
	
	/* Defining timestep, dx = 1/4N since that's what MSMS uses */
	dx = (1.0/(4.0*N));
	
	s = strtod(argv[2],NULL);
	if(s < 0){
		fprintf(stderr,"Allele strength s is negative, not allowed.\n");
		exit(1);
	}
	
	h = strtod(argv[3],NULL);
	if(h < 0 || h > 1){
		fprintf(stderr,"Dominance value must lie between 0 and 1.\n");
		exit(1);
	}
	
	x0 = strtod(argv[4],NULL);
	if(x0 < lim || x0 > (1-lim)){
		fprintf(stderr,"Initial mutant frequency must lie between %0.5lf and %0.5lf.\n",lim,1-lim);
		exit(1);
	}
	
	/* Number of samples/reps to take */
	Nreps = atoi(argv[5]);
	if(Nreps <= 0){
		fprintf(stderr,"Must set positive number of repetitions.\n");
		exit(1);
	}
	
	/* Determining cutoff */
	cutoff = atoi(argv[6]);
	if(!(cutoff == 0 || cutoff == 1)){
		fprintf(stderr,"'Cutoff' switch must equal 0 or 1.\n");
		exit(1);
	}
	
	unsigned int *btime = calloc(Nreps,sizeof(unsigned int));			/* Number of generations needed for ben allele fixation */
	unsigned int *ntime = calloc(Nreps,sizeof(unsigned int));			/* Number of generations needed for neut allele loss */
	unsigned int *ttime = calloc(Nreps,sizeof(unsigned int));			/* Number of generations needed for whole processes */
	double *genotype = calloc(3,sizeof(double));						/* Genotype frequencies */
	
	/* create a generator chosen by the 
    environment variable GSL_RNG_TYPE */
     
	gsl_rng_env_setup();
	if (!getenv("GSL_RNG_SEED")) gsl_rng_default_seed = time(0);
	T = gsl_rng_default;
	r = gsl_rng_alloc(T);
	ofp_sd = fopen("Seed.dat","w");
	fprintf(ofp_sd,"%lu\n",gsl_rng_default_seed);
	fclose(ofp_sd);
	
	mkdir("Traj/", 0777);
	
	/* Executing simulation */
	for(i = 0; i < Nreps; i++){
		
		if(Nreps > 100){
			if(i%100 == 0){
				printf("Starting run %d\n",i);	
			}
		}
		
		double *XFreq = calloc(INITTS,sizeof(double));			/* Beneficial allele frequency */
		double *NFreq = calloc(INITTS,sizeof(double));			/* Neutral allele frequency */

		/* Simulating beneficial trajectories */
		bentraj(XFreq, btime, genotype, x0, i, N, s, h, r);
		
		/* Simulation neutral trajectory */
		neutraj(NFreq, ntime, genotype, x0, i, N, cutoff, r);
		
		*(ttime + i) = (*(btime+i)) + (*(ntime+i) + 2 + es);
/*		printf("%d %d %d\n",*(btime+i),*(ntime+i),*(ttime+i));	*/
		
		/* Defining memory for AFreq */
		double **AFreq = calloc(3,sizeof(double *));					/* Allele frequency over time */
		for(a = 0; a < 3; a++){											/* Assigning space for each run */
			AFreq[a] = calloc(*(ttime + i),sizeof(double));
		}
		
		/* Merging trajectories 
		Timing is set up so tfix = 0
		And discrete generations is scaled by timestep, dx
		*/
		
		/* First add ten timesteps for x = 0
		So trajectory fully covered in MSMS */
		acc = 0;
		for(j = 0; j < es; j++){
			*((*(AFreq + 0)) + j) = ((*(ttime + i) - es - 1)*dx) + (es - j);
			*((*(AFreq + 1)) + j) = 1.0;
			*((*(AFreq + 2)) + j) = 0.0;
		}
		for(j = *(ntime + i); j >= 0; j--){
			*((*(AFreq + 0)) + acc + es) = (*(ttime + i) - acc - es - 1)*dx;
			*((*(AFreq + 1)) + acc + es) = 1.0-(*(NFreq + j));
			*((*(AFreq + 2)) + acc + es) = *(NFreq + j);
			acc++;
		}
		for(j = 0; j <= *(btime + i); j++){
			*((*(AFreq + 0)) + acc + es) = (*(ttime + i) - acc - es - 1)*dx;
			*((*(AFreq + 1)) + acc + es) = 1.0-(*(XFreq + j));
			*((*(AFreq + 2)) + acc + es) = *(XFreq + j);
			acc++;
		}
		/*
		*(ttime + i) = (acc-1);	
		printf("%d\n",*(ttime+i));
		*/
		
		free(NFreq);
		free(XFreq);
		
		/* Printout of trajectories */
		trajprint(AFreq, ttime, i);
		
		for(a = 0; a < 3; a++){
	 		free(AFreq[a]);
		}
		free(AFreq);
		
	}
	
	gsl_rng_free(r);
	free(genotype);
	free(ttime);
	free(ntime);
	free(btime);
		
	return 0;
	
}	/* End of main program */

/* End of File */
```
